# The post-translational modification landscape of commercial beers

**DOI:** 10.1101/2021.01.27.427706

**Authors:** Edward D. Kerr, Christopher H. Caboche, Cassandra L. Pegg, Toan K. Phung, Claudia Gonzalez Viejo, Sigfredo Fuentes, Mark T. Howes, Kate Howell, Benjamin L. Schulz

## Abstract

Beer is one of the most popular beverages worldwide. As a product of variable agricultural ingredients and processes, beer has high molecular complexity. We used DIA/SWATH-MS to investigate the proteomic complexity and diversity of 23 commercial Australian beers. While the overall complexity of the beer proteome was modest, with contributions from barley and yeast proteins, we uncovered a very high diversity of post-translational modifications (PTMs), especially proteolysis, glycation, and glycosylation. Proteolysis was widespread throughout barley proteins, but showed clear site-specificity. Oligohexose modifications were common on lysines in barley proteins, consistent with glycation by maltooligosaccharides released from starch during malting or mashing. *O*-glycosylation consistent with oligomannose was abundant on secreted yeast glycoproteins. We developed and used data analysis pipelines to efficiently extract and quantify site-specific PTMs from SWATH-MS data, and showed incorporating these features into proteomic analyses extended analytical precision. We found that the key differentiator of the beer glyco/proteome was the brewery, with beer from independent breweries having a distinct profile to beer from multinational breweries. Within a given brewery, beer styles also had distinct glyco/proteomes. Targeting our analyses to beers from a single brewery, Newstead Brewing Co., allowed us to identify beer style-specific features of the glyco/proteome. Specifically, we found that proteins in darker beers tended to have low glycation and high proteolysis. Finally, we objectively quantified features of foam formation and stability, and showed that these quality properties correlated with the concentration of abundant surface-active proteins from barley and yeast.

## Introduction

Beer is one of the most popular beverages, with ~1.95 billion hectolitres produced annually worldwide (1, 2). Beer brewing is a highly controlled, well understood industrial process. Barley (*Hordeum vulgare* L. subsp. *vulgare*) is typically the primary ingredient in brewing. Grains are malted with controlled, partial germination, allowing enzymes to be synthesised and the husk to open, and kilned to limit enzyme activity on the nutrient rich endosperm and add flavour through the Maillard reaction (non-enzymatic browning) (1, 3–6). Malt is milled to open the grains and then mashed, where grain is mixed with warm water to solubilise starch and proteins and allow enzymes to degrade them into smaller sugars and free amino nitrogen (FAN) (3, 5–7). Wort, the liquid portion, is separated from the spent grain and is boiled with addition of hops (*Humulus lupulus*) to sterilise the wort and to provide bitterness and flavour (1, 3, 5, 6, 8, 9). After the boiled wort is cooled it is fermented with the addition of yeast, which consumes the sugar and FAN, producing ethanol and other flavour compounds as a by-product of its growth (5, 6). The fermented wort is then matured, packaged, and sold as beer to consumers.

Beers come in many diverse styles, all with different flavours and characteristics. These differences arise from differences during the brewing process. Different kilning parameters or performing additional roasting can change the colour and flavours of the malt which carry over into the beer; heavily roasted malts are used to make porters, stouts, and other dark beers. Hops added during the boil or in fermentation can add bitterness or fruity/citrusy flavours, as in India Pale Ales (IPAs) and other hop-forward beers. Fermentation using different yeast or bacteria can also affect many characteristics of the beer. Fermentation with *Saccharomyces cerevisiae* results in an ale, while *Saccharomyces pastorianus* produces a lager. Different yeast strains can produce different amounts of specific esters and other flavour compounds, subtly changing the flavour of the final beer.

Many proteins in beer are modified, including with proteolysis, glycosylation, and glycation. Proteolysis from malt proteases is abundant in mashing, and many proteolytically clipped proteins remain present in the mature beer (10, 11). Yeast secrete proteins during fermentation, many of which are glycosylated with high mannose *N*-linked and oligomannose *O*-linked glycans (12). Yeast glycoproteins have been previously observed in sparkling wine (13, 14), and can impact bubble formation and stability (13, 15–18). Glycation is a nonenzymatic modification of proteins with reducing sugars via the Maillard reaction (19). This involves the reaction of an aldehyde group of a reducing sugar such as glucose with an amine group in a protein, such as the N-terminus or internal Lys or Arg (19). The reaction forms a reactive Schiff’s base which undergoes Amadori rearrangement, producing intermediates with highly reactive carbonyl groups, α-dicarbonyls, like glyoxal, methylglyoxal, and 3-deoxylglucosone (19–21). These α-dicarbonyl compounds can react with amino acids and proteins forming stable advanced glycation end products (AGEs) (20, 21). AGEs are important in contributing to malt and beer flavour and colour (19). Kilning during the malting process leads to the production of AGEs (20, 21), which may also continue during the mash and boil (10, 22, 23).

The proteomes of beer and the brewing process have been studied with a variety of techniques (10, 11, 24–28). However, their diverse and complex PTMs remain underexplored. Here, we used Data Independent Acquisition (DIA) / Sequential Window Acquisition of all THeoretical Mass Spectra (SWATH) LC-MS/MS with bioinformatic workflows for PTM identification and measurement to explore the underlying protein biochemistry of diverse commercial beers.

## Methods

### Sample preparation

23 unique commercial beers were purchased in Brisbane in December 2017 (Brewery A: Session Ale, Pale Ale, IPA, Porter, Amber Ale, Golden Ale; Brewery B: Lager-1, Lager-2, Lager-3, Pale Ale, Dark Ale, Porter; Brewery C: Pale Ale, IPA; Brewery D: Lager-1, Lager-2, Lager-3, Pale Ale-1, Pale Ale-2; Brewery E: Pale Ale; Brewery F: Pale Ale; Brewery G: IPA, XPA; Breweries B and D were classified as multinational, and Breweries A, C, E, F, and G were classified as independent). Samples of these beers were prepared for mass spectrometry in technical triplicate as previously described (10). Proteins from 50 μL of beer were precipitated by addition of 1 mL 1:1 methanol/acetone, incubation at −20°C for 16 h, and centrifugation at 18,000 rcf at room temperature for 10 min. The supernatant was discarded and proteins were digested by resuspension in 100 μL 100 mM ammonium acetate with 10 mM dithiothreitol and 0.5 μg trypsin (Proteomics grade, Sigma), and incubation at 37 °C with shaking for 16 h.

### Mass spectrometry

Peptides were desalted with C18 ZipTips (Millipore) and measured by LC-ESI-MS/MS using a Prominence nanoLC system (Shimadzu) and a TripleTof 5600 mass spectrometer with a Nanospray III interface (SCIEX) as previously described (29). Approximately 1 μg or 0.2 μg desalted peptides, were injected for data dependent acquisition (DDA) or data independent acquisition (DIA), respectively. Peptides were separated on a VYDAC EVEREST reversed-phase C18 HPLC column (300 Å pore size, 5 μm particle size, 150 μm i.d. x 150 mm) at a flow rate of 1 μl/min with a linear gradient of 10-60% buffer B over 14 min, with buffer A (1% acetonitrile and 0.1% formic acid) and buffer B (80% acetonitrile and 0.1% formic acid), for a total run time of 24 min per sample. LC parameters were identical for DDA and DIA, and DDA and DIA MS parameters were set as previously described (30).

### Data analysis

Peptides and proteins were identified using ProteinPilot 5.0.1 (SCIEX), searching against a database containing all high confidence proteins from transcripts from the barley genome (31) (GCA_901482405.1, downloaded 28 April 2017; 248,180 proteins), all predicted proteins from S288C *S. cerevisiae* (yeast) (Saccharomyces Genome Database (SGD), downloaded December 2017; 6,726 proteins), and contaminant proteins (custom database; 298 proteins), with settings: sample type, identification; cysteine alkylation, none; instrument, TripleTof 5600; species, none; ID focus, biological modifications; enzyme, trypsin; search effort, thorough ID.

Glycopeptides were identified by searching DDA files using Byonic (Protein Metrics, v. 2.13.2). Cleavage specificity was set as C-terminal to Arg/Lys and non-specific (either terminus could disagree), a maximum of two missed cleavages were allowed, and mass tolerances of 50 ppm and 75 ppm were applied to precursor and fragment ions, respectively. Variable modifications set as “Common 1” allowed each modification to be present on a peptide once and included deamidated Asn. Mono-oxidised Met was set as “Common 2”, which allowed the modification to be present twice on a peptide. To control search effort and false discovery rate, a maximum of two common modifications were allowed per peptide. To investigate the masses of modifications present on peptides, *Wildcard* searches were conducted against a focused protein FASTA file with the 138 barley and 47 yeast proteins identified by ProteinPilot search. The search allowed any mass between −40 and +200 on any peptide (including modified peptides). Once common modification masses had been deduced from the *Wildcard* searches, two specific PTM searches were conducted, one for yeast and one for barley with the additional parameters described below.

The FASTA protein file for the yeast search contained the S288C yeast proteome (SGD, downloaded December 2017; 6,726 proteins). The setting “Rare 1” was used, which allowed each modification to be present once on a peptide and included the *N*-linked monosaccharide compositions HexNAc_1-2_ and HexNAc_2_Hex_1-10_ at the consensus sequence N-X-S/T, and the *O*-linked monosaccharide compositions Hex_1_-Hex_10_ at any Ser or Thr residue (HexNAc, *N*-acetylhexosamine; Hex, hexose). A maximum of two common modifications and one rare modification were allowed per peptide. The focused FASTA protein file for the barley searches contained 138 barley proteins identified from ProteinPilot. The “Rare 1” setting included the monosaccharide compositions Hex_1_-Hex_10_ at any Lys residue. A maximum of two common modifications and one rare modification were allowed per peptide. Unique yeast glycopeptides and barley glycated peptides identified by the Byonic searches were manually inspected and validated (Validated spectra in Supplementary Material S1-Glycosylation and S2-Glycation). The highest confidence unique glycopeptide identifications across all searches were used to create a glycopeptide library for DIA/SWATH-MS analyses as previously described (32).

Peptides identified by the ProteinPilot search were combined with glycopeptides identified by the Byonic searches to create one ion library. The abundance of peptide fragments, peptides, and proteins was determined using PeakView 2.2 (SCIEX), with settings: shared peptides, allowed; peptide confidence threshold, 99%; false discovery rate, 1%; XIC extraction window, 6 min; XIC width, 75 ppm. Identified barley proteins were matched against UniProtKB (downloaded 2 December 2017; 555,318 total entries), using BLAST+ as previously described (33). GlypNirO, a previously described Python script was modified and used to calculate the occupancy and proportion of each glycan at each site/peptide (32). A Python script, ClipNirO, was developed to calculate the proportion of the abundance of each physiological proteolytic peptide matched to each full tryptic peptide (https://github.com/bschulzlab/proteolysis-normalization, Supplementary Material S3). For protein-centric analyses, protein abundances were re-calculated by removing all peptide intensities that did not pass an FDR cut-off of 1% using a Python script as previously described (33). Protein abundances were recalculated as the sum of all peptides from that protein and normalised to either the total protein abundance in each sample or to the abundance of trypsin self-digest peptides, as previously described (10). Principal component analysis (PCA) was performed using Python, the machine learning library Scikit-learn (0.19.1), and the data visualisation package Plotly (1.12.2). Protein and sample clustering was performed using Cluster 3.0 (34), implementing a hierarchical, uncentered correlation, and complete linkage.

### Foam and pouring measurement

Beers from Brewery A were analysed for foam-related paraments using an automated robotic pourer, RoboBEER, adapted for canned beers, as previously described (35), with data processed as described (36). RoboBEER is able to pour 80 ± 10 mL of beers in a constant manner while videos are recorded using a smartphone camera to be further analysed using computer vision algorithms developed in Matlab R2018b (Mathworks, Inc). Parameters including maximum volume of foam, total lifetime of foam and bubble size distribution in the foam classified as small, medium and large were obtained.

### Data availability

The mass spectrometry proteomics data have been deposited to the ProteomeXchange Consortium via the PRIDE partner repository with the dataset identifier PXD023116 (username, password: 5QTcVlLc) (19).

## Results and Discussion

### Identification and quantification of beer protein PTMs

We set out to use DIA/SWATH-MS to profile the proteomes of diverse Australian commercial beers, with the aim of identifying molecular markers that could distinguish beer styles and which contribute to beer sensory qualities. We obtained 23 diverse Australian commercial beers and performed LC-MS/MS bottom-up proteomics with DDA for identification and DIA/SWATH for quantification in technical triplicate. Using ProteinPilot to search a combined yeast and barley high confident protein database, we identified 49 yeast and 139 barley proteins. These modest numbers of identified proteins are broadly consistent with previous reports of the beer proteome (11, 25). However, we identified extensive physiological non-tryptic proteolysis that extensively expanded the complexity of the beer proteome. We measured 405 unique semi- and non-tryptic proteolytic cleavage events in 136 barley proteins within beer (Supplementary Table S1). In the most abundant protein identified, non-specific lipid transfer protein 1 (NLTP1), we found 25 unique physiological cleavage events, and in an abundant glutenin subunit protein (GLT3) we found 68 cleavage events (Supplementary Table S1). This extensive proteolytic clipping of proteins most likely occurs during the mashing stage of beer production, where it controls the stability of proteins and hence their concentration in the final beer (10). Nonetheless, we were surprised to measure such a high diversity of proteolytically defined proteoforms in beer.

Recent analyses of the sparkling wine proteome identified abundant glycoproteins from yeast (13), which we suspected might also be present in beer, as it is also a fermented beverage most commonly made using *S. cerevisiae*. Other PTMs have also been previously qualitatively reported on beer proteins, including abundant physiological proteolysis and glycation (10, 21, 23, 37, 38). Indeed, our inspection of LC-MS/MS data of tryptic digests of beer proteomes identified abundant putative glycosylation events with oligohexose, consistent with either oligomannose *O*-glycosylation of yeast proteins or maltooligosaccharide glycation of barley proteins. To identify peptides modified with glycosylation or other PTMs we used a Byonic and DIA/SWATH quantification workflow (32). Byonic wildcard searches identified modification of yeast proteins with oligoHex on Ser/Thr consistent with *O*-mannosylation, and of barley proteins with oligoHex on Lys consistent with non-enzymatic glycation. We did not detect the presence of any other PTMs by these wildcard searches.

We identified extensive glycation events on barley proteins that expanded the complexity of the beer proteome. We measured 111 unique glycation events on 18 barley proteins within beer (Supplementary Table S2). Interestingly, 56 of the 111 glycation events were found on a single protein, NLTP1. We identified oligoHex glycans on lysines, ranging from Hex_1_ to Hex_7_, with Hex_1_ and Hex_2_ being the most common with 52 and 37 events respectively on different peptides (Supplementary Table S2). The extensive glycation we identified likely results from Maillard reactions between proteins and maltooligosaccharides derived from beta- and alpha-amylase digestion of starch during malting, mashing, and boiling. In addition to glycation we found extensive glycosylation events on yeast proteins in beer. Inclusion of Byonic database search strategies to identify glycosylated peptides allowed us to identify many proteins not identified with a standalone ProteinPilot workflow (Supplementary Table S3). We identified 21 unique glycosylated proteins with a total of 60 unique glycosylation events (Supplementary Table S3). OligoHex *O*-glycans ranging from Hex_1_ to Hex_8_ were the most common modification identified, along with a single HexNAc_1_ *N*-glycosylation event (Supplementary Table S3). The HexNAc_1_ modification was detected at an *N*-glycosylation sequon on peptide R_50_-CDTLVG<u>N_57_LT</u>IGGGLK_65_-T from yeast cell wall mannoprotein Pst1. Yeast *N*-glycans are typically high mannose structures (30), and so the structure we identified on Pst1 is consistent with extensive glycosidase trimming by unknown enzymes during fermentation or beer storage. Together, these results showed that, as with sparkling wine, there was extensive glycosylation on secreted yeast proteins in beers (13), and emphasized the value of considering PTMs such as glycosylation, glycation, and proteolysis in discovery and quantitative LC-MS/MS proteomics workflows.

### The PTM profile of commercial beers

To include proteolysis, glycosylation, and glycation PTMs in our proteomic analysis, and to quantify the site-specific extent of these modifications, we constructed glyco/peptide DIA/SWATH ion libraries via a ProteinPilot/Byonic-to-Peakview workflow (13, 32) (Fig. 1A). We used the previously described GlypNirO workflow to measure site-specific glycosylation and glycation (32), and developed and used ClipNirO, a modified version of this workflow to measure physiological proteolytic events mapped to full tryptic peptides (Supplementary Material S3).

**Figure 1.**
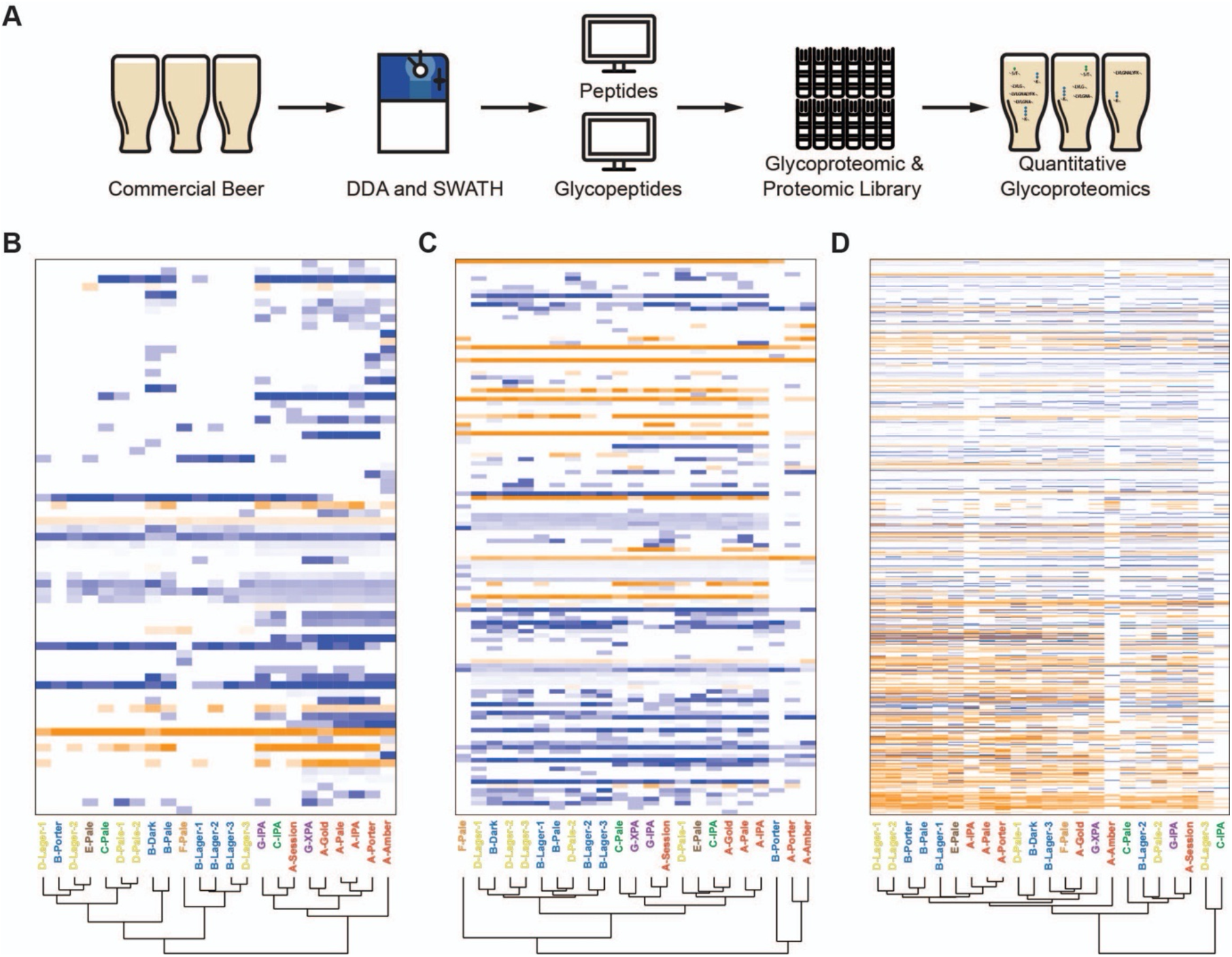
Site-specific glycosylation, glycation, and proteolysis differentiate beer proteomes. **(A)** Overview of the workflow for identification, measurement, and site-specific analysis of modified and unmodified peptides for quantitative glyco/proteomics. Clustered heat map of normalised site-specific **(B)** glycosylation, **(C)** glycation, and **(D)** proteolysis. Modified peptides were normalised to their peptide family. Unmodified peptides shown in orange, modified forms shown directly below in blue. Dendrograms show sample clustering by brewery (A-G) and beer style. Heatmaps use log_10_ protein abundance normalised to total protein abundance in each sample.

We used GlypNirO to measure site-specific modification at 60 unique glycopeptides from yeast proteins (Fig. 1B and Supplementary Table S4). Clustered heatmap analysis showed two main clades clustered by brewery, with abundant yeast *O*-glycopeptides in beers from independent breweries A, G, and C, less abundant yeast *O*-glycopeptides in beers brewed by multinational companies and larger breweries B, D, and F (Fig. 1B). This differentiation of beers by manufacturer based on the yeast *O*-glycoproteome is likely due to differences in the yeast strains used; larger breweries tend to use their own proprietary in-house yeast strains, whereas independent breweries commonly use commercially sourced yeast. Lager and ale yeast may also contribute to this difference, since our analysis did not include a lager produced by an independent brewery. However, there was no consistent separation between lagers and ales from multinational breweries based on their yeast *O*-glycoproteome (Fig. 1B), suggesting any difference between the *O*-glycoproteomes of lager and ale yeasts was not a key differentiator.

We next used GlypNirO to compare site-specific glycation from barley proteins in our set of 23 beers (Fig. 1C and Supplementary Table S5). This analysis identified a predominant clustering based on beer style. Nested within the clade consisting of the majority of beers, two sub-clades resolved, lagers and ales (Fig. 1C). However, as we did not analyse a lager from an independent brewery it is unclear if this differentiation is driven by style or manufacture scale. Separation was observed between B-Porter, A-Porter, and A-Amber from the other lighter coloured lagers and ales (Fig. 1C). These darker beers contained minimal glycated peptides (Fig. 1C and Supplementary Table S5). Dark beers are made from a combination of base malt (usually pale malt) and specialty malts including roasted malts. Roasted malts are roasted at high temperatures after kilning to imbue them with dark colours and rich flavours (1). This roasting process should cause most glycated amino acids that were formed during kilning to be degraded, resulting in very little glycation on either free amino acids or proteins (20). However, these dark beers should still contain substantial amounts of pale malt, and it is therefore unclear why this is not reflected in the presence of glycation in their glycoproteomes. Additional processes or interactions from the components of roasted malt may be at play in controlling the dark beer proteome.

Using ClipNirO, we next investigated the variance within the proteolytic proteome (Fig. 1D and Supplementary Table S6). Clustering based on site-specific proteolysis did not identify any obvious characteristics, although some contribution from brewery was apparent (Fig. 1D). Extent of proteolysis is likely caused by several factors including the mash program and amount of protein-rich base malt used. The lack of obvious clustering is likely due to variation in mash program and malt used between brewery and style.

To compare the global contribution of the glyco/proteome to variance between diverse commercial beers, we compared the base proteome and the integrated glyco/proteome (Fig. 2A – D). This comparison showed that it was necessary to integrate glycosylation, glycation, and proteolysis PTMs for complete proteomic analyses, as including measurement of these modifications (Fig. 2B and D) altered the measured proteomes of individual beers and changed how they clustered relative to beers of different styles and breweries. PCA of the integrated glyco/proteome showed clear clustering of beers by brewery (Fig. 2B), apart from the outlier darker beers B-Porter, A-Porter, and A-Amber, as well as F-Pale. Beers from breweries A and G clustered together and separated almost completely from the other breweries (Fig. 2B). Beers from breweries B, D, and E also clustered together. Closer inspection of clustering based on glyco/proteomes showed that within each manufacturer, beers tended to cluster based on style. For example, the three lagers from brewery B clustered closely and separately from the pale ale, dark ale, and porter from the same brewery (Fig. 2D). Together, this global glyco/proteomic analysis of commercial beers showed the key molecular differentiators were brewery and beer style, likely reflecting a combination of differences including the malt bill, mashing parameters, and the yeast used for fermentation.

**Figure 2.**
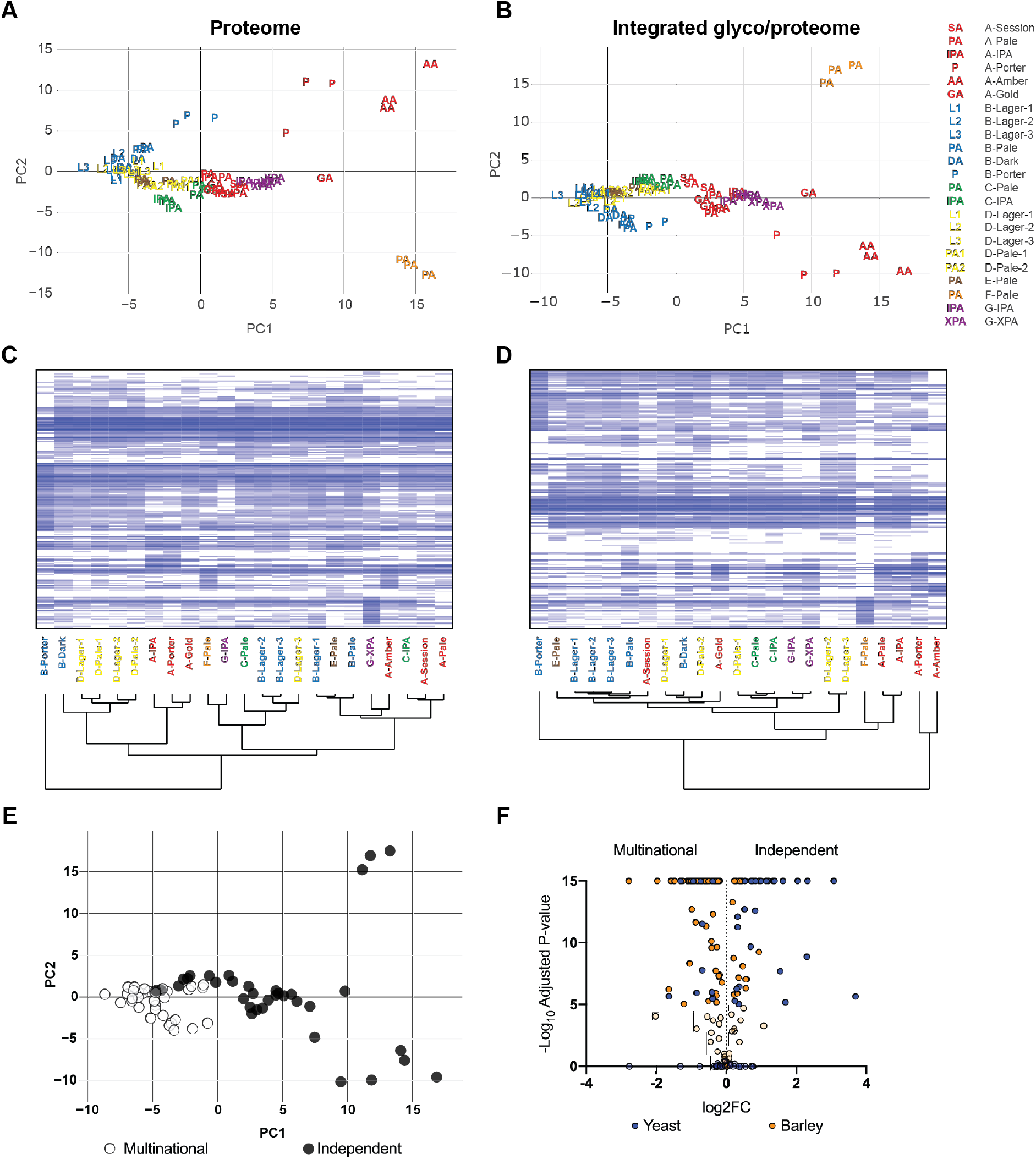
The global glyco/proteomes of commercial beers. **(A)** PCA of beer proteomes without or **(B)** with consideration of glycation and glycosylation. **(C)** Clustered heat map of beer proteomes without or **(D)** with consideration of glycation and glycosylation. Dendrograms show sample clustering by brewery (A-G) and beer style. PCAs and heatmaps use log_10_ protein abundance normalised to total protein abundance in each sample. (E) PCA of beer glyco/proteomes as in (B) but with beers classified as either produced by multinational (white circles) or independent (black circles) breweries. **(F)** Volcano plot of the comparison of the glyco/proteomes of beers produced by multinational or independent breweries. Blue, yeast proteins; orange, barley proteins; 100% opacity, significantly different proteins; 20% opacity, non-significantly different proteins; positive Log_2_FC, significantly more abundant in beers from independent breweries; negative Log_2_FC, significantly more abundant in beers from multinational breweries.

Inspection of PCA of beer glyco/proteomes (Fig. 2B) suggested that there was a clear distinction between beers from independent and multinational breweries. Beers from multinational breweries clustered tightly, with little variance, while in contrast beers from independent breweries separated from multinational beers and showed substantial internal variance (Fig. 2E). To understand the underlying bases for this distinction between independent and multinational breweries we directly compared the glyco/proteomes of beers of all styles from each group. This analysis showed that many proteins were significantly different between multinational and independent beers (Fig. 2F and Supplementary Table S7). A key difference was the overall relative abundance of yeast and barley proteins, with 26 yeast proteins significantly more abundant in independent beers and only 11 more abundant in multinational beers, but only 22 barley proteins significantly more abundant in independent beers and 73 more abundant in multinational beers (Fig. 2F and Supplementary Table S7). Critically, most of these differentially abundant yeast proteins were heavily *O*-glycosylated seripauperins that were only identifiable as glycopeptides using our integrated glyco/proteomic workflow. The differentiation of beers produced by multinational and independent breweries by global glyco/proteome agreed with the clustering observed in our site-specific glycosylation analysis (Fig. 1B) and is consistent with differences in yeast strains used for fermentation by these breweries, or the lack of lagers made by independent breweries.

### Understanding the nuanced proteomic differences of beers from the same brewery

Our global glyco/proteomic analysis of diverse commercial beers showed they tended to primarily cluster by brewery. In order to investigate in more detail how beer style affected the glyco/proteome we therefore focussed on six beers from a single brewery (Brewery A), Newstead Brewing Co. from Brisbane, Australia: Pale Ale, India Pale Ale, Session Ale, Golden Ale, Amber Ale, and Porter (Fig. 3). We analysed the complete glyco/proteome and site-specific glycosylation, glycation, and proteolysis profiles of these beers. Based on PCA and clustered heat map analysis of the global glyco/proteome we observed close clustering of Pale Ale and IPA, together with Golden Ale and Session Ale, while the Porter and Amber Ale were clearly distinct from each other and the other beers (Fig. 3A and B). This clustering correlates with the amount of pale malt used in each beer; above 80% of the total malt in the Session Ale, Pale Ale, IPA, and Golden Ale, but much lower in the Porter or Amber Ale.

**Figure 3.**
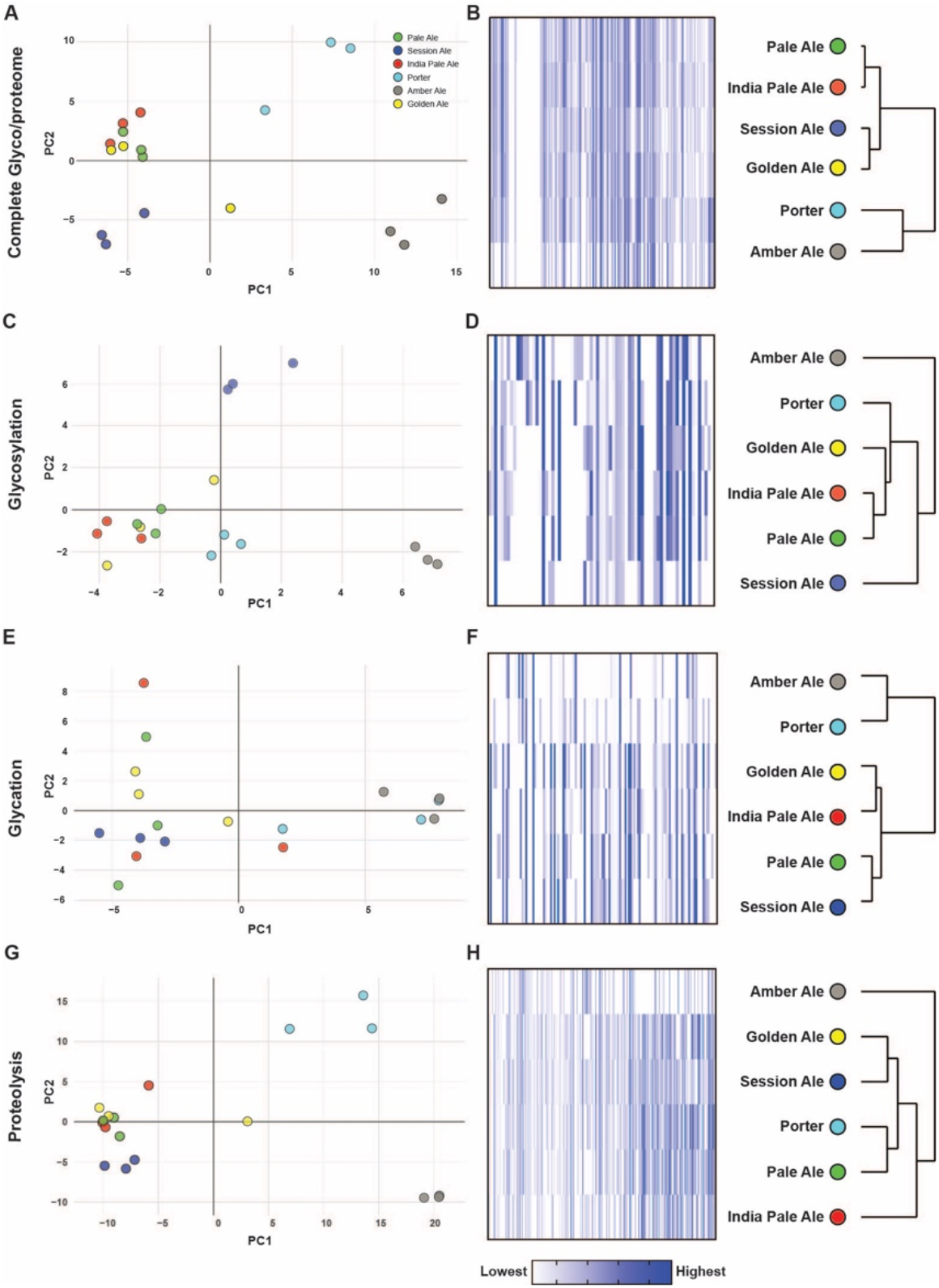
Analysis of beers from Newstead Brewing Co. **(A)** PCA and **(B)** heat map of the complete glyco/proteome. **(C)** PCA and **(D)** Heat map of the site-specific glycoproteome. **(E)** PCA and **(F)** heat map of the site-specific glycated proteome. **(G)** PCA and **(H)** heat map of the site-specific proteolytic proteome. PCAs and heatmaps use log_10_ protein abundance normalised to total protein abundance in each sample. Dendrograms show sample clustering by beer style.

We next focussed on site-specific modification profiles. As with the global glyco/proteome, analysis of site-specific glycosylation profiles showed clustering of Pale Ale, IPA, and Golden Ale, and separation of Porter and Amber Ale (Fig. 3C and D). In contrast, Session Ale showed clear separation from other beers based on site-specific glycosylation. To understand the reasons for this difference, we calculated the total extent of glycosylation as the total abundance of yeast glycopeptides, and the average site-specific glycosylation occupancy in each beer (Fig. 4A – C). This showed that there were no differences in site-specific glycosylation occupancy or glycan composition in Session Ale compared to the other beers (Fig. 4B), exemplified by the glycoforms of R-V_94_ITGVPWYSTR_104_-L from Pau15 (Fig. 4C), but rather that there was low overall abundance of *O*-glycosylated seripauperins from yeast in this beer (Fig. 4A and Supplementary Table S8).

**Figure 4.**
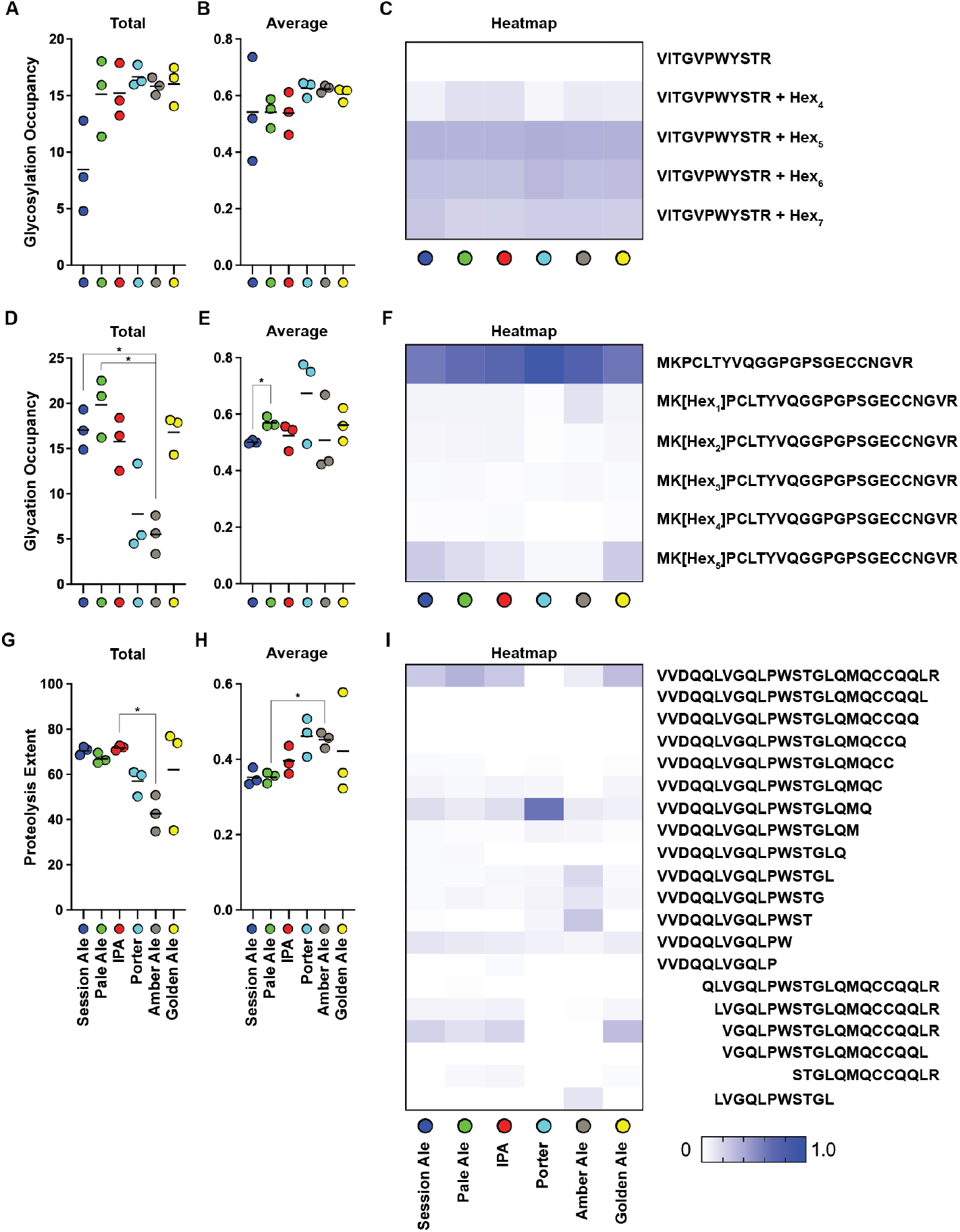
Site-specific occupancy and overall extent of PTMs in beers from Newstead Brewing Co. **(A)** Total and **(B)** average glycosylation occupancy. **(C)** Heat map of normalised abundance of unmodified and glycosylated forms of tryptic peptide R-V_94_ITGVPWYSTR_104_-L from Pau15. Hex modification location was not determined. **(D)** Total and **(E)** average glycation occupancy. **(F)** Heat map of normalised abundance of unmodified and glycated forms of tryptic peptide K-M_36_KPCLTYVQGGPGPSGECCNGVR_58_-S from NLTP1. (G) Total and **(H)** average extent of proteolysis. **(I)** Heat map of normalised abundance of unmodified and proteolytically cleaved forms of full tryptic peptide R-V_56_VDQQLVGQLPWSTGLQMQCCQQLR_80_-D from GLT3. Column graphs were of either total of average PTM occupancy/extent, calculated as either sum or average of all modified peptides normalised to their peptide family, respectively. Heat maps show modified peptides normalised to their peptide family. Bars show mean, n=3, Error bars show SEM, * indicates statistically significant (p < 0.05).

Site-specific glycation profile analysis clustered beers similarly to the global glyco/proteome, with Amber Ale and Porter separating from the other beers (Fig. 3E and F). Although peptides in Amber Ale and Porter were glycated at equivalent occupancy to other beers (Fig. 4D and F and Supplementary Table S9), the differentiation of Amber Ale and Porter was due to the significantly lower levels of detectable glycated peptides in these beers (Fig. 4D). Session Ale and Pale Ale were significantly different in site-specific glycation, but the difference was small (Fig. 4E). These results showed that the levels of glycated proteins were low in beers with a darker malt profile, as observed in the dark beer proteomes from other breweries (Fig. 1C).

Finally, we investigated the site-specific extent of proteolysis. Again, Amber Ale and Porter were separated from other beers, which clustered together (Figure 3G and H). Normalised to within each peptide group, Porter and Amber Ale had high levels of proteolysis (Fig. 4H). For example, cleaved peptide R-V_56_VDQQLVGQLPWSTGLQMQ_74_-C from GLT3 was significantly more abundant in Porter compared to all other beers (Fig. 4I and Supplementary Table 10), and cleaved peptide R-V_56_VDQQLVGQLPWST_69_-G was significantly more abundant in Amber Ale compared to all other beers besides Porter (Fig. 4I and Supplementary Table 10). In contrast, the full tryptic peptide that had not been subjected to proteolytic clipping had lower relative abundance in Porter and Amber Ale. Despite their increased extent of site-specific proteolysis, Porter and Amber Ale had lower total abundance of proteolytically clipped peptides, reflecting the lower overall abundance of proteolytically clipped proteins (Fig. 4G). In summary, we observed a high extent of site-specific proteolysis in dark beers, but with low overall levels of proteolytically clipped proteins. These observations are consistent with the increased proteolysis during the mash in these darker beers destabilising proteins, resulting in their loss from the finished beer (10).

### Glyco/proteome correlates of foam formation and stability

Barley non-specific Lipid Transfer Proteins (NLTPs), serpins, and yeast seripauperins have all been previously reported to be important for foam formation and stability (15, 16, 39–41), and are also heavily modified by glycosylation and glycation (16, 17, 23, 42) (Fig. 1). We implemented the previously described RoboBEER workflow (35) to obtain quantitative parameters describing foam and bubble formation and stability of the selected beers from Newstead Brewing Co, and correlated these foam characteristics with the abundance of NLTPs, serpins, and seripauperins using linear regression (Fig. 5). We tested for correlations between the abundance of each of these classes of protein and the: maximum volume of foam produced (mL); total lifetime of foam (s); foam drainage (mL s^−1^); and the number of small, medium, and large bubbles produced in the foam. Only a very few significant linear relationships were found between protein abundance and foaming characteristics. Seripauperin abundance was significantly negatively correlated with the total lifetime of foam (R^2^ = 0.71) and with the number of small bubbles (R^2^ = 0.38) (Fig. 5 and Supplementary Table S11). That is, low levels of seripauperins were associated with a large amount of stable foam with small bubbles. Although not significant, NLTP levels showed a trend towards a positive correlation with these foam characteristics, while we found no evidence of a correlation between serpins and foam properties.

**Figure 5.**
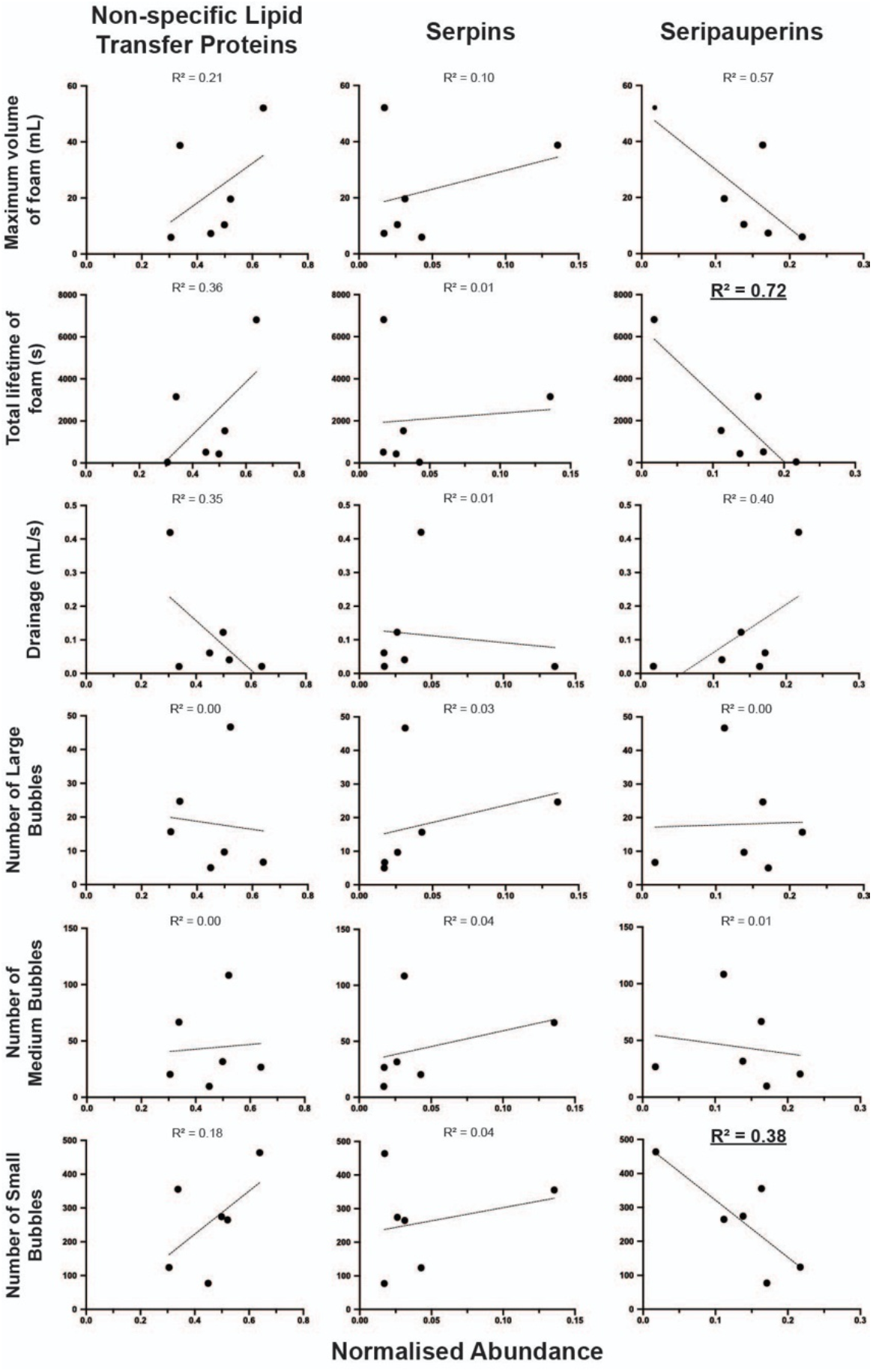
Linear regression analysis of barley and yeast glycoprotein abundance and beer foam characteristics. Correlations of summed normalised abundance of Non-specific Lipid Transfer Proteins (NLTPs), Serpins, and Seripauperins with: maximum volume of foam; total lifetime of foam; foam drainage; and number of large, medium, small bubbles. Values, mean. Line, linear regression with coefficient of determination, R^2^. Bold and underlined, significant (p < 0.05).

## Conclusion

We found extensive PTM complexity and diversity in the proteomes of commercial beers, especially proteolysis from barley proteases, *O*-glycosylation of secreted yeast glycoproteins, and glycation of barley proteins with maltooligosaccharides. The beer glyco/proteome was defined primarily by brewery and then by beer style, suggesting that manufacturing process parameters are a key contributor to the beer proteome. We also identified substantial differences between beers produced by multinational and independent breweries driven predominantly by the contribution of yeast proteins and glycoproteins. Key foam quality parameters correlated with features of the beer glyco/proteome, especially the abundance of *O*-glycosylated yeast seripauperins, confirming the importance of the proteome and its PTMs in determining the quality of beer, and emphasising that yeast is not only of critical importance in producing alcohol and flavours in beer, but is also critical for controlling the highly PTM-modified beer proteome.

## Acknowledgements

We thank Dr Amanda Nouwens and Peter Josh at The University of Queensland, School of Chemistry and Molecular Biosciences Mass Spectrometry Facility for their assistance and expertise. Edward D. Kerr was funded by an Advance Queensland PhD Scholarship. Christopher H. Caboche was funded by a Wine Australia PhD Scholarship.

