## Supplementary Material S1 for "The post-translational modification landscape of commercial beers"

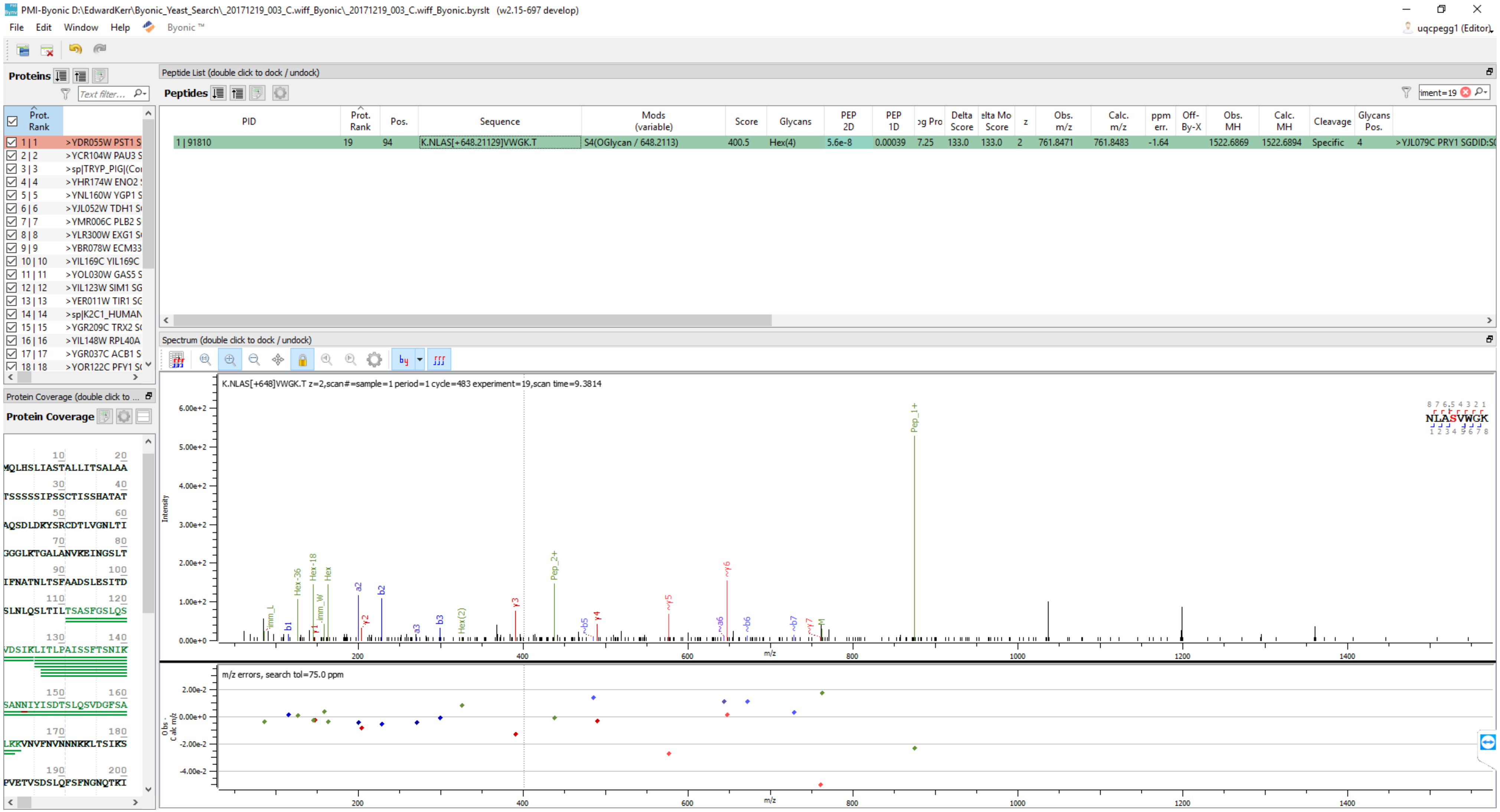
PMI-Byonic D:\EdwardKerr\Byonic_Yeast_Search_20171219_003_C.wiff_Byonic_20171219_003_C.wiff_Byonic.byrslt (w2.15-697 develop)
File Edit Window Help Byonic ™
uqcpegg1 (Editor)
Proteins
Peptide List (double click to dock / undock)
Peptides
PID
Prot. Rank
Pos.
Sequence
Mods (variable)
Score
Glycans
PEP 2D
PEP 1D
Delta Score
z
Obs. m/z
Calc. m/z
ppm err.
Off-By-X
Obs. MH
Calc. MH
Cleavage
Glycans Pos.
1 | 91810
19
94
K.NLAS[+648.21129]VWGK.T
S4(OGlycan / 648.2113)
400.5
Hex(4)
5.6e-8
0.00039
7.25
133.0
133.0
2
761.8471
761.8483
-1.64
1522.6869
1522.6894
Specific
4
>YJL079C PRY1 SGDID:S
1 | 1 >YDR055W PST1 S
2 | 2 >YCR104W PAU3 S
3 | 3 >sp|TRYP_PIG|(Co
4 | 4 >YHR174W ENO2
5 | 5 >YNL160W YGP1 S
6 | 6 >YJL052W TDH1 S
7 | 7 >YMR006C PLB2 S
8 | 8 >YLR300W EXG1 S
9 | 9 >YBR078W ECM33
10 | 10 >YIL169C YIL169C
11 | 11 >YOL030W GAS5 S
12 | 12 >YIL123W SIM1 SG
13 | 13 >YER011W TIR1 SG
14 | 14 >sp|K2C1_HUMAN
15 | 15 >YGR209C TRX2 S
16 | 16 >YIL148W RPL40A
17 | 17 >YGR037C ACB1 S
18 | 18 >YOR122C PFY1 S
Protein Coverage
MQLHSLIASTALLITSALAA
TSSSSSIPSSCTISSHATAT
AQSDLDKYSRCDTLVGNLTI
GGLKTGALANVKEINGSLT
IFNATNLTSFAADSLESITD
SLNLQSLTILTSASFGSLQS
VDSIKLITLPAISSFTSNIK
SANNIYISDTSLQSVDGFSA
LKKVNVFNVNNNKKLTSIKS
PVETVSDSLQFSFNGNQTKI
Spectrum (double click to dock / undock)
K.NLAS[+648]VWGK.T z=2,scan#=sample=1 period=1 cycle=483 experiment=19,scan time=9.3814
Intensity
m/z
NLASVWGK
Pep_1+
Pep_2+
Hex-36
Hex-18
Hex
Hex(2)
a2
b2
b1
b3
a3
y2
y1
y3
y4
~y5
~y6
~a6
~b5
~b6
~b7
~y7
m/z errors, search tol=75.0 ppm
Obs. - Calc m/z

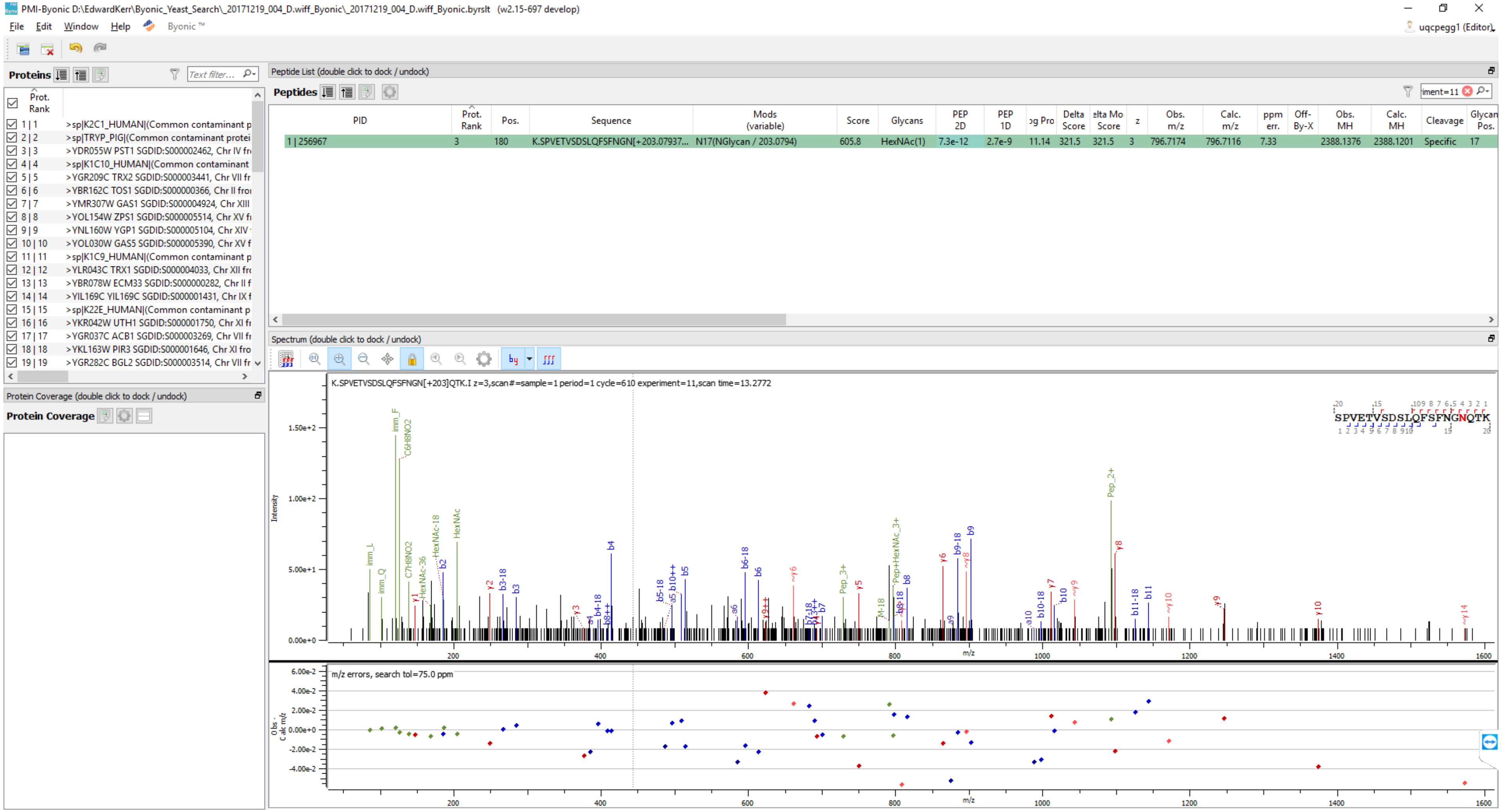
PMI-Byonic D:\EdwardKerr\Byonic_Yeast_Search_20171219_004_D.wiff_Byonic_20171219_004_D.wiff_Byonic.byrslt (w2.15-697 develop)
File Edit Window Help Byonic ™
uqcpegg1 (Editor)
Proteins
Peptide List (double click to dock / undock)
Peptides
PID
Prot. Rank
Pos.
Sequence
Mods (variable)
Score
Glycans
PEP 2D
PEP 1D
Delta Score
Obs. m/z
Calc. m/z
ppm err.
Off-By-X
Obs. MH
Calc. MH
Cleavage
Glycan Pos.
1 | 256967 3 180 K.SPVETVSDSLQFSFNGN[+203.07937... N17(NGlycan / 203.0794) 605.8 HexNAc(1) 7.3e-12 2.7e-9 11.14 321.5 321.5 3 796.7174 796.7116 7.33 2388.1376 2388.1201 Specific 17
>sp|K2C1_HUMAN|(Common contaminant p
>sp|TRYP_PIG|(Common contaminant protei
>YDR055W PST1 SGDID:S000002462, Chr IV fr
>sp|K1C10_HUMAN|(Common contaminant
>YGR209C TRX2 SGDID:S000003441, Chr VII fr
>YBR162C TOS1 SGDID:S000000366, Chr II fro
>YMR307W GAS1 SGDID:S000004924, Chr XIII
>YOL154W ZPS1 SGDID:S000005514, Chr XV fr
>YNL160W YGP1 SGDID:S000005104, Chr XIV
>YOL030W GAS5 SGDID:S000005390, Chr XV f
>sp|K1C9_HUMAN|(Common contaminant p
>YLR043C TRX1 SGDID:S000004033, Chr XII fr
>YBR078W ECM33 SGDID:S000000282, Chr II f
>YIL169C YIL169C SGDID:S000001431, Chr IX f
>sp|K22E_HUMAN|(Common contaminant p
>YKR042W UTH1 SGDID:S000001750, Chr XI fr
>YGR037C ACB1 SGDID:S000003269, Chr VII fr
>YKL163W PIR3 SGDID:S000001646, Chr XI fro
>YGR282C BGL2 SGDID:S000003514, Chr VII fr
Protein Coverage (double click to dock / undock)
Protein Coverage
Spectrum (double click to dock / undock)
K.SPVETVSDSLQFSFNGN[+203]QTK.I z=3,scan#=sample=1 period=1 cycle=610 experiment=11,scan time=13.2772
SPVETVSDSLQFSFNGNQTK
Intensity
m/z
Pep_2+
HexNAc
HexNAc-18
Pep+HexNAc_3+
m/z errors, search tol=75.0 ppm
Obs - Calc m/z

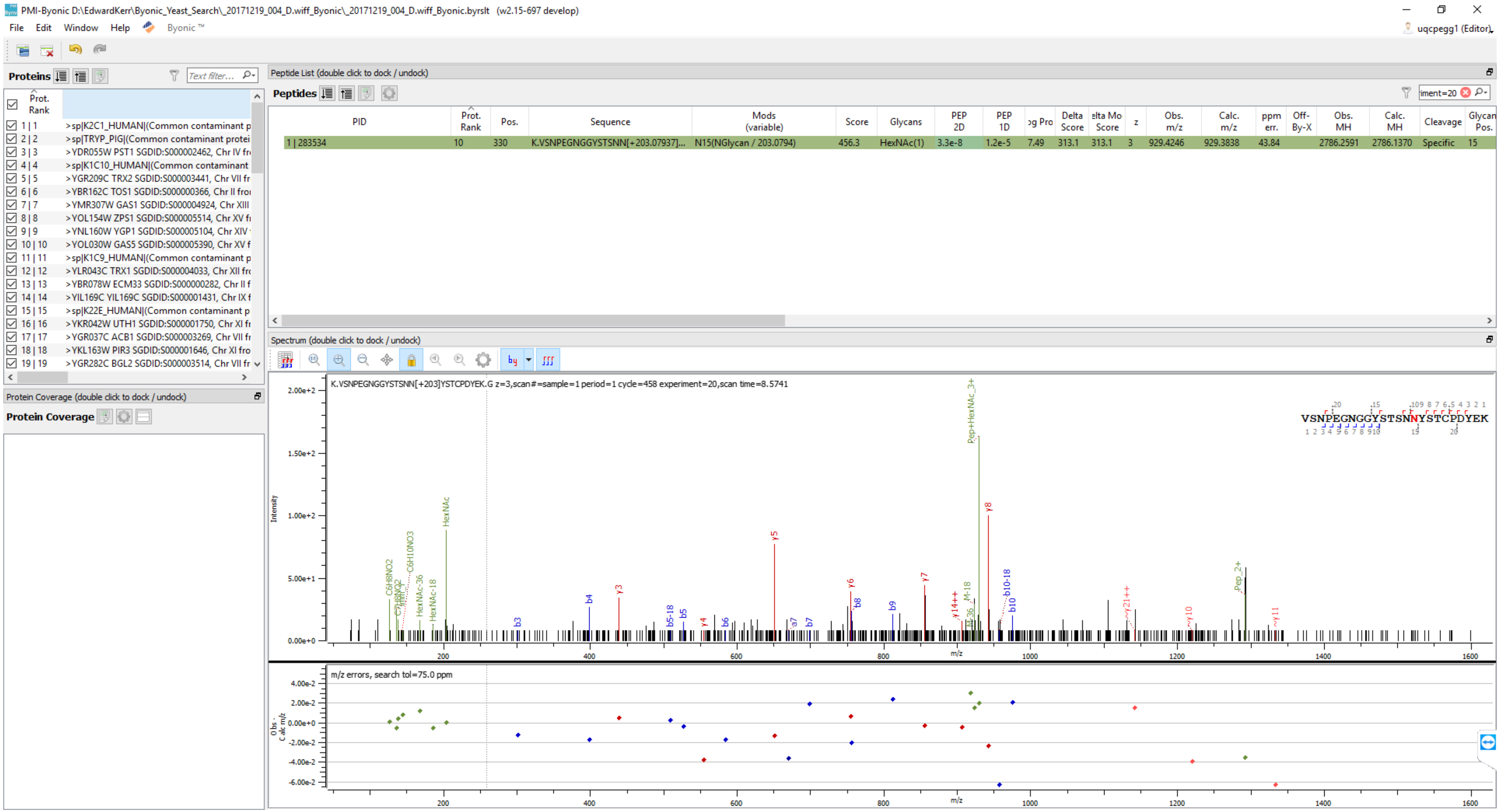
PMI-Byonic D:\EdwardKerr\Byonic_Yeast_Search_20171219_004_D.wiff_Byonic_20171219_004_D.wiff_Byonic.byrslt (w2.15-697 develop)
File Edit Window Help Byonic ™
uqcpegg1 (Editor)
Proteins
Text filter...
Prot. Rank
1 | 1 >sp|K2C1_HUMAN|(Common contaminant p
2 | 2 >sp|TRYP_PIG|(Common contaminant protei
3 | 3 >YDR055W PST1 SGDID:S000002462, Chr IV fr
4 | 4 >sp|K1C10_HUMAN|(Common contaminant
5 | 5 >YGR209C TRX2 SGDID:S000003441, Chr VII fr
6 | 6 >YBR162C TOS1 SGDID:S000000366, Chr II fro
7 | 7 >YMR307W GAS1 SGDID:S000004924, Chr XIII
8 | 8 >YOL154W ZPS1 SGDID:S000005514, Chr XV fr
9 | 9 >YNL160W YGP1 SGDID:S000005104, Chr XIV
10 | 10 >YOL030W GAS5 SGDID:S000005390, Chr XV f
11 | 11 >sp|K1C9_HUMAN|(Common contaminant p
12 | 12 >YLR043C TRX1 SGDID:S000004033, Chr XII fr
13 | 13 >YBR078W ECM33 SGDID:S000000282, Chr II f
14 | 14 >YIL169C YIL169C SGDID:S000001431, Chr IX f
15 | 15 >sp|K22E_HUMAN|(Common contaminant p
16 | 16 >YKR042W UTH1 SGDID:S000001750, Chr XI fr
17 | 17 >YGR037C ACB1 SGDID:S000003269, Chr VII fr
18 | 18 >YKL163W PIR3 SGDID:S000001646, Chr XI fro
19 | 19 >YGR282C BGL2 SGDID:S000003514, Chr VII fr
Protein Coverage (double click to dock / undock)
Protein Coverage
Peptide List (double click to dock / undock)
Peptides
PID | Prot. Rank | Pos. | Sequence | Mods (variable) | Score | Glycans | PEP 2D | PEP 1D | og Pro | Delta Score | elta Mo Score | z | Obs. m/z | Calc. m/z | ppm err. | Off-By-X | Obs. MH | Calc. MH | Cleavage | Glycan Pos.
1 | 283534 | 10 | 330 | K.VSNPEGNGGYSTSNN[+203.07937]... | N15(NGlycan / 203.0794) | 456.3 | HexNAc(1) | 3.3e-8 | 1.2e-5 | 7.49 | 313.1 | 313.1 | 3 | 929.4246 | 929.3838 | 43.84 | | 2786.2591 | 2786.1370 | Specific | 15
Spectrum (double click to dock / undock)
K.VSNPEGNGGYSTSNN[+203]YSTCPDYEK.G z=3,scan#=sample=1 period=1 cycle=458 experiment=20,scan time=8.5741
VSNPEGNGGYSTSNNYSTCPDYEK
Intensity
m/z
Pep+HexNAc_3+
HexNAc
y8
y5
m/z errors, search tol=75.0 ppm
Obs - Calc m/z

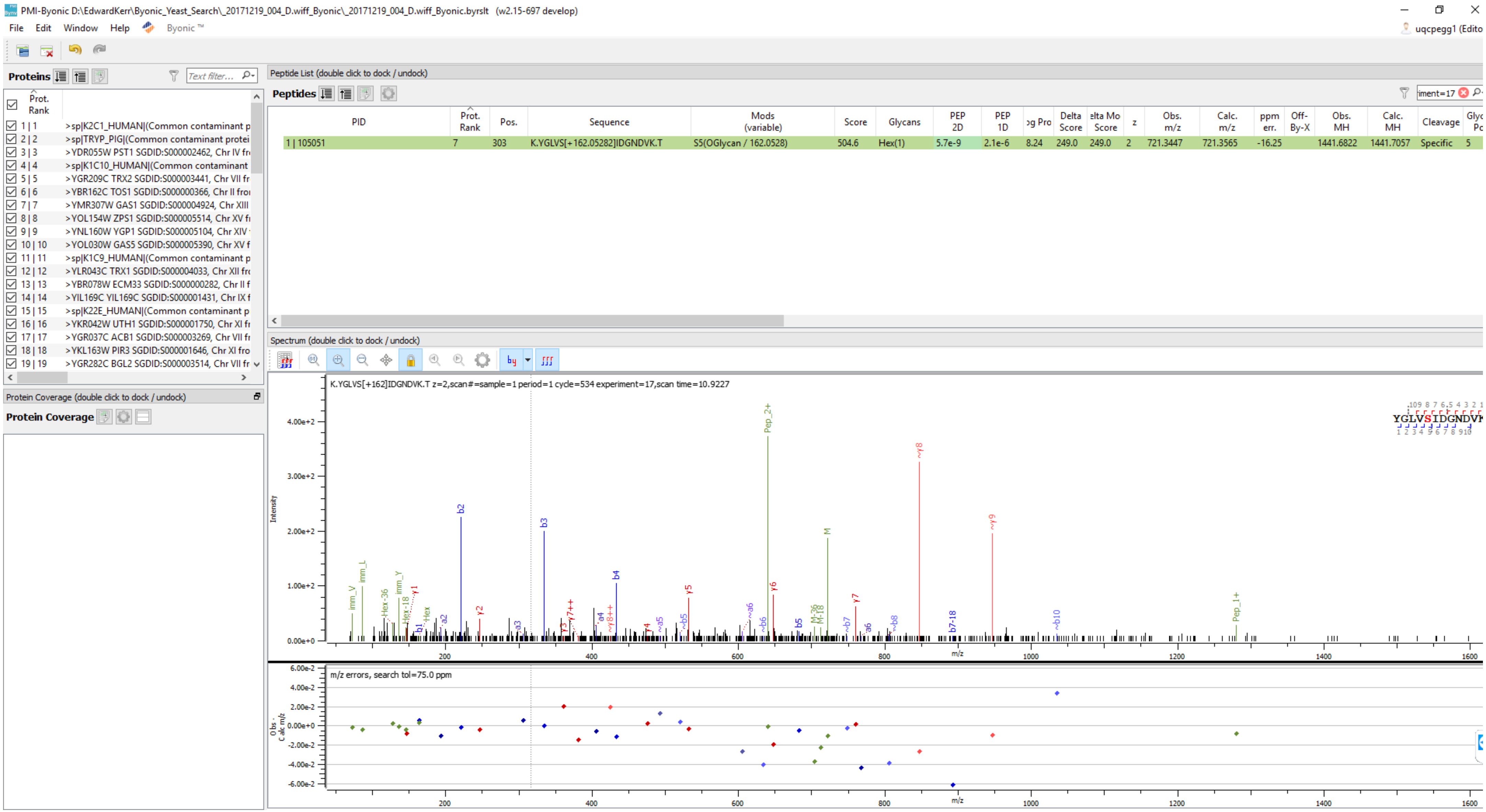
PMI-Byonic D:\EdwardKerr\Byonic_Yeast_Search_20171219_004_D.wiff_Byonic_20171219_004_D.wiff_Byonic.byrslt (w2.15-697 develop)
File Edit Window Help Byonic™
Proteins
Prot. Rank
1 | 1 >sp|K2C1_HUMAN|(Common contaminant p
2 | 2 >sp|TRYP_PIG|(Common contaminant protei
3 | 3 >YDR055W PST1 SGDID:S000002462, Chr IV fr
4 | 4 >sp|K1C10_HUMAN|(Common contaminant
5 | 5 >YGR209C TRX2 SGDID:S000003441, Chr VII fr
6 | 6 >YBR162C TOS1 SGDID:S000000366, Chr II fro
7 | 7 >YMR307W GAS1 SGDID:S000004924, Chr XIII
8 | 8 >YOL154W ZPS1 SGDID:S000005514, Chr XV fr
9 | 9 >YNL160W YGP1 SGDID:S000005104, Chr XIV
10 | 10 >YOL030W GAS5 SGDID:S000005390, Chr XV f
11 | 11 >sp|K1C9_HUMAN|(Common contaminant p
12 | 12 >YLR043C TRX1 SGDID:S000004033, Chr XII fr
13 | 13 >YBR078W ECM33 SGDID:S000000282, Chr II f
14 | 14 >YIL169C YIL169C SGDID:S000001431, Chr IX f
15 | 15 >sp|K22E_HUMAN|(Common contaminant p
16 | 16 >YKR042W UTH1 SGDID:S000001750, Chr XI fr
17 | 17 >YGR037C ACB1 SGDID:S000003269, Chr VII fr
18 | 18 >YKL163W PIR3 SGDID:S000001646, Chr XI fro
19 | 19 >YGR282C BGL2 SGDID:S000003514, Chr VII fr
Protein Coverage (double click to dock / undock)
Protein Coverage
Peptide List (double click to dock / undock)
Peptides
PID | Prot. Rank | Pos. | Sequence | Mods (variable) | Score | Glycans | PEP 2D | PEP 1D | og Pro | Delta Score | elta Mo Score | z | Obs. m/z | Calc. m/z | ppm err. | Off-By-X | Obs. MH | Calc. MH | Cleavage | Glyc Pc
1 | 105051 | 7 | 303 | K.YGLVS[+162.05282]IDGNDVK.T | S5(OGlycan / 162.0528) | 504.6 | Hex(1) | 5.7e-9 | 2.1e-6 | 8.24 | 249.0 | 249.0 | 2 | 721.3447 | 721.3565 | -16.25 | 1441.6822 | 1441.7057 | Specific | 5
Spectrum (double click to dock / undock)
K.YGLVS[+162]IDGNDVK.T z=2,scan#=sample=1 period=1 cycle=534 experiment=17,scan time=10.9227
Intensity
m/z
m/z errors, search tol=75.0 ppm

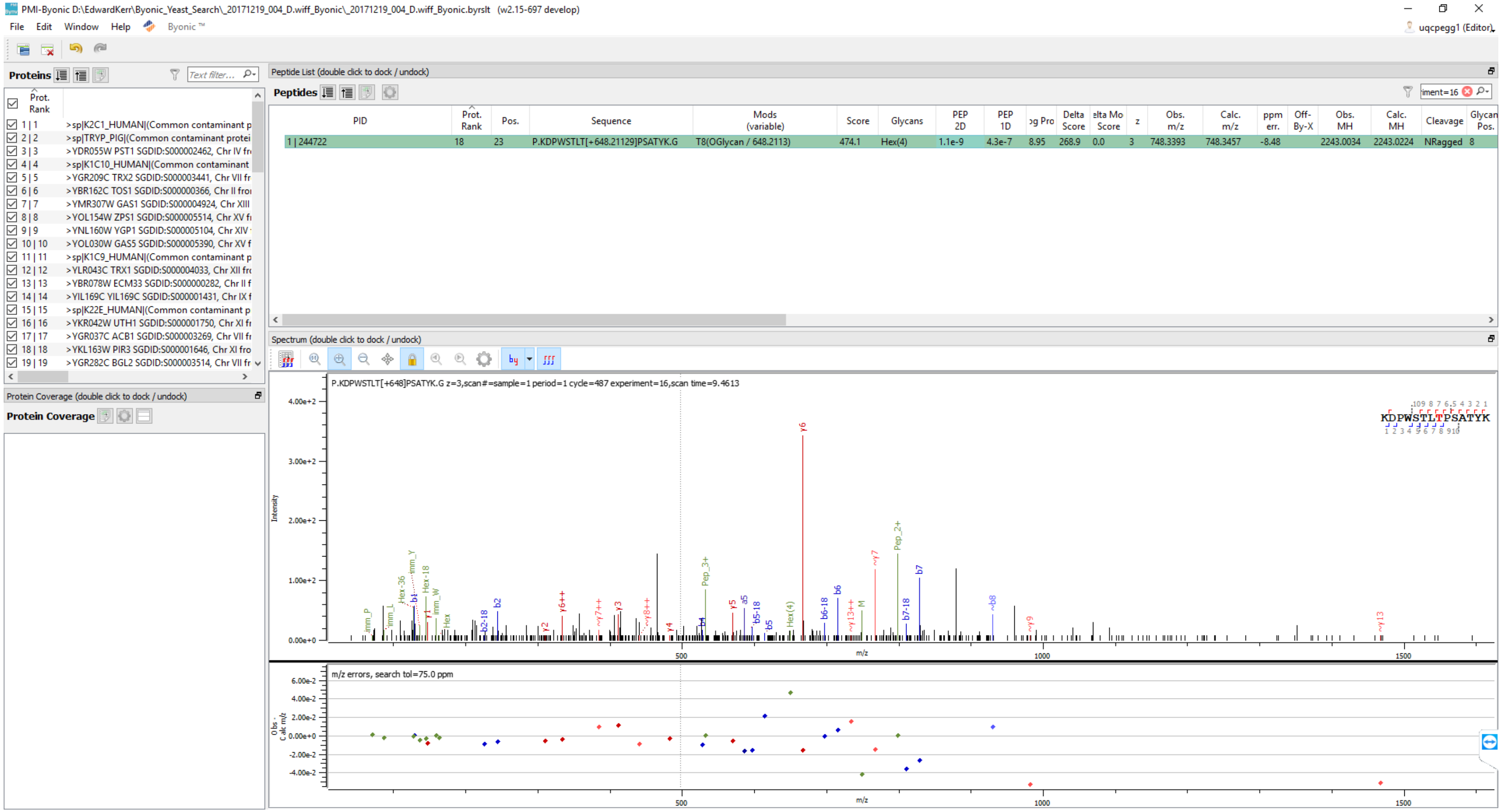
PMI-Byonic D:\EdwardKerr\Byonic_Yeast_Search_20171219_004_D.wiff_Byonic_20171219_004_D.wiff_Byonic.byrslt (w2.15-697 develop)
File Edit Window Help Byonic™
uqcpegg1 (Editor)
Proteins
Prot. Rank
1 | 1 >sp|K2C1_HUMAN|(Common contaminant p
2 | 2 >sp|TRYP_PIG|(Common contaminant protei
3 | 3 >YDR055W PST1 SGDID:S000002462, Chr IV fr
4 | 4 >sp|K1C10_HUMAN|(Common contaminant
5 | 5 >YGR209C TRX2 SGDID:S000003441, Chr VII fr
6 | 6 >YBR162C TOS1 SGDID:S000000366, Chr II fro
7 | 7 >YMR307W GAS1 SGDID:S000004924, Chr XIII
8 | 8 >YOL154W ZPS1 SGDID:S000005514, Chr XV fr
9 | 9 >YNL160W YGP1 SGDID:S000005104, Chr XIV
10 | 10 >YOL030W GAS5 SGDID:S000005390, Chr XV f
11 | 11 >sp|K1C9_HUMAN|(Common contaminant p
12 | 12 >YLR043C TRX1 SGDID:S000004033, Chr XII fr
13 | 13 >YBR078W ECM33 SGDID:S000000282, Chr II f
14 | 14 >YIL169C YIL169C SGDID:S000001431, Chr IX f
15 | 15 >sp|K22E_HUMAN|(Common contaminant p
16 | 16 >YKR042W UTH1 SGDID:S000001750, Chr XI fr
17 | 17 >YGR037C ACB1 SGDID:S000003269, Chr VII fr
18 | 18 >YKL163W PIR3 SGDID:S000001646, Chr XI fro
19 | 19 >YGR282C BGL2 SGDID:S000003514, Chr VII fr
Protein Coverage (double click to dock / undock)
Protein Coverage
Peptide List (double click to dock / undock)
Peptides
PID | Prot. Rank | Pos. | Sequence | Mods (variable) | Score | Glycans | PEP 2D | PEP 1D | Log Prob | Delta Score | Delta Mod Score | z | Obs. m/z | Calc. m/z | ppm err. | Off-By-X | Obs. MH | Calc. MH | Cleavage | Glycan Pos.
1 | 244722 | 18 | 23 | P.KDPWSTLT[+648.21129]PSATYK.G | T8(OGlycan / 648.2113) | 474.1 | Hex(4) | 1.1e-9 | 4.3e-7 | 8.95 | 268.9 | 0.0 | 3 | 748.3393 | 748.3457 | -8.48 | | 2243.0034 | 2243.0224 | NRagged | 8
Spectrum (double click to dock / undock)
P.KDPWSTLT[+648]PSATYK.G z=3,scan#=sample=1 period=1 cycle=487 experiment=16,scan time=9.4613
KDPWSTLTPSATYK
Intensity
4.00e+2
3.00e+2
2.00e+2
1.00e+2
0.00e+0
m/z
500
1000
1500
m/z errors, search tol=75.0 ppm
Obs - Calc m/z
6.00e-2
4.00e-2
2.00e-2
0.00e+0
-2.00e-2
-4.00e-2

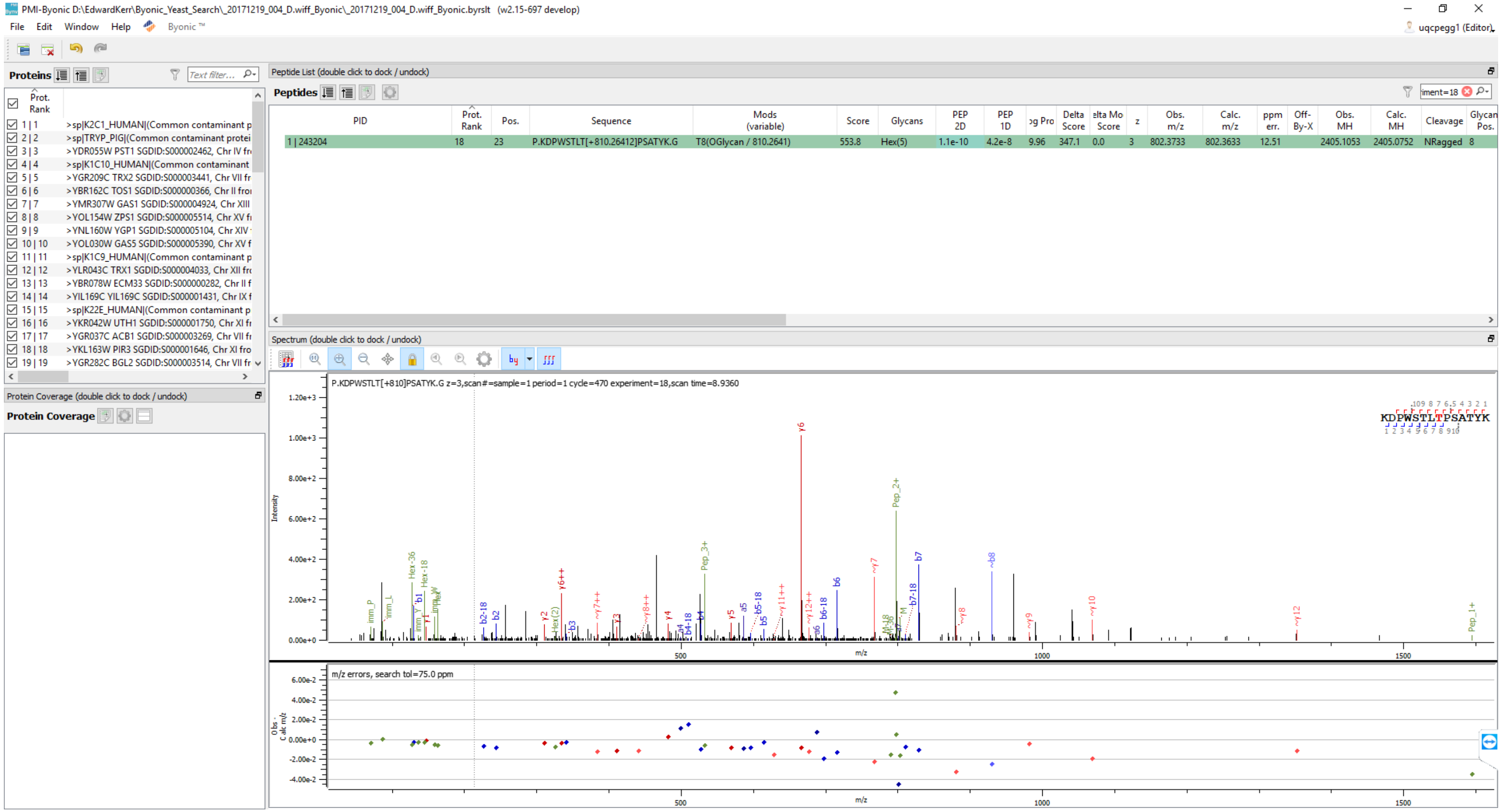
PMI-Byonic D:\EdwardKerr\Byonic_Yeast_Search_20171219_004_D.wiff_Byonic_20171219_004_D.wiff_Byonic.byrslt (w2.15-697 develop)
File Edit Window Help Byonic ™
uqcpegg1 (Editor)
Proteins
Prot. Rank
1 | 1 >sp|K2C1_HUMAN|(Common contaminant p
2 | 2 >sp|TRYP_PIG|(Common contaminant protei
3 | 3 >YDR055W PST1 SGDID:S000002462, Chr IV fr
4 | 4 >sp|K1C10_HUMAN|(Common contaminant
5 | 5 >YGR209C TRX2 SGDID:S000003441, Chr VII fr
6 | 6 >YBR162C TOS1 SGDID:S000000366, Chr II fro
7 | 7 >YMR307W GAS1 SGDID:S000004924, Chr XIII
8 | 8 >YOL154W ZPS1 SGDID:S000005514, Chr XV fr
9 | 9 >YNL160W YGP1 SGDID:S000005104, Chr XIV
10 | 10 >YOL030W GAS5 SGDID:S000005390, Chr XV f
11 | 11 >sp|K1C9_HUMAN|(Common contaminant p
12 | 12 >YLR043C TRX1 SGDID:S000004033, Chr XII fr
13 | 13 >YBR078W ECM33 SGDID:S000000282, Chr II f
14 | 14 >YIL169C YIL169C SGDID:S000001431, Chr IX f
15 | 15 >sp|K22E_HUMAN|(Common contaminant p
16 | 16 >YKR042W UTH1 SGDID:S000001750, Chr XI fr
17 | 17 >YGR037C ACB1 SGDID:S000003269, Chr VII fr
18 | 18 >YKL163W PIR3 SGDID:S000001646, Chr XI fro
19 | 19 >YGR282C BGL2 SGDID:S000003514, Chr VII fr
Protein Coverage (double click to dock / undock)
Protein Coverage
Peptide List (double click to dock / undock)
Peptides
PID | Prot. Rank | Pos. | Sequence | Mods (variable) | Score | Glycans | PEP 2D | PEP 1D | Log Prob | Delta Score | Delta Mod Score | z | Obs. m/z | Calc. m/z | ppm err. | Off-By-X | Obs. MH | Calc. MH | Cleavage | Glycan Pos.
1 | 243204 | 18 | 23 | P.KDPWSTLT[+810.26412]PSATYK.G | T8(OGlycan / 810.2641) | 553.8 | Hex(5) | 1.1e-10 | 4.2e-8 | 9.96 | 347.1 | 0.0 | 3 | 802.3733 | 802.3633 | 12.51 | 2405.1053 | 2405.0752 | NRagged | 8
Spectrum (double click to dock / undock)
P.KDPWSTLT[+810]PSATYK.G z=3,scan#=sample=1 period=1 cycle=470 experiment=18,scan time=8.9360
KDPWSTLTPSATYK
Intensity
m/z
500
1000
1500
y6
Pep_2+
b7
~b8
Pep_3+
y6++
Hex-36
Hex-18
m/z errors, search tol=75.0 ppm
Obs - Calc m/z

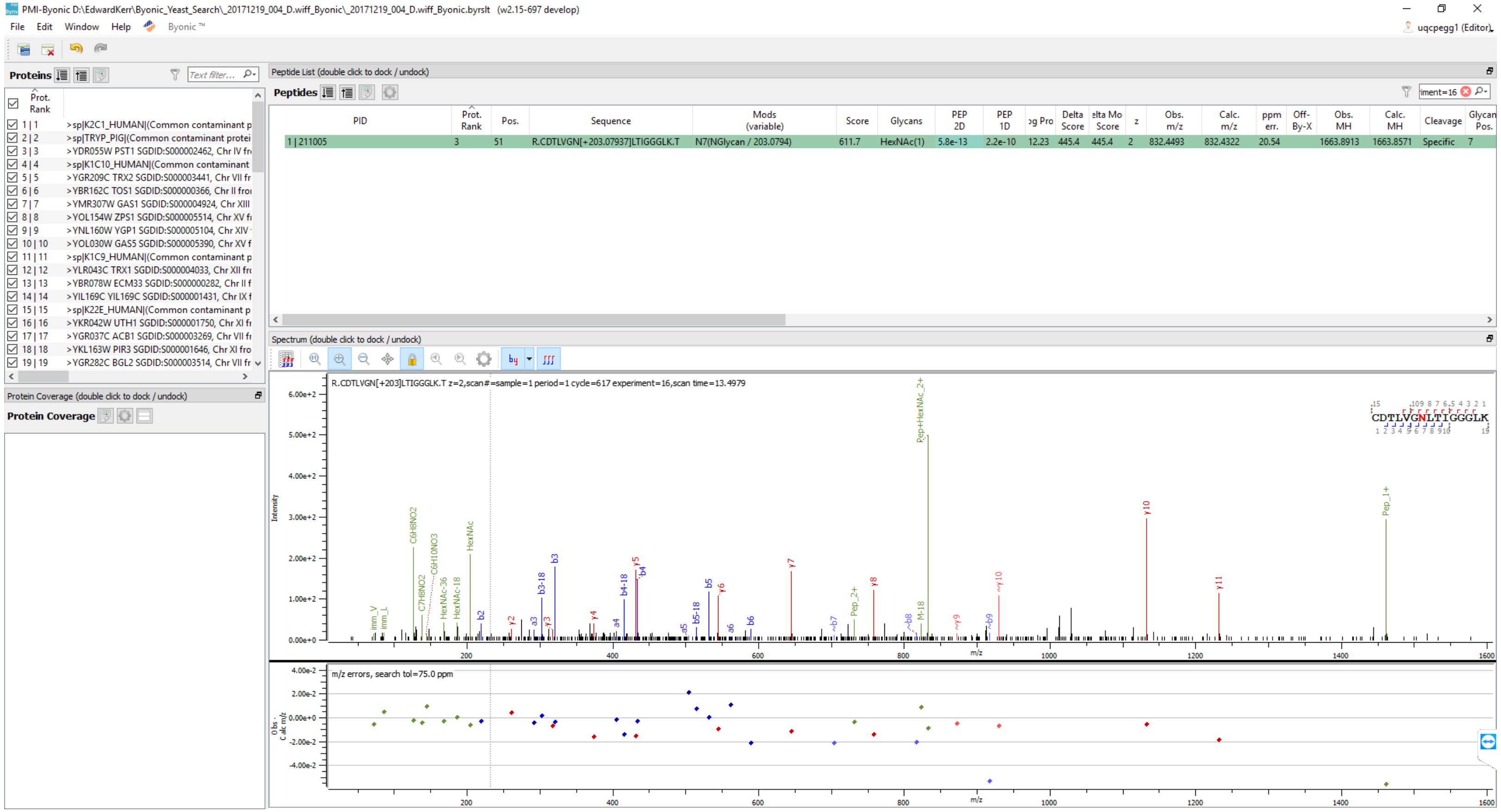
PMI-Byonic D:\EdwardKerr\Byonic_Yeast_Search_20171219_004_D.wiff_Byonic_20171219_004_D.wiff_Byonic.byrslt (w2.15-697 develop)
File Edit Window Help Byonic ™
Proteins
Prot. Rank
1 | 1 >sp|K2C1_HUMAN|(Common contaminant p
2 | 2 >sp|TRYP_PIG|(Common contaminant protei
3 | 3 >YDR055W PST1 SGDID:S000002462, Chr IV fr
4 | 4 >sp|K1C10_HUMAN|(Common contaminant
5 | 5 >YGR209C TRX2 SGDID:S000003441, Chr VII fr
6 | 6 >YBR162C TOS1 SGDID:S000000366, Chr II fro
7 | 7 >YMR307W GAS1 SGDID:S000004924, Chr XIII
8 | 8 >YOL154W ZPS1 SGDID:S000005514, Chr XV fr
9 | 9 >YNL160W YGP1 SGDID:S000005104, Chr XIV
10 | 10 >YOL030W GAS5 SGDID:S000005390, Chr XV f
11 | 11 >sp|K1C9_HUMAN|(Common contaminant p
12 | 12 >YLR043C TRX1 SGDID:S000004033, Chr XII fr
13 | 13 >YBR078W ECM33 SGDID:S000000282, Chr II f
14 | 14 >YIL169C YIL169C SGDID:S000001431, Chr IX f
15 | 15 >sp|K22E_HUMAN|(Common contaminant p
16 | 16 >YKR042W UTH1 SGDID:S000001750, Chr XI fr
17 | 17 >YGR037C ACB1 SGDID:S000003269, Chr VII fr
18 | 18 >YKL163W PIR3 SGDID:S000001646, Chr XI fro
19 | 19 >YGR282C BGL2 SGDID:S000003514, Chr VII fr
Protein Coverage (double click to dock / undock)
Protein Coverage
Peptide List (double click to dock / undock)
Peptides
PID | Prot. Rank | Pos. | Sequence | Mods (variable) | Score | Glycans | PEP 2D | PEP 1D | og Pro | Delta Score | elta Mo Score | z | Obs. m/z | Calc. m/z | ppm err. | Off-By-X | Obs. MH | Calc. MH | Cleavage | Glycan Pos.
1 | 211005 | 3 | 51 | R.CDTLVGN[+203.07937]LTIGGGLK.T | N7(NGlycan / 203.0794) | 611.7 | HexNAc(1) | 5.8e-13 | 2.2e-10 | 12.23 | 445.4 | 445.4 | 2 | 832.4493 | 832.4322 | 20.54 | | 1663.8913 | 1663.8571 | Specific | 7
Spectrum (double click to dock / undock)
R.CDTLVGN[+203]LTIGGGLK.T z=2,scan#=sample=1 period=1 cycle=617 experiment=16,scan time=13.4979
CDTLVGNLTIGGGLK
Intensity
m/z
m/z errors, search tol=75.0 ppm
Obs - Calc m/z

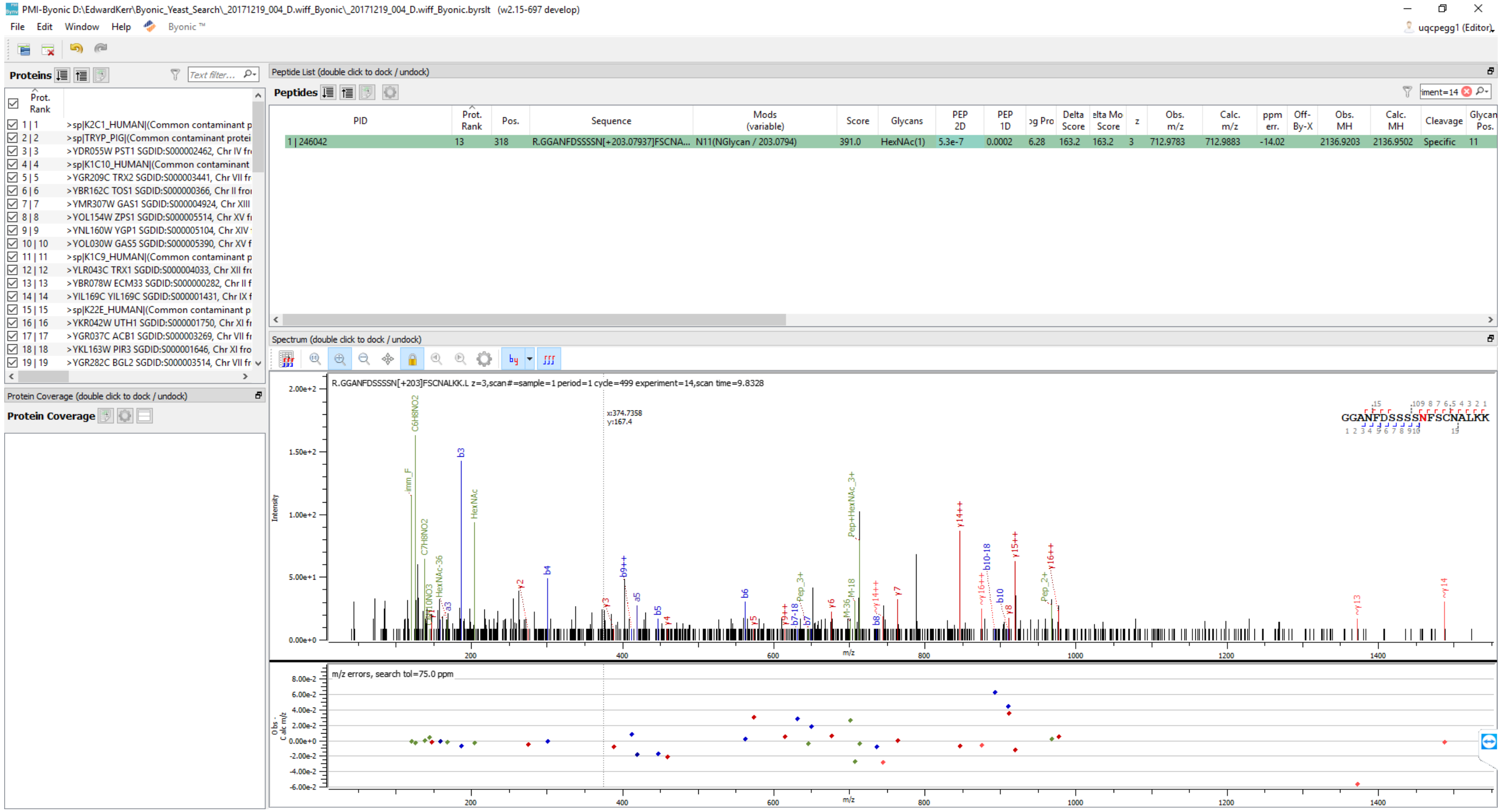
PMI-Byonic D:\EdwardKerr\Byonic_Yeast_Search_20171219_004_D.wiff_Byonic_20171219_004_D.wiff_Byonic.byrslt (w2.15-697 develop)
File Edit Window Help Byonic ™
uqcpegg1 (Editor)
Proteins
Prot. Rank
1 | 1 >sp|K2C1_HUMAN|(Common contaminant p
2 | 2 >sp|TRYP_PIG|(Common contaminant protei
3 | 3 >YDR055W PST1 SGDID:S000002462, Chr IV fr
4 | 4 >sp|K1C10_HUMAN|(Common contaminant
5 | 5 >YGR209C TRX2 SGDID:S000003441, Chr VII fr
6 | 6 >YBR162C TOS1 SGDID:S000000366, Chr II fro
7 | 7 >YMR307W GAS1 SGDID:S000004924, Chr XIII
8 | 8 >YOL154W ZPS1 SGDID:S000005514, Chr XV fi
9 | 9 >YNL160W YGP1 SGDID:S000005104, Chr XIV
10 | 10 >YOL030W GAS5 SGDID:S000005390, Chr XV f
11 | 11 >sp|K1C9_HUMAN|(Common contaminant p
12 | 12 >YLR043C TRX1 SGDID:S000004033, Chr XII fr
13 | 13 >YBR078W ECM33 SGDID:S000000282, Chr II f
14 | 14 >YIL169C YIL169C SGDID:S000001431, Chr IX f
15 | 15 >sp|K22E_HUMAN|(Common contaminant p
16 | 16 >YKR042W UTH1 SGDID:S000001750, Chr XI fr
17 | 17 >YGR037C ACB1 SGDID:S000003269, Chr VII fr
18 | 18 >YKL163W PIR3 SGDID:S000001646, Chr XI fro
19 | 19 >YGR282C BGL2 SGDID:S000003514, Chr VII fr
Protein Coverage (double click to dock / undock)
Protein Coverage
Peptide List (double click to dock / undock)
Peptides
PID | Prot. Rank | Pos. | Sequence | Mods (variable) | Score | Glycans | PEP 2D | PEP 1D | og Pro | Delta Score | elta Mo Score | z | Obs. m/z | Calc. m/z | ppm err. | Off-By-X | Obs. MH | Calc. MH | Cleavage | Glycan Pos.
1 | 246042 | 13 | 318 | R.GGANFDSSSSN[+203.07937]FSCNA... | N11(NGlycan / 203.0794) | 391.0 | HexNAc(1) | 5.3e-7 | 0.0002 | 6.28 | 163.2 | 163.2 | 3 | 712.9783 | 712.9883 | -14.02 | | 2136.9203 | 2136.9502 | Specific | 11
Spectrum (double click to dock / undock)
R.GGANFDSSSSN[+203]FSCNALKK.L z=3,scan#=sample=1 period=1 cycle=499 experiment=14,scan time=9.8328
x:374.7358
y:167.4
GGANFDSSSSNFSCNALKK
Intensity
2.00e+2
1.50e+2
1.00e+2
5.00e+1
0.00e+0
m/z
200
400
600
800
1000
1200
1400
m/z errors, search tol=75.0 ppm
8.00e-2
6.00e-2
4.00e-2
2.00e-2
0.00e+0
-2.00e-2
-4.00e-2
-6.00e-2

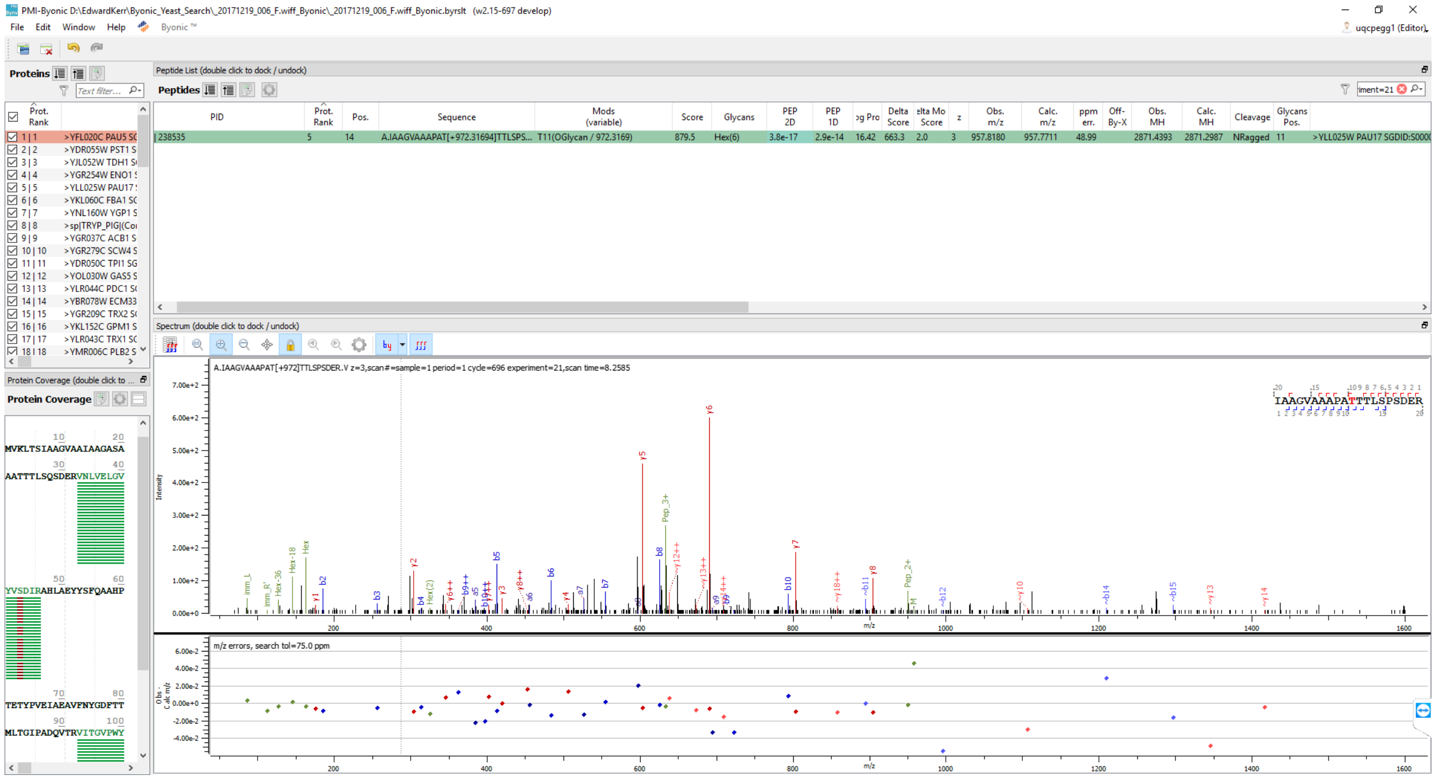
PMI-Byonic D:\EdwardKerr\Byonic_Yeast_Search_20171219_006_F.wiff_Byonic_20171219_006_F.wiff_Byonic.byrslt (w2.15-697 develop)
File Edit Window Help Byonic™
uqcpegg1 (Editor)
Proteins
Peptide List (double click to dock / undock)
Peptides
PID
Prot. Rank
Pos.
Sequence
Mods (variable)
Score
Glycans
PEP 2D
PEP 1D
Delta Score
z
Obs. m/z
Calc. m/z
ppm err.
Obs. MH
Calc. MH
Cleavage
Glycans Pos.
238535 5 14 A.IAAGVAAAPAT[+972.31694]TTLSPS... T11(OGlycan / 972.3169) 879.5 Hex(6) 3.8e-17 2.9e-14 16.42 663.3 2.0 3 957.8180 957.7711 48.99 2871.4393 2871.2987 NRagged 11 >YLL025W PAU17 SGDID:S000
>YFL020C PAU5
>YDR055W PST1
>YJL052W TDH1
>YGR254W ENO1
>YLL025W PAU17
>YKL060C FBA1
>YNL160W YGP1
>sp|TRYP_PIG|(Co
>YGR037C ACB1
>YGR279C SCW4
>YDR050C TPI1
>YOL030W GAS5
>YLR044C PDC1
>YBR078W ECM33
>YGR209C TRX2
>YKL152C GPM1
>YLR043C TRX1
>YMR006C PLB2
Spectrum (double click to dock / undock)
A.IAAGVAAAPAT[+972]TTLSPSDER.V z=3,scan#=sample=1 period=1 cycle=696 experiment=21,scan time=8.2585
IAAGVAAAPATTTLSPSDER
Protein Coverage (double click to ...
Protein Coverage
MVKLTSIAAGVAAIAAGASA
AATTTLSQSDERVNLVELGV
YVSDIRAHLAEYYSFQAAHP
TETYPVEIAEAVFNYGDFTT
MLTGIPADQVTRVITGVPWY
Intensity
m/z
m/z errors, search tol=75.0 ppm
y5
y6
y7
b8
b10
Pep_3+
Pep_2+

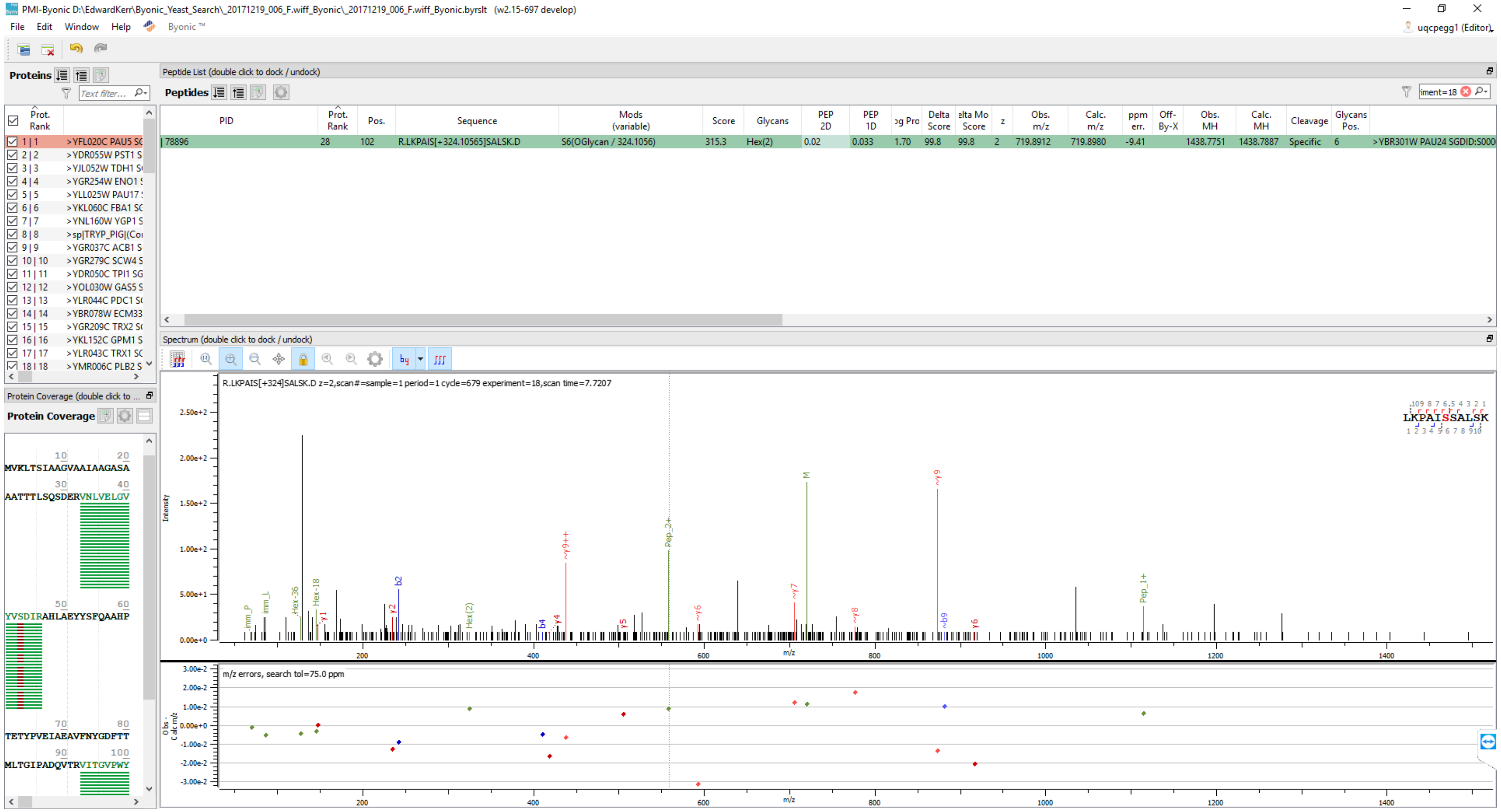
PMI-Byonic D:\EdwardKerr\Byonic_Yeast_Search_20171219_006_F.wiff_Byonic_20171219_006_F.wiff_Byonic.byrslt (w2.15-697 develop)
File Edit Window Help Byonic ™
uqcpegg1 (Editor)
Proteins
Peptide List (double click to dock / undock)
Peptides
PID
Prot. Rank
Pos.
Sequence
Mods (variable)
Score
Glycans
PEP 2D
PEP 1D
Delta Score
z
Obs. m/z
Calc. m/z
ppm err.
Off-By-X
Obs. MH
Calc. MH
Cleavage
Glycans Pos.
78896 28 102 R.LKPAIS[+324.10565]SALSK.D S6(OGlycan / 324.1056) 315.3 Hex(2) 0.02 0.033 1.70 99.8 99.8 2 719.8912 719.8980 -9.41 1438.7751 1438.7887 Specific 6 >YBR301W PAU24 SGDID:S000
Spectrum (double click to dock / undock)
R.LKPAIS[+324]SALSK.D z=2,scan#=sample=1 period=1 cycle=679 experiment=18,scan time=7.7207
LKPAISSALSK
Intensity
m/z
m/z errors, search tol=75.0 ppm
Protein Coverage (double click to ...
Protein Coverage
MVKLTSIAAGVAAIAAGASA
AATTTLSQSDERVNLVELGV
YVSDIRAHLAEYYSFQAAHP
TETYPVEIAEAVFNYGDFTT
MLTGIPADQVTRVITGVPWY

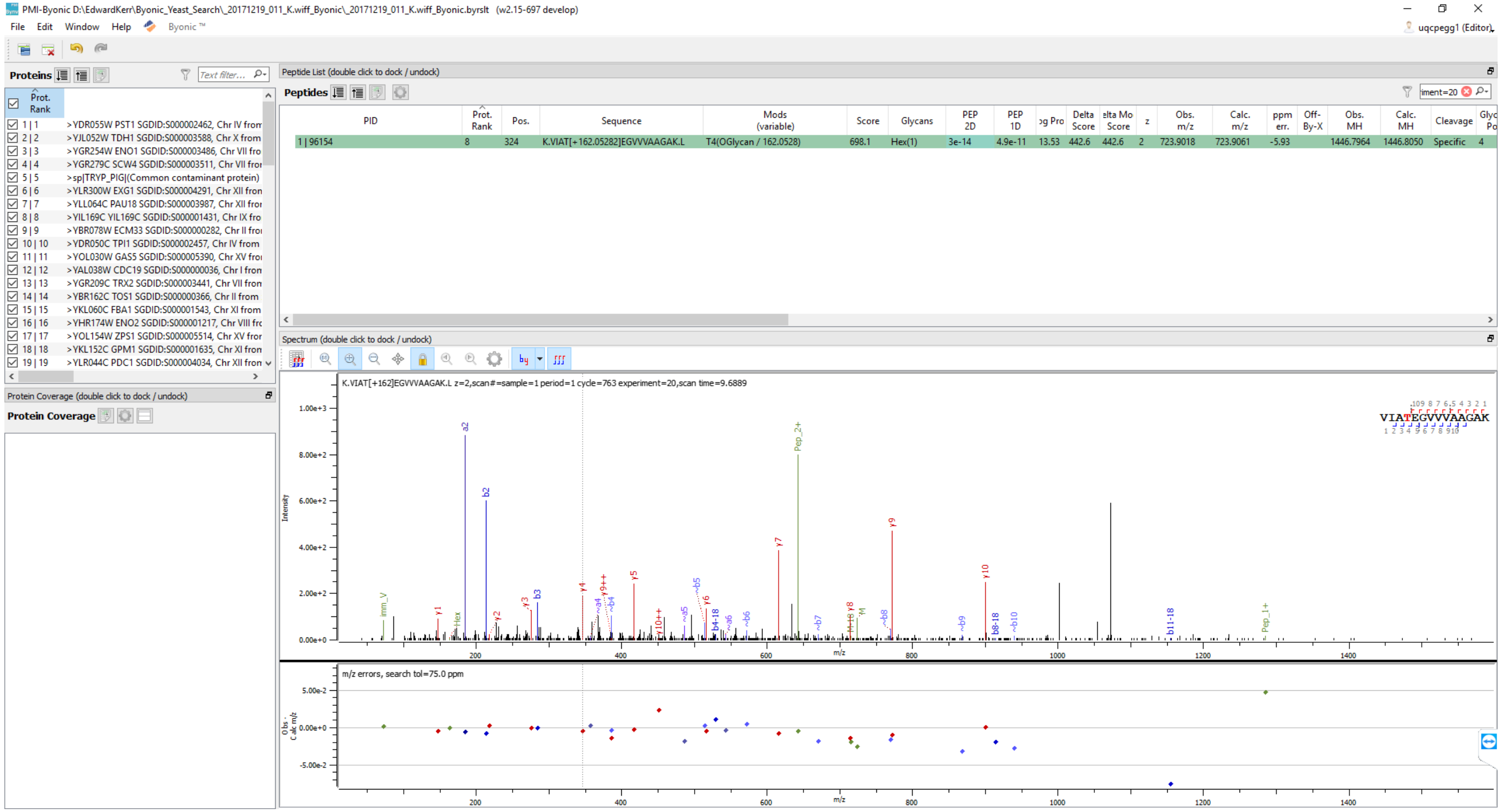

PMI-Byonic D:\EdwardKerr\Byonic_Yeast_Search_20171219_011_K.wiff_Byonic_20171219_011_K.wiff_Byonic.byrslt (w2.15-697 develop)
File Edit Window Help Byonic ™
uqcpegg1 (Editor)
Proteins
Prot. Rank
1 | 1 >YDR055W PST1 SGDID:S000002462, Chr IV from
2 | 2 >YJL052W TDH1 SGDID:S000003588, Chr X from
3 | 3 >YGR254W ENO1 SGDID:S000003486, Chr VII fro
4 | 4 >YGR279C SCW4 SGDID:S000003511, Chr VII fro
5 | 5 >sp|TRYP_PIG|(Common contaminant protein)
6 | 6 >YLR300W EXG1 SGDID:S000004291, Chr XII fron
7 | 7 >YLL064C PAU18 SGDID:S000003987, Chr XII fro
8 | 8 >YIL169C YIL169C SGDID:S000001431, Chr IX fro
9 | 9 >YBR078W ECM33 SGDID:S000000282, Chr II fro
10 | 10 >YDR050C TPI1 SGDID:S000002457, Chr IV from
11 | 11 >YOL030W GAS5 SGDID:S000005390, Chr XV fro
12 | 12 >YAL038W CDC19 SGDID:S000000036, Chr I from
13 | 13 >YGR209C TRX2 SGDID:S000003441, Chr VII from
14 | 14 >YBR162C TOS1 SGDID:S000000366, Chr II from
15 | 15 >YKL060C FBA1 SGDID:S000001543, Chr XI from
16 | 16 >YHR174W ENO2 SGDID:S000001217, Chr VIII fro
17 | 17 >YOL154W ZPS1 SGDID:S000005514, Chr XV fro
18 | 18 >YKL152C GPM1 SGDID:S000001635, Chr XI from
19 | 19 >YLR044C PDC1 SGDID:S000004034, Chr XII from
Protein Coverage (double click to dock / undock)
Protein Coverage
Peptide List (double click to dock / undock)
Peptides
iment=20
PID | Prot. Rank | Pos. | Sequence | Mods (variable) | Score | Glycans | PEP 2D | PEP 1D | og Pro | Delta Score | elta Mo Score | z | Obs. m/z | Calc. m/z | ppm err. | Off-By-X | Obs. MH | Calc. MH | Cleavage | Glyc Po
1 | 96154 | 8 | 324 | K.VIAT[+162.05282]EGVVVAAGAK.L | T4(OGlycan / 162.0528) | 698.1 | Hex(1) | 3e-14 | 4.9e-11 | 13.53 | 442.6 | 442.6 | 2 | 723.9018 | 723.9061 | -5.93 | | 1446.7964 | 1446.8050 | Specific | 4
Spectrum (double click to dock / undock)
K.VIAT[+162]EGVVVAAGAK.L z=2,scan#=sample=1 period=1 cycle=763 experiment=20,scan time=9.6889
VIATEGVVVAAGAK
Intensity
m/z
m/z errors, search tol=75.0 ppm

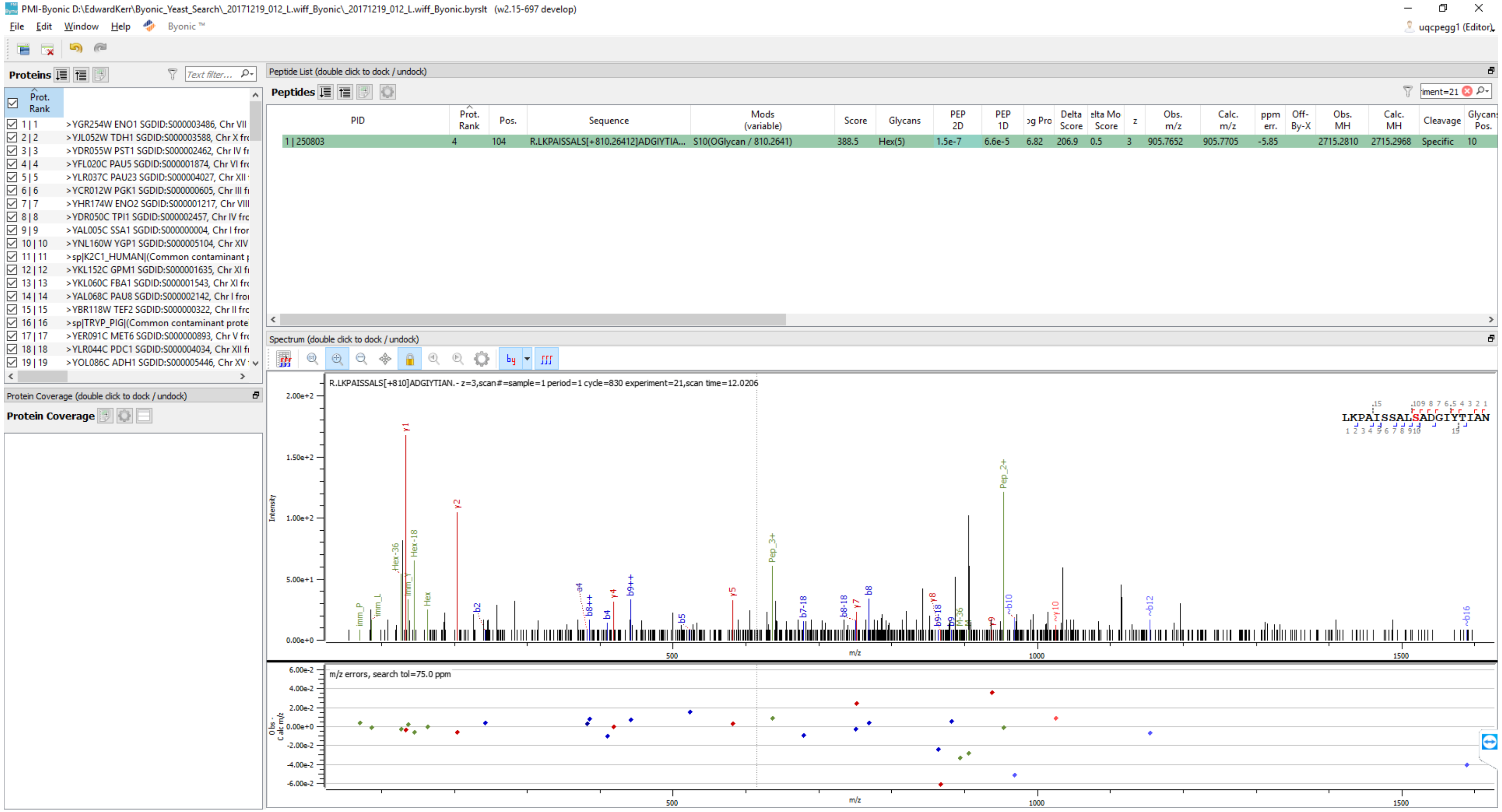
PMI-Byonic D:\EdwardKerr\Byonic_Yeast_Search_20171219_012_L.wiff_Byonic_20171219_012_L.wiff_Byonic.byrslt (w2.15-697 develop)
File Edit Window Help Byonic ™
uqcpegg1 (Editor)
Proteins
Text filter...
Prot. Rank
1 | 1 >YGR254W ENO1 SGDID:S000003486, Chr VII
2 | 2 >YJL052W TDH1 SGDID:S000003588, Chr X
3 | 3 >YDR055W PST1 SGDID:S000002462, Chr IV
4 | 4 >YFL020C PAU5 SGDID:S000001874, Chr VI
5 | 5 >YLR037C PAU23 SGDID:S000004027, Chr XII
6 | 6 >YCR012W PGK1 SGDID:S000000605, Chr III
7 | 7 >YHR174W ENO2 SGDID:S000001217, Chr VIII
8 | 8 >YDR050C TPI1 SGDID:S000002457, Chr IV
9 | 9 >YAL005C SSA1 SGDID:S000000004, Chr I
10 | 10 >YNL160W YGP1 SGDID:S000005104, Chr XIV
11 | 11 >sp|K2C1_HUMAN|(Common contaminant
12 | 12 >YKL152C GPM1 SGDID:S000001635, Chr XI
13 | 13 >YKL060C FBA1 SGDID:S000001543, Chr XI
14 | 14 >YAL068C PAU8 SGDID:S000002142, Chr I
15 | 15 >YBR118W TEF2 SGDID:S000000322, Chr II
16 | 16 >sp|TRYP_PIG|(Common contaminant prote
17 | 17 >YER091C MET6 SGDID:S000000893, Chr V
18 | 18 >YLR044C PDC1 SGDID:S000004034, Chr XII
19 | 19 >YOL086C ADH1 SGDID:S000005446, Chr XV
Protein Coverage (double click to dock / undock)
Protein Coverage
Peptide List (double click to dock / undock)
Peptides
iment=21
PID | Prot. Rank | Pos. | Sequence | Mods (variable) | Score | Glycans | PEP 2D | PEP 1D | og Pro | Delta Score | elta Mo Score | z | Obs. m/z | Calc. m/z | ppm err. | Off-By-X | Obs. MH | Calc. MH | Cleavage | Glycans Pos.
1 | 250803 | 4 | 104 | R.LKPAISSALS[+810.26412]ADGIYTIA... | S10(OGlycan / 810.2641) | 388.5 | Hex(5) | 1.5e-7 | 6.6e-5 | 6.82 | 206.9 | 0.5 | 3 | 905.7652 | 905.7705 | -5.85 | | 2715.2810 | 2715.2968 | Specific | 10
Spectrum (double click to dock / undock)
R.LKPAISSALS[+810]ADGIYTIAN.- z=3,scan#=sample=1 period=1 cycle=830 experiment=21,scan time=12.0206
LKPAISSALSADGIYTIAN
Intensity
2.00e+2
1.50e+2
1.00e+2
5.00e+1
0.00e+0
y1
y2
Hex-36
Hex-18
Hex
imm_P
imm_L
b2
a4
b8++
b4
y4
b9++
b5
y5
Pep_3+
b7-18
b8-18
y7
b8
y8
b9-18
b9
M-36
y9
~b10
Pep_2+
~y10
~b12
~b16
500
m/z
1000
1500
m/z errors, search tol=75.0 ppm
Obs - Calc m/z
6.00e-2
4.00e-2
2.00e-2
0.00e+0
-2.00e-2
-4.00e-2
-6.00e-2
500
m/z
1000
1500

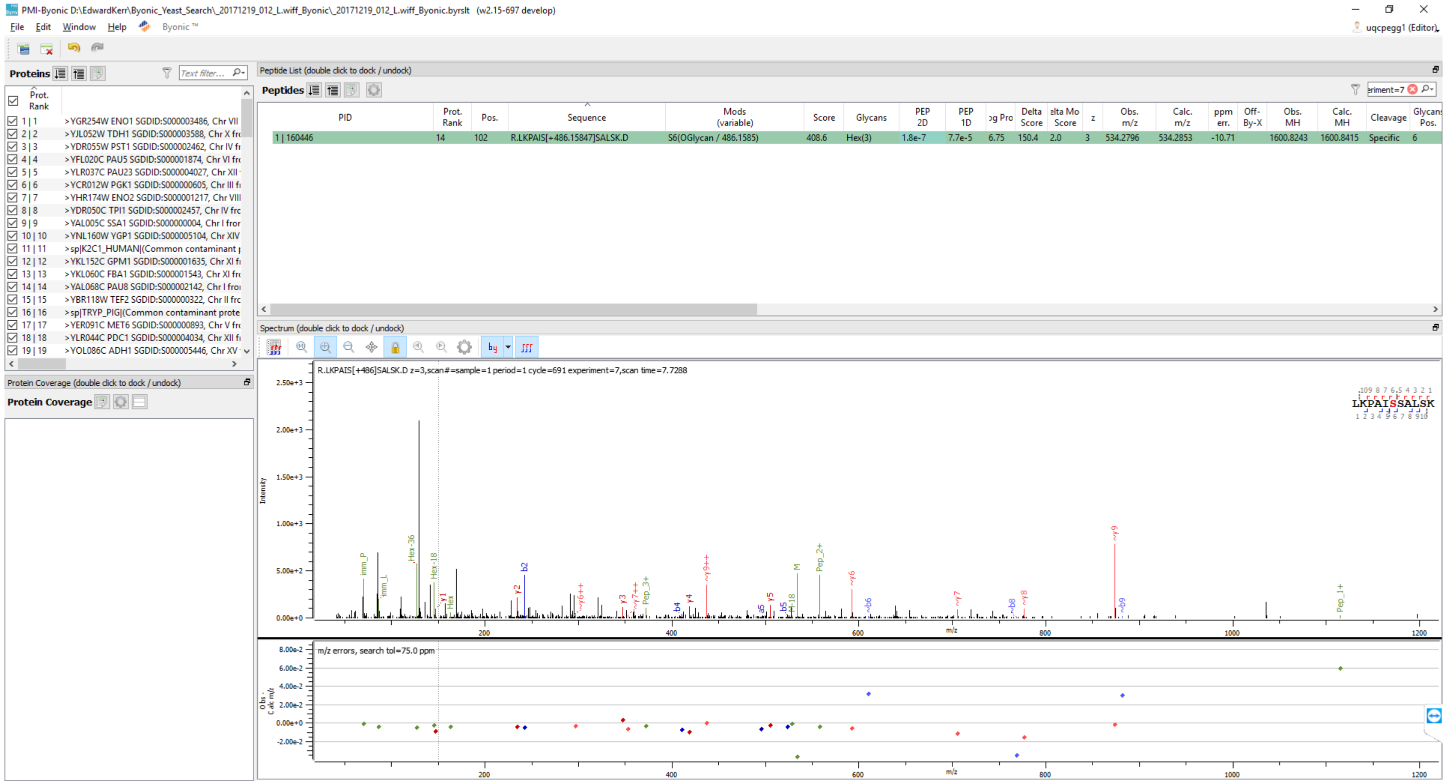
PMI-Byonic D:\EdwardKerr\Byonic_Yeast_Search_20171219_012_L.wiff_Byonic_20171219_012_L.wiff_Byonic.byrslt (w2.15-697 develop)
File Edit Window Help Byonic ™
uqcpegg1 (Editor)
Proteins
Prot. Rank
1 | 1 >YGR254W ENO1 SGDID:S000003486, Chr VII
2 | 2 >YJL052W TDH1 SGDID:S000003588, Chr X fr
3 | 3 >YDR055W PST1 SGDID:S000002462, Chr IV fr
4 | 4 >YFL020C PAU5 SGDID:S000001874, Chr VI fr
5 | 5 >YLR037C PAU23 SGDID:S000004027, Chr XII
6 | 6 >YCR012W PGK1 SGDID:S000000605, Chr III fr
7 | 7 >YHR174W ENO2 SGDID:S000001217, Chr VIII
8 | 8 >YDR050C TPI1 SGDID:S000002457, Chr IV fr
9 | 9 >YAL005C SSA1 SGDID:S000000004, Chr I fror
10 | 10 >YNL160W YGP1 SGDID:S000005104, Chr XIV
11 | 11 >sp|K2C1_HUMAN|(Common contaminant
12 | 12 >YKL152C GPM1 SGDID:S000001635, Chr XI fr
13 | 13 >YKL060C FBA1 SGDID:S000001543, Chr XI fr
14 | 14 >YAL068C PAU8 SGDID:S000002142, Chr I fror
15 | 15 >YBR118W TEF2 SGDID:S000000322, Chr II fr
16 | 16 >sp|TRYP_PIG|(Common contaminant prote
17 | 17 >YER091C MET6 SGDID:S000000893, Chr V fr
18 | 18 >YLR044C PDC1 SGDID:S000004034, Chr XII fr
19 | 19 >YOL086C ADH1 SGDID:S000005446, Chr XV
Protein Coverage (double click to dock / undock)
Protein Coverage
Peptide List (double click to dock / undock)
Peptides
PID | Prot. Rank | Pos. | Sequence | Mods (variable) | Score | Glycans | PEP 2D | PEP 1D | og Pro | Delta Score | elta Mo Score | z | Obs. m/z | Calc. m/z | ppm err. | Off-By-X | Obs. MH | Calc. MH | Cleavage | Glycans Pos.
1 | 160446 | 14 | 102 | R.LKPAIS[+486.15847]SALSK.D | S6(OGlycan / 486.1585) | 408.6 | Hex(3) | 1.8e-7 | 7.7e-5 | 6.75 | 150.4 | 2.0 | 3 | 534.2796 | 534.2853 | -10.71 | | 1600.8243 | 1600.8415 | Specific | 6
Spectrum (double click to dock / undock)
R.LKPAIS[+486]SALSK.D z=3,scan#=sample=1 period=1 cycle=691 experiment=7,scan time=7.7288
LKPAISSALSK
Intensity
m/z
m/z errors, search tol=75.0 ppm

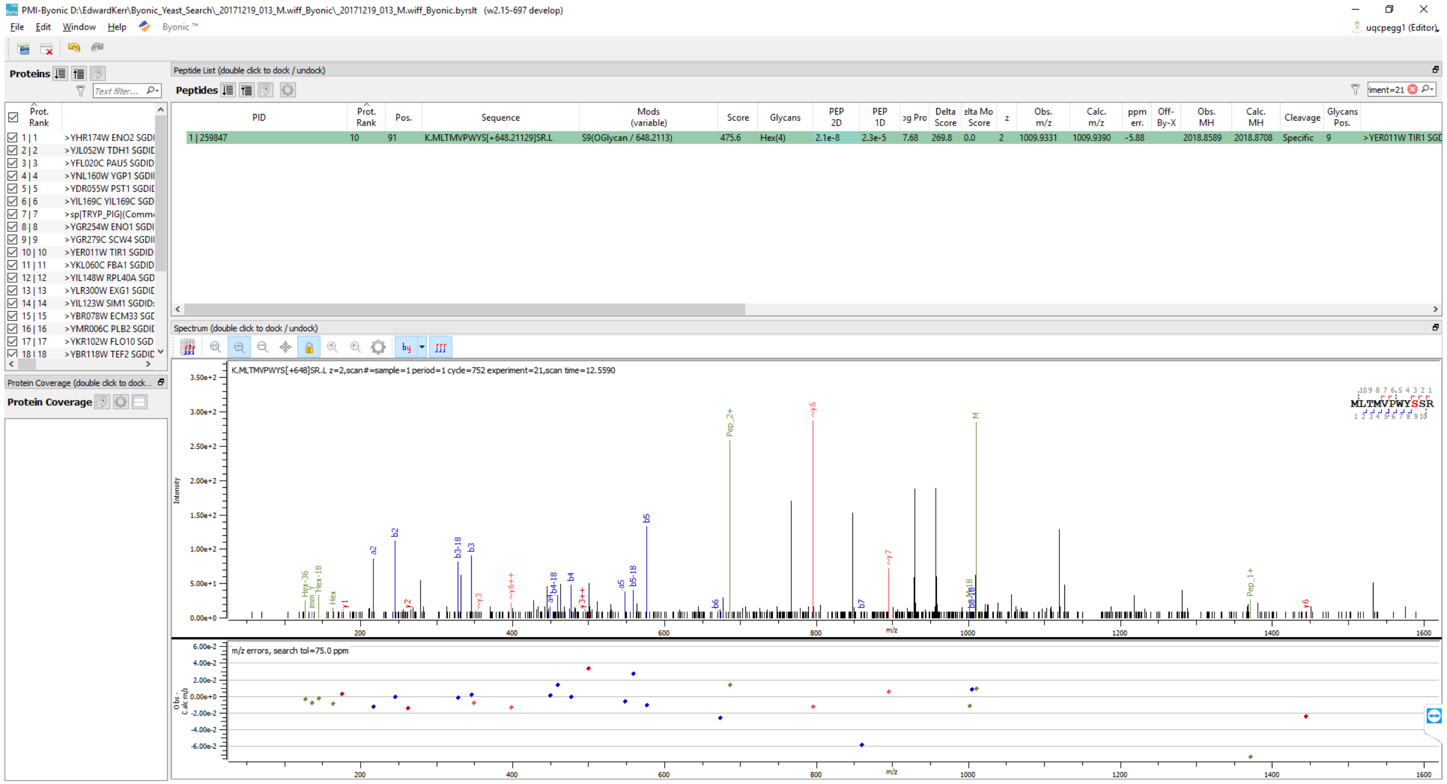

PMI-Byonic D:\EdwardKerr\Byonic_Yeast_Search_20171219_013_M.wiff_Byonic_20171219_013_M.wiff_Byonic.byrslt (w2.15-697 develop)
File Edit Window Help Byonic ™
uqcpegg1 (Editor)
Proteins
Peptide List (double click to dock / undock)
Peptides
Prot. Rank
>YHR174W ENO2 SGDI
>YJL052W TDH1 SGDI
>YFL020C PAU5 SGDID
>YNL160W YGP1 SGDI
>YDR055W PST1 SGDI
>YIL169C YIL169C SGD
>sp|TRYP_PIG|(Comm
>YGR254W ENO1 SGDI
>YGR279C SCW4 SGDI
>YER011W TIR1 SGDID
>YKL060C FBA1 SGDID
>YIL148W RPL40A SGD
>YLR300W EXG1 SGDI
>YIL123W SIM1 SGDID
>YBR078W ECM33 SGI
>YMR006C PLB2 SGDI
>YKR102W FLO10 SGD
>YBR118W TEF2 SGDI
PID | Prot. Rank | Pos. | Sequence | Mods (variable) | Score | Glycans | PEP 2D | PEP 1D | Log Prob | Delta Score | Delta Mod Score | z | Obs. m/z | Calc. m/z | ppm err. | Off-By-X | Obs. MH | Calc. MH | Cleavage | Glycans Pos.
1 | 259847 | 10 | 91 | K.MLTMVPWYS[+648.21129]SR.L | S9(OGlycan / 648.2113) | 475.6 | Hex(4) | 2.1e-8 | 2.3e-5 | 7.68 | 269.8 | 0.0 | 2 | 1009.9331 | 1009.9390 | -5.88 | | 2018.8589 | 2018.8708 | Specific | 9 | >YER011W TIR1 SGD
Spectrum (double click to dock / undock)
K.MLTMVPWYS[+648]SR.L z=2,scan#=sample=1 period=1 cycle=752 experiment=21,scan time=12.5590
Protein Coverage (double click to dock...)
Protein Coverage
Intensity
m/z
M
Pep_2+
~y6
b5
b2
a2
b3
b3-18
b4
b4-18
b5-18
a5
b6
b7
~y7
~y6++
~y3
y3++
Pep_1+
y6
Hex-36
Hex-18
Hex
y1
y2
MLTMVPWYSSR
m/z errors, search tol=75.0 ppm
Obs-Calc m/z

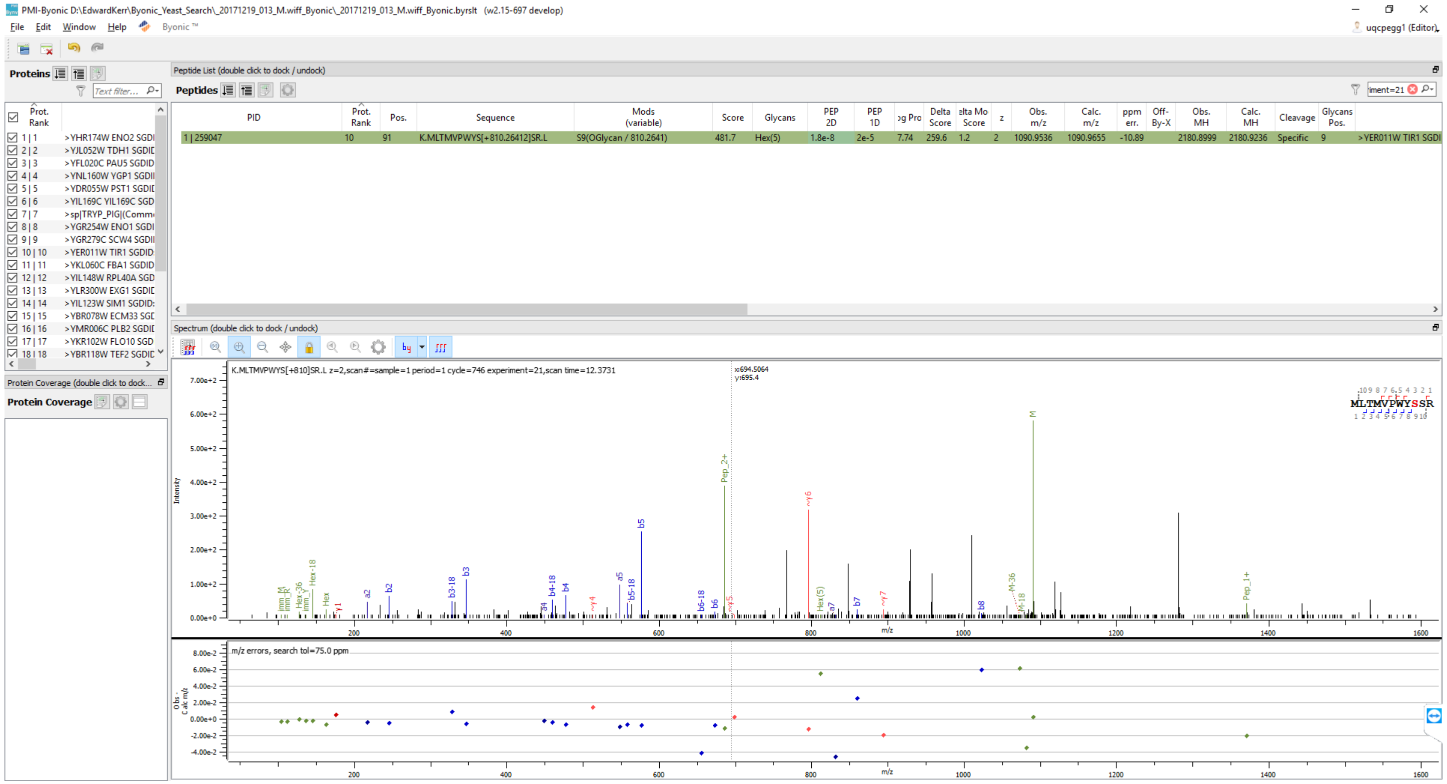
PMI-Byonic D:\EdwardKerr\Byonic_Yeast_Search_20171219_013_M.wiff_Byonic_20171219_013_M.wiff_Byonic.byrslt (w2.15-697 develop)
File Edit Window Help
Proteins
Peptide List (double click to dock / undock)
Peptides
Spectrum (double click to dock / undock)
Protein Coverage (double click to dock...
Protein Coverage
K.MLTMVPWYS[+810]SR.L z=2,scan#=sample=1 period=1 cycle=746 experiment=21,scan time=12.3731
m/z errors, search tol=75.0 ppm
Intensity
m/z

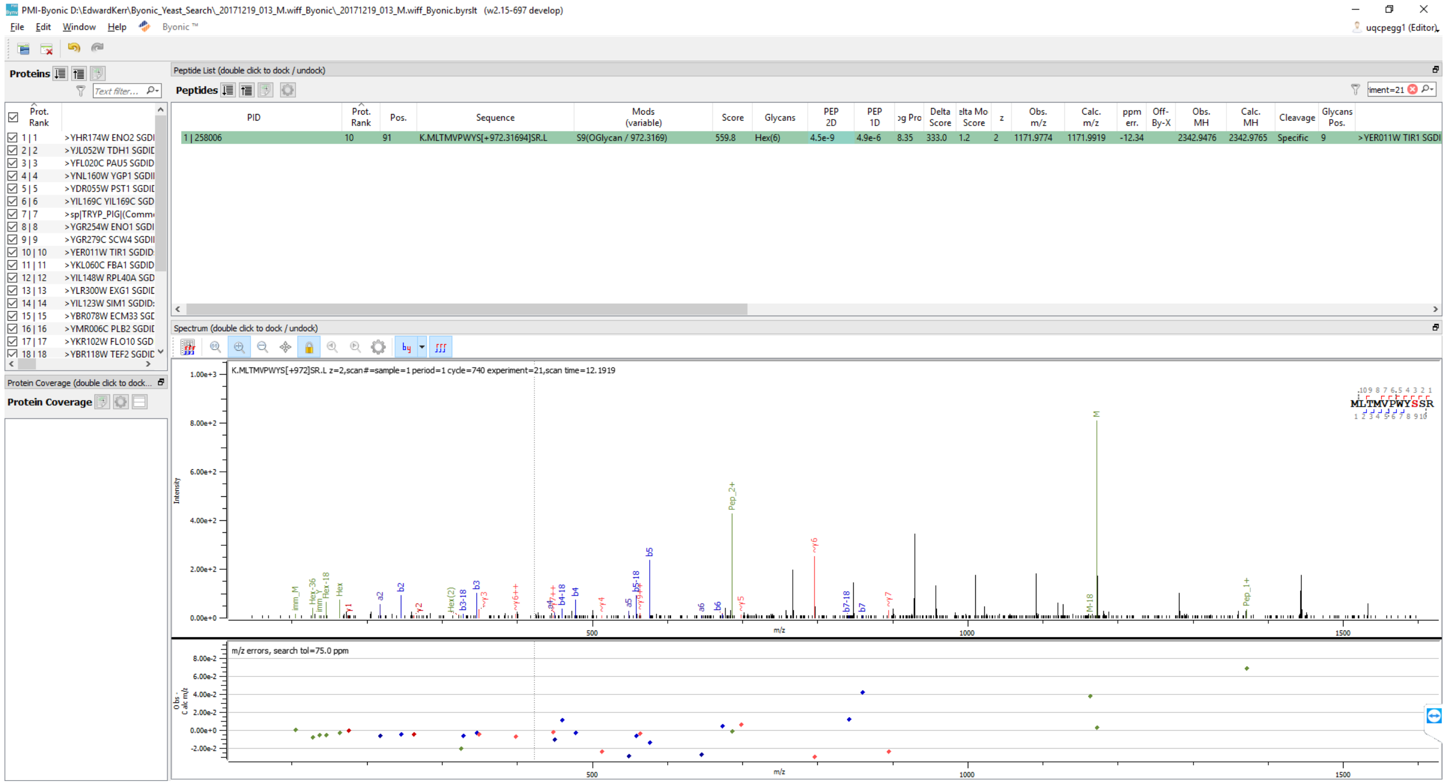
PMI-Byonic D:\EdwardKerr\Byonic_Yeast_Search_20171219_013_M.wiff_Byonic_20171219_013_M.wiff_Byonic.byrslt (w2.15-697 develop)
File Edit Window Help Byonic ™
uqcpegg1 (Editor)
Proteins
Peptide List (double click to dock / undock)
Peptides
PID | Prot. Rank | Pos. | Sequence | Mods (variable) | Score | Glycans | PEP 2D | PEP 1D | Delta Score | z | Obs. m/z | Calc. m/z | ppm err. | Off-By-X | Obs. MH | Calc. MH | Cleavage | Glycans Pos.
1 | 258006 | 10 | 91 | K.MLTMVPWYS[+972.31694]SR.L | S9(OGlycan / 972.3169) | 559.8 | Hex(6) | 4.5e-9 | 4.9e-6 | 8.35 | 333.0 | 1.2 | 2 | 1171.9774 | 1171.9919 | -12.34 | 2342.9476 | 2342.9765 | Specific | 9 | >YER011W TIR1 SGDI
>YHR174W ENO2 SGDI
>YJL052W TDH1 SGDI
>YFL020C PAU5 SGDID
>YNL160W YGP1 SGDI
>YDR055W PST1 SGDI
>YIL169C YIL169C SGD
>sp|TRYP_PIG|(Comm
>YGR254W ENO1 SGDI
>YGR279C SCW4 SGDI
>YER011W TIR1 SGDID
>YKL060C FBA1 SGDID
>YIL148W RPL40A SGD
>YLR300W EXG1 SGDI
>YIL123W SIM1 SGDID
>YBR078W ECM33 SGI
>YMR006C PLB2 SGDI
>YKR102W FLO10 SGD
>YBR118W TEF2 SGDI
Protein Coverage (double click to dock...
Protein Coverage
Spectrum (double click to dock / undock)
K.MLTMVPWYS[+972]SR.L z=2,scan#=sample=1 period=1 cycle=740 experiment=21,scan time=12.1919
MLTMVPWYSSR
Intensity
m/z
m/z errors, search tol=75.0 ppm

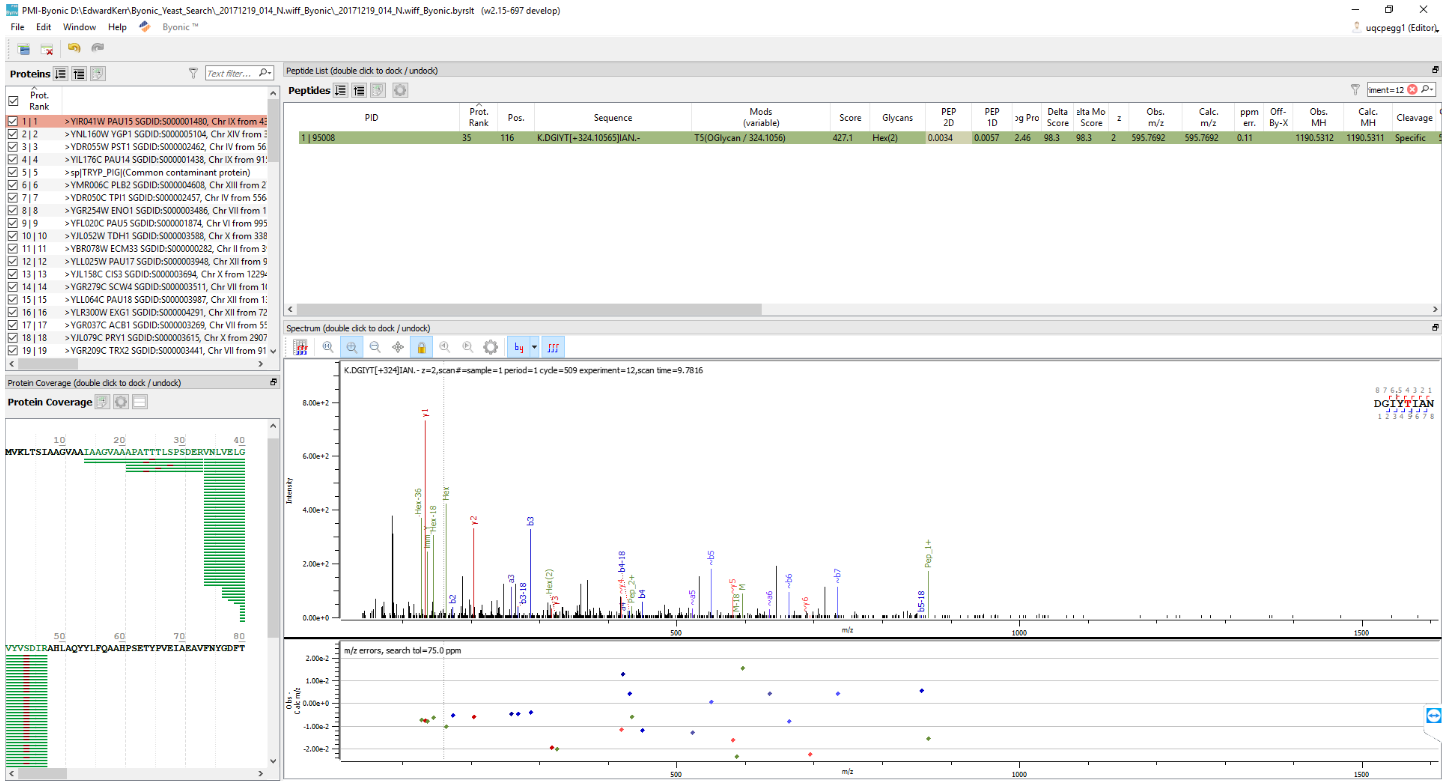

PMI-Byonic D:\EdwardKerr\Byonic_Yeast_Search_20171219_014_N.wiff_Byonic_20171219_014_N.wiff_Byonic.byrslt (v2.15-697 develop)
File Edit Window Help Byonic ™
uqcpegg1 (Editor)
Proteins
Text filter...
Prot. Rank
1 | 1 >YIR041W PAU15 SGDID:S000001480, Chr IX from 43
2 | 2 >YNL160W YGP1 SGDID:S000005104, Chr XIV from 3
3 | 3 >YDR055W PST1 SGDID:S000002462, Chr IV from 56
4 | 4 >YIL176C PAU14 SGDID:S000001438, Chr IX from 91
5 | 5 >sp|TRYP_PIG|(Common contaminant protein)
6 | 6 >YMR006C PLB2 SGDID:S000004608, Chr XIII from 2
7 | 7 >YDR050C TPI1 SGDID:S000002457, Chr IV from 556
8 | 8 >YGR254W ENO1 SGDID:S000003486, Chr VII from 1
9 | 9 >YFL020C PAU5 SGDID:S000001874, Chr VI from 995
10 | 10 >YJL052W TDH1 SGDID:S000003588, Chr X from 338
11 | 11 >YBR078W ECM33 SGDID:S000000282, Chr II from 3
12 | 12 >YLL025W PAU17 SGDID:S000003948, Chr XII from 9
13 | 13 >YJL158C CIS3 SGDID:S000003694, Chr X from 1229
14 | 14 >YGR279C SCW4 SGDID:S000003511, Chr VII from 1
15 | 15 >YLL064C PAU18 SGDID:S000003987, Chr XII from 1
16 | 16 >YLR300W EXG1 SGDID:S000004291, Chr XII from 72
17 | 17 >YGR037C ACB1 SGDID:S000003269, Chr VII from 5
18 | 18 >YJL079C PRY1 SGDID:S000003615, Chr X from 2907
19 | 19 >YGR209C TRX2 SGDID:S000003441, Chr VII from 91
Protein Coverage (double click to dock / undock)
Protein Coverage
MVKLTSIAAGVAAIAAGVAAAPATTTLSPSDERVNLVELG
VYVSDIRAHLAQYYLFQAAHPSETYPVEIAEAVFNYGDFT
Peptide List (double click to dock / undock)
Peptides
PID | Prot. Rank | Pos. | Sequence | Mods (variable) | Score | Glycans | PEP 2D | PEP 1D | og Pro | Delta Score | elta Mo Score | z | Obs. m/z | Calc. m/z | ppm err. | Off-By-X | Obs. MH | Calc. MH | Cleavage
1 | 95008 | 35 | 116 | K.DGIYT[+324.10565]IAN.- | T5(OGlycan / 324.1056) | 427.1 | Hex(2) | 0.0034 | 0.0057 | 2.46 | 98.3 | 98.3 | 2 | 595.7692 | 595.7692 | 0.11 | | 1190.5312 | 1190.5311 | Specific
Spectrum (double click to dock / undock)
K.DGIYT[+324]IAN. - z=2,scan#=sample=1 period=1 cycle=509 experiment=12,scan time=9.7816
DGIYTIAN
Intensity
m/z
m/z errors, search tol=75.0 ppm

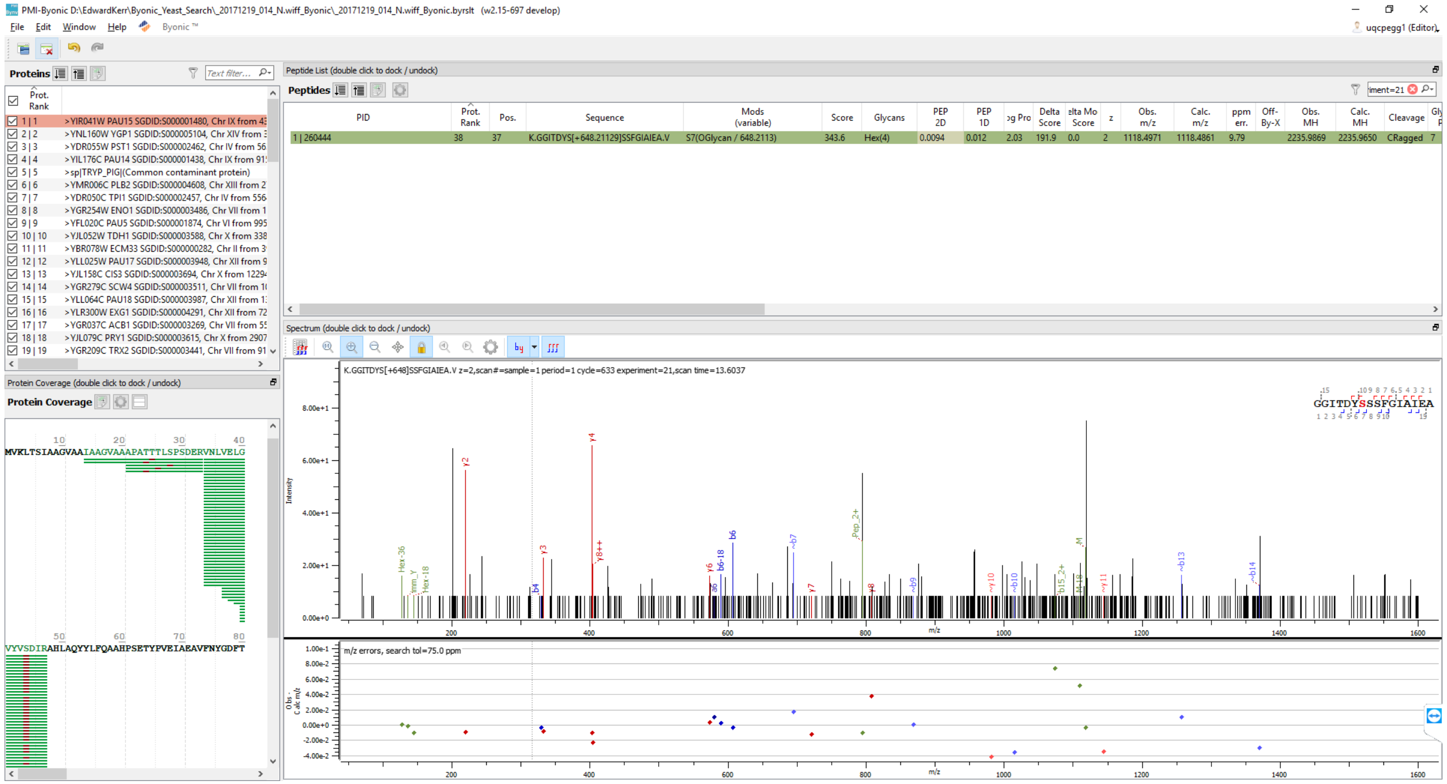
PMI-Byonic D:\EdwardKerr\Byonic_Yeast_Search_20171219_014_N.wiff_Byonic_20171219_014_N.wiff_Byonic.byrslt (w2.15-697 develop)
File Edit Window Help Byonic ™
uqcpegg1 (Editor)
Proteins
Text filter...
Prot. Rank
1 | 1 >YIR041W PAU15 SGDID:S000001480, Chr IX from 43
2 | 2 >YNL160W YGP1 SGDID:S000005104, Chr XIV from 3
3 | 3 >YDR055W PST1 SGDID:S000002462, Chr IV from 56
4 | 4 >YIL176C PAU14 SGDID:S000001438, Chr IX from 91
5 | 5 >sp|TRYP_PIG|(Common contaminant protein)
6 | 6 >YMR006C PLB2 SGDID:S000004608, Chr XIII from 2
7 | 7 >YDR050C TPI1 SGDID:S000002457, Chr IV from 556
8 | 8 >YGR254W ENO1 SGDID:S000003486, Chr VII from 1
9 | 9 >YFL020C PAU5 SGDID:S000001874, Chr VI from 995
10 | 10 >YJL052W TDH1 SGDID:S000003588, Chr X from 338
11 | 11 >YBR078W ECM33 SGDID:S000000282, Chr II from 3
12 | 12 >YLL025W PAU17 SGDID:S000003948, Chr XII from 9
13 | 13 >YJL158C CIS3 SGDID:S000003694, Chr X from 1229
14 | 14 >YGR279C SCW4 SGDID:S000003511, Chr VII from 1
15 | 15 >YLL064C PAU18 SGDID:S000003987, Chr XII from 1
16 | 16 >YLR300W EXG1 SGDID:S000004291, Chr XII from 72
17 | 17 >YGR037C ACB1 SGDID:S000003269, Chr VII from 5
18 | 18 >YJL079C PRY1 SGDID:S000003615, Chr X from 2907
19 | 19 >YGR209C TRX2 SGDID:S000003441, Chr VII from 91
Protein Coverage (double click to dock / undock)
Protein Coverage
10 20 30 40
MVKLTSIAAGVAAIAAGVAAAPATTTLSPSDERVNLVELG
50 60 70 80
VYVSDIRAHLAQYYLFQAAHPSETYPVEIAEAVFNYGDFT
Peptide List (double click to dock / undock)
Peptides
iment=21
PID | Prot. Rank | Pos. | Sequence | Mods (variable) | Score | Glycans | PEP 2D | PEP 1D | og Pro | Delta Score | elta Mo Score | z | Obs. m/z | Calc. m/z | ppm err. | Off-By-X | Obs. MH | Calc. MH | Cleavage | Gly
1 | 260444 | 38 | 37 | K.GGITDYS[+648.21129]SSFGIAIEA.V | S7(OGlycan / 648.2113) | 343.6 | Hex(4) | 0.0094 | 0.012 | 2.03 | 191.9 | 0.0 | 2 | 1118.4971 | 1118.4861 | 9.79 | | 2235.9869 | 2235.9650 | CRagged | 7
Spectrum (double click to dock / undock)
K.GGITDYS[+648]SSFGIAIEA.V z=2,scan#=sample=1 period=1 cycle=633 experiment=21,scan time=13.6037
GGITDYSSSFGIAIEA
Intensity
8.00e+1
6.00e+1
4.00e+1
2.00e+1
0.00e+0
y2
y4
y3
y8++
b4
y6
b6-18
b6
~b7
y7
Pep_2+
~b9
~y10
~b10
~M
~y11
~b13
~b14
Hex-36
Hex-18
200 400 600 800 1000 1200 1400 1600
m/z
m/z errors, search tol=75.0 ppm
1.00e-1
8.00e-2
6.00e-2
4.00e-2
2.00e-2
0.00e+0
-2.00e-2
-4.00e-2
200 400 600 800 1000 1200 1400 1600
m/z

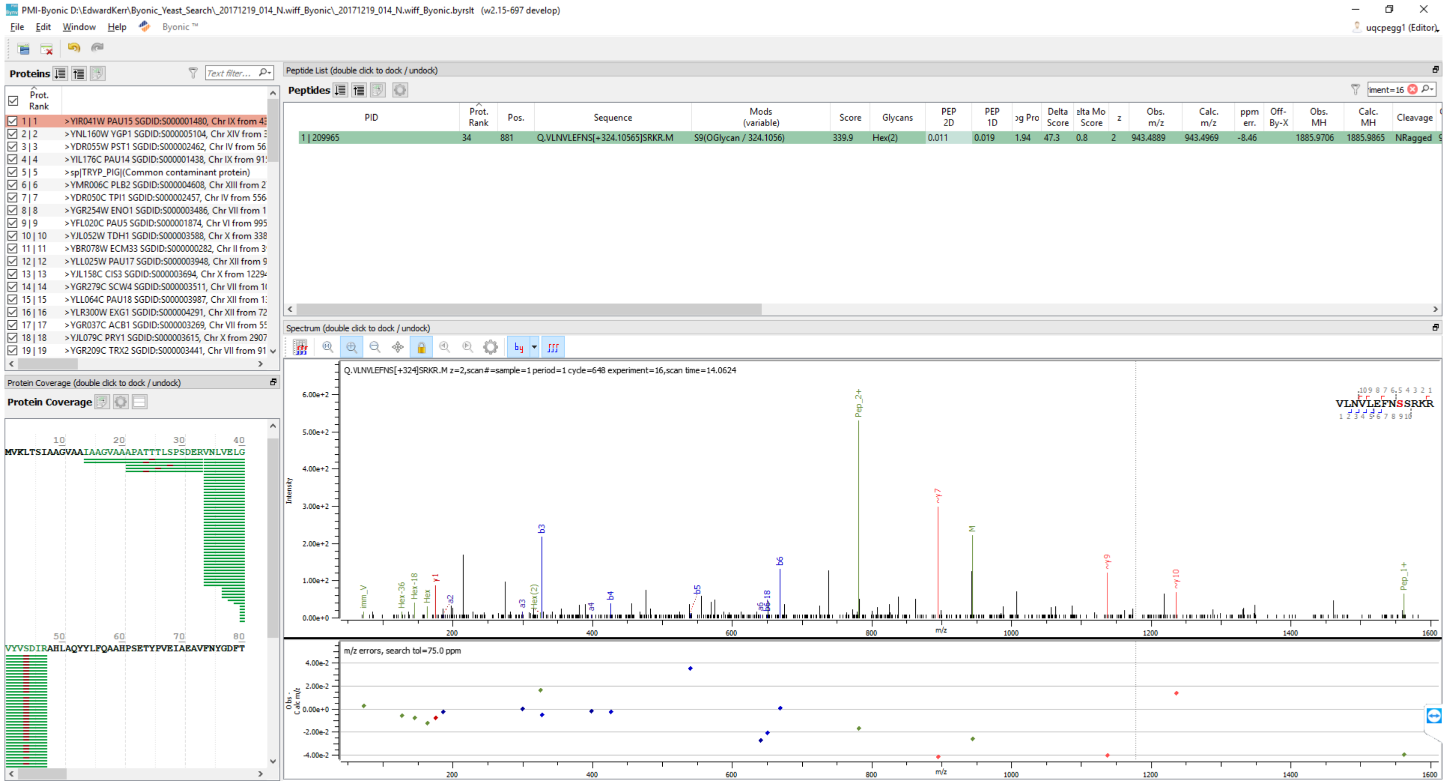

PMI-Byonic D:\EdwardKerr\Byonic_Yeast_Search_20171219_014_N.wiff_Byonic_20171219_014_N.wiff_Byonic.byrslt (w2.15-697 develop)
File Edit Window Help Byonic ™
Proteins
Prot. Rank
1 | 1 >YIR041W PAU15 SGDID:S000001480, Chr IX from 4
2 | 2 >YNL160W YGP1 SGDID:S000005104, Chr XIV from
3 | 3 >YDR055W PST1 SGDID:S000002462, Chr IV from 56
4 | 4 >YIL176C PAU14 SGDID:S000001438, Chr IX from 91
5 | 5 >sp|TRYP_PIG|(Common contaminant protein)
6 | 6 >YMR006C PLB2 SGDID:S000004608, Chr XIII from 2
7 | 7 >YDR050C TPI1 SGDID:S000002457, Chr IV from 556
8 | 8 >YGR254W ENO1 SGDID:S000003486, Chr VII from 1
9 | 9 >YFL020C PAU5 SGDID:S000001874, Chr VI from 995
Peptide List (double click to dock / undock)
Peptides
PID
Prot. Rank
Pos.
Sequence
Mods (variable)
Score
Glycans
1 | 209965
34
881
Q.VLNVLEFNS[+324.10565]SRKR.M
S9(OGlycan / 324.1056)
339.9
Hex(2)
Spectrum (double click to dock / undock)
Protein Coverage (double click to dock / undock)
Protein Coverage
Intensity
m/z
m/z errors, search tol=75.0 ppm

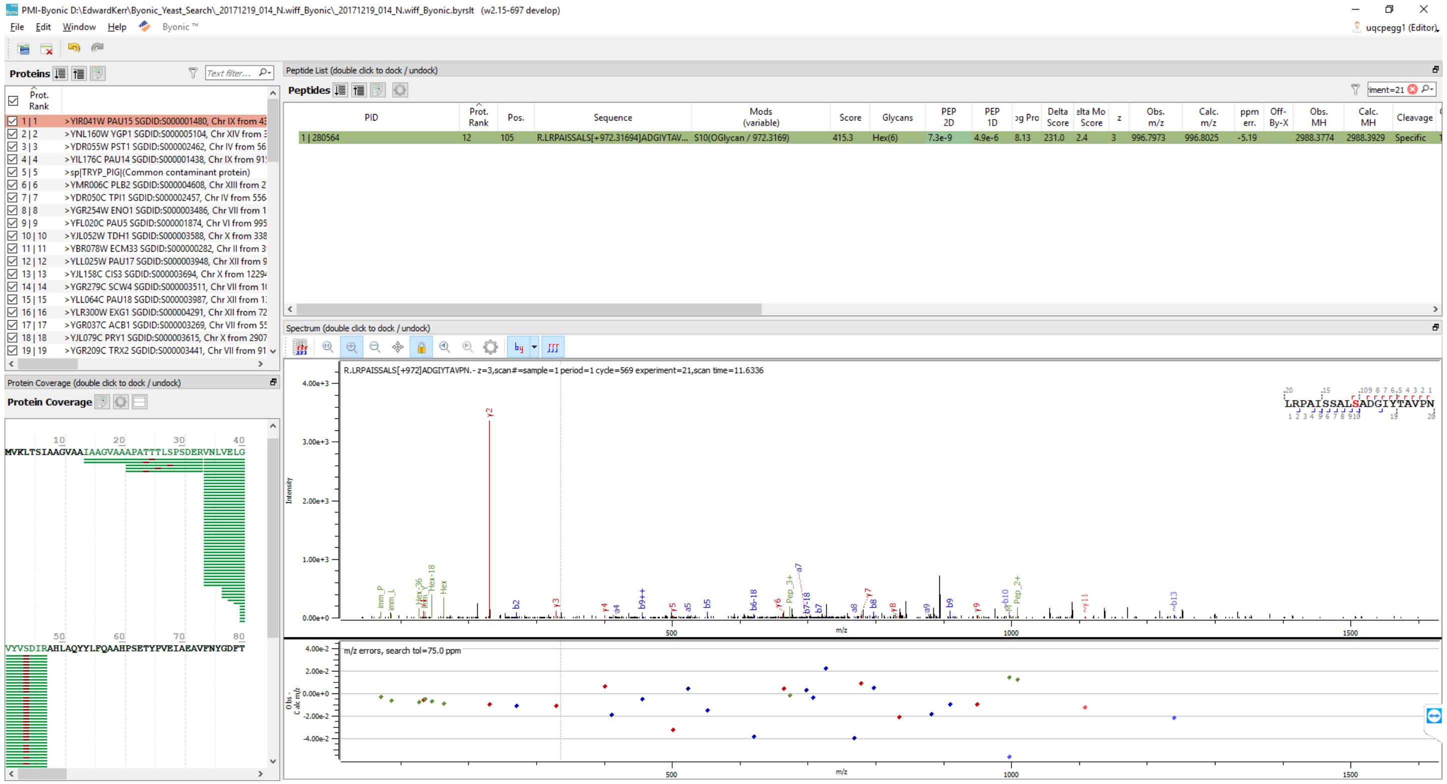
PMI-Byonic D:\EdwardKerr\Byonic_Yeast_Search_20171219_014_N.wiff_Byonic_20171219_014_N.wiff_Byonic.byrslt (w2.15-697 develop)
File Edit Window Help Byonic ™
uqcpegg1 (Editor)
Proteins
Prot. Rank
1 | 1 >YIR041W PAU15 SGDID:S000001480, Chr IX from 4
2 | 2 >YNL160W YGP1 SGDID:S000005104, Chr XIV from
3 | 3 >YDR055W PST1 SGDID:S000002462, Chr IV from 56
4 | 4 >YIL176C PAU14 SGDID:S000001438, Chr IX from 91
5 | 5 >sp|TRYP_PIG|(Common contaminant protein)
6 | 6 >YMR006C PLB2 SGDID:S000004608, Chr XIII from 2
7 | 7 >YDR050C TPI1 SGDID:S000002457, Chr IV from 556
8 | 8 >YGR254W ENO1 SGDID:S000003486, Chr VII from 1
Peptide List (double click to dock / undock)
Peptides
PID | Prot. Rank | Pos. | Sequence | Mods (variable) | Score | Glycans | PEP 2D | PEP 1D | Delta Score | z | Obs. m/z | Calc. m/z | ppm err. | Obs. MH | Calc. MH | Cleavage
1 | 280564 | 12 | 105 | R.LRPAISSALS[+972.31694]ADGIYTAV... | S10(OGlycan / 972.3169) | 415.3 | Hex(6) | 7.3e-9 | 4.9e-6 | 8.13 | 231.0 | 2.4 | 3 | 996.7973 | 996.8025 | -5.19 | 2988.3774 | 2988.3929 | Specific
Spectrum (double click to dock / undock)
R.LRPAISSALS[+972]ADGIYTAVPN.- z=3,scan#=sample=1 period=1 cycle=569 experiment=21,scan time=11.6336
LRPAISSALSADGIYTAVPN
Intensity
m/z
m/z errors, search tol=75.0 ppm
Protein Coverage (double click to dock / undock)
Protein Coverage
MVKLTSIAAGVAAIAAGVAAAPATTTLSPSDERVNLVELG
VYVSDIRAHLAQYYLFQAAHPSETYPVEIAEAVFNYGDFT

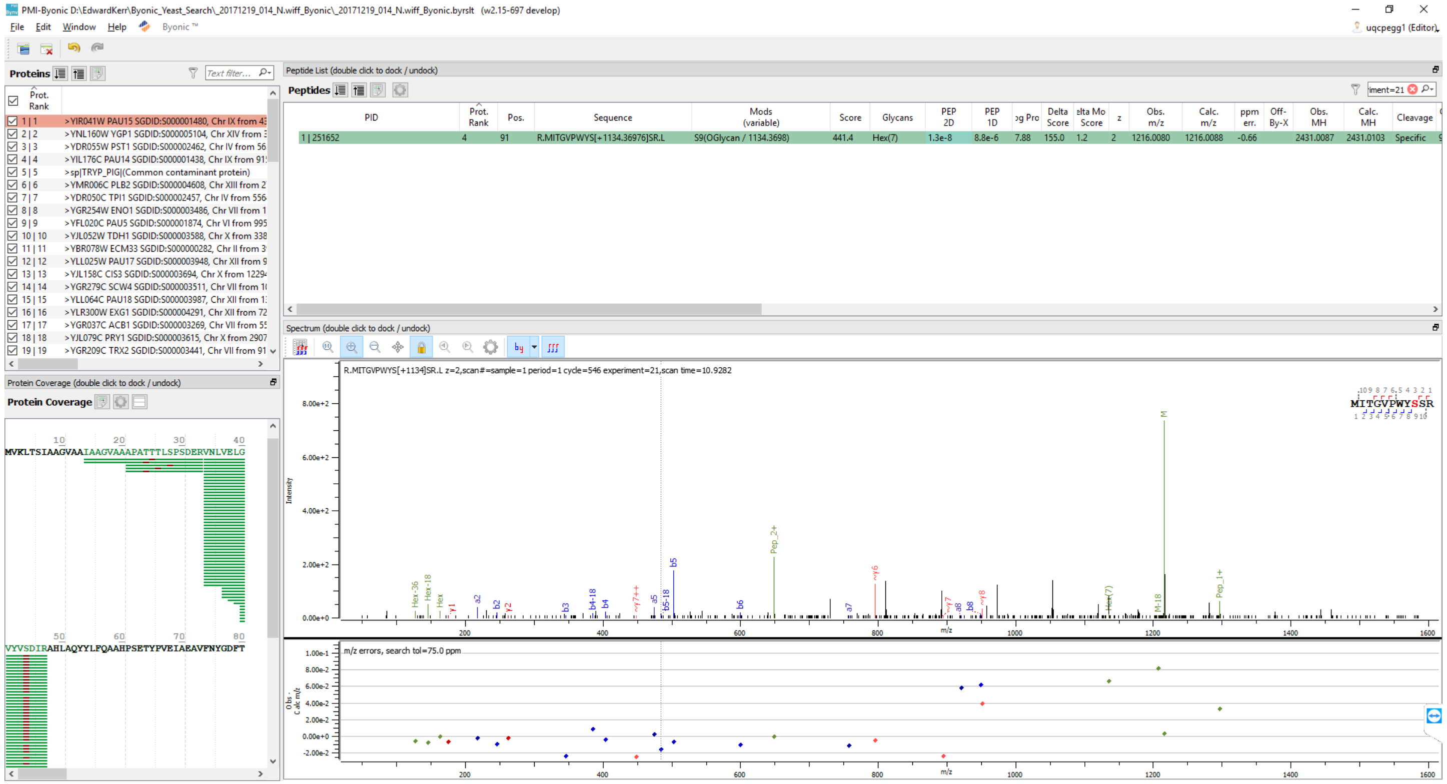
PMI-Byonic D:\EdwardKerr\Byonic_Yeast_Search_20171219_014_N.wiff_Byonic_20171219_014_N.wiff_Byonic.byrslt (w2.15-697 develop)
File Edit Window Help Byonic™
uqcpegg1 (Editor)
Proteins
Prot. Rank
1 | 1 >YIR041W PAU15 SGDID:S000001480, Chr IX from 43
2 | 2 >YNL160W YGP1 SGDID:S000005104, Chr XIV from 3
3 | 3 >YDR055W PST1 SGDID:S000002462, Chr IV from 56
4 | 4 >YIL176C PAU14 SGDID:S000001438, Chr IX from 91
5 | 5 >sp|TRYP_PIG|(Common contaminant protein)
6 | 6 >YMR006C PLB2 SGDID:S000004608, Chr XIII from 2
7 | 7 >YDR050C TPI1 SGDID:S000002457, Chr IV from 556
8 | 8 >YGR254W ENO1 SGDID:S000003486, Chr VII from 1
9 | 9 >YFL020C PAU5 SGDID:S000001874, Chr VI from 995
10 | 10 >YJL052W TDH1 SGDID:S000003588, Chr X from 338
11 | 11 >YBR078W ECM33 SGDID:S000000282, Chr II from 3
12 | 12 >YLL025W PAU17 SGDID:S000003948, Chr XII from 9
13 | 13 >YJL158C CIS3 SGDID:S000003694, Chr X from 1229
14 | 14 >YGR279C SCW4 SGDID:S000003511, Chr VII from 1
15 | 15 >YLL064C PAU18 SGDID:S000003987, Chr XII from 1
16 | 16 >YLR300W EXG1 SGDID:S000004291, Chr XII from 72
17 | 17 >YGR037C ACB1 SGDID:S000003269, Chr VII from 5
18 | 18 >YJL079C PRY1 SGDID:S000003615, Chr X from 2907
19 | 19 >YGR209C TRX2 SGDID:S000003441, Chr VII from 91
Protein Coverage (double click to dock / undock)
Protein Coverage
MVKLTSIAAGVAAIAAGVAAAPATTTLSPSDERVNLVELG
VYVSDIRAHLAQYYLFQAAHPSETYPVEIAEAVFNYGDFT
Peptide List (double click to dock / undock)
Peptides
PID | Prot. Rank | Pos. | Sequence | Mods (variable) | Score | Glycans | PEP 2D | PEP 1D | og Pro | Delta Score | elta Mo Score | z | Obs. m/z | Calc. m/z | ppm err. | Off-By-X | Obs. MH | Calc. MH | Cleavage
1 | 251652 | 4 | 91 | R.MITGVPWYS[+1134.36976]SR.L | S9(OGlycan / 1134.3698) | 441.4 | Hex(7) | 1.3e-8 | 8.8e-6 | 7.88 | 155.0 | 1.2 | 2 | 1216.0080 | 1216.0088 | -0.66 | 2431.0087 | 2431.0103 | Specific
Spectrum (double click to dock / undock)
R.MITGVPWYS[+1134]SR.L z=2,scan#=sample=1 period=1 cycle=546 experiment=21,scan time=10.9282
MITGVPWYSSR
Intensity
m/z
m/z errors, search tol=75.0 ppm
Obs. - Calc. m/z

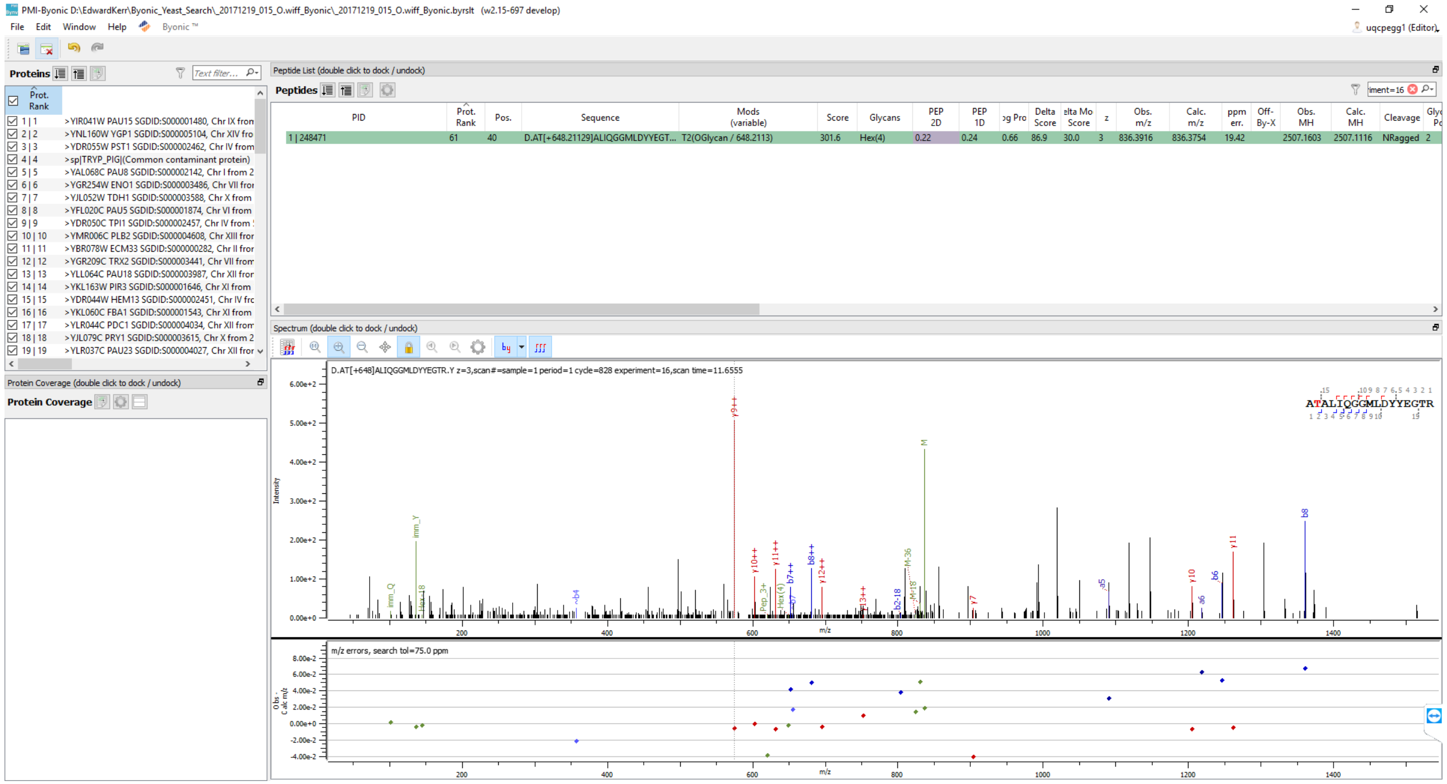
PMI-Byonic D:\EdwardKerr\Byonic_Yeast_Search_20171219_015_O.wiff_Byonic_20171219_015_O.wiff_Byonic.byrslt (w2.15-697 develop)
File Edit Window Help Byonic ™
uqcpegg1 (Editor)
Proteins
Peptide List (double click to dock / undock)
Peptides
PID | Prot. Rank | Pos. | Sequence | Mods (variable) | Score | Glycans | PEP 2D | PEP 1D | Delta Score | z | Obs. m/z | Calc. m/z | ppm err. | Off-By-X | Obs. MH | Calc. MH | Cleavage
1 | 248471 | 61 | 40 | D.AT[+648.21129]ALIQGGMLDYYEGT... | T2(OGlycan / 648.2113) | 301.6 | Hex(4) | 0.22 | 0.24 | 0.66 | 86.9 | 30.0 | 3 | 836.3916 | 836.3754 | 19.42 | | 2507.1603 | 2507.1116 | NRagged | 2
Spectrum (double click to dock / undock)
D.AT[+648]ALIQGGMLDYYEGTR.Y z=3,scan#=sample=1 period=1 cycle=828 experiment=16,scan time=11.6555
Intensity
m/z
m/z errors, search tol=75.0 ppm
Protein Coverage (double click to dock / undock)
Protein Coverage

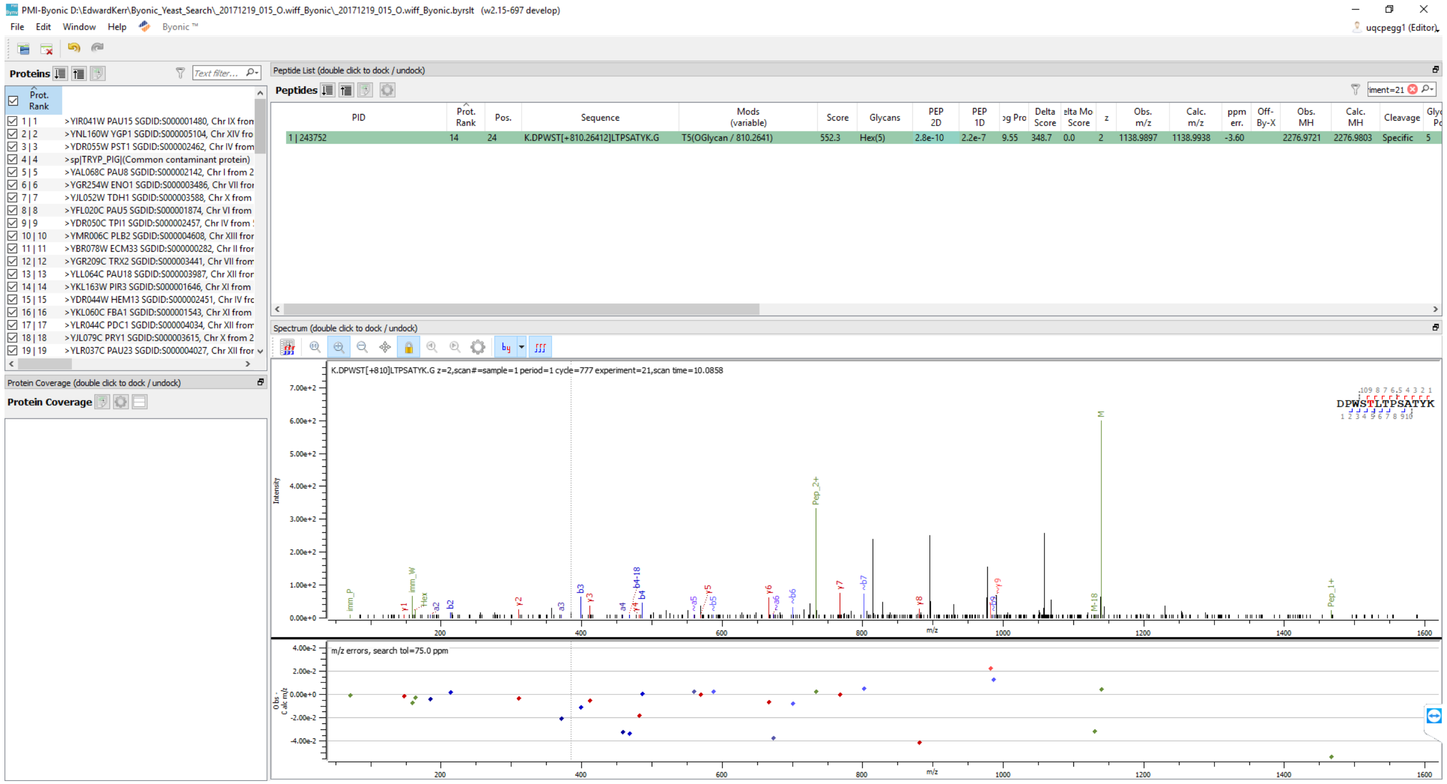
PMI-Byonic D:\EdwardKerr\Byonic_Yeast_Search_20171219_015_O.wiff_Byonic_20171219_015_O.wiff_Byonic.byrslt (w2.15-697 develop)
File Edit Window Help Byonic ™
uqcpegg1 (Editor)
Proteins
Prot. Rank
1 | 1 >YIR041W PAU15 SGDID:S000001480, Chr IX from
2 | 2 >YNL160W YGP1 SGDID:S000005104, Chr XIV fro
3 | 3 >YDR055W PST1 SGDID:S000002462, Chr IV from
4 | 4 >sp|TRYP_PIG|(Common contaminant protein)
5 | 5 >YAL068C PAU8 SGDID:S000002142, Chr I from 2
6 | 6 >YGR254W ENO1 SGDID:S000003486, Chr VII fro
7 | 7 >YJL052W TDH1 SGDID:S000003588, Chr X from
8 | 8 >YFL020C PAU5 SGDID:S000001874, Chr VI from
9 | 9 >YDR050C TPI1 SGDID:S000002457, Chr IV from
10 | 10 >YMR006C PLB2 SGDID:S000004608, Chr XIII fro
11 | 11 >YBR078W ECM33 SGDID:S000000282, Chr II fro
12 | 12 >YGR209C TRX2 SGDID:S000003441, Chr VII from
13 | 13 >YLL064C PAU18 SGDID:S000003987, Chr XII fro
14 | 14 >YKL163W PIR3 SGDID:S000001646, Chr XI from
15 | 15 >YDR044W HEM13 SGDID:S000002451, Chr IV fro
16 | 16 >YKL060C FBA1 SGDID:S000001543, Chr XI from
17 | 17 >YLR044C PDC1 SGDID:S000004034, Chr XII from
18 | 18 >YJL079C PRY1 SGDID:S000003615, Chr X from 2
19 | 19 >YLR037C PAU23 SGDID:S000004027, Chr XII fro
Protein Coverage (double click to dock / undock)
Protein Coverage
Peptide List (double click to dock / undock)
Peptides
PID | Prot. Rank | Pos. | Sequence | Mods (variable) | Score | Glycans | PEP 2D | PEP 1D | Delta Score | Delta Mod Score | z | Obs. m/z | Calc. m/z | ppm err. | Off-By-X | Obs. MH | Calc. MH | Cleavage
1 | 243752 | 14 | 24 | K.DPWST[+810.26412]LTPSATYK.G | T5(OGlycan / 810.2641) | 552.3 | Hex(5) | 2.8e-10 | 2.2e-7 | 9.55 | 348.7 | 0.0 | 2 | 1138.9897 | 1138.9938 | -3.60 | 2276.9721 | 2276.9803 | Specific | 5
Spectrum (double click to dock / undock)
K.DPWST[+810]LTPSATYK.G z=2,scan#=sample=1 period=1 cycle=777 experiment=21,scan time=10.0858
DPWSTLTPSATYK
Intensity
m/z
M
Pep_2+
Pep_1+
M-18
b4-18
m/z errors, search tol=75.0 ppm
Obs - Calc m/z

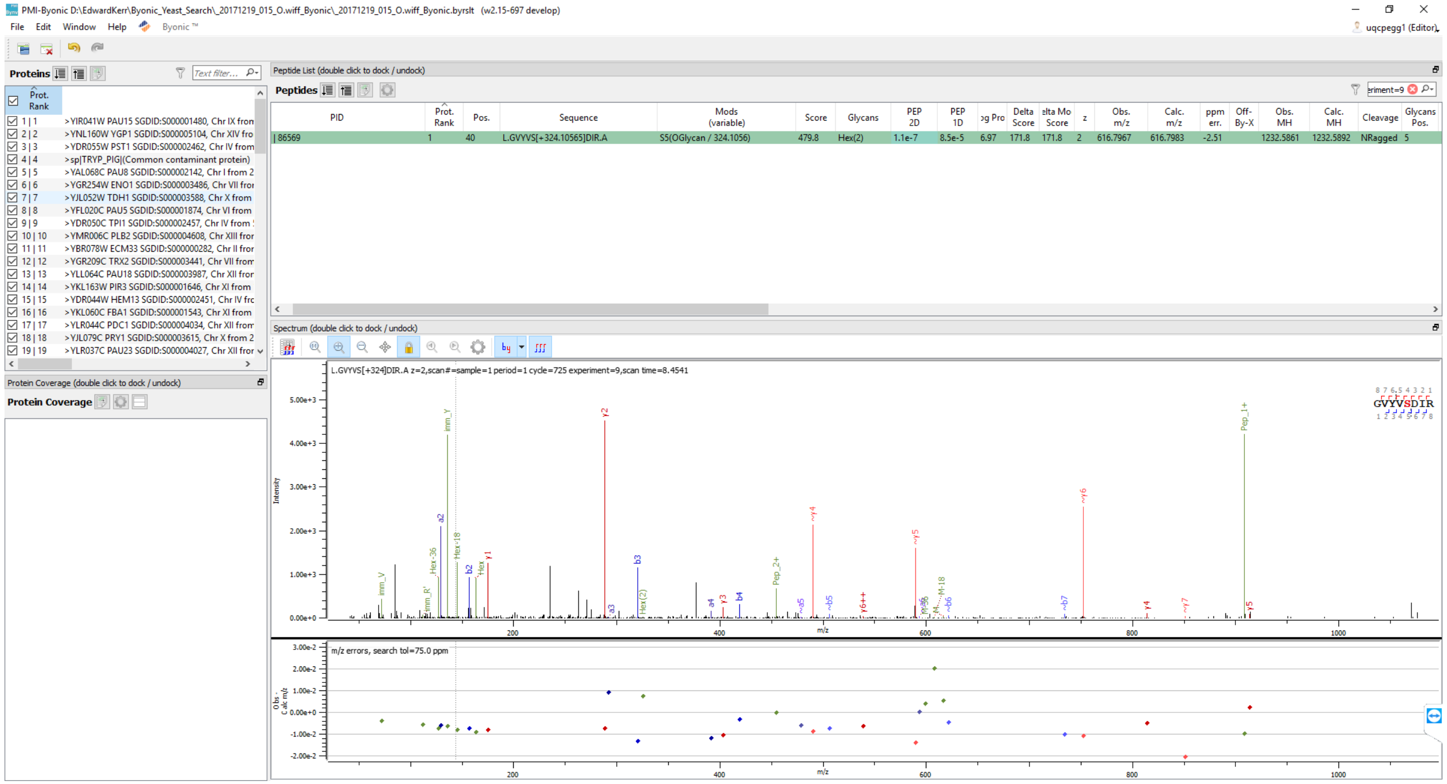

PMI-Byonic D:\EdwardKerr\Byonic_Yeast_Search_20171219_015_O.wiff_Byonic_20171219_015_O.wiff_Byonic.byrslt (w2.15-697 develop)
File Edit Window Help Byonic ™
uqcpegg1 (Editor)
Proteins
Text filter...
Prot. Rank
1 | 1 >YIR041W PAU15 SGDID:S000001480, Chr IX from
2 | 2 >YNL160W YGP1 SGDID:S000005104, Chr XIV fro
3 | 3 >YDR055W PST1 SGDID:S000002462, Chr IV from
4 | 4 >sp|TRYP_PIG|(Common contaminant protein)
5 | 5 >YAL068C PAU8 SGDID:S000002142, Chr I from 2
6 | 6 >YGR254W ENO1 SGDID:S000003486, Chr VII fro
7 | 7 >YJL052W TDH1 SGDID:S000003588, Chr X from
8 | 8 >YFL020C PAU5 SGDID:S000001874, Chr VI from
9 | 9 >YDR050C TPI1 SGDID:S000002457, Chr IV from
10 | 10 >YMR006C PLB2 SGDID:S000004608, Chr XIII fro
11 | 11 >YBR078W ECM33 SGDID:S000000282, Chr II fro
12 | 12 >YGR209C TRX2 SGDID:S000003441, Chr VII from
13 | 13 >YLL064C PAU18 SGDID:S000003987, Chr XII fro
14 | 14 >YKL163W PIR3 SGDID:S000001646, Chr XI from
15 | 15 >YDR044W HEM13 SGDID:S000002451, Chr IV fro
16 | 16 >YKL060C FBA1 SGDID:S000001543, Chr XI from
17 | 17 >YLR044C PDC1 SGDID:S000004034, Chr XII from
18 | 18 >YJL079C PRY1 SGDID:S000003615, Chr X from 2
19 | 19 >YLR037C PAU23 SGDID:S000004027, Chr XII fro
Protein Coverage (double click to dock / undock)
Protein Coverage
Peptide List (double click to dock / undock)
Peptides
eriment=9
PID | Prot. Rank | Pos. | Sequence | Mods (variable) | Score | Glycans | PEP 2D | PEP 1D | og Pro | Delta Score | elta Mo Score | z | Obs. m/z | Calc. m/z | ppm err. | Off-By-X | Obs. MH | Calc. MH | Cleavage | Glycans Pos.
| 86569 | 1 | 40 | L.GVYVS[+324.10565]DIR.A | S5(OGlycan / 324.1056) | 479.8 | Hex(2) | 1.1e-7 | 8.5e-5 | 6.97 | 171.8 | 171.8 | 2 | 616.7967 | 616.7983 | -2.51 | | 1232.5861 | 1232.5892 | NRagged | 5
Spectrum (double click to dock / undock)
L.GVYVS[+324]DIR.A z=2,scan#=sample=1 period=1 cycle=725 experiment=9,scan time=8.4541
Intensity
5.00e+3
4.00e+3
3.00e+3
2.00e+3
1.00e+3
0.00e+0
imm_Y
y2
Pep_1+
~y6
~y4
~y5
a2
b3
b2
y1
Pep_2+
GVYVSDIR
200
400
600
800
1000
m/z
m/z errors, search tol=75.0 ppm
3.00e-2
2.00e-2
1.00e-2
0.00e+0
-1.00e-2
-2.00e-2
Obs - Calc m/z

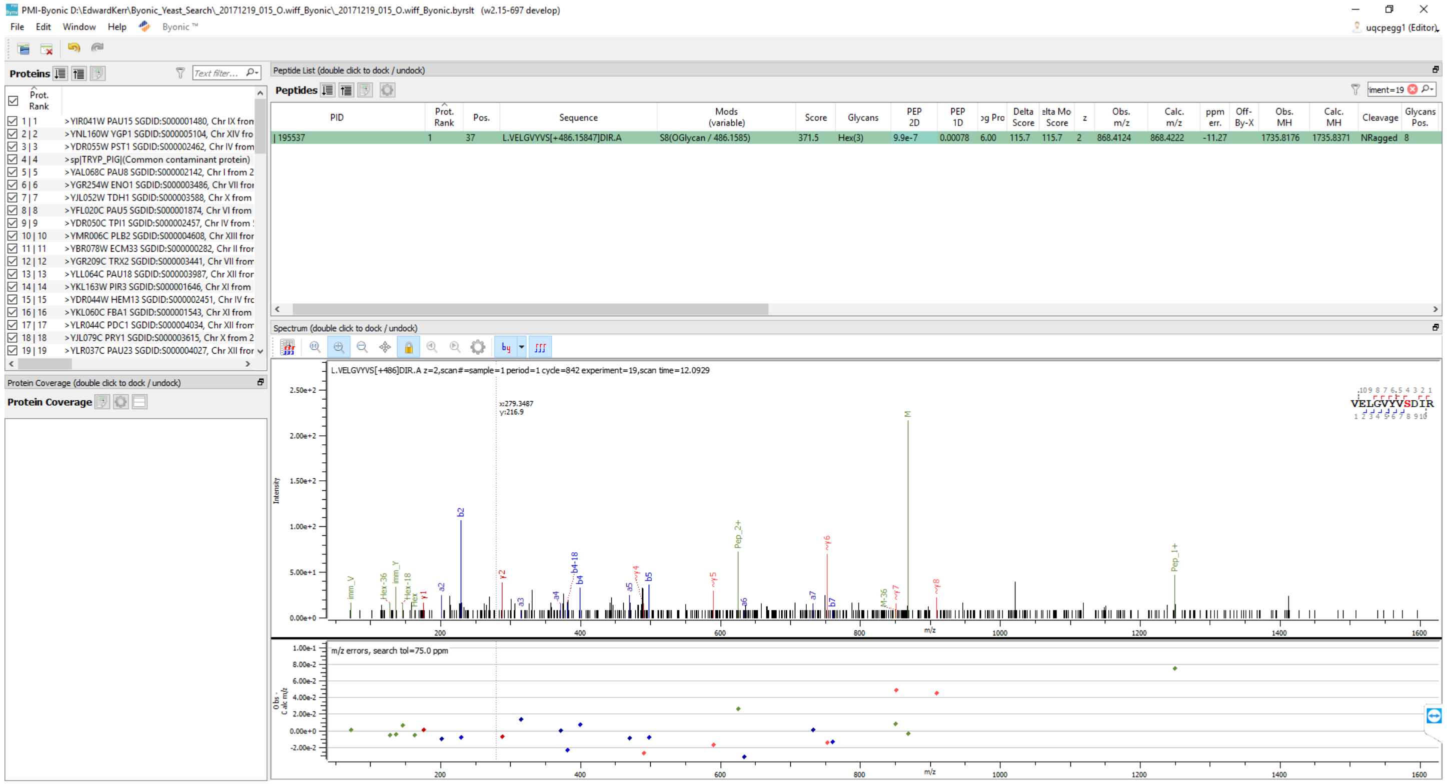
PMI-Byonic D:\EdwardKerr\Byonic_Yeast_Search_20171219_015_O.wiff_Byonic_20171219_015_O.wiff_Byonic.byrslt (w2.15-697 develop)
File Edit Window Help Byonic ™
uqcpegg1 (Editor)
Proteins
Text filter...
Prot. Rank
1 | 1 >YIR041W PAU15 SGDID:S000001480, Chr IX from
2 | 2 >YNL160W YGP1 SGDID:S000005104, Chr XIV fro
3 | 3 >YDR055W PST1 SGDID:S000002462, Chr IV from
4 | 4 >sp|TRYP_PIG|(Common contaminant protein)
5 | 5 >YAL068C PAU8 SGDID:S000002142, Chr I from 2
6 | 6 >YGR254W ENO1 SGDID:S000003486, Chr VII fron
7 | 7 >YJL052W TDH1 SGDID:S000003588, Chr X from
8 | 8 >YFL020C PAU5 SGDID:S000001874, Chr VI from
9 | 9 >YDR050C TPI1 SGDID:S000002457, Chr IV from
10 | 10 >YMR006C PLB2 SGDID:S000004608, Chr XIII fror
11 | 11 >YBR078W ECM33 SGDID:S000000282, Chr II fror
12 | 12 >YGR209C TRX2 SGDID:S000003441, Chr VII from
13 | 13 >YLL064C PAU18 SGDID:S000003987, Chr XII fror
14 | 14 >YKL163W PIR3 SGDID:S000001646, Chr XI from
15 | 15 >YDR044W HEM13 SGDID:S000002451, Chr IV frc
16 | 16 >YKL060C FBA1 SGDID:S000001543, Chr XI from
17 | 17 >YLR044C PDC1 SGDID:S000004034, Chr XII from
18 | 18 >YJL079C PRY1 SGDID:S000003615, Chr X from 2
19 | 19 >YLR037C PAU23 SGDID:S000004027, Chr XII fror
Protein Coverage (double click to dock / undock)
Protein Coverage
Peptide List (double click to dock / undock)
Peptides
iment=19
PID | Prot. Rank | Pos. | Sequence | Mods (variable) | Score | Glycans | PEP 2D | PEP 1D | og Pro | Delta Score | elta Mo Score | z | Obs. m/z | Calc. m/z | ppm err. | Off-By-X | Obs. MH | Calc. MH | Cleavage | Glycans Pos.
| 195537 | 1 | 37 | L.VELGVYVS[+486.15847]DIR.A | S8(OGlycan / 486.1585) | 371.5 | Hex(3) | 9.9e-7 | 0.00078 | 6.00 | 115.7 | 115.7 | 2 | 868.4124 | 868.4222 | -11.27 | | 1735.8176 | 1735.8371 | NRagged | 8
Spectrum (double click to dock / undock)
L.VELGVYVS[+486]DIR.A z=2,scan#=sample=1 period=1 cycle=842 experiment=19,scan time=12.0929
VELGVYVSDIR
Intensity
m/z
Pep_2+
Pep_1+
m/z errors, search tol=75.0 ppm

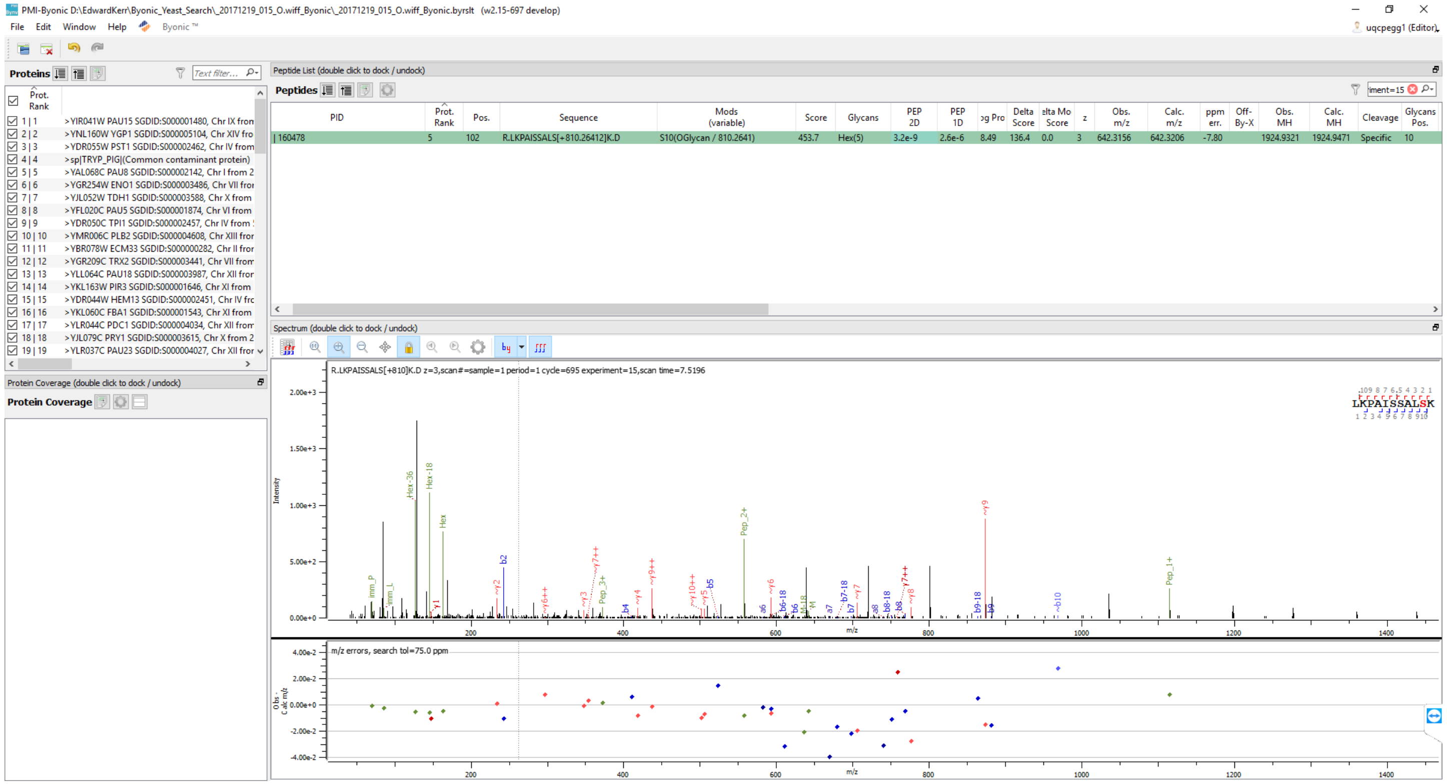
PMI-Byonic D:\EdwardKerr\Byonic_Yeast_Search_20171219_015_O.wiff_Byonic_20171219_015_O.wiff_Byonic.byrslt (w2.15-697 develop)
File Edit Window Help Byonic ™
uqcpegg1 (Editor)
Proteins
Prot. Rank
1 | 1 >YIR041W PAU15 SGDID:S000001480, Chr IX from
2 | 2 >YNL160W YGP1 SGDID:S000005104, Chr XIV fro
3 | 3 >YDR055W PST1 SGDID:S000002462, Chr IV from
4 | 4 >sp|TRYP_PIG|(Common contaminant protein)
5 | 5 >YAL068C PAU8 SGDID:S000002142, Chr I from 2
6 | 6 >YGR254W ENO1 SGDID:S000003486, Chr VII fro
7 | 7 >YJL052W TDH1 SGDID:S000003588, Chr X from
8 | 8 >YFL020C PAU5 SGDID:S000001874, Chr VI from
9 | 9 >YDR050C TPI1 SGDID:S000002457, Chr IV from
10 | 10 >YMR006C PLB2 SGDID:S000004608, Chr XIII fro
11 | 11 >YBR078W ECM33 SGDID:S000000282, Chr II fro
12 | 12 >YGR209C TRX2 SGDID:S000003441, Chr VII fro
13 | 13 >YLL064C PAU18 SGDID:S000003987, Chr XII fro
14 | 14 >YKL163W PIR3 SGDID:S000001646, Chr XI from
15 | 15 >YDR044W HEM13 SGDID:S000002451, Chr IV fro
16 | 16 >YKL060C FBA1 SGDID:S000001543, Chr XI from
17 | 17 >YLR044C PDC1 SGDID:S000004034, Chr XII fro
18 | 18 >YJL079C PRY1 SGDID:S000003615, Chr X from 2
19 | 19 >YLR037C PAU23 SGDID:S000004027, Chr XII fro
Protein Coverage (double click to dock / undock)
Protein Coverage
Peptide List (double click to dock / undock)
Peptides
PID | Prot. Rank | Pos. | Sequence | Mods (variable) | Score | Glycans | PEP 2D | PEP 1D | og Pro | Delta Score | elta Mo Score | z | Obs. m/z | Calc. m/z | ppm err. | Off-By-X | Obs. MH | Calc. MH | Cleavage | Glycans Pos.
160478 | 5 | 102 | R.LKPAISSALS[+810.26412]K.D | S10(OGlycan / 810.2641) | 453.7 | Hex(5) | 3.2e-9 | 2.6e-6 | 8.49 | 136.4 | 0.0 | 3 | 642.3156 | 642.3206 | -7.80 | | 1924.9321 | 1924.9471 | Specific | 10
Spectrum (double click to dock / undock)
R.LKPAISSALS[+810]K.D z=3,scan#=sample=1 period=1 cycle=695 experiment=15,scan time=7.5196
LKPAISSALSK
Intensity
m/z
m/z errors, search tol=75.0 ppm
Obs - Calc m/z

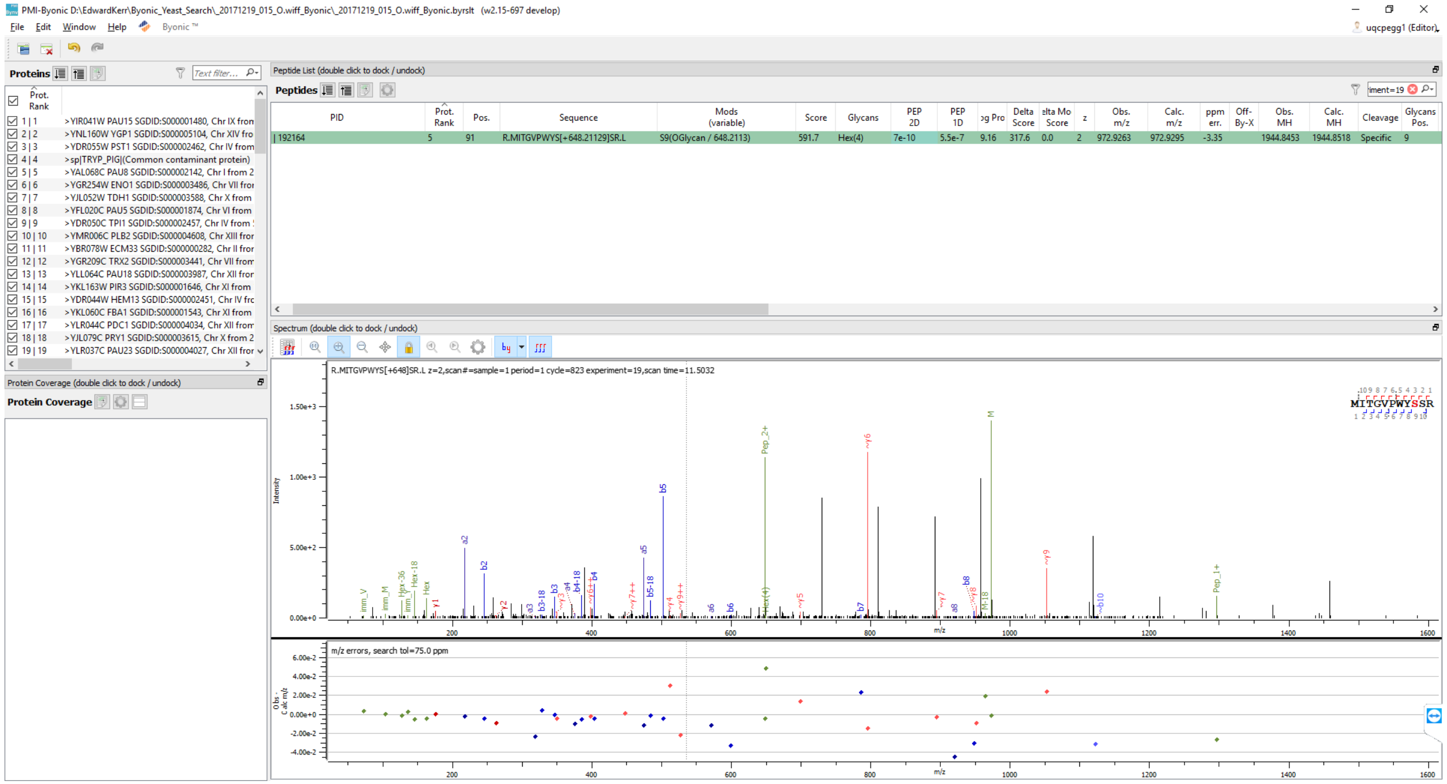
PMI-Byonic D:\EdwardKerr\Byonic_Yeast_Search_20171219_015_O.wiff_Byonic_20171219_015_O.wiff_Byonic.byrslt (w2.15-697 develop)
File Edit Window Help Byonic ™
uqcpegg1 (Editor)
Proteins
Prot. Rank
1 | 1 >YIR041W PAU15 SGDID:S000001480, Chr IX from
2 | 2 >YNL160W YGP1 SGDID:S000005104, Chr XIV fro
3 | 3 >YDR055W PST1 SGDID:S000002462, Chr IV from
4 | 4 >sp|TRYP_PIG|(Common contaminant protein)
5 | 5 >YAL068C PAU8 SGDID:S000002142, Chr I from 2
6 | 6 >YGR254W ENO1 SGDID:S000003486, Chr VII fro
7 | 7 >YJL052W TDH1 SGDID:S000003588, Chr X from
8 | 8 >YFL020C PAU5 SGDID:S000001874, Chr VI from
9 | 9 >YDR050C TPI1 SGDID:S000002457, Chr IV from
10 | 10 >YMR006C PLB2 SGDID:S000004608, Chr XIII fro
11 | 11 >YBR078W ECM33 SGDID:S000000282, Chr II fro
12 | 12 >YGR209C TRX2 SGDID:S000003441, Chr VII fro
13 | 13 >YLL064C PAU18 SGDID:S000003987, Chr XII fro
14 | 14 >YKL163W PIR3 SGDID:S000001646, Chr XI from
15 | 15 >YDR044W HEM13 SGDID:S000002451, Chr IV fro
16 | 16 >YKL060C FBA1 SGDID:S000001543, Chr XI from
17 | 17 >YLR044C PDC1 SGDID:S000004034, Chr XII fro
18 | 18 >YJL079C PRY1 SGDID:S000003615, Chr X from 2
19 | 19 >YLR037C PAU23 SGDID:S000004027, Chr XII fro
Protein Coverage (double click to dock / undock)
Protein Coverage
Peptide List (double click to dock / undock)
Peptides
PID | Prot. Rank | Pos. | Sequence | Mods (variable) | Score | Glycans | PEP 2D | PEP 1D | Log Prob | Delta Score | Delta Mod Score | z | Obs. m/z | Calc. m/z | ppm err. | Off-By-X | Obs. MH | Calc. MH | Cleavage | Glycans Pos.
192164 | 5 | 91 | R.MITGVPWYS[+648.21129]SR.L | S9(OGlycan / 648.2113) | 591.7 | Hex(4) | 7e-10 | 5.5e-7 | 9.16 | 317.6 | 0.0 | 2 | 972.9263 | 972.9295 | -3.35 | | 1944.8453 | 1944.8518 | Specific | 9
Spectrum (double click to dock / undock)
R.MITGVPWYS[+648]SR.L z=2,scan#=sample=1 period=1 cycle=823 experiment=19,scan time=11.5032
MITGVPWYSSR
Intensity
m/z
m/z errors, search tol=75.0 ppm
Obs - Calc m/z

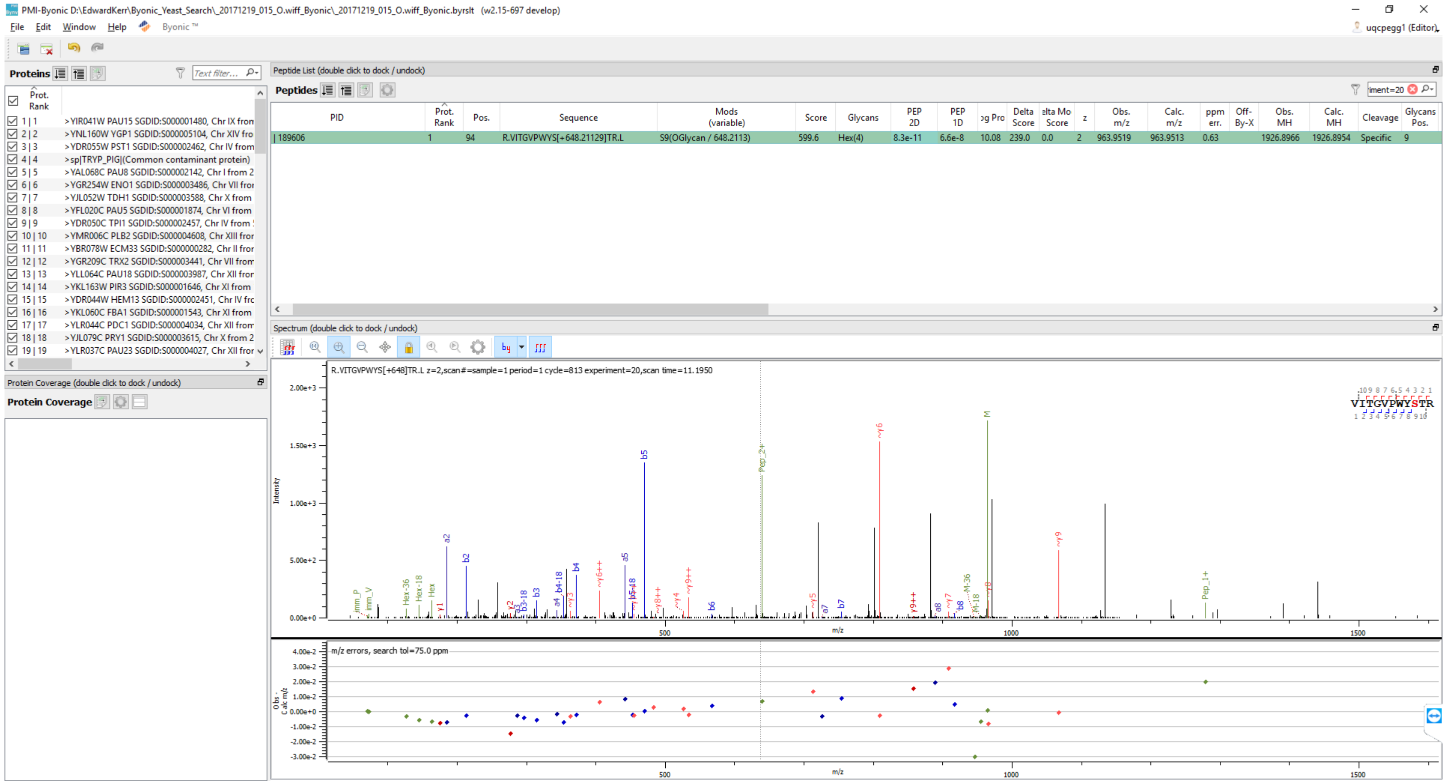
PMI-Byonic D:\EdwardKerr\Byonic_Yeast_Search_20171219_015_O.wiff_Byonic_20171219_015_O.wiff_Byonic.byrslt (w2.15-697 develop)
File Edit Window Help Byonic ™
uqcpegg1 (Editor)
Proteins
Text filter...
Prot. Rank
1 | 1 >YIR041W PAU15 SGDID:S000001480, Chr IX from
2 | 2 >YNL160W YGP1 SGDID:S000005104, Chr XIV fro
3 | 3 >YDR055W PST1 SGDID:S000002462, Chr IV from
4 | 4 >sp|TRYP_PIG|(Common contaminant protein)
5 | 5 >YAL068C PAU8 SGDID:S000002142, Chr I from 2
6 | 6 >YGR254W ENO1 SGDID:S000003486, Chr VII fror
7 | 7 >YJL052W TDH1 SGDID:S000003588, Chr X from
8 | 8 >YFL020C PAU5 SGDID:S000001874, Chr VI from
9 | 9 >YDR050C TPI1 SGDID:S000002457, Chr IV from
10 | 10 >YMR006C PLB2 SGDID:S000004608, Chr XIII fror
11 | 11 >YBR078W ECM33 SGDID:S000000282, Chr II fror
12 | 12 >YGR209C TRX2 SGDID:S000003441, Chr VII from
13 | 13 >YLL064C PAU18 SGDID:S000003987, Chr XII fror
14 | 14 >YKL163W PIR3 SGDID:S000001646, Chr XI from
15 | 15 >YDR044W HEM13 SGDID:S000002451, Chr IV frc
16 | 16 >YKL060C FBA1 SGDID:S000001543, Chr XI from
17 | 17 >YLR044C PDC1 SGDID:S000004034, Chr XII from
18 | 18 >YJL079C PRY1 SGDID:S000003615, Chr X from 2
19 | 19 >YLR037C PAU23 SGDID:S000004027, Chr XII fror
Protein Coverage (double click to dock / undock)
Protein Coverage
Peptide List (double click to dock / undock)
Peptides
iment=20
PID | Prot. Rank | Pos. | Sequence | Mods (variable) | Score | Glycans | PEP 2D | PEP 1D | og Pro | Delta Score | elta Mo Score | z | Obs. m/z | Calc. m/z | ppm err. | Off-By-X | Obs. MH | Calc. MH | Cleavage | Glycans Pos.
| 189606 | 1 | 94 | R.VITGVPWYS[+648.21129]TR.L | S9(OGlycan / 648.2113) | 599.6 | Hex(4) | 8.3e-11 | 6.6e-8 | 10.08 | 239.0 | 0.0 | 2 | 963.9519 | 963.9513 | 0.63 | | 1926.8966 | 1926.8954 | Specific | 9
Spectrum (double click to dock / undock)
R.VITGVPWYS[+648]TR.L z=2,scan#=sample=1 period=1 cycle=813 experiment=20,scan time=11.1950
VITGVPWYSTR
Intensity
m/z
m/z errors, search tol=75.0 ppm
Obs - Calc m/z

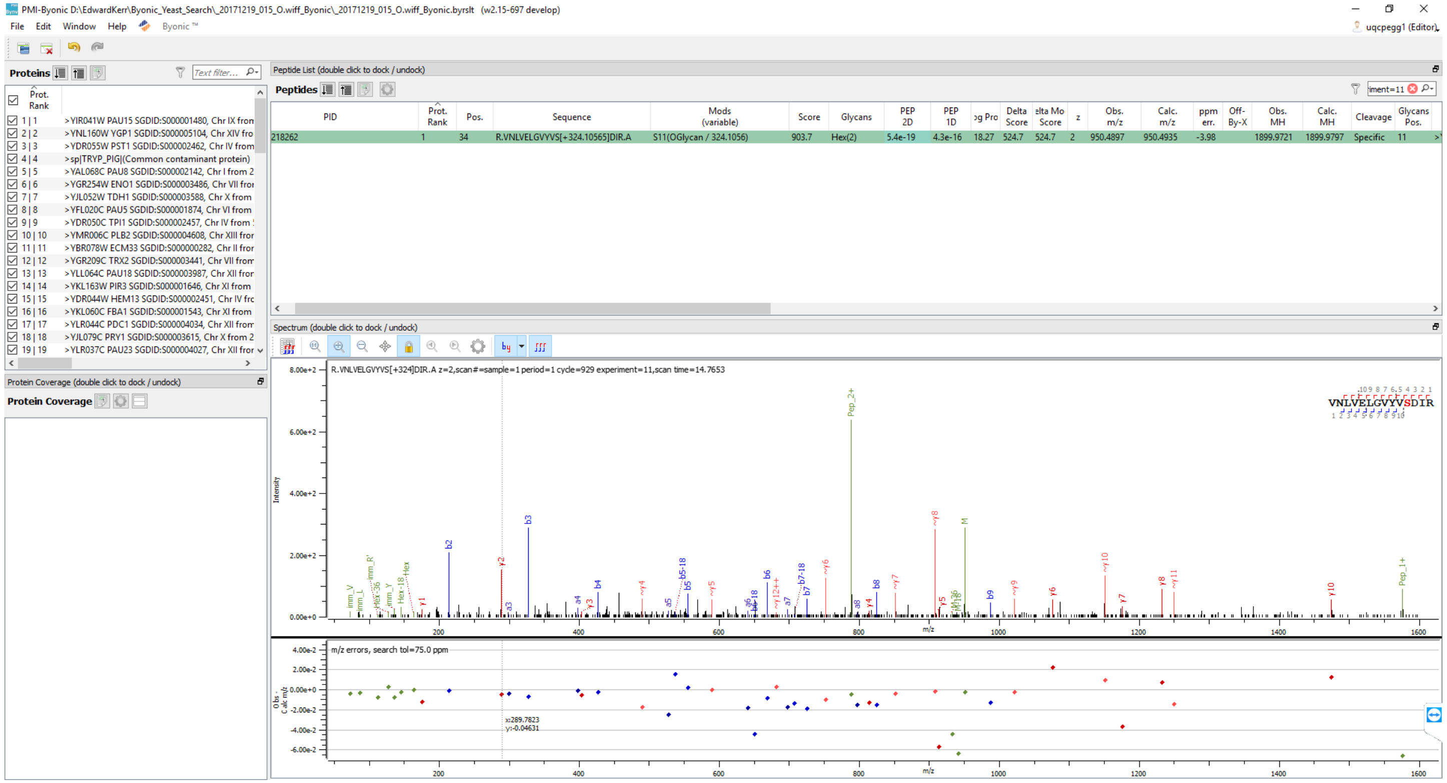
PMI-Byonic D:\EdwardKerr\Byonic_Yeast_Search_20171219_015_O.wiff_Byonic_20171219_015_O.wiff_Byonic.byrslt (w2.15-697 develop)
File Edit Window Help Byonic ™
uqcpegg1 (Editor)
Proteins
Prot. Rank
1 | 1 >YIR041W PAU15 SGDID:S000001480, Chr IX from
2 | 2 >YNL160W YGP1 SGDID:S000005104, Chr XIV fro
3 | 3 >YDR055W PST1 SGDID:S000002462, Chr IV from
4 | 4 >sp|TRYP_PIG|(Common contaminant protein)
5 | 5 >YAL068C PAU8 SGDID:S000002142, Chr I from 2
6 | 6 >YGR254W ENO1 SGDID:S000003486, Chr VII fro
7 | 7 >YJL052W TDH1 SGDID:S000003588, Chr X from
8 | 8 >YFL020C PAU5 SGDID:S000001874, Chr VI from
9 | 9 >YDR050C TPI1 SGDID:S000002457, Chr IV from
10 | 10 >YMR006C PLB2 SGDID:S000004608, Chr XIII fro
11 | 11 >YBR078W ECM33 SGDID:S000000282, Chr II fro
12 | 12 >YGR209C TRX2 SGDID:S000003441, Chr VII from
13 | 13 >YLL064C PAU18 SGDID:S000003987, Chr XII fro
14 | 14 >YKL163W PIR3 SGDID:S000001646, Chr XI from
15 | 15 >YDR044W HEM13 SGDID:S000002451, Chr IV fro
16 | 16 >YKL060C FBA1 SGDID:S000001543, Chr XI from
17 | 17 >YLR044C PDC1 SGDID:S000004034, Chr XII from
18 | 18 >YJL079C PRY1 SGDID:S000003615, Chr X from 2
19 | 19 >YLR037C PAU23 SGDID:S000004027, Chr XII fro
Protein Coverage (double click to dock / undock)
Protein Coverage
Peptide List (double click to dock / undock)
Peptides
PID | Prot. Rank | Pos. | Sequence | Mods (variable) | Score | Glycans | PEP 2D | PEP 1D | og Pro | Delta Score | elta Mo Score | z | Obs. m/z | Calc. m/z | ppm err. | Off-By-X | Obs. MH | Calc. MH | Cleavage | Glycans Pos.
218262 | 1 | 34 | R.VNLVELGVYVS[+324.10565]DIR.A | S11(OGlycan / 324.1056) | 903.7 | Hex(2) | 5.4e-19 | 4.3e-16 | 18.27 | 524.7 | 524.7 | 2 | 950.4897 | 950.4935 | -3.98 | | 1899.9721 | 1899.9797 | Specific | 11
Spectrum (double click to dock / undock)
R.VNLVELGVYVS[+324]DIR.A z=2,scan#=sample=1 period=1 cycle=929 experiment=11,scan time=14.7653
VNLVELGVYVSDIR
Intensity
m/z
m/z errors, search tol=75.0 ppm
x:289.7823
y:-0.04631

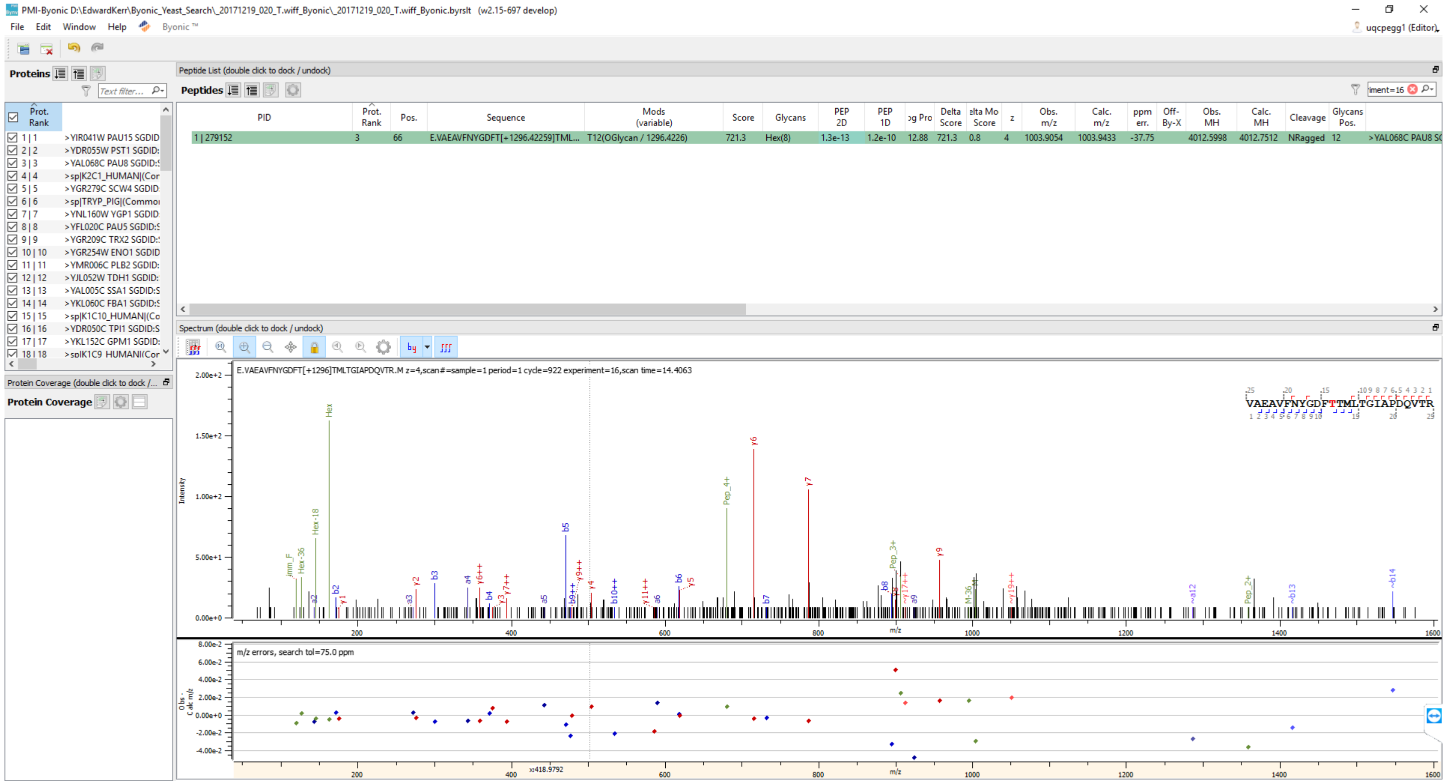
PMI-Byonic D:\EdwardKerr\Byonic_Yeast_Search_20171219_020_T.wiff_Byonic_20171219_020_T.wiff_Byonic.byrslt (w2.15-697 develop)
File Edit Window Help Byonic ™
uqcpegg1 (Editor)
Proteins
Peptide List (double click to dock / undock)
Peptides
PID
Prot. Rank
Pos.
Sequence
Mods (variable)
Score
Glycans
PEP 2D
PEP 1D
Delta Score
z
Obs. m/z
Calc. m/z
ppm err.
Off-By-X
Obs. MH
Calc. MH
Cleavage
Glycans Pos.
1 | 279152
E.VAEAVFNYGDFT[+1296.42259]TML...
T12(OGlycan / 1296.4226)
721.3
Hex(8)
1.3e-13
1.2e-10
12.88
0.8
1003.9054
1003.9433
-37.75
4012.5998
4012.7512
NRagged
>YAL068C PAU8
>YIR041W PAU15 SGDID
>YDR055W PST1 SGDID:
>YAL068C PAU8 SGDID:
>sp|K2C1_HUMAN|(Cor
>YGR279C SCW4 SGDID
>sp|TRYP_PIG|(Commo
>YNL160W YGP1 SGDID
>YFL020C PAU5 SGDID:
>YGR209C TRX2 SGDID:
>YGR254W ENO1 SGDID
>YMR006C PLB2 SGDID:
>YJL052W TDH1 SGDID:
>YAL005C SSA1 SGDID:
>YKL060C FBA1 SGDID:
>sp|K1C10_HUMAN|(Co
>YDR050C TPI1 SGDID:
>YKL152C GPM1 SGDID:
>sp|K1C9_HUMAN|(Cor
Protein Coverage (double click to dock / ...
Protein Coverage
Spectrum (double click to dock / undock)
E.VAEAVFNYGDFT[+1296]TMLTGIAPDQVTR.M z=4,scan#=sample=1 period=1 cycle=922 experiment=16,scan time=14.4063
VAEAVFNYGDFTTMLTGIAPDQVTR
Intensity
m/z
m/z errors, search tol=75.0 ppm
x:418.9792

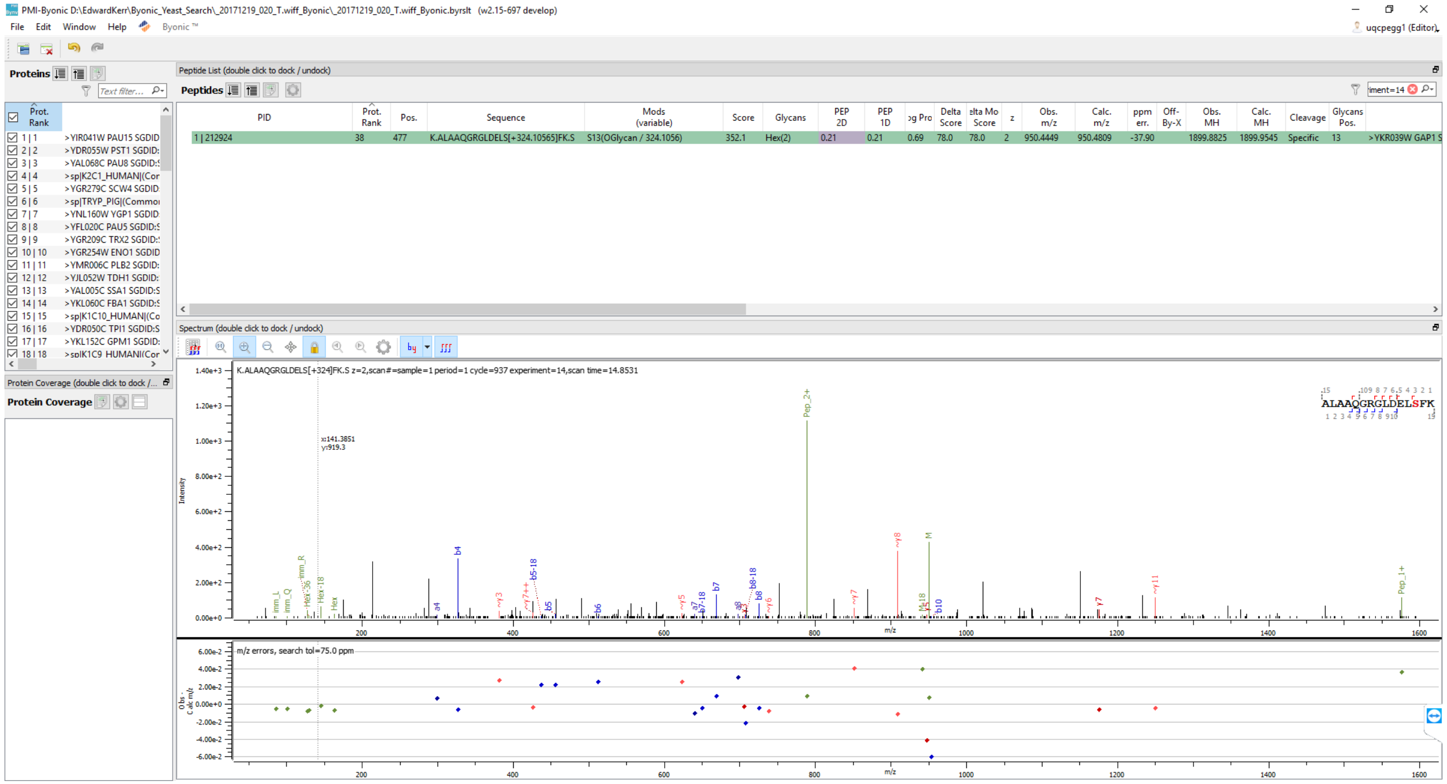
PMI-Byonic D:\EdwardKerr\Byonic_Yeast_Search_20171219_020_T.wiff_Byonic_20171219_020_T.wiff_Byonic.byrslt (w2.15-697 develop)
File Edit Window Help Byonic ™
Proteins
Peptide List (double click to dock / undock)
Peptides
PID
Prot. Rank
Pos.
Sequence
Mods (variable)
Score
Glycans
PEP 2D
PEP 1D
Delta Score
z
Obs. m/z
Calc. m/z
ppm err.
Off-By-X
Obs. MH
Calc. MH
Cleavage
Glycans Pos.
1 | 212924 38 477 K.ALAAQGRGLDELS[+324.10565]FK.S S13(OGlycan / 324.1056) 352.1 Hex(2) 0.21 0.21 0.69 78.0 78.0 2 950.4449 950.4809 -37.90 1899.8825 1899.9545 Specific 13 >YKR039W GAP1 S
Spectrum (double click to dock / undock)
K.ALAAQGRGLDELS[+324]FK.S z=2,scan#=sample=1 period=1 cycle=937 experiment=14,scan time=14.8531
Protein Coverage (double click to dock / ...)
Protein Coverage
Intensity
m/z
m/z errors, search tol=75.0 ppm
ALAAQGRGLDELSFK

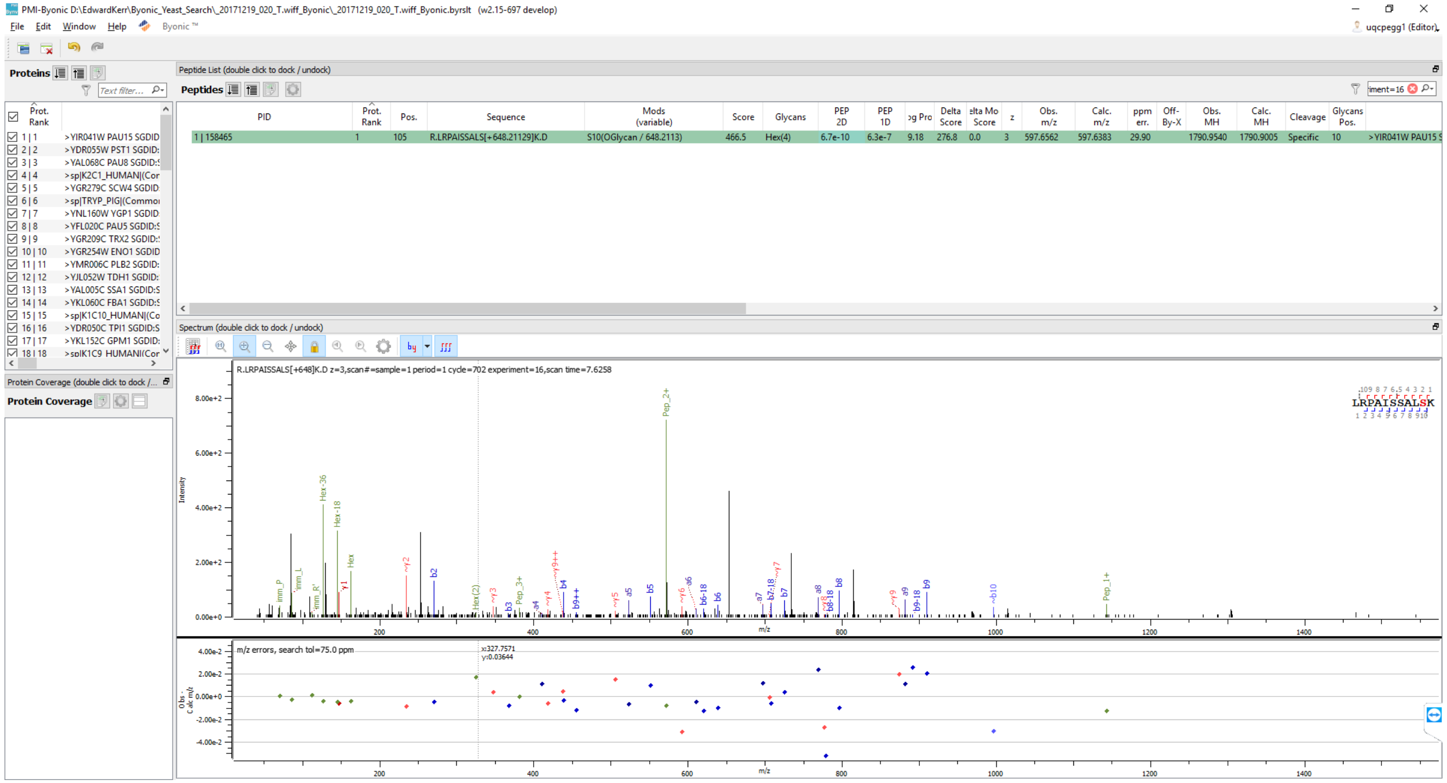
PMI-Byonic D:\EdwardKerr\Byonic_Yeast_Search_20171219_020_T.wiff_Byonic_20171219_020_T.wiff_Byonic.byrslt (w2.15-697 develop)
File Edit Window Help Byonic ™
uqcpegg1 (Editor)
Proteins
Peptide List (double click to dock / undock)
Peptides
Prot. Rank
1 | 1 >YIR041W PAU15 SGDID
2 | 2 >YDR055W PST1 SGDID:
3 | 3 >YAL068C PAU8 SGDID:
4 | 4 >sp|K2C1_HUMAN|(Cor
5 | 5 >YGR279C SCW4 SGDID
6 | 6 >sp|TRYP_PIG|(Commo
7 | 7 >YNL160W YGP1 SGDID
8 | 8 >YFL020C PAU5 SGDID:
9 | 9 >YGR209C TRX2 SGDID:
10 | 10 >YGR254W ENO1 SGDID
11 | 11 >YMR006C PLB2 SGDID:
12 | 12 >YJL052W TDH1 SGDID:
13 | 13 >YAL005C SSA1 SGDID:
14 | 14 >YKL060C FBA1 SGDID:
15 | 15 >sp|K1C10_HUMAN|(Co
16 | 16 >YDR050C TPI1 SGDID:
17 | 17 >YKL152C GPM1 SGDID:
18 | 18 >sp|K1C9_HUMAN|(Cor
PID | Prot. Rank | Pos. | Sequence | Mods (variable) | Score | Glycans | PEP 2D | PEP 1D | og Pro | Delta Score | elta Mo Score | z | Obs. m/z | Calc. m/z | ppm err. | Off-By-X | Obs. MH | Calc. MH | Cleavage | Glycans Pos.
1 | 158465 | 1 | 105 | R.LRPAISSALS[+648.21129]K.D | S10(OGlycan / 648.2113) | 466.5 | Hex(4) | 6.7e-10 | 6.3e-7 | 9.18 | 276.8 | 0.0 | 3 | 597.6562 | 597.6383 | 29.90 | | 1790.9540 | 1790.9005 | Specific | 10 | >YIR041W PAU15 S
Spectrum (double click to dock / undock)
R.LRPAISSALS[+648]K.D z=3,scan#=sample=1 period=1 cycle=702 experiment=16,scan time=7.6258
Protein Coverage (double click to dock / ...)
Protein Coverage
Intensity
m/z
m/z errors, search tol=75.0 ppm
LRPAISSALSK

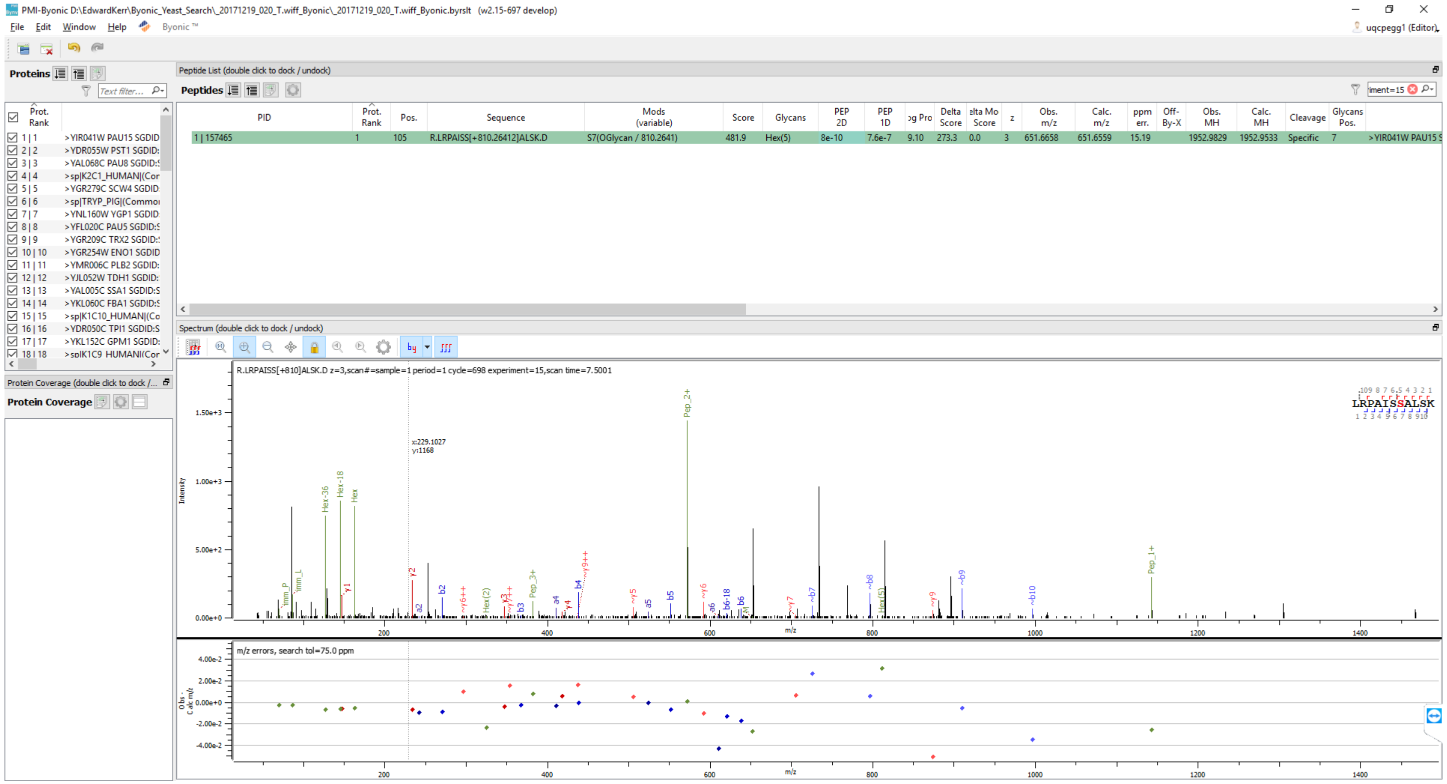
PMI-Byonic D:\EdwardKerr\Byonic_Yeast_Search_20171219_020_T.wiff_Byonic_20171219_020_T.wiff_Byonic.byrslt (w2.15-697 develop)
File Edit Window Help Byonic ™
uqcpegg1 (Editor)
Proteins
Peptide List (double click to dock / undock)
Peptides
PID
Prot. Rank
Pos.
Sequence
Mods (variable)
Score
Glycans
PEP 2D
PEP 1D
Delta Score
z
Obs. m/z
Calc. m/z
ppm err.
Off-By-X
Obs. MH
Calc. MH
Cleavage
Glycans Pos.
1 | 157465
105
R.LRPAISS[+810.26412]ALSK.D
S7(OGlycan / 810.2641)
481.9
Hex(5)
8e-10
7.6e-7
9.10
273.3
0.0
3
651.6658
651.6559
15.19
1952.9829
1952.9533
Specific
7
>YIR041W PAU15
>YDR055W PST1
>YAL068C PAU8
>YGR279C SCW4
>YNL160W YGP1
>YFL020C PAU5
>YGR209C TRX2
>YGR254W ENO1
>YMR006C PLB2
>YJL052W TDH1
>YAL005C SSA1
>YKL060C FBA1
>YDR050C TPI1
>YKL152C GPM1
Protein Coverage (double click to dock / undock)
Protein Coverage
Spectrum (double click to dock / undock)
R.LRPAISS[+810]ALSK.D z=3,scan#=sample=1 period=1 cycle=698 experiment=15,scan time=7.5001
LRPAISSALSK
Intensity
Pep_2+
Pep_1+
Hex-36
Hex-18
Hex
m/z
m/z errors, search tol=75.0 ppm

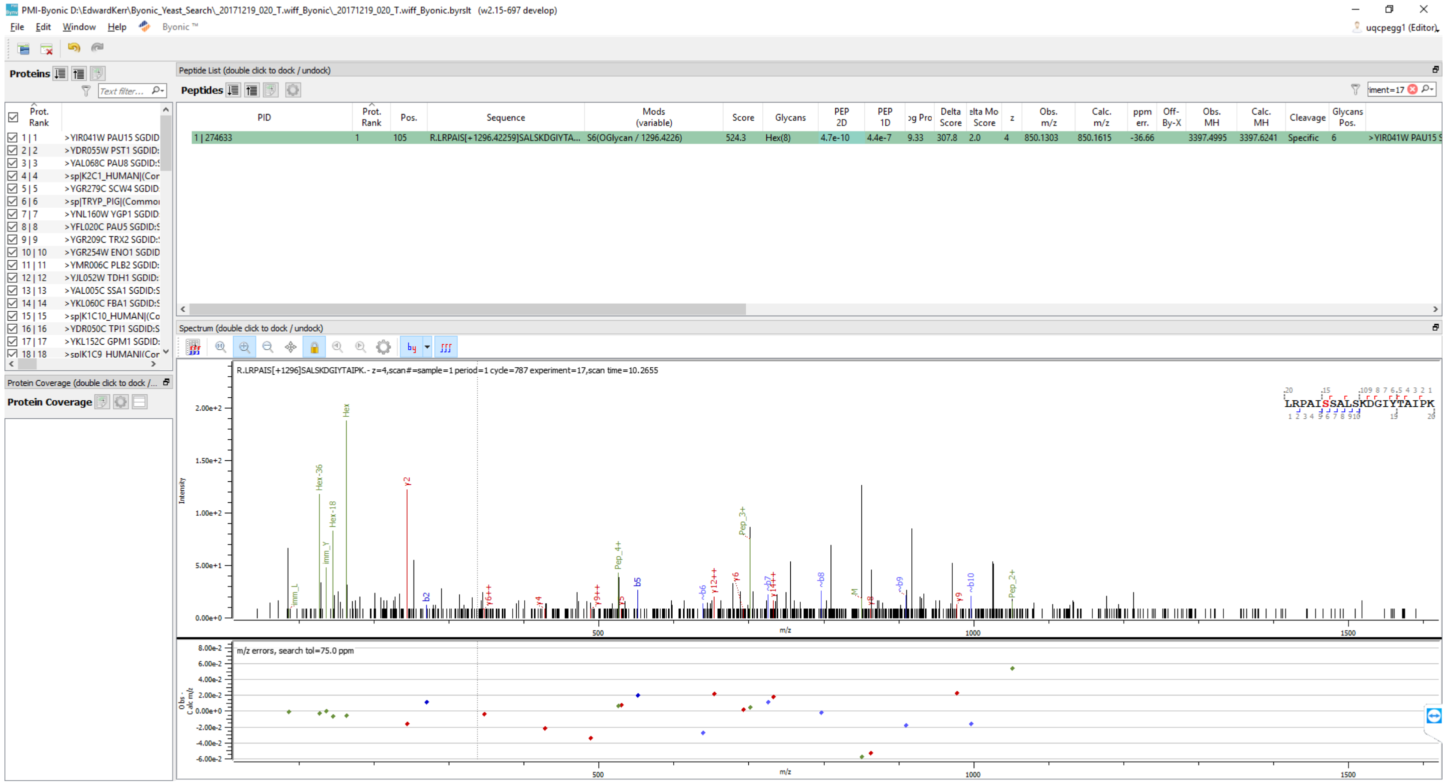

PMI-Byonic D:\EdwardKerr\Byonic_Yeast_Search_20171219_020_T.wiff_Byonic_20171219_020_T.wiff_Byonic.byrslt (w2.15-697 develop)
File Edit Window Help Byonic ™
uqcpegg1 (Editor)
Proteins
Peptide List (double click to dock / undock)
Peptides
PID
Prot. Rank
Pos.
Sequence
Mods (variable)
Score
Glycans
PEP 2D
PEP 1D
Delta Score
z
Obs. m/z
Calc. m/z
ppm err.
Off-By-X
Obs. MH
Calc. MH
Cleavage
Glycans Pos.
1 | 274633
1
105
R.LRPAIS[+1296.42259]SALSKDGIYTA...
S6(OGlycan / 1296.4226)
524.3
Hex(8)
4.7e-10
4.4e-7
9.33
307.8
2.0
4
850.1303
850.1615
-36.66
3397.4995
3397.6241
Specific
6
>YIR041W PAU15
Spectrum (double click to dock / undock)
R.LRPAIS[+1296]SALSKDGIYTAIPK.- z=4,scan#=sample=1 period=1 cycle=787 experiment=17,scan time=10.2655
LRPAISSALSKDGIYTAIPK
Intensity
m/z
m/z errors, search tol=75.0 ppm
Protein Coverage (double click to dock / undock)
Protein Coverage

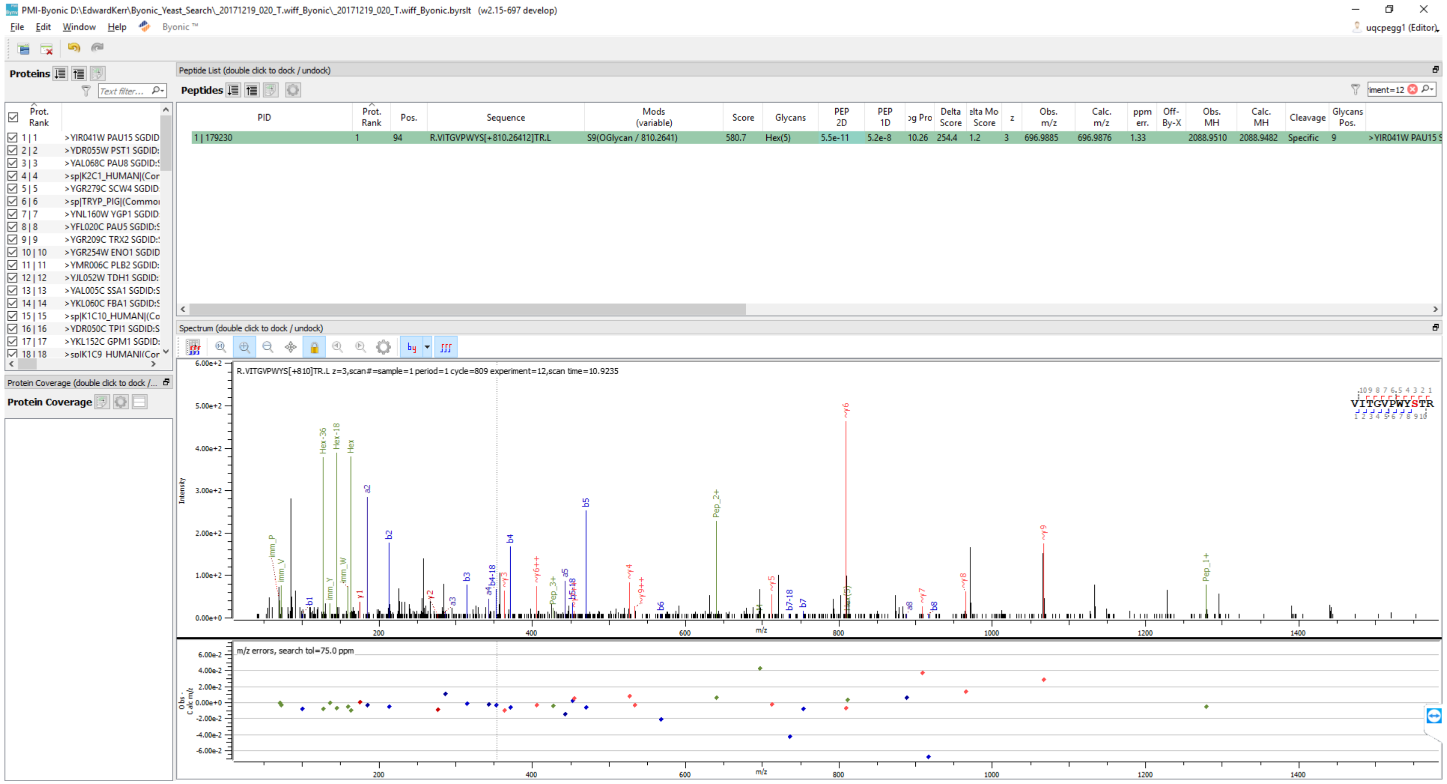
PMI-Byonic D:\EdwardKerr\Byonic_Yeast_Search_20171219_020_T.wiff_Byonic_20171219_020_T.wiff_Byonic.byrslt (v2.15-697 develop)
File Edit Window Help Byonic™
Proteins
Peptide List (double click to dock / undock)
Peptides
PID | Prot. Rank | Pos. | Sequence | Mods (variable) | Score | Glycans | PEP 2D | PEP 1D | Delta Score | z | Obs. m/z | Calc. m/z | ppm err. | Obs. MH | Calc. MH | Cleavage | Glycans Pos.
1 | 179230 | 1 | 94 | R.VITGVPWYS[+810.26412]TR.L | S9(OGlycan / 810.2641) | 580.7 | Hex(5) | 5.5e-11 | 5.2e-8 | 10.26 | 254.4 | 1.2 | 3 | 696.9885 | 696.9876 | 1.33 | 2088.9510 | 2088.9482 | Specific | 9 | >YIR041W PAU15
Spectrum (double click to dock / undock)
R.VITGVPWYS[+810]TR.L z=3,scan#=sample=1 period=1 cycle=809 experiment=12,scan time=10.9235
VITGVPWYSTR
m/z errors, search tol=75.0 ppm
Protein Coverage

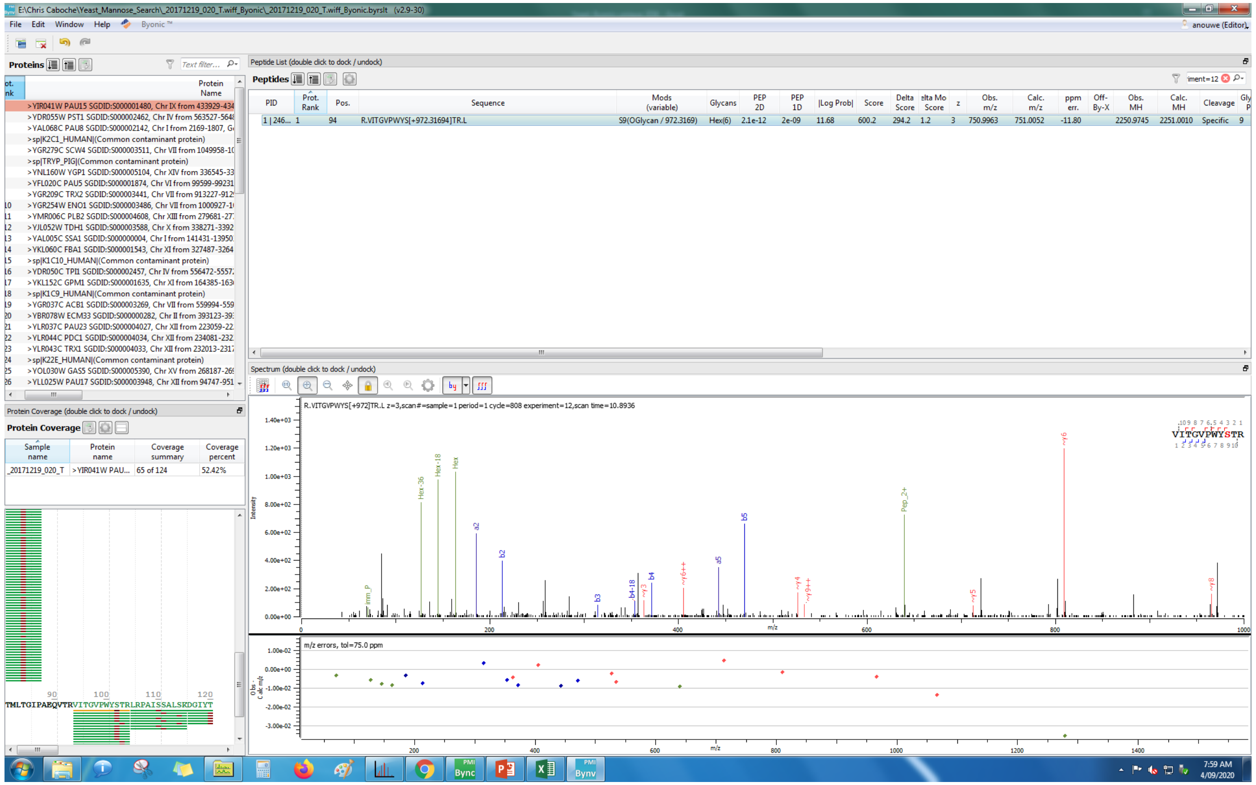
E:\Chris Caboche\Yeast_Mannose_Search_20171219_020_T.wiff_Byonic_20171219_020_T.wiff_Byonic.byrslt (v2.9-30)
File Edit Window Help Byonic™
Proteins
Peptide List (double click to dock / undock)
Peptides
PID
Prot. Rank
Pos.
Sequence
Mods (variable)
Glycans
PEP 2D
PEP 1D
|Log Prob|
Score
Delta Score
Obs. m/z
Calc. m/z
ppm err.
Obs. MH
Calc. MH
Cleavage
R.VITGVPWYS[+972.31694]TR.L
S9(OGlycan / 972.3169)
Hex(6)
Specific
Spectrum (double click to dock / undock)
R.VITGVPWYS[+972]TR.L z=3,scan#=sample=1 period=1 cycle=808 experiment=12,scan time=10.8936
VITGVPWYSTR
Intensity
m/z
m/z errors, tol=75.0 ppm
Protein Coverage (double click to dock / undock)
Protein Coverage
Sample name
Protein name
Coverage summary
Coverage percent
_20171219_020_T
>YIR041W PAU...
65 of 124
52.42%
TMLTGIPAEQVTRVITGVPWYSTRLRPAISSALSKDGIYT

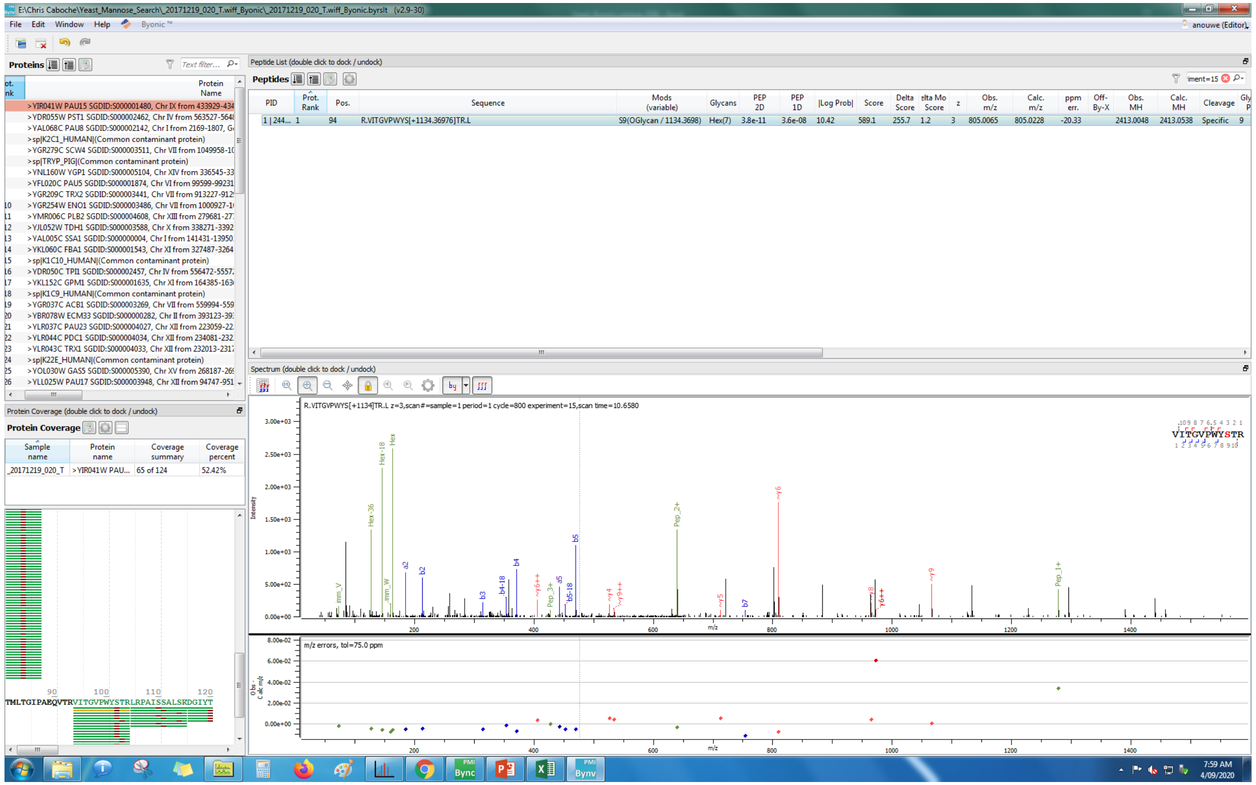

E:\Chris Caboche\Yeast_Mannose_Search_20171219_020_T.wiff_Byonic_20171219_020_T.wiff_Byonic.byrslt (v2.9-30)
File Edit Window Help Byonic™
Proteins
Peptide List (double click to dock / undock)
Peptides
PID | Prot. Rank | Pos. | Sequence | Mods (variable) | Glycans | PEP 2D | PEP 1D | |Log Prob| | Score | Delta Score | Delta Mod Score | z | Obs. m/z | Calc. m/z | ppm err. | Off-By-X | Obs. MH | Calc. MH | Cleavage
1 | 244... | 1 | 94 | R.VITGVPWYS[+1134.36976]TR.L | S9(OGlycan / 1134.3698) | Hex(7) | 3.8e-11 | 3.6e-08 | 10.42 | 589.1 | 255.7 | 1.2 | 3 | 805.0065 | 805.0228 | -20.33 | | 2413.0048 | 2413.0538 | Specific
Spectrum (double click to dock / undock)
R.VITGVPWYS[+1134]TR.L z=3,scan#=sample=1 period=1 cycle=800 experiment=15,scan time=10.6580
VITGVPWYSTR
m/z errors, tol=75.0 ppm
Protein Coverage (double click to dock / undock)
Protein Coverage
Sample name | Protein name | Coverage summary | Coverage percent
_20171219_020_T | >YIR041W PAU... | 65 of 124 | 52.42%
TMLTGIPAEQVTRVITGVPWYSTRLRPAISSALSKDGIYT
7:59 AM
4/09/2020

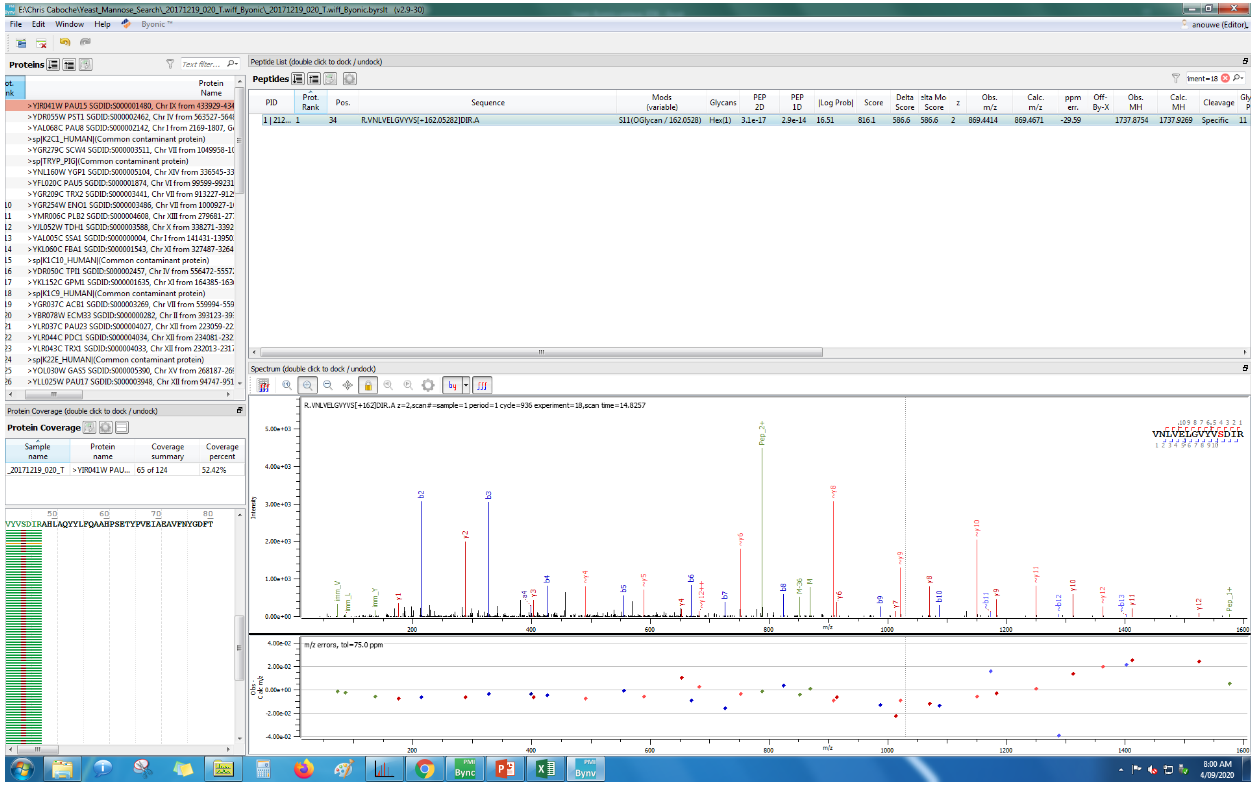
E:\Chris Caboche\Yeast_Mannose_Search_20171219_020_T.wiff_Byonic_20171219_020_T.wiff_Byonic.byrslt (v2.9-30)
Proteins
Peptide List (double click to dock / undock)
Peptides
R.VNLVELGVYVS[+162.05282]DIR.A
S11(OGlycan / 162.0528)
Protein Coverage (double click to dock / undock)
Protein Coverage
Spectrum (double click to dock / undock)
R.VNLVELGVYVS[+162]DIR.A z=2,scan#=sample=1 period=1 cycle=936 experiment=18,scan time=14.8257
VNLVELGVYVSDIR
m/z errors, tol=75.0 ppm

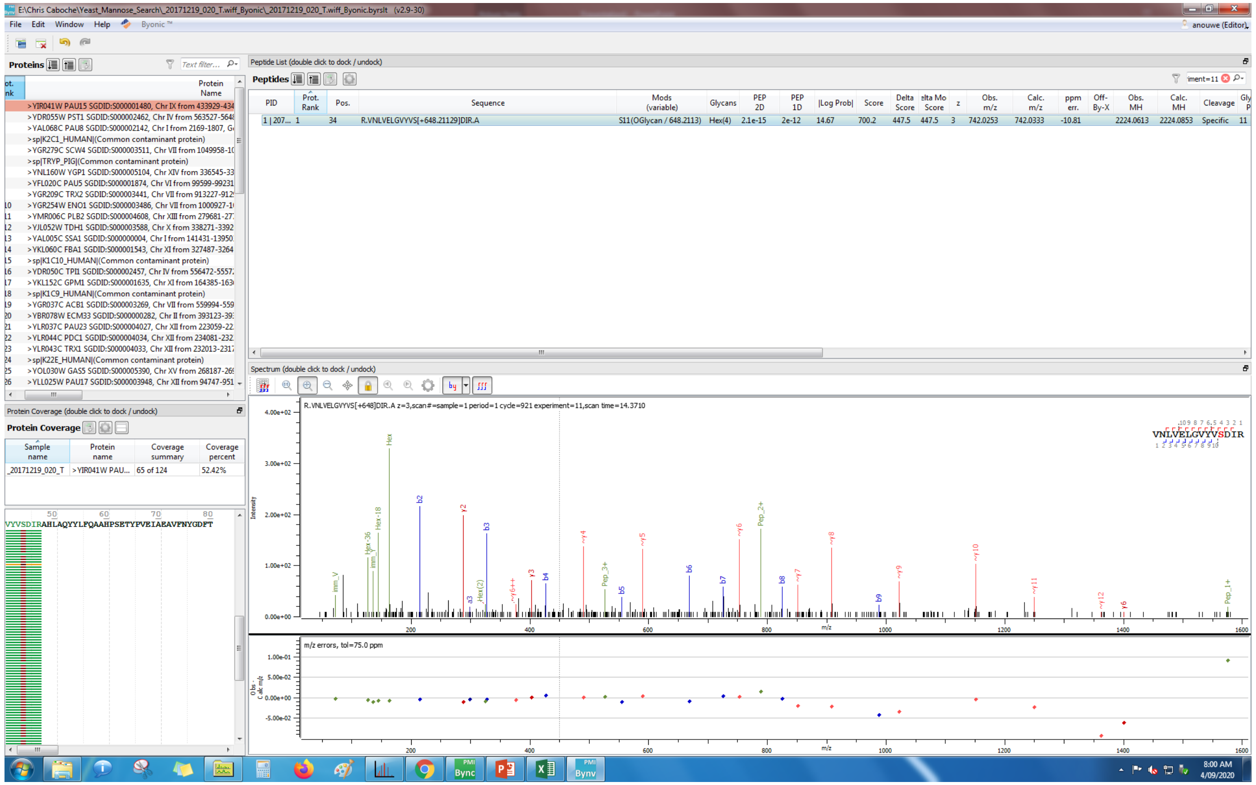
E:\Chris Caboche\Yeast_Mannose_Search_20171219_020_T.wiff_Byonic_20171219_020_T.wiff_Byonic.byrslt (v2.9-30)
File Edit Window Help Byonic™
anouwe (Editor)
Proteins
Peptide List (double click to dock / undock)
Peptides
PID | Prot. Rank | Pos. | Sequence | Mods (variable) | Glycans | PEP 2D | PEP 1D | |Log Prob| | Score | Delta Score | Delta Mod Score | z | Obs. m/z | Calc. m/z | ppm err. | Off-By-X | Obs. MH | Calc. MH | Cleavage
1 | 207... 1 34 R.VNLVELGVYVS[+648.21129]DIR.A S11(OGlycan / 648.2113) Hex(4) 2.1e-15 2e-12 14.67 700.2 447.5 447.5 3 742.0253 742.0333 -10.81 2224.0613 2224.0853 Specific 11
>YIR041W PAU15 SGDID:S000001480, Chr IX from 433929-434
>YDR055W PST1 SGDID:S000002462, Chr IV from 563527-564
>YAL068C PAU8 SGDID:S000002142, Chr I from 2169-1807, G
>sp|K2C1_HUMAN|(Common contaminant protein)
>YGR279C SCW4 SGDID:S000003511, Chr VII from 1049958-1
>sp|TRYP_PIG|(Common contaminant protein)
>YNL160W YGP1 SGDID:S000005104, Chr XIV from 336545-33
>YFL020C PAU5 SGDID:S000001874, Chr VI from 99599-99231
>YGR209C TRX2 SGDID:S000003441, Chr VII from 913227-912
>YGR254W ENO1 SGDID:S000003486, Chr VII from 1000927-1
>YMR006C PLB2 SGDID:S000004608, Chr XIII from 279681-27
Protein Coverage (double click to dock / undock)
Protein Coverage
Sample name | Protein name | Coverage summary | Coverage percent
_20171219_020_T | >YIR041W PAU... | 65 of 124 | 52.42%
VYVSDIRAHLAQYYLFQAAHPSETYPVEIAEAVFNYGDFT
Spectrum (double click to dock / undock)
R.VNLVELGVYVS[+648]DIR.A z=3,scan#=sample=1 period=1 cycle=921 experiment=11,scan time=14.3710
VNLVELGVYVSDIR
Intensity
m/z
m/z errors, tol=75.0 ppm
8:00 AM
4/09/2020

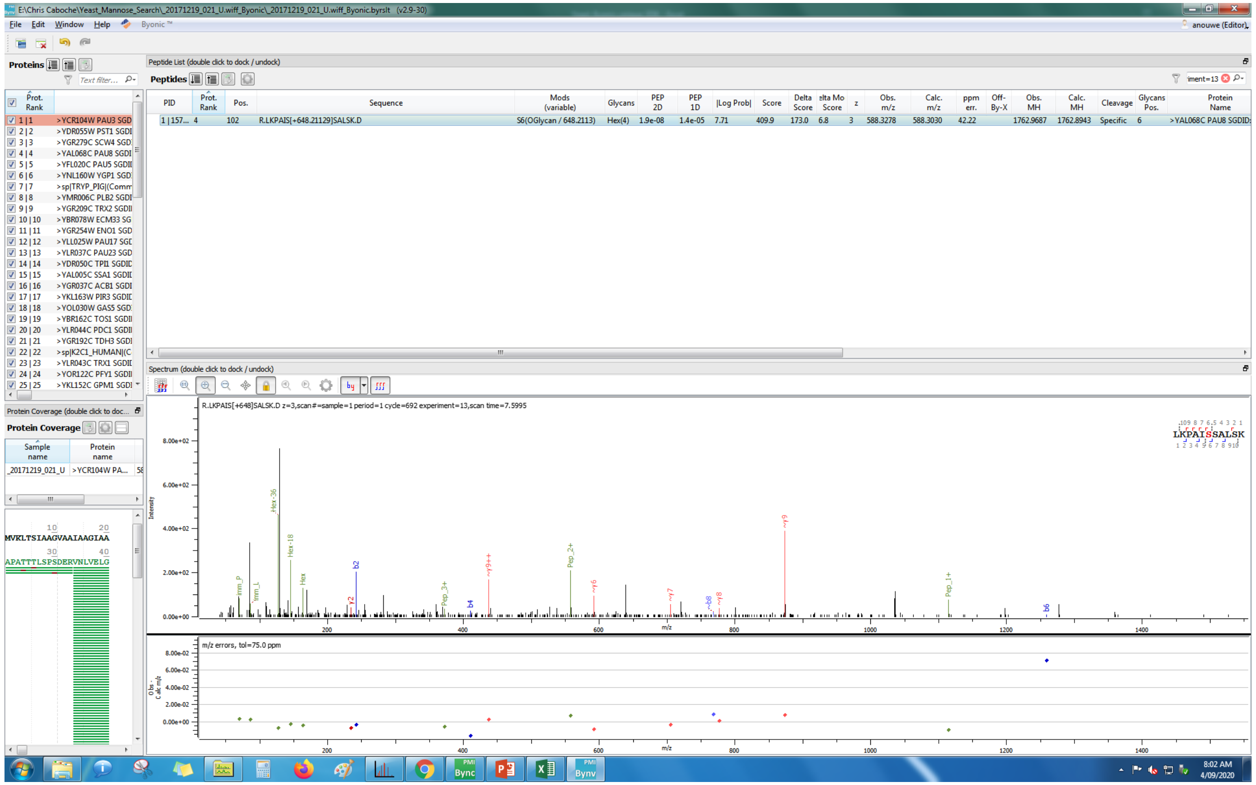

E:\Chris Caboche\Yeast_Mannose_Search_20171219_021_U.wiff_Byonic_20171219_021_U.wiff_Byonic.byrslt (v2.9-30)
File Edit Window Help Byonic™
anouwe (Editor)
Proteins
Peptide List (double click to dock / undock)
Peptides
PID | Prot. Rank | Pos. | Sequence | Mods (variable) | Glycans | PEP 2D | PEP 1D | |Log Prob| | Score | Delta Score | elta Mo Score | z | Obs. m/z | Calc. m/z | ppm err. | Off-By-X | Obs. MH | Calc. MH | Cleavage | Glycans Pos. | Protein Name
1 | 157... 4 102 R.LKPAIS[+648.21129]SALSK.D S6(OGlycan / 648.2113) Hex(4) 1.9e-08 1.4e-05 7.71 409.9 173.0 6.8 3 588.3278 588.3030 42.22 1762.9687 1762.8943 Specific 6 >YAL068C PAU8 SGDID:
Spectrum (double click to dock / undock)
R.LKPAIS[+648]SALSK.D z=3,scan#=sample=1 period=1 cycle=692 experiment=13,scan time=7.5995
LKPAISSALSK
m/z errors, tol=75.0 ppm
Protein Coverage (double click to doc...
Protein Coverage
Sample name
Protein name
MVKLTSIAAGVAAIAAGIAA
APATTTLSPSDERVNLVELG

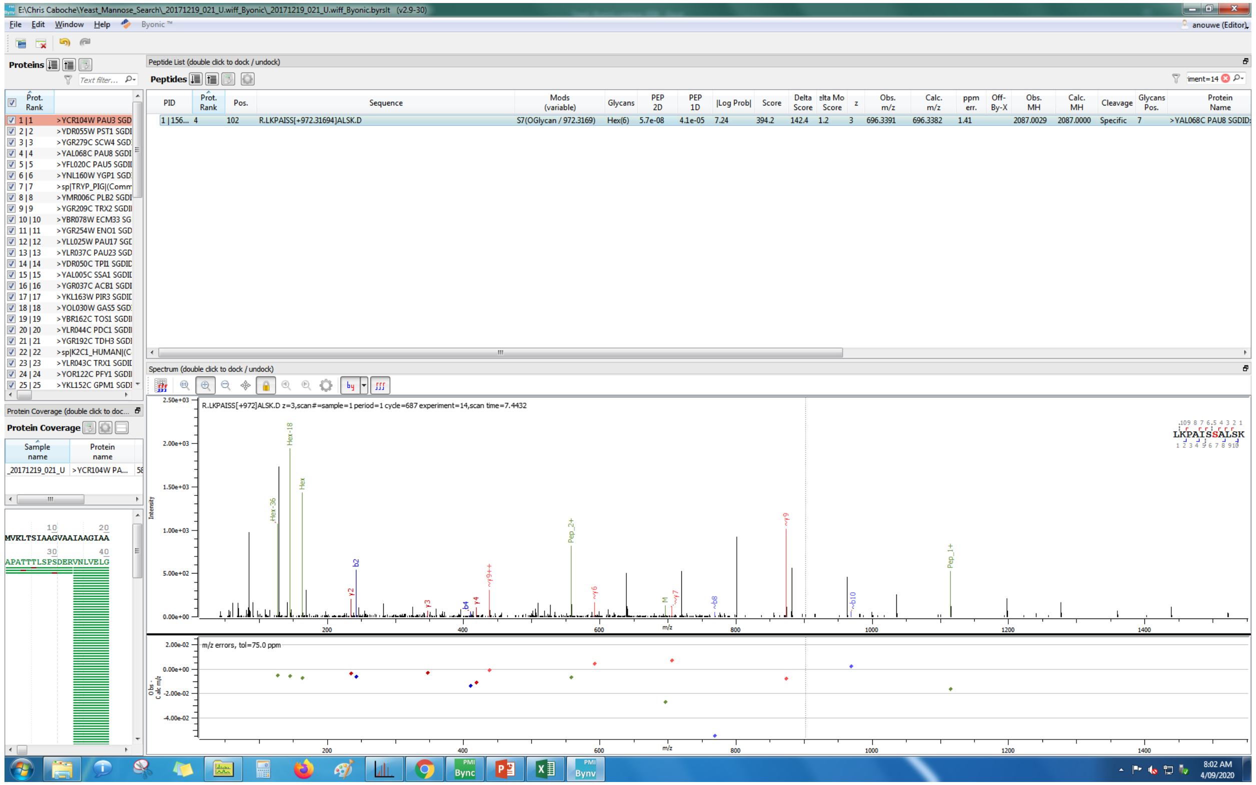
E:\Chris Caboche\Yeast_Mannose_Search_20171219_021_U.wiff_Byonic_20171219_021_U.wiff_Byonic.byrslt (v2.9-30)
File Edit Window Help Byonic™
Proteins
Peptide List (double click to dock / undock)
Peptides
PID Prot. Rank Pos. Sequence Mods (variable) Glycans PEP 2D PEP 1D |Log Prob| Score Delta Score Delta Mod Score z Obs. m/z Calc. m/z ppm err. Off-By-X Obs. MH Calc. MH Cleavage Glycans Pos. Protein Name
1 | 156... 4 102 R.LKPAISS[+972.31694]ALSK.D S7(OGlycan / 972.3169) Hex(6) 5.7e-08 4.1e-05 7.24 394.2 142.4 1.2 3 696.3391 696.3382 1.41 2087.0029 2087.0000 Specific 7 >YAL068C PAU8 SGDID:
Spectrum (double click to dock / undock)
R.LKPAISS[+972]ALSK.D z=3,scan#=sample=1 period=1 cycle=687 experiment=14,scan time=7.4432
LKPAISSALSK
Hex-18
Hex
Hex-36
Pep_2+
Pep_1+
m/z errors, tol=75.0 ppm
Protein Coverage (double click to dock / undock)
Protein Coverage
Sample name
Protein name
_20171219_021_U
>YCR104W PA...
MVKLTSIAAGVAAIAAGIAA
APATTTLSPSDERVNLVELG
8:02 AM
4/09/2020

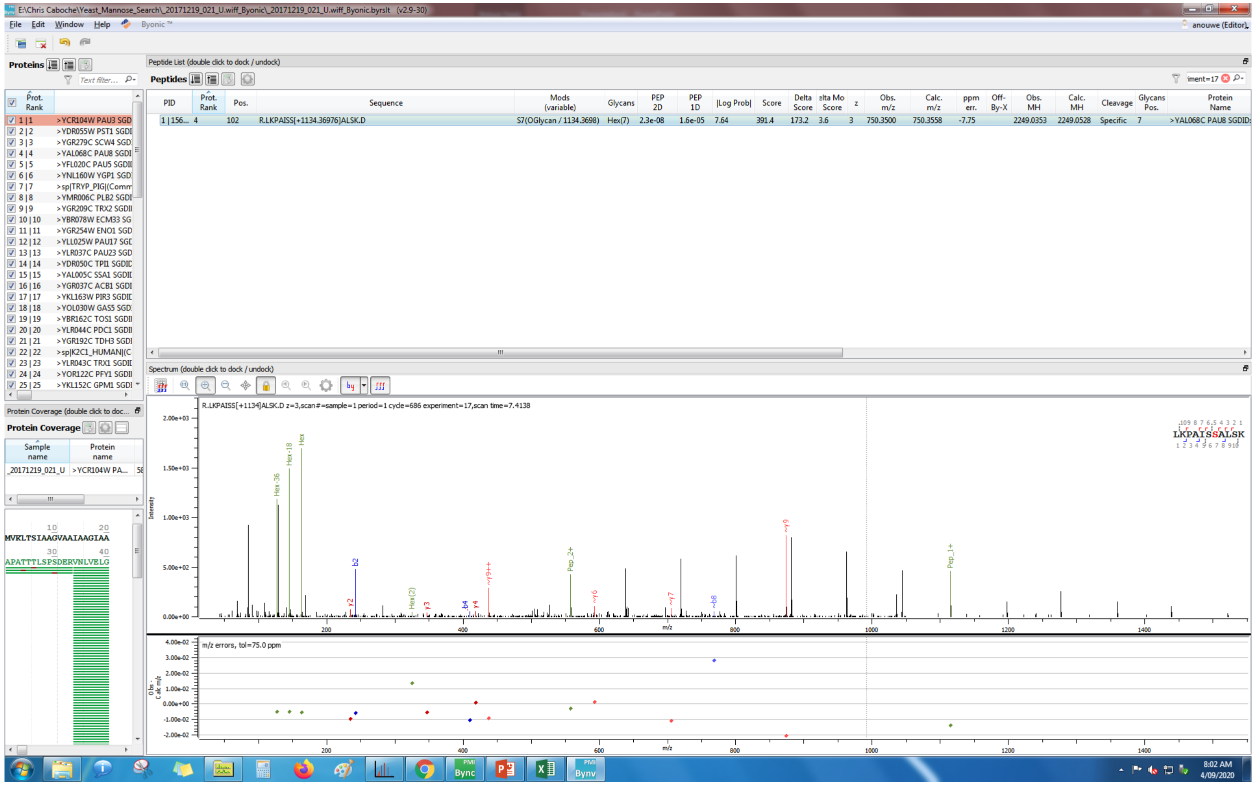
E:\Chris Caboche\Yeast_Mannose_Search_20171219_021_U.wiff_Byonic_20171219_021_U.wiff_Byonic.byrslt (v2.9-30)
File Edit Window Help Byonic ™
anouwe (Editor)
Proteins
Peptide List (double click to dock / undock)
Peptides
PID | Prot. Rank | Pos. | Sequence | Mods (variable) | Glycans | PEP 2D | PEP 1D | |Log Prob| | Score | Delta Score | elta Mo Score | z | Obs. m/z | Calc. m/z | ppm err. | Off-By-X | Obs. MH | Calc. MH | Cleavage | Glycans Pos. | Protein Name
1 | 156... | 4 | 102 | R.LKPAISS[+1134.36976]ALSK.D | S7(OGlycan / 1134.3698) | Hex(7) | 2.3e-08 | 1.6e-05 | 7.64 | 391.4 | 173.2 | 3.6 | 3 | 750.3500 | 750.3558 | -7.75 | | 2249.0353 | 2249.0528 | Specific | 7 | >YAL068C PAU8 SGDID:
1 | 1 >YCR104W PAU3 SGD
2 | 2 >YDR055W PST1 SGDI
3 | 3 >YGR279C SCW4 SGD
4 | 4 >YAL068C PAU8 SGDI
5 | 5 >YFL020C PAU5 SGDI
6 | 6 >YNL160W YGP1 SGD
7 | 7 >sp|TRYP_PIG|(Comm
8 | 8 >YMR006C PLB2 SGDI
Spectrum (double click to dock / undock)
R.LKPAISS[+1134]ALSK.D z=3,scan#=sample=1 period=1 cycle=686 experiment=17,scan time=7.4138
LKPAISSALSK
m/z errors, tol=75.0 ppm
Protein Coverage
MVKLTSIAAGVAAIAAGIAA
APATTTLSPSDERVNLVELG

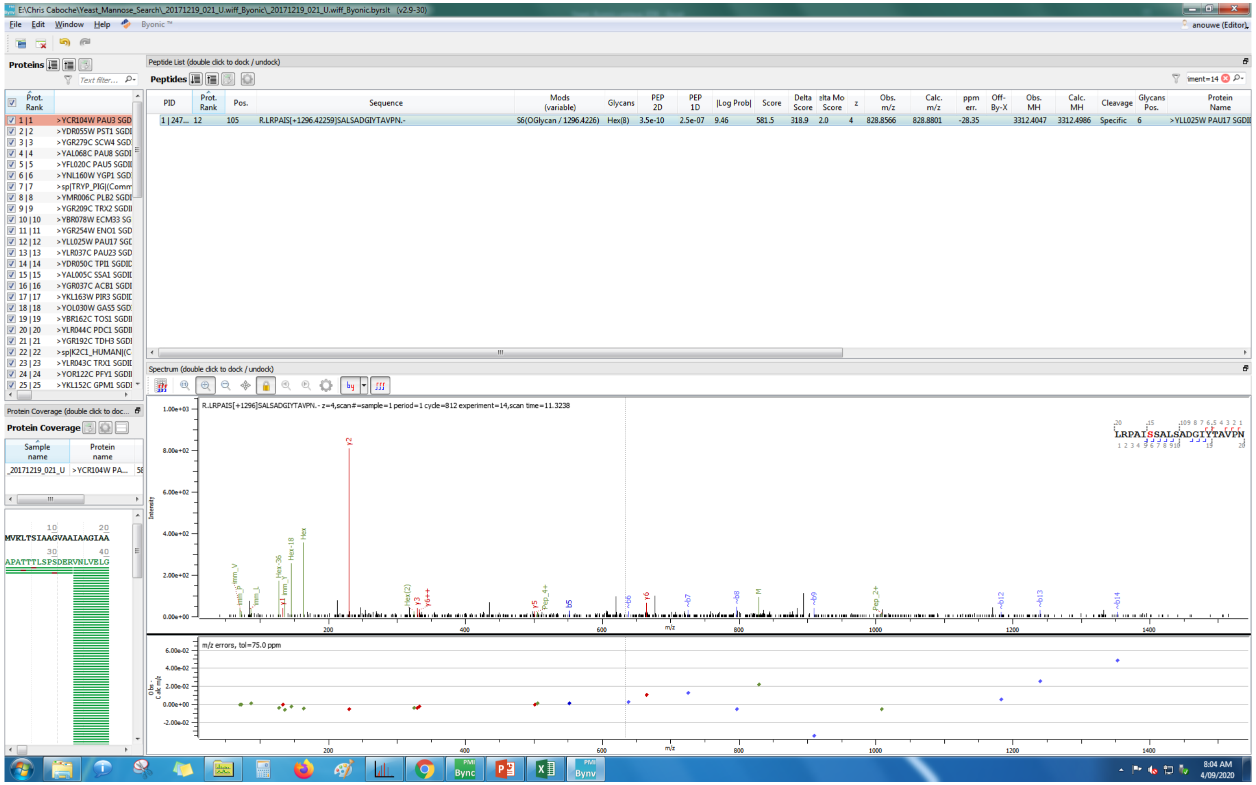
E:\Chris Caboche\Yeast_Mannose_Search_20171219_021_U.wiff_Byonic_20171219_021_U.wiff_Byonic.byrslt (v2.9-30)
File Edit Window Help Byonic
Proteins
Peptide List (double click to dock / undock)
Peptides
PID
Prot. Rank
Pos.
Sequence
Mods (variable)
Glycans
PEP 2D
PEP 1D
|Log Prob|
Score
Delta Score
z
Obs. m/z
Calc. m/z
ppm err.
Off-By-X
Obs. MH
Calc. MH
Cleavage
Glycans Pos.
Protein Name
1 | 247... 12 105 R.LRPAIS[+1296.42259]SALSADGIYTAVPN.- S6(OGlycan / 1296.4226) Hex(8) 3.5e-10 2.5e-07 9.46 581.5 318.9 2.0 4 828.8566 828.8801 -28.35 3312.4047 3312.4986 Specific 6 >YLL025W PAU17 SGDI
Spectrum (double click to dock / undock)
R.LRPAIS[+1296]SALSADGIYTAVPN.- z=4,scan#=sample=1 period=1 cycle=812 experiment=14,scan time=11.3238
LRPAISSALSADGIYTAVPN
m/z errors, tol=75.0 ppm
Protein Coverage (double click to doc...
Protein Coverage
Sample name
Protein name
_20171219_021_U >YCR104W PA...
MVKLTSIAAGVAAIAAGIAA
APATTTLSPSDERVNLVELG
8:04 AM
4/09/2020

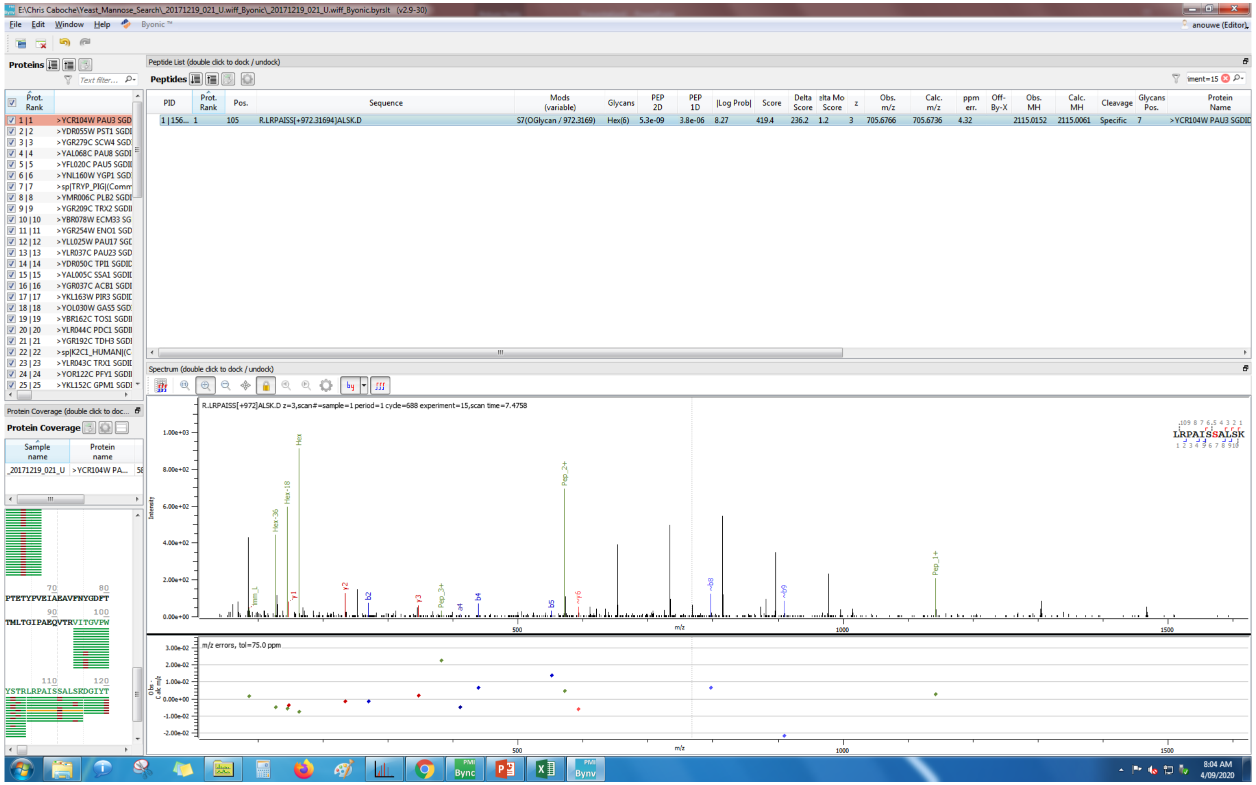
E:\Chris Caboche\Yeast_Mannose_Search_20171219_021_U.wiff_Byonic_20171219_021_U.wiff_Byonic.byrslt (v2.9-30)
Peptide List (double click to dock / undock)
PID | Prot. Rank | Pos. | Sequence | Mods (variable) | Glycans | PEP 2D | PEP 1D | |Log Prob| | Score | Delta Score | Delta Mod Score | z | Obs. m/z | Calc. m/z | ppm err. | Off-By-X | Obs. MH | Calc. MH | Cleavage | Glycans Pos. | Protein Name
1 | 156... 1 105 R.LRPAISS[+972.31694]ALSK.D S7(OGlycan / 972.3169) Hex(6) 5.3e-09 3.8e-06 8.27 419.4 236.2 1.2 3 705.6766 705.6736 4.32 2115.0152 2115.0061 Specific 7 >YCR104W PAU3 SGDID
R.LRPAISS[+972]ALSK.D z=3,scan#=sample=1 period=1 cycle=688 experiment=15,scan time=7.4758
LRPAISSALSK
m/z errors, tol=75.0 ppm

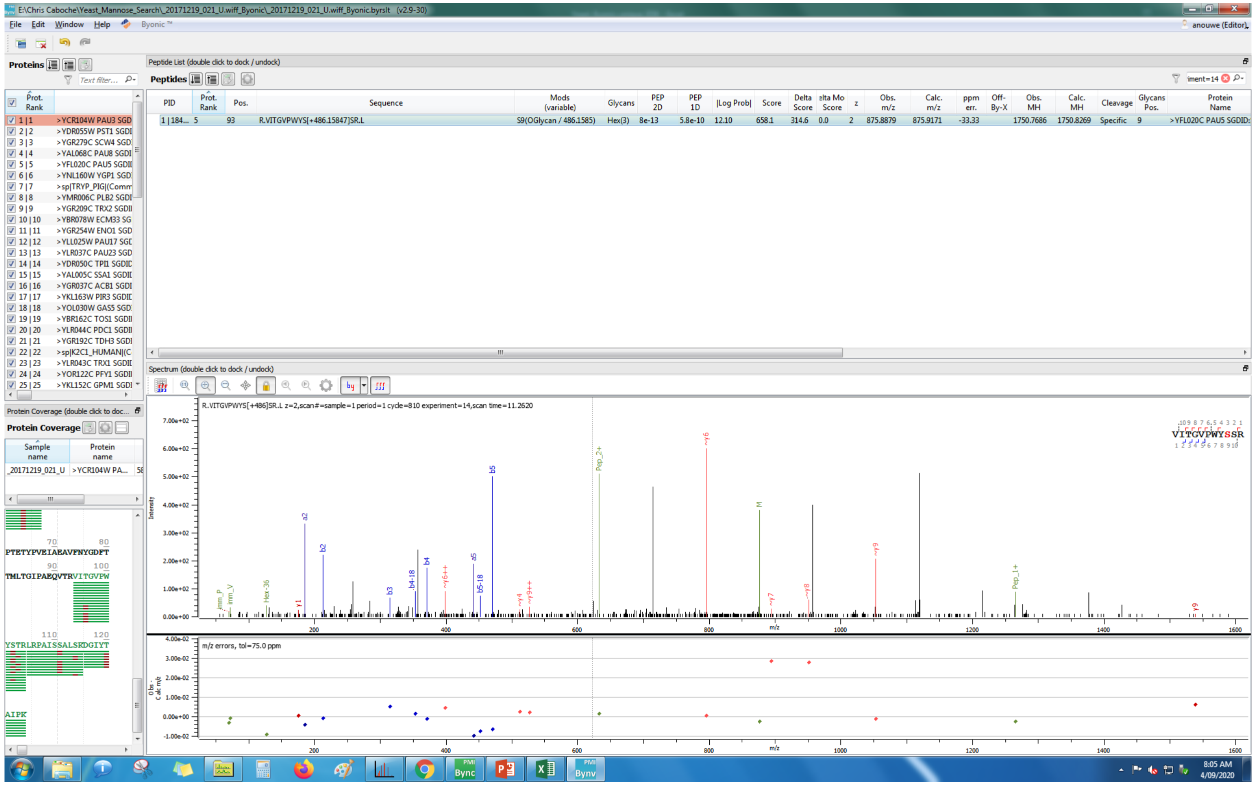
E:\Chris Caboche\Yeast_Mannose_Search_20171219_021_U.wiff_Byonic_20171219_021_U.wiff_Byonic.byrslt (v2.9-30)
File Edit Window Help Byonic™
Proteins
Peptide List (double click to dock / undock)
Peptides
PID Prot. Rank Pos. Sequence Mods (variable) Glycans PEP 2D PEP 1D |Log Prob| Score Delta Score Delta Mod Score z Obs. m/z Calc. m/z ppm err. Off-By-X Obs. MH Calc. MH Cleavage Glycans Pos. Protein Name
1 | 184... 5 93 R.VITGVPWYS[+486.15847]SR.L S9(OGlycan / 486.1585) Hex(3) 8e-13 5.8e-10 12.10 658.1 314.6 0.0 2 875.8879 875.9171 -33.33 1750.7686 1750.8269 Specific 9 >YFL020C PAU5 SGDID:
Spectrum (double click to dock / undock)
R.VITGVPWYS[+486]SR.L z=2,scan#=sample=1 period=1 cycle=810 experiment=14,scan time=11.2620
VITGVPWYSSR
m/z errors, tol=75.0 ppm
Protein Coverage (double click to dock / undock)
Protein Coverage
Sample name
Protein name
_20171219_021_U
>YCR104W PA...
PTETYPVELAEAVFNYGDFT
TMLTGIPAEQVTRVITGVPW
YSTRLRPAISSALSKDGIYT
AIPK
8:05 AM
4/09/2020

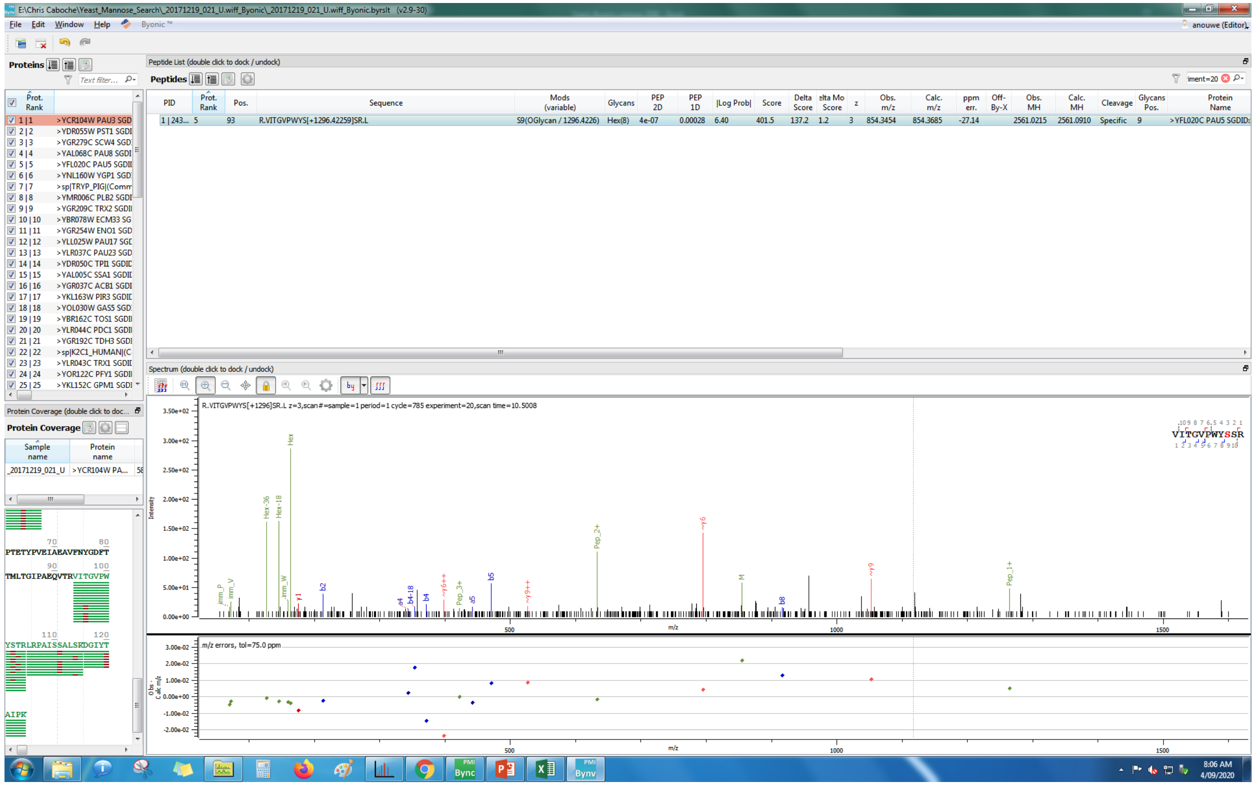
E:\Chris Caboche\Yeast_Mannose_Search_20171219_021_U.wiff_Byonic_20171219_021_U.wiff_Byonic.byrslt (v2.9-30)
File Edit Window Help Byonic ™
anouwe (Editor)
Proteins
Peptide List (double click to dock / undock)
Peptides
Prot. Rank
1 | 1 >YCR104W PAU3 SGD
2 | 2 >YDR055W PST1 SGDI
3 | 3 >YGR279C SCW4 SGD
4 | 4 >YAL068C PAU8 SGDI
5 | 5 >YFL020C PAU5 SGDI
6 | 6 >YNL160W YGP1 SGD
7 | 7 >sp|TRYP_PIG|(Comm
8 | 8 >YMR006C PLB2 SGDI
9 | 9 >YGR209C TRX2 SGDI
PID | Prot. Rank | Pos. | Sequence | Mods (variable) | Glycans | PEP 2D | PEP 1D | |Log Prob| | Score | Delta Score | Delta Mod Score | z | Obs. m/z | Calc. m/z | ppm err. | Off-By-X | Obs. MH | Calc. MH | Cleavage | Glycans Pos. | Protein Name
1 | 243... | 5 | 93 | R.VITGVPWYS[+1296.42259]SR.L | S9(OGlycan / 1296.4226) | Hex(8) | 4e-07 | 0.00028 | 6.40 | 401.5 | 137.2 | 1.2 | 3 | 854.3454 | 854.3685 | -27.14 | | 2561.0215 | 2561.0910 | Specific | 9 | >YFL020C PAU5 SGDID:
Spectrum (double click to dock / undock)
R.VITGVPWYS[+1296]SR.L z=3,scan#=sample=1 period=1 cycle=785 experiment=20,scan time=10.5008
VITGVPWYSSR
Protein Coverage
m/z errors, tol=75.0 ppm

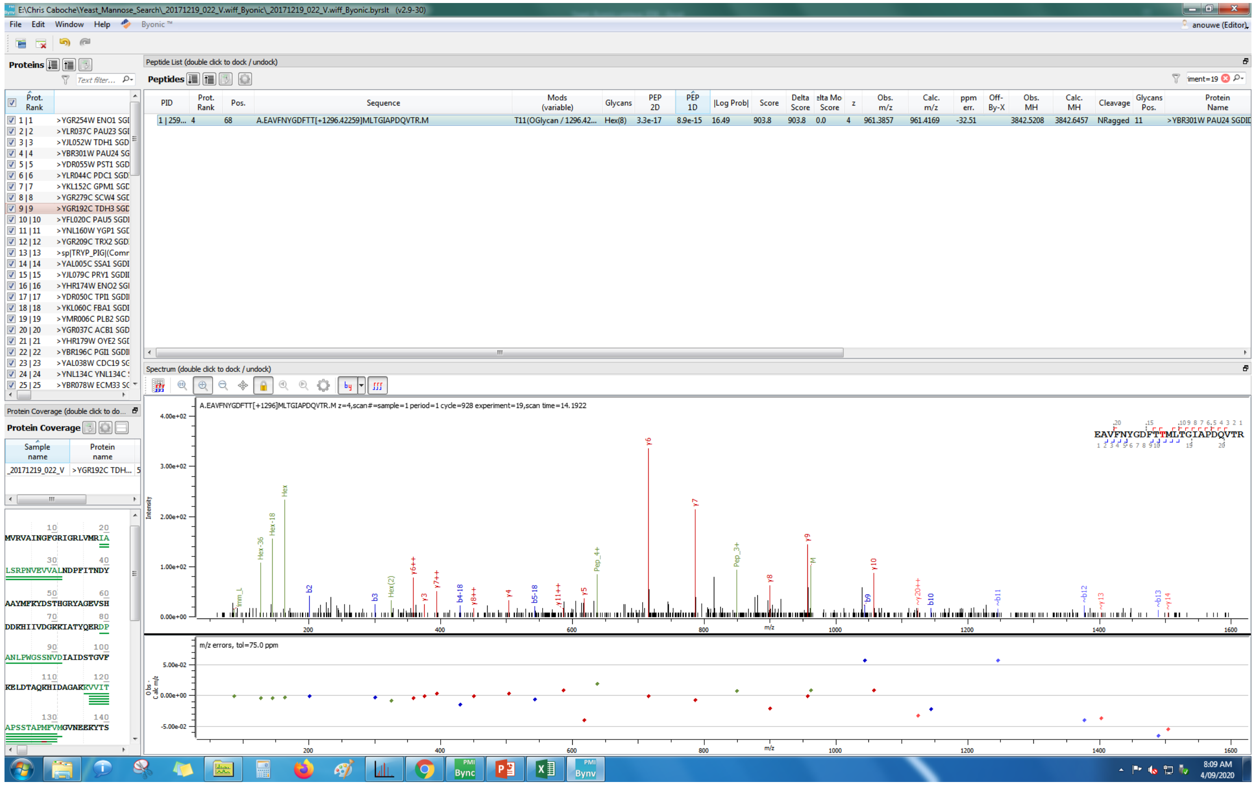
E:\Chris Caboche\Yeast_Mannose_Search_20171219_022_V.wiff_Byonic_20171219_022_V.wiff_Byonic.byrslt (v2.9-30)
Proteins
Peptide List (double click to dock / undock)
Peptides
A.EAVFNYGDFTT[+1296.42259]MLTGIAPDQVTR.M
Spectrum (double click to dock / undock)
A.EAVFNYGDFTT[+1296]MLTGIAPDQVTR.M z=4,scan#=sample=1 period=1 cycle=928 experiment=19,scan time=14.1922
EAVFNYGDFTTMLTGIAPDQVTR
m/z errors, tol=75.0 ppm
Protein Coverage

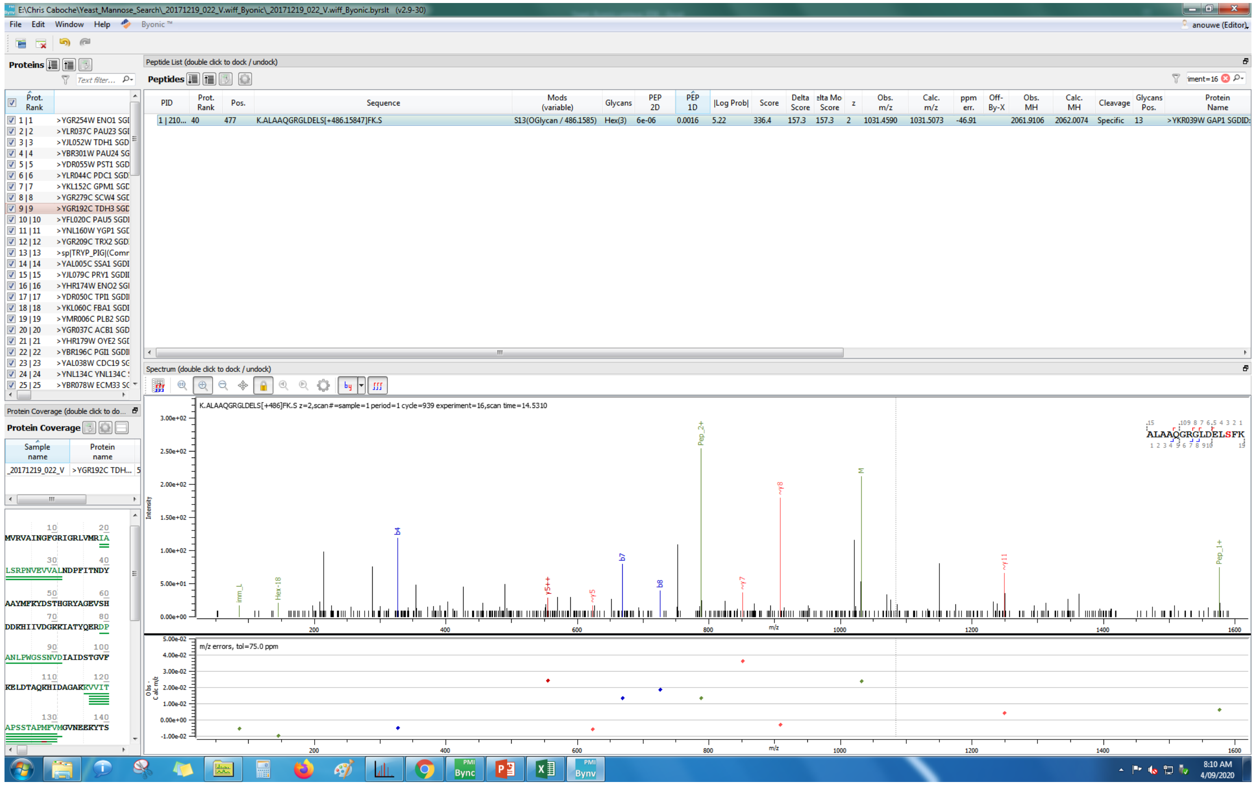
E:\Chris Caboche\Yeast_Mannose_Search_20171219_022_V.wiff_Byonic_20171219_022_V.wiff_Byonic.byrslt (v2.9-30)
File Edit Window Help
Proteins
Peptide List (double click to dock / undock)
Peptides
K.ALAAQGRGLDELS[+486.15847]FK.S
S13(OGlycan / 486.1585)
Hex(3)
>YKR039W GAP1 SGDID:
Spectrum (double click to dock / undock)
K.ALAAQGRGLDELS[+486]FK.S z=2,scan#=sample=1 period=1 cycle=939 experiment=16,scan time=14.5310
m/z errors, tol=75.0 ppm
Protein Coverage (double click to dock / undock)
Protein Coverage
MVRVAINGFGRIGRLVMRIA
LSRPNVEVVALNDPFITNDY
AAYMFKYDSTHGRYAGEVSH
DDKHIIVDGKKIATYQERDP
ANLPWGSSNVDIAIDSTGVF
KELDTAQKHIDAGAKKVVIT
APSSTAPMFVMGVNEEKYTS

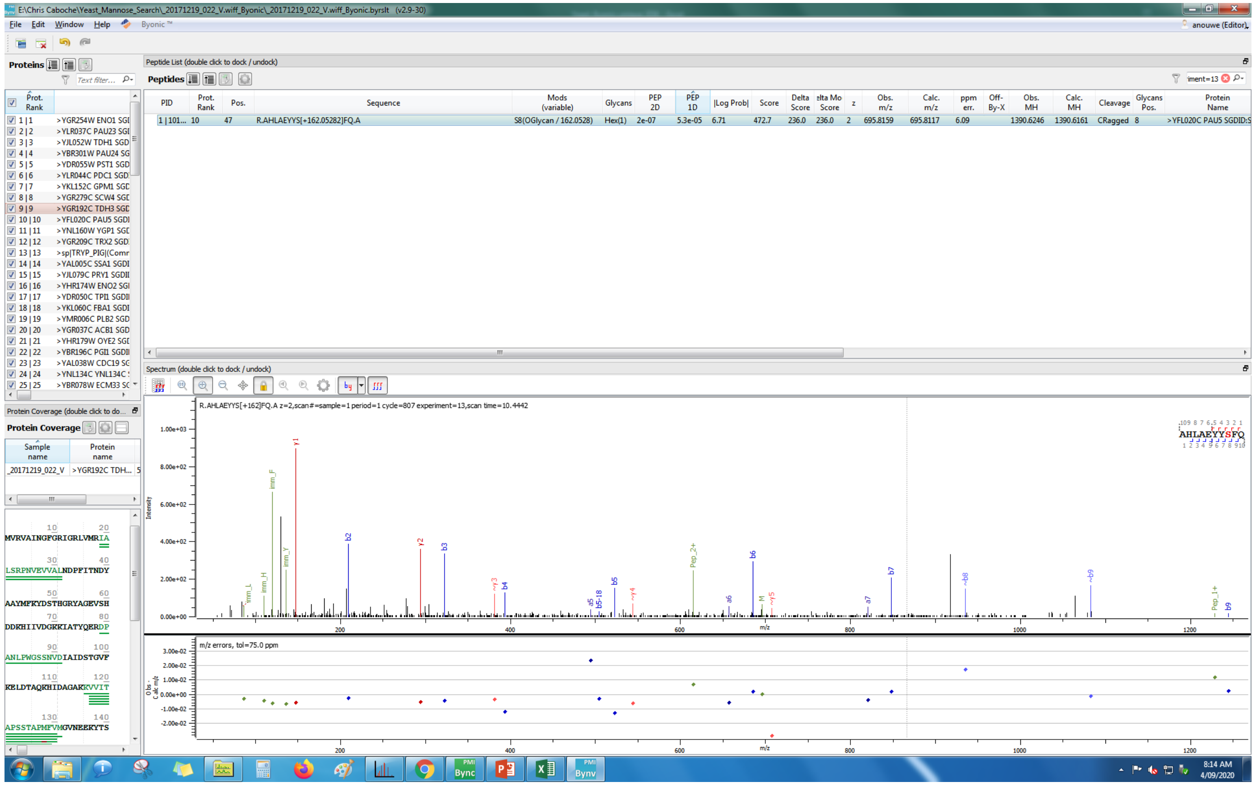
E:\Chris Caboche\Yeast_Mannose_Search_20171219_022_V.wiff_Byonic_20171219_022_V.wiff_Byonic.byrslt (v2.9-30)
File Edit Window Help Byonic™
anouwe (Editor)
Proteins
Peptide List (double click to dock / undock)
Peptides
PID | Prot. Rank | Pos. | Sequence | Mods (variable) | Glycans | PEP 2D | PEP 1D | |Log Prob| | Score | Delta Score | elta Mo Score | z | Obs. m/z | Calc. m/z | ppm err. | Off-By-X | Obs. MH | Calc. MH | Cleavage | Glycans Pos. | Protein Name
1 | 101... 10 47 R.AHLAEYYS[+162.05282]FQ.A S8(OGlycan / 162.0528) Hex(1) 2e-07 5.3e-05 6.71 472.7 236.0 236.0 2 695.8159 695.8117 6.09 1390.6246 1390.6161 CRagged 8 >YFL020C PAU5 SGDID:S
Spectrum (double click to dock / undock)
R.AHLAEYYS[+162]FQ.A z=2,scan#=sample=1 period=1 cycle=807 experiment=13,scan time=10.4442
AHLAEYYSFQ
Protein Coverage
m/z errors, tol=75.0 ppm

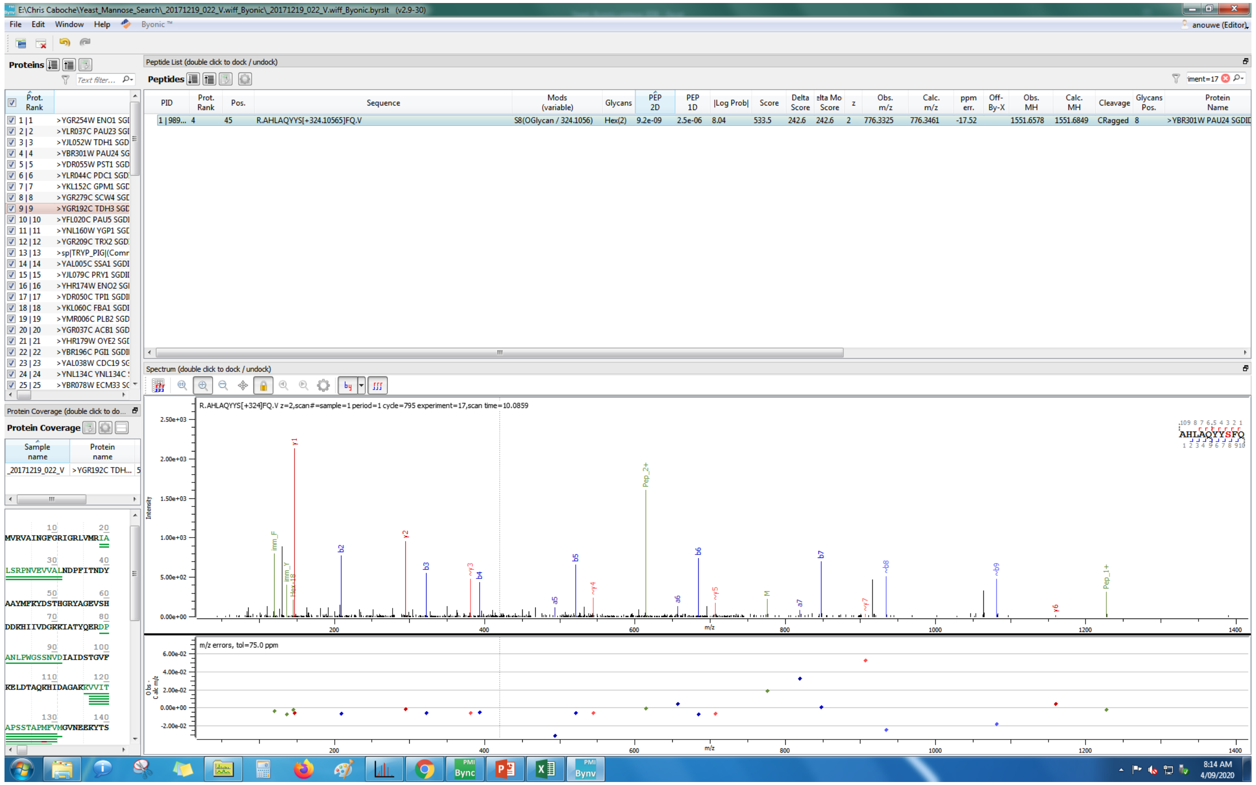
E:\Chris Caboche\Yeast_Mannose_Search_20171219_022_V.wiff_Byonic_20171219_022_V.wiff_Byonic.byrslt (v2.9-30)
File Edit Window Help Byonic™
anouwe (Editor)
Proteins
Peptide List (double click to dock / undock)
Peptides
PID | Prot. Rank | Pos. | Sequence | Mods (variable) | Glycans | PEP 2D | PEP 1D | |Log Prob| | Score | Delta Score | elta Mo Score | z | Obs. m/z | Calc. m/z | ppm err. | Off-By-X | Obs. MH | Calc. MH | Cleavage | Glycans Pos. | Protein Name
1 | 989... | 4 | 45 | R.AHLAQYYS[+324.10565]FQ.V | S8(OGlycan / 324.1056) | Hex(2) | 9.2e-09 | 2.5e-06 | 8.04 | 533.5 | 242.6 | 242.6 | 2 | 776.3325 | 776.3461 | -17.52 | | 1551.6578 | 1551.6849 | CRagged | 8 | >YBR301W PAU24 SGDID
Spectrum (double click to dock / undock)
R.AHLAQYYS[+324]FQ.V z=2,scan#=sample=1 period=1 cycle=795 experiment=17,scan time=10.0859
AHLAQYYSFQ
m/z errors, tol=75.0 ppm
Protein Coverage
Sample name
Protein name
_20171219_022_V
>YGR192C TDH...
8:14 AM
4/09/2020

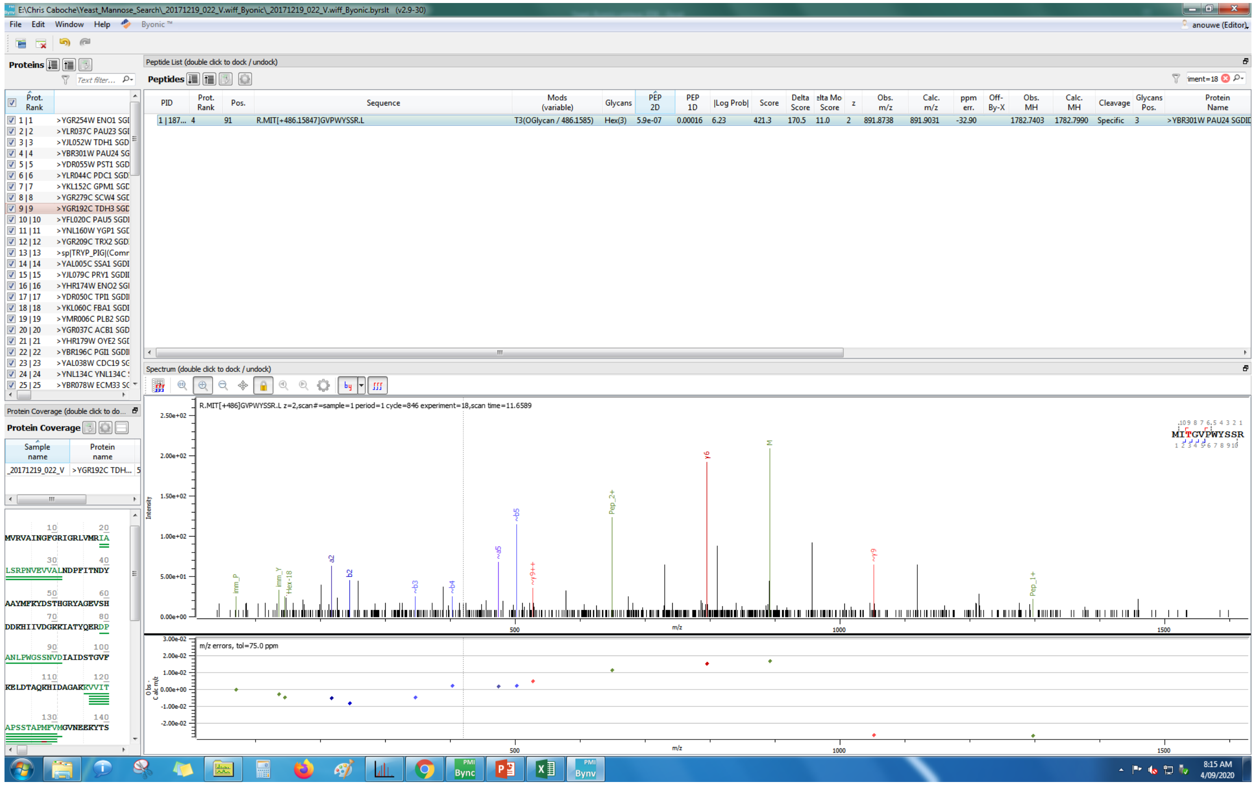
E:\Chris Caboche\Yeast_Mannose_Search_20171219_022_V.wiff_Byonic_20171219_022_V.wiff_Byonic.byrslt (v2.9-30)
File Edit Window Help Byonic™
Proteins
Peptide List (double click to dock / undock)
Peptides
R.MIT[+486.15847]GVPWYSSR.L
T3(OGlycan / 486.1585)
Hex(3)
>YBR301W PAU24 SGDID
Spectrum (double click to dock / undock)
R.MIT[+486]GVPWYSSR.L z=2,scan#=sample=1 period=1 cycle=846 experiment=18,scan time=11.6589
MITGVPWYSSR
m/z errors, tol=75.0 ppm
Protein Coverage
8:15 AM
4/09/2020

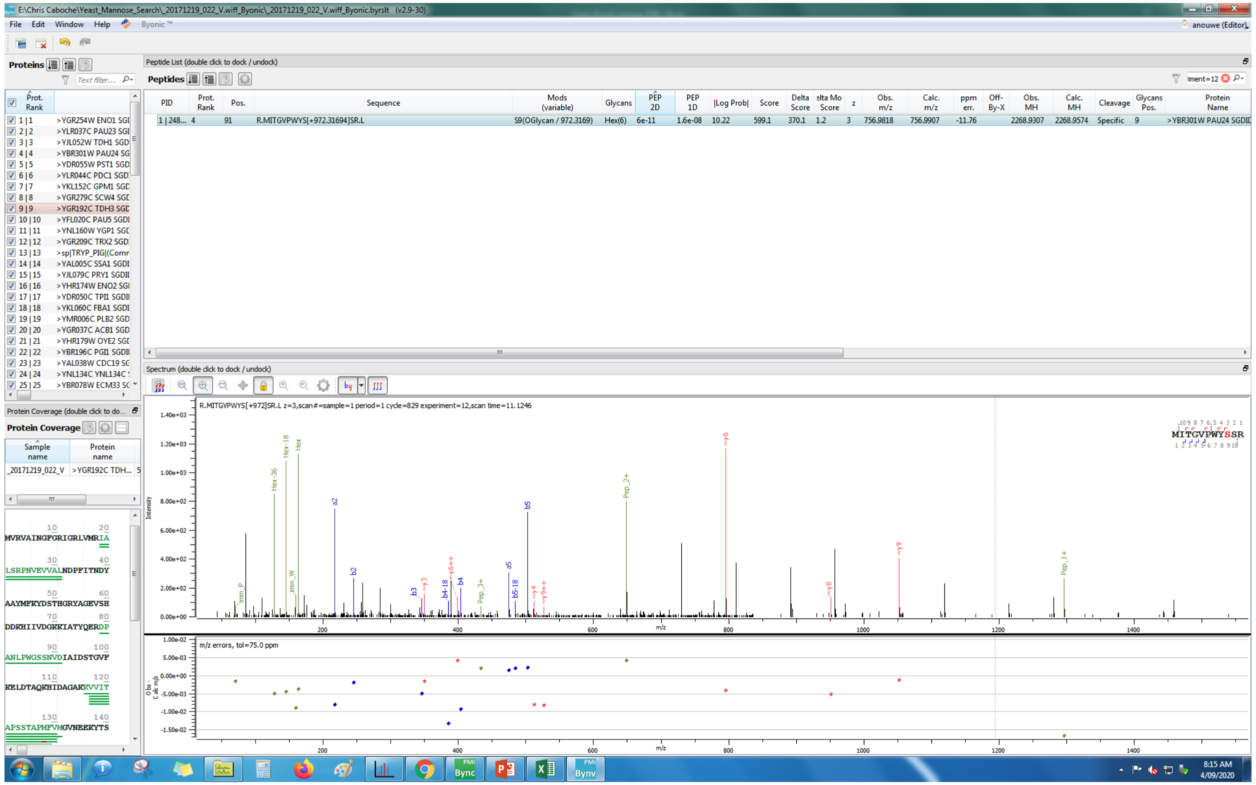

E:\Chris Caboche\Yeast_Mannose_Search_20171219_022_V.wiff_Byonic_20171219_022_V.wiff_Byonic.byrslt (v2.9-30)
File Edit Window Help Byonic ™
anouwe (Editor)
Proteins
Peptide List (double click to dock / undock)
Peptides
PID | Prot. Rank | Pos. | Sequence | Mods (variable) | Glycans | PEP 2D | PEP 1D | |Log Prob| | Score | Delta Score | elta Mo Score | z | Obs. m/z | Calc. m/z | ppm err. | Off-By-X | Obs. MH | Calc. MH | Cleavage | Glycans Pos. | Protein Name
1 | 248... 4 91 R.MITGVPWYS[+972.31694]SR.L S9(OGlycan / 972.3169) Hex(6) 6e-11 1.6e-08 10.22 599.1 370.1 1.2 3 756.9818 756.9907 -11.76 2268.9307 2268.9574 Specific 9 >YBR301W PAU24 SGDID
Spectrum (double click to dock / undock)
R.MITGVPWYS[+972]SR.L z=3,scan#=sample=1 period=1 cycle=829 experiment=12,scan time=11.1246
MITGVPWYSSR
m/z errors, tol=75.0 ppm
Protein Coverage

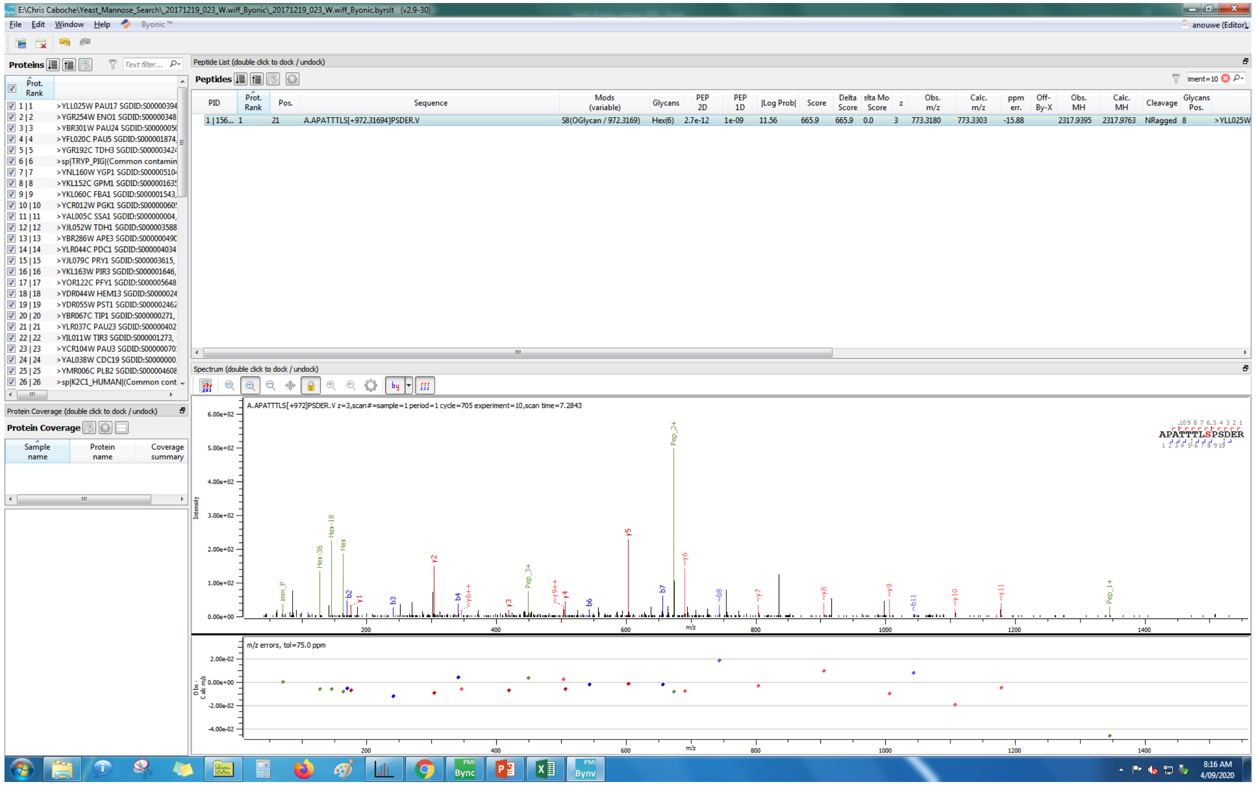
E:\Chris Caboche\Yeast_Mannose_Search_20171219_023_W.wiff_Byonic_20171219_023_W.wiff_Byonic.byrslt (v2.9-30)
File Edit Window Help Byonic™
Proteins
Peptide List (double click to dock / undock)
Peptides
A.APATTTLS[+972.31694]PSDER.V
S8(OGlycan / 972.3169)
Hex(6)
Spectrum (double click to dock / undock)
A.APATTTLS[+972]PSDER.V z=3,scan#=sample=1 period=1 cycle=705 experiment=10,scan time=7.2843
APATTTLSPSDER
m/z errors, tol=75.0 ppm
Protein Coverage (double click to dock / undock)
Protein Coverage
Sample name
Protein name
Coverage summary
8:16 AM
4/09/2020

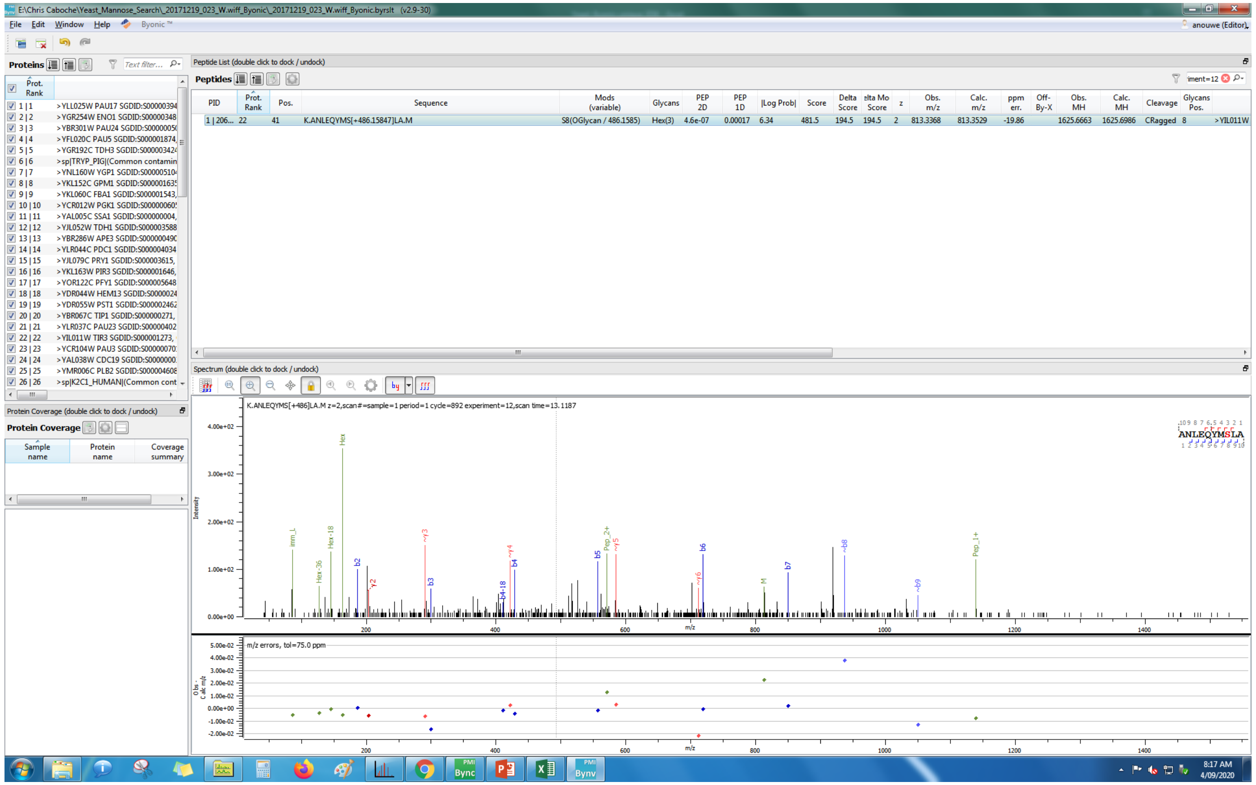

E:\Chris Caboche\Yeast_Mannose_Search_20171219_023_W.wiff_Byonic_20171219_023_W.wiff_Byonic.byrslt (v2.9-30)
File Edit Window Help Byonic™
anouwe (Editor)
Proteins
Peptide List (double click to dock / undock)
Peptides
PID | Prot. Rank | Pos. | Sequence | Mods (variable) | Glycans | PEP 2D | PEP 1D | |Log Prob| | Score | Delta Score | elta Mo Score | z | Obs. m/z | Calc. m/z | ppm err. | Off-By-X | Obs. MH | Calc. MH | Cleavage | Glycans Pos.
1 | 206... 22 41 K.ANLEQYMS[+486.15847]LA.M S8(OGlycan / 486.1585) Hex(3) 4.6e-07 0.00017 6.34 481.5 194.5 194.5 2 813.3368 813.3529 -19.86 1625.6663 1625.6986 CRagged 8 >YIL011W
Protein Coverage (double click to dock / undock)
Protein Coverage
Sample name | Protein name | Coverage summary
Spectrum (double click to dock / undock)
K.ANLEQYMS[+486]LA.M z=2,scan#=sample=1 period=1 cycle=892 experiment=12,scan time=13.1187
ANLEQYMSLA
m/z errors, tol=75.0 ppm
8:17 AM
4/09/2020

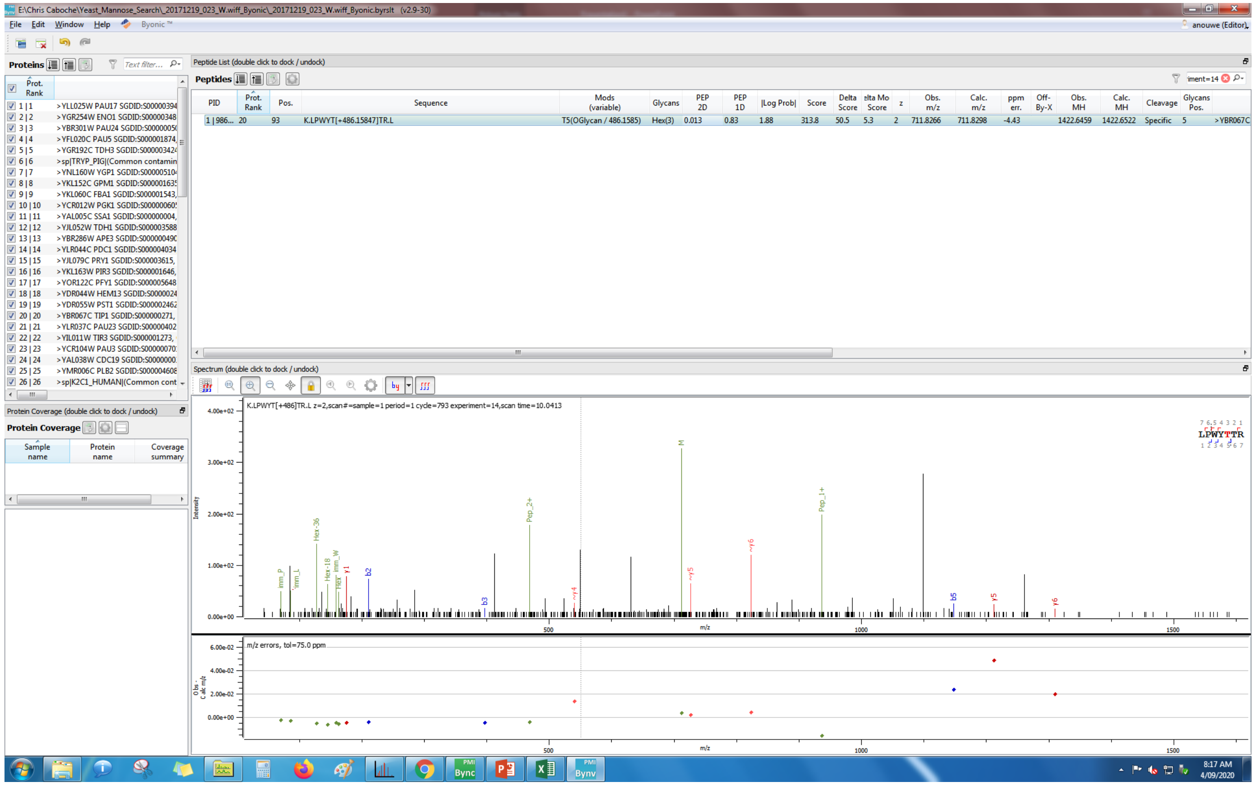
E:\Chris Caboche\Yeast_Mannose_Search_20171219_023_W.wiff_Byonic_20171219_023_W.wiff_Byonic.byrslt (v2.9-30)
File Edit Window Help Byonic™
anouwe (Editor)
Proteins
Peptide List (double click to dock / undock)
Peptides
PID | Prot. Rank | Pos. | Sequence | Mods (variable) | Glycans | PEP 2D | PEP 1D | |Log Prob| | Score | Delta Score | elta Mo Score | z | Obs. m/z | Calc. m/z | ppm err. | Off-By-X | Obs. MH | Calc. MH | Cleavage | Glycans Pos.
1 | 986... 20 93 K.LPWYT[+486.15847]TR.L T5(OGlycan / 486.1585) Hex(3) 0.013 0.83 1.88 313.8 50.5 5.3 2 711.8266 711.8298 -4.43 1422.6459 1422.6522 Specific 5 >YBR067C
Spectrum (double click to dock / undock)
K.LPWYT[+486]TR.L z=2,scan#=sample=1 period=1 cycle=793 experiment=14,scan time=10.0413
LPWYTTR
Intensity
m/z
m/z errors, tol=75.0 ppm
Protein Coverage (double click to dock / undock)
Protein Coverage
Sample name
Protein name
Coverage summary

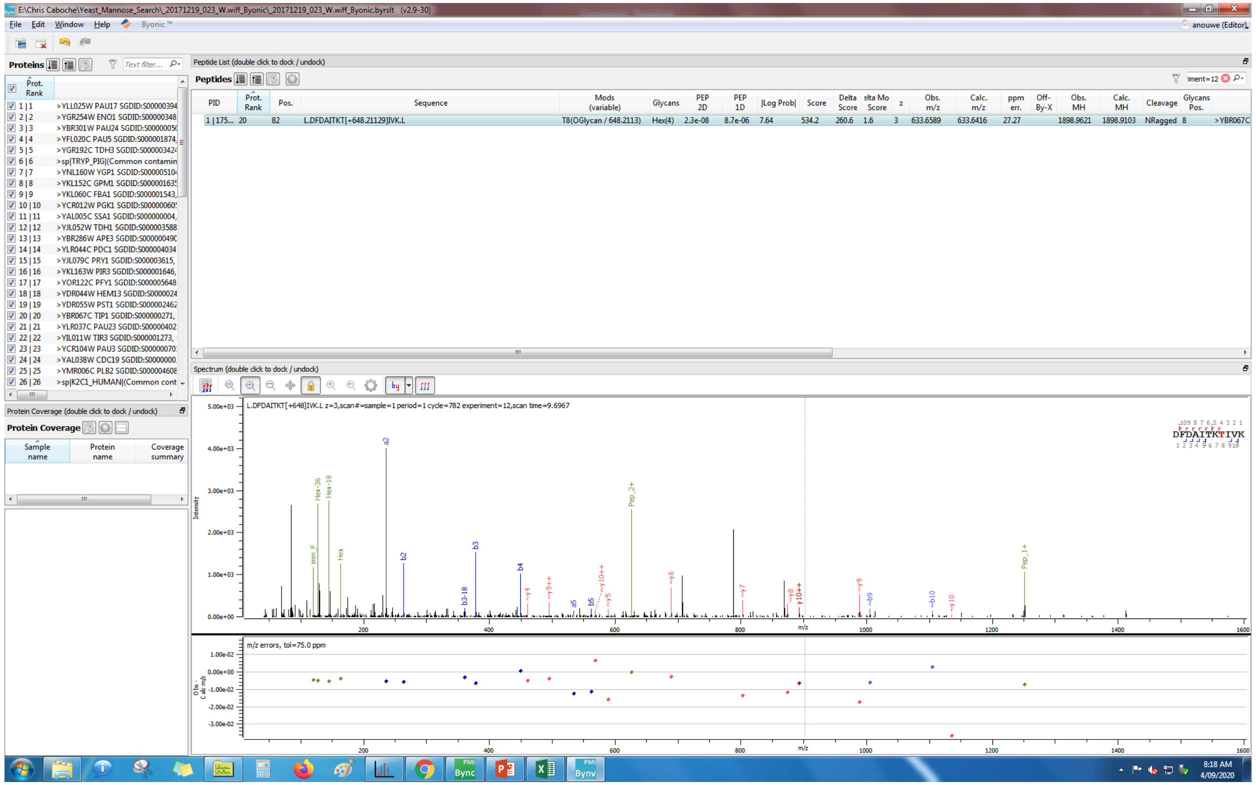
E:\Chris Caboche\Yeast_Mannose_Search_20171219_023_W.wiff_Byonic_20171219_023_W.wiff_Byonic.byrslt (v2.9-30)
Peptide List (double click to dock / undock)
Peptides
PID | Prot. Rank | Pos. | Sequence | Mods (variable) | Glycans | PEP 2D | PEP 1D | |Log Prob| | Score | Delta Score | Delta Mod Score | z | Obs. m/z | Calc. m/z | ppm err. | Off-By-X | Obs. MH | Calc. MH | Cleavage | Glycans Pos.
1 | 175... 20 82 L.DFDAITKT[+648.21129]IVK.L T8(OGlycan / 648.2113) Hex(4) 2.3e-08 8.7e-06 7.64 534.2 260.6 1.6 3 633.6589 633.6416 27.27 1898.9621 1898.9103 NRagged 8 >YBR067C
Spectrum (double click to dock / undock)
L.DFDAITKT[+648]IVK.L z=3,scan#=sample=1 period=1 cycle=782 experiment=12,scan time=9.6967
DFDAITKTIVK
m/z errors, tol=75.0 ppm

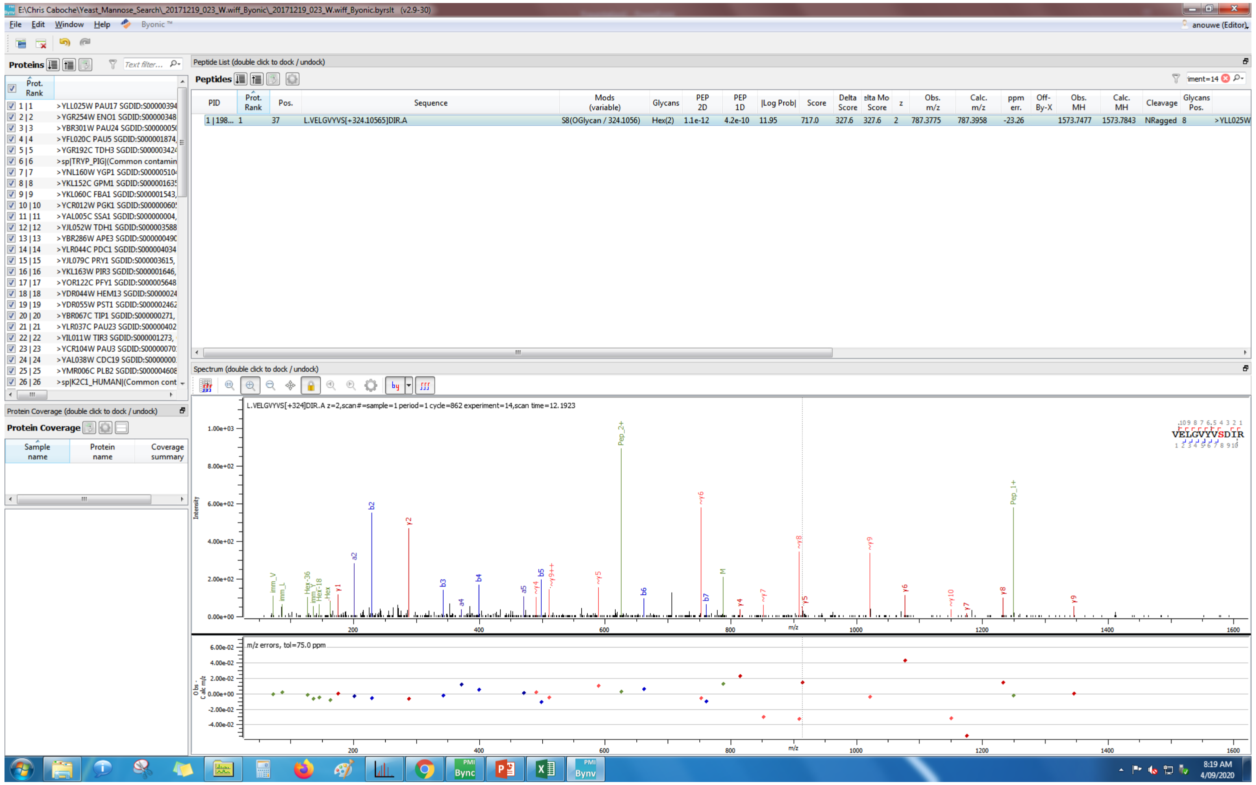
E:\Chris Caboche\Yeast_Mannose_Search_20171219_023_W.wiff_Byonic_20171219_023_W.wiff_Byonic.byrslt (v2.9-30)
File Edit Window Help Byonic™
anouwe (Editor)
Proteins
Peptide List (double click to dock / undock)
Peptides
PID Prot. Rank Pos. Sequence Mods (variable) Glycans PEP 2D PEP 1D |Log Prob| Score Delta Score Delta Mod Score z Obs. m/z Calc. m/z ppm err. Off-By-X Obs. MH Calc. MH Cleavage Glycans Pos.
1 | 198... 1 37 L.VELGVYVS[+324.10565]DIR.A S8(OGlycan / 324.1056) Hex(2) 1.1e-12 4.2e-10 11.95 717.0 327.6 327.6 2 787.3775 787.3958 -23.26 1573.7477 1573.7843 NRagged 8 >YLL025W
Spectrum (double click to dock / undock)
L.VELGVYVS[+324]DIR.A z=2,scan#=sample=1 period=1 cycle=862 experiment=14,scan time=12.1923
VELGVYVSDIR
Protein Coverage (double click to dock / undock)
Protein Coverage
Sample name Protein name Coverage summary
m/z errors, tol=75.0 ppm

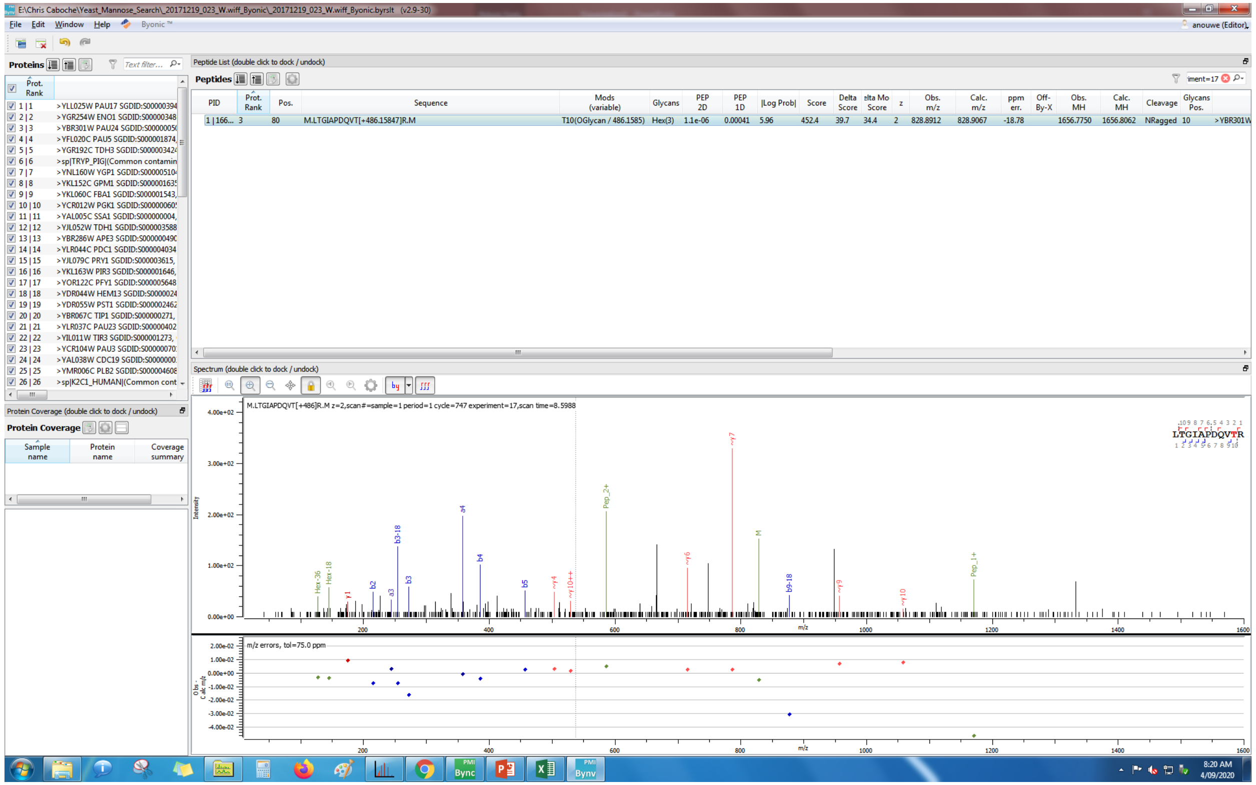
E:\Chris Caboche\Yeast_Mannose_Search_20171219_023_W.wiff_Byonic_20171219_023_W.wiff_Byonic.byrslt (v2.9-30)
File Edit Window Help Byonic™
anouwe (Editor)
Proteins
Peptide List (double click to dock / undock)
Peptides
PID | Prot. Rank | Pos. | Sequence | Mods (variable) | Glycans | PEP 2D | PEP 1D | |Log Prob| | Score | Delta Score | elta Mo Score | z | Obs. m/z | Calc. m/z | ppm err. | Off-By-X | Obs. MH | Calc. MH | Cleavage | Glycans Pos.
1 | 166... 3 80 M.LTGIAPDQVT[+486.15847]R.M T10(OGlycan / 486.1585) Hex(3) 1.1e-06 0.00041 5.96 452.4 39.7 34.4 2 828.8912 828.9067 -18.78 1656.7750 1656.8062 NRagged 10 >YBR301W
Spectrum (double click to dock / undock)
M.LTGIAPDQVT[+486]R.M z=2,scan#=sample=1 period=1 cycle=747 experiment=17,scan time=8.5988
LTGIAPDQVTR
Protein Coverage (double click to dock / undock)
Protein Coverage
Sample name
Protein name
Coverage summary
m/z errors, tol=75.0 ppm
Intensity
m/z

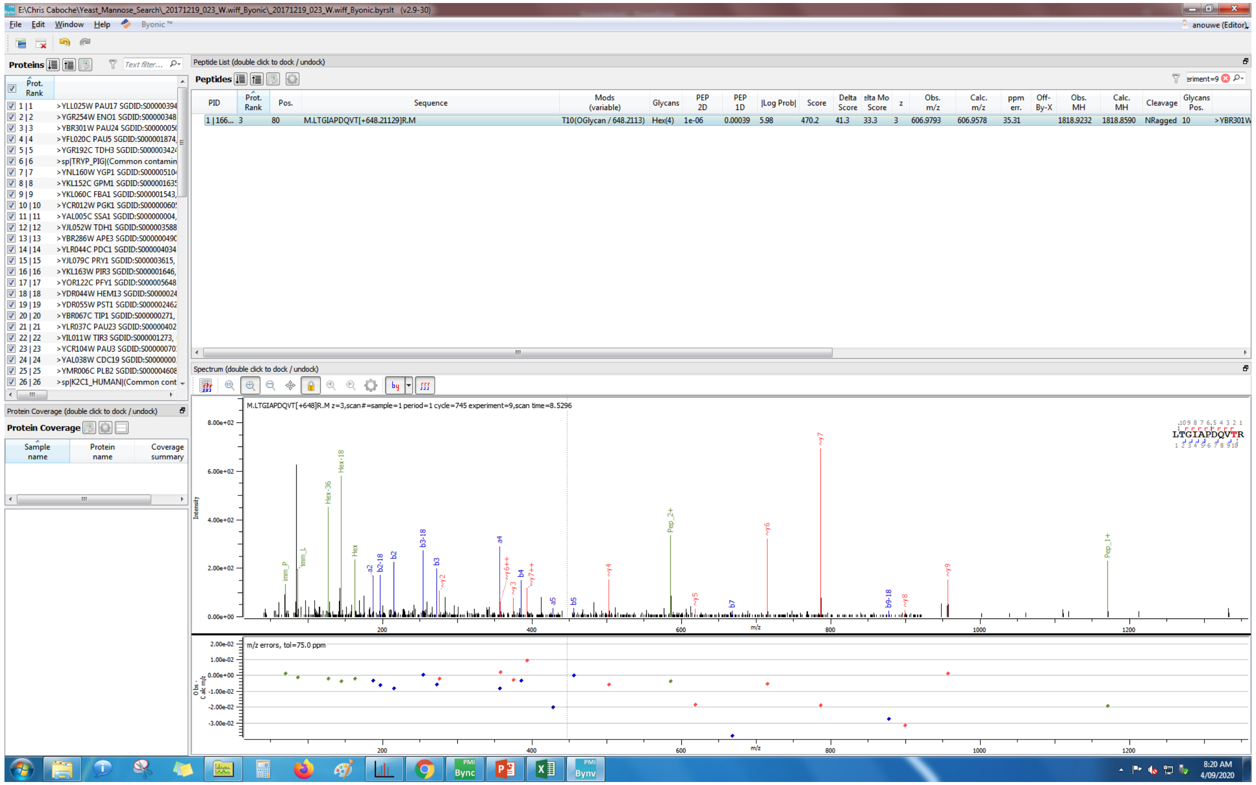
E:\Chris Caboche\Yeast_Mannose_Search_20171219_023_W.wiff_Byonic_20171219_023_W.wiff_Byonic.byrslt (v2.9-30)
Peptide List (double click to dock / undock)
Peptides
PID
Prot. Rank
Pos.
Sequence
Mods (variable)
Glycans
PEP 2D
PEP 1D
|Log Prob|
Score
Delta Score
Obs. m/z
Calc. m/z
ppm err.
Obs. MH
Calc. MH
Cleavage
Glycans Pos.
1 | 166... 3 80 M.LTGIAPDQVT[+648.21129]R.M T10(OGlycan / 648.2113) Hex(4) 1e-06 0.00039 5.98 470.2 41.3 33.3 3 606.9793 606.9578 35.31 1818.9232 1818.8590 NRagged 10 >YBR301W
Spectrum (double click to dock / undock)
M.LTGIAPDQVT[+648]R.M z=3,scan#=sample=1 period=1 cycle=745 experiment=9,scan time=8.5296
LTGIAPDQVTR
m/z errors, tol=75.0 ppm
Protein Coverage

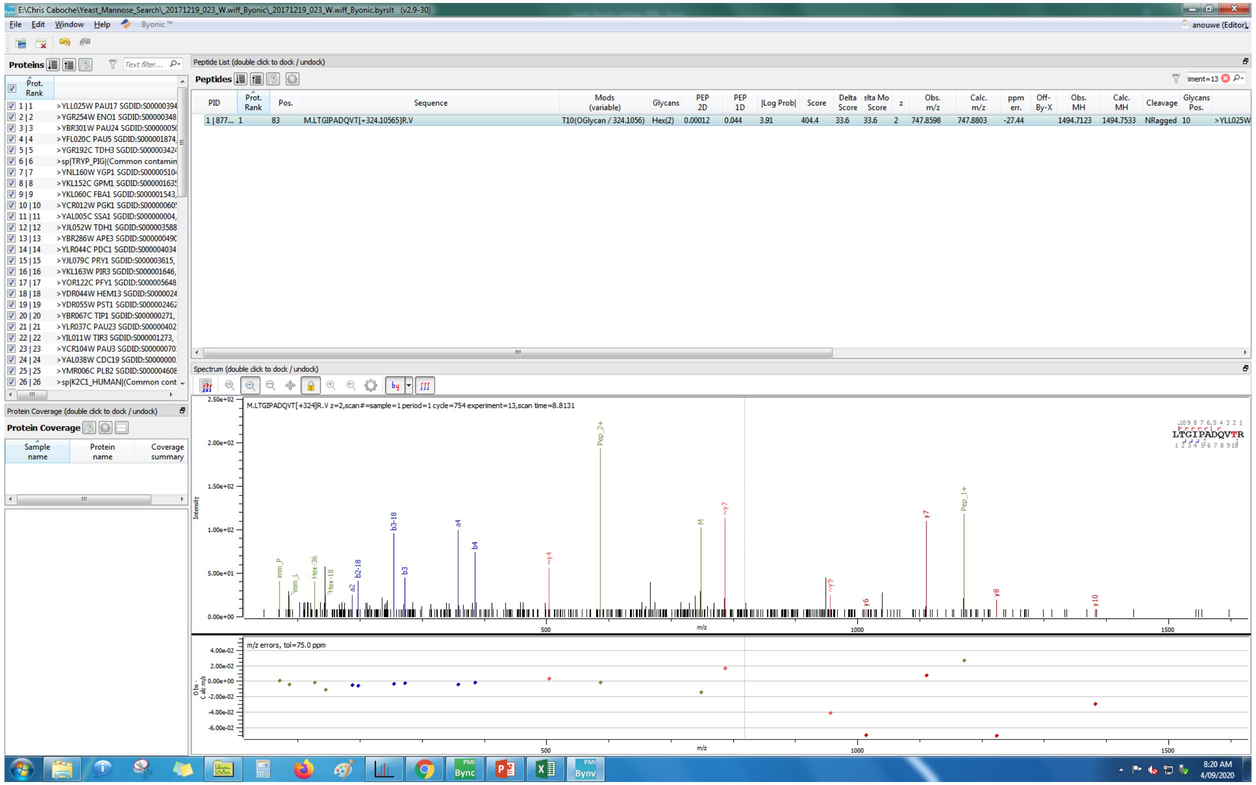

File
Edit
Window
Help
Byonic
Proteins
Peptide List (double click to dock / undock)
Peptides
PID
Prot. Rank
Pos.
Sequence
Mods (variable)
Glycans
PEP 2D
PEP 1D
|Log Prob|
Score
Obs. m/z
Calc. m/z
ppm err.
Obs. MH
Calc. MH
Cleavage
M.LTGIPADQVT[+324.10565]R.V
Protein Coverage (double click to dock / undock)
Protein Coverage
Sample name
Protein name
Coverage summary
Spectrum (double click to dock / undock)
M.LTGIPADQVT[+324]R.V z=2,scan#=sample=1 period=1 cycle=754 experiment=13,scan time=8.8131
LTGIPADQVTR
Pep_2+
Pep_1+
m/z errors, tol=75.0 ppm
m/z

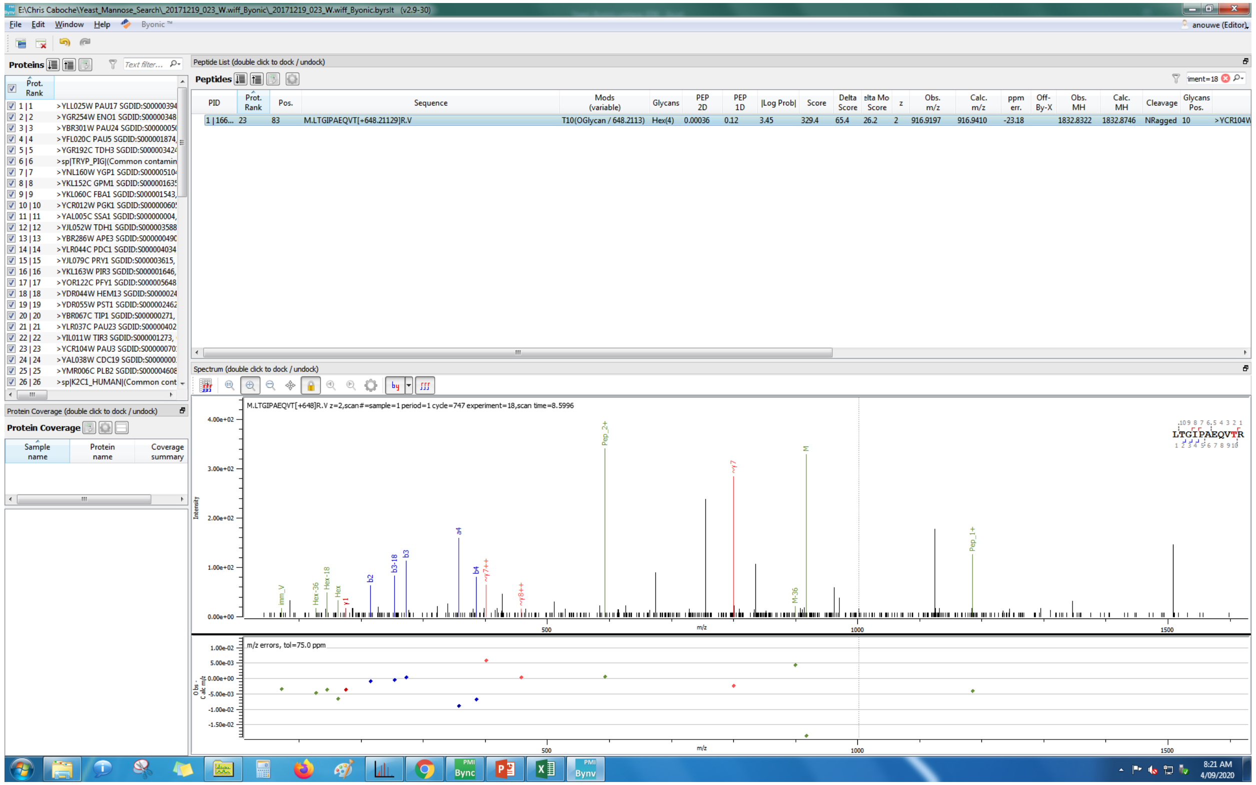

E:\Chris Caboche\Yeast_Mannose_Search_20171219_023_W.wiff_Byonic_20171219_023_W.wiff_Byonic.byrslt (v2.9-30)
File Edit Window Help Byonic™
Proteins
Peptide List (double click to dock / undock)
Peptides
PID
Prot. Rank
Pos.
Sequence
Mods (variable)
Glycans
PEP 2D
PEP 1D
|Log Prob|
Score
Delta Score
z
Obs. m/z
Calc. m/z
ppm err.
Off-By-X
Obs. MH
Calc. MH
Cleavage
Glycans Pos.
M.LTGIPAEQVT[+648.21129]R.V
T10(OGlycan / 648.2113)
Hex(4)
Spectrum (double click to dock / undock)
M.LTGIPAEQVT[+648]R.V z=2,scan#=sample=1 period=1 cycle=747 experiment=18,scan time=8.5996
Protein Coverage (double click to dock / undock)
Protein Coverage
Sample name
Protein name
Coverage summary
Intensity
m/z
m/z errors, tol=75.0 ppm
LTGIPAEQVTR
8:21 AM
4/09/2020

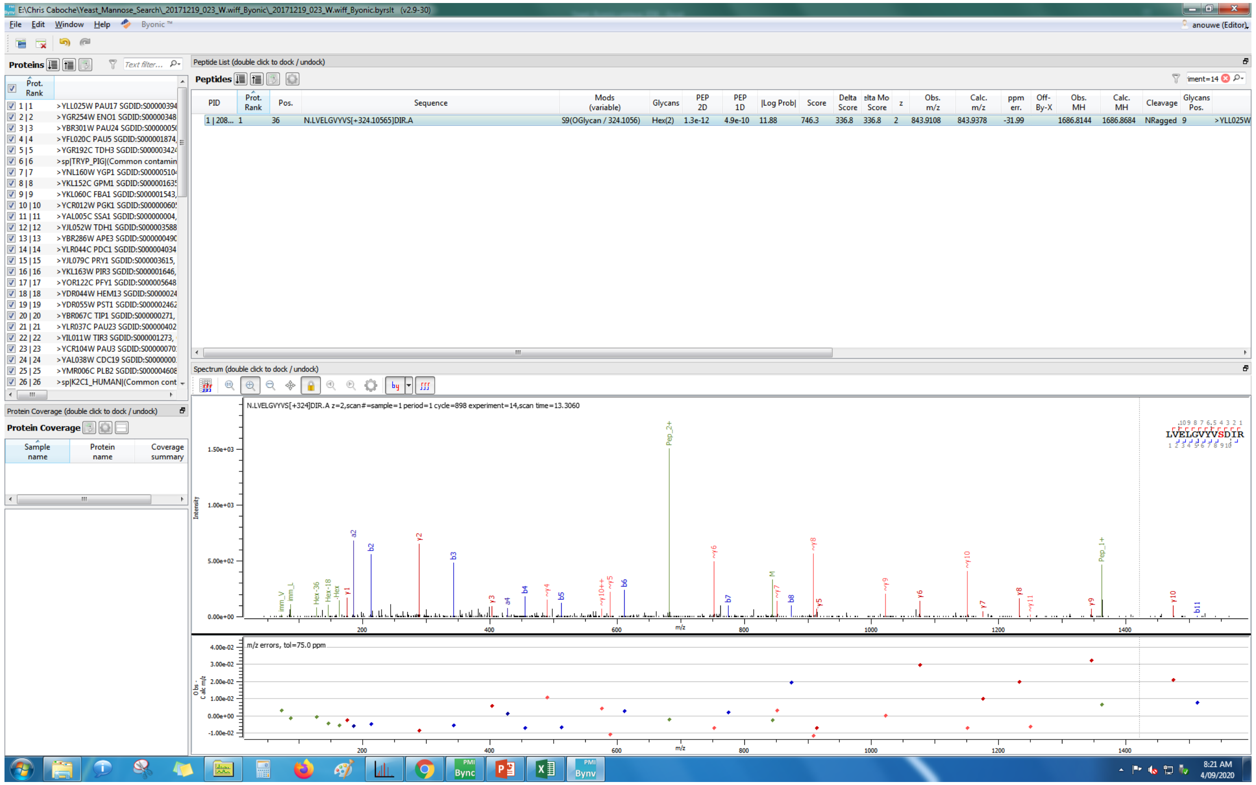
E:\Chris Caboche\Yeast_Mannose_Search_20171219_023_W.wiff_Byonic_20171219_023_W.wiff_Byonic.byrslt (v2.9-30)
Peptide List (double click to dock / undock)
Peptides
PID | Prot. Rank | Pos. | Sequence | Mods (variable) | Glycans | PEP 2D | PEP 1D | |Log Prob| | Score | Delta Score | Delta Mod Score | z | Obs. m/z | Calc. m/z | ppm err. | Off-By-X | Obs. MH | Calc. MH | Cleavage | Glycans Pos.
1 | 208... 1 36 N.LVELGVYVS[+324.10565]DIR.A S9(OGlycan / 324.1056) Hex(2) 1.3e-12 4.9e-10 11.88 746.3 336.8 336.8 2 843.9108 843.9378 -31.99 1686.8144 1686.8684 NRagged 9 >YLL025W
Spectrum (double click to dock / undock)
N.LVELGVYVS[+324]DIR.A z=2,scan#=sample=1 period=1 cycle=898 experiment=14,scan time=13.3060
m/z errors, tol=75.0 ppm
Protein Coverage
Sample name
Protein name
Coverage summary

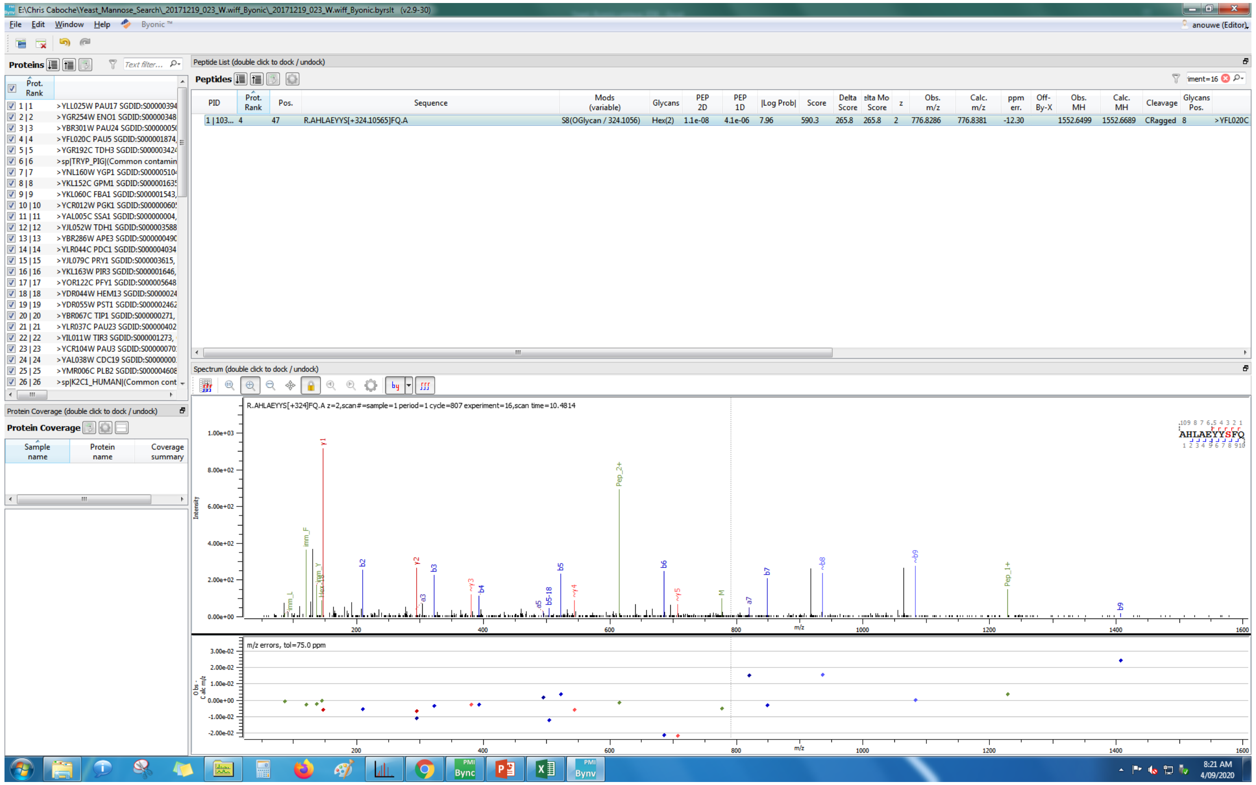
E:\Chris Caboche\Yeast_Mannose_Search_20171219_023_W.wiff_Byonic_20171219_023_W.wiff_Byonic.byrslt (v2.9-30)
File Edit Window Help Byonic™
Proteins
Peptide List (double click to dock / undock)
Peptides
PID | Prot. Rank | Pos. | Sequence | Mods (variable) | Glycans | PEP 2D | PEP 1D | |Log Prob| | Score | Delta Score | Delta Mod Score | z | Obs. m/z | Calc. m/z | ppm err. | Off-By-X | Obs. MH | Calc. MH | Cleavage | Glycans Pos.
1 | 103... 4 47 R.AHLAEYYS[+324.10565]FQ.A S8(OGlycan / 324.1056) Hex(2) 1.1e-08 4.1e-06 7.96 590.3 265.8 265.8 2 776.8286 776.8381 -12.30 1552.6499 1552.6689 CRagged 8 >YFL020C
>YLL025W PAU17 SGDID:S00000394
>YGR254W ENO1 SGDID:S00000348
>YBR301W PAU24 SGDID:S00000005
>YFL020C PAU5 SGDID:S000001874
>YGR192C TDH3 SGDID:S00000342
>sp|TRYP_PIG|(Common contamin
>YNL160W YGP1 SGDID:S00000510
>YKL152C GPM1 SGDID:S00000163
>YKL060C FBA1 SGDID:S000001543
>YCR012W PGK1 SGDID:S00000060
>YAL005C SSA1 SGDID:S000000004
>YJL052W TDH1 SGDID:S000003588
>YBR286W APE3 SGDID:S00000049
>YLR044C PDC1 SGDID:S00000403
>YJL079C PRY1 SGDID:S000003615
>YKL163W PIR3 SGDID:S000001646
>YOR122C PFY1 SGDID:S00000564
>YDR044W HEM13 SGDID:S0000002
>YDR055W PST1 SGDID:S00000246
>YBR067C TIP1 SGDID:S000000271
>YLR037C PAU23 SGDID:S00000040
>YIL011W TIR3 SGDID:S000001273
>YCR104W PAU3 SGDID:S00000007
>YAL038W CDC19 SGDID:S0000000
>YMR006C PLB2 SGDID:S00000460
>sp|K2C1_HUMAN|(Common cont
Protein Coverage (double click to dock / undock)
Protein Coverage
Sample name | Protein name | Coverage summary
Spectrum (double click to dock / undock)
R.AHLAEYYS[+324]FQ.A z=2,scan#=sample=1 period=1 cycle=807 experiment=16,scan time=10.4814
AHLAEYYSFQ
Intensity
m/z
m/z errors, tol=75.0 ppm
8:21 AM
4/09/2020

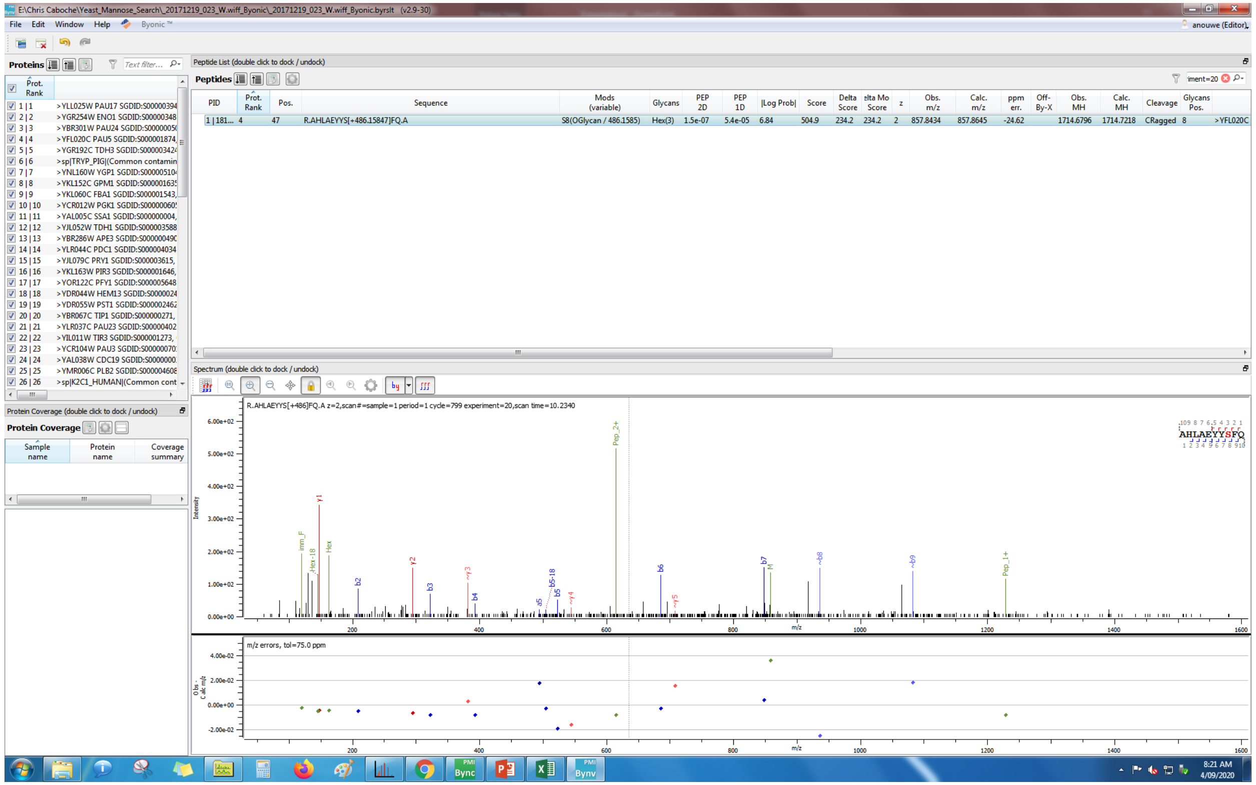

File Edit Window Help Byonic
Proteins
Peptide List (double click to dock / undock)
Peptides
PID
Prot. Rank
Pos.
Sequence
Mods (variable)
Glycans
PEP 2D
PEP 1D
|Log Prob|
Score
Delta Score
Obs. m/z
Calc. m/z
ppm err.
Obs. MH
Calc. MH
Cleavage
R.AHLAEYYS[+486.15847]FQ.A
S8(OGlycan / 486.1585)
Hex(3)
Spectrum (double click to dock / undock)
R.AHLAEYYS[+486]FQ.A z=2,scan#=sample=1 period=1 cycle=799 experiment=20,scan time=10.2340
AHLAEYYSFQ
Intensity
m/z
m/z errors, tol=75.0 ppm
Protein Coverage (double click to dock / undock)
Protein Coverage
Sample name
Protein name
Coverage summary

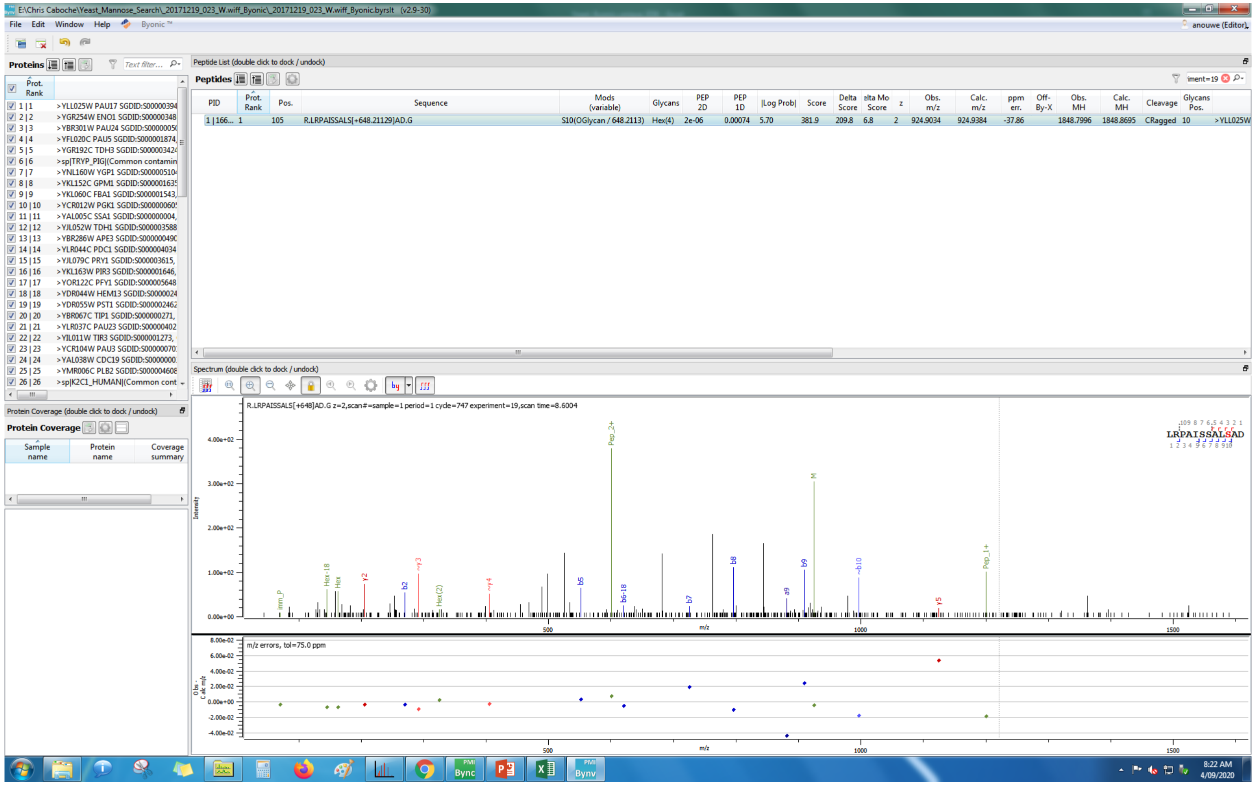
E:\Chris Caboche\Yeast_Mannose_Search_20171219_023_W.wiff_Byonic_20171219_023_W.wiff_Byonic.byrslt (v2.9-30)
File Edit Window Help Byonic™
anouwe (Editor)
Proteins
Peptide List (double click to dock / undock)
Peptides
PID | Prot. Rank | Pos. | Sequence | Mods (variable) | Glycans | PEP 2D | PEP 1D | |Log Prob| | Score | Delta Score | Delta Mod Score | z | Obs. m/z | Calc. m/z | ppm err. | Off-By-X | Obs. MH | Calc. MH | Cleavage | Glycans Pos.
1 | 166... 1 105 R.LRPAISSALS[+648.21129]AD.G S10(OGlycan / 648.2113) Hex(4) 2e-06 0.00074 5.70 381.9 209.8 6.8 2 924.9034 924.9384 -37.86 1848.7996 1848.8695 CRagged 10 >YLL025W
Spectrum (double click to dock / undock)
R.LRPAISSALS[+648]AD.G z=2,scan#=sample=1 period=1 cycle=747 experiment=19,scan time=8.6004
LRPAISSALSAD
Protein Coverage (double click to dock / undock)
Protein Coverage
Sample name
Protein name
Coverage summary
m/z errors, tol=75.0 ppm
Intensity
m/z

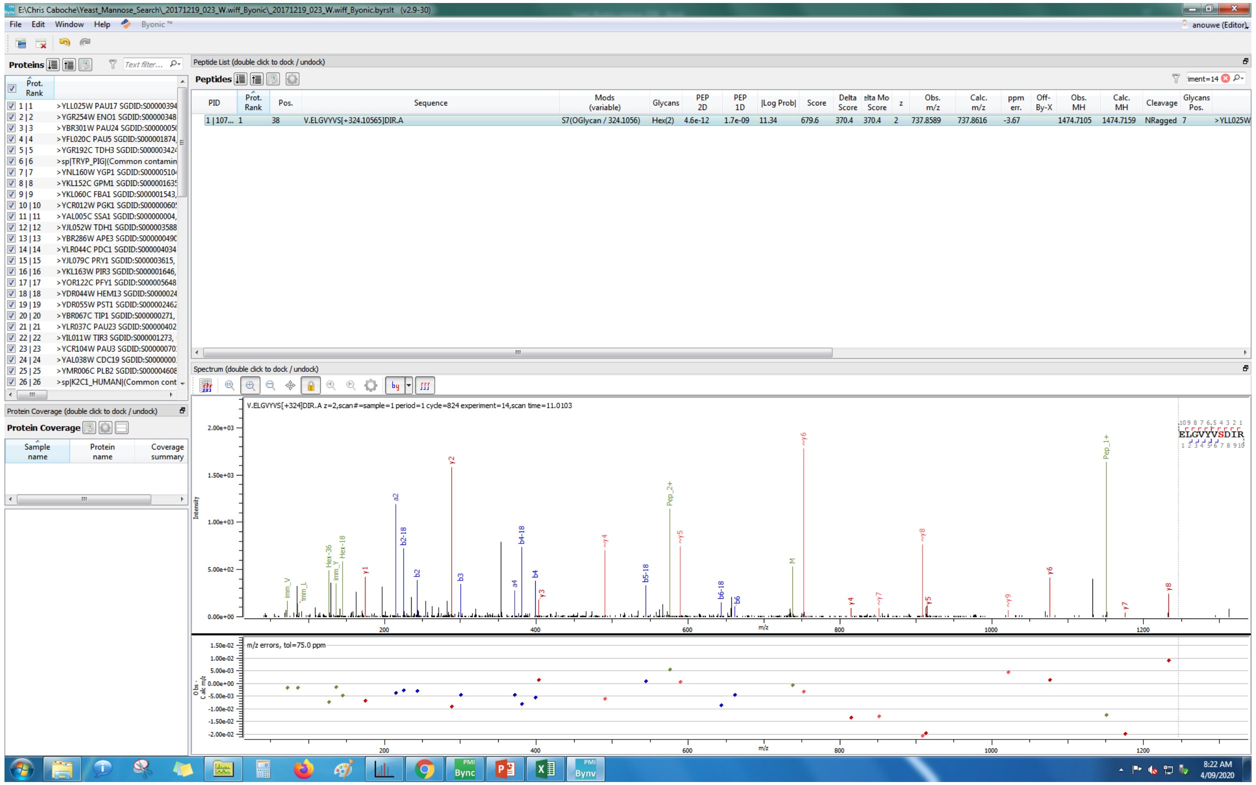
E:\Chris Caboche\Yeast_Mannose_Search_20171219_023_W.wiff_Byonic_20171219_023_W.wiff_Byonic.byrslt (v2.9-30)
File Edit Window Help Byonic™
anouwe (Editor)
Proteins
Peptide List (double click to dock / undock)
Peptides
PID | Prot. Rank | Pos. | Sequence | Mods (variable) | Glycans | PEP 2D | PEP 1D | |Log Prob| | Score | Delta Score | Delta Mod Score | z | Obs. m/z | Calc. m/z | ppm err. | Off-By-X | Obs. MH | Calc. MH | Cleavage | Glycans Pos.
1 | 107... 1 38 V.ELGVYVS[+324.10565]DIR.A S7(OGlycan / 324.1056) Hex(2) 4.6e-12 1.7e-09 11.34 679.6 370.4 370.4 2 737.8589 737.8616 -3.67 1474.7105 1474.7159 NRagged 7 >YLL025W
1 | 1 >YLL025W PAU17 SGDID:S00000394
2 | 2 >YGR254W ENO1 SGDID:S00000348
3 | 3 >YBR301W PAU24 SGDID:S0000005
4 | 4 >YFL020C PAU5 SGDID:S000001874,
5 | 5 >YGR192C TDH3 SGDID:S00000342
6 | 6 >sp|TRYP_PIG|(Common contamin
7 | 7 >YNL160W YGP1 SGDID:S00000510
8 | 8 >YKL152C GPM1 SGDID:S00000163
9 | 9 >YKL060C FBA1 SGDID:S000001543,
Spectrum (double click to dock / undock)
V.ELGVYVS[+324]DIR.A z=2,scan#=sample=1 period=1 cycle=824 experiment=14,scan time=11.0103
ELGVYVSDIR
m/z errors, tol=75.0 ppm
Protein Coverage
Sample name
Protein name
Coverage summary

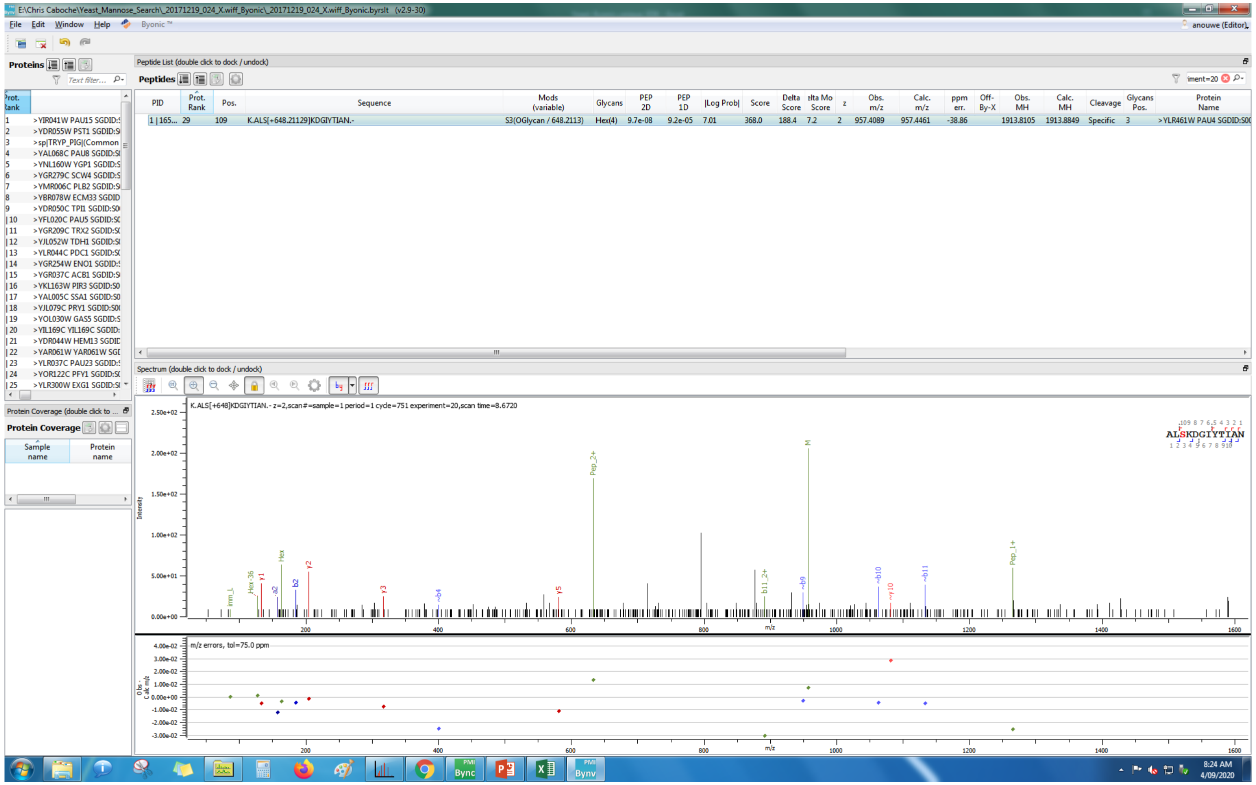
E:\Chris Caboche\Yeast_Mannose_Search_20171219_024_X.wiff_Byonic_20171219_024_X.wiff_Byonic.byrslt (v2.9-30)
File Edit Window Help Byonic ™
anouwe (Editor)
Proteins
Peptide List (double click to dock / undock)
Peptides
PID | Prot. Rank | Pos. | Sequence | Mods (variable) | Glycans | PEP 2D | PEP 1D | |Log Prob| | Score | Delta Score | Delta Mod Score | z | Obs. m/z | Calc. m/z | ppm err. | Off-By-X | Obs. MH | Calc. MH | Cleavage | Glycans Pos. | Protein Name
1 | 165... 29 109 K.ALS[+648.21129]KDGIYTIAN.- S3(OGlycan / 648.2113) Hex(4) 9.7e-08 9.2e-05 7.01 368.0 188.4 7.2 2 957.4089 957.4461 -38.86 1913.8105 1913.8849 Specific 3 >YLR461W PAU4 SGDID:S0
1 >YIR041W PAU15 SGDID
2 >YDR055W PST1 SGDID
3 >sp|TRYP_PIG|(Common
4 >YAL068C PAU8 SGDID
5 >YNL160W YGP1 SGDID
6 >YGR279C SCW4 SGDID
7 >YMR006C PLB2 SGDID
8 >YBR078W ECM33 SGDID
Spectrum (double click to dock / undock)
K.ALS[+648]KDGIYTIAN.- z=2,scan#=sample=1 period=1 cycle=751 experiment=20,scan time=8.6720
ALSKDGIYTIAN
Intensity
m/z
m/z errors, tol=75.0 ppm
8:24 AM
4/09/2020

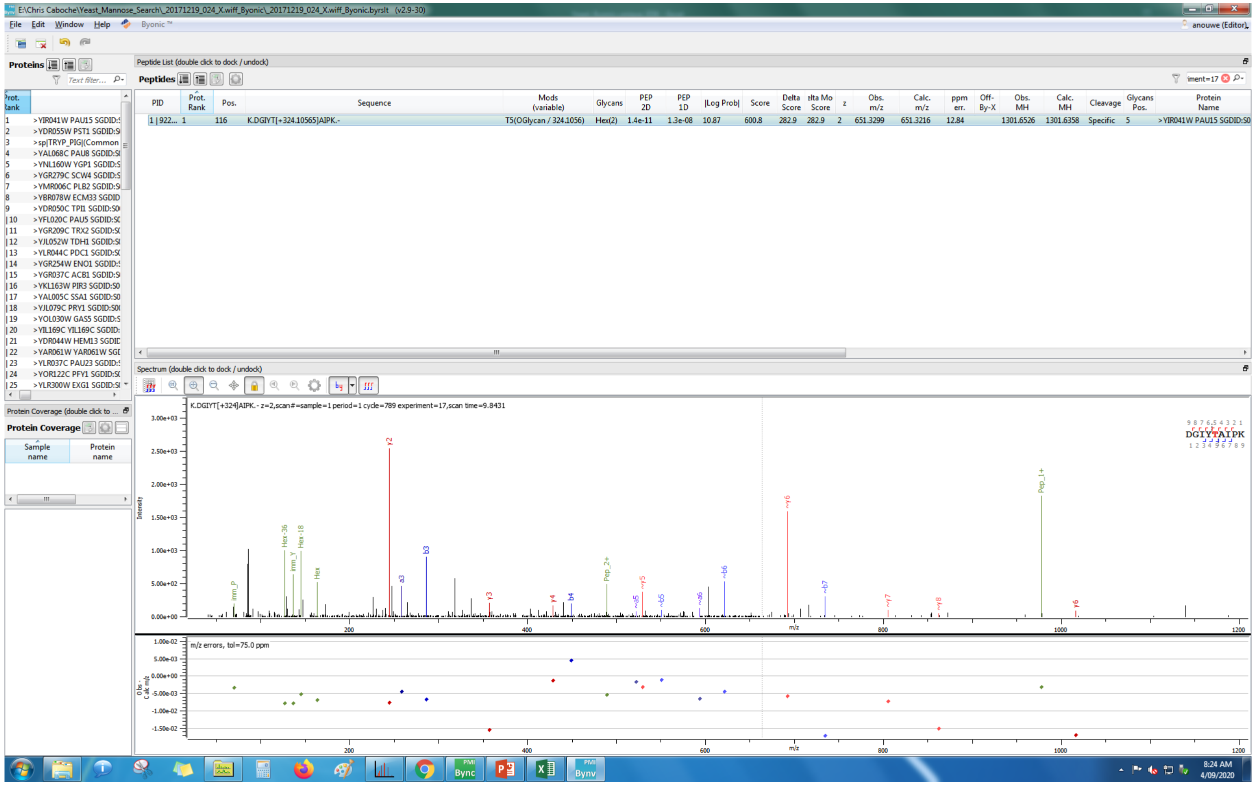

E:\Chris Caboche\Yeast_Mannose_Search_20171219_024_X.wiff_Byonic_20171219_024_X.wiff_Byonic.byrslt (v2.9-30)
File Edit Window Help Byonic ™
anouwe (Editor)
Proteins
Peptide List (double click to dock / undock)
Peptides
PID | Prot. Rank | Pos. | Sequence | Mods (variable) | Glycans | PEP 2D | PEP 1D | |Log Prob| | Score | Delta Score | elta Mo Score | z | Obs. m/z | Calc. m/z | ppm err. | Off-By-X | Obs. MH | Calc. MH | Cleavage | Glycans Pos. | Protein Name
1 | 922... | 1 | 116 | K.DGIYT[+324.10565]AIPK.- | T5(OGlycan / 324.1056) | Hex(2) | 1.4e-11 | 1.3e-08 | 10.87 | 600.8 | 282.9 | 282.9 | 2 | 651.3299 | 651.3216 | 12.84 | | 1301.6526 | 1301.6358 | Specific | 5 | >YIR041W PAU15 SGDID:S0
1 >YIR041W PAU15 SGDID:
2 >YDR055W PST1 SGDID:
3 >sp|TRYP_PIG|(Common
4 >YAL068C PAU8 SGDID:
5 >YNL160W YGP1 SGDID:
6 >YGR279C SCW4 SGDID:
7 >YMR006C PLB2 SGDID:
8 >YBR078W ECM33 SGDID
Spectrum (double click to dock / undock)
K.DGIYT[+324]AIPK.- z=2,scan#=sample=1 period=1 cycle=789 experiment=17,scan time=9.8431
DGIYTAIPK
Protein Coverage
Sample name
Protein name
m/z errors, tol=75.0 ppm
8:24 AM
4/09/2020

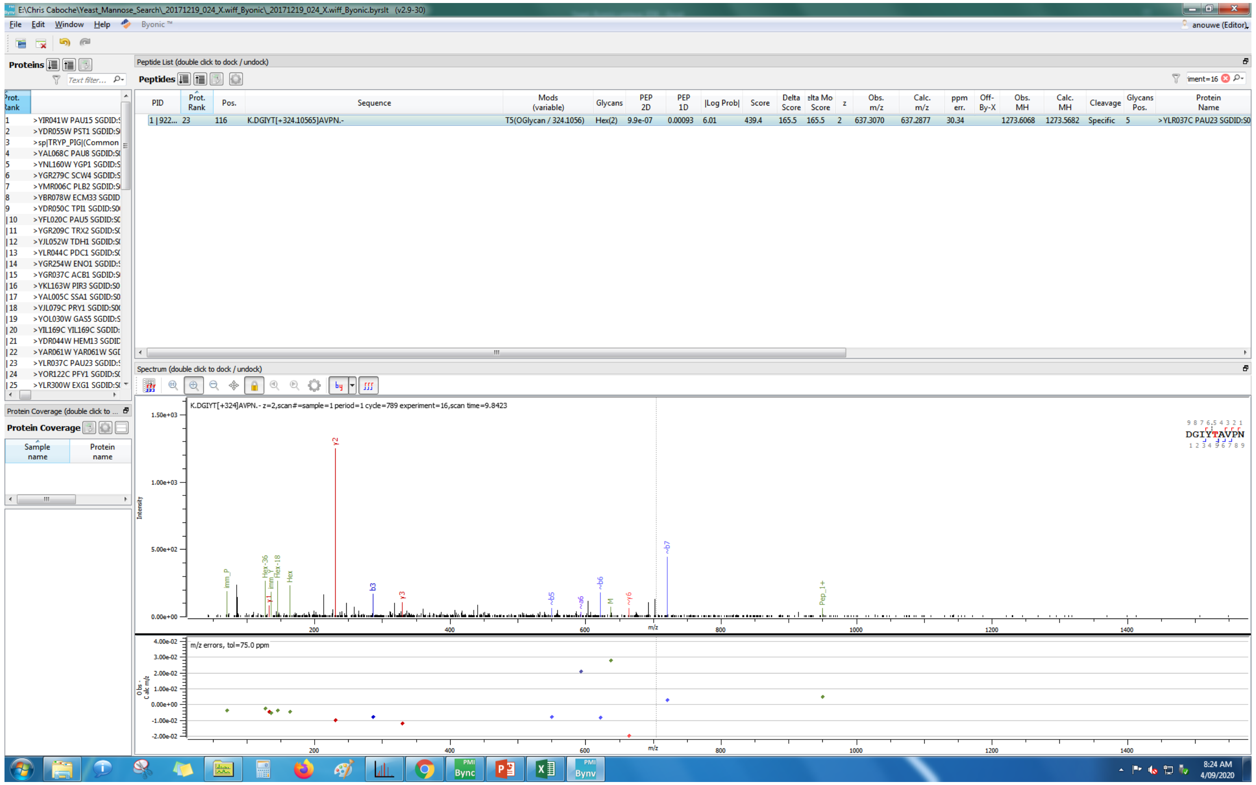
E:\Chris Caboche\Yeast_Mannose_Search_20171219_024_X.wiff_Byonic_20171219_024_X.wiff_Byonic.byrslt (v2.9-30)
File Edit Window Help Byonic™
anouwe (Editor)
Proteins
Peptide List (double click to dock / undock)
Peptides
PID | Prot. Rank | Pos. | Sequence | Mods (variable) | Glycans | PEP 2D | PEP 1D | |Log Prob| | Score | Delta Score | elta Mo Score | z | Obs. m/z | Calc. m/z | ppm err. | Off-By-X | Obs. MH | Calc. MH | Cleavage | Glycans Pos. | Protein Name
1 | 922... 23 116 K.DGIYT[+324.10565]AVPN.- T5(OGlycan / 324.1056) Hex(2) 9.9e-07 0.00093 6.01 439.4 165.5 165.5 2 637.3070 637.2877 30.34 1273.6068 1273.5682 Specific 5 >YLR037C PAU23 SGDID:S0
Spectrum (double click to dock / undock)
K.DGIYT[+324]AVPN.- z=2,scan#=sample=1 period=1 cycle=789 experiment=16,scan time=9.8423
DGIYTAVPN
Protein Coverage
Sample name
Protein name
m/z errors, tol=75.0 ppm
8:24 AM
4/09/2020

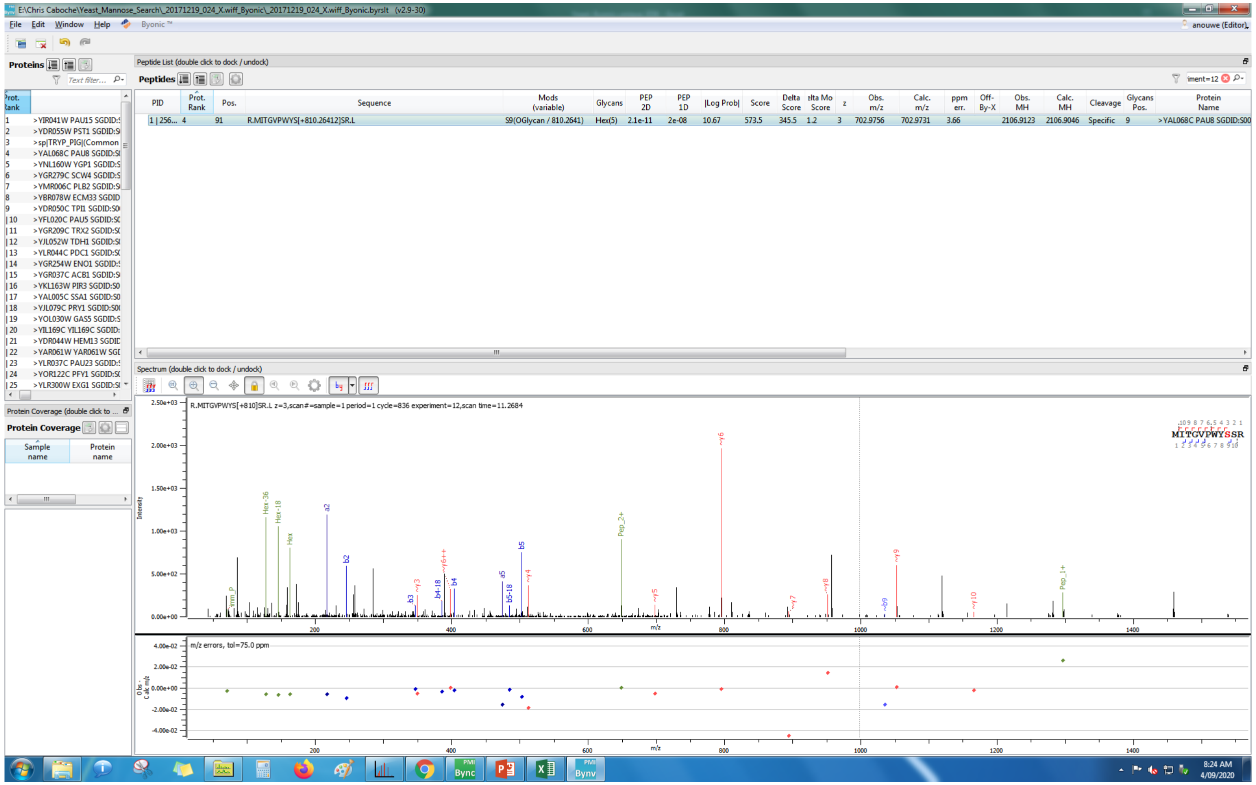
E:\Chris Caboche\Yeast_Mannose_Search_20171219_024_X.wiff_Byonic_20171219_024_X.wiff_Byonic.byrslt (v2.9-30)
Peptide List (double click to dock / undock)
R.MITGVPWYS[+810.26412]SR.L
S9(OGlycan / 810.2641)
Hex(5)
R.MITGVPWYS[+810]SR.L z=3,scan#=sample=1 period=1 cycle=836 experiment=12,scan time=11.2684
m/z errors, tol=75.0 ppm

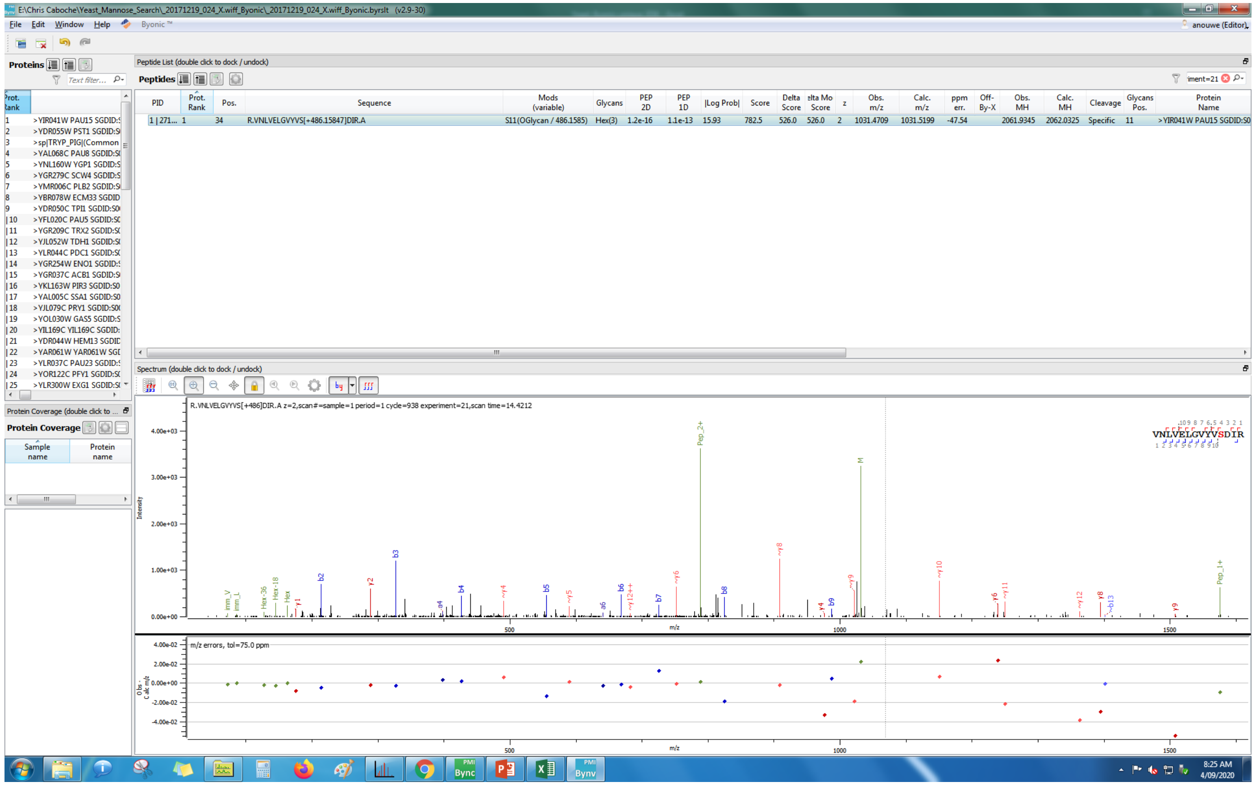
E:\Chris Caboche\Yeast_Mannose_Search_20171219_024_X.wiff_Byonic_20171219_024_X.wiff_Byonic.byrslt (v2.9-30)
File Edit Window Help Byonic™
anouwe (Editor)
Proteins
Peptide List (double click to dock / undock)
Peptides
PID Prot. Rank Pos. Sequence Mods (variable) Glycans PEP 2D PEP 1D |Log Prob| Score Delta Score Delta Mod Score z Obs. m/z Calc. m/z ppm err. Off-By-X Obs. MH Calc. MH Cleavage Glycans Pos. Protein Name
1 | 271... 1 34 R.VNLVELGVYVS[+486.15847]DIR.A S11(OGlycan / 486.1585) Hex(3) 1.2e-16 1.1e-13 15.93 782.5 526.0 526.0 2 1031.4709 1031.5199 -47.54 2061.9345 2062.0325 Specific 11 >YIR041W PAU15 SGDID:S0
Spectrum (double click to dock / undock)
R.VNLVELGVYVS[+486]DIR.A z=2,scan#=sample=1 period=1 cycle=938 experiment=21,scan time=14.4212
VNLVELGVYVSDIR
Protein Coverage
m/z errors, tol=75.0 ppm
8:25 AM
4/09/2020
