## Supplementary Material S2 for "The post-translational modification landscape of commercial beers"

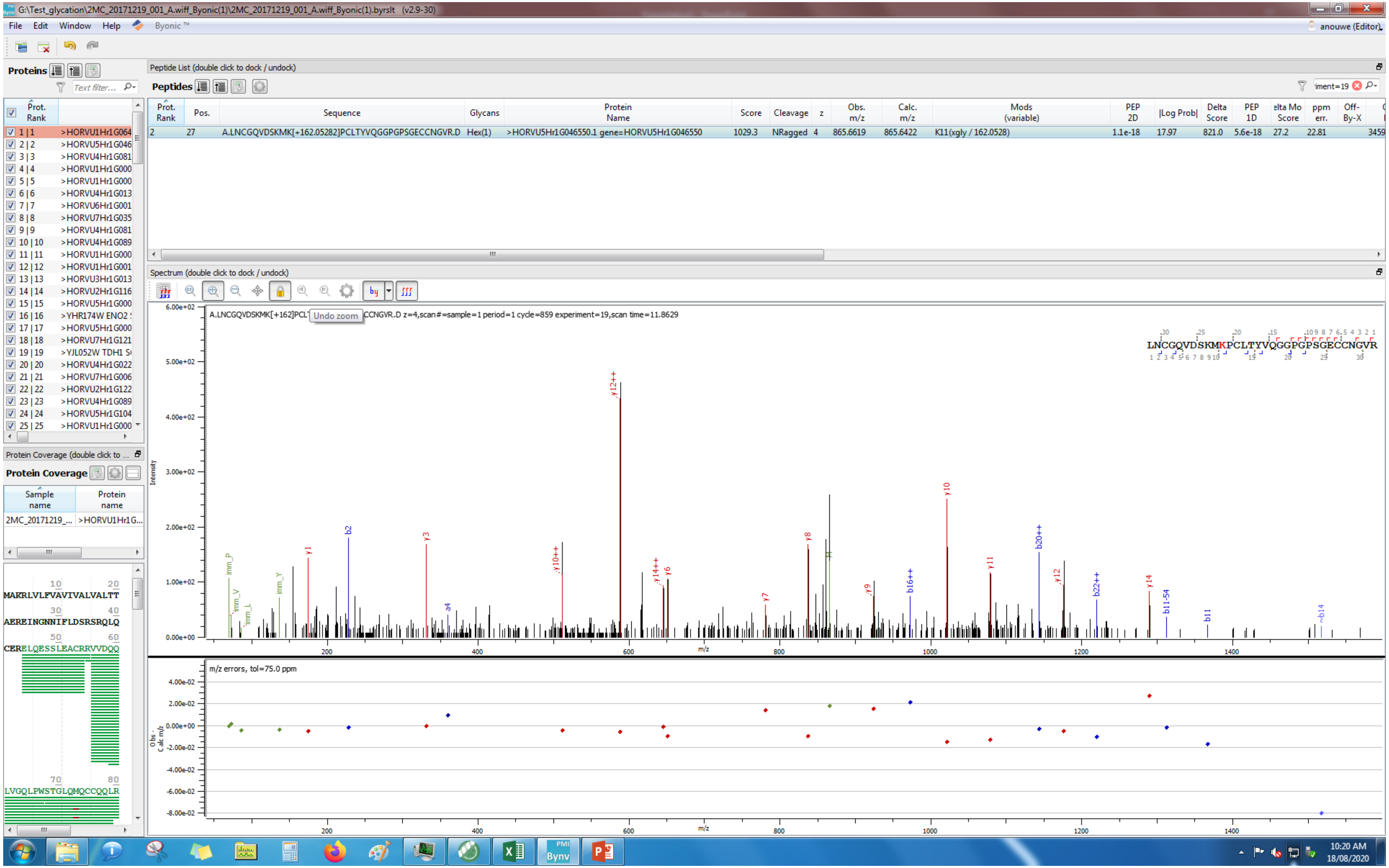
G:\Test_glycation\2MC_20171219_001_A.wiff_Byonic(1)\2MC_20171219_001_A.wiff_Byonic(1).byrslt (v2.9-30)
File Edit Window Help Byonic™
anouwe (Editor)
Proteins
Peptide List (double click to dock / undock)
Peptides
Prot. Rank | Pos. | Sequence | Glycans | Protein Name | Score | Cleavage | z | Obs. m/z | Calc. m/z | Mods (variable) | PEP 2D | |Log Prob| | Delta Score | PEP 1D | ppm err. | Off-By-X
2 | 27 | A.LNCGQVDSKMK[+162.05282]PCLTYVQGGPGPSGECCNGVR.D | Hex(1) | >HORVU5Hr1G046550.1 gene=HORVU5Hr1G046550 | 1029.3 | NRagged | 4 | 865.6619 | 865.6422 | K11(xgly / 162.0528) | 1.1e-18 | 17.97 | 821.0 | 5.6e-18 | 27.2 | 22.81 | 3459
Spectrum (double click to dock / undock)
A.LNCGQVDSKMK[+162]PCL Undo zoom CCNGVR.D z=4,scan#=sample=1 period=1 cycle=859 experiment=19,scan time=11.8629
LNCGQVDSKMKPCLTYVQGGPGPSGECCNGVR
Intensity
m/z
m/z errors, tol=75.0 ppm
Protein Coverage (double click to ...
Protein Coverage
Sample name | Protein name
2MC_20171219_... | >HORVU1Hr1G...
MAKRLVLPVAVIVALVALTT
AEREINGNNIFLDSRSRQLQ
CERELQESSLEACRRVVDQQ
LVGQLPWSTGLQMQCCQQLR
10:20 AM
18/08/2020

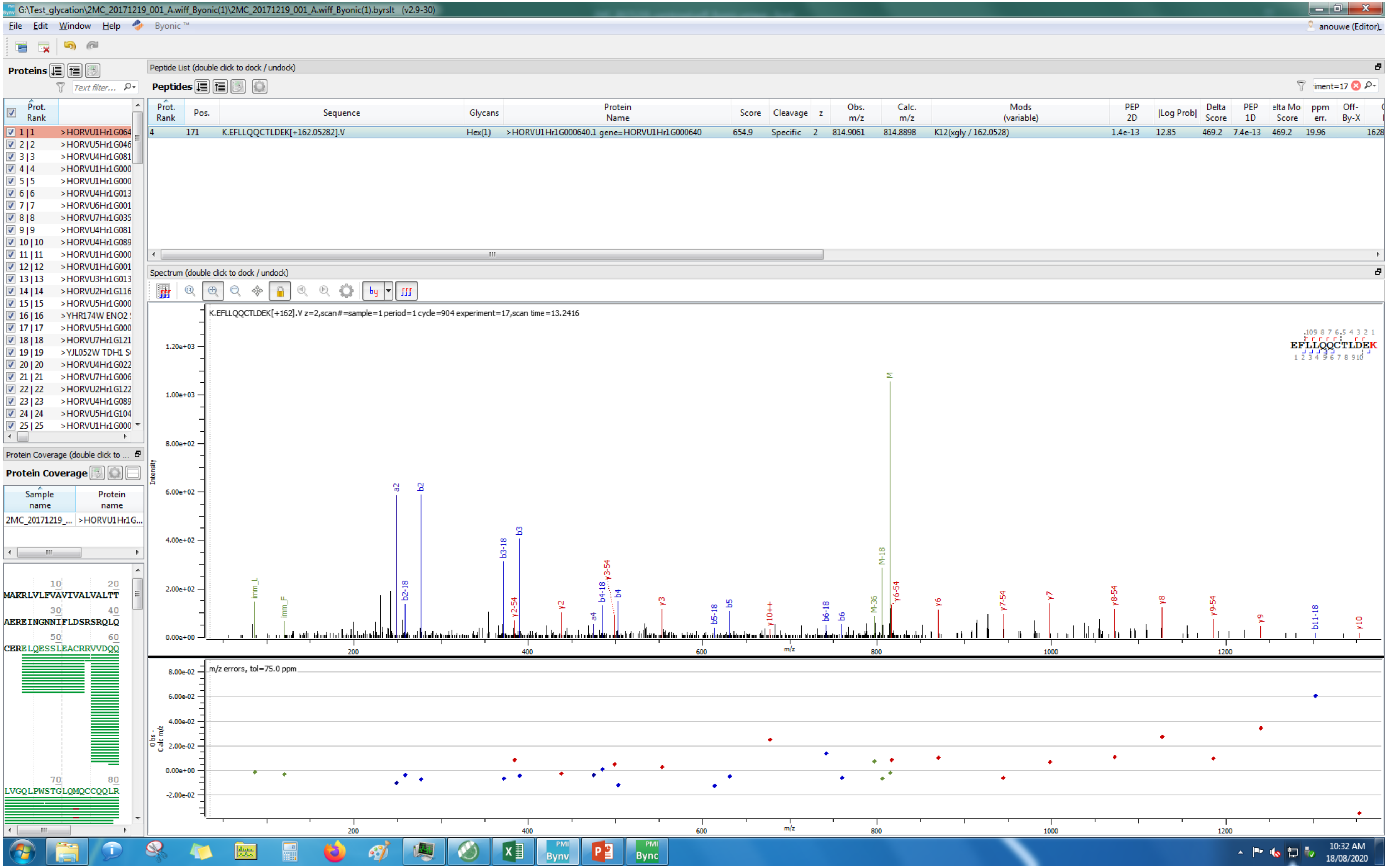
G:\Test_glycation\2MC_20171219_001_A.wiff_Byonic(1)\2MC_20171219_001_A.wiff_Byonic(1).byrslt (v2.9-30)
File Edit Window Help Byonic ™
anouwe (Editor)
Proteins
Peptide List (double click to dock / undock)
Peptides
Prot. Rank | Pos. | Sequence | Glycans | Protein Name | Score | Cleavage | z | Obs. m/z | Calc. m/z | Mods (variable) | PEP 2D | |Log Prob| | Delta Score | PEP 1D | Delta Mod Score | ppm err. | Off-By-X
4 | 171 | K.EFLLQQCTLDEK[+162.05282].V | Hex(1) | >HORVU1Hr1G000640.1 gene=HORVU1Hr1G000640 | 654.9 | Specific | 2 | 814.9061 | 814.8898 | K12(xgly / 162.0528) | 1.4e-13 | 12.85 | 469.2 | 7.4e-13 | 469.2 | 19.96 | 1628
Spectrum (double click to dock / undock)
K.EFLLQQCTLDEK[+162].V z=2,scan#=sample=1 period=1 cycle=904 experiment=17,scan time=13.2416
EFLLQQCTLDEK
Intensity
m/z
m/z errors, tol=75.0 ppm
Protein Coverage (double click to ...
Protein Coverage
Sample name
Protein name
2MC_20171219_...
>HORVU1Hr1G...
MAKRLVLFVAVIVALVALTT
AERE INGNNIFLDSRSRQLQ
CERELQESSLEACRRVVDQQ
LVGQLPWSTGLQMQCCQQLR
10:32 AM
18/08/2020

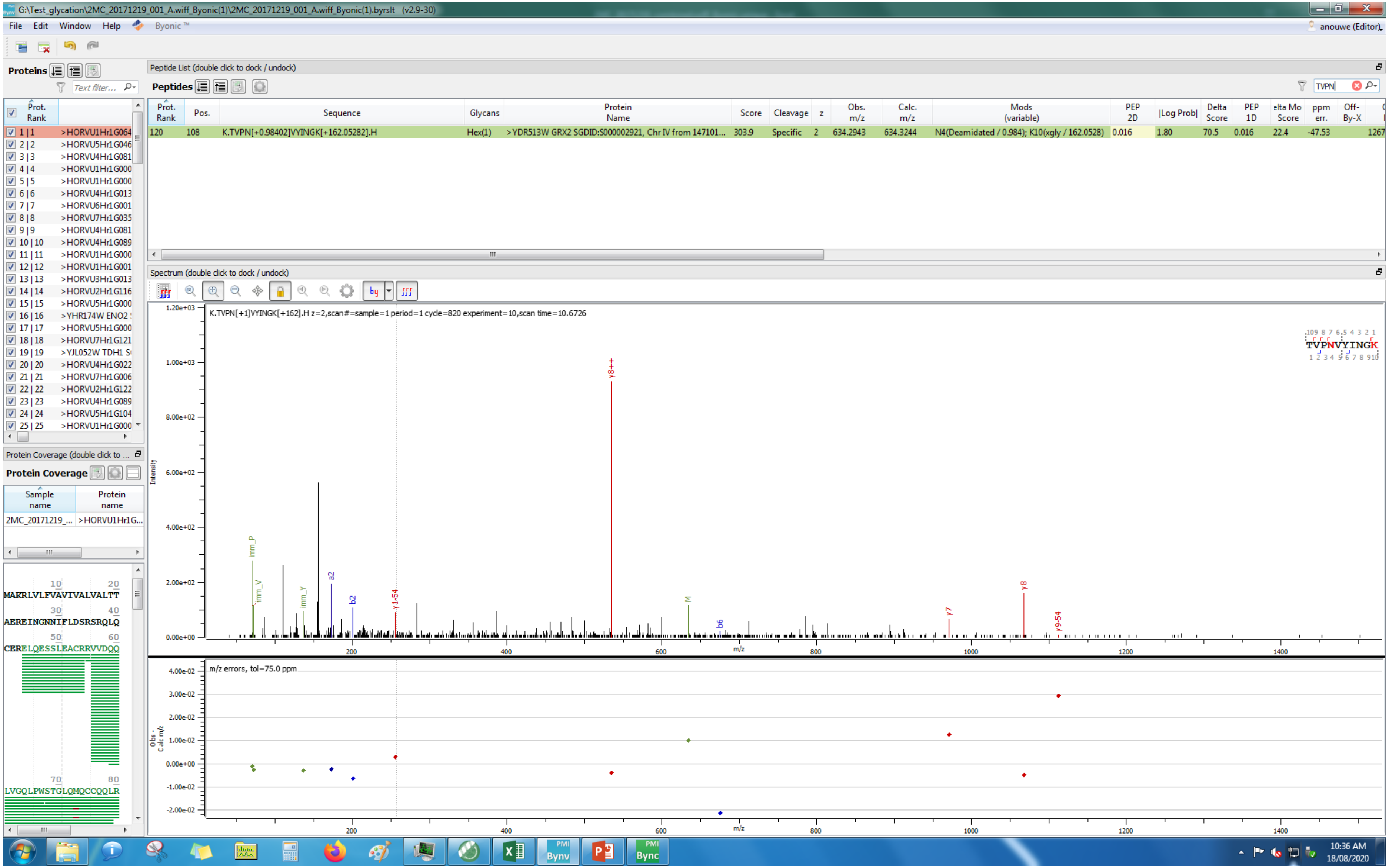
G:\Test_glycation\2MC_20171219_001_A.wiff_Byonic(1)\2MC_20171219_001_A.wiff_Byonic(1).byrslt (v2.9-30)
File Edit Window Help Byonic™
anouwe (Editor)
Proteins
Peptide List (double click to dock / undock)
Peptides
TVPN
Prot. Rank | Pos. | Sequence | Glycans | Protein Name | Score | Cleavage | z | Obs. m/z | Calc. m/z | Mods (variable) | PEP 2D | |Log Prob| | Delta Score | PEP 1D | Delta Mod Score | ppm err. | Off-By-X
120 | 108 | K.TVPN[+0.98402]VYINGK[+162.05282].H | Hex(1) | >YDR513W GRX2 SGDID:S000002921, Chr IV from 147101... | 303.9 | Specific | 2 | 634.2943 | 634.3244 | N4(Deamidated / 0.984); K10(xgly / 162.0528) | 0.016 | 1.80 | 70.5 | 0.016 | 22.4 | -47.53 | 1267
Spectrum (double click to dock / undock)
K.TVPN[+1]VYINGK[+162].H z=2,scan#=sample=1 period=1 cycle=820 experiment=10,scan time=10.6726
TVPNVYINGK
Intensity
m/z
y8++
y8
y7
y9-54
y1-54
imm_P
imm_V
imm_Y
a2
b2
b6
M
m/z errors, tol=75.0 ppm
Obs - Calc m/z
Protein Coverage (double click to ...
Protein Coverage
Sample name | Protein name
2MC_20171219_... | >HORVU1Hr1G...
MAKRLVLFVAVIVALVALTT
AEREINGNNIFLDSRSRQLQ
CERELQESSLEACRRVVDQQ
LVGQLPWSTGLQMQCCQQLR
10:36 AM
18/08/2020

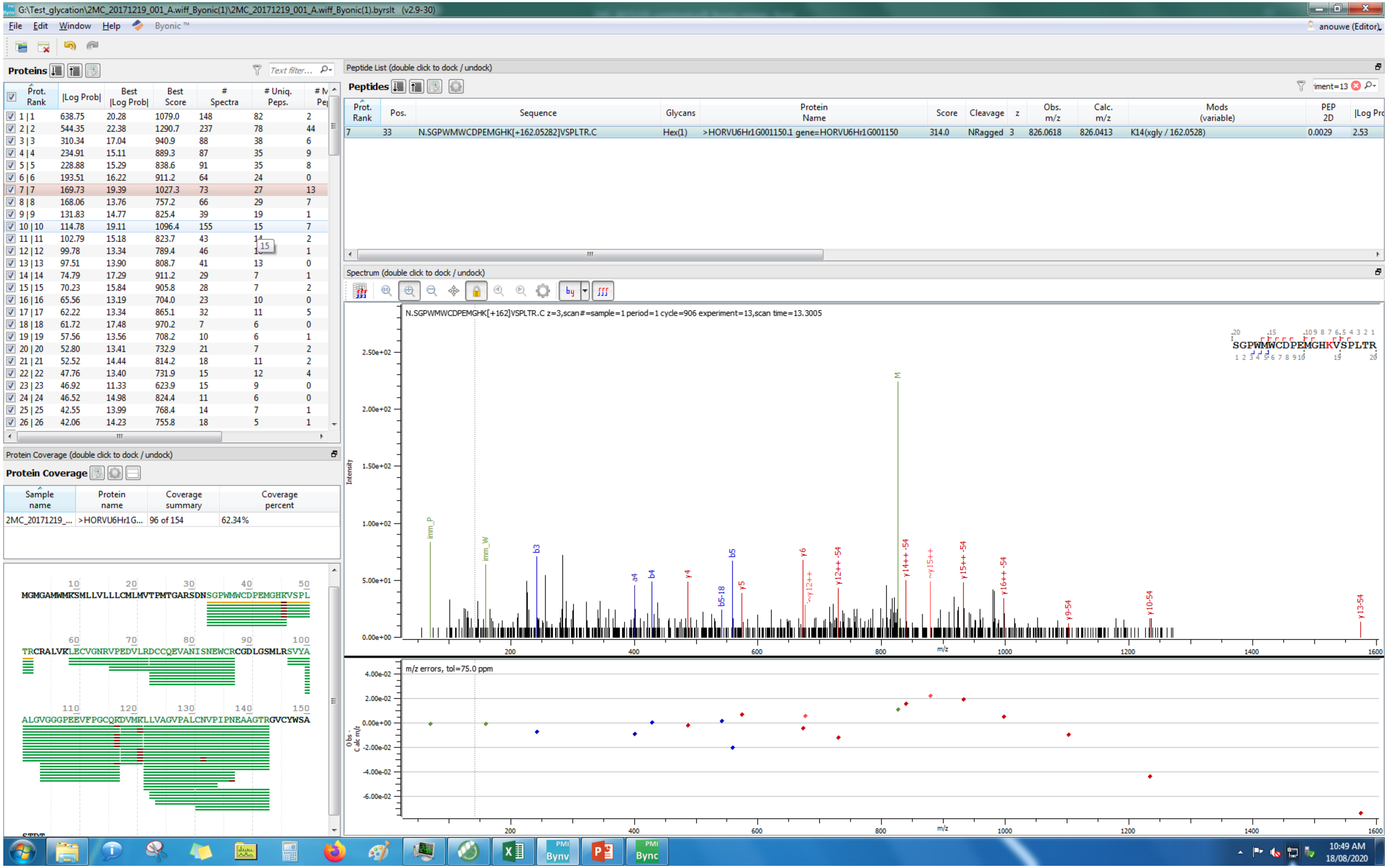
G:\Test_glycation\2MC_20171219_001_A.wiff_Byonic(1)\2MC_20171219_001_A.wiff_Byonic(1).byrslt (v2.9-30)
File Edit Window Help Byonic™
anouwe (Editor)
Proteins
Peptide List (double click to dock / undock)
Peptides
N.SGPWMWCDPEMGHK[+162.05282]VSPLTR.C
Hex(1)
>HORVU6Hr1G001150.1 gene=HORVU6Hr1G001150
314.0
NRagged
826.0618
826.0413
K14(xgly / 162.0528)
0.0029
2.53
Spectrum (double click to dock / undock)
N.SGPWMWCDPEMGHK[+162]VSPLTR.C z=3,scan#=sample=1 period=1 cycle=906 experiment=13,scan time=13.3005
SGPWMWCDPEMGHKVSPLTR
Intensity
m/z
m/z errors, tol=75.0 ppm
Protein Coverage (double click to dock / undock)
Protein Coverage
Sample name
Protein name
Coverage summary
Coverage percent
2MC_20171219_...
>HORVU6Hr1G...
96 of 154
62.34%
MGMGAMWMKSMLLVLLLCMLMVTPMTGARSDNSGPWMWCDPEMGHKVSPL
TRCRALVKLECVGNRVPEDVLRDCCQEVANISNEWCRCGDLGSMLRSVYA
ALGVGGGPEEVFPGCQKDVMKLLVAGVPALCNVPIPNEAAGTRGVCYWSA
10:49 AM
18/08/2020

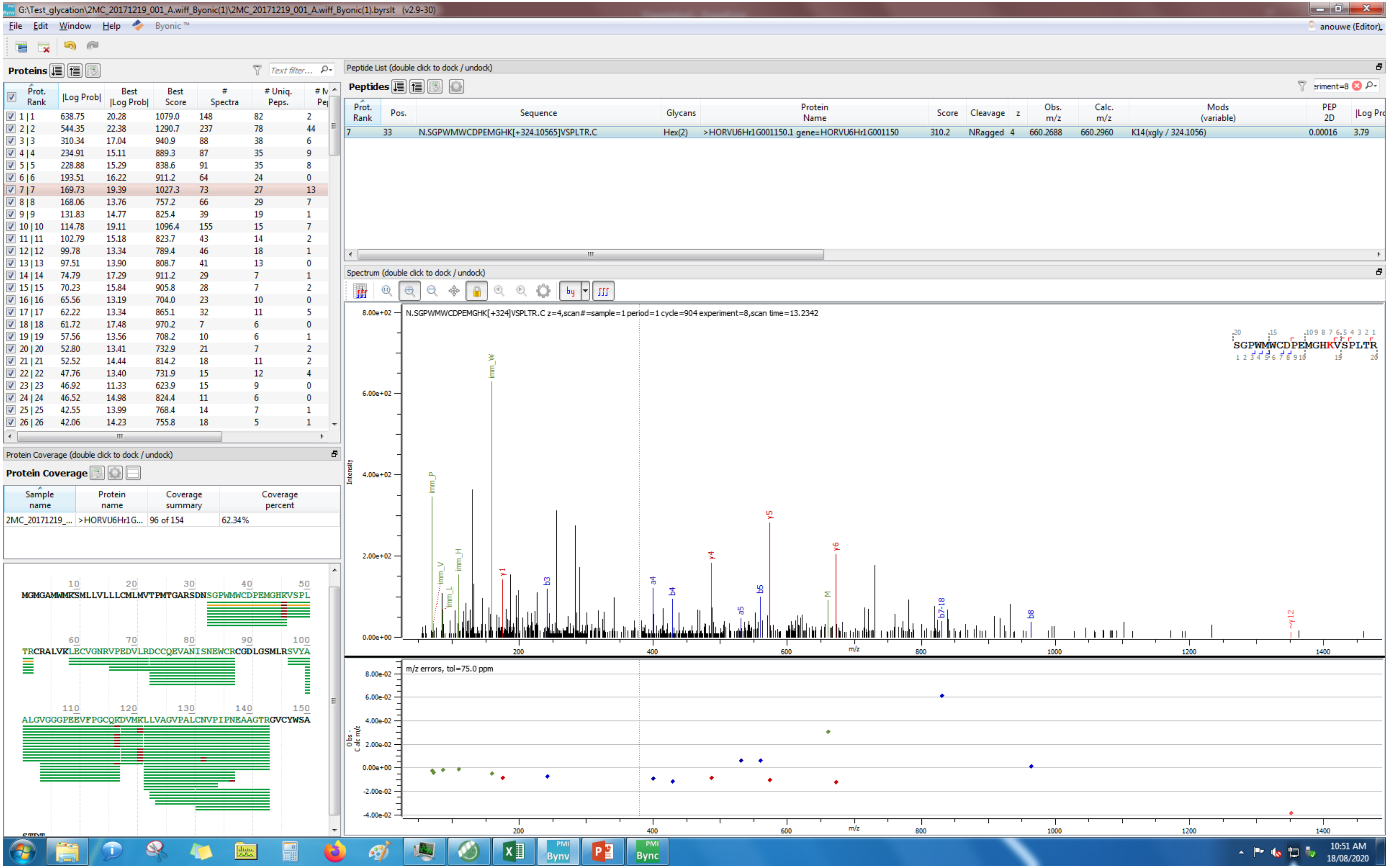
G:\Test_glycation\2MC_20171219_001_A.wiff_Byonic(1)\2MC_20171219_001_A.wiff_Byonic(1).byrslt (v2.9-30)
File Edit Window Help Byonic
Proteins
Peptide List (double click to dock / undock)
Peptides
7 33 N.SGPWMWCDPEMGHK[+324.10565]VSPLTR.C Hex(2) >HORVU6Hr1G001150.1 gene=HORVU6Hr1G001150 310.2 NRagged 4 660.2688 660.2960 K14(xgly / 324.1056) 0.00016 3.79
Spectrum (double click to dock / undock)
N.SGPWMWCDPEMGHK[+324]VSPLTR.C z=4,scan#=sample=1 period=1 cycle=904 experiment=8,scan time=13.2342
SGPWMWCDPEMGHKVSPLTR
m/z errors, tol=75.0 ppm
Protein Coverage (double click to dock / undock)
Protein Coverage
2MC_20171219_... >HORVU6Hr1G... 96 of 154 62.34%
MGMGAMWMKSMLLVLLLCMLMVTPMTGARSDNSGPWMWCDPEMGHKVSPL
TRCRALVKLECVGNRVPEDVLRDCCQEVANISNEWCRCGDLGSMLRSVYA
ALGVGGGPEEVFPGCQKDVMKLLVAGVPALCNVPIPNEAAGTRGVCYWSA
10:51 AM
18/08/2020

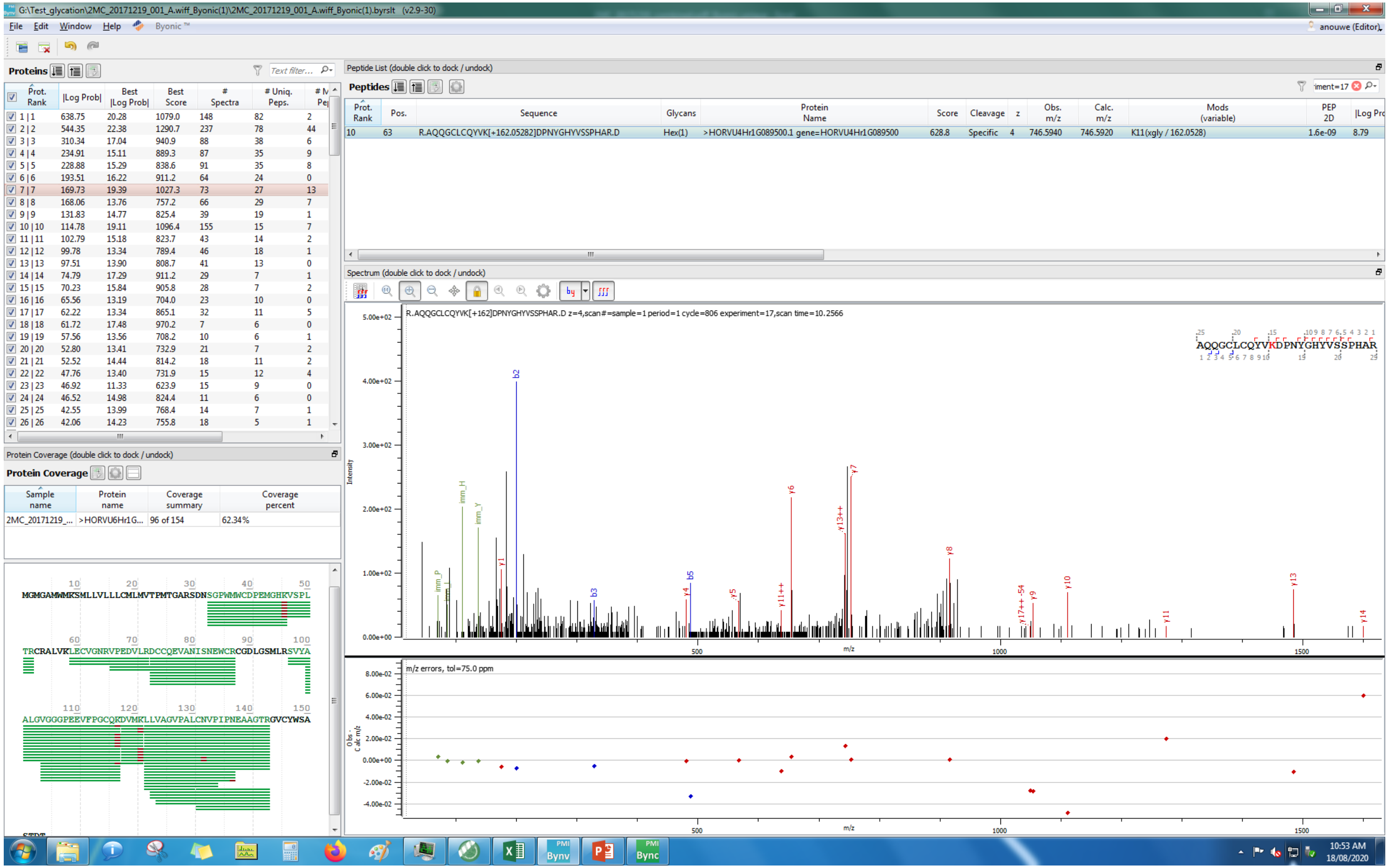

G:\Test_glycation\2MC_20171219_001_A.wiff_Byonic(1)\2MC_20171219_001_A.wiff_Byonic(1).byrslt (v2.9-30)
File Edit Window Help Byonic™
anouwe (Editor)
Proteins
Prot. Rank | |Log Prob| | Best |Log Prob| | Best Score | # Spectra | # Uniq. Peps.
1 | 1 638.75 20.28 1079.0 148 82 2
2 | 2 544.35 22.38 1290.7 237 78 44
3 | 3 310.34 17.04 940.9 88 38 6
4 | 4 234.91 15.11 889.3 87 35 9
5 | 5 228.88 15.29 838.6 91 35 8
6 | 6 193.51 16.22 911.2 64 24 0
7 | 7 169.73 19.39 1027.3 73 27 13
8 | 8 168.06 13.76 757.2 66 29 7
9 | 9 131.83 14.77 825.4 39 19 1
10 | 10 114.78 19.11 1096.4 155 15 7
11 | 11 102.79 15.18 823.7 43 14 2
12 | 12 99.78 13.34 789.4 46 18 1
13 | 13 97.51 13.90 808.7 41 13 0
14 | 14 74.79 17.29 911.2 29 7 1
15 | 15 70.23 15.84 905.8 28 7 2
16 | 16 65.56 13.19 704.0 23 10 0
17 | 17 62.22 13.34 865.1 32 11 5
18 | 18 61.72 17.48 970.2 7 6 0
19 | 19 57.56 13.56 708.2 10 6 1
20 | 20 52.80 13.41 732.9 21 7 2
21 | 21 52.52 14.44 814.2 18 11 2
22 | 22 47.76 13.40 731.9 15 12 4
23 | 23 46.92 11.33 623.9 15 9 0
24 | 24 46.52 14.98 824.4 11 6 0
25 | 25 42.55 13.99 768.4 14 7 1
26 | 26 42.06 14.23 755.8 18 5 1
Protein Coverage (double click to dock / undock)
Protein Coverage
Sample name | Protein name | Coverage summary | Coverage percent
2MC_20171219_... >HORVU6Hr1G... 96 of 154 62.34%
MGMGAMWMKSMLLVLLLCMLMVTPMTGARSDNSGPWMWCDPEMGHKVSPL
TRCRALVKLECVGNRVPEDVLRDCCQEVANISNEWCRCGDLGSMLRSVYA
ALGVGGGPEEVFPGCQKDVMKLLVAGVPALCNVPIPNEAAGTRGVCYWSA
Peptide List (double click to dock / undock)
Peptides
Prot. Rank | Pos. | Sequence | Glycans | Protein Name | Score | Cleavage | z | Obs. m/z | Calc. m/z | Mods (variable) | PEP 2D | |Log Prob|
10 63 R.AQQGCLCQYVK[+162.05282]DPNYGHYVSSPHAR.D Hex(1) >HORVU4Hr1G089500.1 gene=HORVU4Hr1G089500 628.8 Specific 4 746.5940 746.5920 K11(xgly / 162.0528) 1.6e-09 8.79
Spectrum (double click to dock / undock)
R.AQQGCLCQYVK[+162]DPNYGHYVSSPHAR.D z=4,scan#=sample=1 period=1 cycle=806 experiment=17,scan time=10.2566
AQQGCLCQYVKDPNYGHYVSSPHAR
Intensity
m/z
m/z errors, tol=75.0 ppm
10:53 AM
18/08/2020

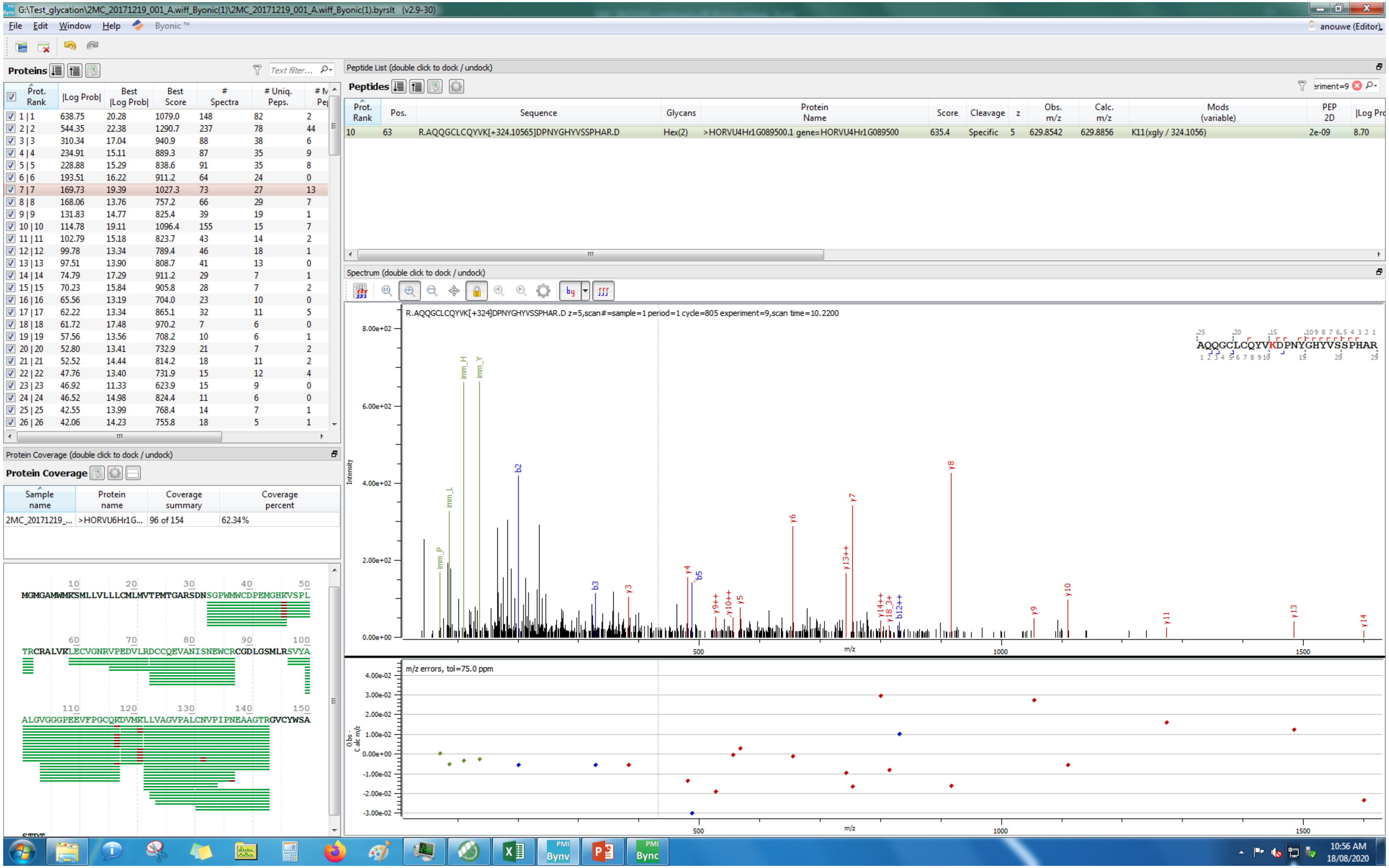
G:\Test_glycation\2MC_20171219_001_A.wiff_Byonic(1)\2MC_20171219_001_A.wiff_Byonic(1).byrslt (v2.9-30)
File Edit Window Help Byonic™
anouwe (Editor)
Proteins
Peptide List (double click to dock / undock)
Peptides
Prot. Rank | Pos. | Sequence | Glycans | Protein Name | Score | Cleavage | z | Obs. m/z | Calc. m/z | Mods (variable) | PEP 2D | |Log Prob|
10 | 63 | R.AQQGCLCQYVK[+324.10565]DPNYGHYVSSPHAR.D | Hex(2) | >HORVU4Hr1G089500.1 gene=HORVU4Hr1G089500 | 635.4 | Specific | 5 | 629.8542 | 629.8856 | K11(xgly / 324.1056) | 2e-09 | 8.70
Spectrum (double click to dock / undock)
R.AQQGCLCQYVK[+324]DPNYGHYVSSPHAR.D z=5,scan#=sample=1 period=1 cycle=805 experiment=9,scan time=10.2200
AQQGCLCQYVKDPNYGHYVSSPHAR
m/z errors, tol=75.0 ppm
Protein Coverage (double click to dock / undock)
Protein Coverage
Sample name | Protein name | Coverage summary | Coverage percent
2MC_20171219_... | >HORVU6Hr1G... | 96 of 154 | 62.34%
MGMGAMWMKSMLLVLLLCMLMVTPMTGARSDNSGPWMWCDPEMGHKVSPL
TRCRALVKLECVGNRVPEDVLRDCCQEVANISNEWCRCGDLGSMLRSVYA
ALGVGGPEEVFPGCQKDVMKLLVAGVPALCNVPIPNEAAGTRGVCYWSA
10:56 AM
18/08/2020

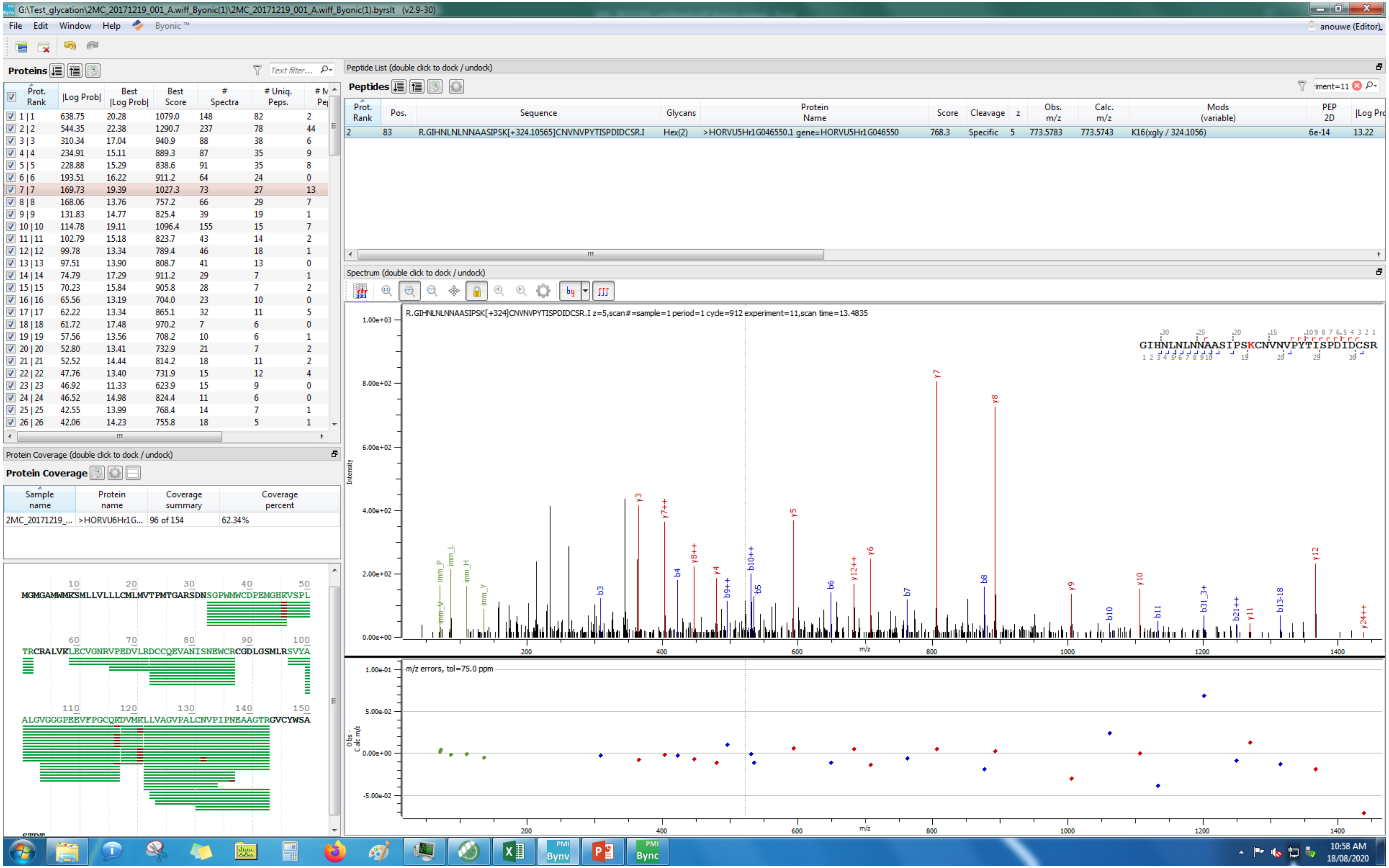


| Prot. Rank | \|Log Prob\| | Best \|Log Prob\| | Best Score | # Spectra | # Uniq. Peps. | # M Pe |
|---|---|---|---|---|---|---|
| 1 \| 1 | 638.75 | 20.28 | 1079.0 | 148 | 82 | 2 |
| 2 \| 2 | 544.35 | 22.38 | 1290.7 | 237 | 78 | 44 |
| 3 \| 3 | 310.34 | 17.04 | 940.9 | 88 | 38 | 6 |
| 4 \| 4 | 234.91 | 15.11 | 889.3 | 87 | 35 | 9 |
| 5 \| 5 | 228.88 | 15.29 | 838.6 | 91 | 35 | 8 |
| 6 \| 6 | 193.51 | 16.22 | 911.2 | 64 | 24 | 0 |
| 7 \| 7 | 169.73 | 19.39 | 1027.3 | 73 | 27 | 13 |
| 8 \| 8 | 168.06 | 13.76 | 757.2 | 66 | 29 | 7 |
| 9 \| 9 | 131.83 | 14.77 | 825.4 | 39 | 19 | 1 |
| 10 \| 10 | 114.78 | 19.11 | 1096.4 | 155 | 15 | 7 |
| 11 \| 11 | 102.79 | 15.18 | 823.7 | 43 | 14 | 2 |
| 12 \| 12 | 99.78 | 13.34 | 789.4 | 46 | 18 | 1 |
| 13 \| 13 | 97.51 | 13.90 | 808.7 | 41 | 13 | 0 |
| 14 \| 14 | 74.79 | 17.29 | 911.2 | 29 | 7 | 1 |
| 15 \| 15 | 70.23 | 15.84 | 905.8 | 28 | 7 | 2 |
| 16 \| 16 | 65.56 | 13.19 | 704.0 | 23 | 10 | 0 |
| 17 \| 17 | 62.22 | 13.34 | 865.1 | 32 | 11 | 5 |
| 18 \| 18 | 61.72 | 17.48 | 970.2 | 7 | 6 | 0 |
| 19 \| 19 | 57.56 | 13.56 | 708.2 | 10 | 6 | 1 |
| 20 \| 20 | 52.80 | 13.41 | 732.9 | 21 | 7 | 2 |
| 21 \| 21 | 52.52 | 14.44 | 814.2 | 18 | 11 | 2 |
| 22 \| 22 | 47.76 | 13.40 | 731.9 | 15 | 12 | 4 |
| 23 \| 23 | 46.92 | 11.33 | 623.9 | 15 | 9 | 0 |
| 24 \| 24 | 46.52 | 14.98 | 824.4 | 11 | 6 | 0 |
| 25 \| 25 | 42.55 | 13.99 | 768.4 | 14 | 7 | 1 |
| 26 \| 26 | 42.06 | 14.23 | 755.8 | 18 | 5 | 1 |

| Prot. Rank | Pos. | Sequence | Glycans | Protein Name | Score | Cleavage | z | Obs. m/z | Calc. m/z | Mods (variable) | PEP 2D | \|Log Pro |
|---|---|---|---|---|---|---|---|---|---|---|---|---|
| 2 | 83 | R.GIHNLNLNNAASIPSK[+324.10565]CNVNVPYTISPDIDCSR.I | Hex(2) | >HORVU5Hr1G046550.1 gene=HORVU5Hr1G046550 | 768.3 | Specific | 5 | 773.5783 | 773.5743 | K16(xgly / 324.1056) | 6e-14 | 13.22 |

| Sample name | Protein name | Coverage summary | Coverage percent |
|---|---|---|---|
| 2MC_20171219_... | >HORVU6Hr1G... | 96 of 154 | 62.34% |

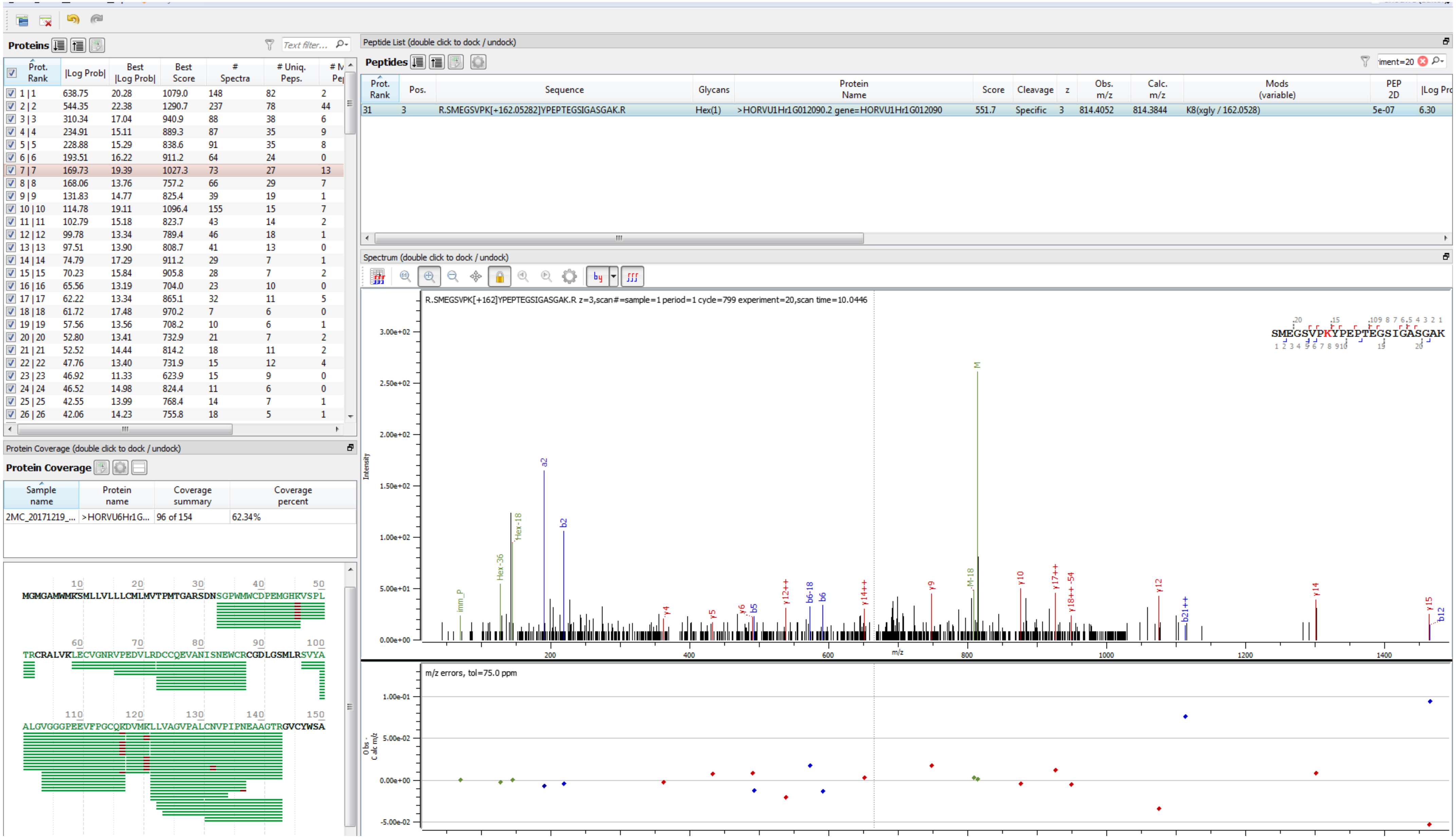

Proteins
Text filter...
Prot. Rank | |Log Prob| | Best |Log Prob| | Best Score | # Spectra | # Uniq. Peps.
1 | 1 638.75 20.28 1079.0 148 82 2
2 | 2 544.35 22.38 1290.7 237 78 44
3 | 3 310.34 17.04 940.9 88 38 6
4 | 4 234.91 15.11 889.3 87 35 9
5 | 5 228.88 15.29 838.6 91 35 8
6 | 6 193.51 16.22 911.2 64 24 0
7 | 7 169.73 19.39 1027.3 73 27 13
8 | 8 168.06 13.76 757.2 66 29 7
9 | 9 131.83 14.77 825.4 39 19 1
10 | 10 114.78 19.11 1096.4 155 15 7
11 | 11 102.79 15.18 823.7 43 14 2
12 | 12 99.78 13.34 789.4 46 18 1
13 | 13 97.51 13.90 808.7 41 13 0
14 | 14 74.79 17.29 911.2 29 7 1
15 | 15 70.23 15.84 905.8 28 7 2
16 | 16 65.56 13.19 704.0 23 10 0
17 | 17 62.22 13.34 865.1 32 11 5
18 | 18 61.72 17.48 970.2 7 6 0
19 | 19 57.56 13.56 708.2 10 6 1
20 | 20 52.80 13.41 732.9 21 7 2
21 | 21 52.52 14.44 814.2 18 11 2
22 | 22 47.76 13.40 731.9 15 12 4
23 | 23 46.92 11.33 623.9 15 9 0
24 | 24 46.52 14.98 824.4 11 6 0
25 | 25 42.55 13.99 768.4 14 7 1
26 | 26 42.06 14.23 755.8 18 5 1
Protein Coverage (double click to dock / undock)
Protein Coverage
Sample name | Protein name | Coverage summary | Coverage percent
2MC_20171219_... | >HORVU6Hr1G... | 96 of 154 | 62.34%
MGMGAMWMKSMLLVLLLCMLMVTPMTGARSDNSGPWMWCDPEMGHKVSPL
TRCRALVKLECVGNRVPEDVLRDCCQEVANISNEWCRCGDLGSMLRSVYA
ALGVGGGPEEVFPGCQRDVMKLLVAGVPALCNVPIPNEAAGTRGVCYWSA
Peptide List (double click to dock / undock)
Peptides
Prot. Rank | Pos. | Sequence | Glycans | Protein Name | Score | Cleavage | z | Obs. m/z | Calc. m/z | Mods (variable) | PEP 2D | |Log Pro
31 | 3 | R.SMEGSVPK[+162.05282]YPEPTEGSIGASGAK.R | Hex(1) | >HORVU1Hr1G012090.2 gene=HORVU1Hr1G012090 | 551.7 | Specific | 3 | 814.4052 | 814.3844 | K8(xgly / 162.0528) | 5e-07 | 6.30
Spectrum (double click to dock / undock)
R.SMEGSVPK[+162]YPEPTEGSIGASGAK.R z=3,scan#=sample=1 period=1 cycle=799 experiment=20,scan time=10.0446
SMEGSVPKYPEPTEGSIGASGAK
Intensity
m/z
m/z errors, tol=75.0 ppm
Obs - Calc m/z

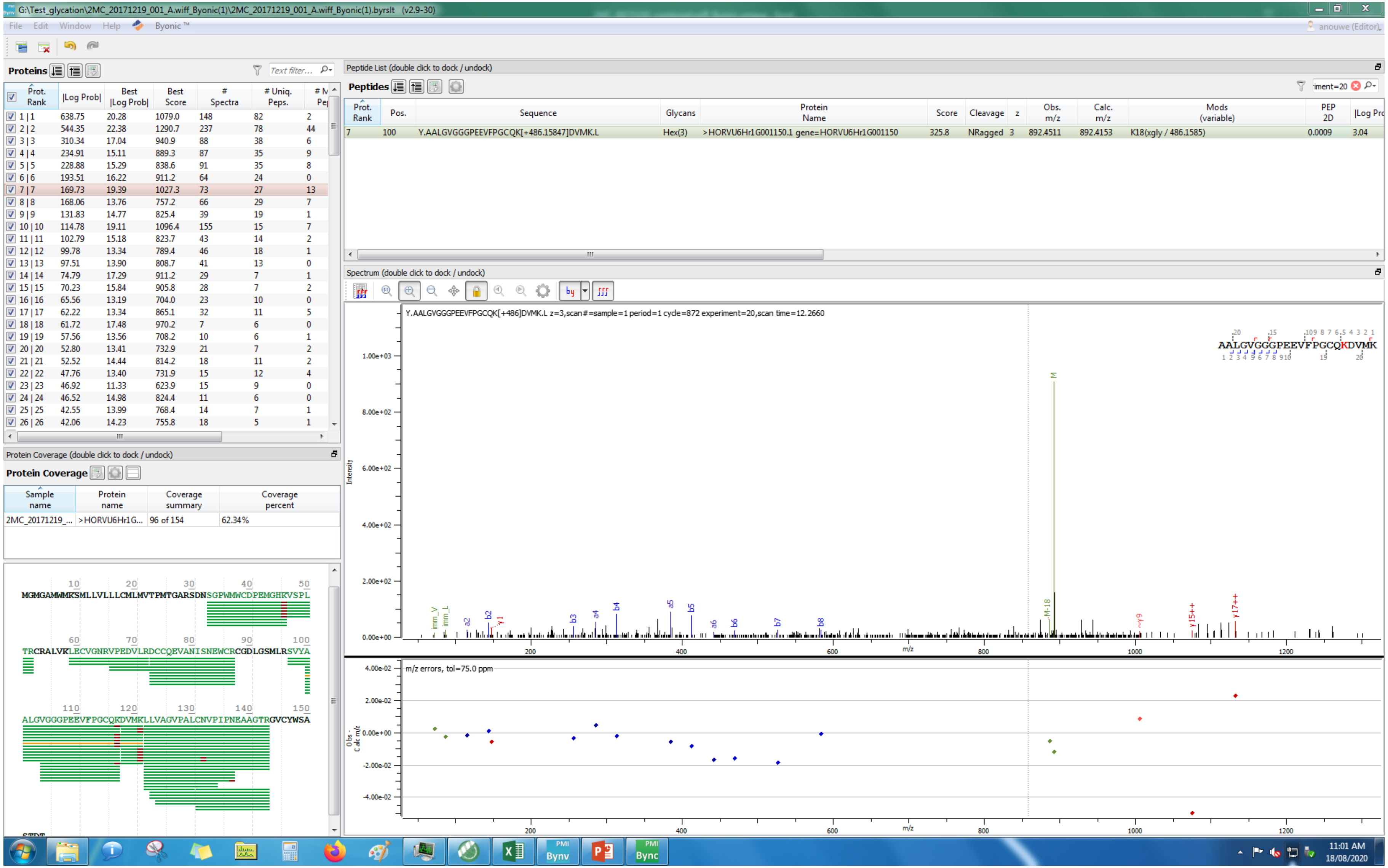
G:\Test_glycation\2MC_20171219_001_A.wiff_Byonic(1)\2MC_20171219_001_A.wiff_Byonic(1).byrslt (v2.9-30)
File Edit Window Help Byonic™
Proteins
Prot. Rank | |Log Prob| | Best |Log Prob| | Best Score | # Spectra | # Uniq. Peps.
1 | 1 638.75 20.28 1079.0 148 82 2
2 | 2 544.35 22.38 1290.7 237 78 44
3 | 3 310.34 17.04 940.9 88 38 6
4 | 4 234.91 15.11 889.3 87 35 9
5 | 5 228.88 15.29 838.6 91 35 8
6 | 6 193.51 16.22 911.2 64 24 0
7 | 7 169.73 19.39 1027.3 73 27 13
8 | 8 168.06 13.76 757.2 66 29 7
9 | 9 131.83 14.77 825.4 39 19 1
10 | 10 114.78 19.11 1096.4 155 15 7
11 | 11 102.79 15.18 823.7 43 14 2
12 | 12 99.78 13.34 789.4 46 18 1
13 | 13 97.51 13.90 808.7 41 13 0
14 | 14 74.79 17.29 911.2 29 7 1
15 | 15 70.23 15.84 905.8 28 7 2
16 | 16 65.56 13.19 704.0 23 10 0
17 | 17 62.22 13.34 865.1 32 11 5
18 | 18 61.72 17.48 970.2 7 6 0
19 | 19 57.56 13.56 708.2 10 6 1
20 | 20 52.80 13.41 732.9 21 7 2
21 | 21 52.52 14.44 814.2 18 11 2
22 | 22 47.76 13.40 731.9 15 12 4
23 | 23 46.92 11.33 623.9 15 9 0
24 | 24 46.52 14.98 824.4 11 6 0
25 | 25 42.55 13.99 768.4 14 7 1
26 | 26 42.06 14.23 755.8 18 5 1
Peptide List (double click to dock / undock)
Peptides
Prot. Rank | Pos. | Sequence | Glycans | Protein Name | Score | Cleavage | z | Obs. m/z | Calc. m/z | Mods (variable) | PEP 2D | |Log Prob|
7 100 Y.AALGVGGGPEEVFPGCQK[+486.15847]DVMK.L Hex(3) >HORVU6Hr1G001150.1 gene=HORVU6Hr1G001150 325.8 NRagged 3 892.4511 892.4153 K18(xgly / 486.1585) 0.0009 3.04
Spectrum (double click to dock / undock)
Y.AALGVGGGPEEVFPGCQK[+486]DVMK.L z=3,scan#=sample=1 period=1 cycle=872 experiment=20,scan time=12.2660
AALGVGGGPEEVFPGCQKDVMK
Intensity
m/z
m/z errors, tol=75.0 ppm
Protein Coverage (double click to dock / undock)
Protein Coverage
Sample name | Protein name | Coverage summary | Coverage percent
2MC_20171219_... >HORVU6Hr1G... 96 of 154 62.34%
MGMGAMWMKSMLLVLLLCMLMVTPMTGARSDNSGPWMWCDPEMGHKVSPL
TRCRALVKLECVGNRVPEDVLRDCCQEVANISNEWCRCGDLGSMLRSVYA
ALGVGGGPEEVFPGCQKDVMKLLVAGVPALCNVPIPNEAAGTRGVCYWSA
11:01 AM
18/08/2020

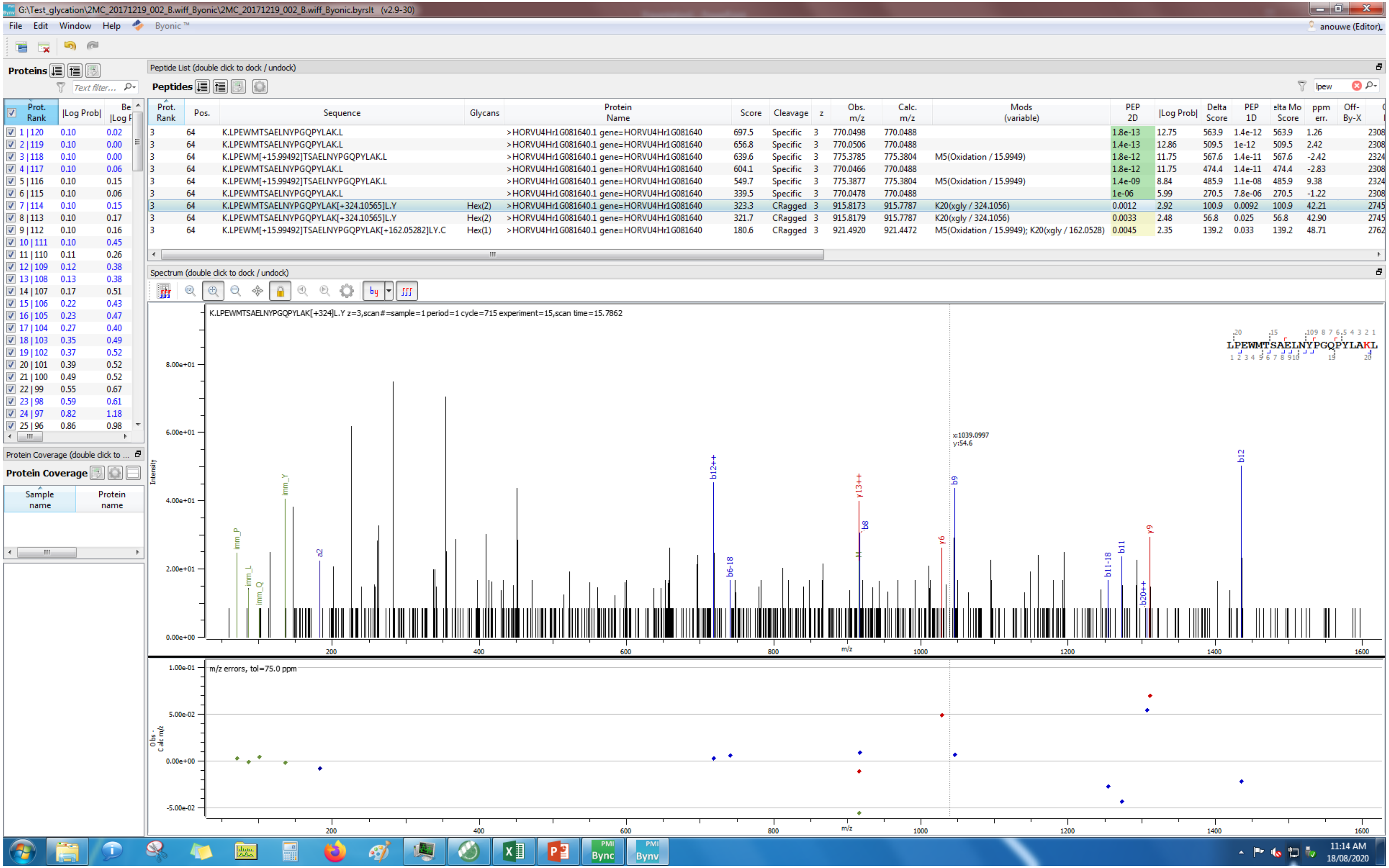
G:\Test_glycation\2MC_20171219_002_B.wiff_Byonic\2MC_20171219_002_B.wiff_Byonic.byrslt (v2.9-30)
File Edit Window Help Byonic™
anouwe (Editor)
Proteins
Peptide List (double click to dock / undock)
Peptides
Prot. Rank | Pos. | Sequence | Glycans | Protein Name | Score | Cleavage | z | Obs. m/z | Calc. m/z | Mods (variable) | PEP 2D | |Log Prob| | Delta Score | PEP 1D | Delta Mod Score | ppm err. | Off-By-X
3 64 K.LPEWMTSAELNYPGQPYLAK.L >HORVU4Hr1G081640.1 gene=HORVU4Hr1G081640 697.5 Specific 3 770.0498 770.0488 1.8e-13 12.75 563.9 1.4e-12 563.9 1.26
3 64 K.LPEWMTSAELNYPGQPYLAK.L >HORVU4Hr1G081640.1 gene=HORVU4Hr1G081640 656.8 Specific 3 770.0506 770.0488 1.4e-13 12.86 509.5 1e-12 509.5 2.42
3 64 K.LPEWM[+15.99492]TSAELNYPGQPYLAK.L >HORVU4Hr1G081640.1 gene=HORVU4Hr1G081640 639.6 Specific 3 775.3785 775.3804 M5(Oxidation / 15.9949) 1.8e-12 11.75 567.6 1.4e-11 567.6 -2.42
3 64 K.LPEWMTSAELNYPGQPYLAK.L >HORVU4Hr1G081640.1 gene=HORVU4Hr1G081640 604.1 Specific 3 770.0466 770.0488 1.8e-12 11.75 474.4 1.4e-11 474.4 -2.83
3 64 K.LPEWM[+15.99492]TSAELNYPGQPYLAK.L >HORVU4Hr1G081640.1 gene=HORVU4Hr1G081640 549.7 Specific 3 775.3877 775.3804 M5(Oxidation / 15.9949) 1.4e-09 8.84 485.9 1.1e-08 485.9 9.38
3 64 K.LPEWMTSAELNYPGQPYLAK.L >HORVU4Hr1G081640.1 gene=HORVU4Hr1G081640 339.5 Specific 3 770.0478 770.0488 1e-06 5.99 270.5 7.8e-06 270.5 -1.22
3 64 K.LPEWMTSAELNYPGQPYLAK[+324.10565]L.Y Hex(2) >HORVU4Hr1G081640.1 gene=HORVU4Hr1G081640 323.3 CRagged 3 915.8173 915.7787 K20(xgly / 324.1056) 0.0012 2.92 100.9 0.0092 100.9 42.21
3 64 K.LPEWMTSAELNYPGQPYLAK[+324.10565]L.Y Hex(2) >HORVU4Hr1G081640.1 gene=HORVU4Hr1G081640 321.7 CRagged 3 915.8179 915.7787 K20(xgly / 324.1056) 0.0033 2.48 56.8 0.025 56.8 42.90
3 64 K.LPEWM[+15.99492]TSAELNYPGQPYLAK[+162.05282]LY.C Hex(1) >HORVU4Hr1G081640.1 gene=HORVU4Hr1G081640 180.6 CRagged 3 921.4920 921.4472 M5(Oxidation / 15.9949); K20(xgly / 162.0528) 0.0045 2.35 139.2 0.033 139.2 48.71
Spectrum (double click to dock / undock)
K.LPEWMTSAELNYPGQPYLAK[+324]L.Y z=3,scan#=sample=1 period=1 cycle=715 experiment=15,scan time=15.7862
LPEWMTSAELNYPGQPYLAKL
Intensity
m/z
m/z errors, tol=75.0 ppm
Protein Coverage
11:14 AM
18/08/2020

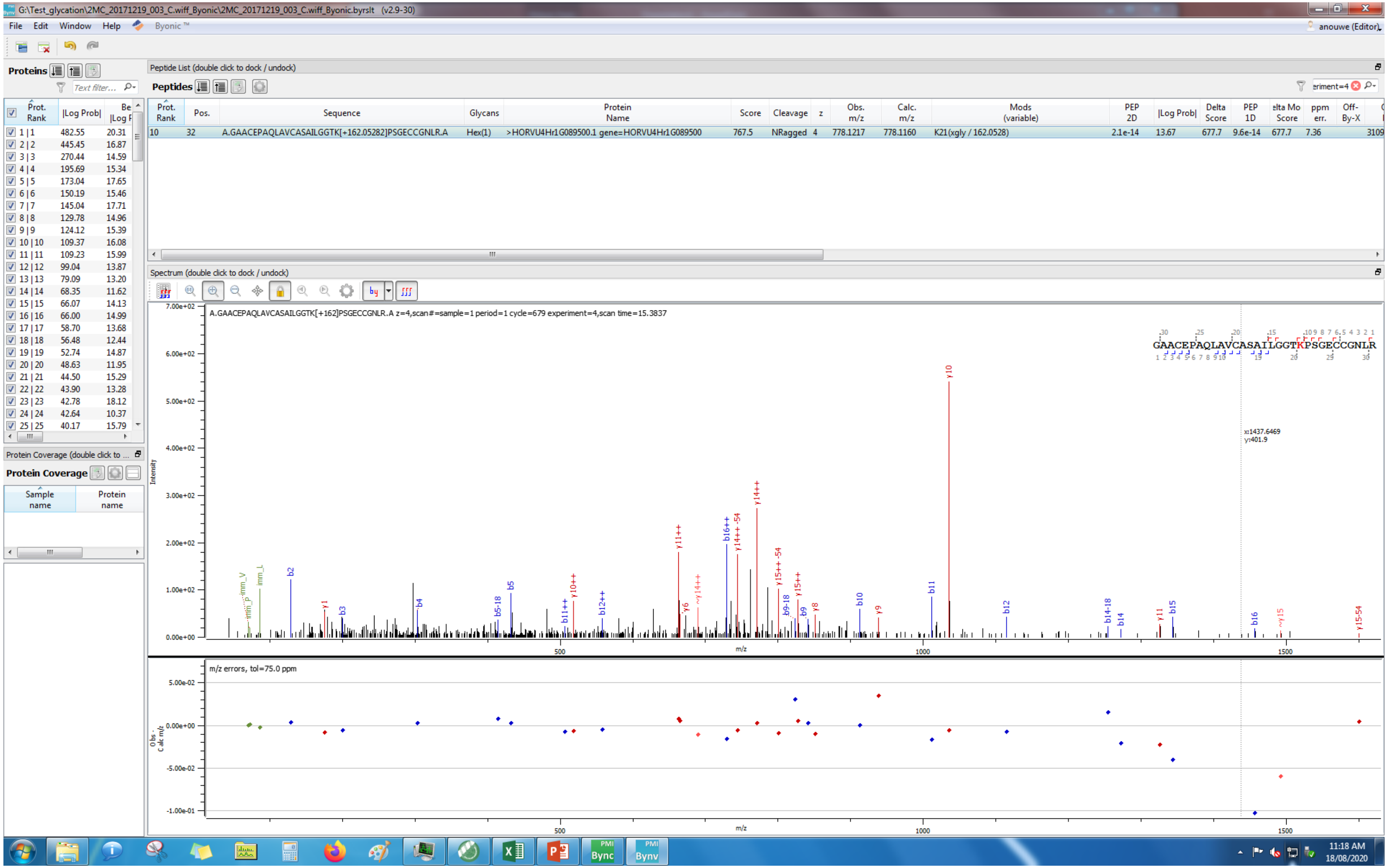
G:\Test_glycation\2MC_20171219_003_C.wiff_Byonic\2MC_20171219_003_C.wiff_Byonic.byrslt (v2.9-30)
File Edit Window Help Byonic™
anouwe (Editor)
Proteins
Peptide List (double click to dock / undock)
Peptides
Prot. Rank | Pos. | Sequence | Glycans | Protein Name | Score | Cleavage | z | Obs. m/z | Calc. m/z | Mods (variable) | PEP 2D | |Log Prob| | Delta Score | PEP 1D | Delta Mod Score | ppm err. | Off-By-X
10 | 32 | A.GAACEPAQLAVCASAILGGTK[+162.05282]PSGECCGNLR.A | Hex(1) | >HORVU4Hr1G089500.1 gene=HORVU4Hr1G089500 | 767.5 | NRagged | 4 | 778.1217 | 778.1160 | K21(xgly / 162.0528) | 2.1e-14 | 13.67 | 677.7 | 9.6e-14 | 677.7 | 7.36 | 3109
Spectrum (double click to dock / undock)
A.GAACEPAQLAVCASAILGGTK[+162]PSGECCGNLR.A z=4,scan#=sample=1 period=1 cycle=679 experiment=4,scan time=15.3837
GAACEPAQLAVCASAILGGTKPSGECCGNLR
x:1437.6469
y:401.9
Intensity
m/z
m/z errors, tol=75.0 ppm
Obs - Calc m/z
Protein Coverage (double click to ...
Protein Coverage
Sample name
Protein name
11:18 AM
18/08/2020

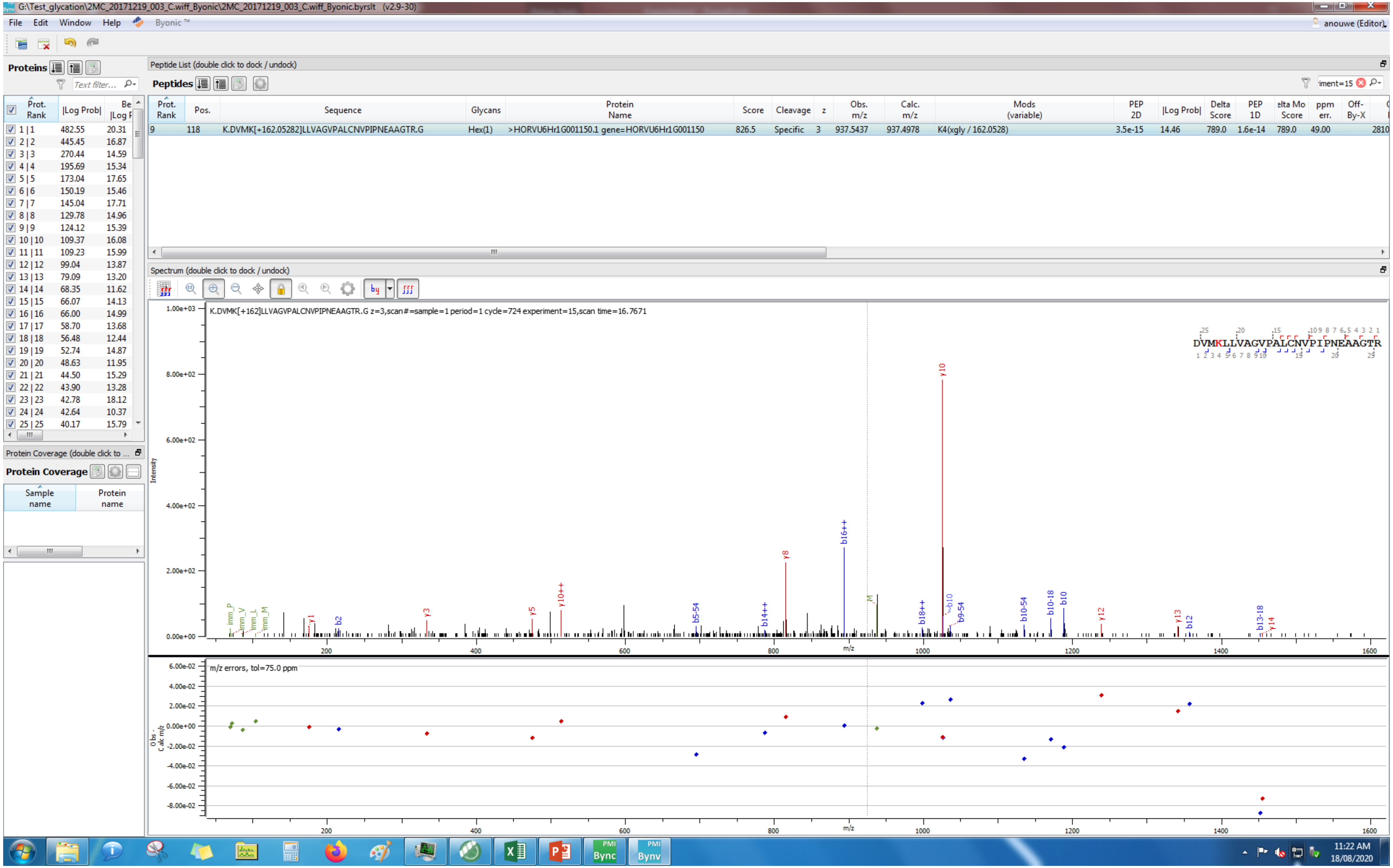

G:\Test_glycation\2MC_20171219_003_C.wiff_Byonic\2MC_20171219_003_C.wiff_Byonic.byrslt (v2.9-30)
File Edit Window Help Byonic™
anouwe (Editor)
Proteins
Peptide List (double click to dock / undock)
Peptides
Prot. Rank | Pos. | Sequence | Glycans | Protein Name | Score | Cleavage | z | Obs. m/z | Calc. m/z | Mods (variable) | PEP 2D | |Log Prob| | Delta Score | PEP 1D | Delta Mod Score | ppm err. | Off-By-X
9 | 118 | K.DVMK[+162.05282]LLVAGVPALCNVPIPNEAAGTR.G | Hex(1) | >HORVU6Hr1G001150.1 gene=HORVU6Hr1G001150 | 826.5 | Specific | 3 | 937.5437 | 937.4978 | K4(xgly / 162.0528) | 3.5e-15 | 14.46 | 789.0 | 1.6e-14 | 789.0 | 49.00
Spectrum (double click to dock / undock)
K.DVMK[+162]LLVAGVPALCNVPIPNEAAGTR.G z=3,scan#=sample=1 period=1 cycle=724 experiment=15,scan time=16.7671
DVMKLLVAGVPALCNVPIPNEAAGTR
Intensity
m/z
m/z errors, tol=75.0 ppm
Obs - Calc m/z
Protein Coverage (double click to dock / undock)
Protein Coverage
Sample name
Protein name
11:22 AM
18/08/2020

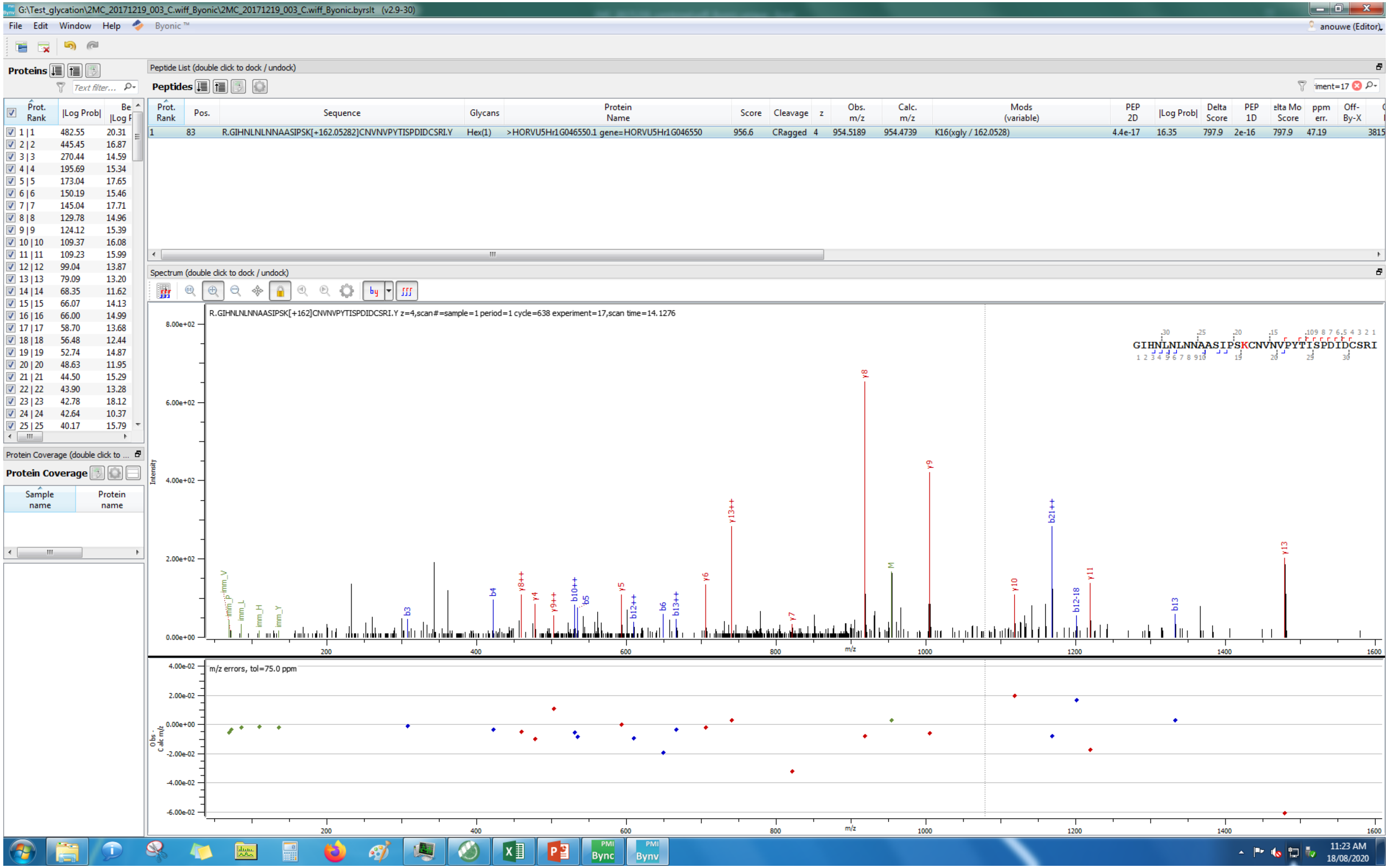
G:\Test_glycation\2MC_20171219_003_C.wiff_Byonic\2MC_20171219_003_C.wiff_Byonic.byrslt (v2.9-30)
File Edit Window Help Byonic™
anouwe (Editor)
Proteins
Peptide List (double click to dock / undock)
Peptides
R.GIHNLNLNNAASIPSK[+162.05282]CNVNVPYTISPDIDCSRI.Y
Hex(1)
>HORVU5Hr1G046550.1 gene=HORVU5Hr1G046550
956.6
CRagged
954.5189
954.4739
K16(xgly / 162.0528)
Spectrum (double click to dock / undock)
R.GIHNLNLNNAASIPSK[+162]CNVNVPYTISPDIDCSRI.Y z=4,scan#=sample=1 period=1 cycle=638 experiment=17,scan time=14.1276
Intensity
m/z
m/z errors, tol=75.0 ppm
Protein Coverage
11:23 AM
18/08/2020

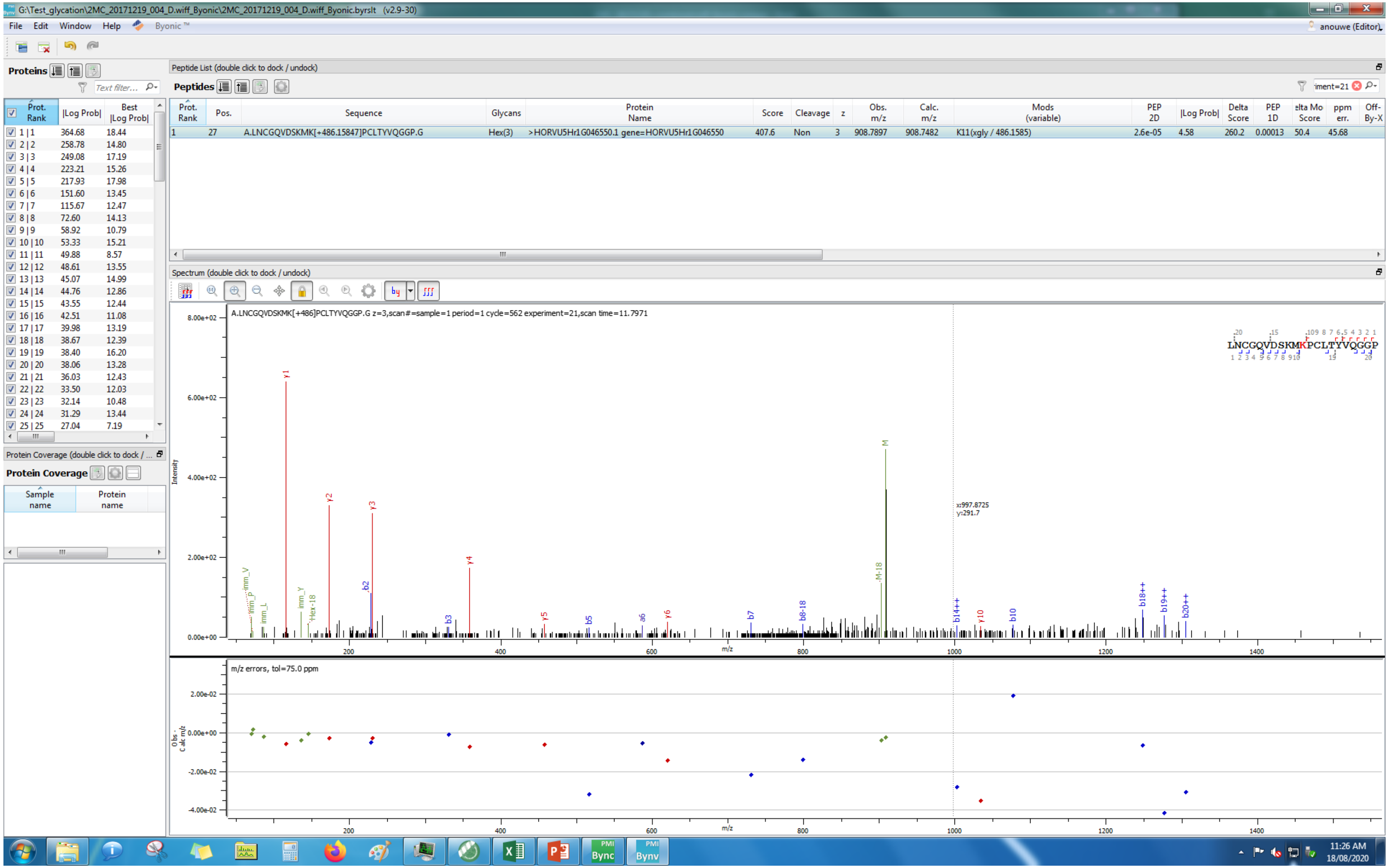
G:\Test_glycation\2MC_20171219_004_D.wiff_Byonic\2MC_20171219_004_D.wiff_Byonic.byrslt (v2.9-30)
File Edit Window Help Byonic™
anouwe (Editor)
Proteins
Peptide List (double click to dock / undock)
Peptides
Prot. Rank | Pos. | Sequence | Glycans | Protein Name | Score | Cleavage | z | Obs. m/z | Calc. m/z | Mods (variable) | PEP 2D | |Log Prob| | Delta Score | PEP 1D | Delta Mod Score | ppm err. | Off-By-X
1 | 27 | A.LNCGQVDSKMK[+486.15847]PCLTYVQGGP.G | Hex(3) | >HORVU5Hr1G046550.1 gene=HORVU5Hr1G046550 | 407.6 | Non | 3 | 908.7897 | 908.7482 | K11(xgly / 486.1585) | 2.6e-05 | 4.58 | 260.2 | 0.00013 | 50.4 | 45.68
Spectrum (double click to dock / undock)
A.LNCGQVDSKMK[+486]PCLTYVQGGP.G z=3,scan#=sample=1 period=1 cycle=562 experiment=21,scan time=11.7971
LNCGQVDSKMKPCLTYVQGGP
Intensity
m/z
x:997.8725
y:291.7
m/z errors, tol=75.0 ppm
Obs - Calc m/z
Protein Coverage (double click to dock / undock)
Protein Coverage
Sample name
Protein name
11:26 AM
18/08/2020

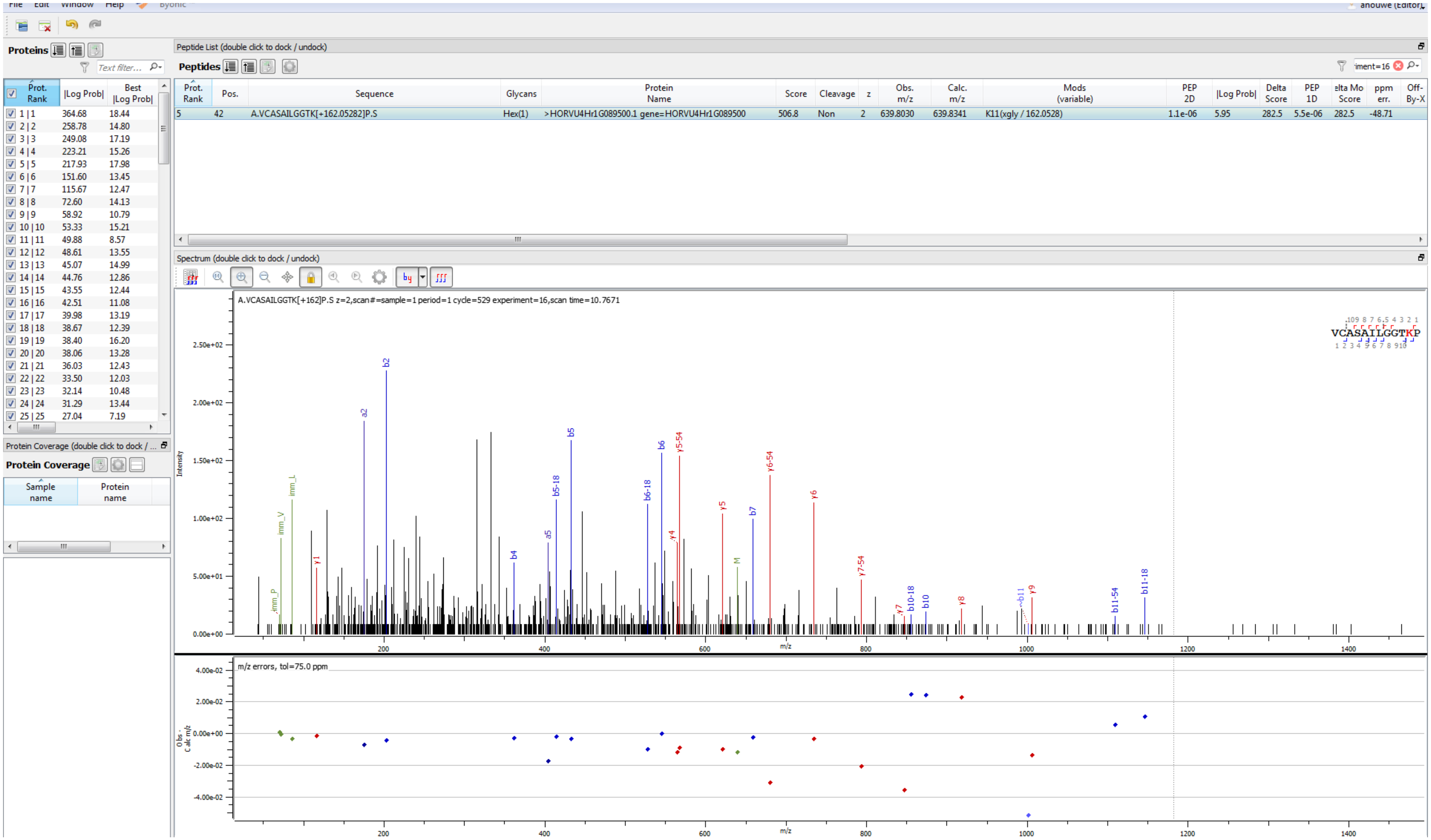
File Edit Window Help Byonic
Proteins
Text filter...
Prot. Rank | |Log Prob| | Best |Log Prob|
1 | 1 364.68 18.44
2 | 2 258.78 14.80
3 | 3 249.08 17.19
4 | 4 223.21 15.26
5 | 5 217.93 17.98
6 | 6 151.60 13.45
7 | 7 115.67 12.47
8 | 8 72.60 14.13
9 | 9 58.92 10.79
10 | 10 53.33 15.21
11 | 11 49.88 8.57
12 | 12 48.61 13.55
13 | 13 45.07 14.99
14 | 14 44.76 12.86
15 | 15 43.55 12.44
16 | 16 42.51 11.08
17 | 17 39.98 13.19
18 | 18 38.67 12.39
19 | 19 38.40 16.20
20 | 20 38.06 13.28
21 | 21 36.03 12.43
22 | 22 33.50 12.03
23 | 23 32.14 10.48
24 | 24 31.29 13.44
25 | 25 27.04 7.19
Protein Coverage (double click to dock / ...
Protein Coverage
Sample name | Protein name
Peptide List (double click to dock / undock)
Peptides
Prot. Rank | Pos. | Sequence | Glycans | Protein Name | Score | Cleavage | z | Obs. m/z | Calc. m/z | Mods (variable) | PEP 2D | |Log Prob| | Delta Score | PEP 1D | elta Mo Score | ppm err. | Off-By-X
5 | 42 | A.VCASAILGGTK[+162.05282]P.S | Hex(1) | >HORVU4Hr1G089500.1 gene=HORVU4Hr1G089500 | 506.8 | Non | 2 | 639.8030 | 639.8341 | K11(xgly / 162.0528) | 1.1e-06 | 5.95 | 282.5 | 5.5e-06 | 282.5 | -48.71
Spectrum (double click to dock / undock)
A.VCASAILGGTK[+162]P.S z=2,scan#=sample=1 period=1 cycle=529 experiment=16,scan time=10.7671
VCASAILGGTKP
Intensity
m/z
m/z errors, tol=75.0 ppm
Obs - Calc m/z

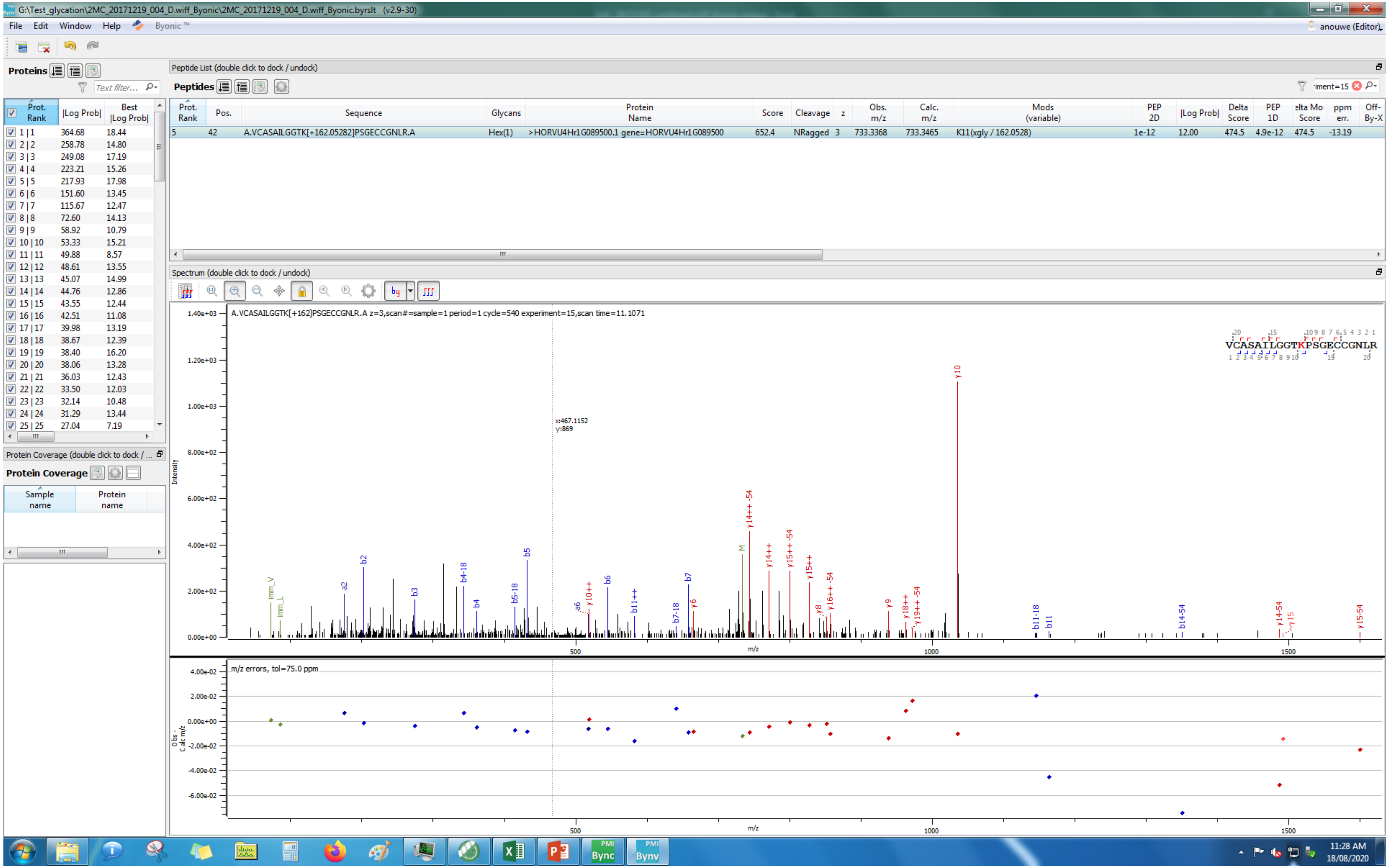
G:\Test_glycation\2MC_20171219_004_D.wiff_Byonic\2MC_20171219_004_D.wiff_Byonic.byrslt (v2.9-30)
File Edit Window Help Byonic™
anouwe (Editor)
Proteins
Peptide List (double click to dock / undock)
Peptides
Prot. Rank | Pos. | Sequence | Glycans | Protein Name | Score | Cleavage | z | Obs. m/z | Calc. m/z | Mods (variable) | PEP 2D | |Log Prob| | Delta Score | PEP 1D | Delta Mod Score | ppm err. | Off-By-X
5 | 42 | A.VCASAILGGTK[+162.05282]PSGECCGNLR.A | Hex(1) | >HORVU4Hr1G089500.1 gene=HORVU4Hr1G089500 | 652.4 | NRagged | 3 | 733.3368 | 733.3465 | K11(xgly / 162.0528) | 1e-12 | 12.00 | 474.5 | 4.9e-12 | 474.5 | -13.19
Prot. Rank | |Log Prob| | Best |Log Prob|
1 | 1 364.68 18.44
2 | 2 258.78 14.80
3 | 3 249.08 17.19
4 | 4 223.21 15.26
5 | 5 217.93 17.98
6 | 6 151.60 13.45
7 | 7 115.67 12.47
8 | 8 72.60 14.13
9 | 9 58.92 10.79
10 | 10 53.33 15.21
11 | 11 49.88 8.57
12 | 12 48.61 13.55
13 | 13 45.07 14.99
14 | 14 44.76 12.86
15 | 15 43.55 12.44
16 | 16 42.51 11.08
17 | 17 39.98 13.19
18 | 18 38.67 12.39
19 | 19 38.40 16.20
20 | 20 38.06 13.28
21 | 21 36.03 12.43
22 | 22 33.50 12.03
23 | 23 32.14 10.48
24 | 24 31.29 13.44
25 | 25 27.04 7.19
Protein Coverage (double click to dock / ...
Protein Coverage
Sample name
Protein name
Spectrum (double click to dock / undock)
A.VCASAILGGTK[+162]PSGECCGNLR.A z=3,scan#=sample=1 period=1 cycle=540 experiment=15,scan time=11.1071
VCASAILGGTKPSGECCGNLR
x:467.1152
y:869
Intensity
m/z
m/z errors, tol=75.0 ppm
Obs - Calc m/z
11:28 AM
18/08/2020

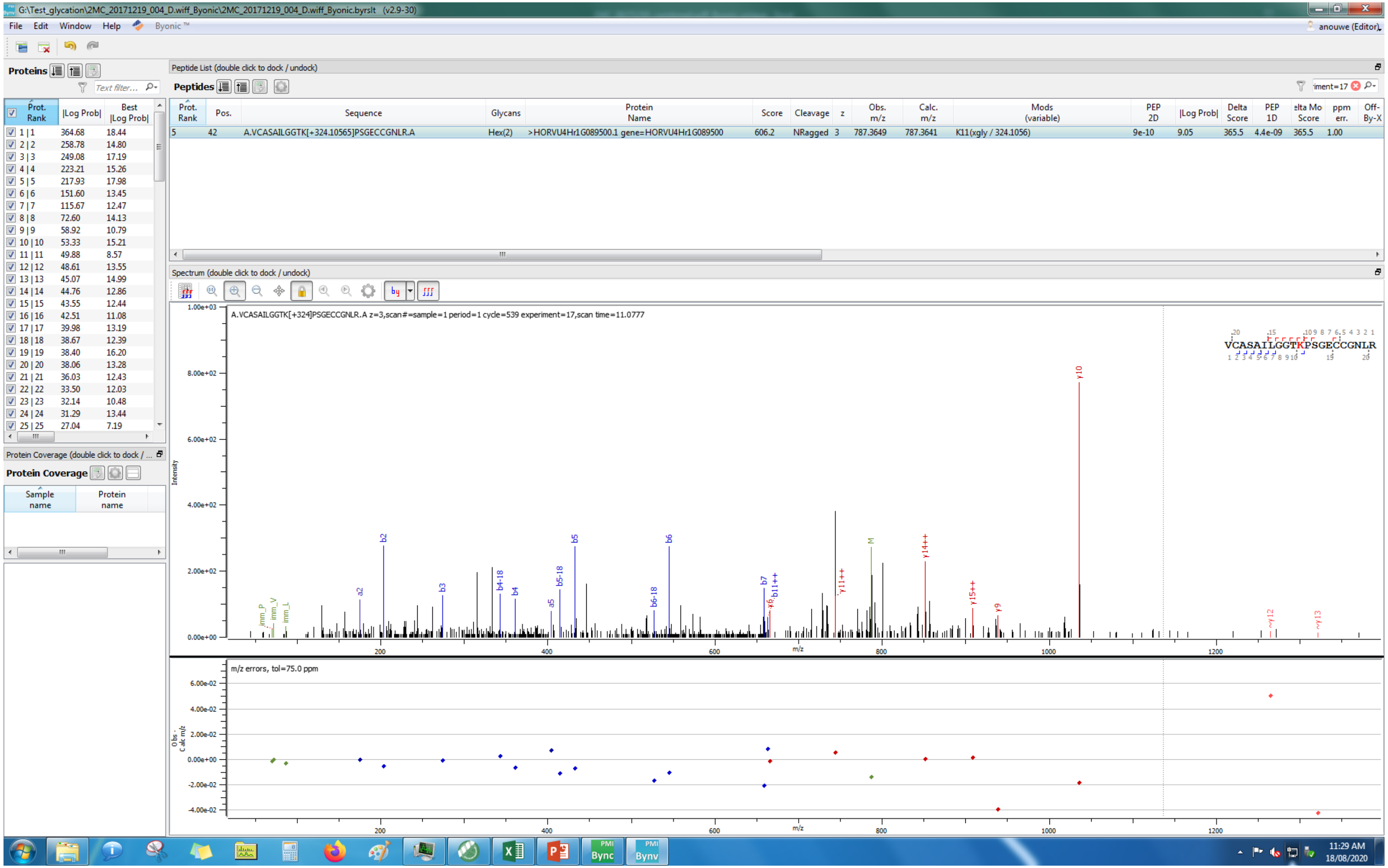
G:\Test_glycation\2MC_20171219_004_D.wiff_Byonic\2MC_20171219_004_D.wiff_Byonic.byrslt (v2.9-30)
File Edit Window Help Byonic™
anouwe (Editor)
Proteins
Prot. Rank | |Log Prob| | Best |Log Prob|
1 | 1 364.68 18.44
2 | 2 258.78 14.80
3 | 3 249.08 17.19
4 | 4 223.21 15.26
5 | 5 217.93 17.98
6 | 6 151.60 13.45
7 | 7 115.67 12.47
8 | 8 72.60 14.13
9 | 9 58.92 10.79
10 | 10 53.33 15.21
11 | 11 49.88 8.57
12 | 12 48.61 13.55
13 | 13 45.07 14.99
14 | 14 44.76 12.86
15 | 15 43.55 12.44
16 | 16 42.51 11.08
17 | 17 39.98 13.19
18 | 18 38.67 12.39
19 | 19 38.40 16.20
20 | 20 38.06 13.28
21 | 21 36.03 12.43
22 | 22 33.50 12.03
23 | 23 32.14 10.48
24 | 24 31.29 13.44
25 | 25 27.04 7.19
Protein Coverage
Sample name | Protein name
Peptide List (double click to dock / undock)
Peptides
Prot. Rank | Pos. | Sequence | Glycans | Protein Name | Score | Cleavage | z | Obs. m/z | Calc. m/z | Mods (variable) | PEP 2D | |Log Prob| | Delta Score | PEP 1D | elta Mo Score | ppm err. | Off-By-X
5 | 42 | A.VCASAILGGTK[+324.10565]PSGECCGNLR.A | Hex(2) | >HORVU4Hr1G089500.1 gene=HORVU4Hr1G089500 | 606.2 | NRagged | 3 | 787.3649 | 787.3641 | K11(xgly / 324.1056) | 9e-10 | 9.05 | 365.5 | 4.4e-09 | 365.5 | 1.00
Spectrum (double click to dock / undock)
A.VCASAILGGTK[+324]PSGECCGNLR.A z=3,scan#=sample=1 period=1 cycle=539 experiment=17,scan time=11.0777
VCASAILGGTKPSGECCGNLR
Intensity
m/z
m/z errors, tol=75.0 ppm
Obs - Calc m/z
11:29 AM
18/08/2020

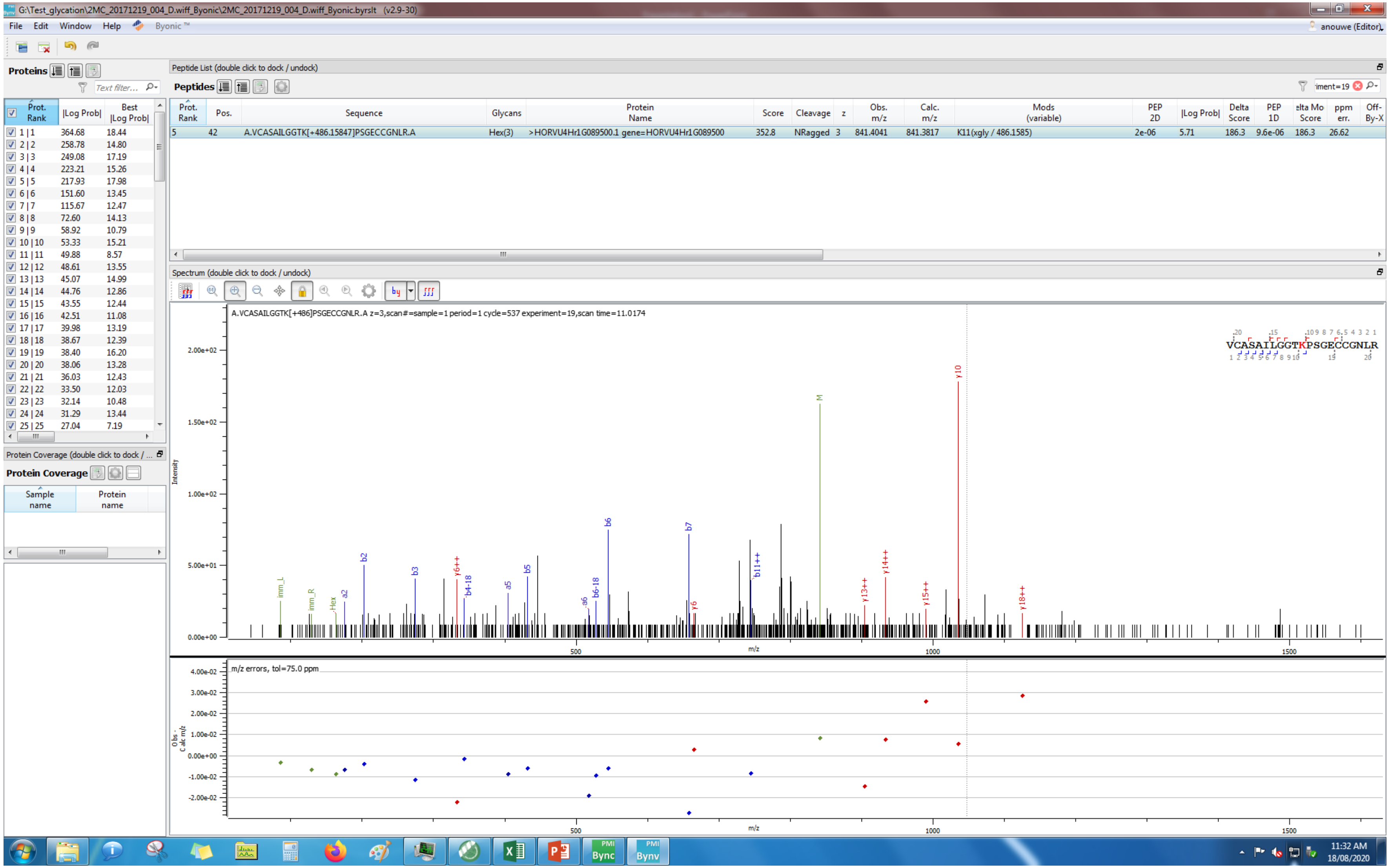
G:\Test_glycation\2MC_20171219_004_D.wiff_Byonic\2MC_20171219_004_D.wiff_Byonic.byrslt (v2.9-30)
File Edit Window Help Byonic
anouwe (Editor)
Proteins
Peptide List (double click to dock / undock)
Peptides
Prot. Rank | Pos. | Sequence | Glycans | Protein Name | Score | Cleavage | z | Obs. m/z | Calc. m/z | Mods (variable) | PEP 2D | |Log Prob| | Delta Score | PEP 1D | elta Mo Score | ppm err. | Off-By-X
5 | 42 | A.VCASAILGGTK[+486.15847]PSGECCGNLR.A | Hex(3) | >HORVU4Hr1G089500.1 gene=HORVU4Hr1G089500 | 352.8 | NRagged | 3 | 841.4041 | 841.3817 | K11(xgly / 486.1585) | 2e-06 | 5.71 | 186.3 | 9.6e-06 | 186.3 | 26.62
Prot. Rank | |Log Prob| | Best |Log Prob|
1 | 1 364.68 18.44
2 | 2 258.78 14.80
3 | 3 249.08 17.19
4 | 4 223.21 15.26
5 | 5 217.93 17.98
6 | 6 151.60 13.45
7 | 7 115.67 12.47
8 | 8 72.60 14.13
9 | 9 58.92 10.79
10 | 10 53.33 15.21
11 | 11 49.88 8.57
12 | 12 48.61 13.55
13 | 13 45.07 14.99
14 | 14 44.76 12.86
15 | 15 43.55 12.44
16 | 16 42.51 11.08
17 | 17 39.98 13.19
18 | 18 38.67 12.39
19 | 19 38.40 16.20
20 | 20 38.06 13.28
21 | 21 36.03 12.43
22 | 22 33.50 12.03
23 | 23 32.14 10.48
24 | 24 31.29 13.44
25 | 25 27.04 7.19
Protein Coverage (double click to dock / ...
Protein Coverage
Sample name | Protein name
Spectrum (double click to dock / undock)
A.VCASAILGGTK[+486]PSGECCGNLR.A z=3,scan#=sample=1 period=1 cycle=537 experiment=19,scan time=11.0174
VCASAILGGTKPSGECCGNLR
Intensity
m/z
m/z errors, tol=75.0 ppm
Obs - Calc m/z
11:32 AM
18/08/2020

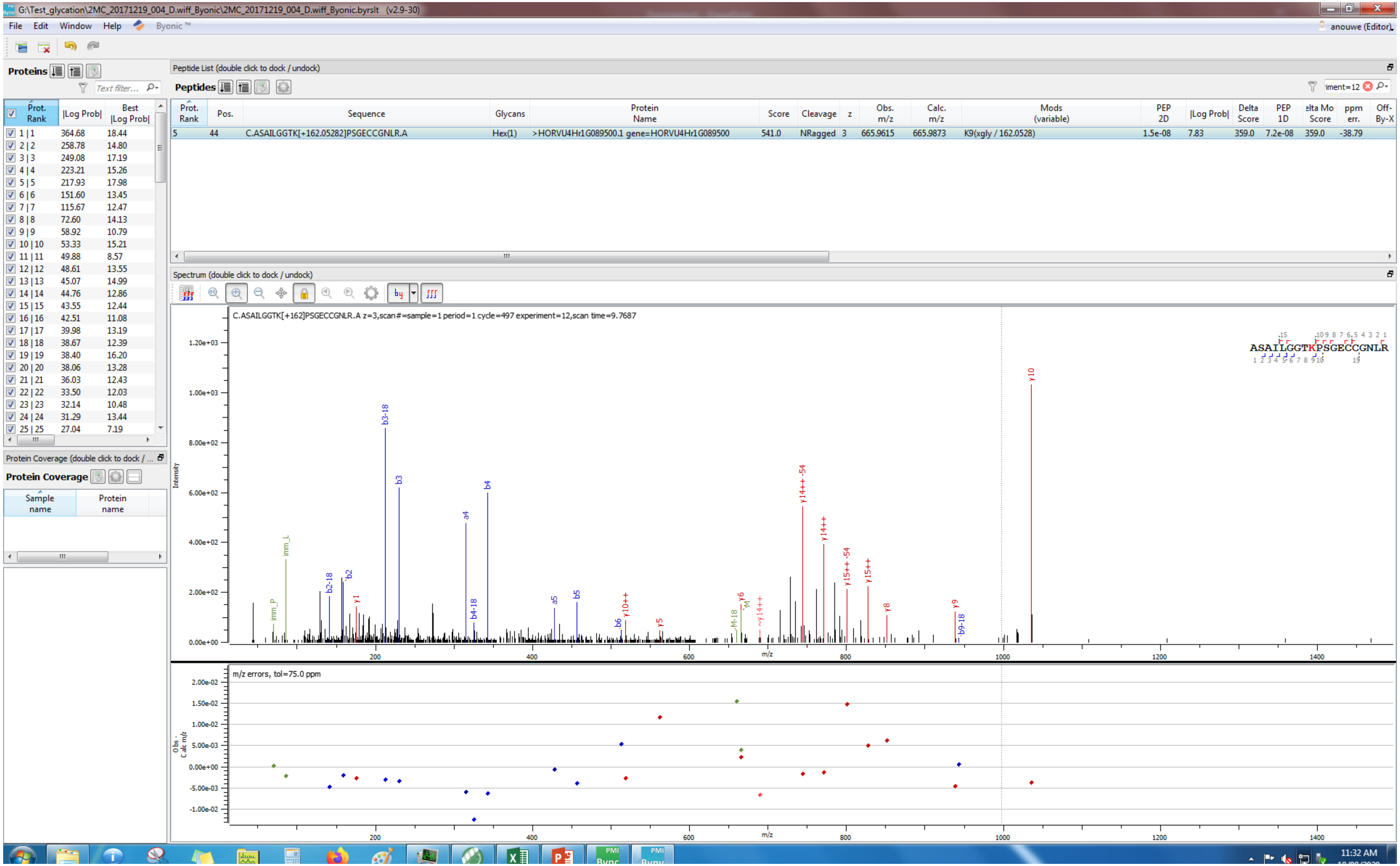
G:\Test_glycation\2MC_20171219_004_D.wiff_Byonic\2MC_20171219_004_D.wiff_Byonic.byrslt (v2.9-30)
File Edit Window Help Byonic™
anouwe (Editor)
Proteins
Prot. Rank | |Log Prob| | Best |Log Prob|
1 | 1 364.68 18.44
2 | 2 258.78 14.80
3 | 3 249.08 17.19
4 | 4 223.21 15.26
5 | 5 217.93 17.98
6 | 6 151.60 13.45
7 | 7 115.67 12.47
8 | 8 72.60 14.13
9 | 9 58.92 10.79
10 | 10 53.33 15.21
11 | 11 49.88 8.57
12 | 12 48.61 13.55
13 | 13 45.07 14.99
14 | 14 44.76 12.86
15 | 15 43.55 12.44
16 | 16 42.51 11.08
17 | 17 39.98 13.19
18 | 18 38.67 12.39
19 | 19 38.40 16.20
20 | 20 38.06 13.28
21 | 21 36.03 12.43
22 | 22 33.50 12.03
23 | 23 32.14 10.48
24 | 24 31.29 13.44
25 | 25 27.04 7.19
Peptide List (double click to dock / undock)
Peptides
Prot. Rank | Pos. | Sequence | Glycans | Protein Name | Score | Cleavage | z | Obs. m/z | Calc. m/z | Mods (variable) | PEP 2D | |Log Prob| | Delta Score | PEP 1D | Delta Mod Score | ppm err. | Off-By-X
5 | 44 | C.ASAILGGTK[+162.05282]PSGECCGNLR.A | Hex(1) | >HORVU4Hr1G089500.1 gene=HORVU4Hr1G089500 | 541.0 | NRagged | 3 | 665.9615 | 665.9873 | K9(xgly / 162.0528) | 1.5e-08 | 7.83 | 359.0 | 7.2e-08 | 359.0 | -38.79
Spectrum (double click to dock / undock)
C.ASAILGGTK[+162]PSGECCGNLR.A z=3,scan#=sample=1 period=1 cycle=497 experiment=12,scan time=9.7687
ASAILGGTKPSGECCGNLR
Intensity
m/z
y10
b3-18
b3
b4
a4
y14++-54
y14++
y15++-54
y15++
imm_L
imm_P
b2-18
b2
y1
a5
b5
b4-18
b6
y10++
y5
M-18
y6
~y14++
y8
y9
b9-18
m/z errors, tol=75.0 ppm
Obs - Calc m/z
Protein Coverage
Sample name | Protein name
11:32 AM

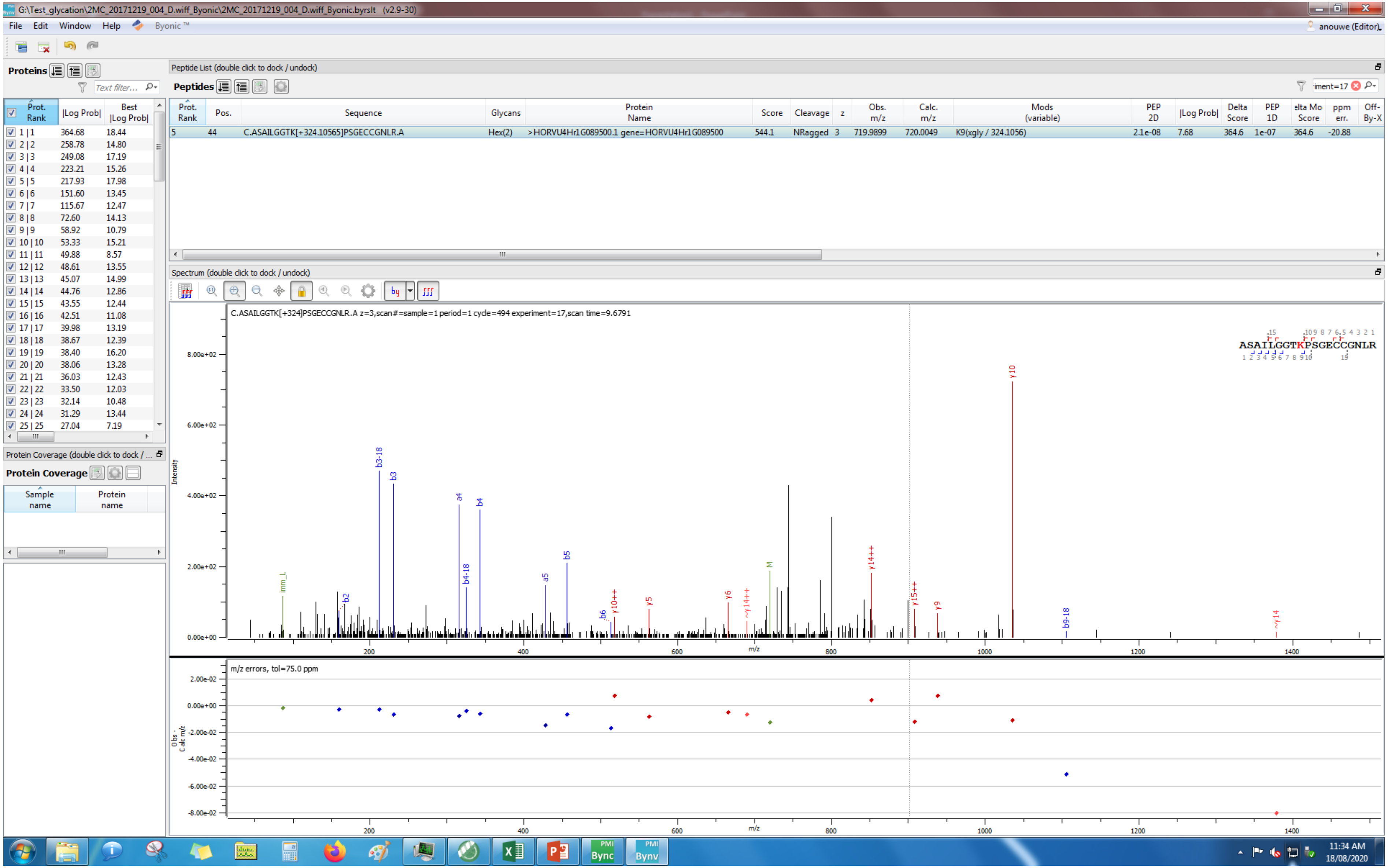
G:\Test_glycation\2MC_20171219_004_D.wiff_Byonic\2MC_20171219_004_D.wiff_Byonic.byrslt (v2.9-30)
File Edit Window Help Byonic™
anouwe (Editor)
Proteins
Text filter...
Prot. Rank | |Log Prob| | Best |Log Prob|
1 | 1 364.68 18.44
2 | 2 258.78 14.80
3 | 3 249.08 17.19
4 | 4 223.21 15.26
5 | 5 217.93 17.98
6 | 6 151.60 13.45
7 | 7 115.67 12.47
8 | 8 72.60 14.13
9 | 9 58.92 10.79
10 | 10 53.33 15.21
11 | 11 49.88 8.57
12 | 12 48.61 13.55
13 | 13 45.07 14.99
14 | 14 44.76 12.86
15 | 15 43.55 12.44
16 | 16 42.51 11.08
17 | 17 39.98 13.19
18 | 18 38.67 12.39
19 | 19 38.40 16.20
20 | 20 38.06 13.28
21 | 21 36.03 12.43
22 | 22 33.50 12.03
23 | 23 32.14 10.48
24 | 24 31.29 13.44
25 | 25 27.04 7.19
Protein Coverage (double click to dock / ...
Protein Coverage
Sample name | Protein name
Peptide List (double click to dock / undock)
Peptides
Prot. Rank | Pos. | Sequence | Glycans | Protein Name | Score | Cleavage | z | Obs. m/z | Calc. m/z | Mods (variable) | PEP 2D | |Log Prob| | Delta Score | PEP 1D | elta Mo Score | ppm err. | Off-By-X
5 44 C.ASAILGGTK[+324.10565]PSGECCGNLR.A Hex(2) >HORVU4Hr1G089500.1 gene=HORVU4Hr1G089500 544.1 NRagged 3 719.9899 720.0049 K9(xgly / 324.1056) 2.1e-08 7.68 364.6 1e-07 364.6 -20.88
Spectrum (double click to dock / undock)
C.ASAILGGTK[+324]PSGECCGNLR.A z=3,scan#=sample=1 period=1 cycle=494 experiment=17,scan time=9.6791
ASAILGGTKPSGECCGNLR
Intensity
m/z
m/z errors, tol=75.0 ppm
Obs. - Calc m/z
11:34 AM
18/08/2020

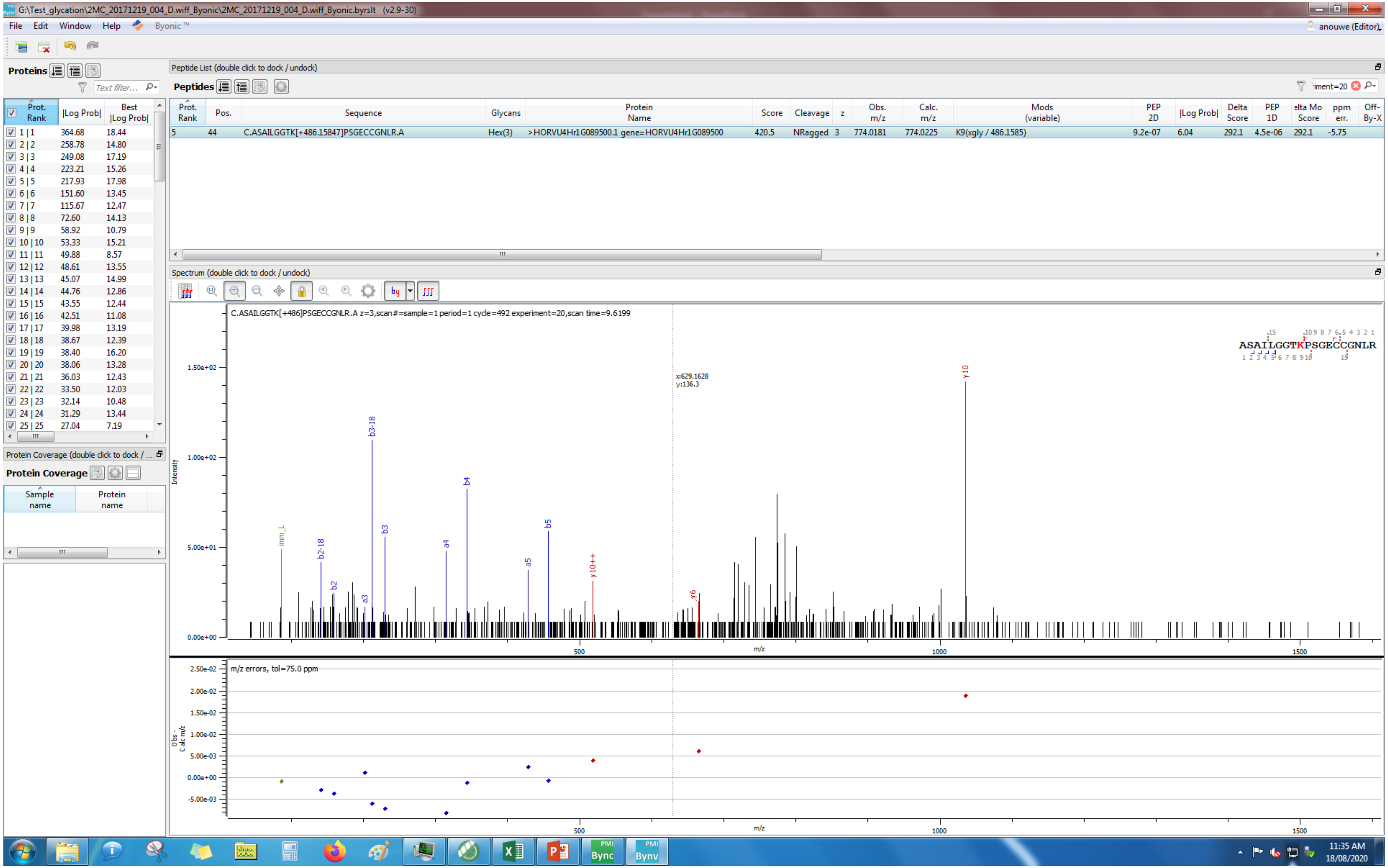
G:\Test_glycation\2MC_20171219_004_D.wiff_Byonic\2MC_20171219_004_D.wiff_Byonic.byrslt (v2.9-30)
File Edit Window Help Byonic™
anouwe (Editor)
Proteins
Peptide List (double click to dock / undock)
Peptides
Prot. Rank | Pos. | Sequence | Glycans | Protein Name | Score | Cleavage | z | Obs. m/z | Calc. m/z | Mods (variable) | PEP 2D | |Log Prob| | Delta Score | PEP 1D | elta Mo Score | ppm err. | Off-By-X
5 | 44 | C.ASAILGGTK[+486.15847]PSGECCGNLR.A | Hex(3) | >HORVU4Hr1G089500.1 gene=HORVU4Hr1G089500 | 420.5 | NRagged | 3 | 774.0181 | 774.0225 | K9(xgly / 486.1585) | 9.2e-07 | 6.04 | 292.1 | 4.5e-06 | 292.1 | -5.75
Spectrum (double click to dock / undock)
C.ASAILGGTK[+486]PSGECCGNLR.A z=3,scan#=sample=1 period=1 cycle=492 experiment=20,scan time=9.6199
ASAILGGTKPSGECCGNLR
x:629.1628
y:136.3
Intensity
m/z
m/z errors, tol=75.0 ppm
Obs - Calc m/z
Protein Coverage (double click to dock / ...
Protein Coverage
Sample name
Protein name
11:35 AM
18/08/2020

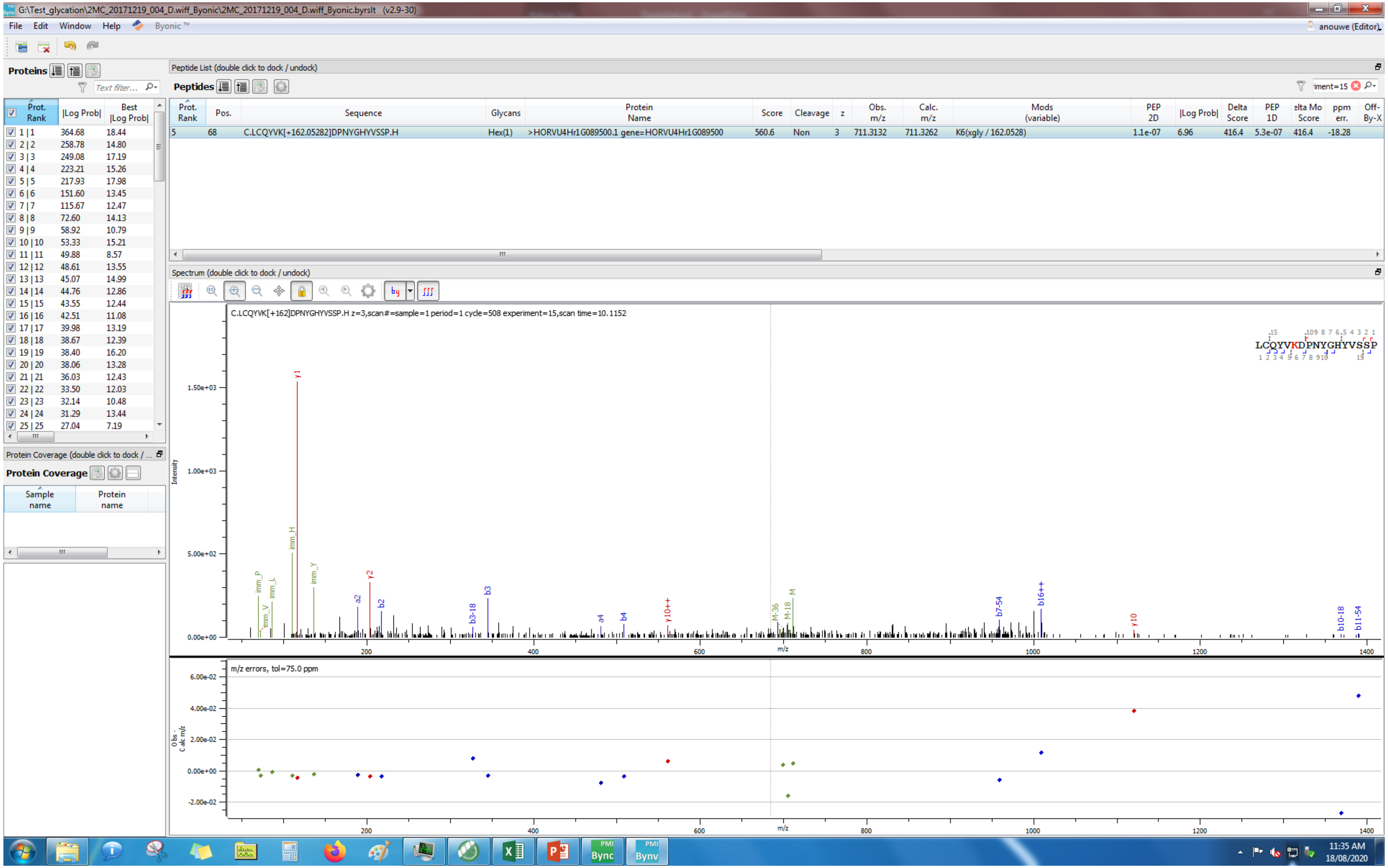

G:\Test_glycation\2MC_20171219_004_D.wiff_Byonic\2MC_20171219_004_D.wiff_Byonic.byrslt (v2.9-30)
File Edit Window Help Byonic
Proteins
Peptide List (double click to dock / undock)
Peptides
Prot. Rank | Pos. | Sequence | Glycans | Protein Name | Score | Cleavage | z | Obs. m/z | Calc. m/z | Mods (variable) | PEP 2D | |Log Prob| | Delta Score | PEP 1D | Delta Mod Score | ppm err. | Off-By-X
5 | 68 | C.LCQYVK[+162.05282]DPNYGHYVSSP.H | Hex(1) | >HORVU4Hr1G089500.1 gene=HORVU4Hr1G089500 | 560.6 | Non | 3 | 711.3132 | 711.3262 | K6(xgly / 162.0528) | 1.1e-07 | 6.96 | 416.4 | 5.3e-07 | 416.4 | -18.28
Spectrum (double click to dock / undock)
C.LCQYVK[+162]DPNYGHYVSSP.H z=3,scan#=sample=1 period=1 cycle=508 experiment=15,scan time=10.1152
LCQYVKDPNYGHYVSSP
Intensity
m/z
m/z errors, tol=75.0 ppm
Obs - Calc m/z
Protein Coverage
Sample name
Protein name
11:35 AM
18/08/2020

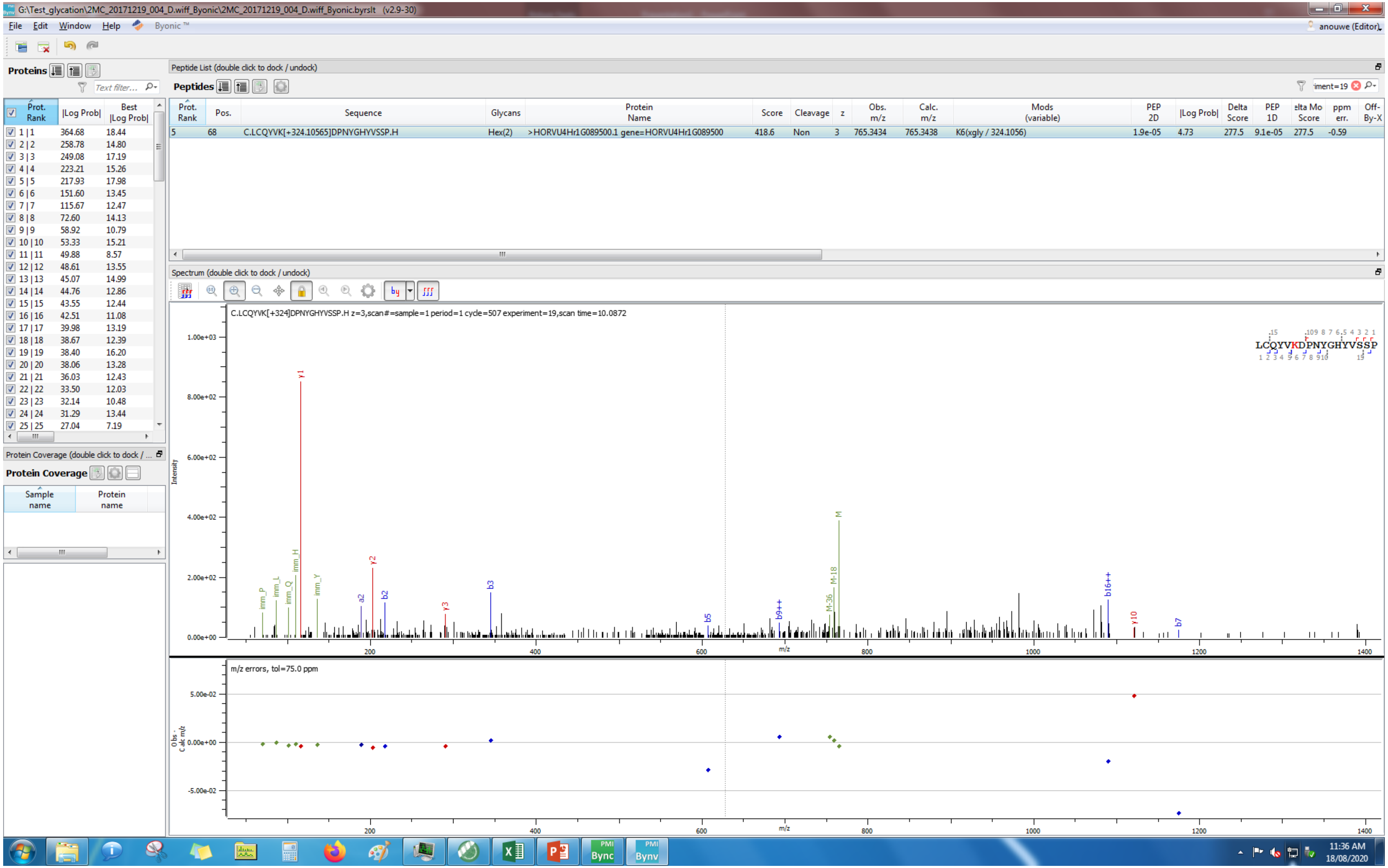
G:\Test_glycation\2MC_20171219_004_D.wiff_Byonic\2MC_20171219_004_D.wiff_Byonic.byrslt (v2.9-30)
File Edit Window Help Byonic
anouwe (Editor)
Proteins
Peptide List (double click to dock / undock)
Peptides
Prot. Rank | Pos. | Sequence | Glycans | Protein Name | Score | Cleavage | z | Obs. m/z | Calc. m/z | Mods (variable) | PEP 2D | |Log Prob| | Delta Score | PEP 1D | elta Mo Score | ppm err. | Off-By-X
5 | 68 | C.LCQYVK[+324.10565]DPNYGHYVSSP.H | Hex(2) | >HORVU4Hr1G089500.1 gene=HORVU4Hr1G089500 | 418.6 | Non | 3 | 765.3434 | 765.3438 | K6(xgly / 324.1056) | 1.9e-05 | 4.73 | 277.5 | 9.1e-05 | 277.5 | -0.59
Spectrum (double click to dock / undock)
C.LCQYVK[+324]DPNYGHYVSSP.H z=3,scan#=sample=1 period=1 cycle=507 experiment=19,scan time=10.0872
LCQYVKDPNYGHYVSSP
m/z errors, tol=75.0 ppm
Protein Coverage (double click to dock / undock)
Protein Coverage
Sample name
Protein name
11:36 AM
18/08/2020

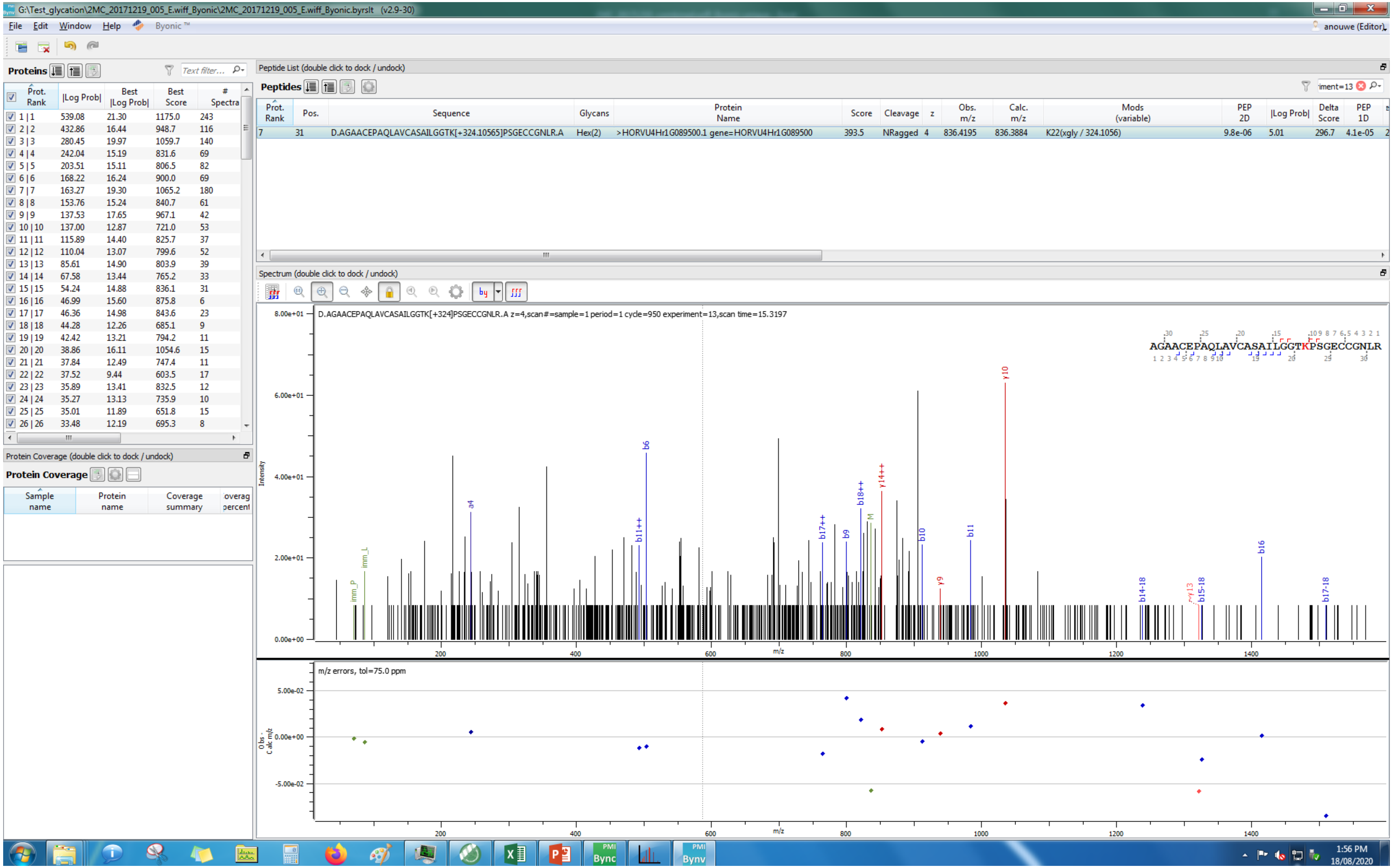

G:\Test_glycation\2MC_20171219_005_E.wiff_Byonic\2MC_20171219_005_E.wiff_Byonic.byrslt (v2.9-30)
File Edit Window Help Byonic ™
anouwe (Editor)
Proteins
Prot. Rank | |Log Prob| | Best |Log Prob| | Best Score | # Spectra
1 | 1 539.08 21.30 1175.0 243
2 | 2 432.86 16.44 948.7 116
3 | 3 280.45 19.97 1059.7 140
4 | 4 242.04 15.19 831.6 69
5 | 5 203.51 15.11 806.5 82
6 | 6 168.22 16.24 900.0 69
7 | 7 163.27 19.30 1065.2 180
8 | 8 153.76 15.24 840.7 61
9 | 9 137.53 17.65 967.1 42
10 | 10 137.00 12.87 721.0 53
11 | 11 115.89 14.40 825.7 37
12 | 12 110.04 13.07 799.6 52
13 | 13 85.61 14.90 803.9 39
14 | 14 67.58 13.44 765.2 33
15 | 15 54.24 14.88 836.1 31
16 | 16 46.99 15.60 875.8 6
17 | 17 46.36 14.98 843.6 23
18 | 18 44.28 12.26 685.1 9
19 | 19 42.42 13.21 794.2 11
20 | 20 38.86 16.11 1054.6 15
21 | 21 37.84 12.49 747.4 11
22 | 22 37.52 9.44 603.5 17
23 | 23 35.89 13.41 832.5 12
24 | 24 35.27 13.13 735.9 10
25 | 25 35.01 11.89 651.8 15
26 | 26 33.48 12.19 695.3 8
Protein Coverage (double click to dock / undock)
Protein Coverage
Sample name | Protein name | Coverage summary | overag percent
Peptide List (double click to dock / undock)
Peptides
Prot. Rank | Pos. | Sequence | Glycans | Protein Name | Score | Cleavage | z | Obs. m/z | Calc. m/z | Mods (variable) | PEP 2D | |Log Prob| | Delta Score | PEP 1D
7 | 31 | D.AGAACEPAQLAVCASAILGGTK[+324.10565]PSGECCGNLR.A | Hex(2) | >HORVU4Hr1G089500.1 gene=HORVU4Hr1G089500 | 393.5 | NRagged | 4 | 836.4195 | 836.3884 | K22(xgly / 324.1056) | 9.8e-06 | 5.01 | 296.7 | 4.1e-05
Spectrum (double click to dock / undock)
D.AGAACEPAQLAVCASAILGGTK[+324]PSGECCGNLR.A z=4,scan#=sample=1 period=1 cycle=950 experiment=13,scan time=15.3197
AGAACEPAQLAVCASAILGGTKPSGECCGNLR
Intensity
m/z
m/z errors, tol=75.0 ppm
Obs - Calc m/z
1:56 PM
18/08/2020

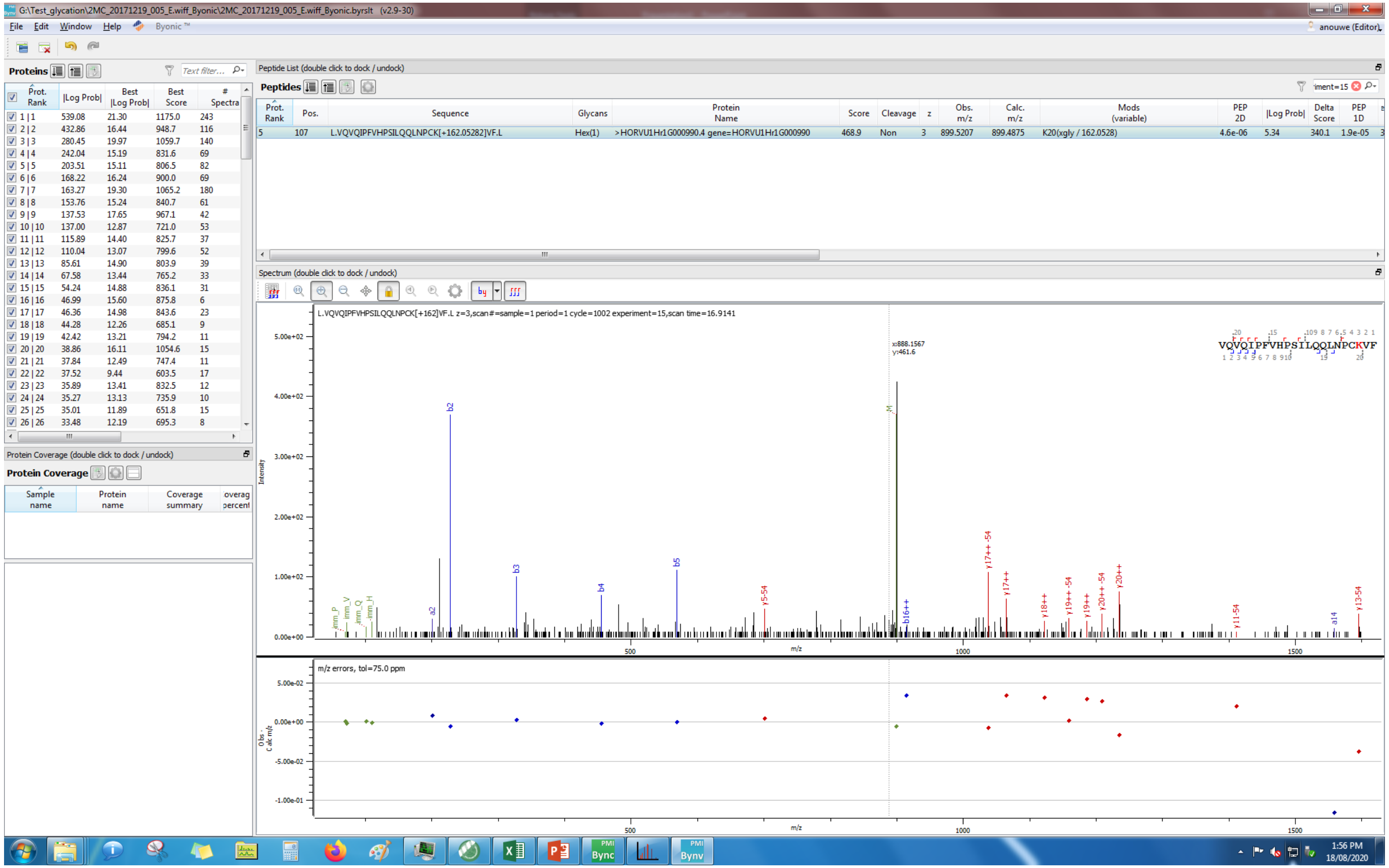
G:\Test_glycation\2MC_20171219_005_E.wiff_Byonic\2MC_20171219_005_E.wiff_Byonic.byrslt (v2.9-30)
File Edit Window Help Byonic™
Proteins
Peptide List (double click to dock / undock)
Peptides
5 107 L.VQVQIPFVHPSILQQLNPCK[+162.05282]VF.L Hex(1) >HORVU1Hr1G000990.4 gene=HORVU1Hr1G000990 468.9 Non 3 899.5207 899.4875 K20(xgly / 162.0528) 4.6e-06 5.34 340.1 1.9e-05
Spectrum (double click to dock / undock)
L.VQVQIPFVHPSILQQLNPCK[+162]VF.L z=3,scan#=sample=1 period=1 cycle=1002 experiment=15,scan time=16.9141
Protein Coverage (double click to dock / undock)
Protein Coverage
m/z errors, tol=75.0 ppm
1:56 PM
18/08/2020

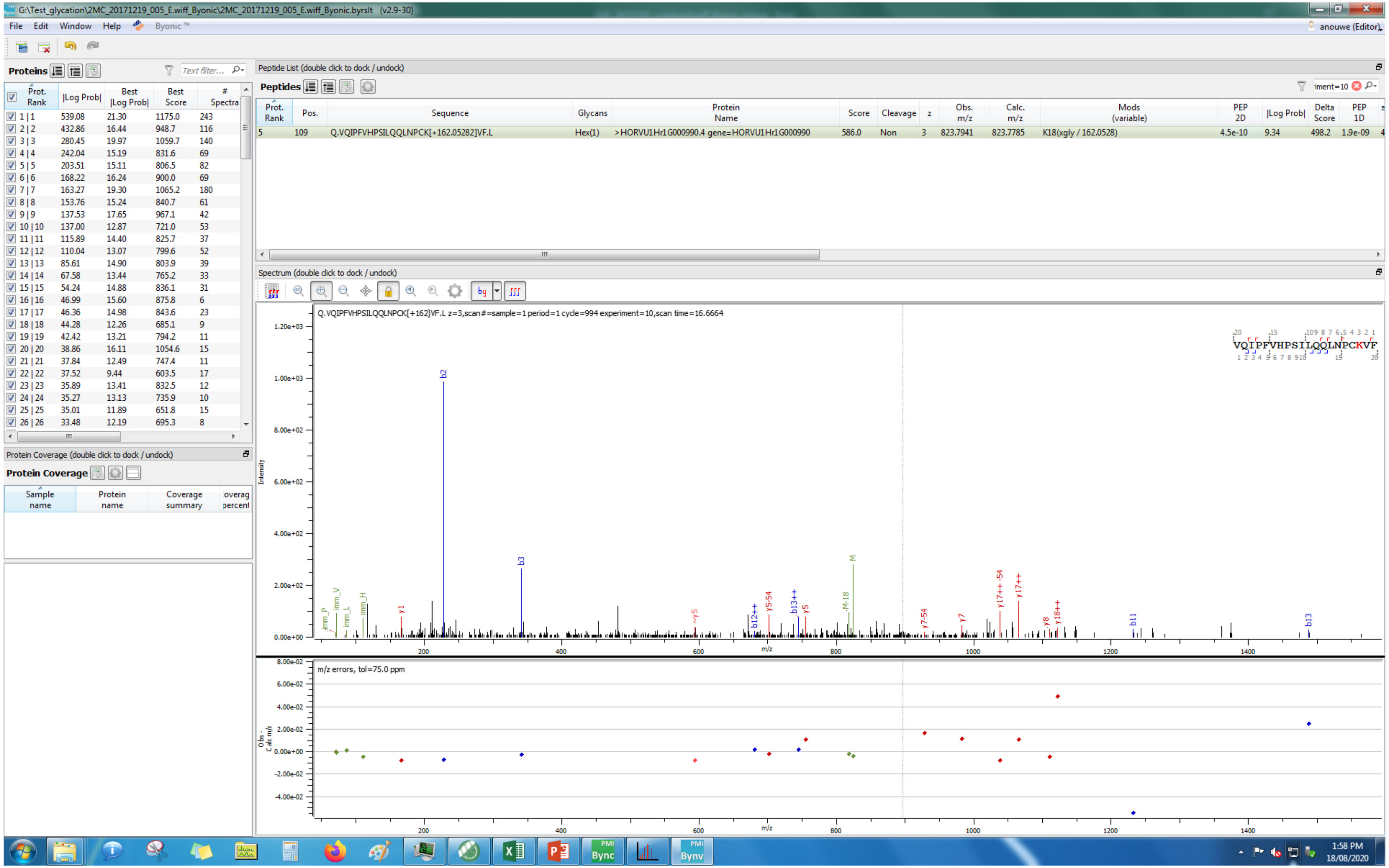
G:\Test_glycation\2MC_20171219_005_E.wiff_Byonic\2MC_20171219_005_E.wiff_Byonic.byrslt (v2.9-30)
File Edit Window Help Byonic
Proteins
Peptide List (double click to dock / undock)
Peptides
5 109 Q.VQIPFVHPSILQQLNPCK[+162.05282]VF.L Hex(1) >HORVU1Hr1G000990.4 gene=HORVU1Hr1G000990 586.0 Non 3 823.7941 823.7785 K18(xgly / 162.0528) 4.5e-10 9.34 498.2 1.9e-09
Spectrum (double click to dock / undock)
Q.VQIPFVHPSILQQLNPCK[+162]VF.L z=3,scan#=sample=1 period=1 cycle=994 experiment=10,scan time=16.6664
VQIPFVHPSILQQLNPCKVF
m/z errors, tol=75.0 ppm
Protein Coverage (double click to dock / undock)
Protein Coverage
Sample name
Protein name
Coverage summary
1:58 PM
18/08/2020

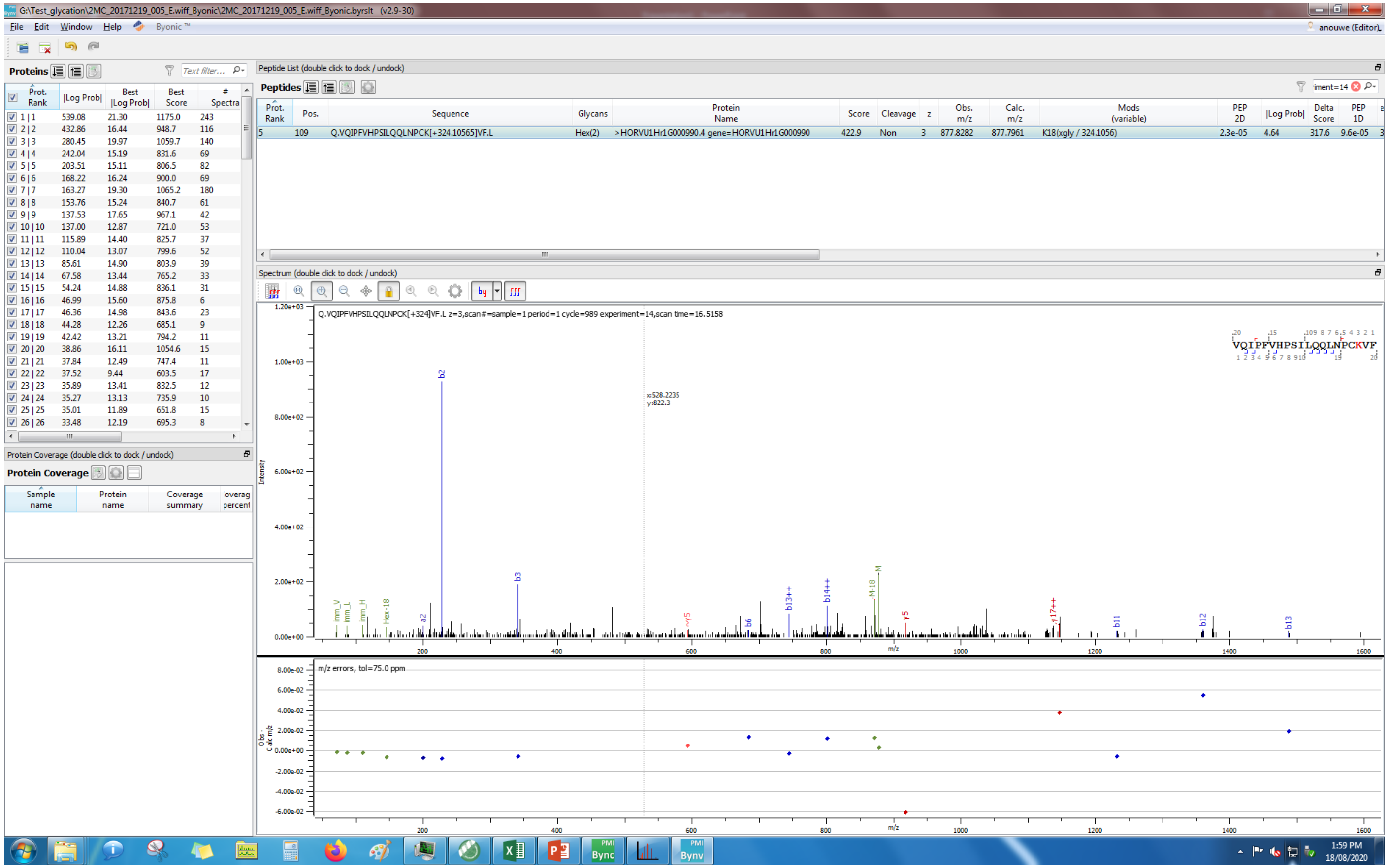
G:\Test_glycation\2MC_20171219_005_E.wiff_Byonic\2MC_20171219_005_E.wiff_Byonic.byrslt (v2.9-30)
File Edit Window Help Byonic™
anouwe (Editor)
Proteins
Text filter...
Prot. Rank |Log Prob| Best |Log Prob| Best Score # Spectra
1 | 1 539.08 21.30 1175.0 243
2 | 2 432.86 16.44 948.7 116
3 | 3 280.45 19.97 1059.7 140
4 | 4 242.04 15.19 831.6 69
5 | 5 203.51 15.11 806.5 82
6 | 6 168.22 16.24 900.0 69
7 | 7 163.27 19.30 1065.2 180
8 | 8 153.76 15.24 840.7 61
9 | 9 137.53 17.65 967.1 42
Peptide List (double click to dock / undock)
Peptides
iment=14
Prot. Rank Pos. Sequence Glycans Protein Name Score Cleavage z Obs. m/z Calc. m/z Mods (variable) PEP 2D |Log Prob| Delta Score PEP 1D
5 109 Q.VQIPFVHPSILQQLNPCK[+324.10565]VF.L Hex(2) >HORVU1Hr1G000990.4 gene=HORVU1Hr1G000990 422.9 Non 3 877.8282 877.7961 K18(xgly / 324.1056) 2.3e-05 4.64 317.6 9.6e-05
Protein Coverage (double click to dock / undock)
Protein Coverage
Sample name Protein name Coverage summary
Spectrum (double click to dock / undock)
Q.VQIPFVHPSILQQLNPCK[+324]VF.L z=3,scan#=sample=1 period=1 cycle=989 experiment=14,scan time=16.5158
VQIPFVHPSILQQLNPCKVF
x:528.2235
y:822.3
Intensity
m/z
m/z errors, tol=75.0 ppm
1:59 PM
18/08/2020

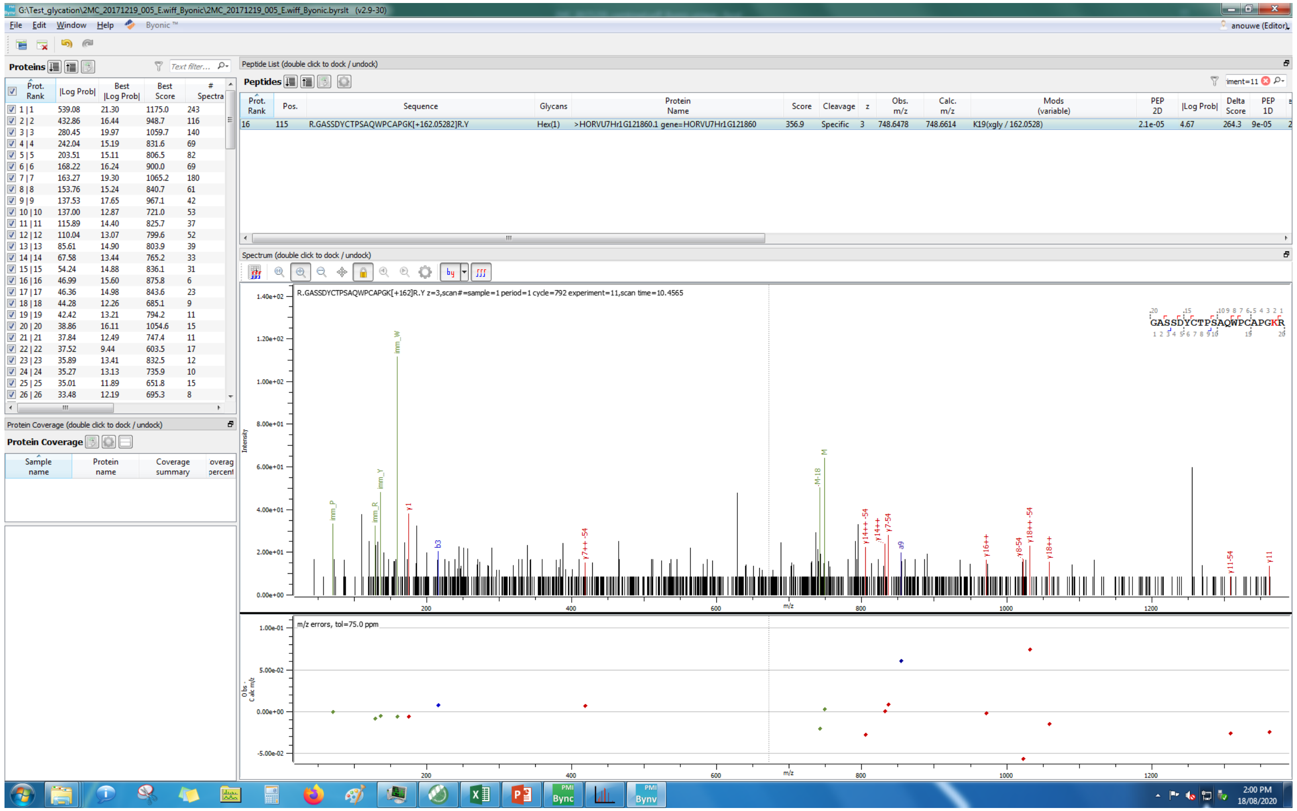

Proteins
Peptide List (double click to dock / undock)
Peptides
R.GASSDYCTPSAQWPCAPGK[+162.05282]R.Y
Spectrum (double click to dock / undock)
Protein Coverage

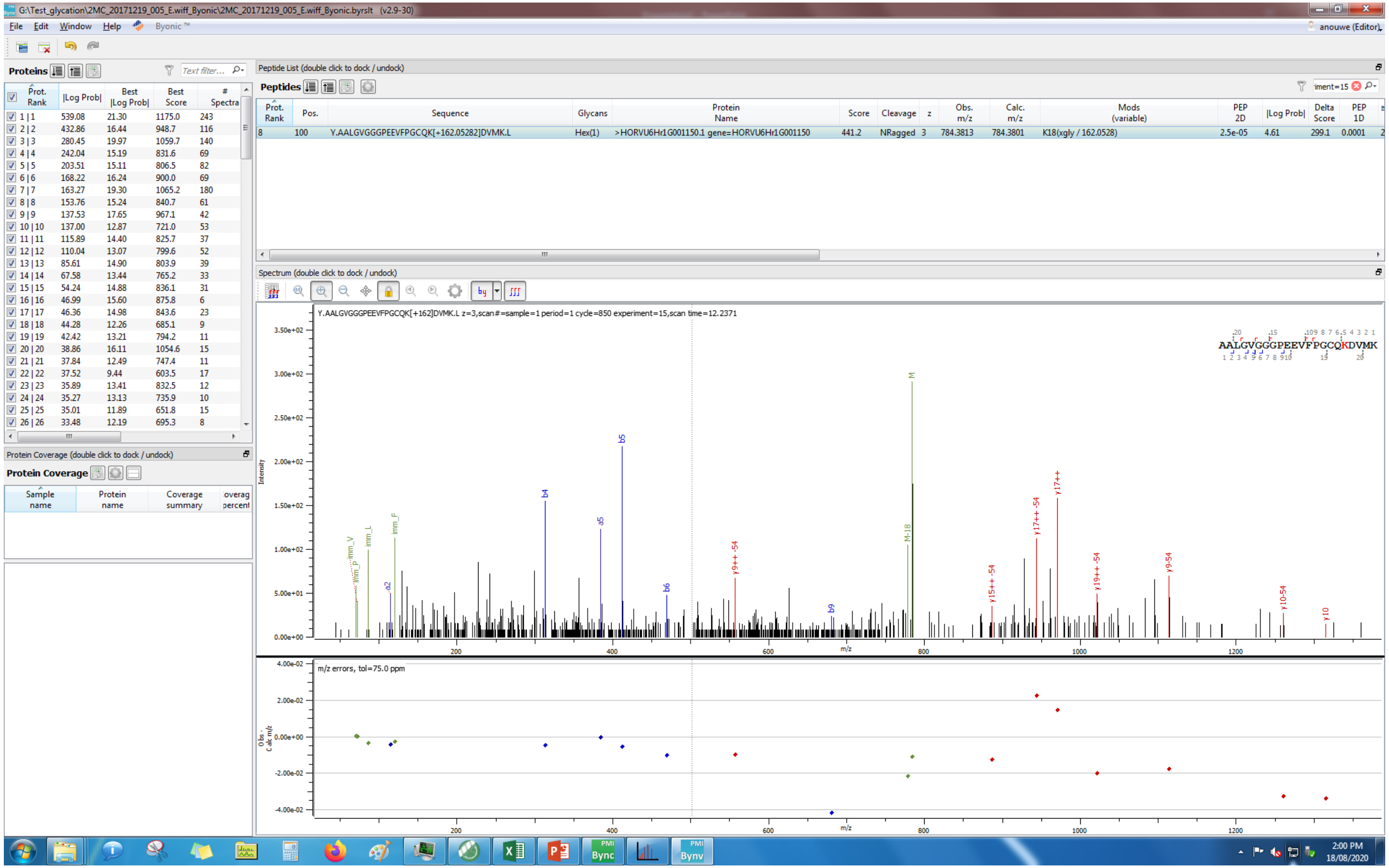
G:\Test_glycation\2MC_20171219_005_E.wiff_Byonic\2MC_20171219_005_E.wiff_Byonic.byrslt (v2.9-30)
File Edit Window Help Byonic™
Proteins
Peptide List (double click to dock / undock)
Peptides
Y.AALGVGGGPEEVFPGCQK[+162.05282]DVMK.L
Hex(1)
>HORVU6Hr1G001150.1 gene=HORVU6Hr1G001150
K18(xgly / 162.0528)
Spectrum (double click to dock / undock)
Y.AALGVGGGPEEVFPGCQK[+162]DVMK.L z=3,scan#=sample=1 period=1 cycle=850 experiment=15,scan time=12.2371
Protein Coverage (double click to dock / undock)
Protein Coverage
m/z errors, tol=75.0 ppm
Intensity
m/z
2:00 PM
18/08/2020

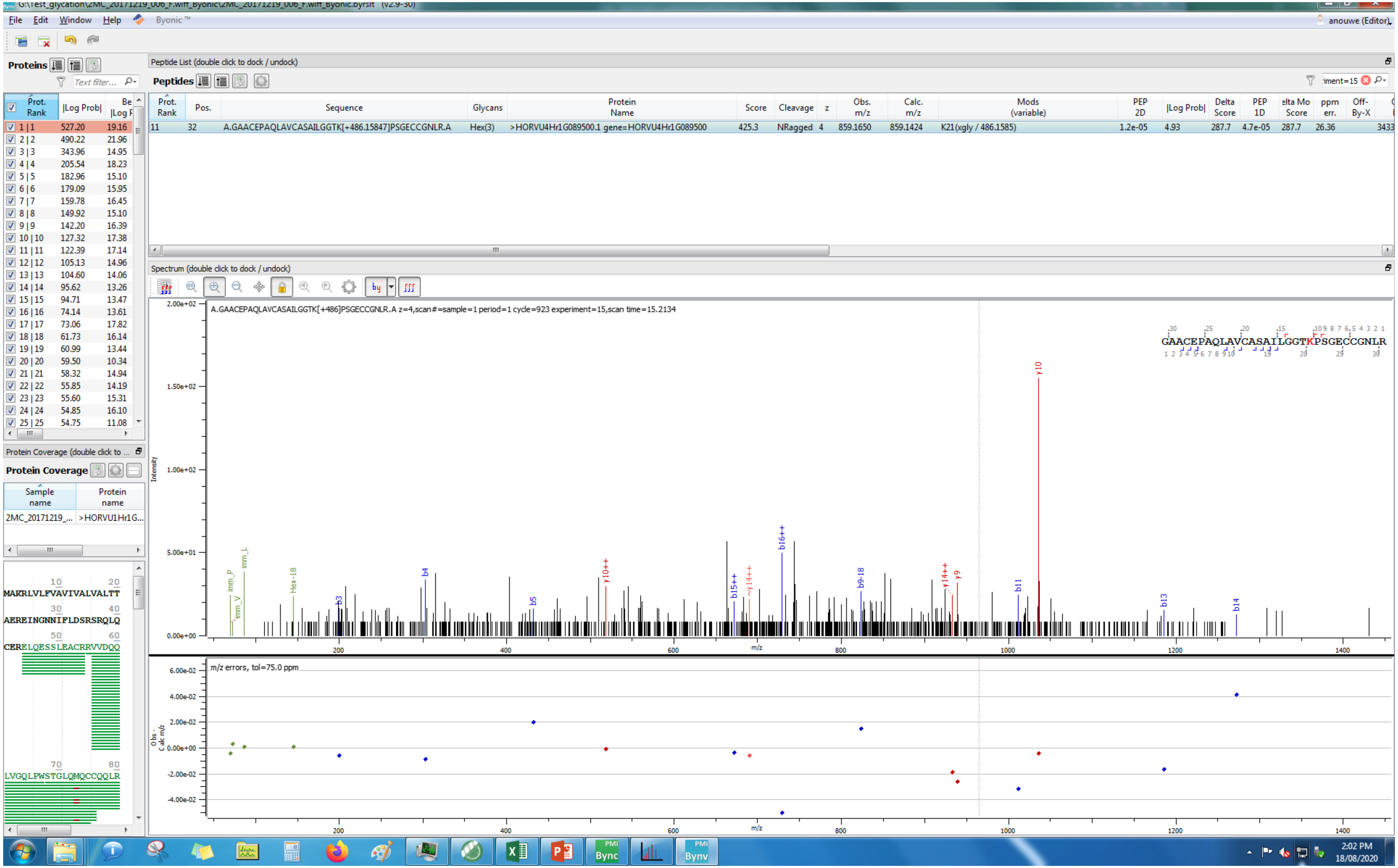
File Edit Window Help Byonic™
anouwe (Editor)
Proteins
Peptide List (double click to dock / undock)
Peptides
Prot. Rank | Pos. | Sequence | Glycans | Protein Name | Score | Cleavage | z | Obs. m/z | Calc. m/z | Mods (variable) | PEP 2D | |Log Prob| | Delta Score | PEP 1D | Delta Mod Score | ppm err. | Off-By-X
11 32 A.GAACEPAQLAVCASAILGGTK[+486.15847]PSGECCGNLR.A Hex(3) >HORVU4Hr1G089500.1 gene=HORVU4Hr1G089500 425.3 NRagged 4 859.1650 859.1424 K21(xgly / 486.1585) 1.2e-05 4.93 287.7 4.7e-05 287.7 26.36
Spectrum (double click to dock / undock)
A.GAACEPAQLAVCASAILGGTK[+486]PSGECCGNLR.A z=4,scan#=sample=1 period=1 cycle=923 experiment=15,scan time=15.2134
GAACEPAQLAVCASAILGGTKPSGECCGNLR
Intensity
m/z
y10
b16++
b4
b5
b3
b9-18
b11
b13
b14
y9
Hex-18
m/z errors, tol=75.0 ppm
Obs - Calc m/z
Protein Coverage (double click to dock / undock)
Protein Coverage
Sample name | Protein name
2MC_20171219_... | >HORVU1Hr1G...
MAKRLVLFVAVIVALVALTT
AEREINGNNIFLDSRSRQLQ
CERELQESSLEACRRVVDQQ
LVGQLPWSTGLQMQCCQQLR
2:02 PM
18/08/2020

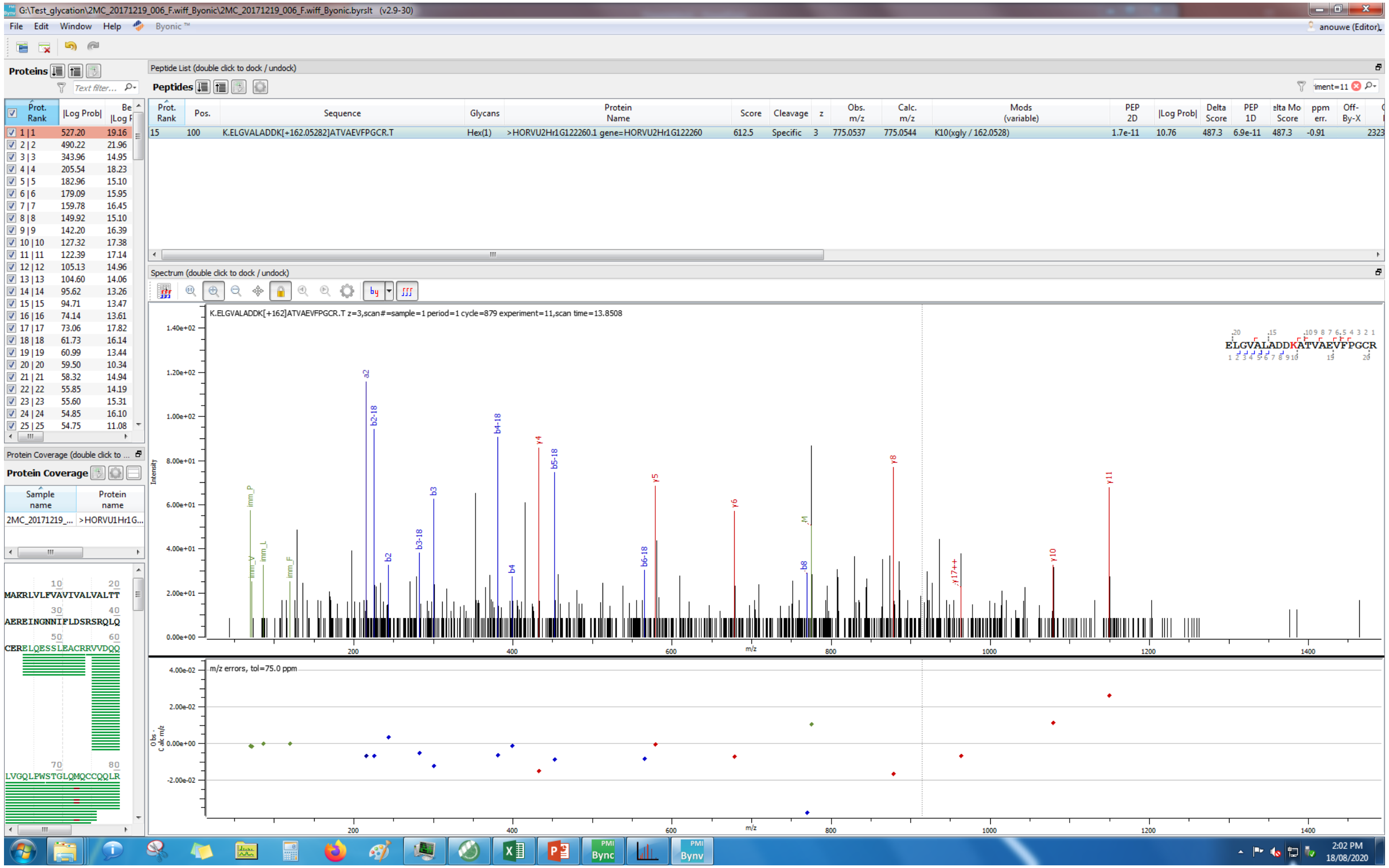
G:\Test_glycation\2MC_20171219_006_F.wiff_Byonic\2MC_20171219_006_F.wiff_Byonic.byrslt (v2.9-30)
File Edit Window Help Byonic™
anouwe (Editor)
Proteins
Peptide List (double click to dock / undock)
Peptides
Prot. Rank | Pos. | Sequence | Glycans | Protein Name | Score | Cleavage | z | Obs. m/z | Calc. m/z | Mods (variable) | PEP 2D | |Log Prob| | Delta Score | PEP 1D | Delta Mod Score | ppm err. | Off-By-X
15 | 100 | K.ELGVALADDK[+162.05282]ATVAEVFPGCR.T | Hex(1) | >HORVU2Hr1G122260.1 gene=HORVU2Hr1G122260 | 612.5 | Specific | 3 | 775.0537 | 775.0544 | K10(xgly / 162.0528) | 1.7e-11 | 10.76 | 487.3 | 6.9e-11 | 487.3 | -0.91
Spectrum (double click to dock / undock)
K.ELGVALADDK[+162]ATVAEVFPGCR.T z=3,scan#=sample=1 period=1 cycle=879 experiment=11,scan time=13.8508
ELGVALADDKATVAEVFPGCR
Intensity
m/z
m/z errors, tol=75.0 ppm
Protein Coverage
Sample name | Protein name
2MC_20171219_... | >HORVU1Hr1G...
MAKRLVLFVAVIVALVALTT
AEREINGNNIFLDSRSRQLQ
CERELQESSLEACRRVVDQQ
LVGQLPWSTGLQMQCCQQLR
2:02 PM
18/08/2020

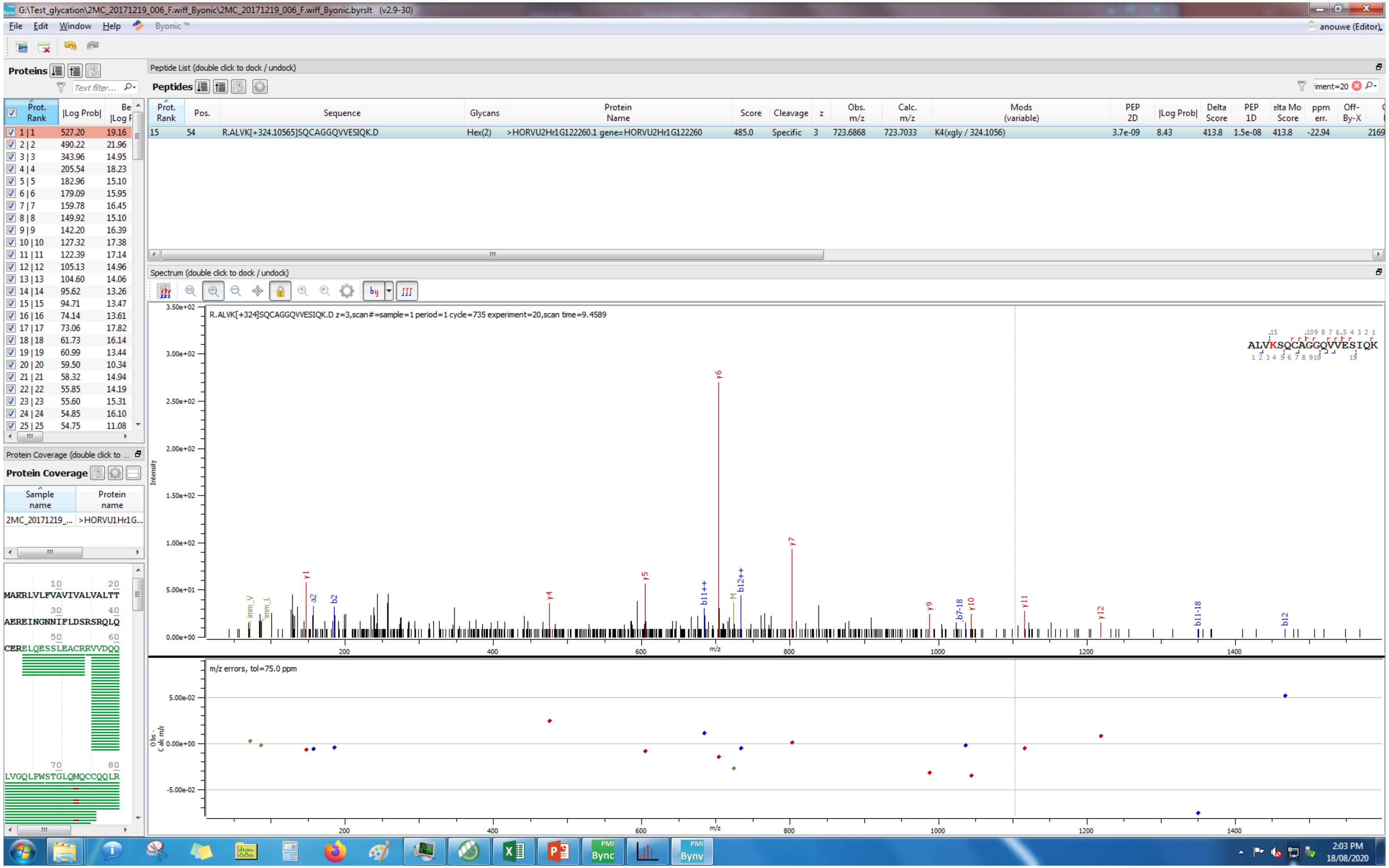
G:\Test_glycation\2MC_20171219_006_F.wiff_Byonic\2MC_20171219_006_F.wiff_Byonic.byrslt (v2.9-30)
File Edit Window Help Byonic™
anouwe (Editor)
Proteins
Peptide List (double click to dock / undock)
Peptides
Prot. Rank | Pos. | Sequence | Glycans | Protein Name | Score | Cleavage | z | Obs. m/z | Calc. m/z | Mods (variable) | PEP 2D | |Log Prob| | Delta Score | PEP 1D | Delta Mod Score | ppm err. | Off-By-X
15 | 54 | R.ALVK[+324.10565]SQCAGGQVVESIQK.D | Hex(2) | >HORVU2Hr1G122260.1 gene=HORVU2Hr1G122260 | 485.0 | Specific | 3 | 723.6868 | 723.7033 | K4(xgly / 324.1056) | 3.7e-09 | 8.43 | 413.8 | 1.5e-08 | 413.8 | -22.94
Spectrum (double click to dock / undock)
R.ALVK[+324]SQCAGGQVVESIQK.D z=3,scan#=sample=1 period=1 cycle=735 experiment=20,scan time=9.4589
ALVKSQCAGGQVVESIQK
m/z errors, tol=75.0 ppm
Protein Coverage
Sample name | Protein name
2MC_20171219_... | >HORVU1Hr1G...
MAKRLVLFVAVIVALVALTT
AEREINGNNIFLDSRSRQLQ
CERELQESSLEACRRVVDQQ
LVGQLPWSTGLQMQCCQQLR
2:03 PM 18/08/2020

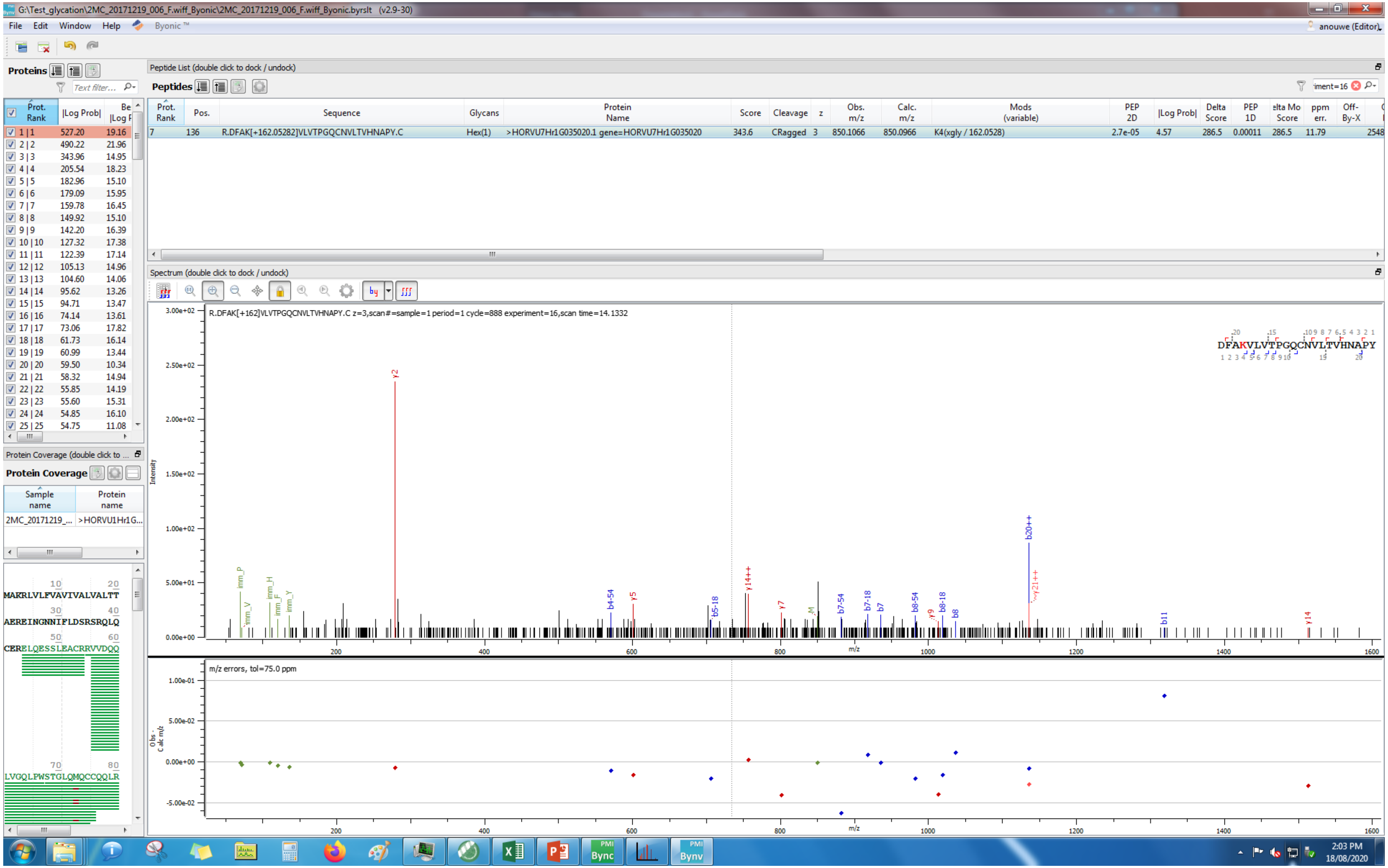

G:\Test_glycation\2MC_20171219_006_F.wiff_Byonic\2MC_20171219_006_F.wiff_Byonic.byrslt (v2.9-30)
File Edit Window Help Byonic™
Proteins
Peptide List (double click to dock / undock)
Peptides
Prot. Rank Pos. Sequence Glycans Protein Name Score Cleavage z Obs. m/z Calc. m/z Mods (variable) PEP 2D |Log Prob| Delta Score PEP 1D Delta Mod Score ppm err. Off-By-X
7 136 R.DFAK[+162.05282]VLVTPGQCNVLTVHNAPY.C Hex(1) >HORVU7Hr1G035020.1 gene=HORVU7Hr1G035020 343.6 CRagged 3 850.1066 850.0966 K4(xgly / 162.0528) 2.7e-05 4.57 286.5 0.00011 286.5 11.79
Spectrum (double click to dock / undock)
R.DFAK[+162]VLVTPGQCNVLTVHNAPY.C z=3,scan#=sample=1 period=1 cycle=888 experiment=16,scan time=14.1332
DFAKVLVTPGQCNVLTVHNAPY
Intensity
m/z
m/z errors, tol=75.0 ppm
Protein Coverage
Sample name
Protein name
2MC_20171219_...
>HORVU1Hr1G...
MAKRLVLFVAVIVALVALTT
AEREINGNNIFLDSRSRQLQ
CERELQESSLEACRRVVDQQ
LVGQLPWSTGLQMQCCQQLR
2:03 PM
18/08/2020

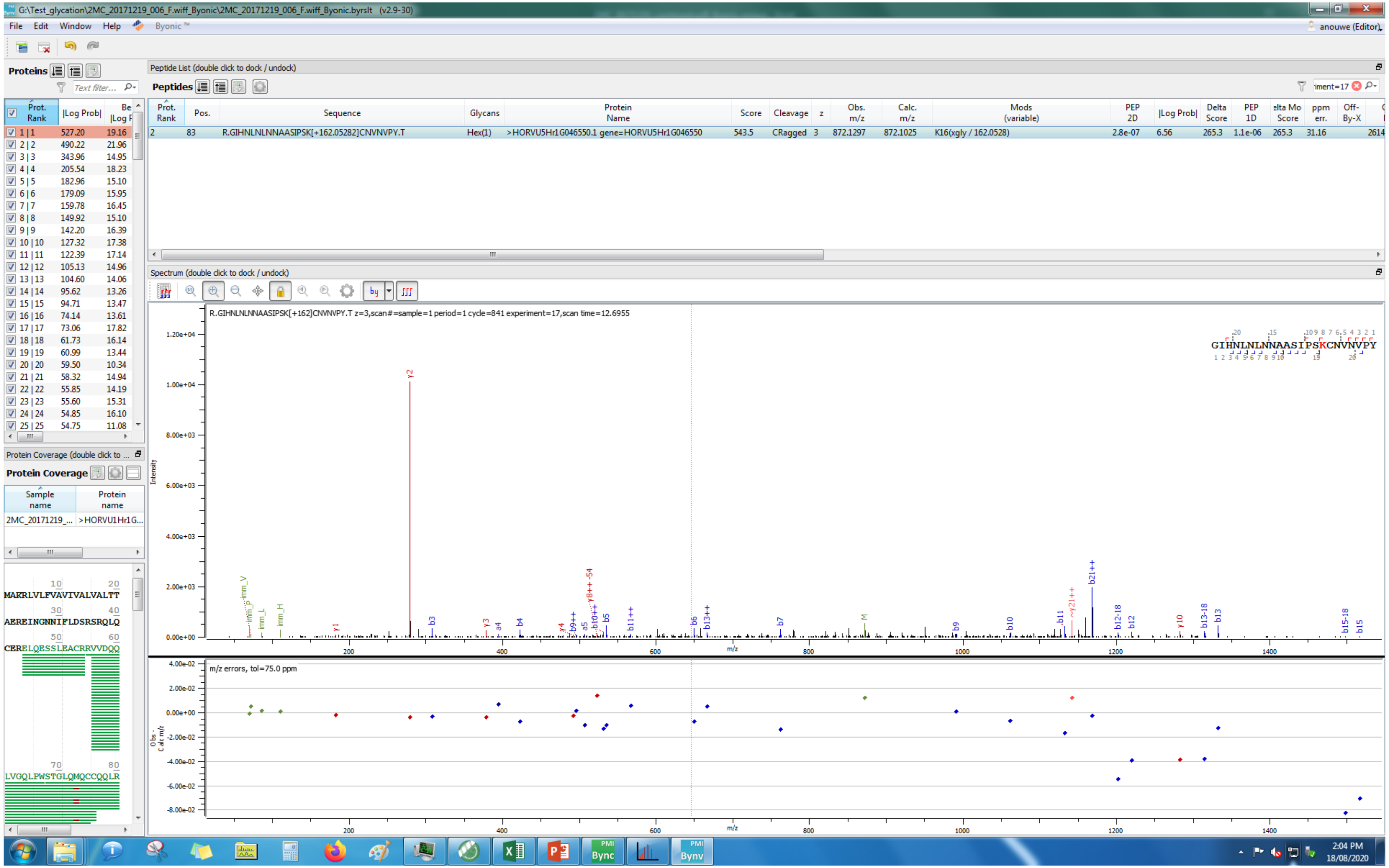
G:\Test_glycation\2MC_20171219_006_F.wiff_Byonic\2MC_20171219_006_F.wiff_Byonic.byrslt (v2.9-30)
File Edit Window Help Byonic™
anouwe (Editor)
Proteins
Peptide List (double click to dock / undock)
Peptides
Prot. Rank | Pos. | Sequence | Glycans | Protein Name | Score | Cleavage | z | Obs. m/z | Calc. m/z | Mods (variable) | PEP 2D | |Log Prob| | Delta Score | PEP 1D | Delta Mod Score | ppm err. | Off-By-X
2 | 83 | R.GIHNLNLNNAASIPSK[+162.05282]CNVNPY.T | Hex(1) | >HORVU5Hr1G046550.1 gene=HORVU5Hr1G046550 | 543.5 | CRagged | 3 | 872.1297 | 872.1025 | K16(xgly / 162.0528) | 2.8e-07 | 6.56 | 265.3 | 1.1e-06 | 265.3 | 31.16 | 2614
Spectrum (double click to dock / undock)
R.GIHNLNLNNAASIPSK[+162]CNVNPY.T z=3,scan#=sample=1 period=1 cycle=841 experiment=17,scan time=12.6955
GIHNLNLNNAASIPSKCNVNPY
Intensity
m/z
m/z errors, tol=75.0 ppm
Obs. - Calc m/z
Protein Coverage (double click to ...
Protein Coverage
Sample name | Protein name
2MC_20171219_... | >HORVU1Hr1G...
MAKRLVLFVAVIVALVALTT
AEREINGNNIFLDSRSRQLQ
CERELQESSLEACRRVVDQQ
LVGQLPWSTGLQMQCCQQLR
2:04 PM
18/08/2020

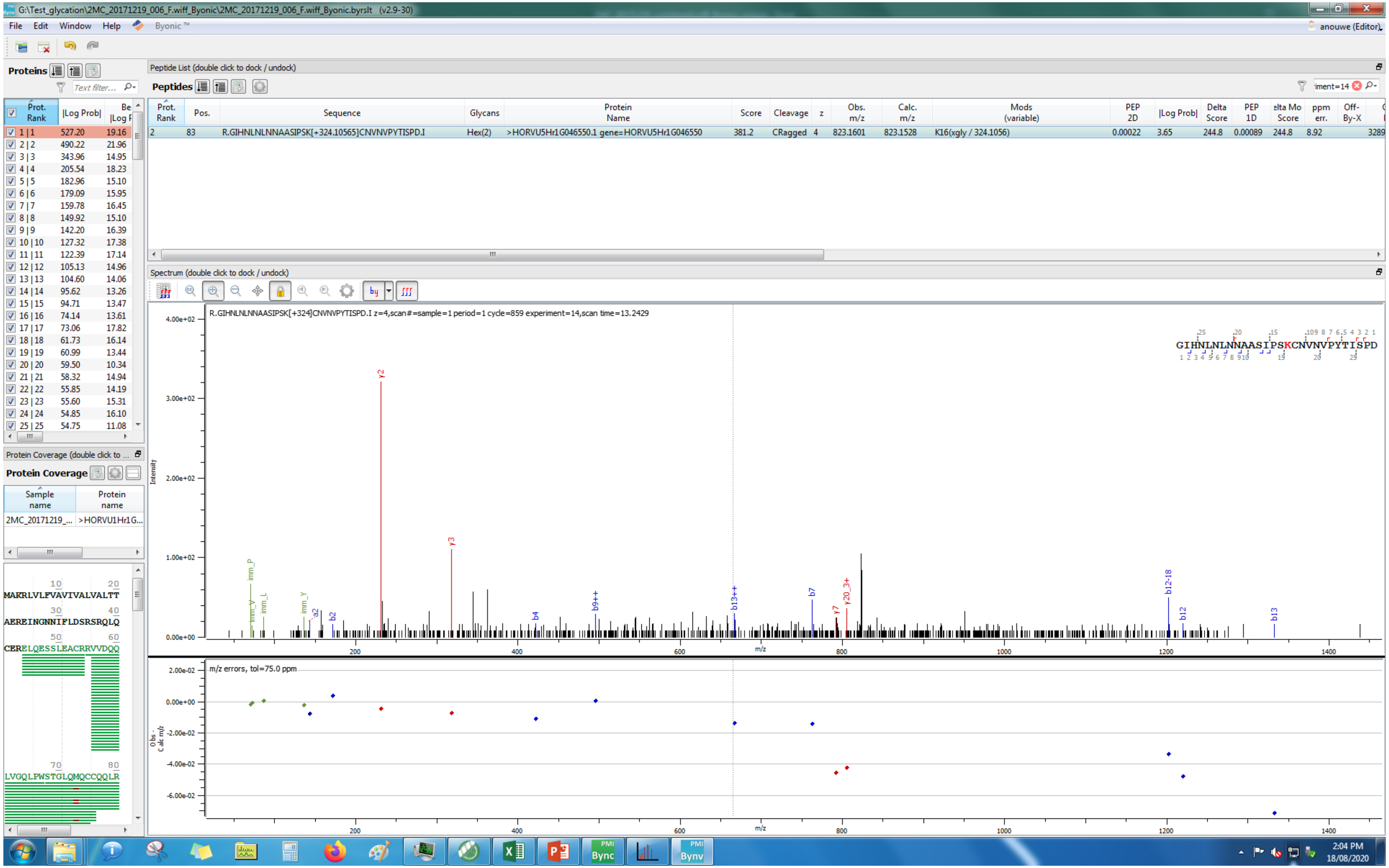
G:\Test_glycation\2MC_20171219_006_F.wiff_Byonic\2MC_20171219_006_F.wiff_Byonic.byrslt (v2.9-30)
File Edit Window Help Byonic™
anouwe (Editor)
Proteins
Peptide List (double click to dock / undock)
Peptides
Prot. Rank | Pos. | Sequence | Glycans | Protein Name | Score | Cleavage | z | Obs. m/z | Calc. m/z | Mods (variable) | PEP 2D | |Log Prob| | Delta Score | PEP 1D | Delta Mod Score | ppm err. | Off-By-X
2 | 83 | R.GIHNLNLNNAASIPSK[+324.10565]CNVNVPYTISPD.I | Hex(2) | >HORVU5Hr1G046550.1 gene=HORVU5Hr1G046550 | 381.2 | CRagged | 4 | 823.1601 | 823.1528 | K16(xgly / 324.1056) | 0.00022 | 3.65 | 244.8 | 0.00089 | 244.8 | 8.92 | 3289
Spectrum (double click to dock / undock)
R.GIHNLNLNNAASIPSK[+324]CNVNVPYTISPD.I z=4,scan#=sample=1 period=1 cycle=859 experiment=14,scan time=13.2429
GIHNLNLNNAASIPSKCNVNVPYTISPD
y2
y3
b2
b4
b9++
b13++
b7
y7
y20_3+
b12-18
b12
b13
imm_P
imm_L
imm_Y
a2
Intensity
m/z
m/z errors, tol=75.0 ppm
Obs - Calc m/z
Protein Coverage (double click to ...
Protein Coverage
Sample name | Protein name
2MC_20171219_... | >HORVU1Hr1G...
MAKRLVLFVAVIVALVALTT
AEREINGNNIFLDSRSRQLQ
CERELQESSLEACRRVVDQQ
LVGQLPWSTGLQMQCCQQLR
2:04 PM
18/08/2020

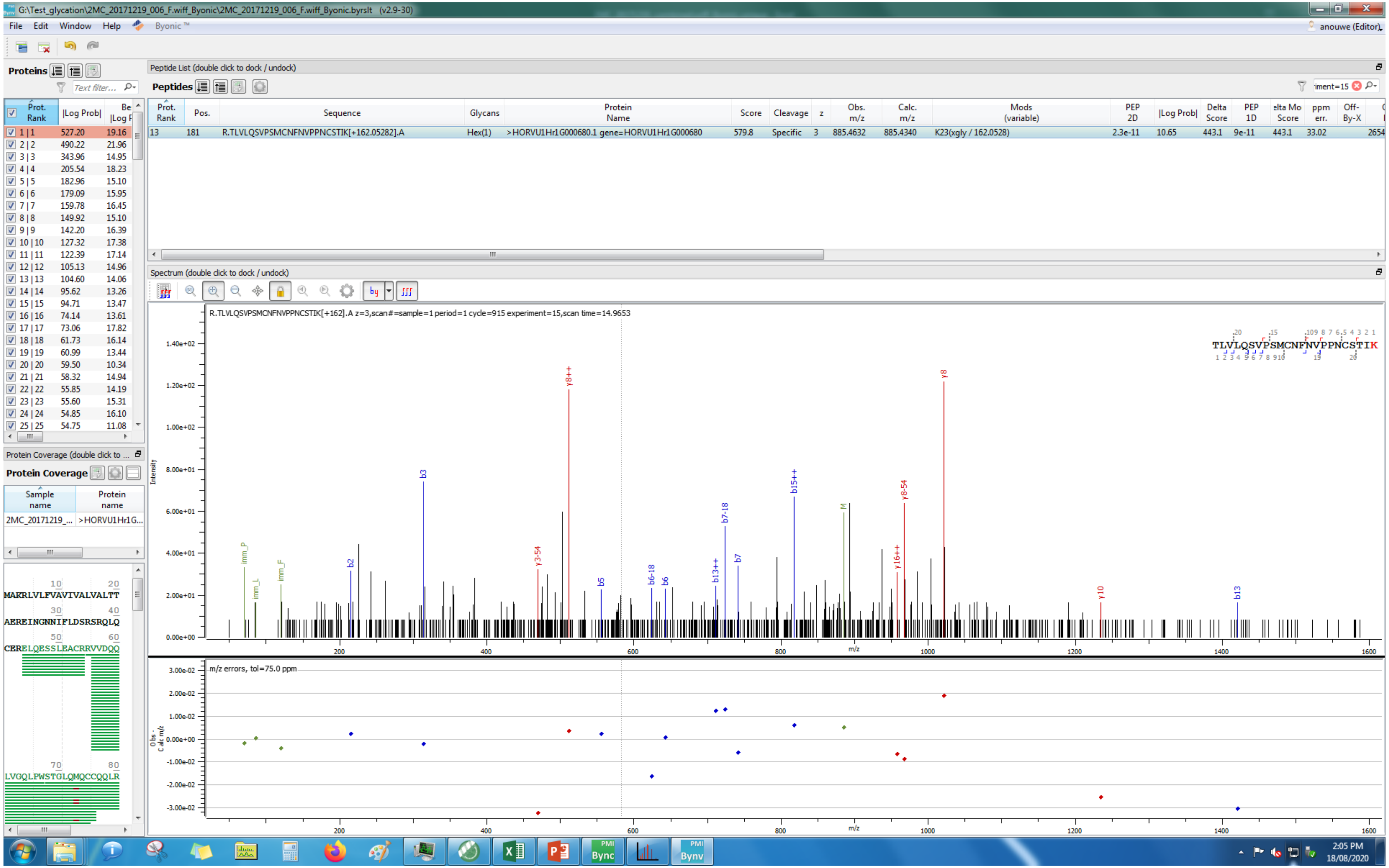
G:\Test_glycation\2MC_20171219_006_F.wiff_Byonic\2MC_20171219_006_F.wiff_Byonic.byrslt (v2.9-30)
File Edit Window Help Byonic™
Proteins
Peptide List (double click to dock / undock)
Peptides
R.TLVLQSVPSMCNFNVPPNCSTIK[+162.05282].A
Hex(1)
>HORVU1Hr1G000680.1 gene=HORVU1Hr1G000680
579.8
Specific
885.4632
885.4340
K23(xgly / 162.0528)
Spectrum (double click to dock / undock)
R.TLVLQSVPSMCNFNVPPNCSTIK[+162].A z=3,scan#=sample=1 period=1 cycle=915 experiment=15,scan time=14.9653
m/z errors, tol=75.0 ppm
Protein Coverage
2MC_20171219_...
>HORVU1Hr1G...
2:05 PM
18/08/2020

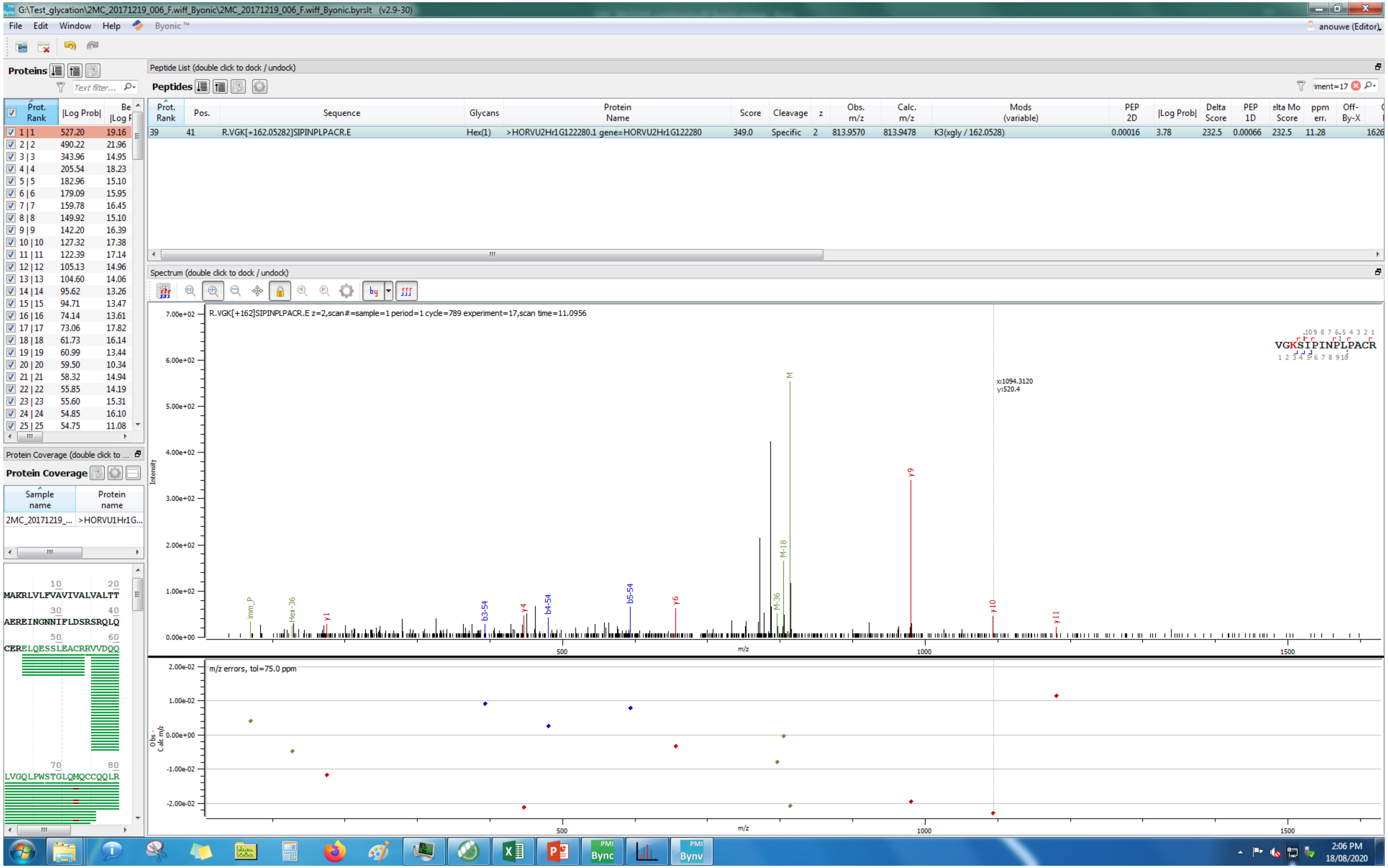
G:\Test_glycation\2MC_20171219_006_F.wiff_Byonic\2MC_20171219_006_F.wiff_Byonic.byrslt (v2.9-30)
File Edit Window Help Byonic™
Proteins
Peptide List (double click to dock / undock)
Peptides
Prot. Rank | Pos. | Sequence | Glycans | Protein Name | Score | Cleavage | z | Obs. m/z | Calc. m/z | Mods (variable) | PEP 2D | |Log Prob| | Delta Score | PEP 1D | Delta Mod Score | ppm err. | Off-By-X
39 | 41 | R.VGK[+162.05282]SIPINPLPACR.E | Hex(1) | >HORVU2Hr1G122280.1 gene=HORVU2Hr1G122280 | 349.0 | Specific | 2 | 813.9570 | 813.9478 | K3(xgly / 162.0528) | 0.00016 | 3.78 | 232.5 | 0.00066 | 232.5 | 11.28
Spectrum (double click to dock / undock)
R.VGK[+162]SIPINPLPACR.E z=2,scan#=sample=1 period=1 cycle=789 experiment=17,scan time=11.0956
VGKSIPINPLPACR
x:1094.3120
y:520.4
Intensity
m/z
m/z errors, tol=75.0 ppm
Protein Coverage (double click to dock / undock)
Protein Coverage
Sample name | Protein name
2MC_20171219_... | >HORVU1Hr1G...
MAKRLVLFVAVIVALVALTT
AEREINGNNIPLDSRSRQLQ
CERELQESSLEACRRVVDQQ
LVGQLPWSTGLQMQCCQQLR
2:06 PM
18/08/2020

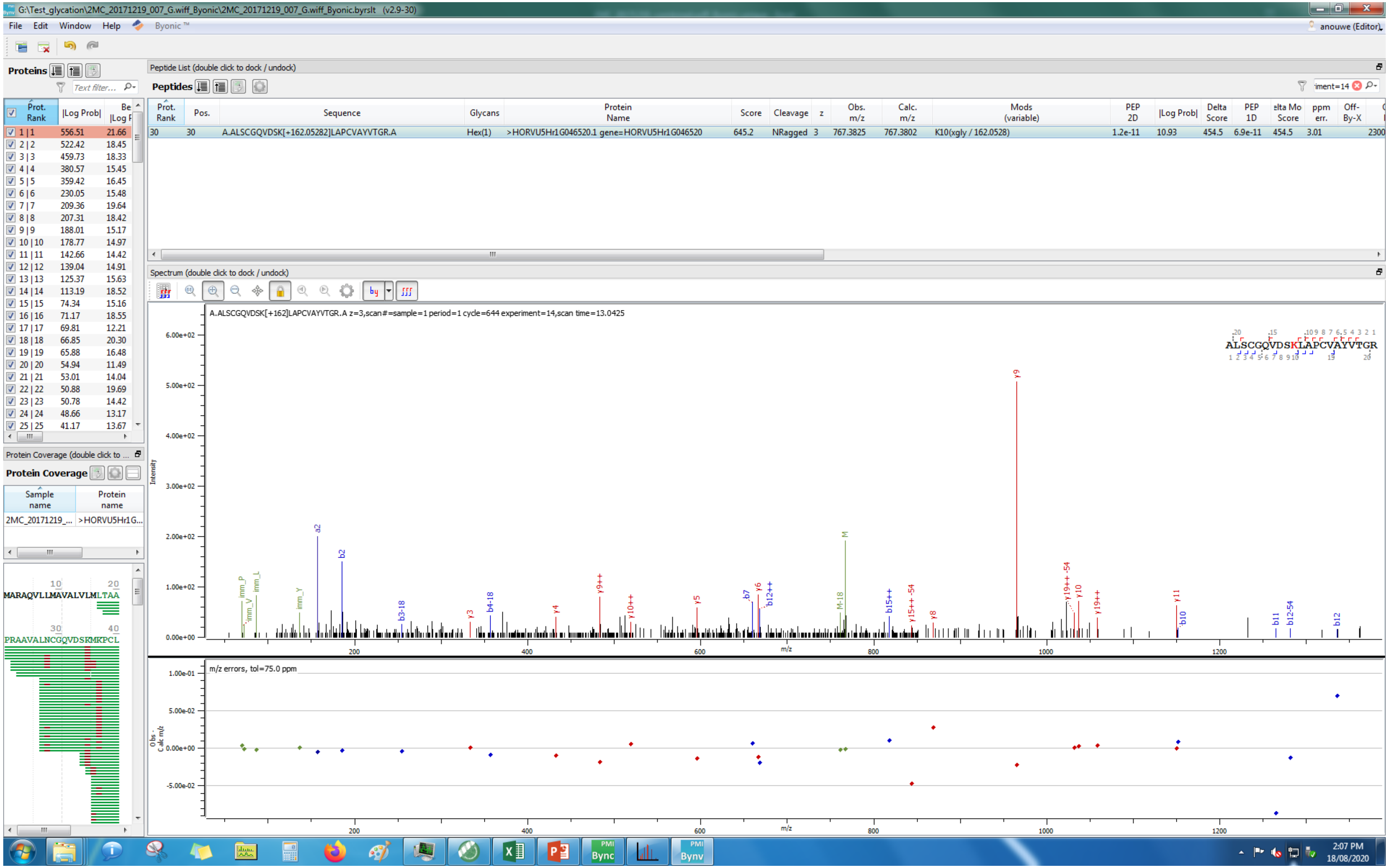
G:\Test_glycation\2MC_20171219_007_G.wiff_Byonic\2MC_20171219_007_G.wiff_Byonic.byrslt (v2.9-30)
File Edit Window Help Byonic™
anouwe (Editor)
Proteins
Peptide List (double click to dock / undock)
Peptides
Prot. Rank | Pos. | Sequence | Glycans | Protein Name | Score | Cleavage | z | Obs. m/z | Calc. m/z | Mods (variable) | PEP 2D | |Log Prob| | Delta Score | PEP 1D | Delta Mod Score | ppm err. | Off-By-X
30 | 30 | A.ALSCGQVDSK[+162.05282]LAPCVAYVTGR.A | Hex(1) | >HORVU5Hr1G046520.1 gene=HORVU5Hr1G046520 | 645.2 | NRagged | 3 | 767.3825 | 767.3802 | K10(xgly / 162.0528) | 1.2e-11 | 10.93 | 454.5 | 6.9e-11 | 454.5 | 3.01
Spectrum (double click to dock / undock)
A.ALSCGQVDSK[+162]LAPCVAYVTGR.A z=3,scan#=sample=1 period=1 cycle=644 experiment=14,scan time=13.0425
ALSCGQVDSKLAPCVAYVTGR
Intensity
m/z
m/z errors, tol=75.0 ppm
Obs - Calc m/z
Protein Coverage
Sample name
Protein name
2MC_20171219_...
>HORVU5Hr1G...
MARAQVLLMAVALVLMLTAA
PRAAVALNCGQVDSKMKPCL
2:07 PM
18/08/2020

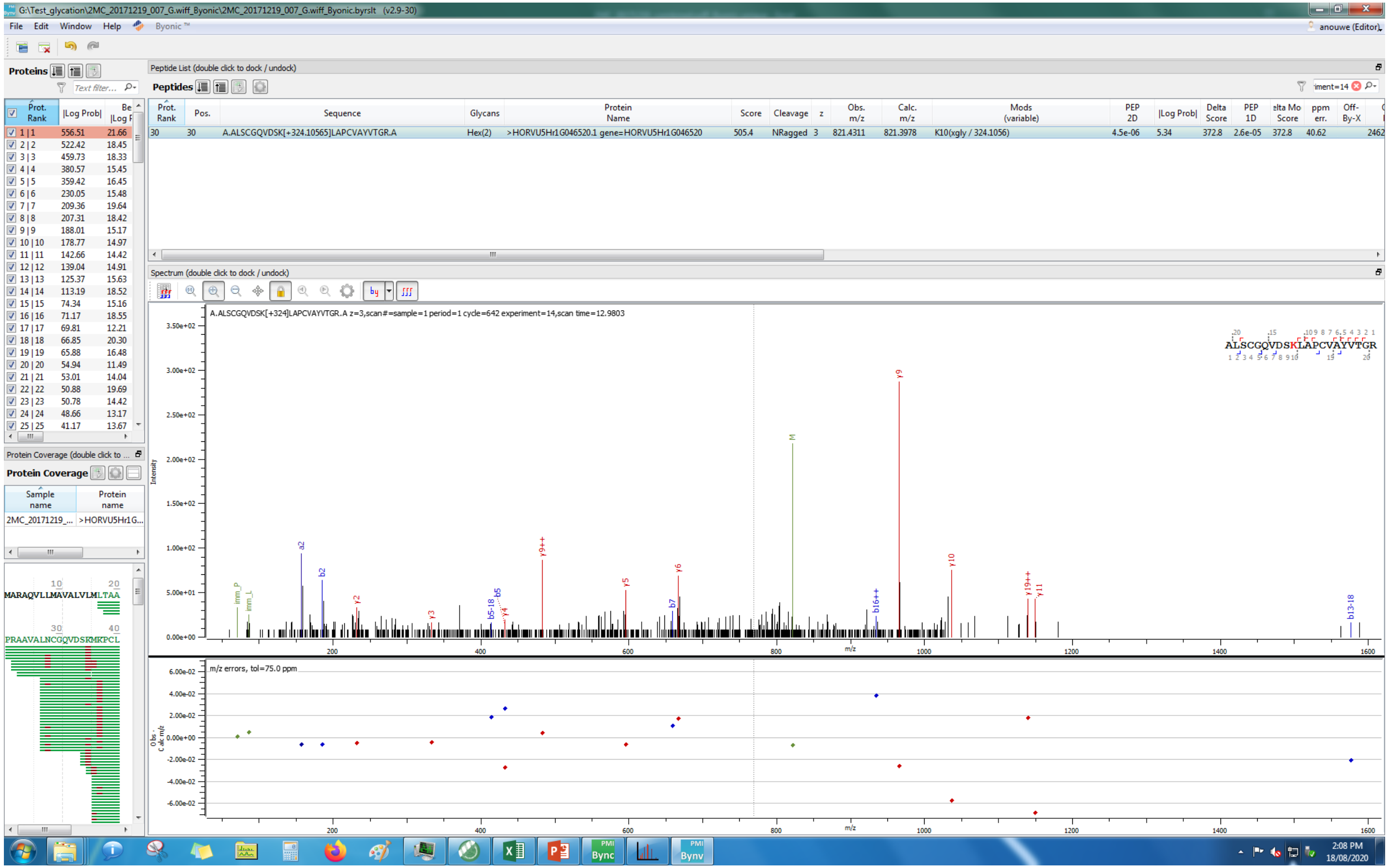
G:\Test_glycation\2MC_20171219_007_G.wiff_Byonic\2MC_20171219_007_G.wiff_Byonic.byrslt (v2.9-30)
File Edit Window Help Byonic ™
Proteins
Peptide List (double click to dock / undock)
Peptides
A.ALSCGQVDSK[+324.10565]LAPCVAYVTGR.A
Hex(2)
>HORVU5Hr1G046520.1 gene=HORVU5Hr1G046520
K10(xgly / 324.1056)
Spectrum (double click to dock / undock)
A.ALSCGQVDSK[+324]LAPCVAYVTGR.A z=3,scan#=sample=1 period=1 cycle=642 experiment=14,scan time=12.9803
ALSCGQVDSKLAPCVAYVTGR
Intensity
m/z
m/z errors, tol=75.0 ppm
Protein Coverage
MARAQVLLMAVALVLMLTAA
PRAAVALNCGQVDSKMKPCL
2:08 PM
18/08/2020

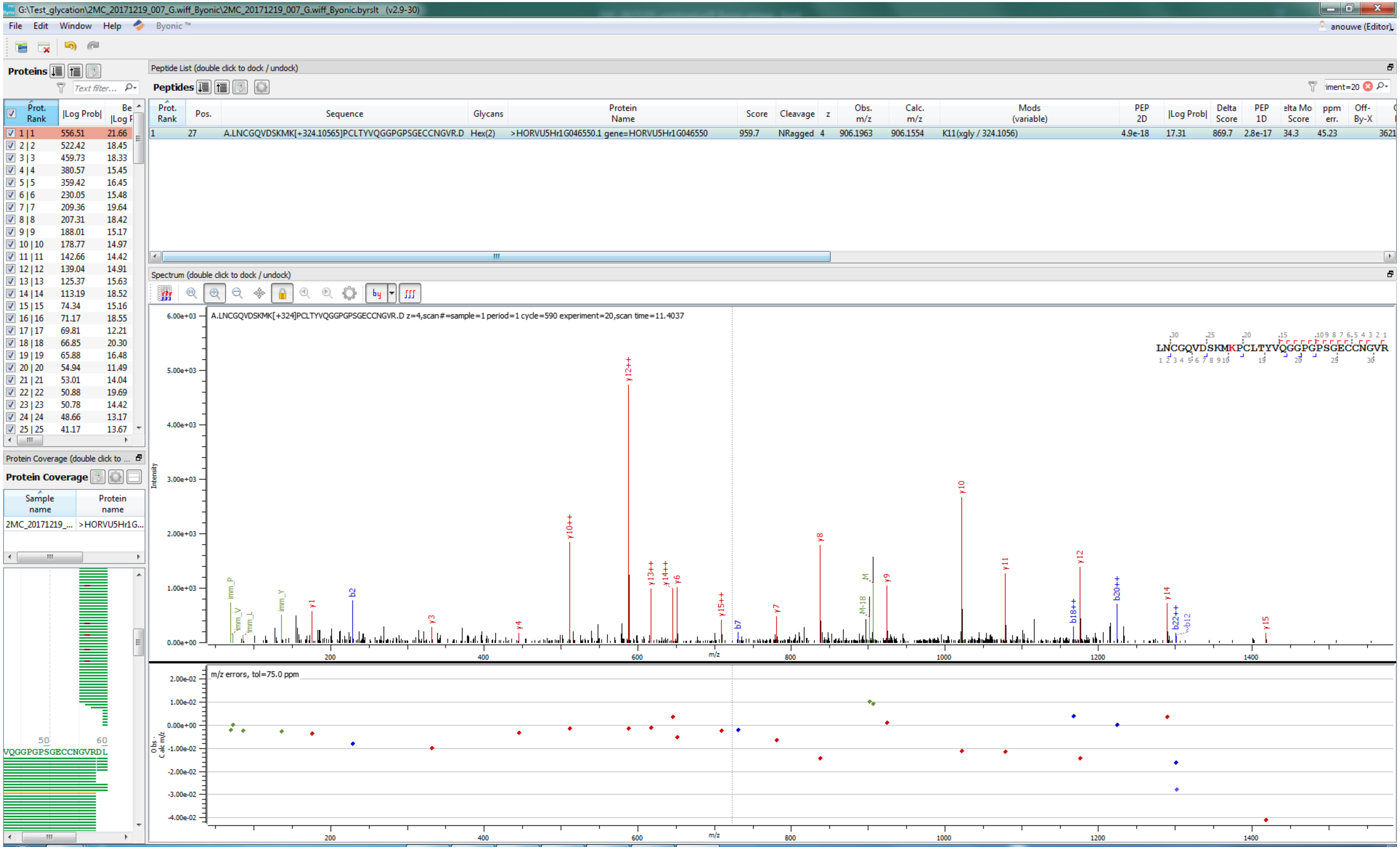
G:\Test_glycation\2MC_20171219_007_G.wiff_Byonic\2MC_20171219_007_G.wiff_Byonic.byrslt (v2.9-30)
File Edit Window Help Byonic™
anouwe (Editor)
Proteins
Peptide List (double click to dock / undock)
Peptides
Prot. Rank
|Log Prob|
Pos.
Sequence
Glycans
Protein Name
Score
Cleavage
z
Obs. m/z
Calc. m/z
Mods (variable)
PEP 2D
Delta Score
PEP 1D
ppm err.
Off-By-X
1 27 A.LNCGQVDSKMK[+324.10565]PCLTYVQGGPGPSGECCNGVR.D Hex(2) >HORVU5Hr1G046550.1 gene=HORVU5Hr1G046550 959.7 NRagged 4 906.1963 906.1554 K11(xgly / 324.1056) 4.9e-18 17.31 869.7 2.8e-17 34.3 45.23 3621
Spectrum (double click to dock / undock)
A.LNCGQVDSKMK[+324]PCLTYVQGGPGPSGECCNGVR.D z=4,scan#=sample=1 period=1 cycle=590 experiment=20,scan time=11.4037
LNCGQVDSKMKPCLTYVQGGPGPSGECCNGVR
Intensity
m/z
m/z errors, tol=75.0 ppm
Obs - Calc m/z
Protein Coverage (double click to ...
Protein Coverage
Sample name
Protein name
2MC_20171219_...
>HORVU5Hr1G...
VQGGPGPSGECCNGVRDL

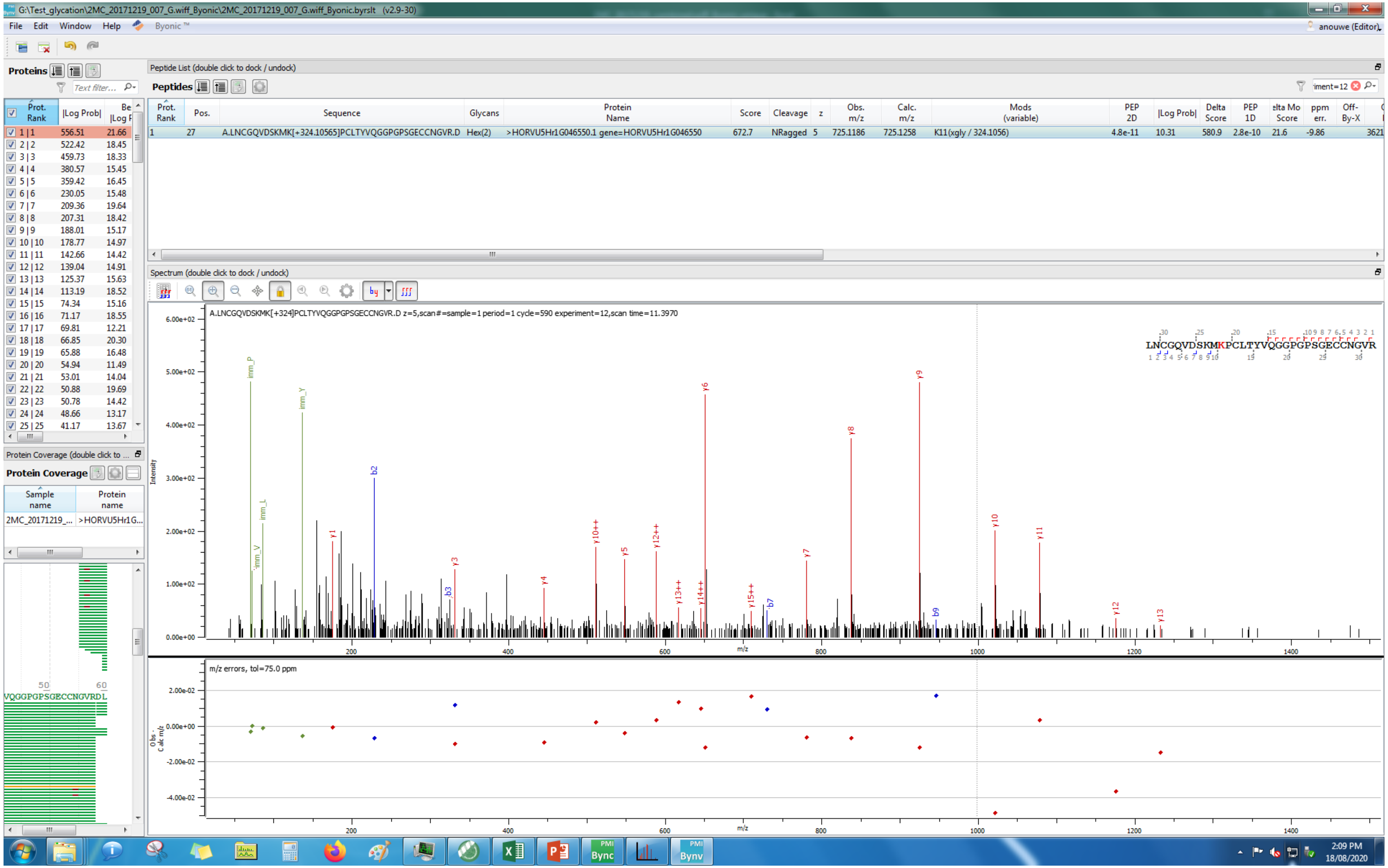
G:\Test_glycation\2MC_20171219_007_G.wiff_Byonic\2MC_20171219_007_G.wiff_Byonic.byrslt (v2.9-30)
File Edit Window Help Byonic™
Proteins
Peptide List (double click to dock / undock)
Peptides
Prot. Rank | Pos. | Sequence | Glycans | Protein Name | Score | Cleavage | z | Obs. m/z | Calc. m/z | Mods (variable) | PEP 2D | |Log Prob| | Delta Score | PEP 1D | ppm err.
1 | 27 | A.LNCGQVDSKMK[+324.10565]PCLTYVQGGPGPSGECCNGVR.D | Hex(2) | >HORVU5Hr1G046550.1 gene=HORVU5Hr1G046550 | 672.7 | NRagged | 5 | 725.1186 | 725.1258 | K11(xgly / 324.1056) | 4.8e-11 | 10.31 | 580.9 | 2.8e-10 | 21.6 | -9.86
Spectrum (double click to dock / undock)
A.LNCGQVDSKMK[+324]PCLTYVQGGPGPSGECCNGVR.D z=5,scan#=sample=1 period=1 cycle=590 experiment=12,scan time=11.3970
LNCGQVDSKMKPCLTYVQGGPGPSGECCNGVR
Intensity
m/z
m/z errors, tol=75.0 ppm
Protein Coverage
Sample name
Protein name
2MC_20171219_...
>HORVU5Hr1G...
VQGGPGPSGECCNGVRDL
anouwe (Editor)
2:09 PM
18/08/2020

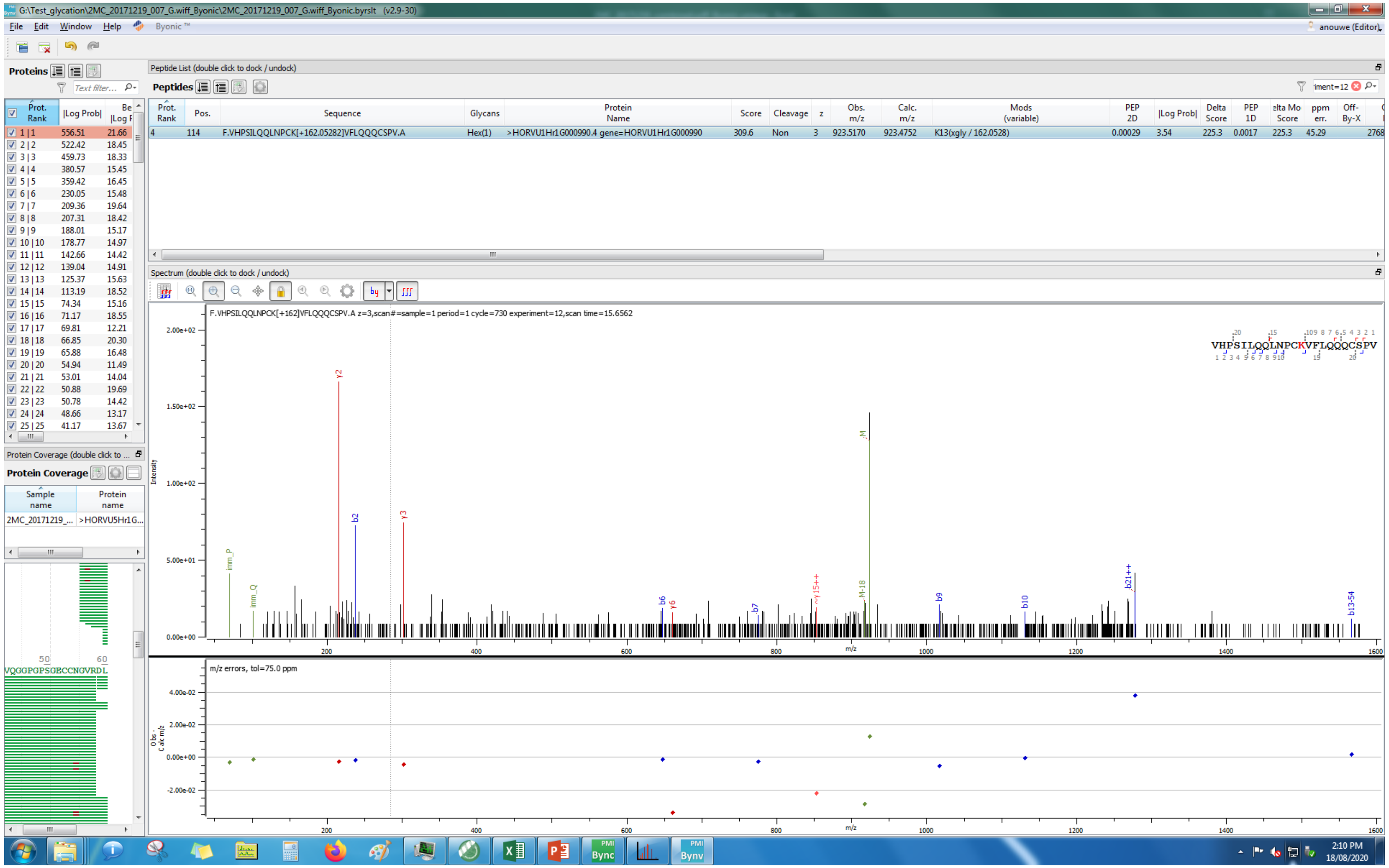
G:\Test_glycation\2MC_20171219_007_G.wiff_Byonic\2MC_20171219_007_G.wiff_Byonic.byrslt (v2.9-30)
File Edit Window Help Byonic™
Proteins
Peptide List (double click to dock / undock)
Peptides
F.VHPSILQQLNPCK[+162.05282]VFLQQQCSPV.A
Hex(1)
>HORVU1Hr1G000990.4 gene=HORVU1Hr1G000990
K13(xgly / 162.0528)
Spectrum (double click to dock / undock)
F.VHPSILQQLNPCK[+162]VFLQQQCSPV.A z=3,scan#=sample=1 period=1 cycle=730 experiment=12,scan time=15.6562
VHPSILQQLNPCKVFLQQQCSPV
m/z errors, tol=75.0 ppm
Protein Coverage
2:10 PM
18/08/2020

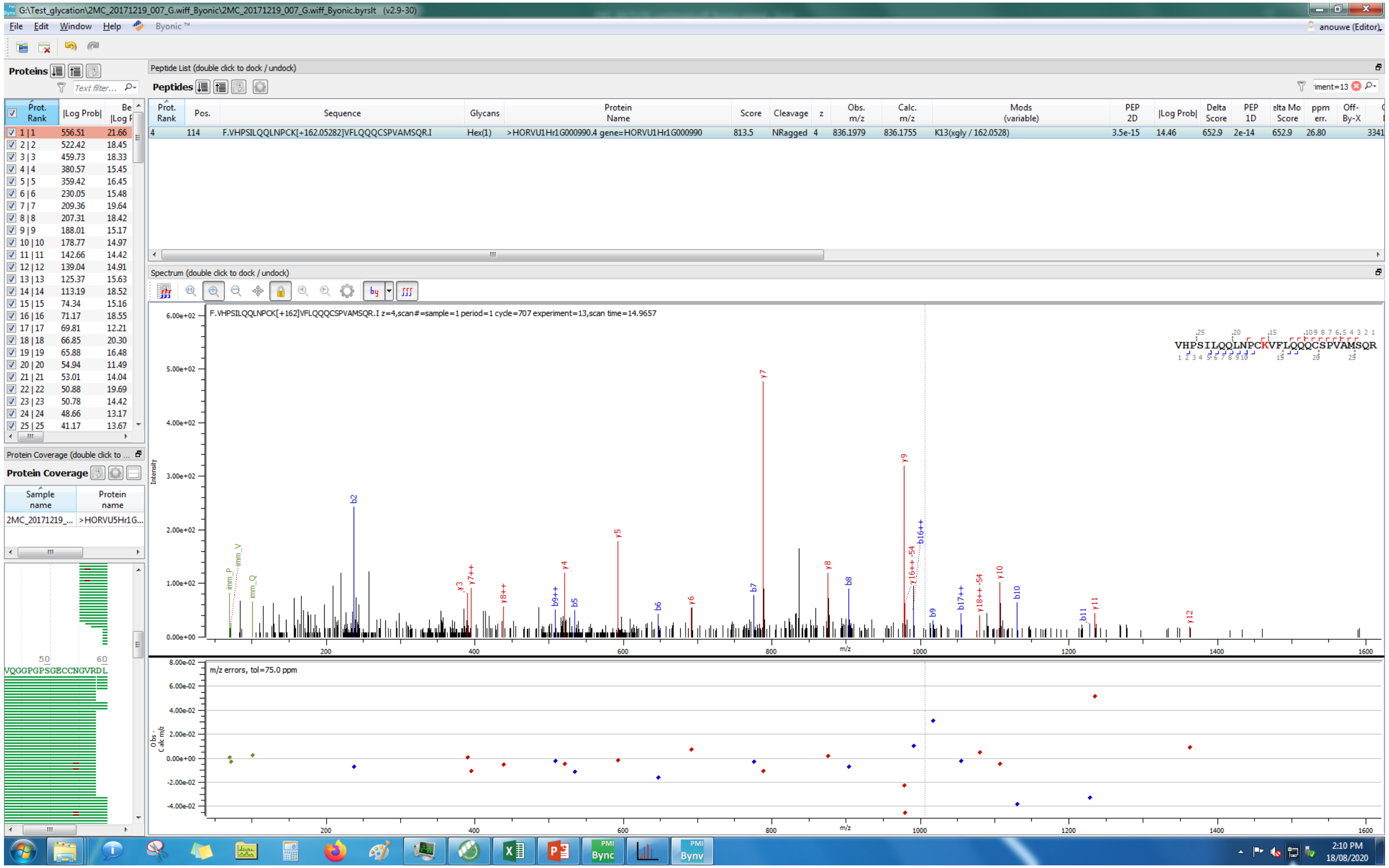

G:\Test_glycation\2MC_20171219_007_G.wiff_Byonic\2MC_20171219_007_G.wiff_Byonic.byrslt (v2.9-30)
File Edit Window Help Byonic ™
anouwe (Editor)
Proteins
Peptide List (double click to dock / undock)
Peptides
Prot. Rank | Pos. | Sequence | Glycans | Protein Name | Score | Cleavage | z | Obs. m/z | Calc. m/z | Mods (variable) | PEP 2D | |Log Prob| | Delta Score | PEP 1D | elta Mo Score | ppm err. | Off-By-X
4 | 114 | F.VHPSILQQLNPCK[+162.05282]VFLQQQCSPVAMSQR.I | Hex(1) | >HORVU1Hr1G000990.4 gene=HORVU1Hr1G000990 | 813.5 | NRagged | 4 | 836.1979 | 836.1755 | K13(xgly / 162.0528) | 3.5e-15 | 14.46 | 652.9 | 2e-14 | 652.9 | 26.80
Spectrum (double click to dock / undock)
F.VHPSILQQLNPCK[+162]VFLQQQCSPVAMSQR.I z=4,scan#=sample=1 period=1 cycle=707 experiment=13,scan time=14.9657
VHPSILQQLNPCKVFLQQQCSPVAMSQR
Intensity
m/z
m/z errors, tol=75.0 ppm
Protein Coverage
2MC_20171219_... >HORVU5Hr1G...
VQGGPGPSGECCNGVRDL

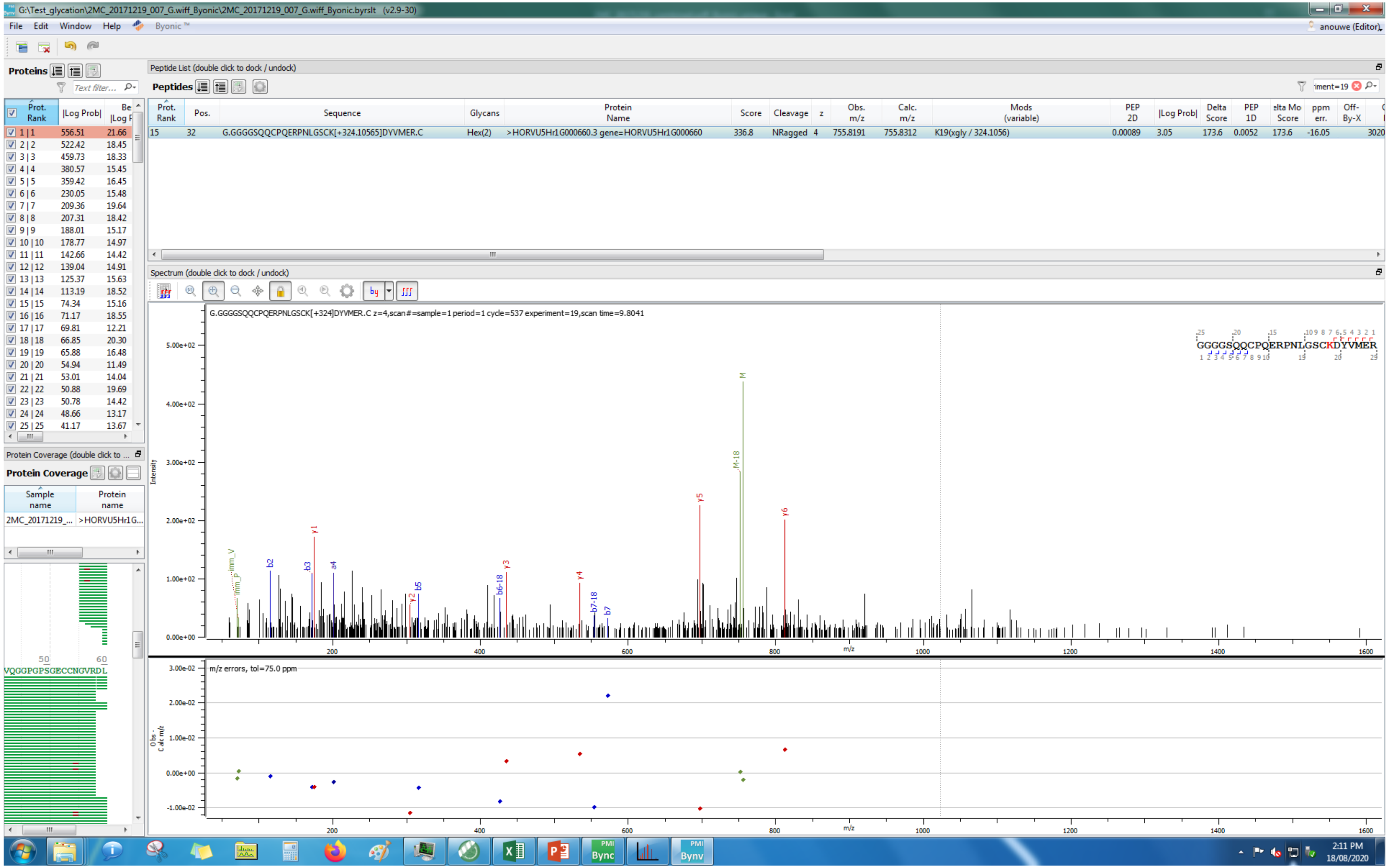
G:\Test_glycation\2MC_20171219_007_G.wiff_Byonic\2MC_20171219_007_G.wiff_Byonic.byrslt (v2.9-30)
File Edit Window Help Byonic ™
anouwe (Editor)
Proteins
Peptide List (double click to dock / undock)
Peptides
Prot. Rank
Pos.
Sequence
Glycans
Protein Name
Score
Cleavage
z
Obs. m/z
Calc. m/z
Mods (variable)
PEP 2D
|Log Prob|
Delta Score
PEP 1D
Off-By-X
15 32 G.GGGGSQQCPQERPNLGSCK[+324.10565]DYVMER.C Hex(2) >HORVU5Hr1G000660.3 gene=HORVU5Hr1G000660 336.8 NRagged 4 755.8191 755.8312 K19(xgly / 324.1056) 0.00089 3.05 173.6 0.0052 173.6 -16.05
Spectrum (double click to dock / undock)
G.GGGGSQQCPQERPNLGSCK[+324]DYVMER.C z=4,scan#=sample=1 period=1 cycle=537 experiment=19,scan time=9.8041
GGGGSQQCPQERPNLGSCKDYVMER
Intensity
m/z
m/z errors, tol=75.0 ppm
Protein Coverage
Sample name
Protein name
2MC_20171219_...
>HORVU5Hr1G...
VQGGPGPSGECCNGVRDL
2:11 PM
18/08/2020

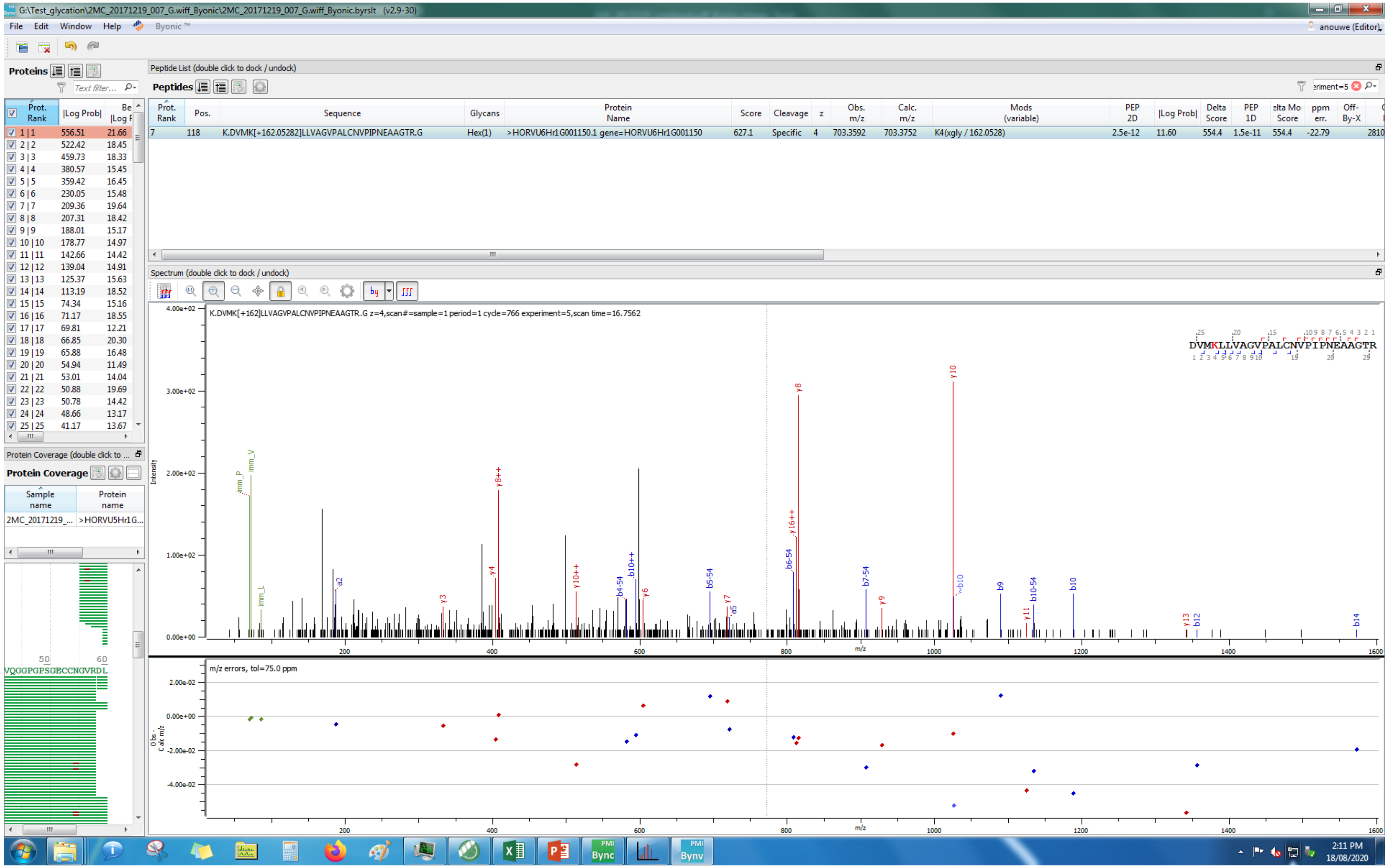
G:\Test_glycation\2MC_20171219_007_G.wiff_Byonic\2MC_20171219_007_G.wiff_Byonic.byrslt (v2.9-30)
File Edit Window Help Byonic
anouwe (Editor)
Proteins
Peptide List (double click to dock / undock)
Peptides
Prot. Rank | Pos. | Sequence | Glycans | Protein Name | Score | Cleavage | z | Obs. m/z | Calc. m/z | Mods (variable) | PEP 2D | |Log Prob| | Delta Score | PEP 1D | elta Mo Score | ppm err. | Off-By-X
7 | 118 | K.DVMK[+162.05282]LLVAGVPALCNVPIPNEAAGTR.G | Hex(1) | >HORVU6Hr1G001150.1 gene=HORVU6Hr1G001150 | 627.1 | Specific | 4 | 703.3592 | 703.3752 | K4(xgly / 162.0528) | 2.5e-12 | 11.60 | 554.4 | 1.5e-11 | 554.4 | -22.79 | 2810
Spectrum (double click to dock / undock)
K.DVMK[+162]LLVAGVPALCNVPIPNEAAGTR.G z=4,scan#=sample=1 period=1 cycle=766 experiment=5,scan time=16.7562
DVMKLLVAGVPALCNVPIPNEAAGTR
Intensity
m/z
m/z errors, tol=75.0 ppm
Protein Coverage
Sample name | Protein name
2MC_20171219_... | >HORVU5Hr1G...
VQGGPGPSGECCNGVRDL
2:11 PM
18/08/2020

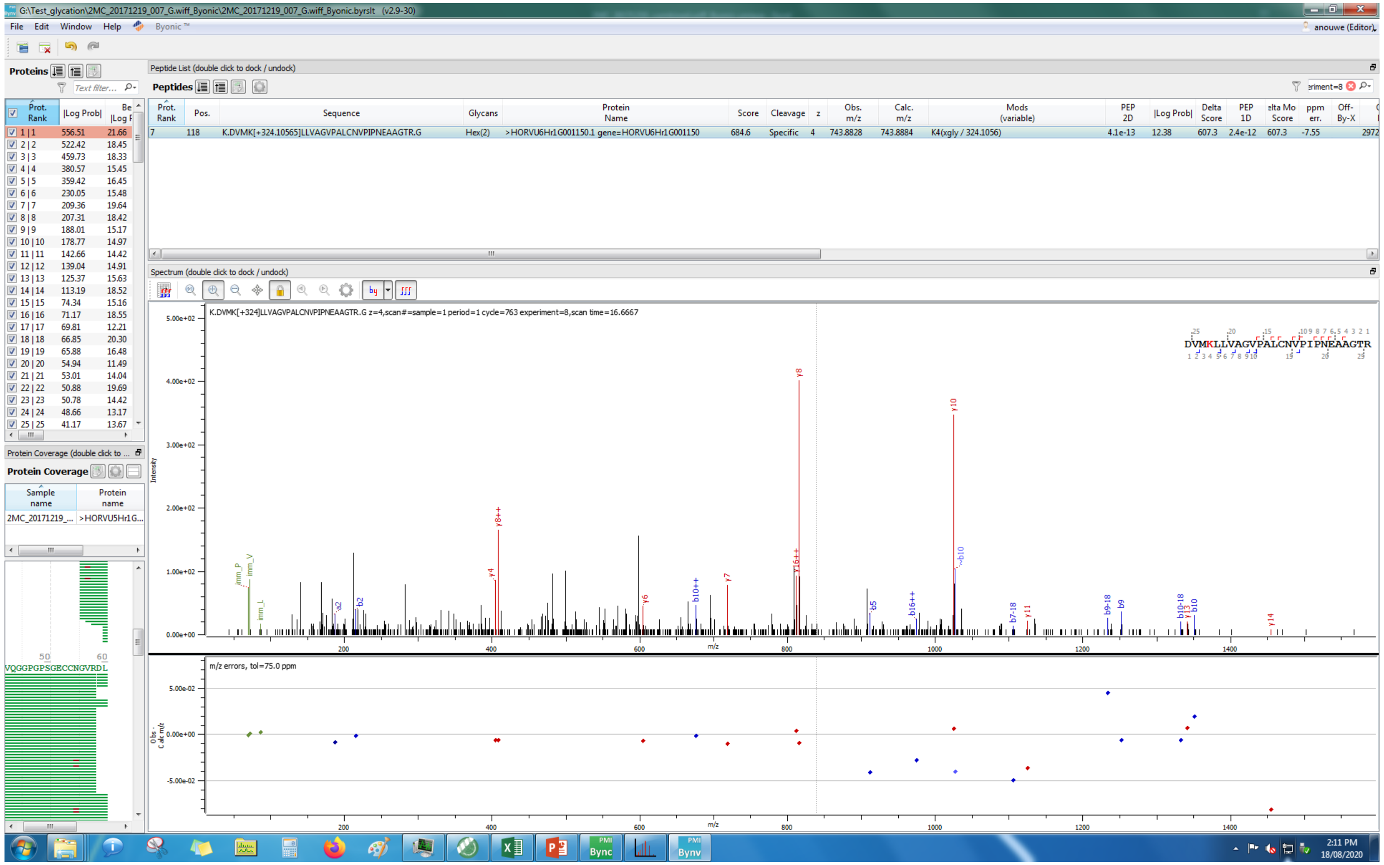
G:\Test_glycation\2MC_20171219_007_G.wiff_Byonic\2MC_20171219_007_G.wiff_Byonic.byrslt (v2.9-30)
File Edit Window Help Byonic ™
Proteins
Peptide List (double click to dock / undock)
Peptides
K.DVMK[+324.10565]LLVAGVPALCNVPIPNEAAGTR.G
Hex(2)
>HORVU6Hr1G001150.1 gene=HORVU6Hr1G001150
K4(xgly / 324.1056)
Spectrum (double click to dock / undock)
K.DVMK[+324]LLVAGVPALCNVPIPNEAAGTR.G z=4,scan#=sample=1 period=1 cycle=763 experiment=8,scan time=16.6667
Protein Coverage
m/z errors, tol=75.0 ppm
2:11 PM
18/08/2020

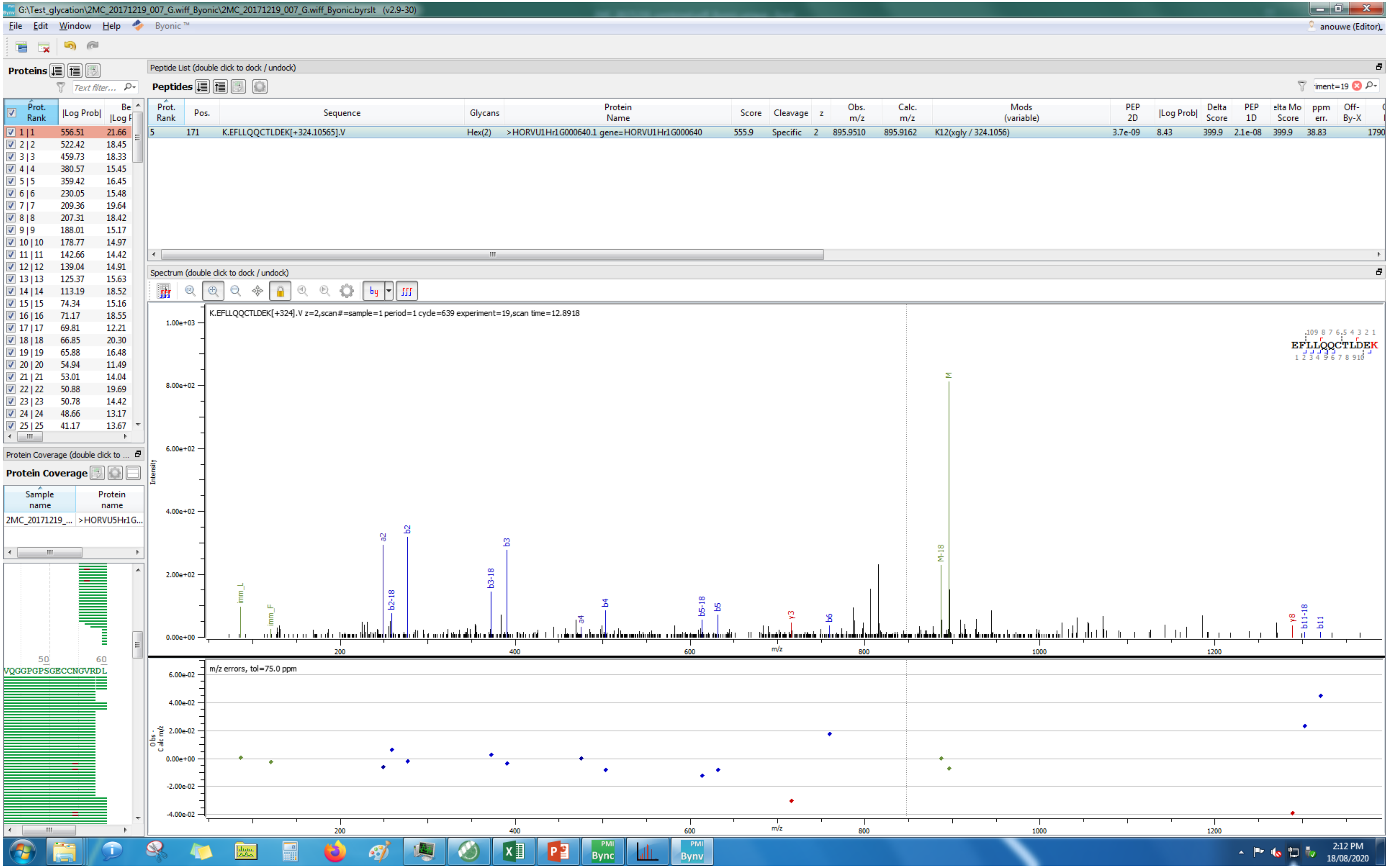
G:\Test_glycation\2MC_20171219_007_G.wiff_Byonic\2MC_20171219_007_G.wiff_Byonic.byrslt (v2.9-30)
File Edit Window Help Byonic™
Proteins
Peptide List (double click to dock / undock)
Peptides
Prot. Rank | Pos. | Sequence | Glycans | Protein Name | Score | Cleavage | z | Obs. m/z | Calc. m/z | Mods (variable) | PEP 2D | |Log Prob| | Delta Score | PEP 1D
5 | 171 | K.EFLLQQCTLDEK[+324.10565].V | Hex(2) | >HORVU1Hr1G000640.1 gene=HORVU1Hr1G000640 | 555.9 | Specific | 2 | 895.9510 | 895.9162 | K12(xgly / 324.1056) | 3.7e-09 | 8.43 | 399.9 | 2.1e-08
Spectrum (double click to dock / undock)
K.EFLLQQCTLDEK[+324].V z=2,scan#=sample=1 period=1 cycle=639 experiment=19,scan time=12.8918
EFLLQQCTLDEK
Intensity
m/z
m/z errors, tol=75.0 ppm
Protein Coverage
Sample name
Protein name
2MC_20171219_...
>HORVU5Hr1G...
VQGGPGPSGECCNGVRDL
2:12 PM
18/08/2020

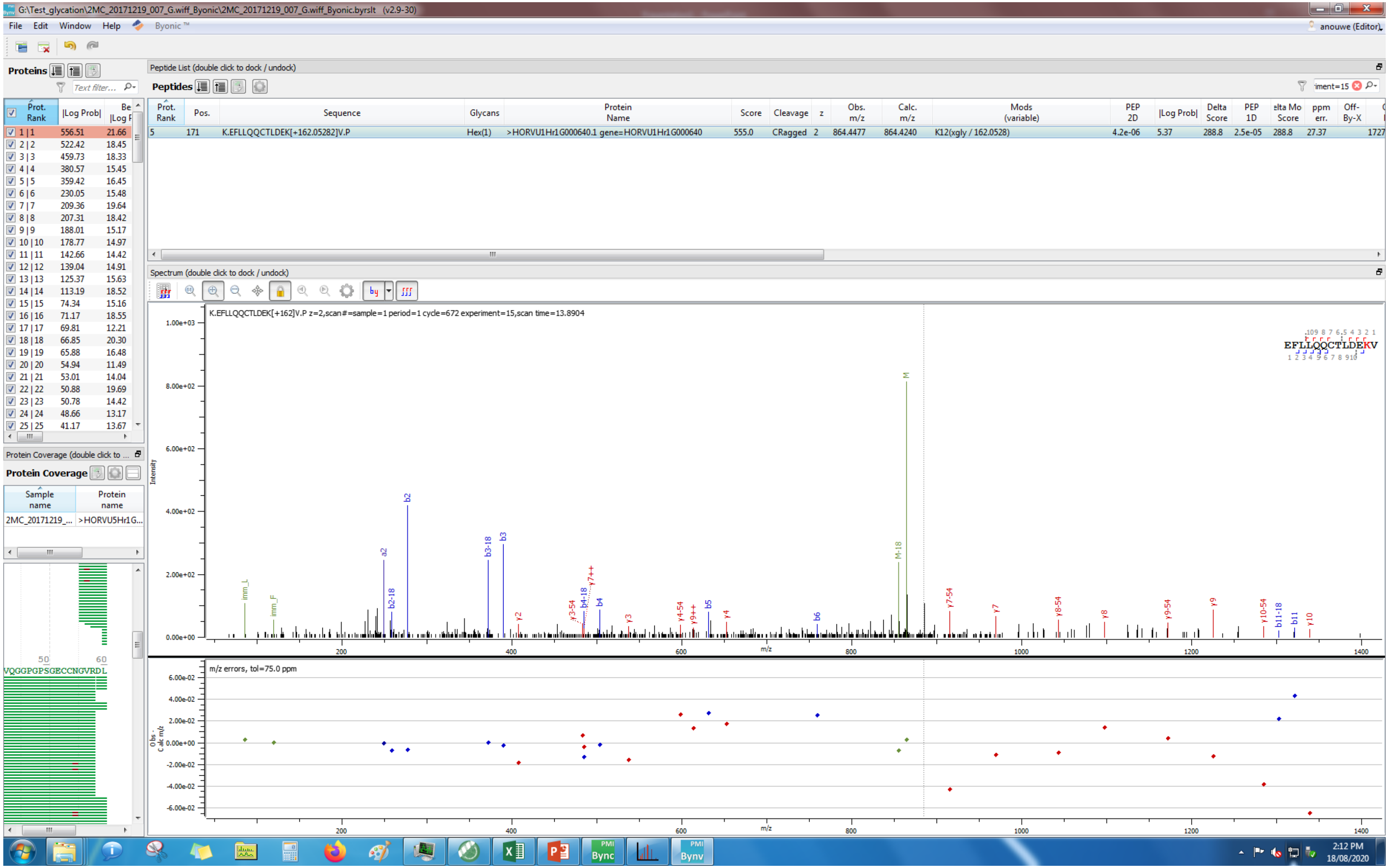
G:\Test_glycation\2MC_20171219_007_G.wiff_Byonic\2MC_20171219_007_G.wiff_Byonic.byrslt (v2.9-30)
File Edit Window Help Byonic™
anouwe (Editor)
Proteins
Peptide List (double click to dock / undock)
Peptides
Prot. Rank | Pos. | Sequence | Glycans | Protein Name | Score | Cleavage | z | Obs. m/z | Calc. m/z | Mods (variable) | PEP 2D | |Log Prob| | Delta Score | PEP 1D | Delta Mod Score | ppm err. | Off-By-X
5 | 171 | K.EFLLQQCTLDEK[+162.05282]V.P | Hex(1) | >HORVU1Hr1G000640.1 gene=HORVU1Hr1G000640 | 555.0 | CRagged | 2 | 864.4477 | 864.4240 | K12(xgly / 162.0528) | 4.2e-06 | 5.37 | 288.8 | 2.5e-05 | 288.8 | 27.37
Spectrum (double click to dock / undock)
K.EFLLQQCTLDEK[+162]V.P z=2,scan#=sample=1 period=1 cycle=672 experiment=15,scan time=13.8904
EFLLQQCTLDEKV
Intensity
m/z
m/z errors, tol=75.0 ppm
Obs - Calc m/z
Protein Coverage (double click to ...
Protein Coverage
Sample name
Protein name
2MC_20171219_...
>HORVU5Hr1G...
VQGGPGPSGECCNGVRDL
2:12 PM
18/08/2020

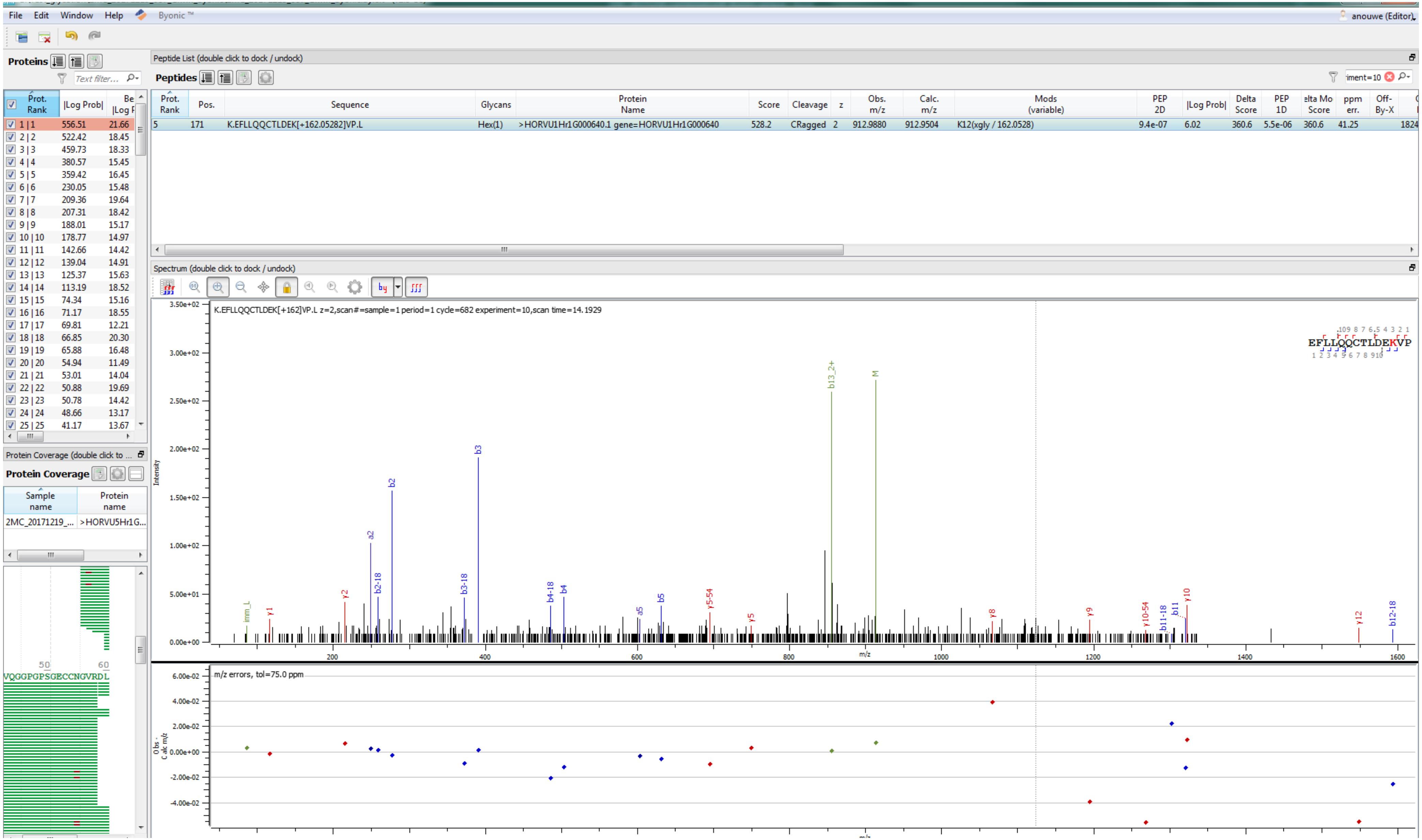
File Edit Window Help Byonic ™
anouwe (Editor)
Proteins
Peptide List (double click to dock / undock)
Peptides
Prot. Rank | Pos. | Sequence | Glycans | Protein Name | Score | Cleavage | z | Obs. m/z | Calc. m/z | Mods (variable) | PEP 2D | |Log Prob| | Delta Score | PEP 1D | Delta Mod Score | ppm err. | Off-By-X
5 | 171 | K.EFLLQQCTLDEK[+162.05282]VP.L | Hex(1) | >HORVU1Hr1G000640.1 gene=HORVU1Hr1G000640 | 528.2 | CRagged | 2 | 912.9880 | 912.9504 | K12(xgly / 162.0528) | 9.4e-07 | 6.02 | 360.6 | 5.5e-06 | 360.6 | 41.25 | 1824
Spectrum (double click to dock / undock)
K.EFLLQQCTLDEK[+162]VP.L z=2,scan#=sample=1 period=1 cycle=682 experiment=10,scan time=14.1929
EFLLQQCTLDEKVP
Intensity
m/z
m/z errors, tol=75.0 ppm
Obs - Calc m/z
Protein Coverage (double click to ...
Protein Coverage
Sample name | Protein name
2MC_20171219_... | >HORVU5Hr1G...
VQGGPGPSGECCNGVRDL

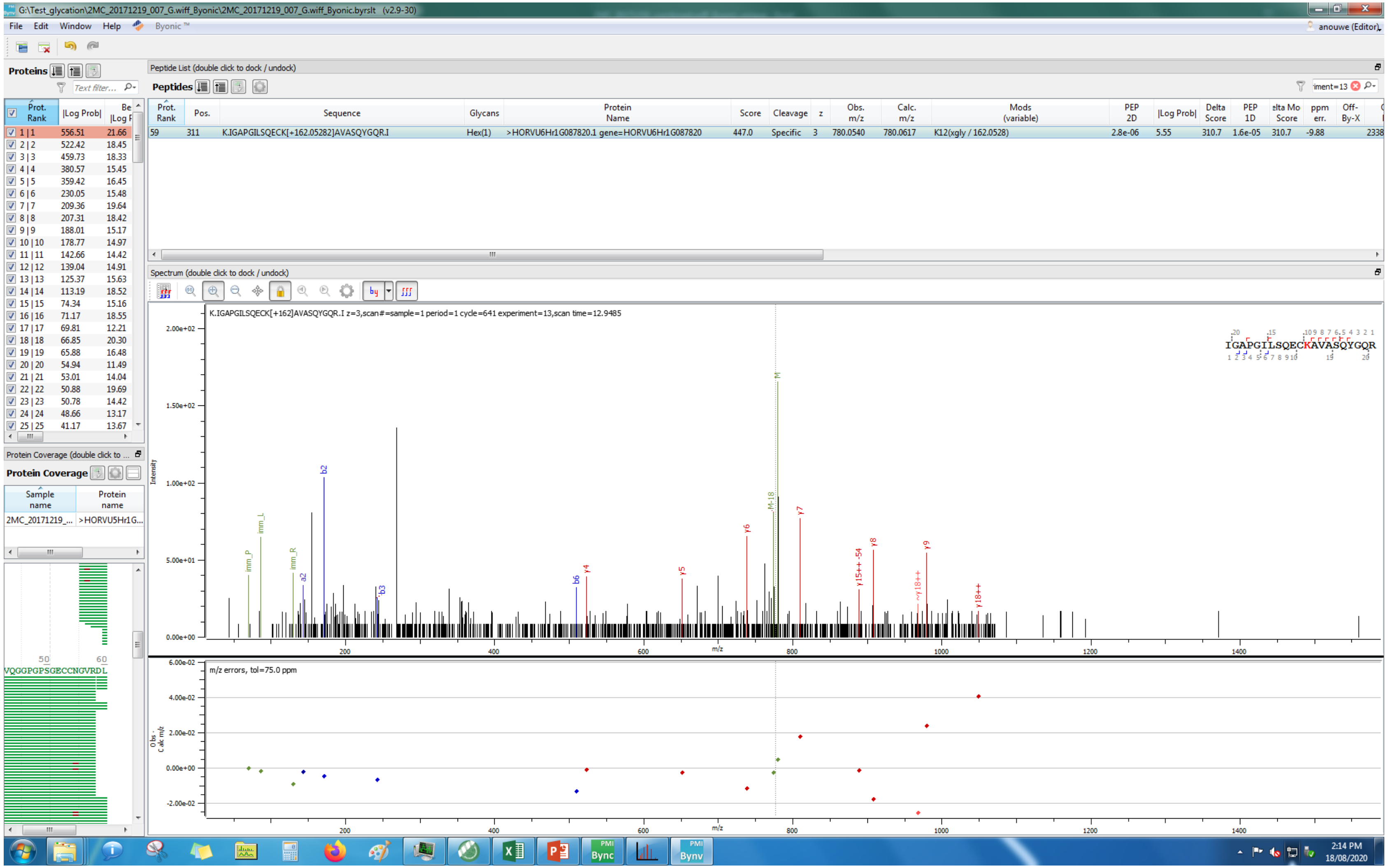
G:\Test_glycation\2MC_20171219_007_G.wiff_Byonic\2MC_20171219_007_G.wiff_Byonic.byrslt (v2.9-30)
File Edit Window Help Byonic ™
anouwe (Editor)
Proteins
Peptide List (double click to dock / undock)
Peptides
Prot. Rank | Pos. | Sequence | Glycans | Protein Name | Score | Cleavage | z | Obs. m/z | Calc. m/z | Mods (variable) | PEP 2D | |Log Prob| | Delta Score | PEP 1D | Delta Mod Score | ppm err. | Off-By-X
59 | 311 | K.IGAPGILSQECK[+162.05282]AVASQYGQR.I | Hex(1) | >HORVU6Hr1G087820.1 gene=HORVU6Hr1G087820 | 447.0 | Specific | 3 | 780.0540 | 780.0617 | K12(xgly / 162.0528) | 2.8e-06 | 5.55 | 310.7 | 1.6e-05 | 310.7 | -9.88 | 2338
Spectrum (double click to dock / undock)
K.IGAPGILSQECK[+162]AVASQYGQR.I z=3,scan#=sample=1 period=1 cycle=641 experiment=13,scan time=12.9485
IGAPGILSQECKAVASQYGQR
Intensity
m/z
m/z errors, tol=75.0 ppm
Obs - Calc m/z
Protein Coverage (double click to ...
Protein Coverage
Sample name | Protein name
2MC_20171219_... | >HORVU5Hr1G...
VQGGPGPSGECCNGVRDL
2:14 PM
18/08/2020

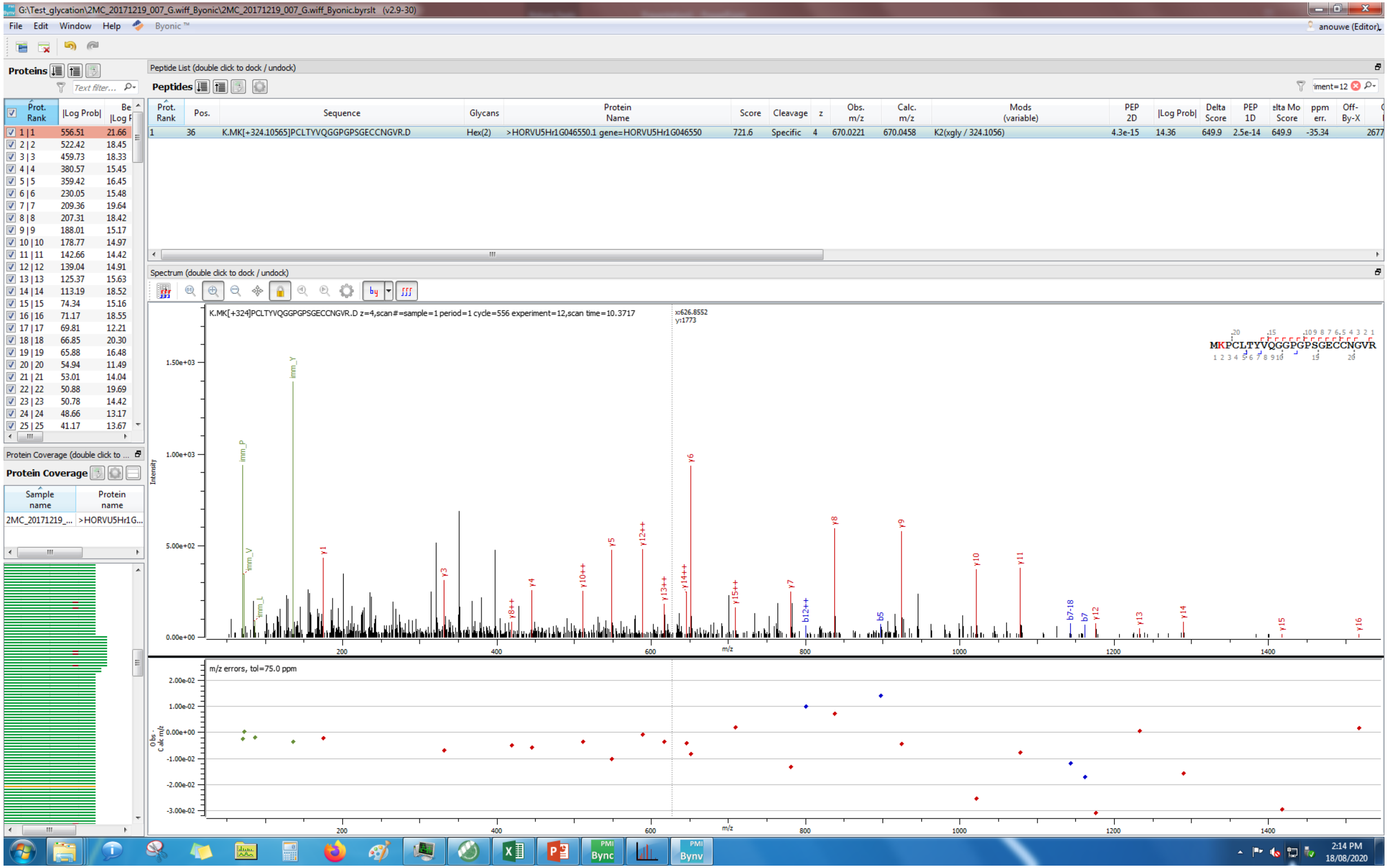
G:\Test_glycation\2MC_20171219_007_G.wiff_Byonic\2MC_20171219_007_G.wiff_Byonic.byrslt (v2.9-30)
Peptide List (double click to dock / undock)
K.MK[+324.10565]PCLTYVQGGPGPSGECCNGVR.D
Hex(2)
>HORVU5Hr1G046550.1 gene=HORVU5Hr1G046550
K2(xgly / 324.1056)
Spectrum (double click to dock / undock)
K.MK[+324]PCLTYVQGGPGPSGECCNGVR.D z=4,scan#=sample=1 period=1 cycle=556 experiment=12,scan time=10.3717
MKPCLTYVQGGPGPSGECCNGVR
m/z errors, tol=75.0 ppm

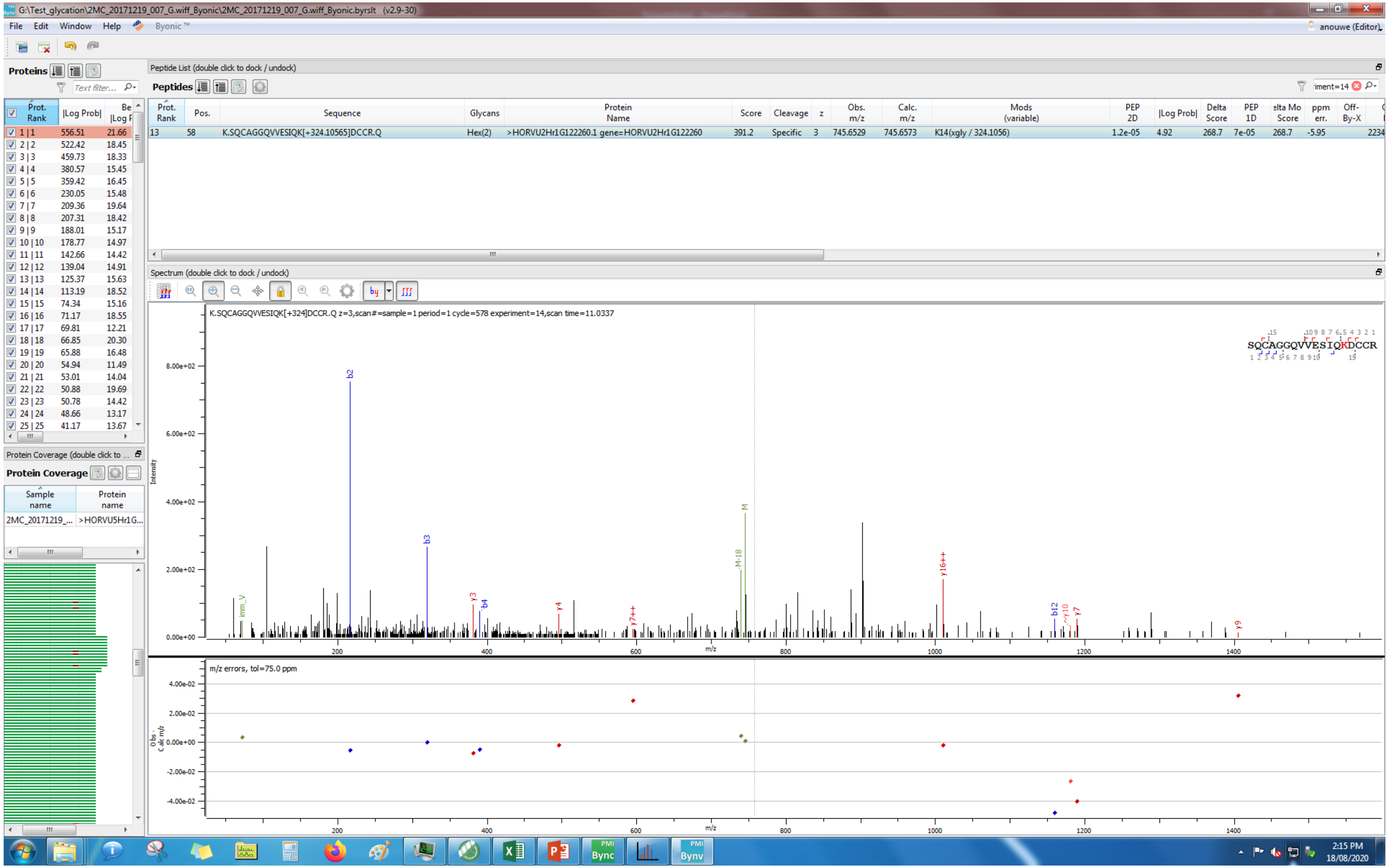
G:\Test_glycation\2MC_20171219_007_G.wiff_Byonic\2MC_20171219_007_G.wiff_Byonic.byrslt (v2.9-30)
File Edit Window Help Byonic™
anouwe (Editor)
Proteins
Peptide List (double click to dock / undock)
Peptides
Prot. Rank | Pos. | Sequence | Glycans | Protein Name | Score | Cleavage | z | Obs. m/z | Calc. m/z | Mods (variable) | PEP 2D | |Log Prob| | Delta Score | PEP 1D | Delta Mod Score | ppm err. | Off-By-X
13 | 58 | K.SQCAGGQVVESIQK[+324.10565]DCCR.Q | Hex(2) | >HORVU2Hr1G122260.1 gene=HORVU2Hr1G122260 | 391.2 | Specific | 3 | 745.6529 | 745.6573 | K14(xgly / 324.1056) | 1.2e-05 | 4.92 | 268.7 | 7e-05 | 268.7 | -5.95 | 2234
Spectrum (double click to dock / undock)
K.SQCAGGQVVESIQK[+324]DCCR.Q z=3,scan#=sample=1 period=1 cycle=578 experiment=14,scan time=11.0337
SQCAGGQVVESIQKDCCR
Intensity
m/z
m/z errors, tol=75.0 ppm
Obs - Calc m/z
Protein Coverage (double click to ...
Protein Coverage
Sample name | Protein name
2MC_20171219_... | >HORVU5Hr1G...
2:15 PM
18/08/2020

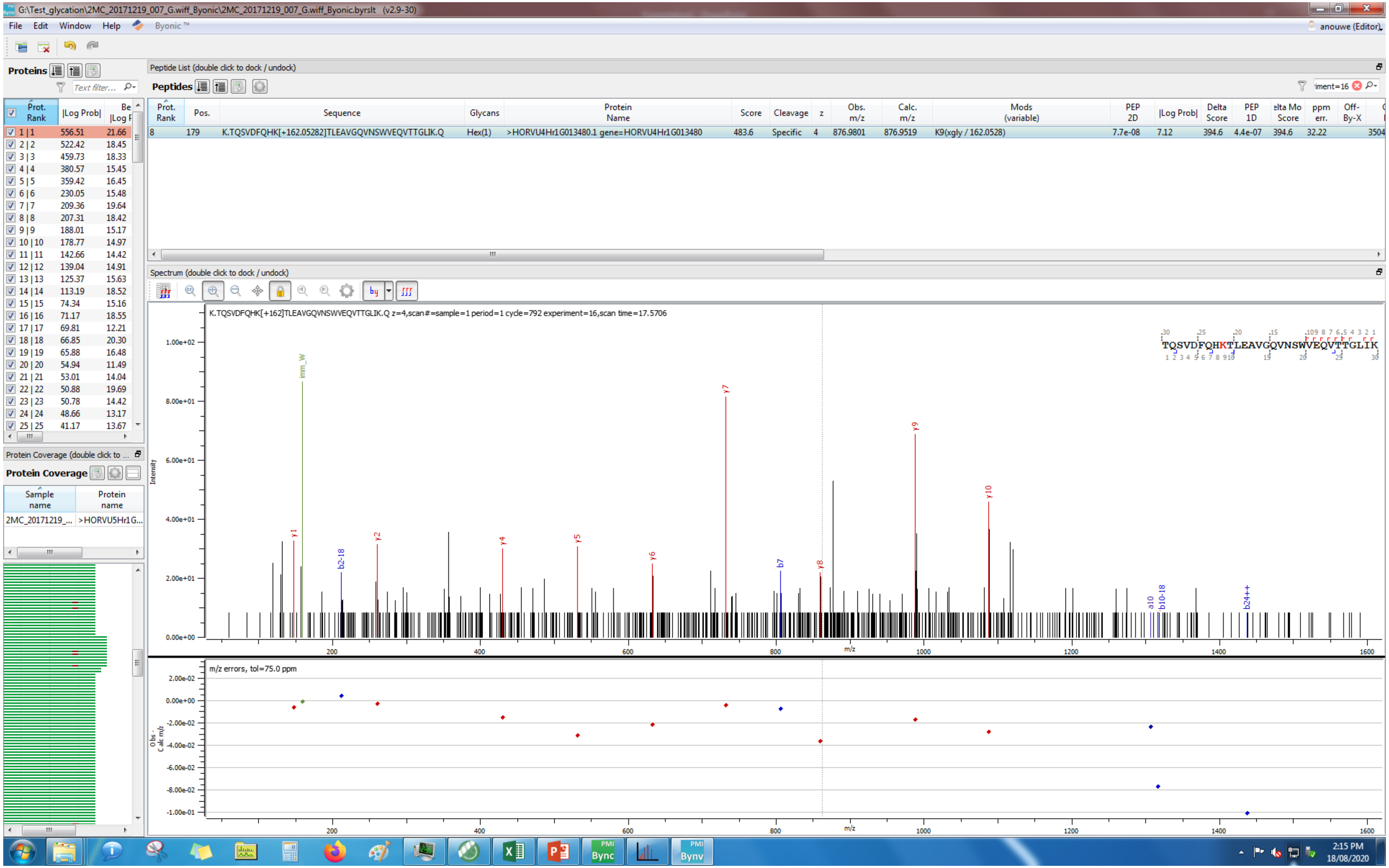
G:\Test_glycation\2MC_20171219_007_G.wiff_Byonic\2MC_20171219_007_G.wiff_Byonic.byrslt (v2.9-30)
File Edit Window Help Byonic™
anouwe (Editor)
Proteins
Peptide List (double click to dock / undock)
Peptides
Prot. Rank | Pos. | Sequence | Glycans | Protein Name | Score | Cleavage | z | Obs. m/z | Calc. m/z | Mods (variable) | PEP 2D | |Log Prob| | Delta Score | PEP 1D | elta Mo Score | ppm err. | Off-By-X
8 | 179 | K.TQSVDFQHK[+162.05282]TLEAVGQVNSWVEQVTTGLIK.Q | Hex(1) | >HORVU4Hr1G013480.1 gene=HORVU4Hr1G013480 | 483.6 | Specific | 4 | 876.9801 | 876.9519 | K9(xgly / 162.0528) | 7.7e-08 | 7.12 | 394.6 | 4.4e-07 | 394.6 | 32.22 | 3504
Prot. Rank | |Log Prob| | Be |Log F
1 | 1 | 556.51 | 21.66
2 | 2 | 522.42 | 18.45
3 | 3 | 459.73 | 18.33
4 | 4 | 380.57 | 15.45
5 | 5 | 359.42 | 16.45
6 | 6 | 230.05 | 15.48
7 | 7 | 209.36 | 19.64
8 | 8 | 207.31 | 18.42
9 | 9 | 188.01 | 15.17
10 | 10 | 178.77 | 14.97
11 | 11 | 142.66 | 14.42
12 | 12 | 139.04 | 14.91
13 | 13 | 125.37 | 15.63
14 | 14 | 113.19 | 18.52
15 | 15 | 74.34 | 15.16
16 | 16 | 71.17 | 18.55
17 | 17 | 69.81 | 12.21
18 | 18 | 66.85 | 20.30
19 | 19 | 65.88 | 16.48
20 | 20 | 54.94 | 11.49
21 | 21 | 53.01 | 14.04
22 | 22 | 50.88 | 19.69
23 | 23 | 50.78 | 14.42
24 | 24 | 48.66 | 13.17
25 | 25 | 41.17 | 13.67
Protein Coverage (double click to ...
Protein Coverage
Sample name | Protein name
2MC_20171219_... | >HORVU5Hr1G...
Spectrum (double click to dock / undock)
K.TQSVDFQHK[+162]TLEAVGQVNSWVEQVTTGLIK.Q z=4,scan#=sample=1 period=1 cycle=792 experiment=16,scan time=17.5706
TQSVDFQHKTLEAVGQVNSWVEQVTTGLIK
Intensity
m/z
imm_W
y1
b2-18
y2
y4
y5
y6
y7
b7
y8
y9
y10
a10
b10-18
b24++
m/z errors, tol=75.0 ppm
Obs - Calc m/z
2:15 PM
18/08/2020

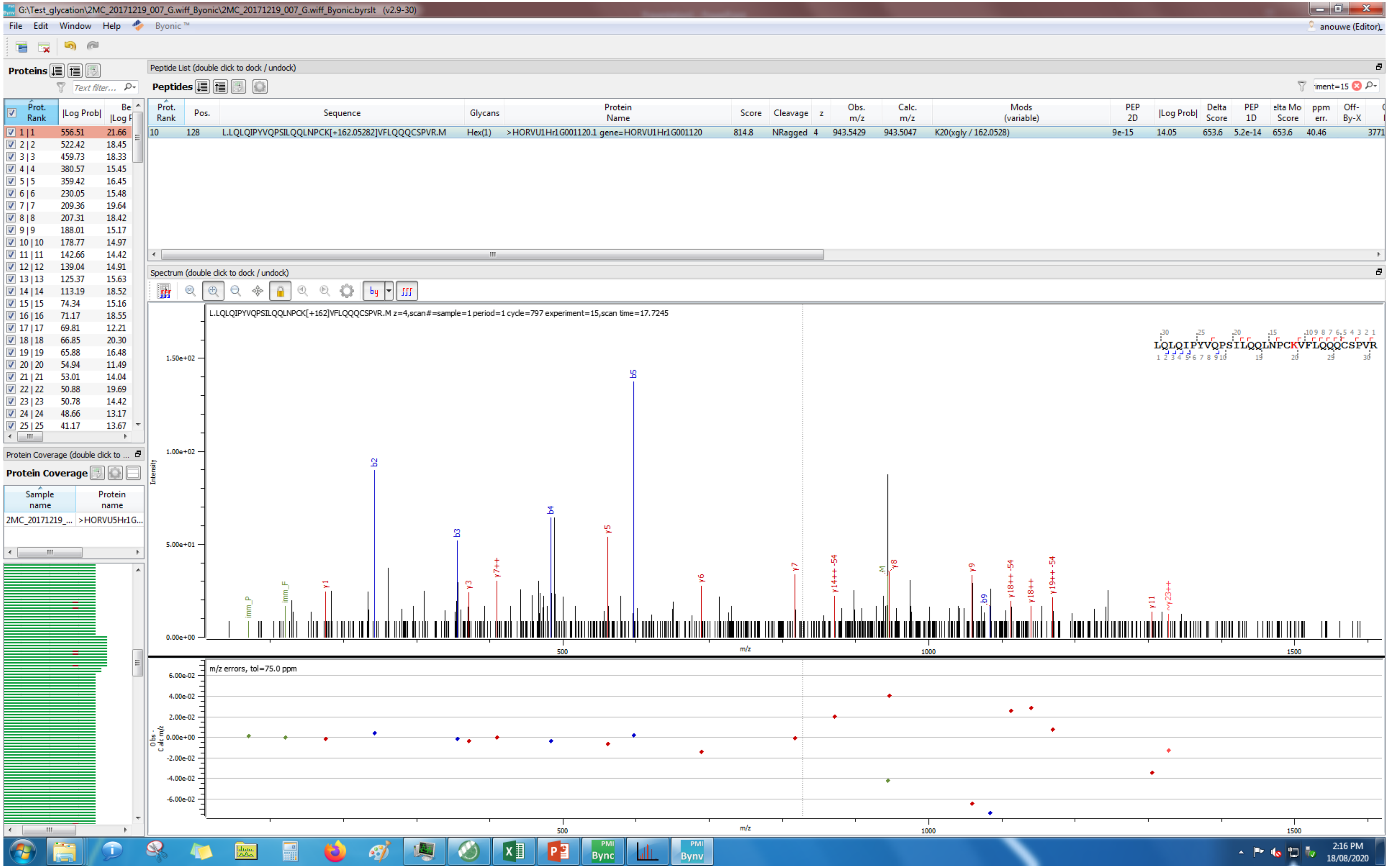
G:\Test_glycation\2MC_20171219_007_G.wiff_Byonic\2MC_20171219_007_G.wiff_Byonic.byrslt (v2.9-30)
File Edit Window Help Byonic™
anouwe (Editor)
Proteins
Peptide List (double click to dock / undock)
Peptides
Prot. Rank | Pos. | Sequence | Glycans | Protein Name | Score | Cleavage | z | Obs. m/z | Calc. m/z | Mods (variable) | PEP 2D | |Log Prob| | Delta Score | PEP 1D | Delta Mod Score | ppm err. | Off-By-X
10 | 128 | L.LQLQIPYVQPSILQQLNPCK[+162.05282]VFLQQQCSPVR.M | Hex(1) | >HORVU1Hr1G001120.1 gene=HORVU1Hr1G001120 | 814.8 | NRagged | 4 | 943.5429 | 943.5047 | K20(xgly / 162.0528) | 9e-15 | 14.05 | 653.6 | 5.2e-14 | 653.6 | 40.46
Spectrum (double click to dock / undock)
L.LQLQIPYVQPSILQQLNPCK[+162]VFLQQQCSPVR.M z=4,scan#=sample=1 period=1 cycle=797 experiment=15,scan time=17.7245
LQLQIPYVQPSILQQLNPCKVFLQQQCSPVR
Intensity
m/z
m/z errors, tol=75.0 ppm
Obs - Calc m/z
Protein Coverage (double click to ...
Protein Coverage
Sample name | Protein name
2MC_20171219_... | >HORVU5Hr1G...
2:16 PM
18/08/2020

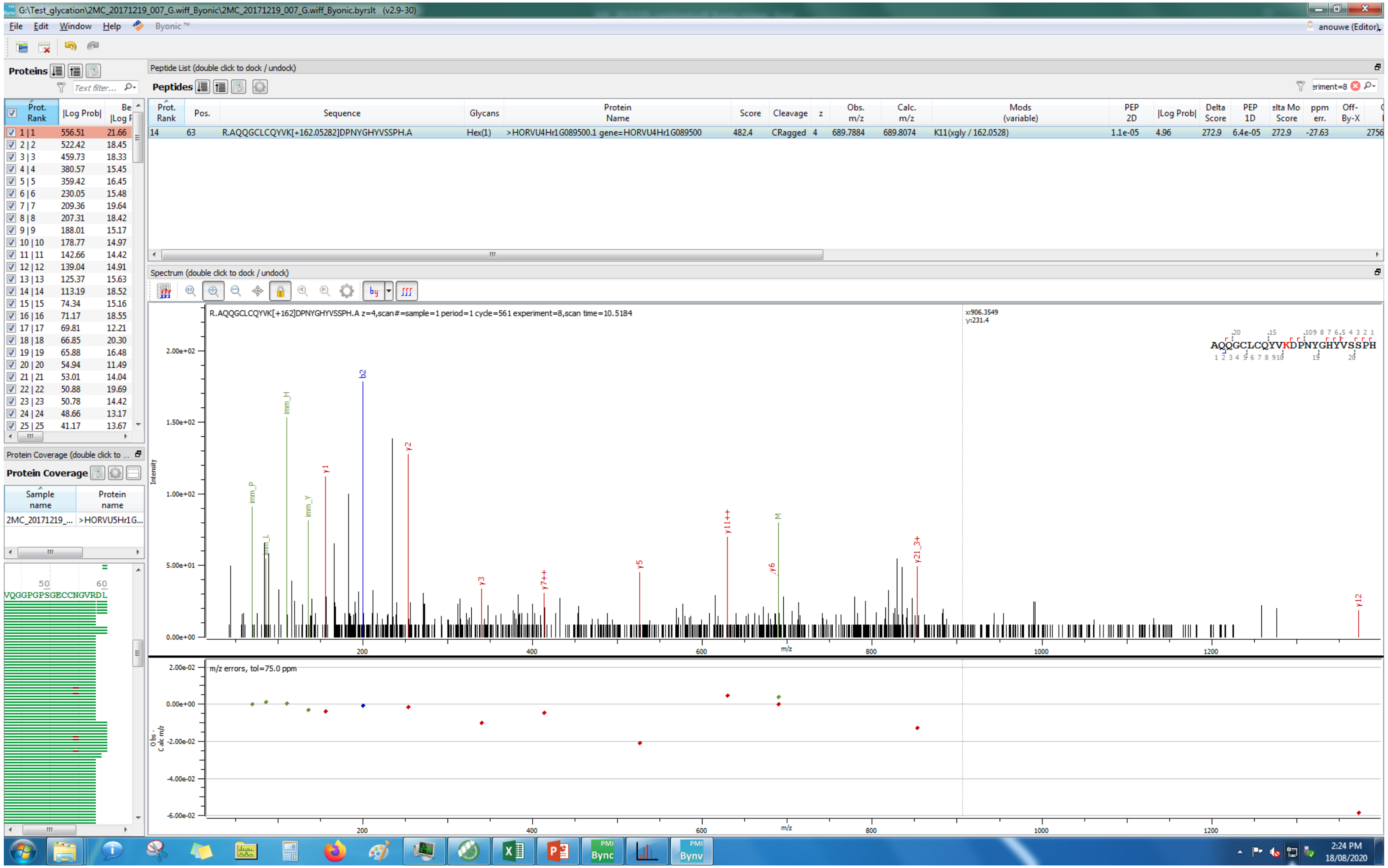
G:\Test_glycation\2MC_20171219_007_G.wiff_Byonic\2MC_20171219_007_G.wiff_Byonic.byrslt (v2.9-30)
File Edit Window Help Byonic
Proteins
Peptide List (double click to dock / undock)
Peptides
Prot. Rank | Pos. | Sequence | Glycans | Protein Name | Score | Cleavage | z | Obs. m/z | Calc. m/z | Mods (variable) | PEP 2D | |Log Prob| | Delta Score | PEP 1D
14 | 63 | R.AQQGCLCQYVK[+162.05282]DPNYGHYVSSPH.A | Hex(1) | >HORVU4Hr1G089500.1 gene=HORVU4Hr1G089500 | 482.4 | CRagged | 4 | 689.7884 | 689.8074 | K11(xgly / 162.0528) | 1.1e-05 | 4.96 | 272.9 | 6.4e-05
Spectrum (double click to dock / undock)
R.AQQGCLCQYVK[+162]DPNYGHYVSSPH.A z=4,scan#=sample=1 period=1 cycle=561 experiment=8,scan time=10.5184
AQQGCLCQYVKDPNYGHYVSSPH
Intensity
m/z
m/z errors, tol=75.0 ppm
Protein Coverage
Sample name
Protein name
2MC_20171219_...
>HORVU5Hr1G...
VQGGPGPSGECCNGVRDL
2:24 PM
18/08/2020

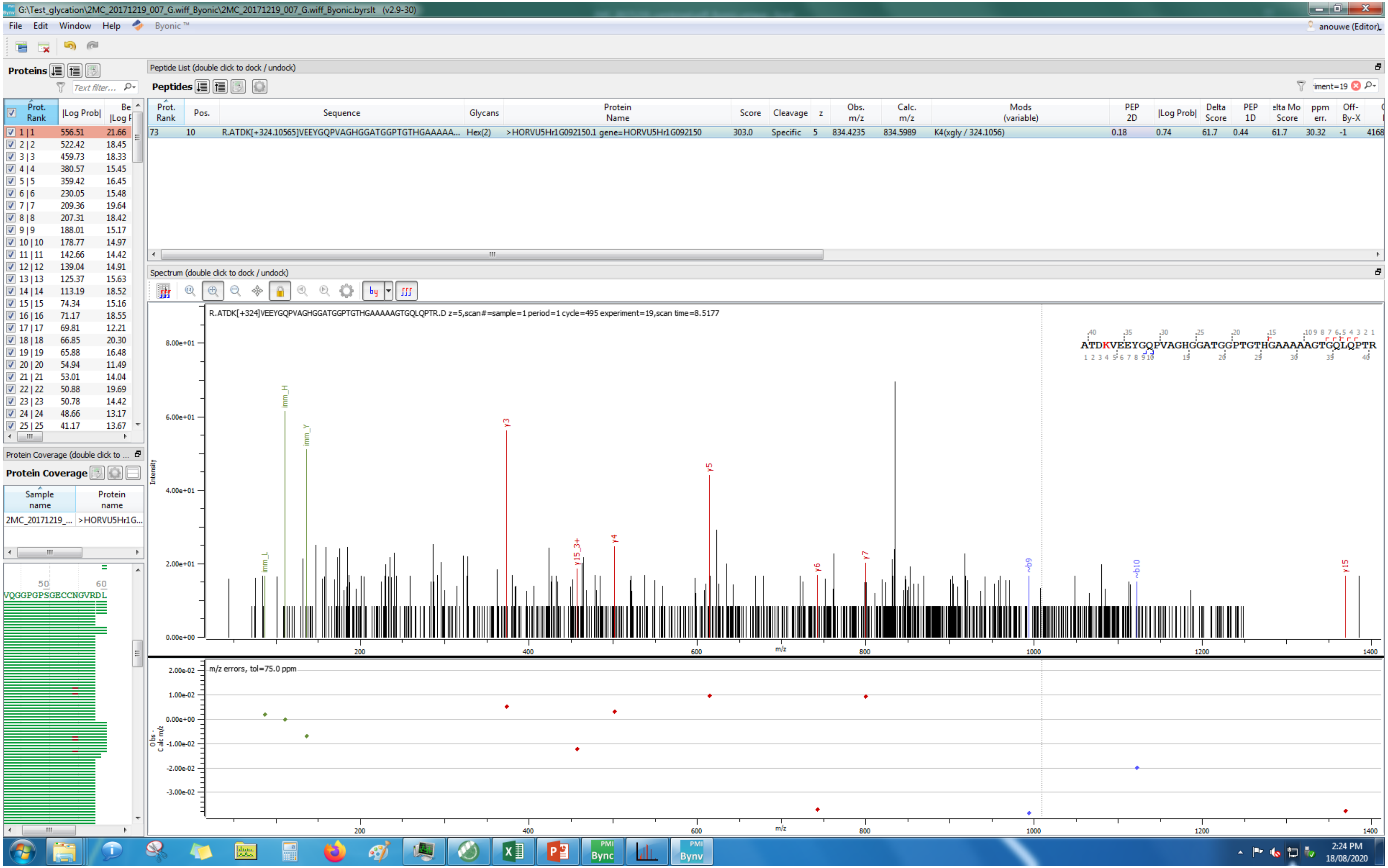
G:\Test_glycation\2MC_20171219_007_G.wiff_Byonic\2MC_20171219_007_G.wiff_Byonic.byrslt (v2.9-30)
File Edit Window Help Byonic™
Proteins
Peptide List (double click to dock / undock)
Peptides
Prot. Rank | Pos. | Sequence | Glycans | Protein Name | Score | Cleavage | z | Obs. m/z | Calc. m/z | Mods (variable) | PEP 2D | |Log Prob| | Delta Score | PEP 1D | Delta Mod Score | ppm err. | Off-By-X
73 | 10 | R.ATDK[+324.10565]VEEYGQPVAGHGGATGGPTGTHGAAAAA... | Hex(2) | >HORVU5Hr1G092150.1 gene=HORVU5Hr1G092150 | 303.0 | Specific | 5 | 834.4235 | 834.5989 | K4(xgly / 324.1056) | 0.18 | 0.74 | 61.7 | 0.44 | 61.7 | 30.32 | -1
Spectrum (double click to dock / undock)
R.ATDK[+324]VEEYGQPVAGHGGATGGPTGTHGAAAAAGTGQLQPTR.D z=5,scan#=sample=1 period=1 cycle=495 experiment=19,scan time=8.5177
ATDKVEEYGQPVAGHGGATGGPTGTHGAAAAAGTGQLQPTR
Intensity
m/z
m/z errors, tol=75.0 ppm
Obs - Calc m/z
Protein Coverage
Sample name
Protein name
2MC_20171219_...
>HORVU5Hr1G...
VQGGPGPSGECCNGVRDL
2:24 PM
18/08/2020

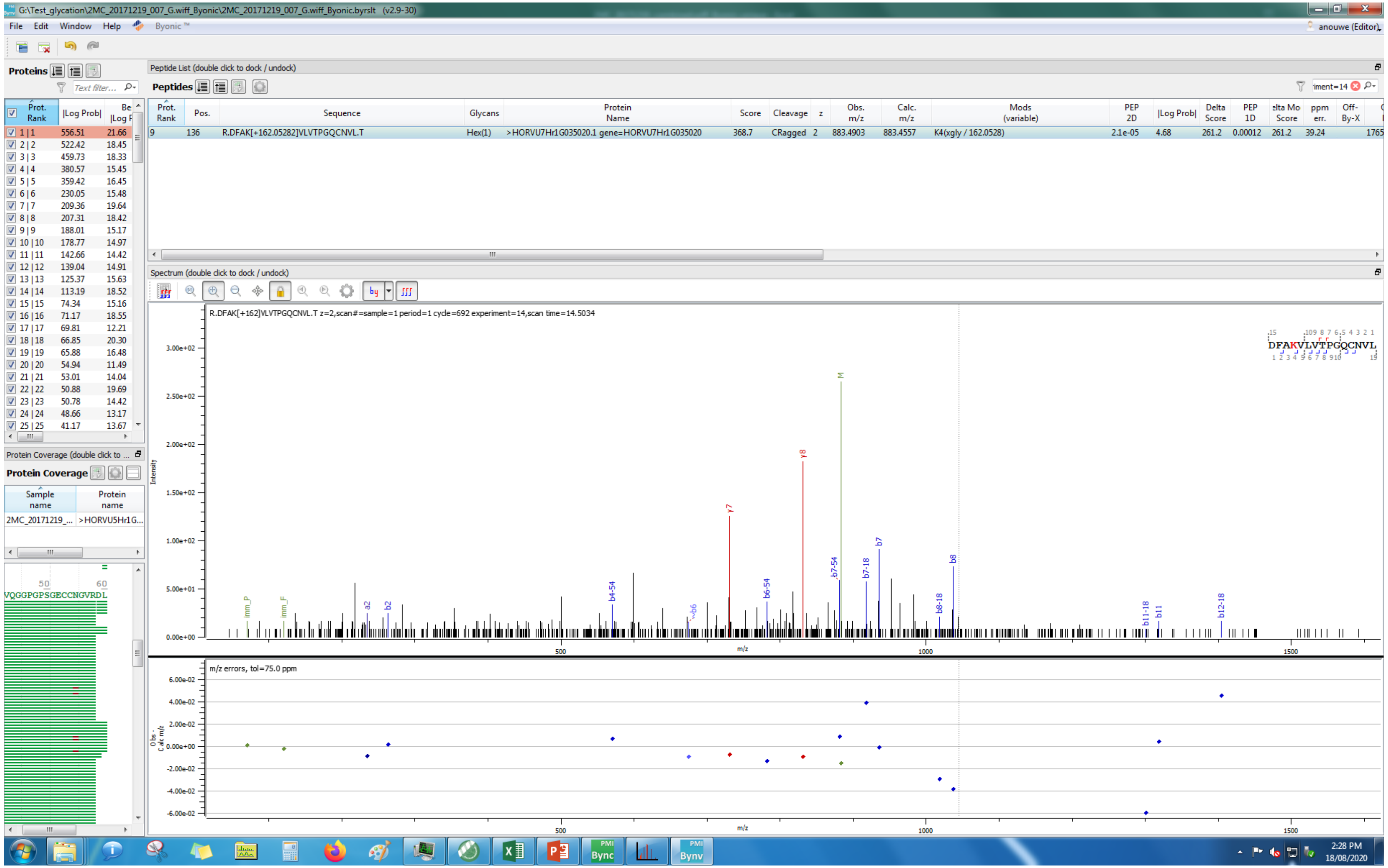
G:\Test_glycation\2MC_20171219_007_G.wiff_Byonic\2MC_20171219_007_G.wiff_Byonic.byrslt (v2.9-30)
File Edit Window Help Byonic ™
anouwe (Editor)
Proteins
Peptide List (double click to dock / undock)
Peptides
Prot. Rank | Pos. | Sequence | Glycans | Protein Name | Score | Cleavage | z | Obs. m/z | Calc. m/z | Mods (variable) | PEP 2D | |Log Prob| | Delta Score | PEP 1D | Delta Mod Score | ppm err. | Off-By-X
9 | 136 | R.DFAK[+162.05282]VLVTPGQCNVL.T | Hex(1) | >HORVU7Hr1G035020.1 gene=HORVU7Hr1G035020 | 368.7 | CRagged | 2 | 883.4903 | 883.4557 | K4(xgly / 162.0528) | 2.1e-05 | 4.68 | 261.2 | 0.00012 | 261.2 | 39.24
Spectrum (double click to dock / undock)
R.DFAK[+162]VLVTPGQCNVL.T z=2,scan#=sample=1 period=1 cycle=692 experiment=14,scan time=14.5034
DFAKVLVTPGQCNVL
Intensity
m/z
m/z errors, tol=75.0 ppm
Obs - Calc m/z
Protein Coverage (double click to ...
Protein Coverage
Sample name | Protein name
2MC_20171219_... | >HORVU5Hr1G...
VQGGPGPSGECCNGVRDL
2:28 PM
18/08/2020

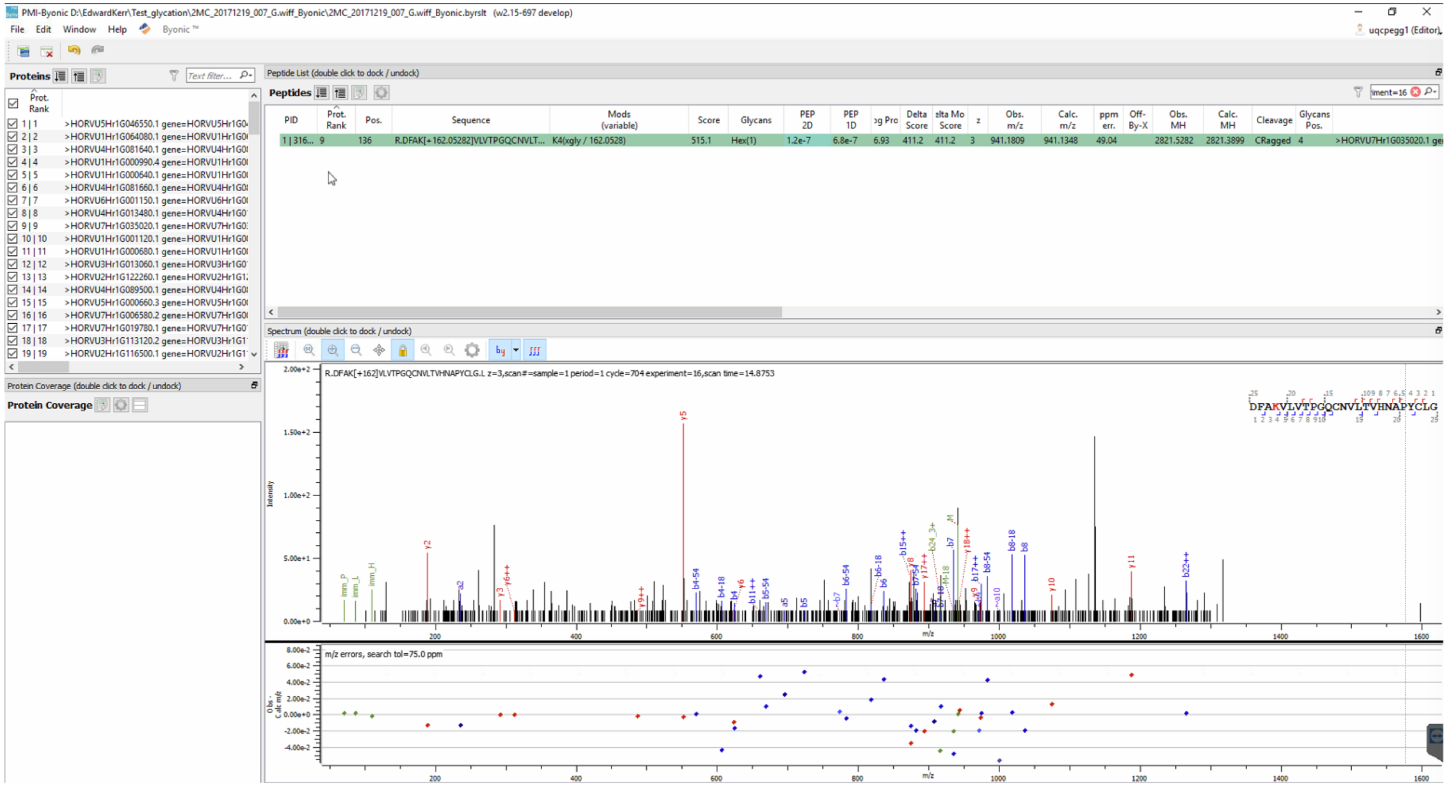
PMI-Byonic D:\EdwardKerr\Test_glycation\2MC_20171219_007_G.wiff_Byonic\2MC_20171219_007_G.wiff_Byonic.byrslt (v2.15-697 develop)
File Edit Window Help Byonic™
Proteins
Peptide List (double click to dock / undock)
Peptides
PID
Prot. Rank
Pos.
Sequence
Mods (variable)
Score
Glycans
PEP 2D
PEP 1D
Delta Score
z
Obs. m/z
Calc. m/z
ppm err.
Off-By-X
Obs. MH
Calc. MH
Cleavage
Glycans Pos.
R.DFAK[+162.05282]VLVTPGQCNVLT...
K4(xgly / 162.0528)
515.1
Hex(1)
1.2e-7
6.8e-7
941.1809
941.1348
49.04
2821.5282
2821.3899
CRagged
Spectrum (double click to dock / undock)
R.DFAK[+162]VLVTPGQCNVLTVHNAPYCLG.L z=3,scan#=sample=1 period=1 cycle=704 experiment=16,scan time=14.8753
DFAKVLVTPGQCNVLTVHNAPYCLG
Intensity
m/z
m/z errors, search tol=75.0 ppm
Protein Coverage (double click to dock / undock)
Protein Coverage

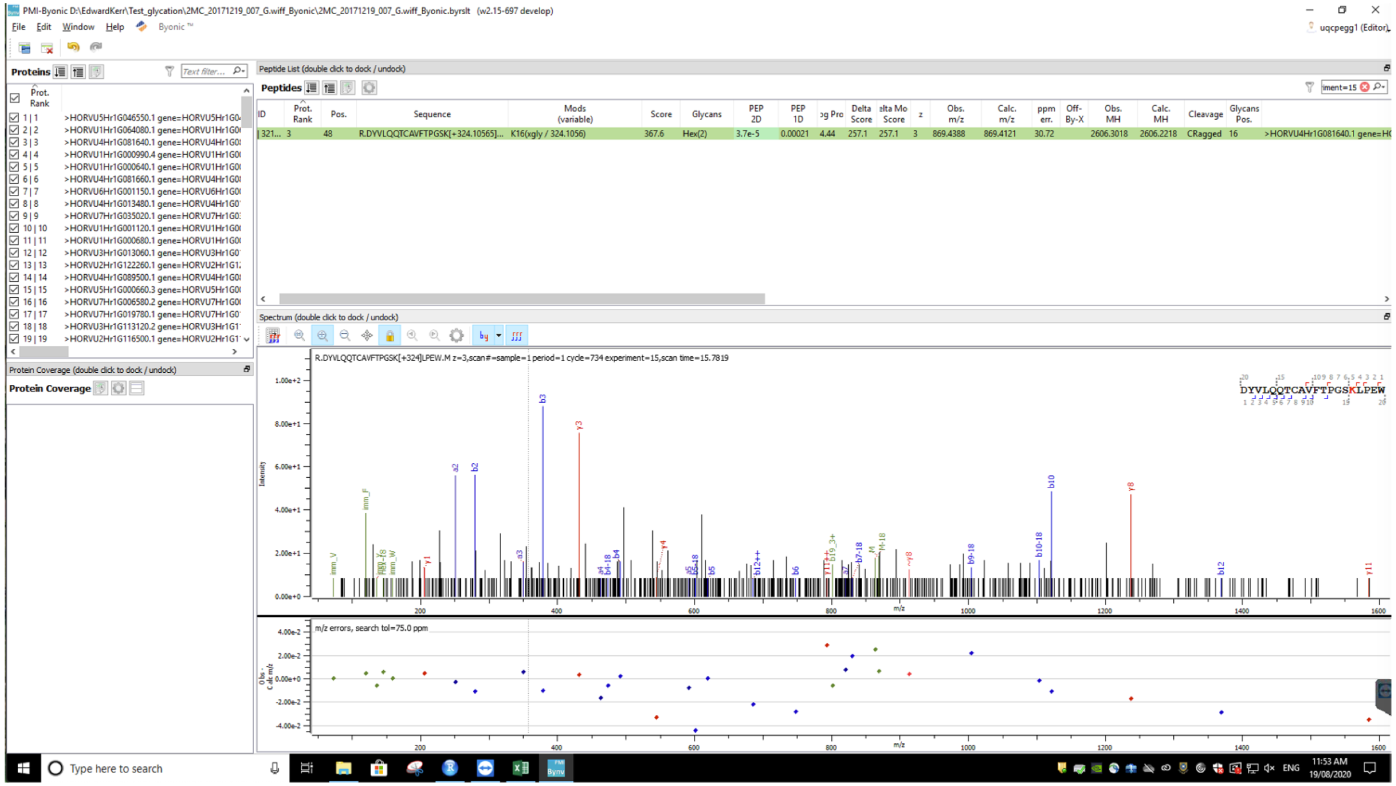

PMI-Byonic
File Edit Window Help Byonic
Proteins
Peptide List (double click to dock / undock)
Peptides
R.DYVLQQTCAVFTPGSK[+324.10565]...
K16(xgly / 324.1056)
Hex(2)
Spectrum (double click to dock / undock)
R.DYVLQQTCAVFTPGSK[+324]LPEW.M z=3,scan#=sample=1 period=1 cycle=734 experiment=15,scan time=15.7819
DYVLQQTCAVFTPGSKLPEW
m/z errors, search tol=75.0 ppm
Protein Coverage (double click to dock / undock)
Protein Coverage
Type here to search

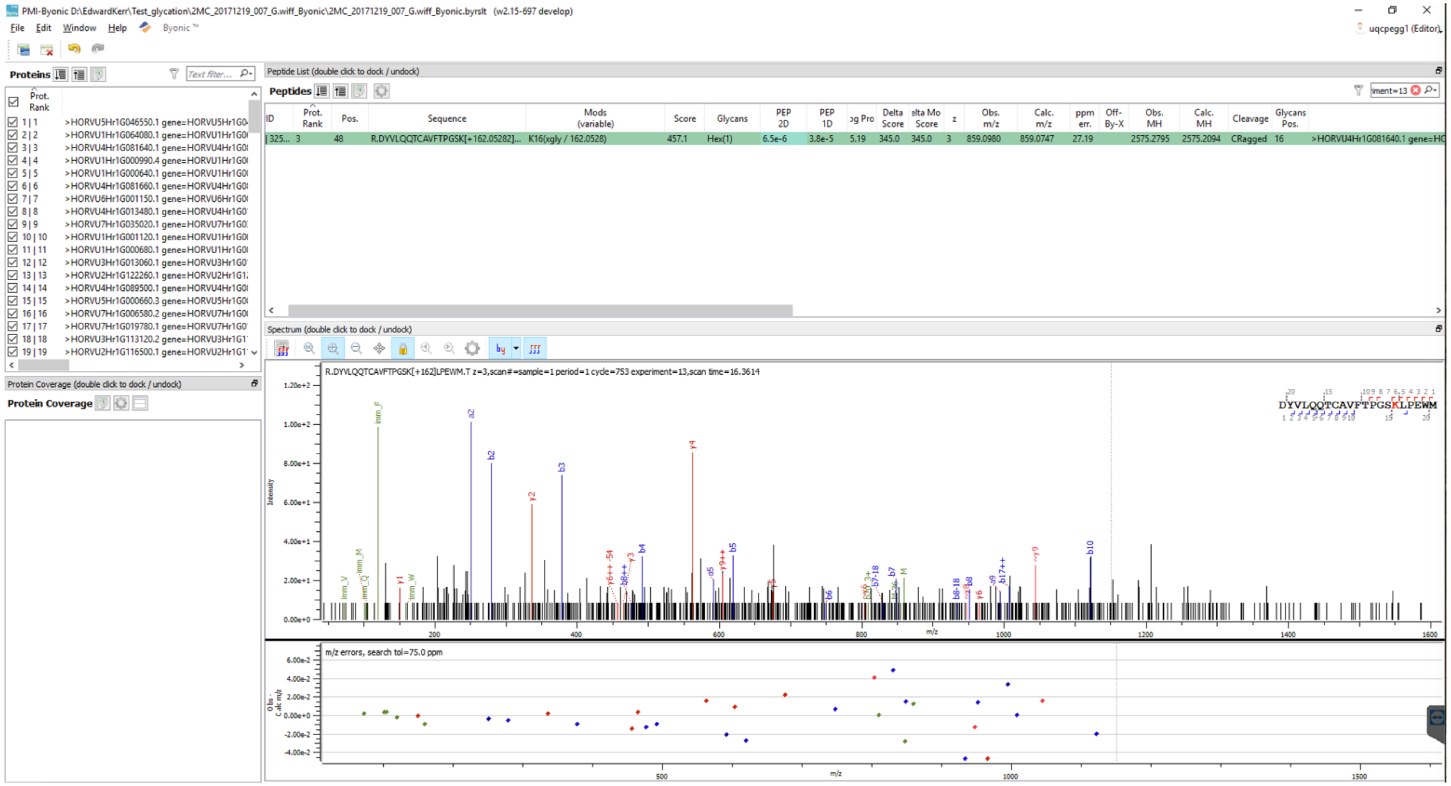
PMI-Byonic D:\EdwardKerr\Test_glycation\2MC_20171219_007_G.wiff_Byonic\2MC_20171219_007_G.wiff_Byonic.byrslt (w2.15-697 develop)
File Edit Window Help Byonic
Proteins
Peptide List (double click to dock / undock)
Peptides
ID
Prot. Rank
Pos.
Sequence
Mods (variable)
Score
Glycans
PEP 2D
PEP 1D
Delta Score
z
Obs. m/z
Calc. m/z
ppm err.
Obs. MH
Calc. MH
Cleavage
Glycans Pos.
R.DYVLQQTCAVFTPGSK[+162.05282]...
K16(xgly / 162.0528)
457.1
Hex(1)
6.5e-6
3.8e-5
5.19
345.0
345.0
3
859.0980
859.0747
27.19
2575.2795
2575.2094
CRagged
16
>HORVU4Hr1G081640.1 gene=HO
Spectrum (double click to dock / undock)
R.DYVLQQTCAVFTPGSK[+162]LPEWM.T z=3,scan#=sample=1 period=1 cycle=753 experiment=13,scan time=16.3614
Intensity
m/z
m/z errors, search tol=75.0 ppm
Protein Coverage (double click to dock / undock)
Protein Coverage

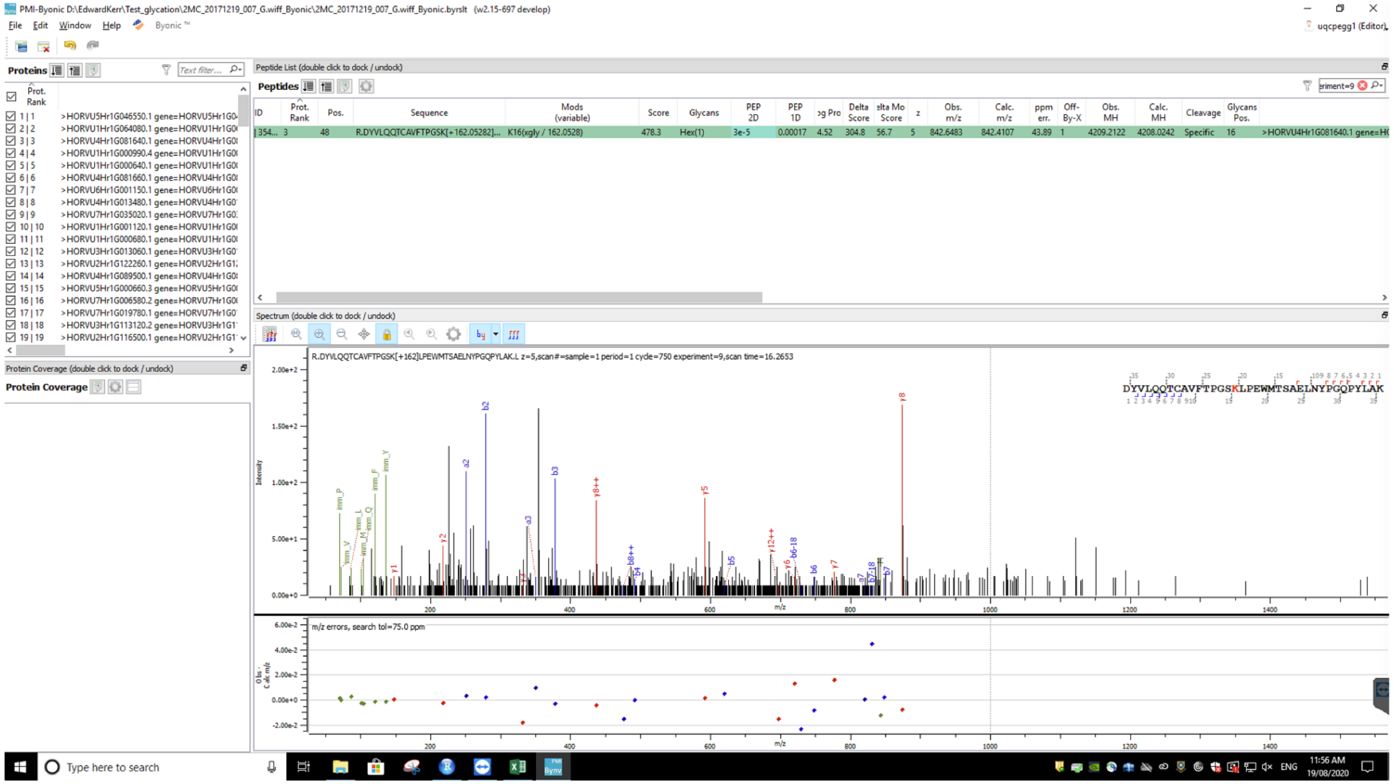
PMI-Byonic D:\EdwardKerr\Test_glycation\2MC_20171219_007_G.wiff_Byonic\2MC_20171219_007_G.wiff_Byonic.byrslt (w2.15-697 develop)
File Edit Window Help Byonic
Proteins
Peptide List (double click to dock / undock)
Peptides
Spectrum (double click to dock / undock)
R.DYVLQQTCAVFTPGSK[+162]LPEWMTSAELNYPGQPYLAK.L z=5,scan#=sample=1 period=1 cycle=750 experiment=9,scan time=16.2653
DYVLQQTCAVFTPGSKLPEWMTSAELNYPGQPYLAK
m/z errors, search tol=75.0 ppm
Protein Coverage (double click to dock / undock)
Protein Coverage
Type here to search
11:56 AM
19/08/2020

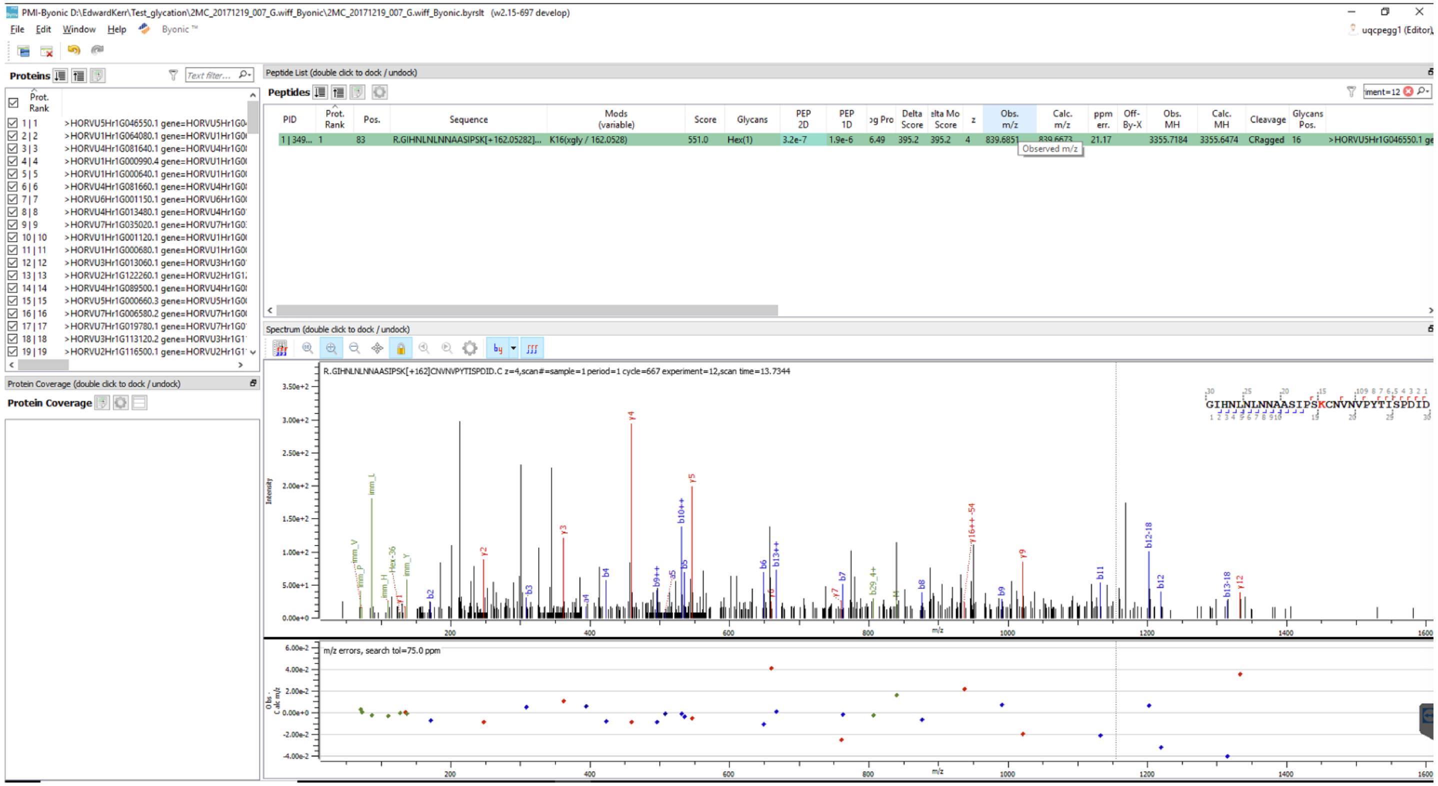
PMI-Byonic D:\EdwardKerr\Test_glycation\2MC_20171219_007_G.wiff_Byonic\2MC_20171219_007_G.wiff_Byonic.byrslt (v2.15-697 develop)
File Edit Window Help Byonic
Proteins
Peptide List (double click to dock / undock)
Peptides
PID
Prot. Rank
Pos.
Sequence
Mods (variable)
Score
Glycans
PEP 2D
PEP 1D
Delta Score
z
Obs. m/z
Calc. m/z
ppm err.
Obs. MH
Calc. MH
Cleavage
Glycans Pos.
R.GIHNLNLNNAASIPSK[+162.05282]...
K16(xgly / 162.0528)
551.0
Hex(1)
3.2e-7
1.9e-6
6.49
395.2
839.6851
21.17
3355.7184
3355.6474
CRagged
Observed m/z
Protein Coverage (double click to dock / undock)
Protein Coverage
Spectrum (double click to dock / undock)
R.GIHNLNLNNAASIPSK[+162]CNVNVPYTISPDID.C z=4,scan#=sample=1 period=1 cycle=667 experiment=12,scan time=13.7344
GIHNLNLNNAASIPSKCNVNVPYTISPDID
Intensity
m/z
m/z errors, search tol=75.0 ppm

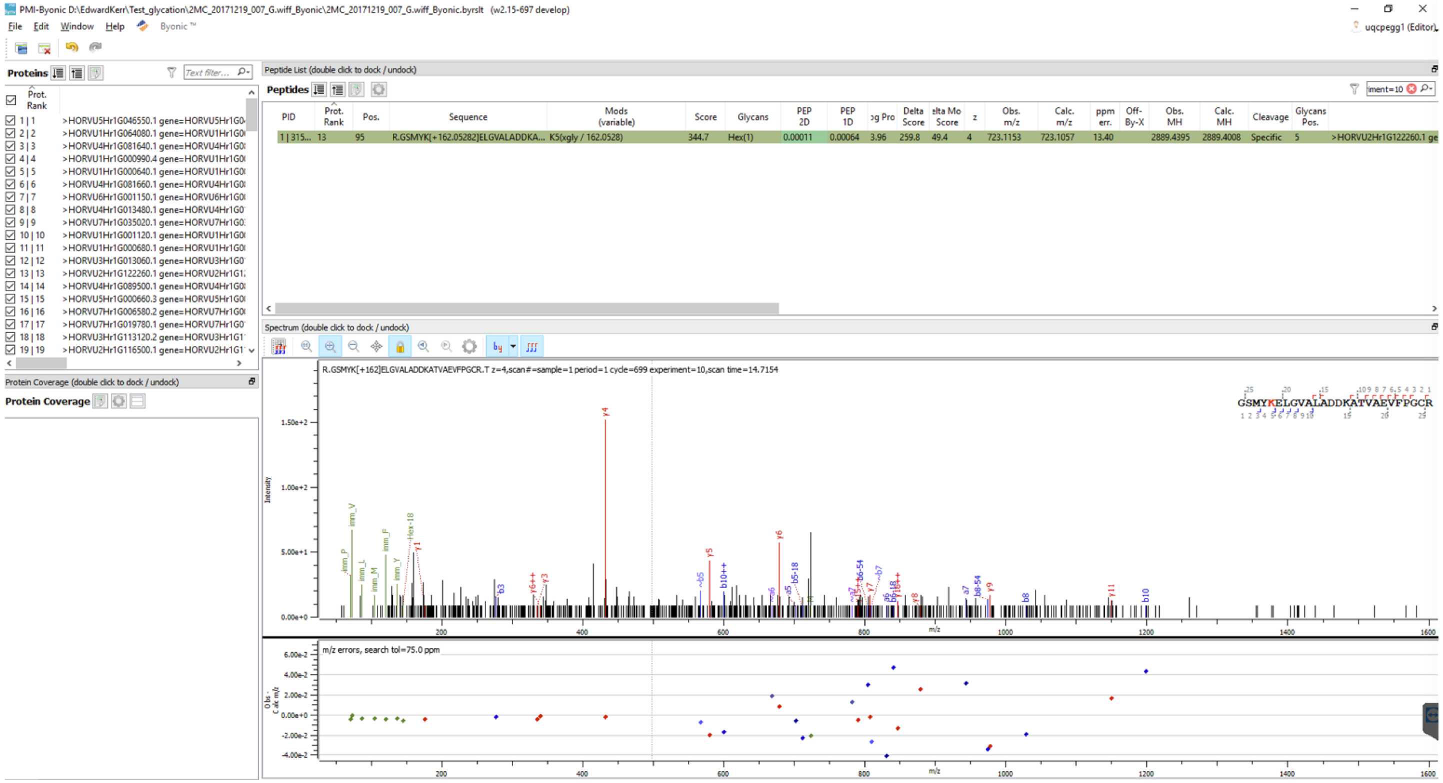
PMI-Byonic D:\EdwardKerr\Test_glycation\2MC_20171219_007_G.wiff_Byonic\2MC_20171219_007_G.wiff_Byonic.byrslt (w2.15-697 develop)
File Edit Window Help Byonic
Proteins
Peptide List (double click to dock / undock)
Peptides
R.GSMYK[+162.05282]ELGVALADDKA... K5(xgly / 162.0528)
Spectrum (double click to dock / undock)
R.GSMYK[+162]ELGVALADDKATVAEVFPGCR.T z=4,scan#=sample=1 period=1 cycle=699 experiment=10,scan time=14.7154
GSMYKELGVALADDKATVAEVFPGCR
m/z errors, search tol=75.0 ppm
Protein Coverage (double click to dock / undock)
Protein Coverage

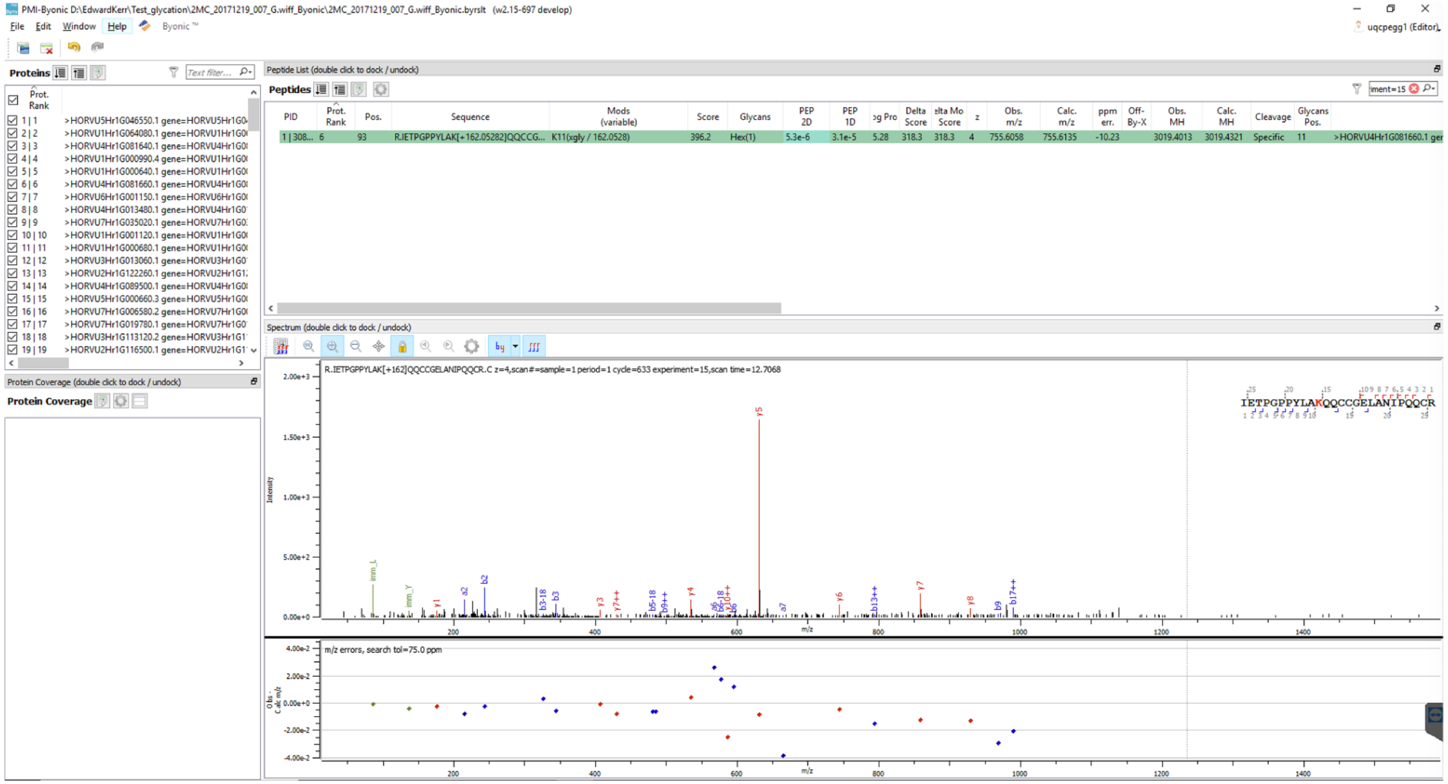
PMI-Byonic D:\EdwardKerr\Test_glycation\2MC_20171219_007_G.wiff_Byonic\2MC_20171219_007_G.wiff_Byonic.byrslt (w2.15-697 develop)
File Edit Window Help Byonic
Proteins
Peptide List (double click to dock / undock)
Peptides
PID Prot. Rank Pos. Sequence Mods (variable) Score Glycans PEP 2D PEP 1D Delta Score z Obs. m/z Calc. m/z ppm err. Off-By-X Obs. MH Calc. MH Cleavage Glycans Pos.
1 | 308... 6 93 R.IETPGPPYLAK[+162.05282]QQCCG... K11(xgly / 162.0528) 396.2 Hex(1) 5.3e-6 3.1e-5 5.28 318.3 318.3 4 755.6058 755.6135 -10.23 3019.4013 3019.4321 Specific 11
Spectrum (double click to dock / undock)
R.IETPGPPYLAK[+162]QQCCGELANIPQQCR.C z=4,scan#=sample=1 period=1 cycle=633 experiment=15,scan time=12.7068
IETPGPPYLAKQQCCGELANIPQQCR
Intensity
m/z
m/z errors, search tol=75.0 ppm
Protein Coverage (double click to dock / undock)
Protein Coverage

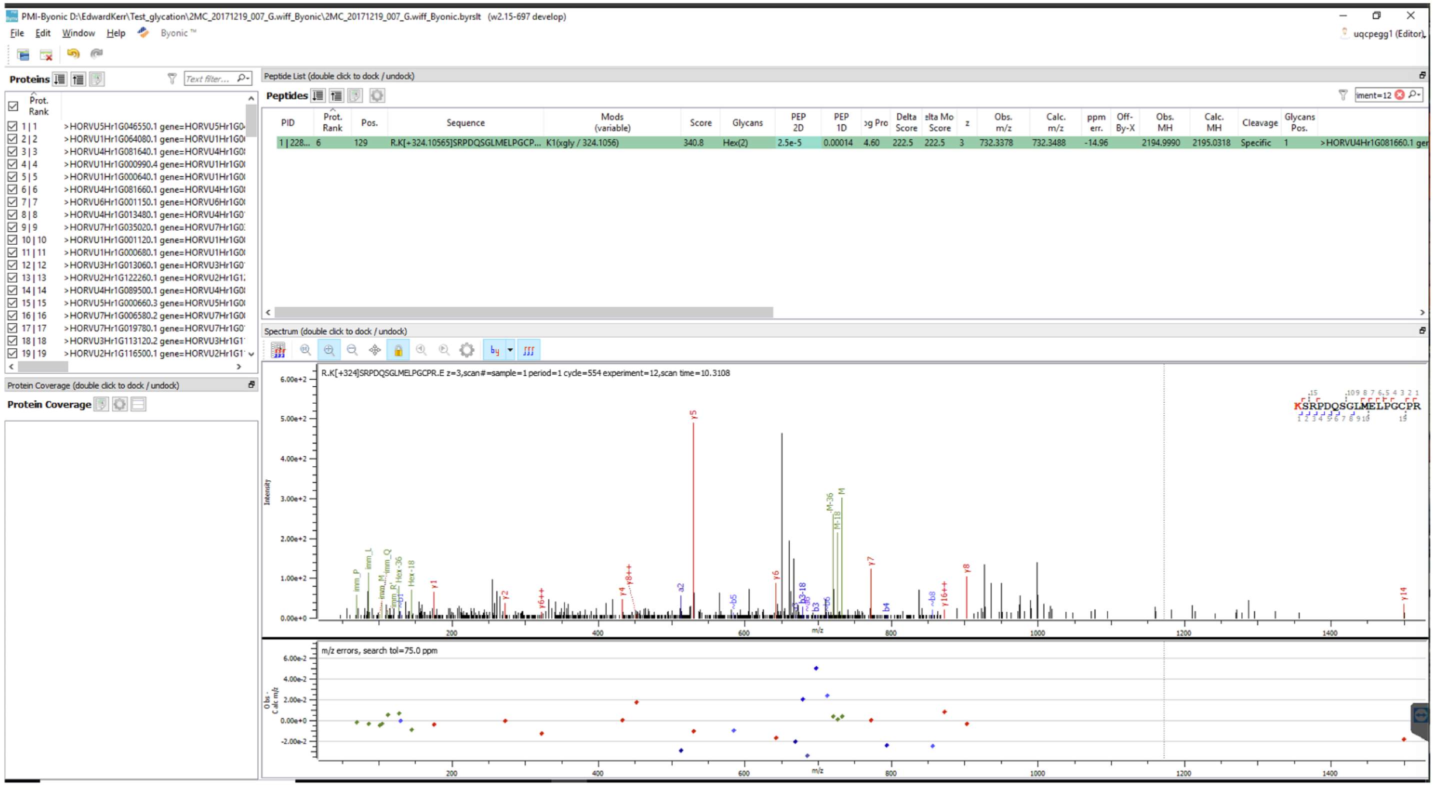
PMI-Byonic D:\EdwardKerr\Test_glycation\2MC_20171219_007_G.wiff_Byonic\2MC_20171219_007_G.wiff_Byonic.byrslt (v2.15-697 develop)
File Edit Window Help Byonic
Proteins
Peptide List (double click to dock / undock)
Peptides
PID
Prot. Rank
Pos.
Sequence
Mods (variable)
Score
Glycans
PEP 2D
PEP 1D
Delta Score
z
Obs. m/z
Calc. m/z
ppm err.
Obs. MH
Calc. MH
Cleavage
Glycans Pos.
1 | 228... 6 129 R.K[+324.10565]SRPDQSGLMELPGCP... K1(xgly / 324.1056) 340.8 Hex(2) 2.5e-5 0.00014 4.60 222.5 222.5 3 732.3378 732.3488 -14.96 2194.9990 2195.0318 Specific 1
Spectrum (double click to dock / undock)
R.K[+324]SRPDQSGLMELPGCPR.E z=3,scan#=sample=1 period=1 cycle=554 experiment=12,scan time=10.3108
Protein Coverage (double click to dock / undock)
Protein Coverage
m/z errors, search tol=75.0 ppm
m/z

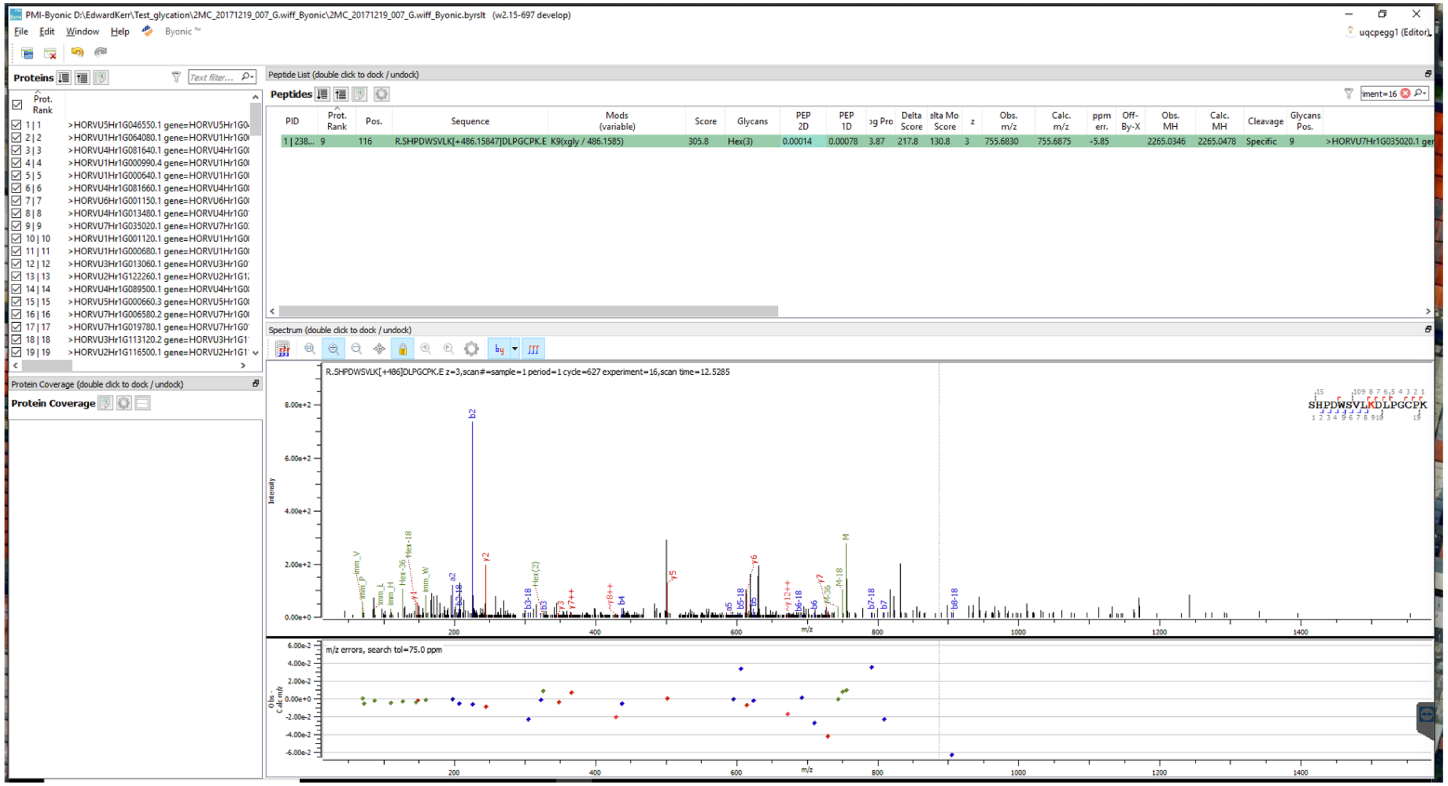
PMI-Byonic D:\EdwardKerr\Test_glycation\2MC_20171219_007_G.wiff_Byonic\2MC_20171219_007_G.wiff_Byonic.byrslt (w2.15-697 develop)
File Edit Window Help Byonic ™
uqcpegg1 (Editor)
Proteins
Peptide List (double click to dock / undock)
Peptides
PID | Prot. Rank | Pos. | Sequence | Mods (variable) | Score | Glycans | PEP 2D | PEP 1D | Delta Score | z | Obs. m/z | Calc. m/z | ppm err. | Off-By-X | Obs. MH | Calc. MH | Cleavage | Glycans Pos.
1 | 238... 9 116 R.SHPDWSVLK[+486.15847]DLPGCPK.E K9(xgly / 486.1585) 305.8 Hex(3) 0.00014 0.00078 3.87 217.8 130.8 3 755.6830 755.6875 -5.85 2265.0346 2265.0478 Specific 9 >HORVU7Hr1G035020.1
Spectrum (double click to dock / undock)
R.SHPDWSVLK[+486]DLPGCPK.E z=3,scan#=sample=1 period=1 cycle=627 experiment=16,scan time=12.5285
Protein Coverage (double click to dock / undock)
Protein Coverage
m/z errors, search tol=75.0 ppm

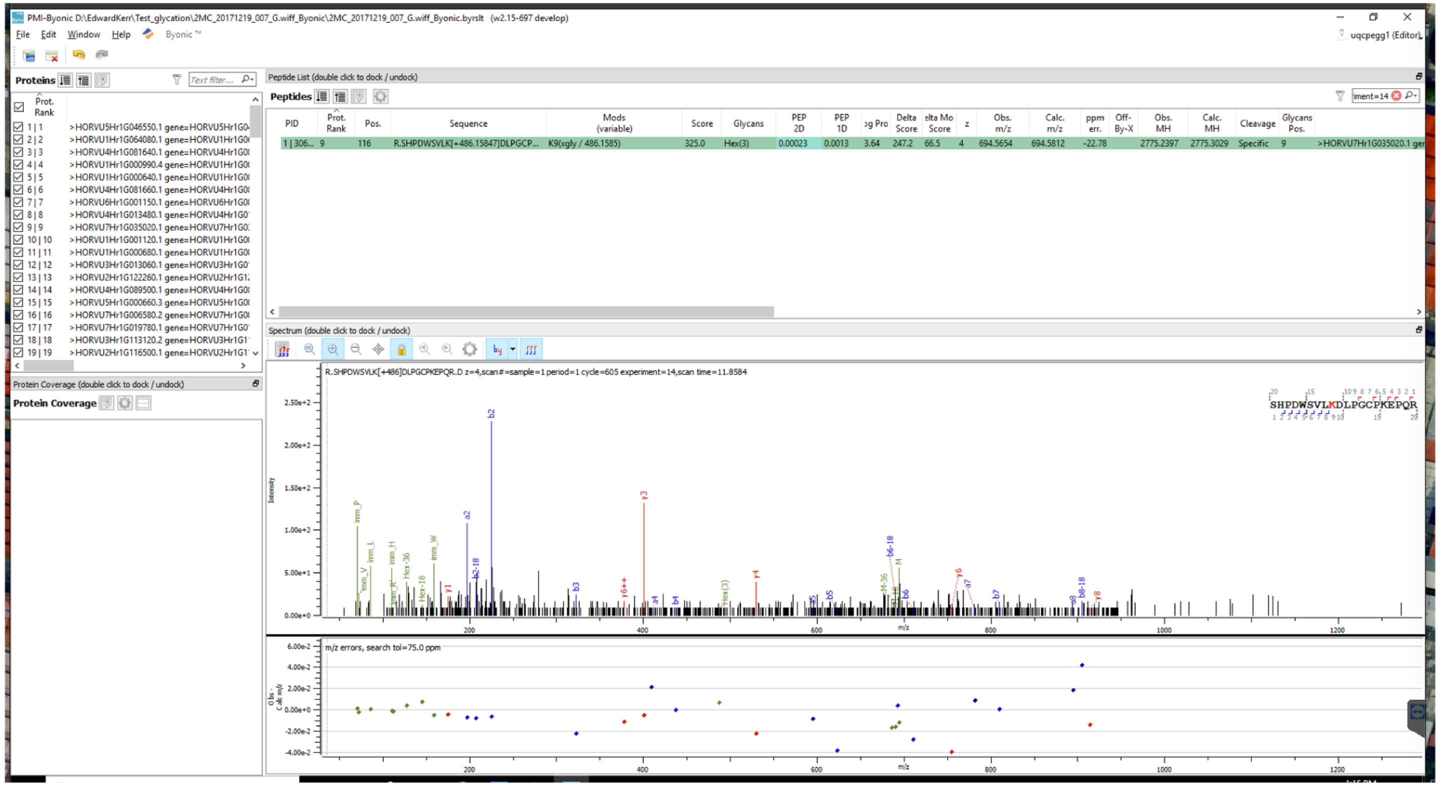

PMI-Byonic D:\EdwardKerr\Test_glycation\2MC_20171219_007_G.wiff_Byonic\2MC_20171219_007_G.wiff_Byonic.byrslt (w2.15-697 develop)
File Edit Window Help Byonic
Proteins
Peptide List (double click to dock / undock)
Peptides
R.SHPDWSVLK[+486.15847]DLPGCP...
K9(xgly / 486.1585)
Spectrum (double click to dock / undock)
R.SHPDWSVLK[+486]DLPGCPKEPQR.D z=4,scan#=sample=1 period=1 cycle=605 experiment=14,scan time=11.8584
SHPDWSVLKDLPGCPKEPQR
m/z errors, search tol=75.0 ppm
Protein Coverage (double click to dock / undock)
Protein Coverage

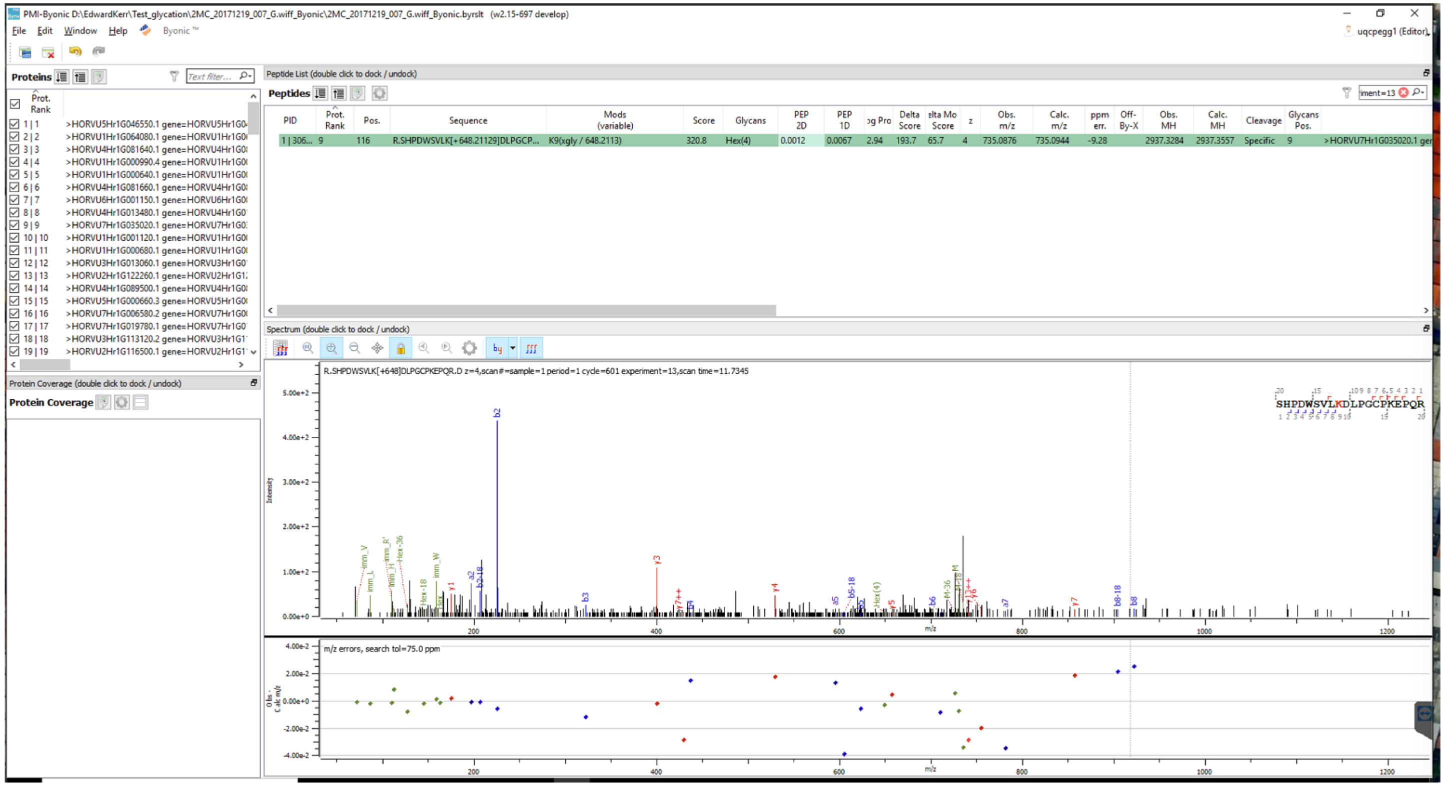
PMI-Byonic D:\EdwardKerr\Test_glycation\2MC_20171219_007_G.wiff_Byonic\2MC_20171219_007_G.wiff_Byonic.byrslt (w2.15-697 develop)
File Edit Window Help Byonic
Proteins
Peptide List (double click to dock / undock)
Peptides
PID
Prot. Rank
Pos.
Sequence
Mods (variable)
Score
Glycans
PEP 2D
PEP 1D
Delta Score
Obs. m/z
Calc. m/z
ppm err.
Off-By-X
Obs. MH
Calc. MH
Cleavage
Glycans Pos.
R.SHPDWSVLK[+648.21129]DLPGCP...
K9(xgly / 648.2113)
320.8
Hex(4)
0.0012
0.0067
735.0876
735.0944
-9.28
2937.3284
2937.3557
Specific
Spectrum (double click to dock / undock)
R.SHPDWSVLK[+648]DLPGCPKEPQR.D z=4,scan#=sample=1 period=1 cycle=601 experiment=13,scan time=11.7345
SHPDWSVLKDLPGCPKEPQR
Intensity
m/z
m/z errors, search tol=75.0 ppm
Protein Coverage (double click to dock / undock)
Protein Coverage

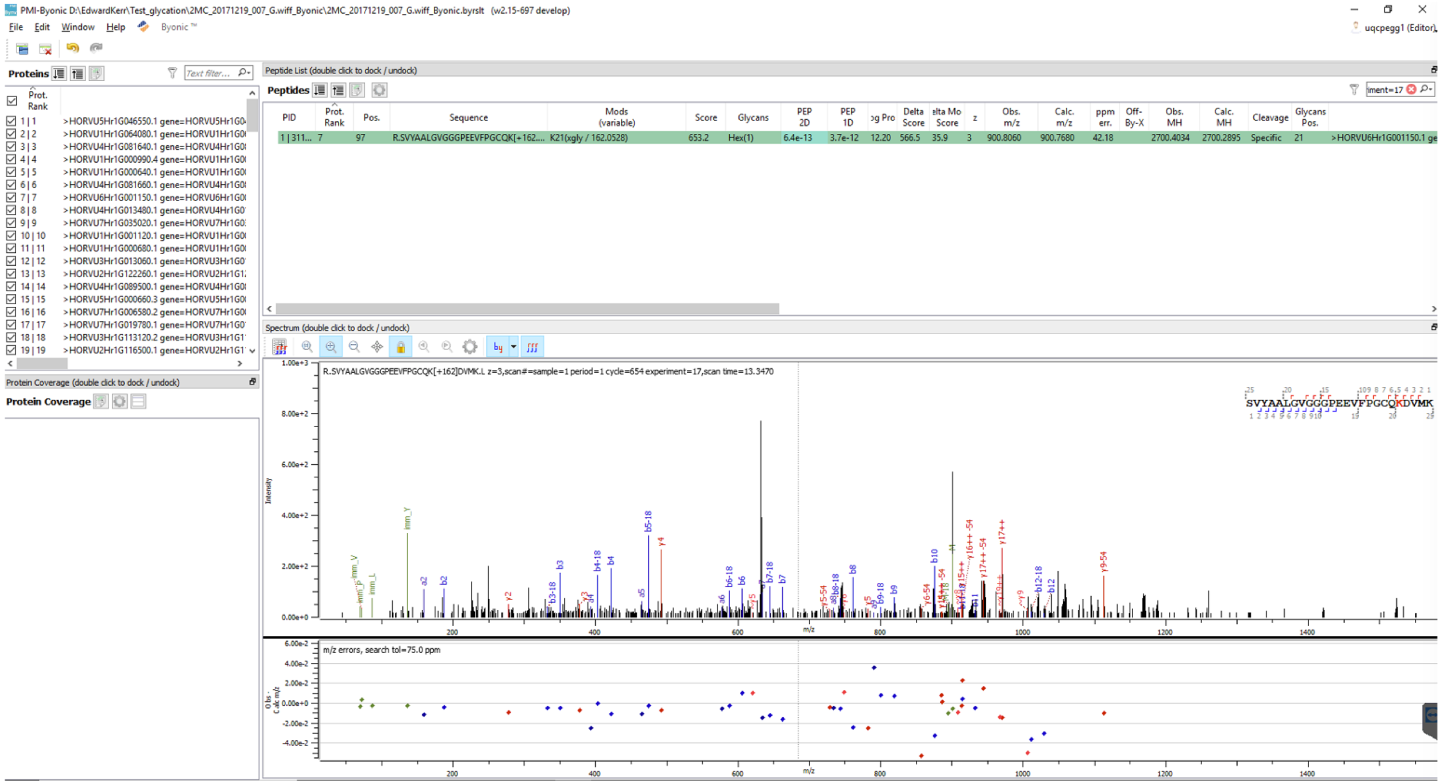
PMI-Byonic D:\EdwardKerr\Test_glycation\2MC_20171219_007_G.wiff_Byonic\2MC_20171219_007_G.wiff_Byonic.byrslt (v2.15-697 develop)
File Edit Window Help Byonic
Proteins
Peptide List (double click to dock / undock)
Peptides
PID
Prot. Rank
Pos.
Sequence
Mods (variable)
Score
Glycans
PEP 2D
PEP 1D
Delta Score
Obs. m/z
Calc. m/z
ppm err.
Obs. MH
Calc. MH
Cleavage
Glycans Pos.
R.SVYAALGVGGGPEEVFPGCQK[+162.... K21(xgly / 162.0528)
Spectrum (double click to dock / undock)
R.SVYAALGVGGGPEEVFPGCQK[+162]DVMK.L z=3,scan#=sample=1 period=1 cycle=654 experiment=17,scan time=13.3470
SVYAALGVGGGPEEVFPGCQKDVMK
Protein Coverage (double click to dock / undock)
Protein Coverage
m/z errors, search tol=75.0 ppm

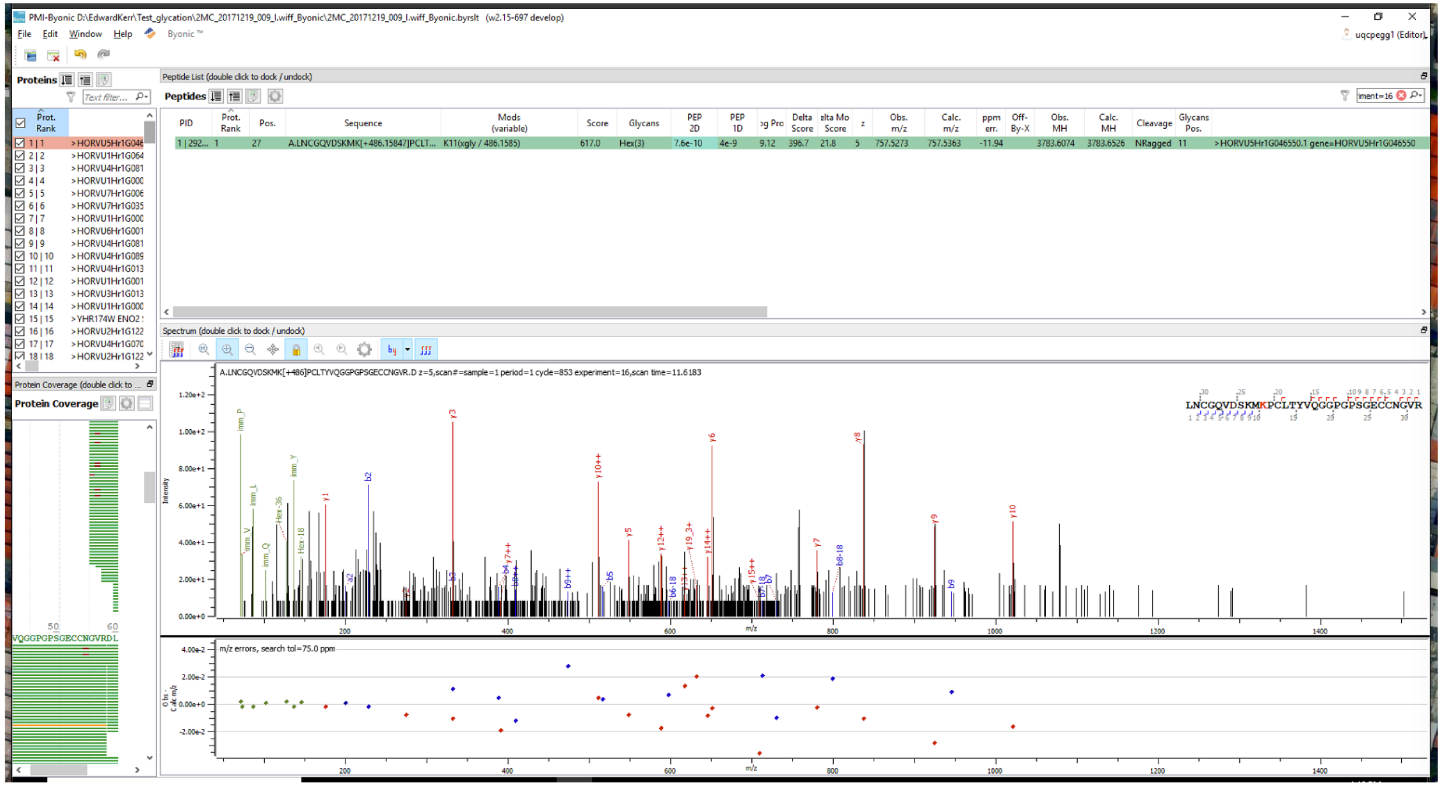
PMI-Byonic D:\EdwardKerr\Test_glycation\2MC_20171219_009_I.wiff_Byonic\2MC_20171219_009_I.wiff_Byonic.byrslt (v2.15-697 develop)
File Edit Window Help Byonic
Proteins
Peptide List (double click to dock / undock)
Peptides
Spectrum (double click to dock / undock)
A.LNCGQVDSKMK[+486]PCLTYVQGGPGPSGECCNGVR.D z=5,scan#=sample=1 period=1 cycle=853 experiment=16,scan time=11.6183
LNCGQVDSKMKPCLTYVQGGPGPSGECCNGVR
Protein Coverage
m/z errors, search tol=75.0 ppm
Intensity

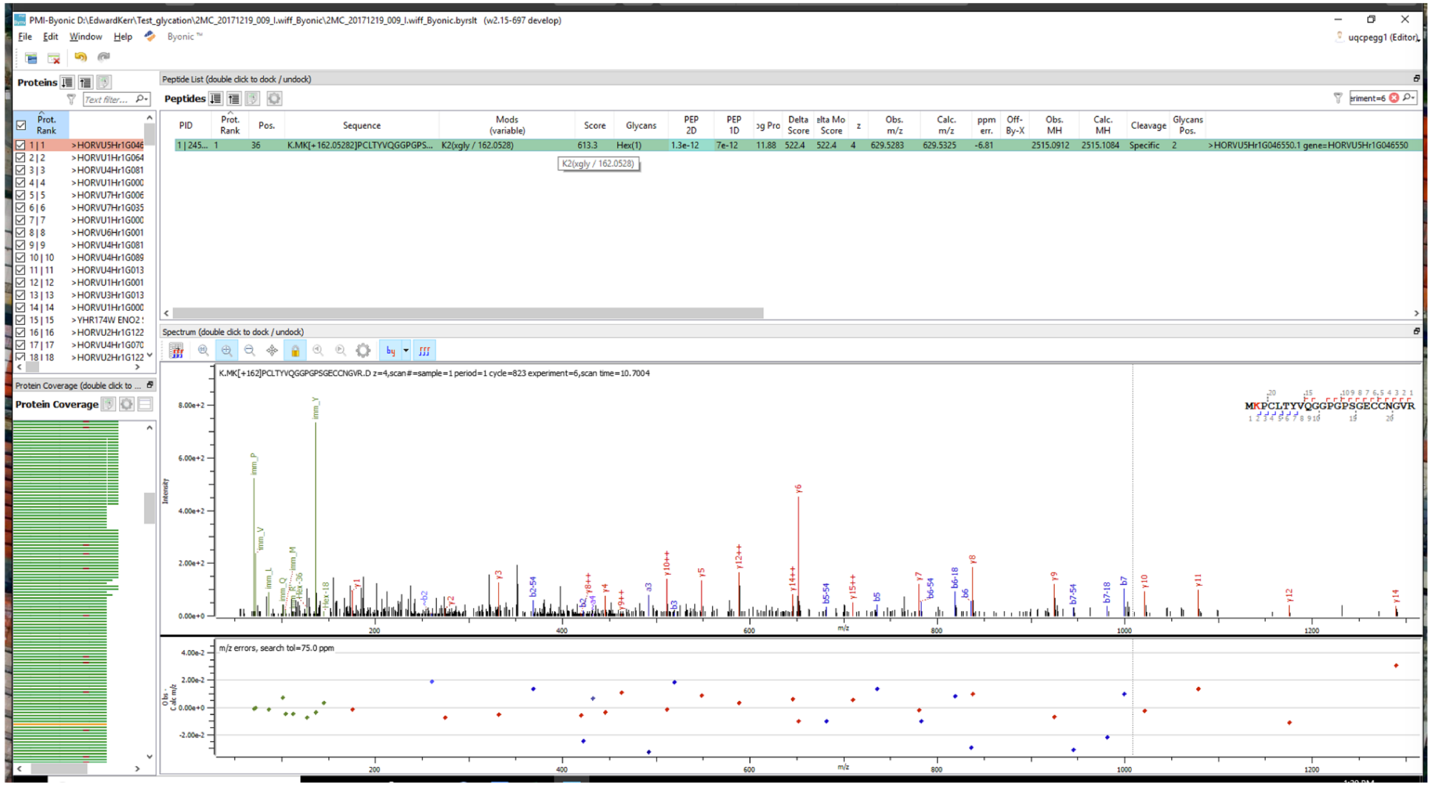
PMI-Byonic D:\EdwardKerr\Test_glycation\2MC_20171219_009_l.wiff_Byonic\2MC_20171219_009_l.wiff_Byonic.byrslt (w2.15-697 develop)
File Edit Window Help
Proteins
Peptide List (double click to dock / undock)
Peptides
PID
Prot. Rank
Pos.
Sequence
Mods (variable)
Score
Glycans
PEP 2D
PEP 1D
Obs. m/z
Calc. m/z
ppm err.
Off-By-X
Obs. MH
Calc. MH
Cleavage
Glycans Pos.
K.MK[+162.05282]PCLTYVQGGPGPS...
K2(xgly / 162.0528)
Hex(1)
Specific
Spectrum (double click to dock / undock)
K.MK[+162]PCLTYVQGGPGPSGECCNGVR.D z=4,scan#=sample=1 period=1 cycle=823 experiment=6,scan time=10.7004
MKPCLTYVQGGPGPSGECCNGVR
Intensity
m/z
m/z errors, search tol=75.0 ppm
Protein Coverage

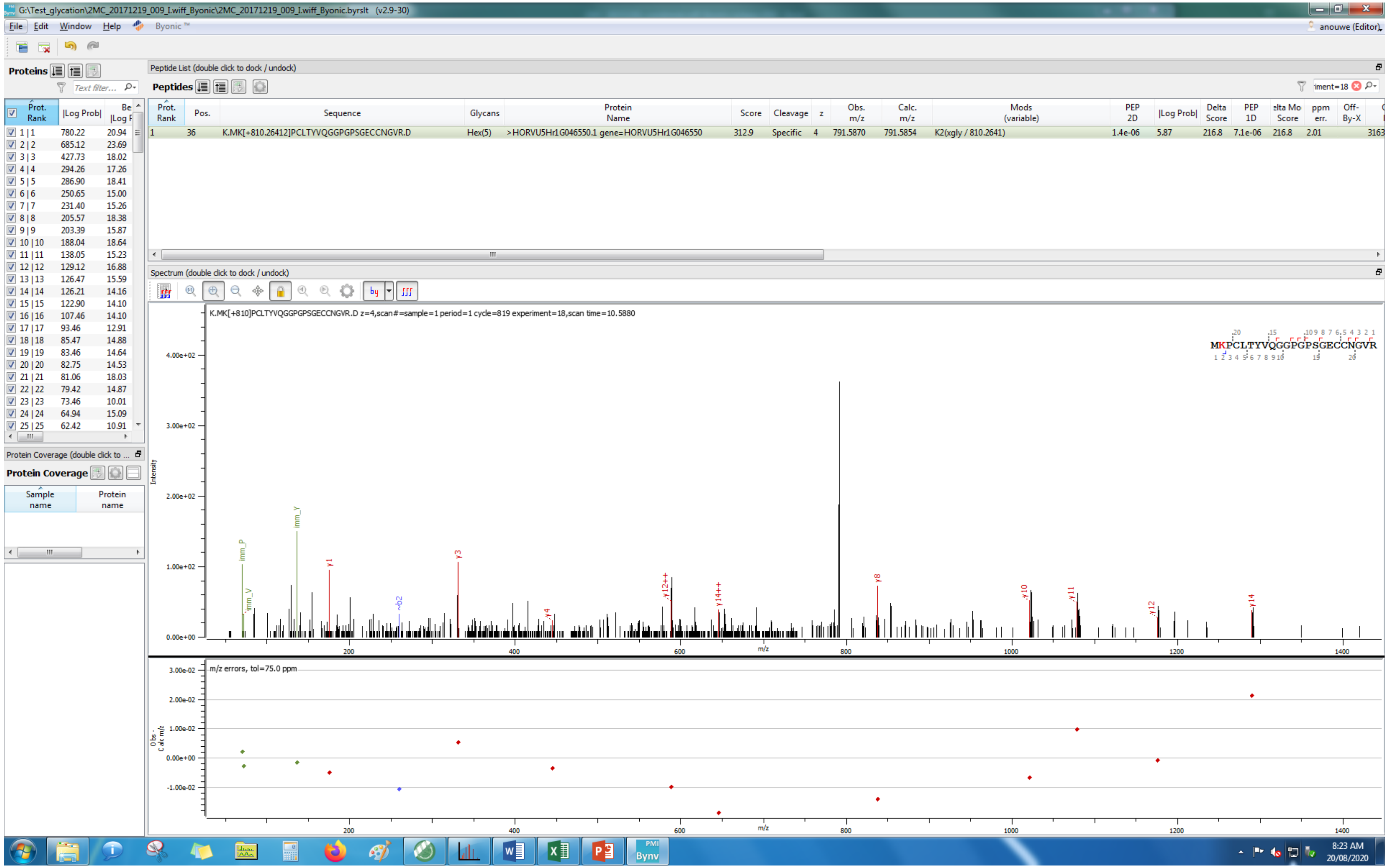
G:\Test_glycation\2MC_20171219_009_I.wiff_Byonic\2MC_20171219_009_I.wiff_Byonic.byrslt (v2.9-30)
File Edit Window Help Byonic™
anouwe (Editor)
Proteins
Peptide List (double click to dock / undock)
Peptides
Prot. Rank | Pos. | Sequence | Glycans | Protein Name | Score | Cleavage | z | Obs. m/z | Calc. m/z | Mods (variable) | PEP 2D | |Log Prob| | Delta Score | PEP 1D | Delta Mod Score | ppm err. | Off-By-X
1 | 36 | K.MK[+810.26412]PCLTYVQGGPGPSGECCNGVR.D | Hex(5) | >HORVU5Hr1G046550.1 gene=HORVU5Hr1G046550 | 312.9 | Specific | 4 | 791.5870 | 791.5854 | K2(xgly / 810.2641) | 1.4e-06 | 5.87 | 216.8 | 7.1e-06 | 216.8 | 2.01
Spectrum (double click to dock / undock)
K.MK[+810]PCLTYVQGGPGPSGECCNGVR.D z=4,scan#=sample=1 period=1 cycle=819 experiment=18,scan time=10.5880
MKPCLTYVQGGPGPSGECCNGVR
Intensity
m/z
m/z errors, tol=75.0 ppm
Obs - Calc m/z
Protein Coverage (double click to ...
Protein Coverage
Sample name
Protein name
8:23 AM
20/08/2020

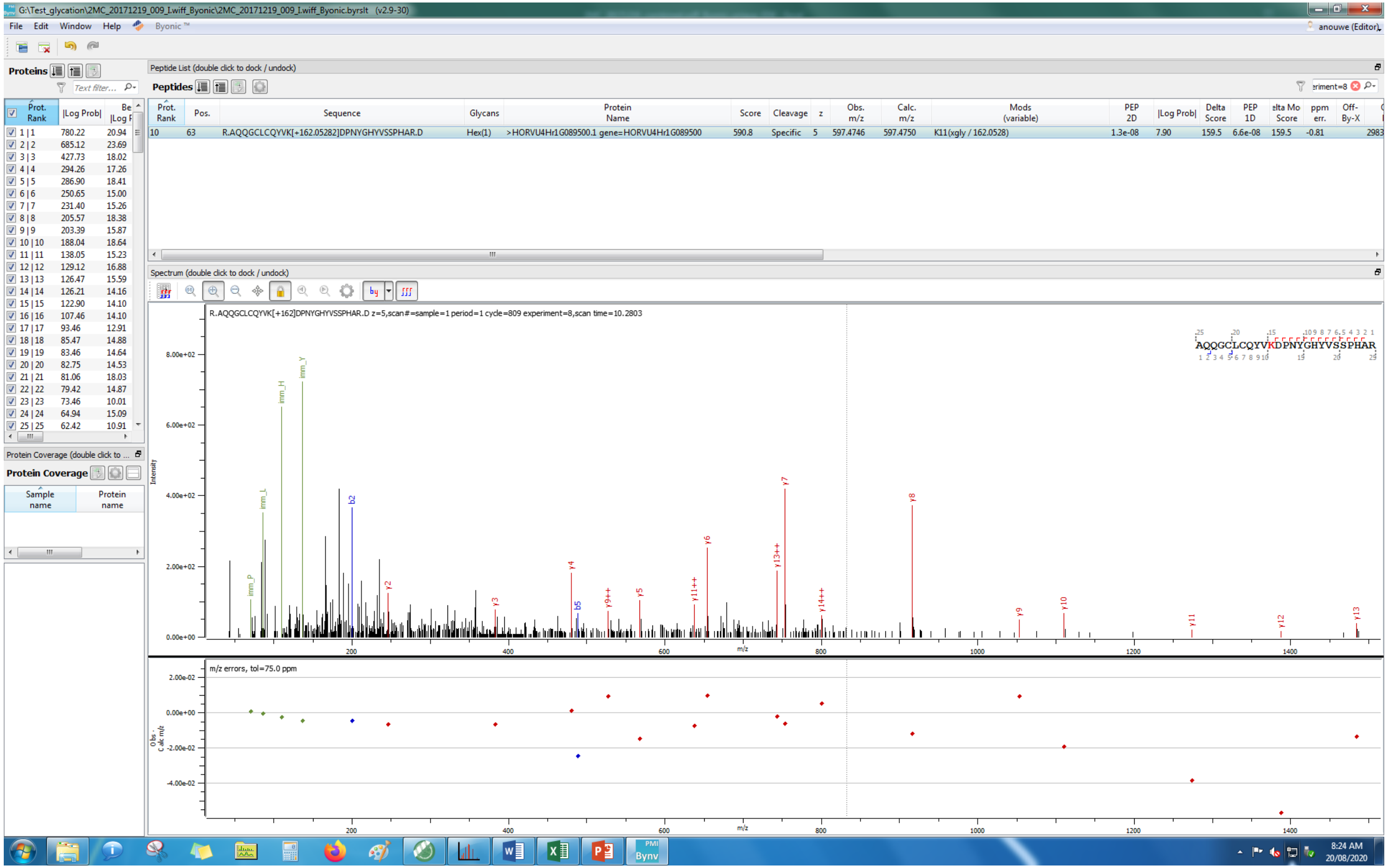
G:\Test_glycation\2MC_20171219_009_I.wiff_Byonic\2MC_20171219_009_I.wiff_Byonic.byrslt (v2.9-30)
File Edit Window Help Byonic™
Proteins
Peptide List (double click to dock / undock)
Peptides
Prot. Rank | Pos. | Sequence | Glycans | Protein Name | Score | Cleavage | z | Obs. m/z | Calc. m/z | Mods (variable) | PEP 2D | |Log Prob| | Delta Score | PEP 1D | Delta Mod Score | ppm err. | Off-By-X
10 | 63 | R.AQQGCLCQYVK[+162.05282]DPNYGHYVSSPHAR.D | Hex(1) | >HORVU4Hr1G089500.1 gene=HORVU4Hr1G089500 | 590.8 | Specific | 5 | 597.4746 | 597.4750 | K11(xgly / 162.0528) | 1.3e-08 | 7.90 | 159.5 | 6.6e-08 | 159.5 | -0.81 | 2983
Spectrum (double click to dock / undock)
R.AQQGCLCQYVK[+162]DPNYGHYVSSPHAR.D z=5,scan#=sample=1 period=1 cycle=809 experiment=8,scan time=10.2803
AQQGCLCQYVKDPNYGHYVSSPHAR
Intensity
m/z
m/z errors, tol=75.0 ppm
Protein Coverage
Sample name
Protein name
8:24 AM
20/08/2020

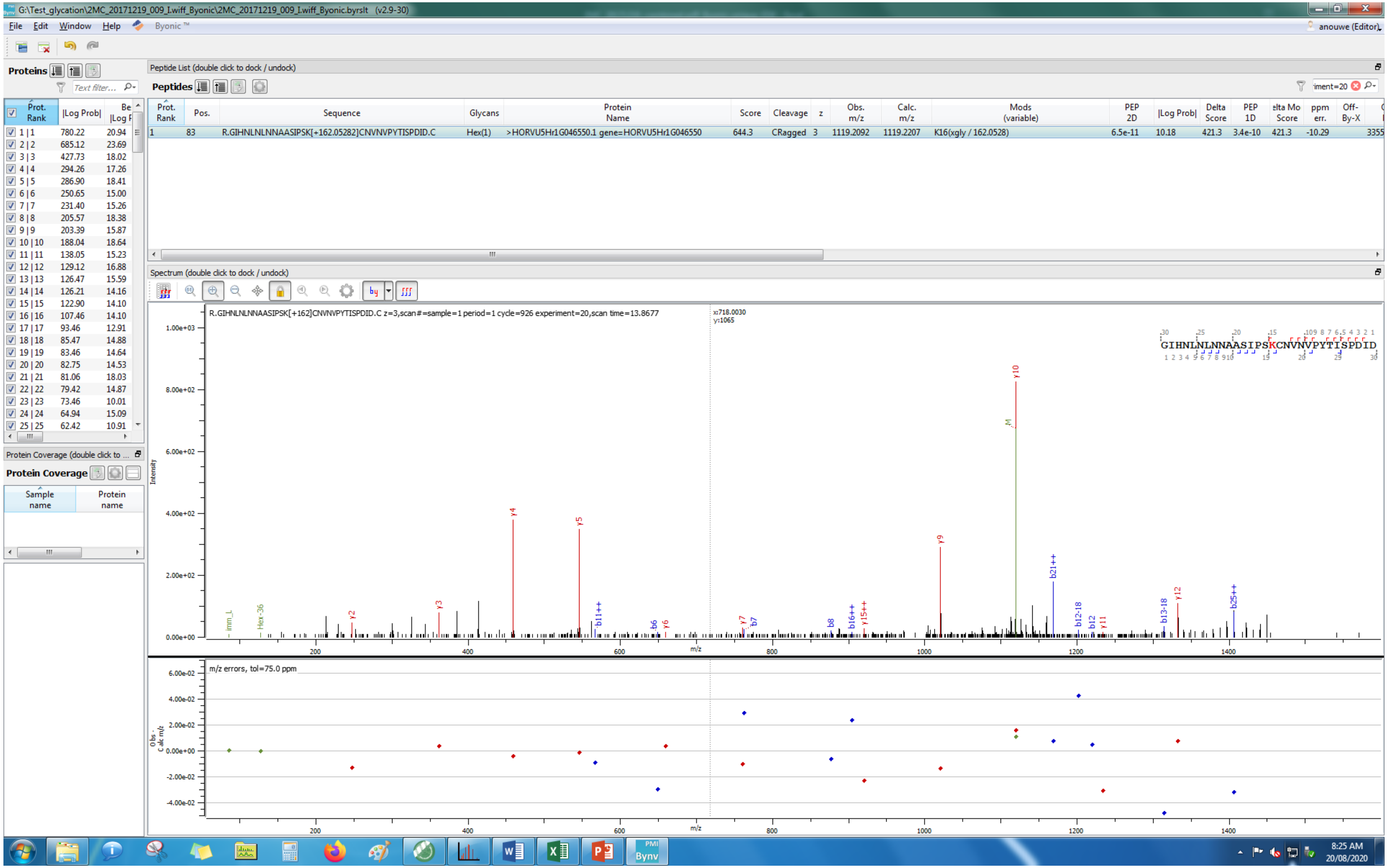
G:\Test_glycation\2MC_20171219_009_I.wiff_Byonic\2MC_20171219_009_I.wiff_Byonic.byrslt (v2.9-30)
File Edit Window Help Byonic ™
anouwe (Editor)
Proteins
Peptide List (double click to dock / undock)
Peptides
Prot. Rank | Pos. | Sequence | Glycans | Protein Name | Score | Cleavage | z | Obs. m/z | Calc. m/z | Mods (variable) | PEP 2D | |Log Prob| | Delta Score | PEP 1D | Delta Mod Score | ppm err. | Off-By-X
1 | 83 | R.GIHNLNLNNAASIPSK[+162.05282]CNVNVPYTISPDID.C | Hex(1) | >HORVU5Hr1G046550.1 gene=HORVU5Hr1G046550 | 644.3 | CRagged | 3 | 1119.2092 | 1119.2207 | K16(xgly / 162.0528) | 6.5e-11 | 10.18 | 421.3 | 3.4e-10 | 421.3 | -10.29 | | 3355
Prot. Rank | |Log Prob| | Best |Log Prob|
1 | 1 780.22 20.94
2 | 2 685.12 23.69
3 | 3 427.73 18.02
4 | 4 294.26 17.26
5 | 5 286.90 18.41
6 | 6 250.65 15.00
7 | 7 231.40 15.26
8 | 8 205.57 18.38
9 | 9 203.39 15.87
10 | 10 188.04 18.64
11 | 11 138.05 15.23
12 | 12 129.12 16.88
13 | 13 126.47 15.59
14 | 14 126.21 14.16
15 | 15 122.90 14.10
16 | 16 107.46 14.10
17 | 17 93.46 12.91
18 | 18 85.47 14.88
19 | 19 83.46 14.64
20 | 20 82.75 14.53
21 | 21 81.06 18.03
22 | 22 79.42 14.87
23 | 23 73.46 10.01
24 | 24 64.94 15.09
25 | 25 62.42 10.91
Protein Coverage
Sample name | Protein name
Spectrum (double click to dock / undock)
R.GIHNLNLNNAASIPSK[+162]CNVNVPYTISPDID.C z=3,scan#=sample=1 period=1 cycle=926 experiment=20,scan time=13.8677
x:718.0030
y:1065
GIHNLNLNNAASIPSKCNVNVPYTISPDID
Intensity
m/z
m/z errors, tol=75.0 ppm
Obs - Calc m/z
y10 y9 y5 y4 y3 y2 b21++ b25++ b13-18 b12-18 y12 b12 y11 b16++ y15++ b8 y7 b7 b6 y6 b11++ Hex-36 imm_L M
8:25 AM
20/08/2020

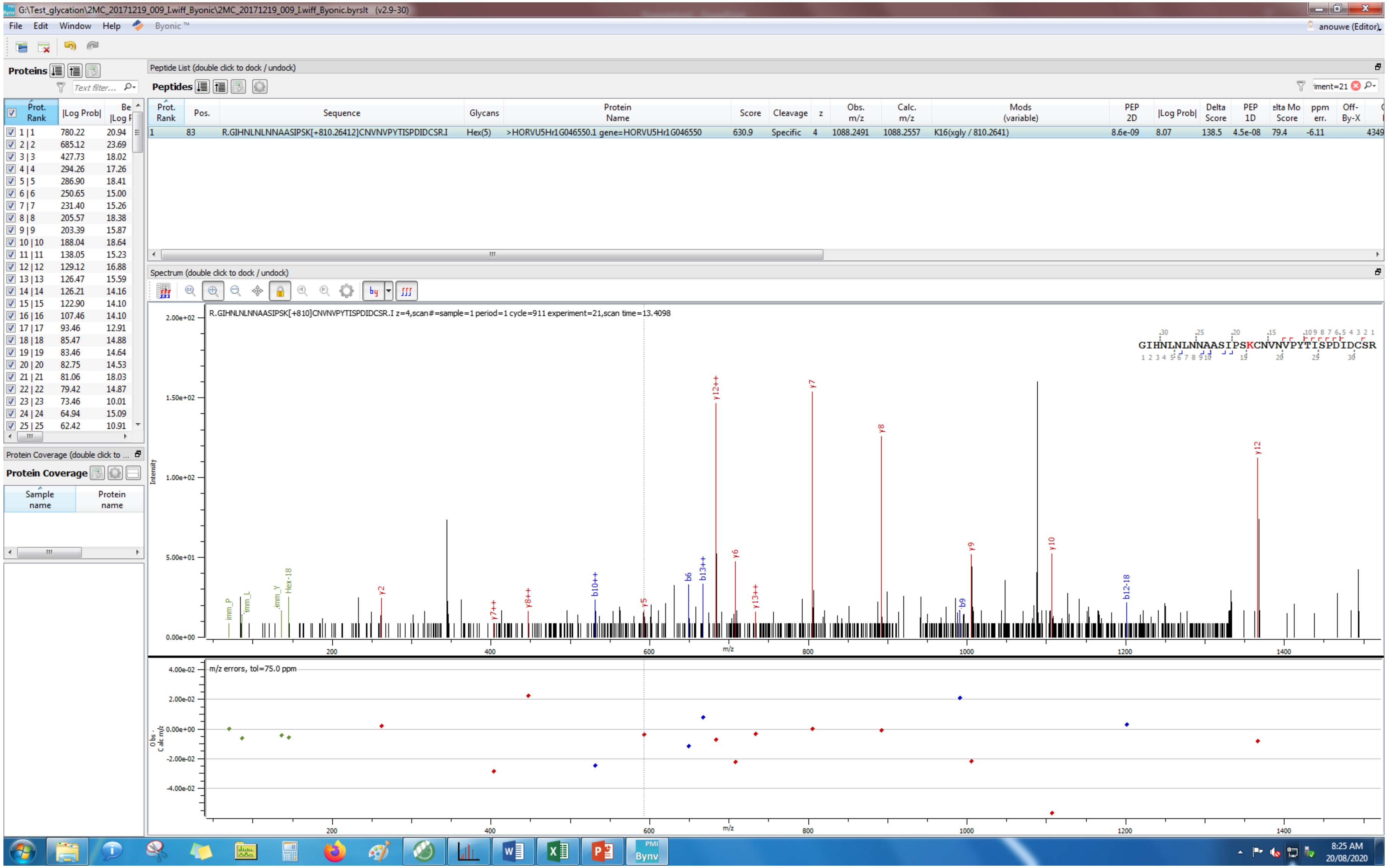
G:\Test_glycation\2MC_20171219_009_I.wiff_Byonic\2MC_20171219_009_I.wiff_Byonic.byrslt (v2.9-30)
File Edit Window Help Byonic™
anouwe (Editor)
Proteins
Peptide List (double click to dock / undock)
Peptides
Prot. Rank | Pos. | Sequence | Glycans | Protein Name | Score | Cleavage | z | Obs. m/z | Calc. m/z | Mods (variable) | PEP 2D | |Log Prob| | Delta Score | PEP 1D | Delta Mod Score | ppm err. | Off-By-X
1 | 83 | R.GIHNLNLNNAASIPSK[+810.26412]CNVNVPYTISPDIDCSR.I | Hex(5) | >HORVU5Hr1G046550.1 gene=HORVU5Hr1G046550 | 630.9 | Specific | 4 | 1088.2491 | 1088.2557 | K16(xgly / 810.2641) | 8.6e-09 | 8.07 | 138.5 | 4.5e-08 | 79.4 | -6.11
Spectrum (double click to dock / undock)
R.GIHNLNLNNAASIPSK[+810]CNVNVPYTISPDIDCSR.I z=4,scan#=sample=1 period=1 cycle=911 experiment=21,scan time=13.4098
GIHNLNLNNAASIPSKCNVNVPYTISPDIDCSR
Intensity
m/z
m/z errors, tol=75.0 ppm
Protein Coverage
Sample name
Protein name

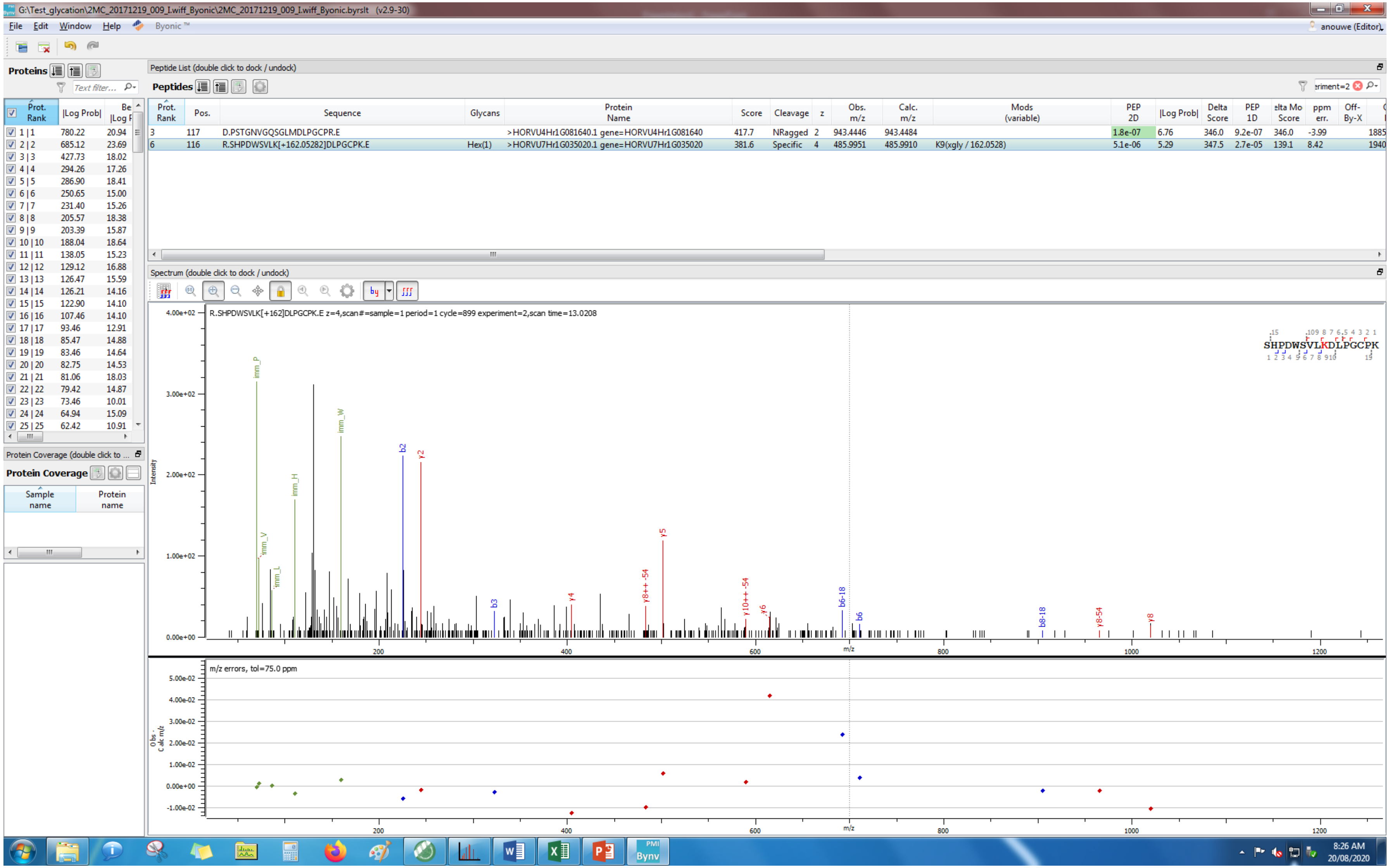
G:\Test_glycation\2MC_20171219_009_I.wiff_Byonic\2MC_20171219_009_I.wiff_Byonic.byrslt (v2.9-30)
File Edit Window Help Byonic™
anouwe (Editor)
Proteins
Prot. Rank |Log Prob| Be |Log P
1 | 1 780.22 20.94
2 | 2 685.12 23.69
3 | 3 427.73 18.02
4 | 4 294.26 17.26
5 | 5 286.90 18.41
6 | 6 250.65 15.00
7 | 7 231.40 15.26
8 | 8 205.57 18.38
9 | 9 203.39 15.87
10 | 10 188.04 18.64
11 | 11 138.05 15.23
12 | 12 129.12 16.88
13 | 13 126.47 15.59
14 | 14 126.21 14.16
15 | 15 122.90 14.10
16 | 16 107.46 14.10
17 | 17 93.46 12.91
18 | 18 85.47 14.88
19 | 19 83.46 14.64
20 | 20 82.75 14.53
21 | 21 81.06 18.03
22 | 22 79.42 14.87
23 | 23 73.46 10.01
24 | 24 64.94 15.09
25 | 25 62.42 10.91
Protein Coverage
Sample name
Protein name
Peptide List (double click to dock / undock)
Peptides
Prot. Rank | Pos. | Sequence | Glycans | Protein Name | Score | Cleavage | z | Obs. m/z | Calc. m/z | Mods (variable) | PEP 2D | |Log Prob| | Delta Score | PEP 1D | elta Mo Score | ppm err. | Off-By-X
3 | 117 | D.PSTGNVGQSGLMDLPGCPR.E | | >HORVU4Hr1G081640.1 gene=HORVU4Hr1G081640 | 417.7 | NRagged | 2 | 943.4446 | 943.4484 | | 1.8e-07 | 6.76 | 346.0 | 9.2e-07 | 346.0 | -3.99 | | 1885
6 | 116 | R.SHPDWSVLK[+162.05282]DLPGCPK.E | Hex(1) | >HORVU7Hr1G035020.1 gene=HORVU7Hr1G035020 | 381.6 | Specific | 4 | 485.9951 | 485.9910 | K9(xgly / 162.0528) | 5.1e-06 | 5.29 | 347.5 | 2.7e-05 | 139.1 | 8.42 | | 1940
Spectrum (double click to dock / undock)
R.SHPDWSVLK[+162]DLPGCPK.E z=4,scan#=sample=1 period=1 cycle=899 experiment=2,scan time=13.0208
SHPDWSVLKDLPGCPK
Intensity
4.00e+02
3.00e+02
2.00e+02
1.00e+02
0.00e+00
imm_P
imm_W
imm_H
imm_V
imm_L
b2
y2
b3
y4
y5
y8++ -54
y10++ -54
y6
b6-18
b6
b8-18
y8-54
y8
200
400
600
800
1000
1200
m/z
m/z errors, tol=75.0 ppm
Obs - Calc m/z
5.00e-02
4.00e-02
3.00e-02
2.00e-02
1.00e-02
0.00e+00
-1.00e-02
8:26 AM
20/08/2020

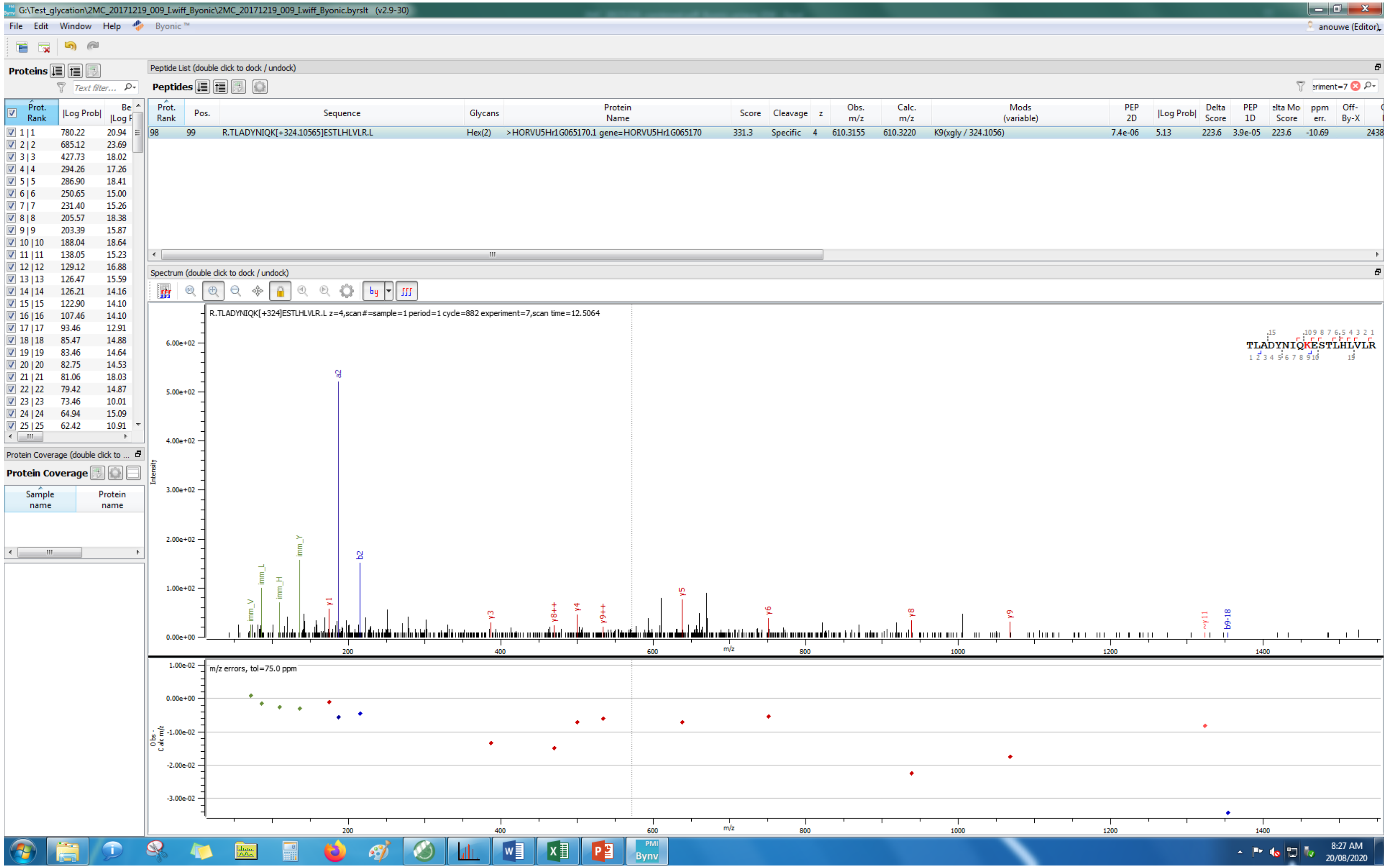
G:\Test_glycation\2MC_20171219_009_I.wiff_Byonic\2MC_20171219_009_I.wiff_Byonic.byrslt (v2.9-30)
File Edit Window Help Byonic™
Proteins
Peptide List (double click to dock / undock)
Peptides
Prot. Rank | Pos. | Sequence | Glycans | Protein Name | Score | Cleavage | z | Obs. m/z | Calc. m/z | Mods (variable) | PEP 2D | |Log Prob| | Delta Score | PEP 1D | ppm err.
98 | 99 | R.TLADYNIQK[+324.10565]ESTLHLVLR.L | Hex(2) | >HORVU5Hr1G065170.1 gene=HORVU5Hr1G065170 | 331.3 | Specific | 4 | 610.3155 | 610.3220 | K9(xgly / 324.1056) | 7.4e-06 | 5.13 | 223.6 | 3.9e-05 | -10.69
Spectrum (double click to dock / undock)
R.TLADYNIQK[+324]ESTLHLVLR.L z=4,scan#=sample=1 period=1 cycle=882 experiment=7,scan time=12.5064
TLADYNIQKESTLHLVLR
Intensity
m/z
m/z errors, tol=75.0 ppm
Obs - Calc m/z
Protein Coverage
Sample name
Protein name
8:27 AM
20/08/2020

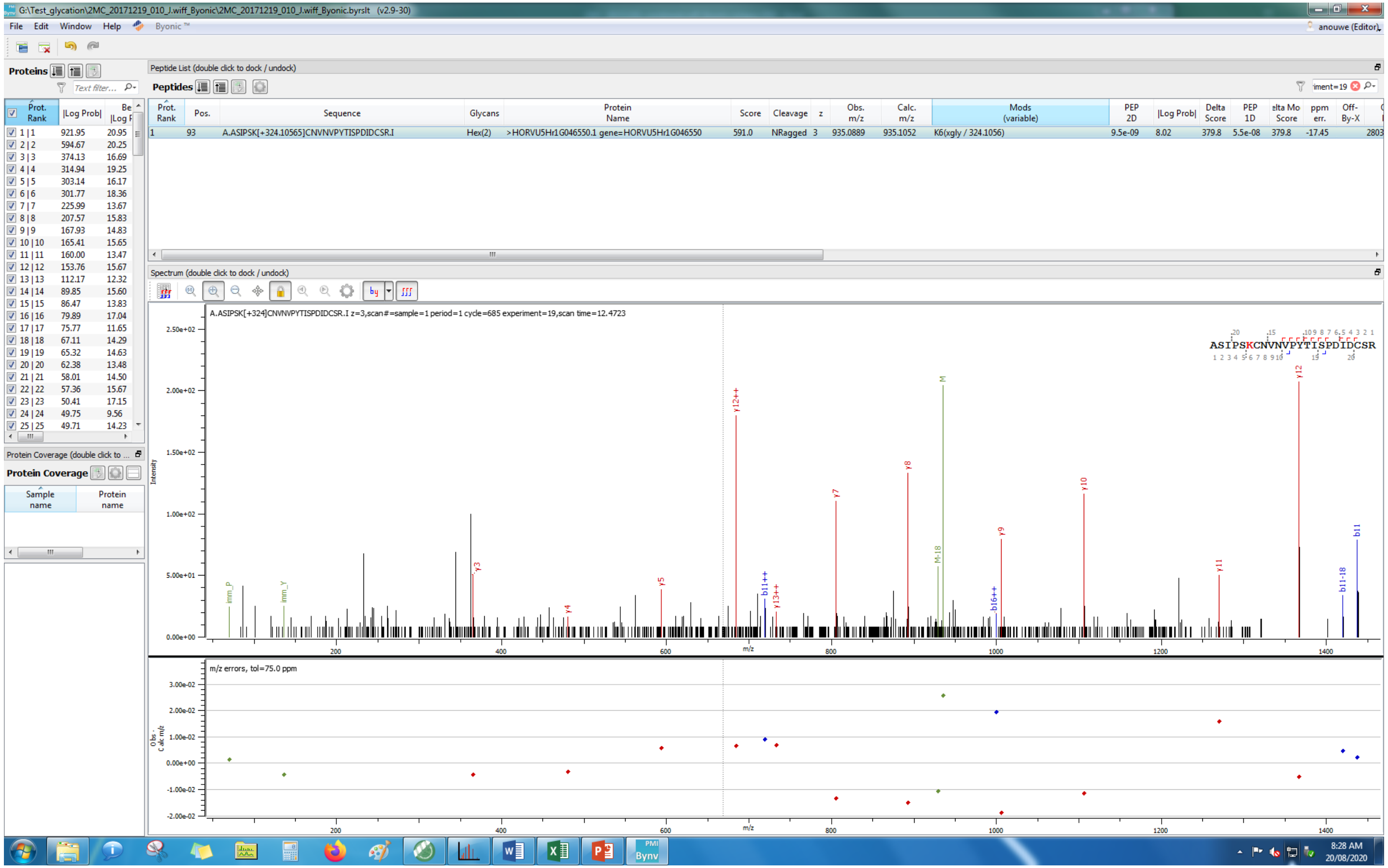
G:\Test_glycation\2MC_20171219_010_J.wiff_Byonic\2MC_20171219_010_J.wiff_Byonic.byrslt (v2.9-30)
File Edit Window Help Byonic™
anouwe (Editor)
Proteins
Peptide List (double click to dock / undock)
Peptides
Prot. Rank | Pos. | Sequence | Glycans | Protein Name | Score | Cleavage | z | Obs. m/z | Calc. m/z | Mods (variable) | PEP 2D | |Log Prob| | Delta Score | PEP 1D | elta Mo Score | ppm err. | Off-By-X
1 | 93 | A.ASIPSK[+324.10565]CNVNVPYTISPDIDCSR.I | Hex(2) | >HORVU5Hr1G046550.1 gene=HORVU5Hr1G046550 | 591.0 | NRagged | 3 | 935.0889 | 935.1052 | K6(xgly / 324.1056) | 9.5e-09 | 8.02 | 379.8 | 5.5e-08 | 379.8 | -17.45 | 2803
Spectrum (double click to dock / undock)
A.ASIPSK[+324]CNVNVPYTISPDIDCSR.I z=3,scan#=sample=1 period=1 cycle=685 experiment=19,scan time=12.4723
ASIPSKCNVNVPYTISPDIDCSR
Intensity
m/z
m/z errors, tol=75.0 ppm
Obs - Calc m/z
Protein Coverage (double click to ...
Protein Coverage
Sample name
Protein name
8:28 AM
20/08/2020

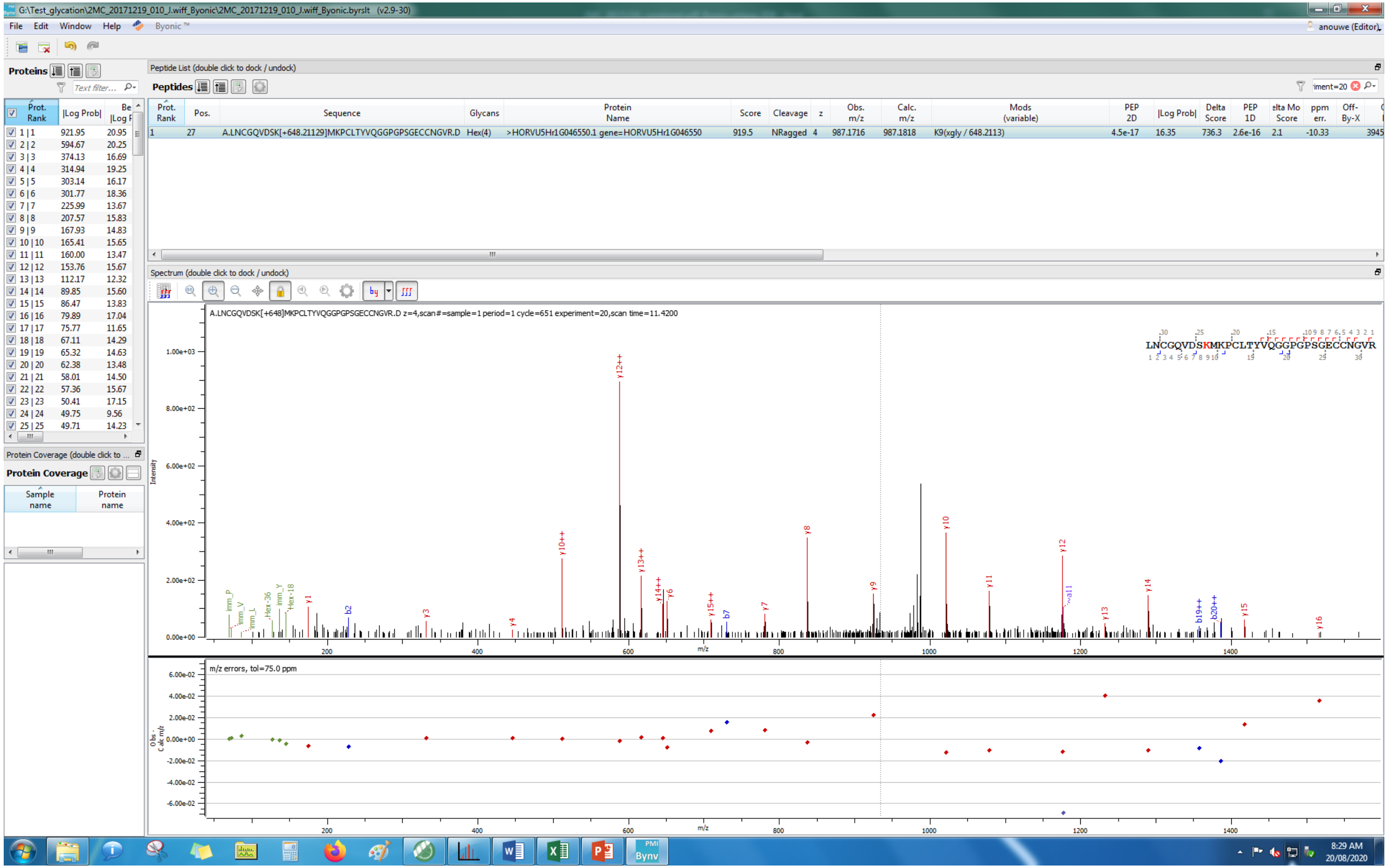
G:\Test_glycation\2MC_20171219_010_I.wiff_Byonic\2MC_20171219_010_I.wiff_Byonic.byrslt (v2.9-30)
File Edit Window Help Byonic ™
anouwe (Editor)
Proteins
Peptide List (double click to dock / undock)
Peptides
Prot. Rank
Pos.
Sequence
Glycans
Protein Name
Score
Cleavage
z
Obs. m/z
Calc. m/z
Mods (variable)
PEP 2D
|Log Prob|
Delta Score
PEP 1D
ppm err.
Off-By-X
A.LNCGQVDSK[+648.21129]MKPCLTYVQGGPGPSGECCNGVR.D
Hex(4)
>HORVU5Hr1G046550.1 gene=HORVU5Hr1G046550
919.5
NRagged
987.1716
987.1818
K9(xgly / 648.2113)
4.5e-17
16.35
736.3
2.6e-16
-10.33
Spectrum (double click to dock / undock)
A.LNCGQVDSK[+648]MKPCLTYVQGGPGPSGECCNGVR.D z=4,scan#=sample=1 period=1 cycle=651 experiment=20,scan time=11.4200
LNCGQVDSKMKPCLTYVQGGPGPSGECCNGVR
Intensity
m/z
m/z errors, tol=75.0 ppm
Protein Coverage (double click to ...
Protein Coverage
Sample name
Protein name
8:29 AM
20/08/2020

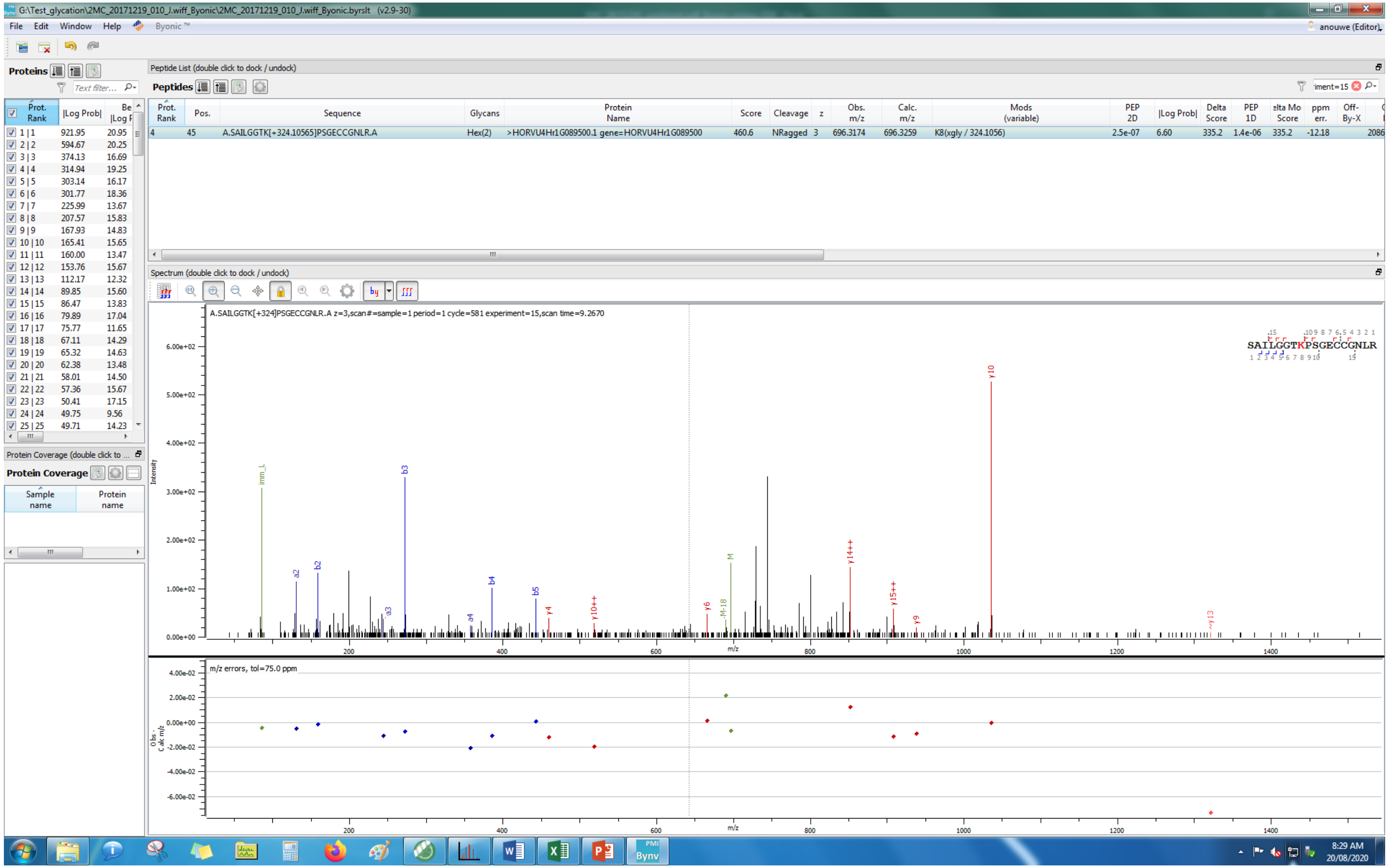
G:\Test_glycation\2MC_20171219_010_J.wiff_Byonic\2MC_20171219_010_J.wiff_Byonic.byrslt (v2.9-30)
File Edit Window Help Byonic™
Proteins
Peptide List (double click to dock / undock)
Peptides
Prot. Rank | Pos. | Sequence | Glycans | Protein Name | Score | Cleavage | z | Obs. m/z | Calc. m/z | Mods (variable) | PEP 2D | |Log Prob| | Delta Score | PEP 1D | Delta Mod Score | ppm err. | Off-By-X
4 | 45 | A.SAILGGTK[+324.10565]PSGECCGNLR.A | Hex(2) | >HORVU4Hr1G089500.1 gene=HORVU4Hr1G089500 | 460.6 | NRagged | 3 | 696.3174 | 696.3259 | K8(xgly / 324.1056) | 2.5e-07 | 6.60 | 335.2 | 1.4e-06 | 335.2 | -12.18
Spectrum (double click to dock / undock)
A.SAILGGTK[+324]PSGECCGNLR.A z=3,scan#=sample=1 period=1 cycle=581 experiment=15,scan time=9.2670
SAILGGTKPSGECCGNLR
Intensity
m/z
m/z errors, tol=75.0 ppm
Protein Coverage
Sample name
Protein name

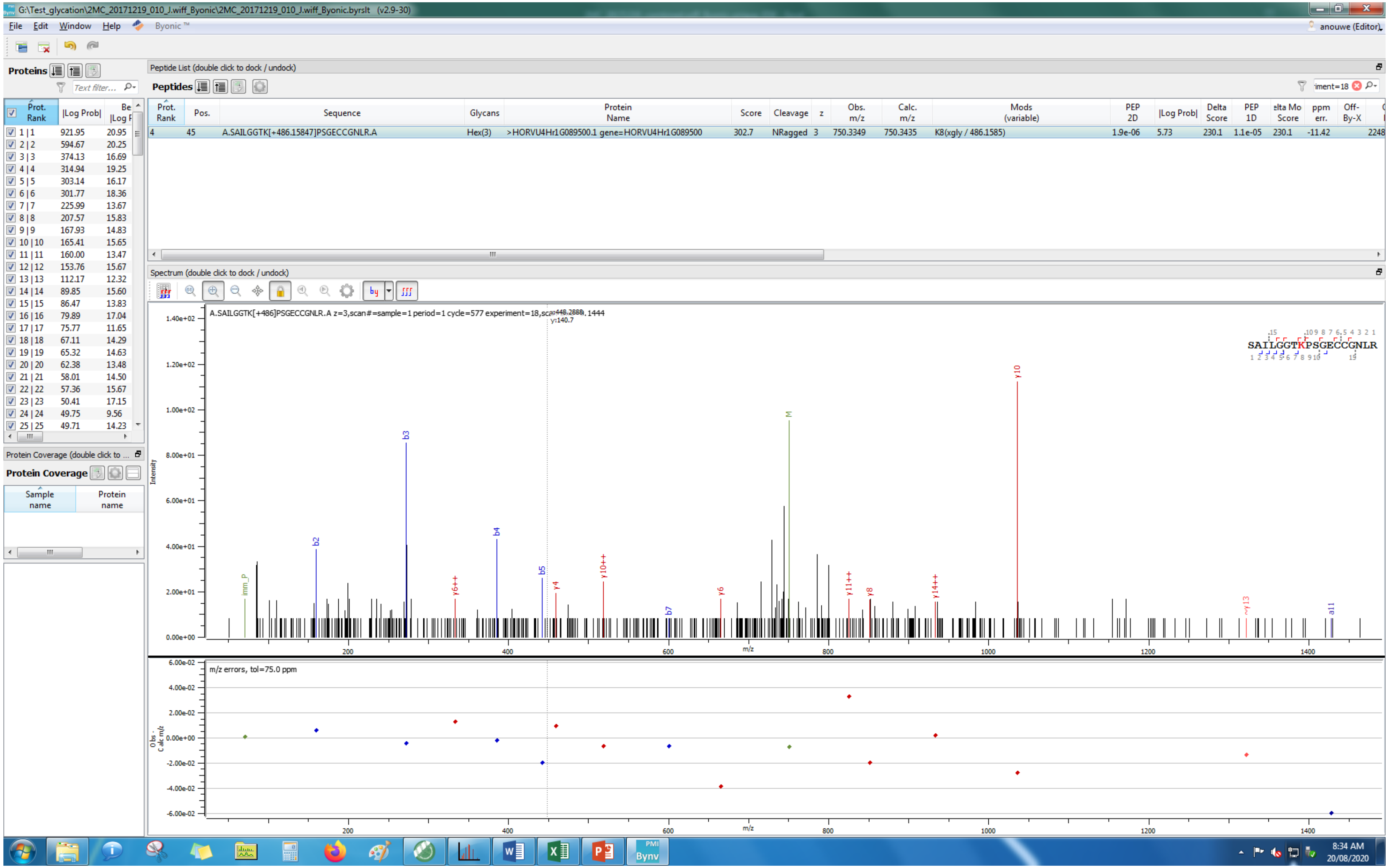
G:\Test_glycation\2MC_20171219_010_J.wiff_Byonic\2MC_20171219_010_J.wiff_Byonic.byrslt (v2.9-30)
File Edit Window Help Byonic ™
anouwe (Editor)
Proteins
Text filter...
Prot. Rank | |Log Prob| | Be |Log P
1 | 1 921.95 20.95
2 | 2 594.67 20.25
3 | 3 374.13 16.69
4 | 4 314.94 19.25
5 | 5 303.14 16.17
6 | 6 301.77 18.36
7 | 7 225.99 13.67
8 | 8 207.57 15.83
9 | 9 167.93 14.83
10 | 10 165.41 15.65
11 | 11 160.00 13.47
12 | 12 153.76 15.67
13 | 13 112.17 12.32
14 | 14 89.85 15.60
15 | 15 86.47 13.83
16 | 16 79.89 17.04
17 | 17 75.77 11.65
18 | 18 67.11 14.29
19 | 19 65.32 14.63
20 | 20 62.38 13.48
21 | 21 58.01 14.50
22 | 22 57.36 15.67
23 | 23 50.41 17.15
24 | 24 49.75 9.56
25 | 25 49.71 14.23
Protein Coverage (double click to ...
Protein Coverage
Sample name
Protein name
Peptide List (double click to dock / undock)
Peptides
iment=18
Prot. Rank | Pos. | Sequence | Glycans | Protein Name | Score | Cleavage | z | Obs. m/z | Calc. m/z | Mods (variable) | PEP 2D | |Log Prob| | Delta Score | PEP 1D | elta Mo Score | ppm err. | Off-By-X
4 45 A.SAILGGTK[+486.15847]PSGECCGNLR.A Hex(3) >HORVU4Hr1G089500.1 gene=HORVU4Hr1G089500 302.7 NRagged 3 750.3349 750.3435 K8(xgly / 486.1585) 1.9e-06 5.73 230.1 1.1e-05 230.1 -11.42 2248
Spectrum (double click to dock / undock)
A.SAILGGTK[+486]PSGECCGNLR.A z=3,scan#=sample=1 period=1 cycle=577 experiment=18,sc
SAILGGTKPSGECCGNLR
Intensity
m/z
imm_P b2 b3 b4 b5 y6++ y4 y10++ b7 y6 M y11++ y8 y14++ y10 ~y13 a11
m/z errors, tol=75.0 ppm
Obs - Calc m/z
8:34 AM
20/08/2020

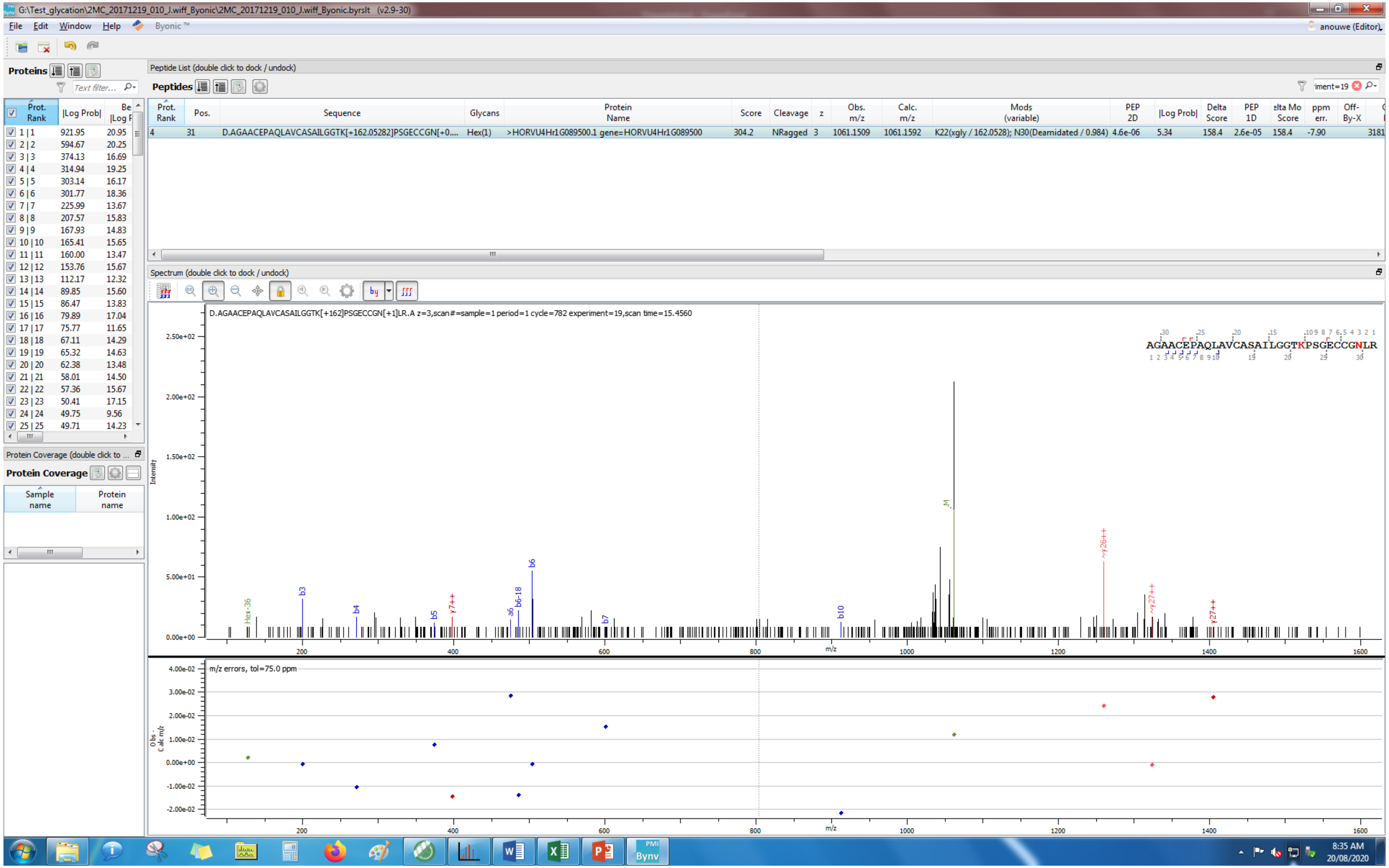
G:\Test_glycation\2MC_20171219_010_J.wiff_Byonic\2MC_20171219_010_J.wiff_Byonic.byrslt (v2.9-30)
File Edit Window Help Byonic™
anouwe (Editor)
Proteins
Peptide List (double click to dock / undock)
Peptides
Prot. Rank | Pos. | Sequence | Glycans | Protein Name | Score | Cleavage | z | Obs. m/z | Calc. m/z | Mods (variable) | PEP 2D | |Log Prob| | Delta Score | PEP 1D | Delta Mod Score | ppm err. | Off-By-X
4 | 31 | D.AGAACEPAQLAVCASAILGGTK[+162.05282]PSGECCGN[+0.... | Hex(1) | >HORVU4Hr1G089500.1 gene=HORVU4Hr1G089500 | 304.2 | NRagged | 3 | 1061.1509 | 1061.1592 | K22(xgly / 162.0528); N30(Deamidated / 0.984) | 4.6e-06 | 5.34 | 158.4 | 2.6e-05 | 158.4 | -7.90 | 3181
Prot. Rank | |Log Prob| | Best |Log Prob|
1 | 1 | 921.95 | 20.95
2 | 2 | 594.67 | 20.25
3 | 3 | 374.13 | 16.69
4 | 4 | 314.94 | 19.25
5 | 5 | 303.14 | 16.17
6 | 6 | 301.77 | 18.36
7 | 7 | 225.99 | 13.67
8 | 8 | 207.57 | 15.83
9 | 9 | 167.93 | 14.83
10 | 10 | 165.41 | 15.65
11 | 11 | 160.00 | 13.47
12 | 12 | 153.76 | 15.67
13 | 13 | 112.17 | 12.32
14 | 14 | 89.85 | 15.60
15 | 15 | 86.47 | 13.83
16 | 16 | 79.89 | 17.04
17 | 17 | 75.77 | 11.65
18 | 18 | 67.11 | 14.29
19 | 19 | 65.32 | 14.63
20 | 20 | 62.38 | 13.48
21 | 21 | 58.01 | 14.50
22 | 22 | 57.36 | 15.67
23 | 23 | 50.41 | 17.15
24 | 24 | 49.75 | 9.56
25 | 25 | 49.71 | 14.23
Protein Coverage (double click to ...
Protein Coverage
Sample name
Protein name
Spectrum (double click to dock / undock)
D.AGAACEPAQLAVCASAILGGTK[+162]PSGECCGN[+1]LR.A z=3,scan#=sample=1 period=1 cycle=782 experiment=19,scan time=15.4560
AGAACEPAQLAVCASAILGGTKPSGECCGNLR
Intensity
Hex-36
b3
b4
b5
y7++
a6
b6-18
b6
b7
b10
M
y26++
y27++
y27++
m/z
m/z errors, tol=75.0 ppm
Obs - Calc m/z
8:35 AM
20/08/2020

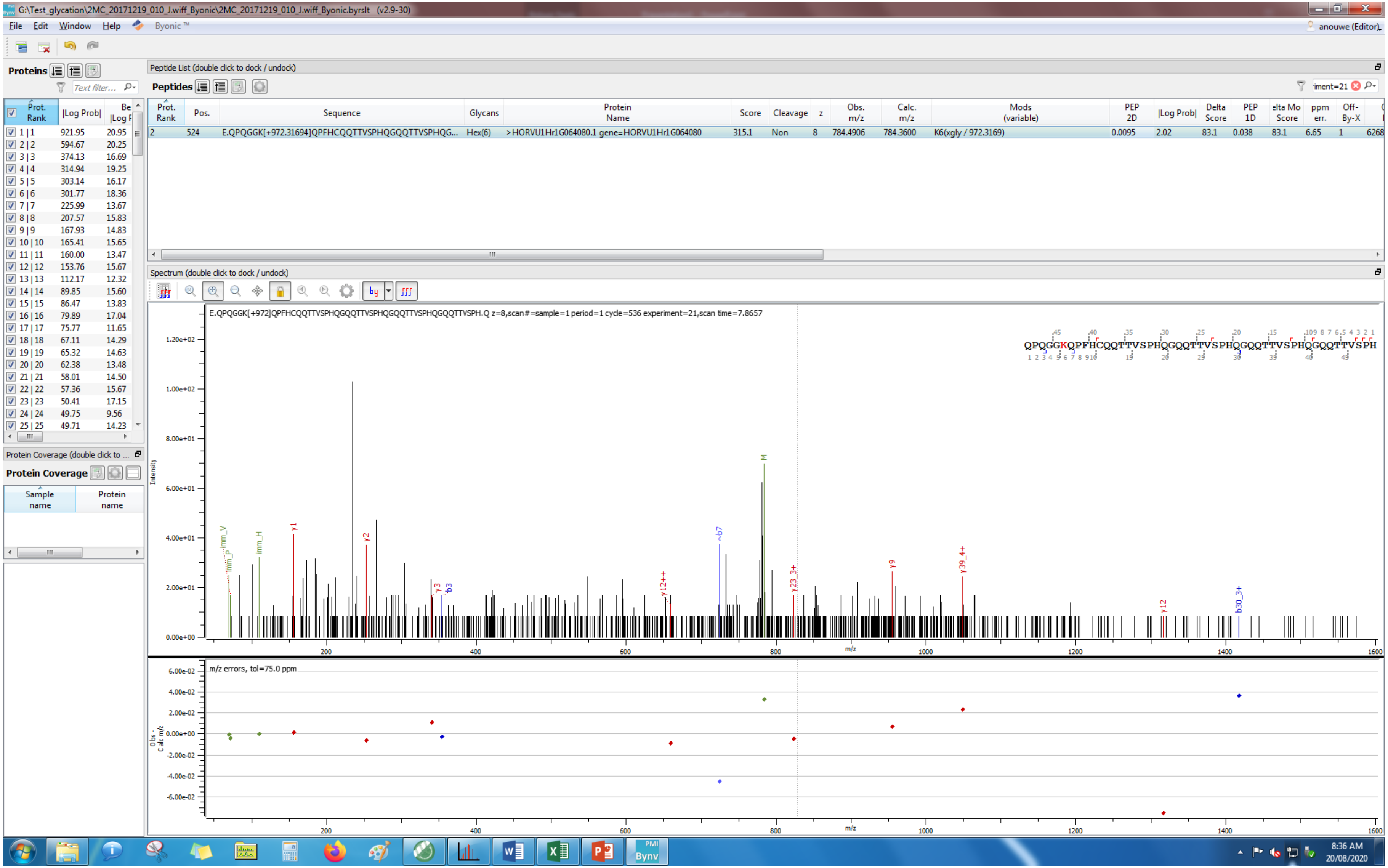
G:\Test_glycation\2MC_20171219_010_J.wiff_Byonic\2MC_20171219_010_J.wiff_Byonic.byrslt (v2.9-30)
File Edit Window Help Byonic ™
anouwe (Editor)
Proteins
Peptide List (double click to dock / undock)
Peptides
Prot. Rank | Pos. | Sequence | Glycans | Protein Name | Score | Cleavage | z | Obs. m/z | Calc. m/z | Mods (variable) | PEP 2D | |Log Prob| | Delta Score | PEP 1D | Delta Mod Score | ppm err. | Off-By-X
2 | 524 | E.QPQGGK[+972.31694]QPFHCQQTTVSPHQGQQTTVSPHQG... | Hex(6) | >HORVU1Hr1G064080.1 gene=HORVU1Hr1G064080 | 315.1 | Non | 8 | 784.4906 | 784.3600 | K6(xgly / 972.3169) | 0.0095 | 2.02 | 83.1 | 0.038 | 83.1 | 6.65 | 1
Spectrum (double click to dock / undock)
E.QPQGGK[+972]QPFHCQQTTVSPHQGQQTTVSPHQGQQTTVSPHQGQQTTVSPH.Q z=8,scan#=sample=1 period=1 cycle=536 experiment=21,scan time=7.8657
QPQGGKQPFHCQQTTVSPHQGQQTTVSPHQGQQTTVSPHQGQQTTVSPH
Intensity
m/z
m/z errors, tol=75.0 ppm
Obs - Calc m/z
Protein Coverage (double click to ...
Protein Coverage
Sample name
Protein name
8:36 AM
20/08/2020

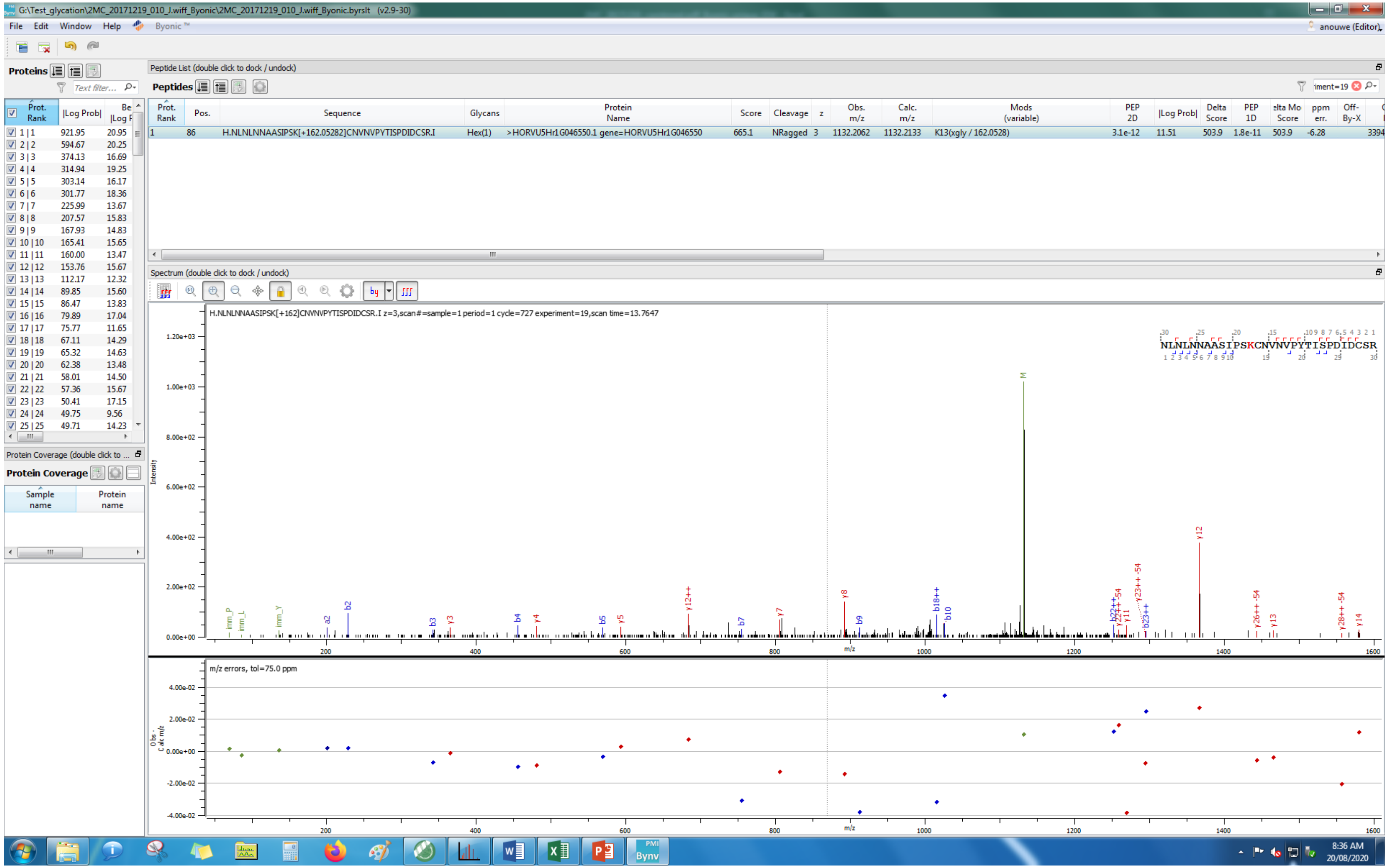
G:\Test_glycation\2MC_20171219_010_J.wiff_Byonic\2MC_20171219_010_J.wiff_Byonic.byrslt (v2.9-30)
File Edit Window Help Byonic
Proteins
Peptide List (double click to dock / undock)
Peptides
Prot. Rank
Pos.
Sequence
Glycans
Protein Name
Score
Cleavage
z
Obs. m/z
Calc. m/z
Mods (variable)
PEP 2D
|Log Prob|
Delta Score
PEP 1D
ppm err.
1 86 H.NLNLNNAASIPSK[+162.05282]CNVNVPYTISPDIDCSR.I Hex(1) >HORVU5Hr1G046550.1 gene=HORVU5Hr1G046550 665.1 NRagged 3 1132.2062 1132.2133 K13(xgly / 162.0528) 3.1e-12 11.51 503.9 1.8e-11 503.9 -6.28
Spectrum (double click to dock / undock)
H.NLNLNNAASIPSK[+162]CNVNVPYTISPDIDCSR.I z=3,scan#=sample=1 period=1 cycle=727 experiment=19,scan time=13.7647
NLNLNNAASIPSKCNVNVPYTISPDIDCSR
Intensity
m/z
m/z errors, tol=75.0 ppm
Protein Coverage (double click to ...
Protein Coverage
Sample name
Protein name
8:36 AM
20/08/2020

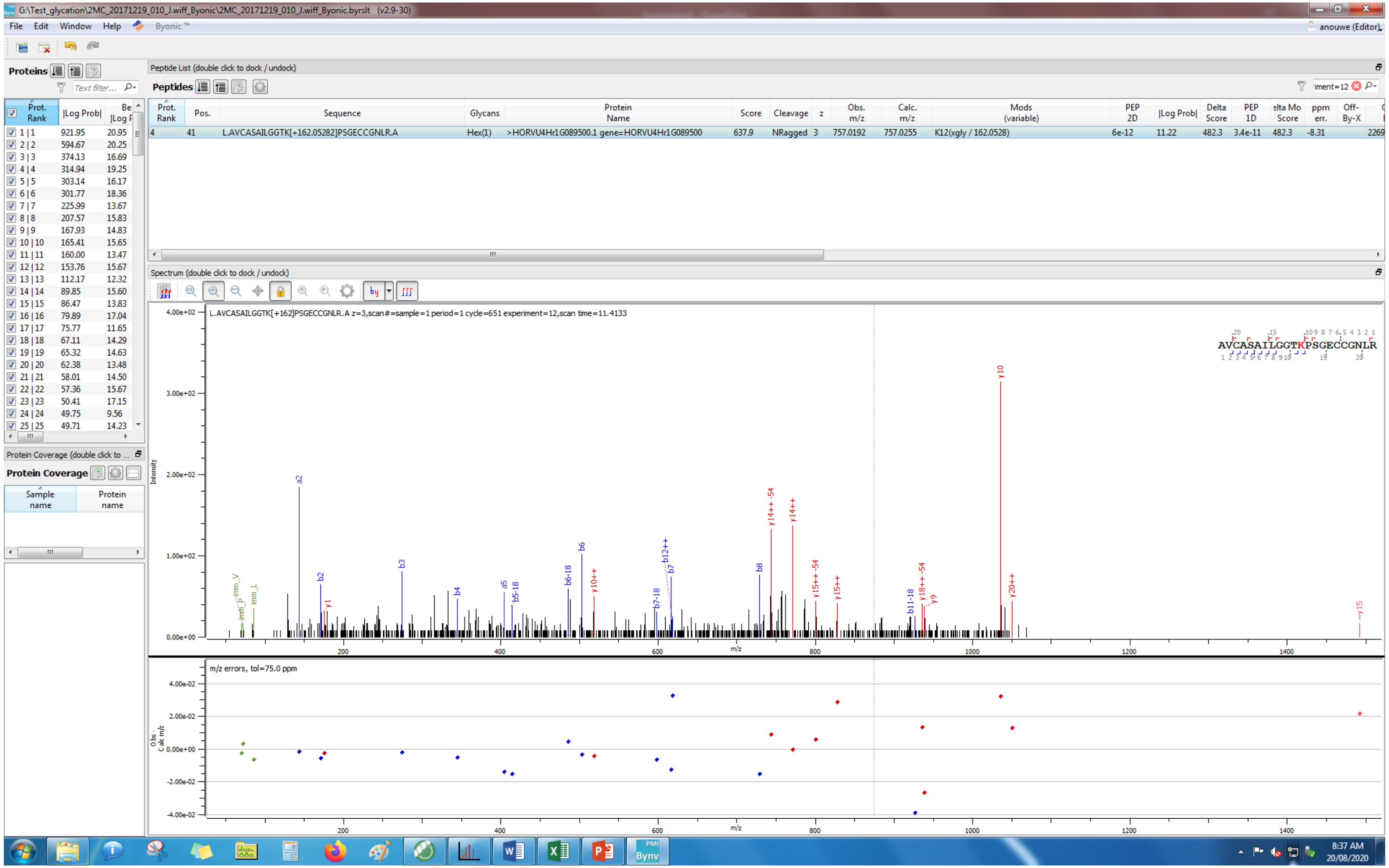
G:\Test_glycation\2MC_20171219_010_J.wiff_Byonic\2MC_20171219_010_J.wiff_Byonic.byrslt (v2.9-30)
File Edit Window Help Byonic ™
anouwe (Editor)
Proteins
Peptide List (double click to dock / undock)
Peptides
Prot. Rank | Pos. | Sequence | Glycans | Protein Name | Score | Cleavage | z | Obs. m/z | Calc. m/z | Mods (variable) | PEP 2D | |Log Prob| | Delta Score | PEP 1D | Delta Mod Score | ppm err. | Off-By-X
4 | 41 | L.AVCASAILGGTK[+162.05282]PSGECCGNLR.A | Hex(1) | >HORVU4Hr1G089500.1 gene=HORVU4Hr1G089500 | 637.9 | NRagged | 3 | 757.0192 | 757.0255 | K12(xgly / 162.0528) | 6e-12 | 11.22 | 482.3 | 3.4e-11 | 482.3 | -8.31 | 2269
Spectrum (double click to dock / undock)
L.AVCASAILGGTK[+162]PSGECCGNLR.A z=3,scan#=sample=1 period=1 cycle=651 experiment=12,scan time=11.4133
AVCASAILGGTKPSGECCGNLR
Intensity
m/z
m/z errors, tol=75.0 ppm
Protein Coverage
Sample name
Protein name
8:37 AM
20/08/2020

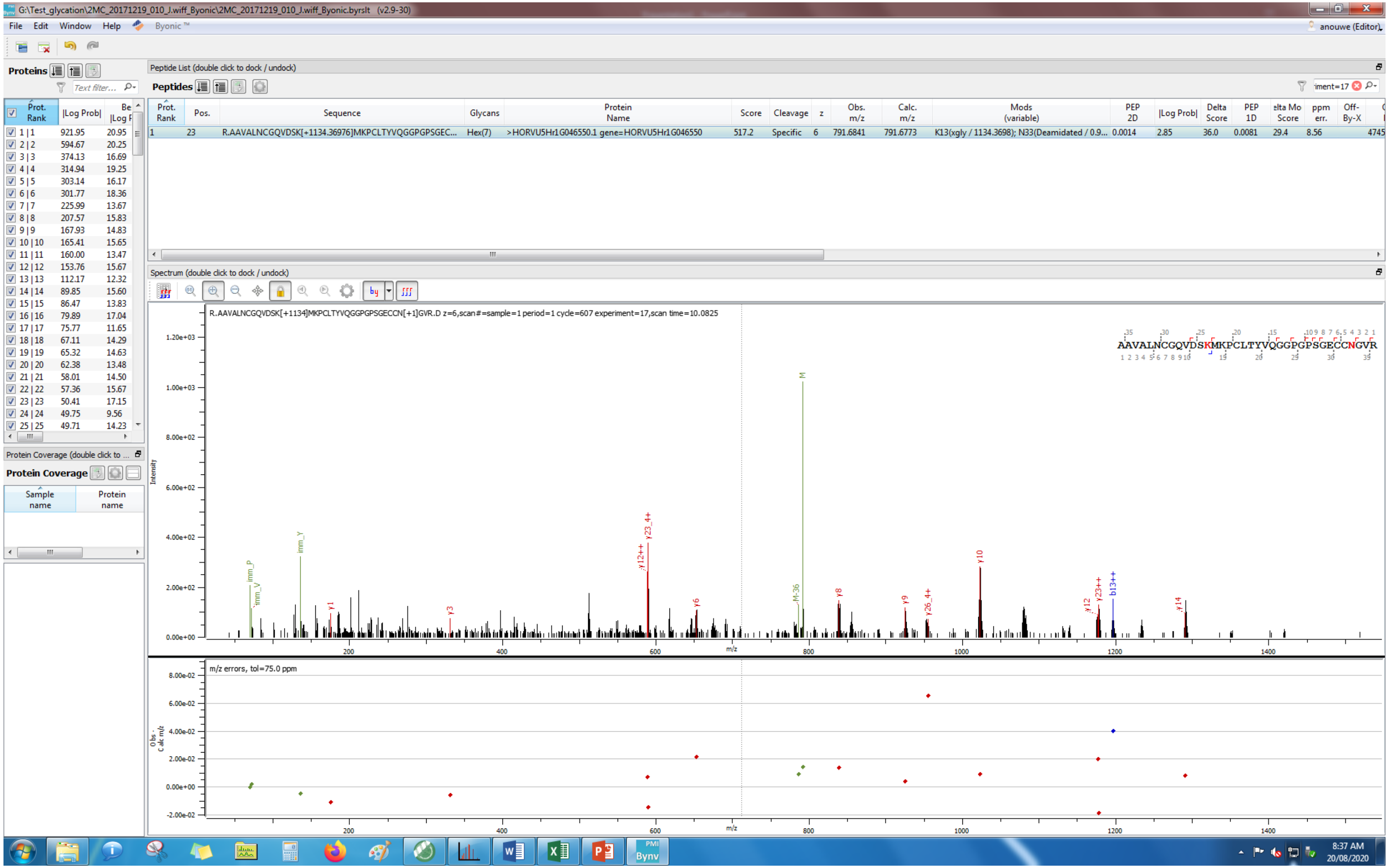
G:\Test_glycation\2MC_20171219_010_J.wiff_Byonic\2MC_20171219_010_J.wiff_Byonic.byrslt (v2.9-30)
File Edit Window Help Byonic ™
anouwe (Editor)
Proteins
Peptide List (double click to dock / undock)
Peptides
R.AAVALNCGQVDSK[+1134.36976]MKPCLTYVQGGPGPSGEC...
Hex(7)
>HORVU5Hr1G046550.1 gene=HORVU5Hr1G046550
K13(xgly / 1134.3698); N33(Deamidated / 0.9...
Spectrum (double click to dock / undock)
R.AAVALNCGQVDSK[+1134]MKPCLTYVQGGPGPSGECCN[+1]GVR.D z=6,scan#=sample=1 period=1 cycle=607 experiment=17,scan time=10.0825
AAVALNCGQVDSKMKPCLTYVQGGPGPSGECCNGVR
m/z errors, tol=75.0 ppm
Protein Coverage
Sample name
Protein name
8:37 AM
20/08/2020

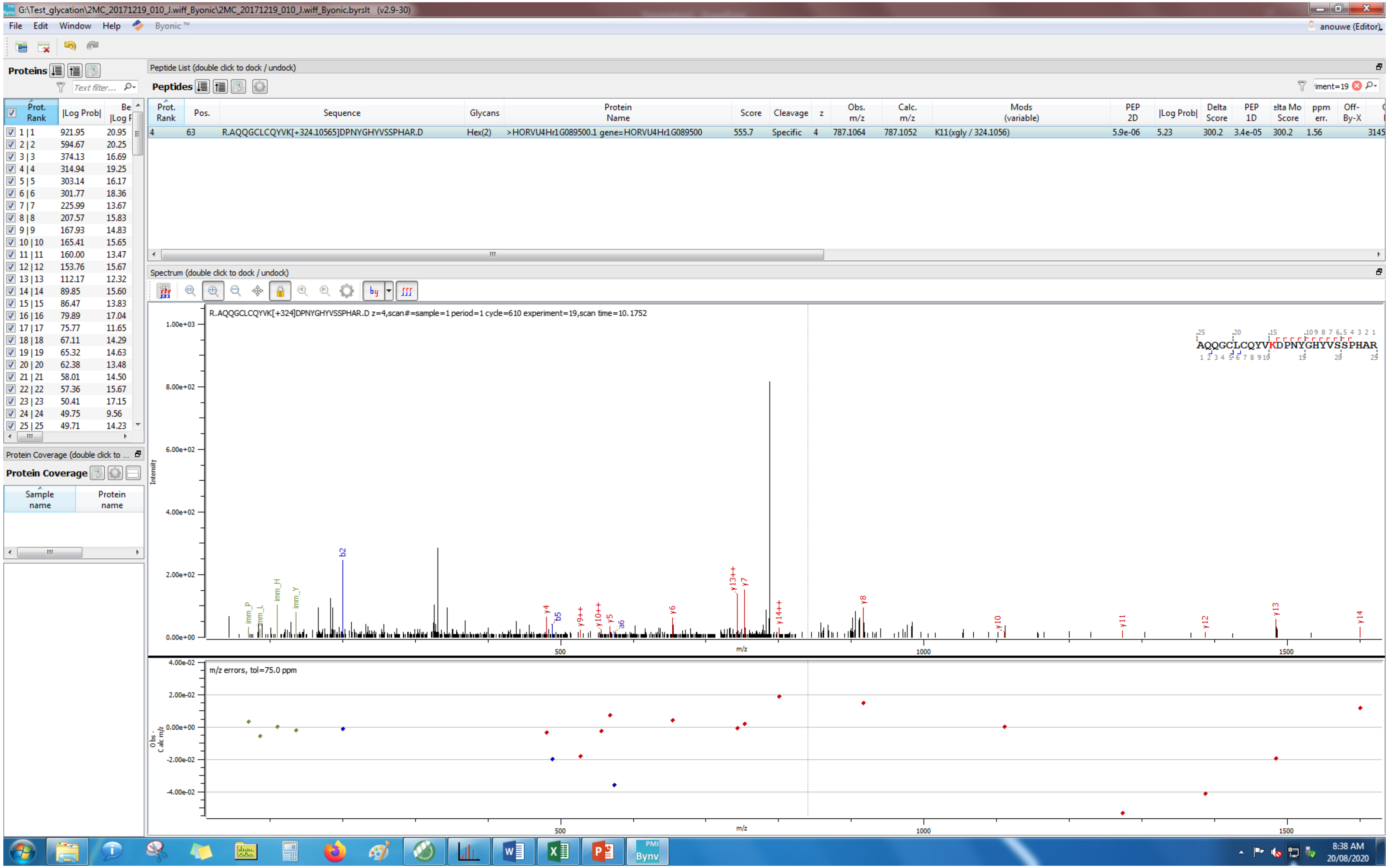

G:\Test_glycation\2MC_20171219_010_J.wiff_Byonic\2MC_20171219_010_J.wiff_Byonic.byrslt (v2.9-30)
File Edit Window Help Byonic™
anouwe (Editor)
Proteins
Peptide List (double click to dock / undock)
Peptides
Prot. Rank | Pos. | Sequence | Glycans | Protein Name | Score | Cleavage | z | Obs. m/z | Calc. m/z | Mods (variable) | PEP 2D | |Log Prob| | Delta Score | PEP 1D | Delta Mod Score | ppm err. | Off-By-X
4 | 63 | R.AQQGCLCQYVK[+324.10565]DPNYGHYVSSPHAR.D | Hex(2) | >HORVU4Hr1G089500.1 gene=HORVU4Hr1G089500 | 555.7 | Specific | 4 | 787.1064 | 787.1052 | K11(xgly / 324.1056) | 5.9e-06 | 5.23 | 300.2 | 3.4e-05 | 300.2 | 1.56
Spectrum (double click to dock / undock)
R.AQQGCLCQYVK[+324]DPNYGHYVSSPHAR.D z=4,scan#=sample=1 period=1 cycle=610 experiment=19,scan time=10.1752
AQQGCLCQYVKDPNYGHYVSSPHAR
Intensity
m/z
m/z errors, tol=75.0 ppm
Protein Coverage (double click to ...
Protein Coverage
Sample name
Protein name
8:38 AM
20/08/2020

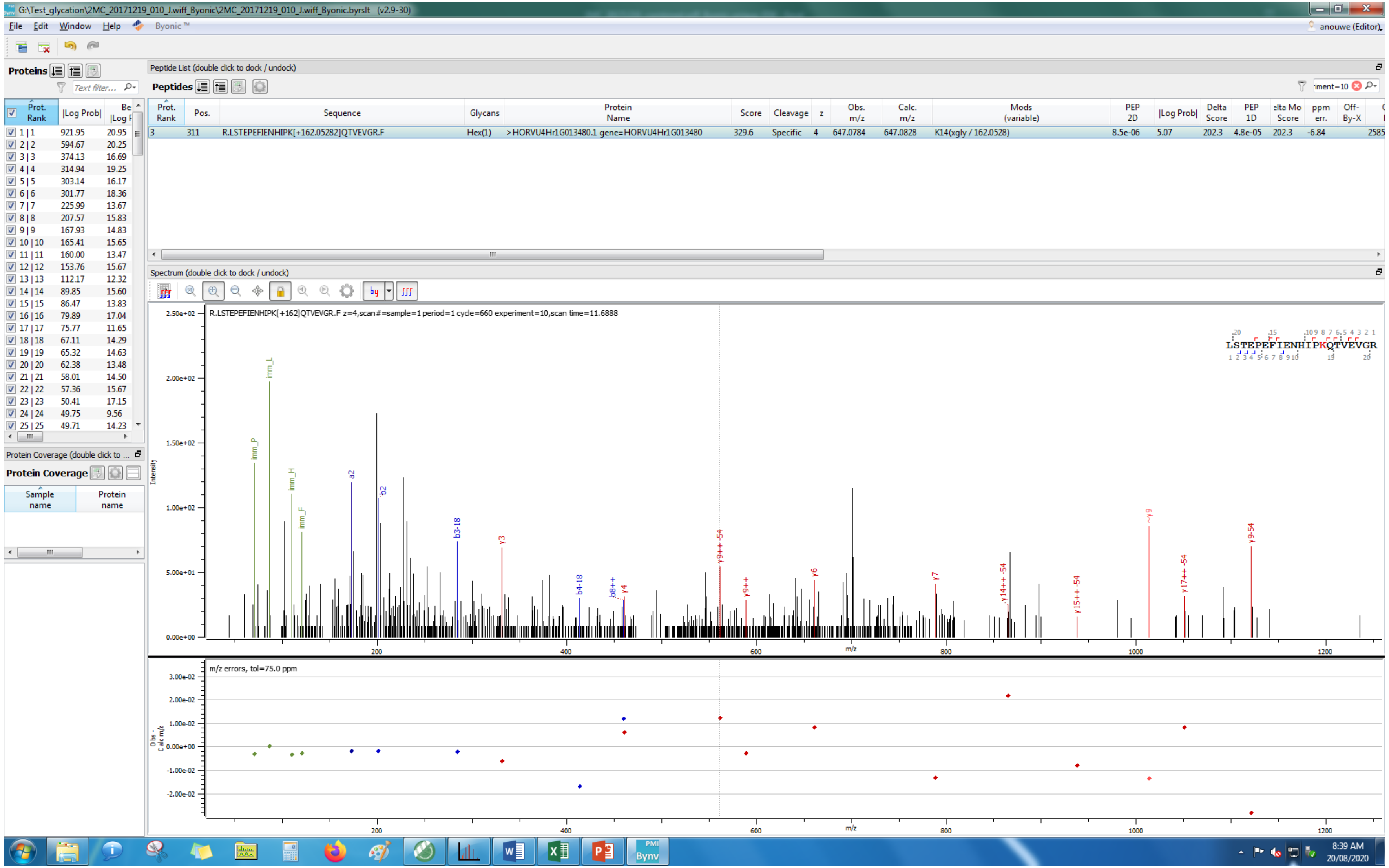

G:\Test_glycation\2MC_20171219_010_J.wiff_Byonic\2MC_20171219_010_J.wiff_Byonic.byrslt (v2.9-30)
File Edit Window Help Byonic
anouwe (Editor)
Proteins
Peptide List (double click to dock / undock)
Peptides
Prot. Rank | Pos. | Sequence | Glycans | Protein Name | Score | Cleavage | z | Obs. m/z | Calc. m/z | Mods (variable) | PEP 2D | |Log Prob| | Delta Score | PEP 1D | Delta Mod Score | ppm err. | Off-By-X
3 | 311 | R.LSTEPEFIENHIPK[+162.05282]QTVEVGR.F | Hex(1) | >HORVU4Hr1G013480.1 gene=HORVU4Hr1G013480 | 329.6 | Specific | 4 | 647.0784 | 647.0828 | K14(xgly / 162.0528) | 8.5e-06 | 5.07 | 202.3 | 4.8e-05 | 202.3 | -6.84
Spectrum (double click to dock / undock)
R.LSTEPEFIENHIPK[+162]QTVEVGR.F z=4,scan#=sample=1 period=1 cycle=660 experiment=10,scan time=11.6888
LSTEPEFIENHIPKQTVEVGR
Intensity
m/z
m/z errors, tol=75.0 ppm
Obs - Calc m/z
Protein Coverage
Sample name
Protein name
8:39 AM
20/08/2020

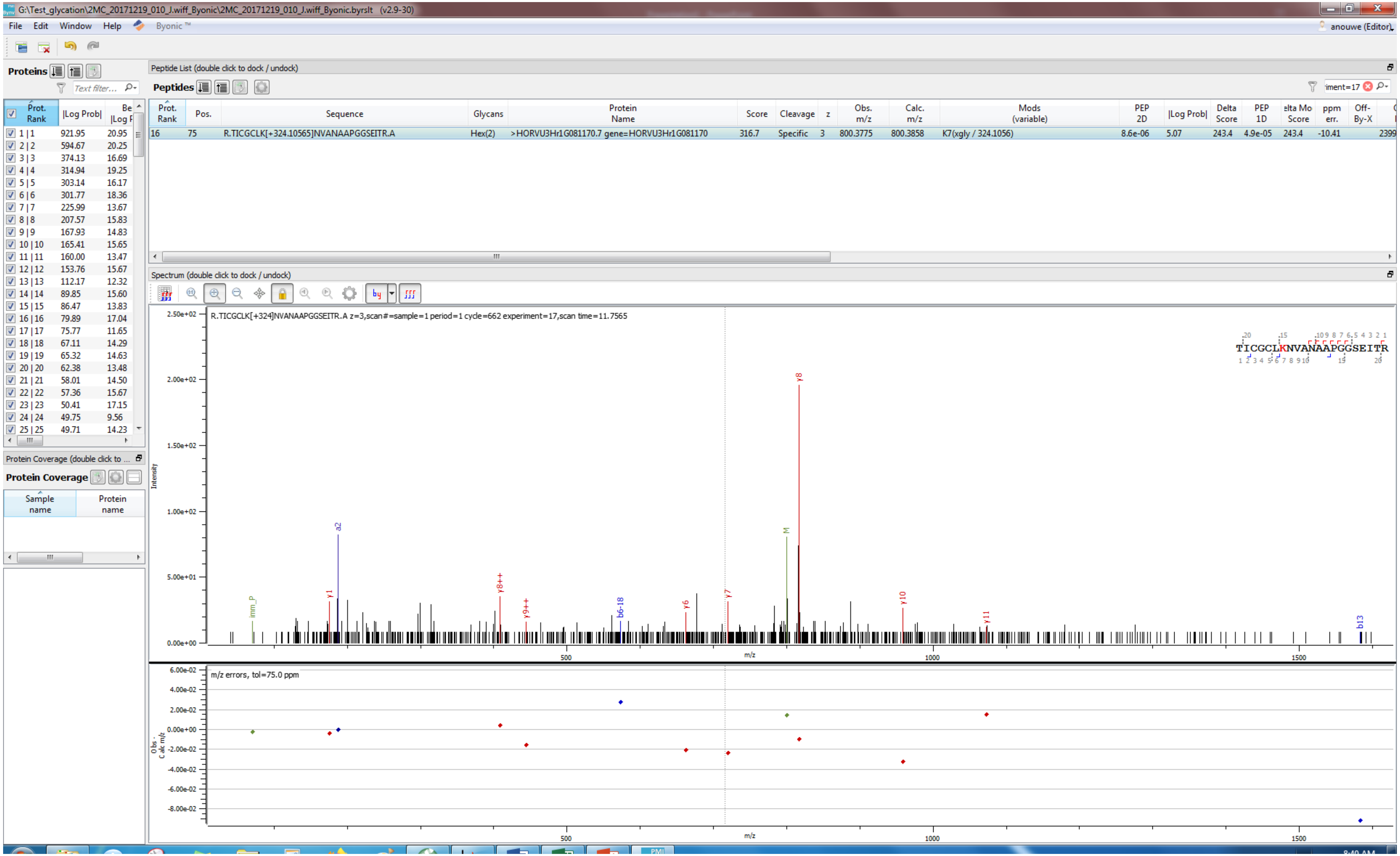
G:\Test_glycation\2MC_20171219_010_J.wiff_Byonic\2MC_20171219_010_J.wiff_Byonic.byrslt (v2.9-30)
File Edit Window Help Byonic™
anouwe (Editor)
Proteins
Peptide List (double click to dock / undock)
Peptides
Prot. Rank | Pos. | Sequence | Glycans | Protein Name | Score | Cleavage | z | Obs. m/z | Calc. m/z | Mods (variable) | PEP 2D | |Log Prob| | Delta Score | PEP 1D | elta Mo Score | ppm err. | Off-By-X
16 | 75 | R.TICGCLK[+324.10565]NVANAAPGGSEITR.A | Hex(2) | >HORVU3Hr1G081170.7 gene=HORVU3Hr1G081170 | 316.7 | Specific | 3 | 800.3775 | 800.3858 | K7(xgly / 324.1056) | 8.6e-06 | 5.07 | 243.4 | 4.9e-05 | 243.4 | -10.41 | 2399
Prot. Rank | |Log Prob|
1 | 1 921.95 20.95
2 | 2 594.67 20.25
3 | 3 374.13 16.69
4 | 4 314.94 19.25
5 | 5 303.14 16.17
6 | 6 301.77 18.36
7 | 7 225.99 13.67
8 | 8 207.57 15.83
9 | 9 167.93 14.83
10 | 10 165.41 15.65
11 | 11 160.00 13.47
12 | 12 153.76 15.67
13 | 13 112.17 12.32
14 | 14 89.85 15.60
15 | 15 86.47 13.83
16 | 16 79.89 17.04
17 | 17 75.77 11.65
18 | 18 67.11 14.29
19 | 19 65.32 14.63
20 | 20 62.38 13.48
21 | 21 58.01 14.50
22 | 22 57.36 15.67
23 | 23 50.41 17.15
24 | 24 49.75 9.56
25 | 25 49.71 14.23
Protein Coverage
Sample name
Protein name
Spectrum (double click to dock / undock)
R.TICGCLK[+324]NVANAAPGGSEITR.A z=3,scan#=sample=1 period=1 cycle=662 experiment=17,scan time=11.7565
TICGCLKNVANAAPGGSEITR
Intensity
m/z
y8
a2
M
imm_P
y1
y8++
y9++
b6-18
y6
y7
y10
y11
b13
500
1000
1500
m/z errors, tol=75.0 ppm
Obs - Calc m/z

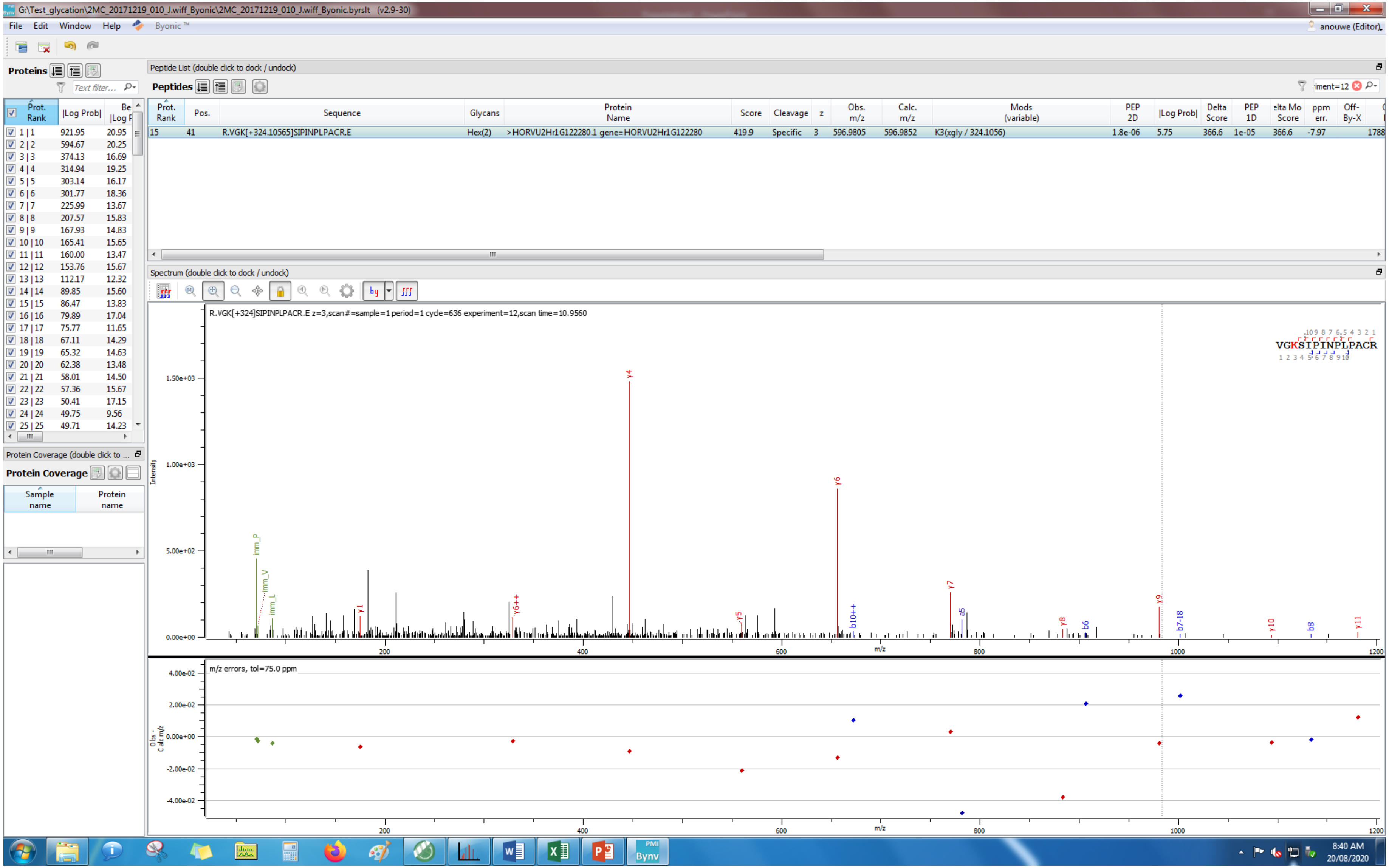
G:\Test_glycation\2MC_20171219_010_J.wiff_Byonic\2MC_20171219_010_J.wiff_Byonic.byrslt (v2.9-30)
File Edit Window Help Byonic
anouwe (Editor)
Proteins
Peptide List (double click to dock / undock)
Peptides
Prot. Rank | Pos. | Sequence | Glycans | Protein Name | Score | Cleavage | z | Obs. m/z | Calc. m/z | Mods (variable) | PEP 2D | |Log Prob| | Delta Score | PEP 1D | Delta Mod Score | ppm err. | Off-By-X
15 | 41 | R.VGK[+324.10565]SIPINPLPACR.E | Hex(2) | >HORVU2Hr1G122280.1 gene=HORVU2Hr1G122280 | 419.9 | Specific | 3 | 596.9805 | 596.9852 | K3(xgly / 324.1056) | 1.8e-06 | 5.75 | 366.6 | 1e-05 | 366.6 | -7.97 | 1788
Prot. Rank | |Log Prob| | Best |Log Prob|
1 | 1 | 921.95 | 20.95
2 | 2 | 594.67 | 20.25
3 | 3 | 374.13 | 16.69
4 | 4 | 314.94 | 19.25
5 | 5 | 303.14 | 16.17
6 | 6 | 301.77 | 18.36
7 | 7 | 225.99 | 13.67
8 | 8 | 207.57 | 15.83
9 | 9 | 167.93 | 14.83
10 | 10 | 165.41 | 15.65
11 | 11 | 160.00 | 13.47
12 | 12 | 153.76 | 15.67
13 | 13 | 112.17 | 12.32
14 | 14 | 89.85 | 15.60
15 | 15 | 86.47 | 13.83
16 | 16 | 79.89 | 17.04
17 | 17 | 75.77 | 11.65
18 | 18 | 67.11 | 14.29
19 | 19 | 65.32 | 14.63
20 | 20 | 62.38 | 13.48
21 | 21 | 58.01 | 14.50
22 | 22 | 57.36 | 15.67
23 | 23 | 50.41 | 17.15
24 | 24 | 49.75 | 9.56
25 | 25 | 49.71 | 14.23
Protein Coverage
Sample name
Protein name
Spectrum (double click to dock / undock)
R.VGK[+324]SIPINPLPACR.E z=3,scan#=sample=1 period=1 cycle=636 experiment=12,scan time=10.9560
VGKSIPINPLPACR
Intensity
m/z
m/z errors, tol=75.0 ppm
Obs - Calc m/z
8:40 AM
20/08/2020

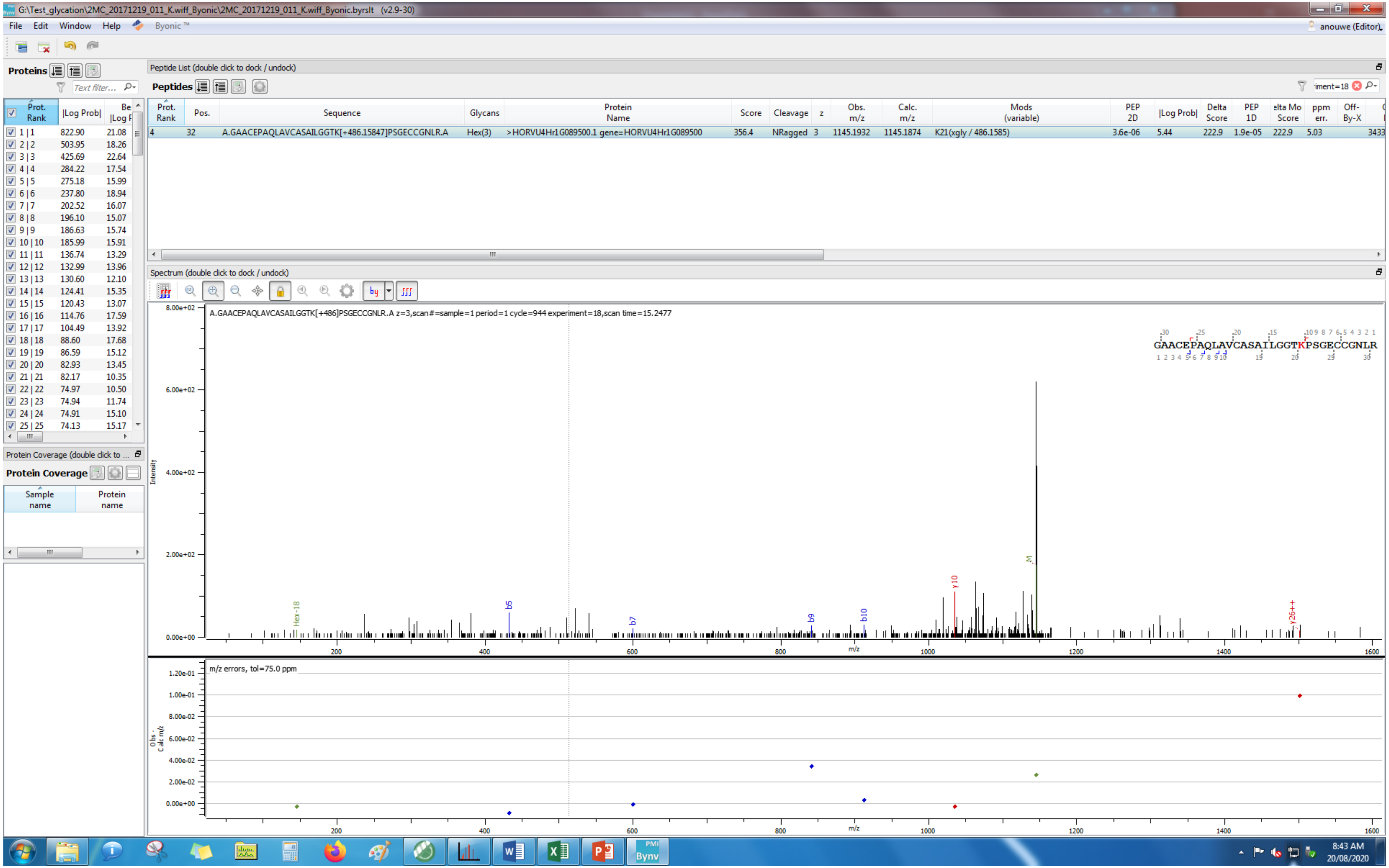
G:\Test_glycation\2MC_20171219_011_K.wiff_Byonic\2MC_20171219_011_K.wiff_Byonic.byrslt (v2.9-30)
File Edit Window Help Byonic
Proteins
Peptide List (double click to dock / undock)
Peptides
Prot. Rank | Pos. | Sequence | Glycans | Protein Name | Score | Cleavage | z | Obs. m/z | Calc. m/z | Mods (variable) | PEP 2D | |Log Prob| | Delta Score | PEP 1D | ppm err.
4 | 32 | A.GAACEPAQLAVCASAILGGTK[+486.15847]PSGECCGNLR.A | Hex(3) | >HORVU4Hr1G089500.1 gene=HORVU4Hr1G089500 | 356.4 | NRagged | 3 | 1145.1932 | 1145.1874 | K21(xgly / 486.1585) | 3.6e-06 | 5.44 | 222.9 | 1.9e-05 | 222.9 | 5.03
Spectrum (double click to dock / undock)
A.GAACEPAQLAVCASAILGGTK[+486]PSGECCGNLR.A z=3,scan#=sample=1 period=1 cycle=944 experiment=18,scan time=15.2477
GAACEPAQLAVCASAILGGTKPSGECCGNLR
Intensity
m/z
m/z errors, tol=75.0 ppm
Protein Coverage (double click to ...
Protein Coverage
Sample name
Protein name
8:43 AM
20/08/2020

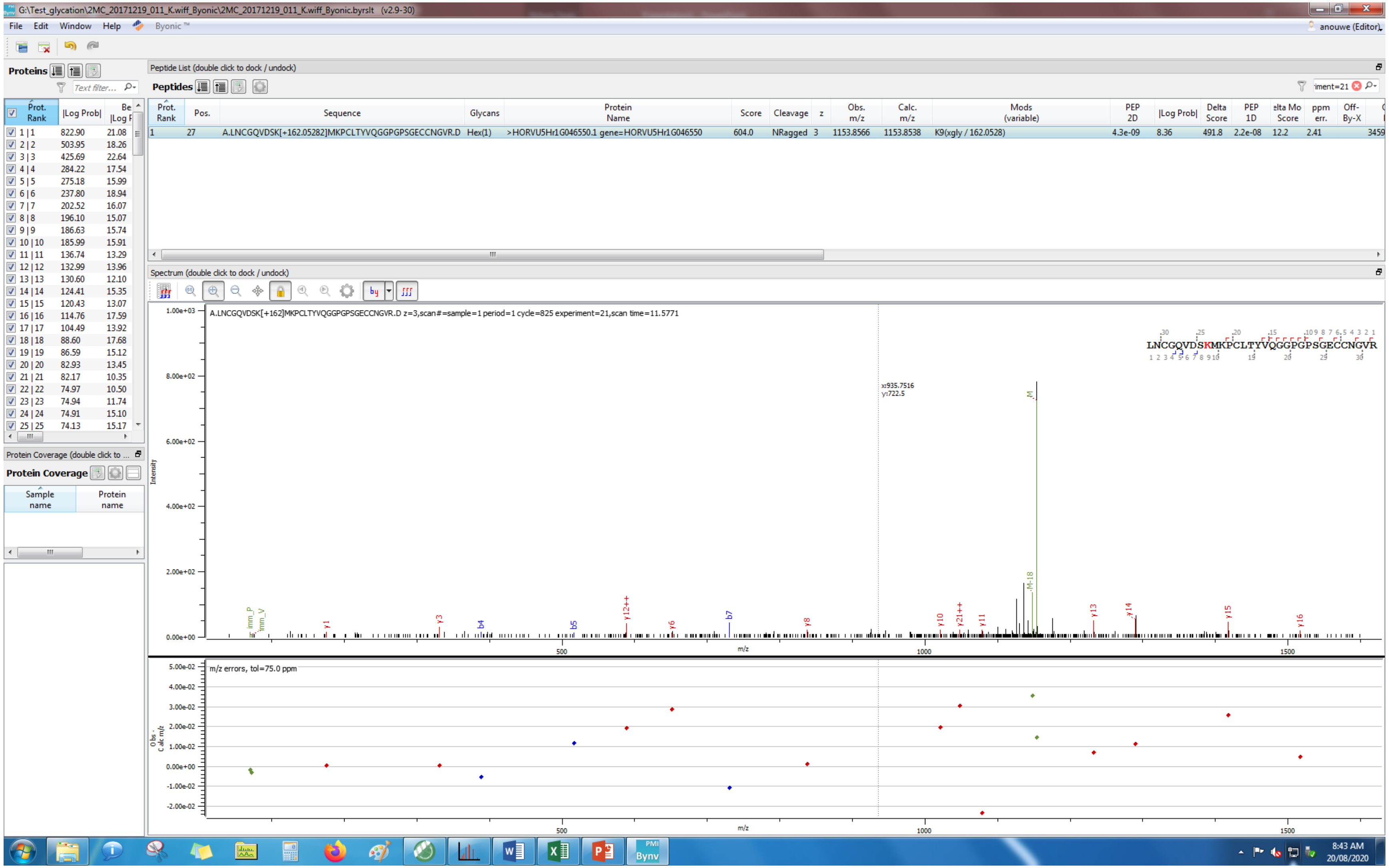
G:\Test_glycation\2MC_20171219_011_K.wiff_Byonic\2MC_20171219_011_K.wiff_Byonic.byrslt (v2.9-30)
File Edit Window Help Byonic ™
anouwe (Editor)
Proteins
Peptide List (double click to dock / undock)
Peptides
Prot. Rank | Pos. | Sequence | Glycans | Protein Name | Score | Cleavage | z | Obs. m/z | Calc. m/z | Mods (variable) | PEP 2D | |Log Prob| | Delta Score | PEP 1D | Delta Mod Score | ppm err. | Off-By-X
1 | 27 | A.LNCGQVDSK[+162.05282]MKPCLTYVQGGPGPSGECCNGVR.D | Hex(1) | >HORVU5Hr1G046550.1 gene=HORVU5Hr1G046550 | 604.0 | NRagged | 3 | 1153.8566 | 1153.8538 | K9(xgly / 162.0528) | 4.3e-09 | 8.36 | 491.8 | 2.2e-08 | 12.2 | 2.41
Spectrum (double click to dock / undock)
A.LNCGQVDSK[+162]MKPCLTYVQGGPGPSGECCNGVR.D z=3,scan#=sample=1 period=1 cycle=825 experiment=21,scan time=11.5771
Intensity
m/z
m/z errors, tol=75.0 ppm
Protein Coverage (double click to ...
Protein Coverage
Sample name
Protein name
8:43 AM
20/08/2020

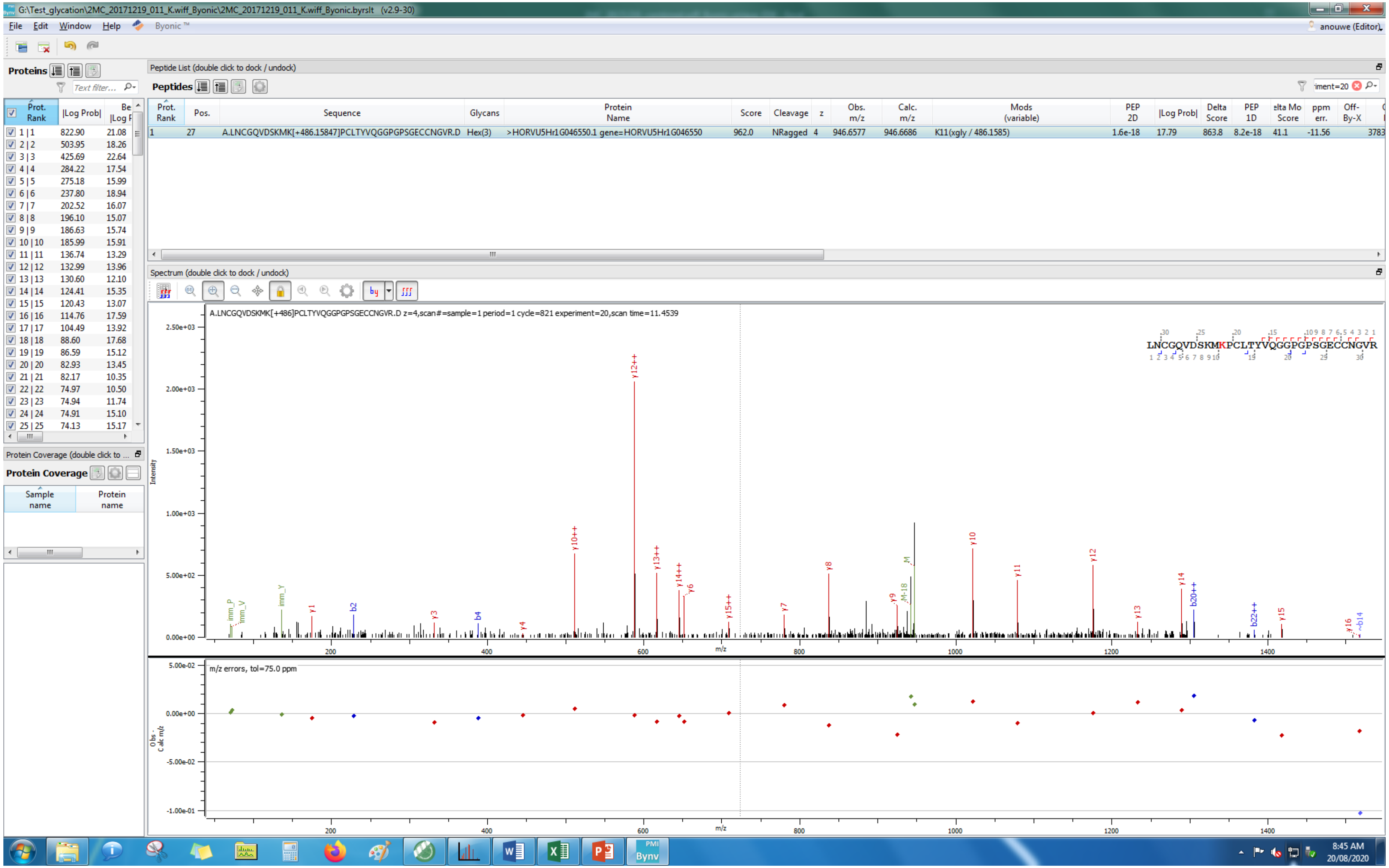

G:\Test_glycation\2MC_20171219_011_K.wiff_Byonic\2MC_20171219_011_K.wiff_Byonic.byrslt (v2.9-30)
File Edit Window Help Byonic™
anouwe (Editor)
Proteins
Peptide List (double click to dock / undock)
Peptides
Prot. Rank | Pos. | Sequence | Glycans | Protein Name | Score | Cleavage | z | Obs. m/z | Calc. m/z | Mods (variable) | PEP 2D | |Log Prob| | Delta Score | PEP 1D | ppm err.
1 | 27 | A.LNCGQVDSKMK[+486.15847]PCLTYVQGGPGPSGECCNGVR.D | Hex(3) | >HORVU5Hr1G046550.1 gene=HORVU5Hr1G046550 | 962.0 | NRagged | 4 | 946.6577 | 946.6686 | K11(xgly / 486.1585) | 1.6e-18 | 17.79 | 863.8 | 8.2e-18 | 41.1 | -11.56
Spectrum (double click to dock / undock)
A.LNCGQVDSKMK[+486]PCLTYVQGGPGPSGECCNGVR.D z=4,scan#=sample=1 period=1 cycle=821 experiment=20,scan time=11.4539
LNCGQVDSKMKPCLTYVQGGPGPSGECCNGVR
Intensity
m/z
m/z errors, tol=75.0 ppm
Obs - Calc m/z
Protein Coverage
Sample name
Protein name
8:45 AM
20/08/2020

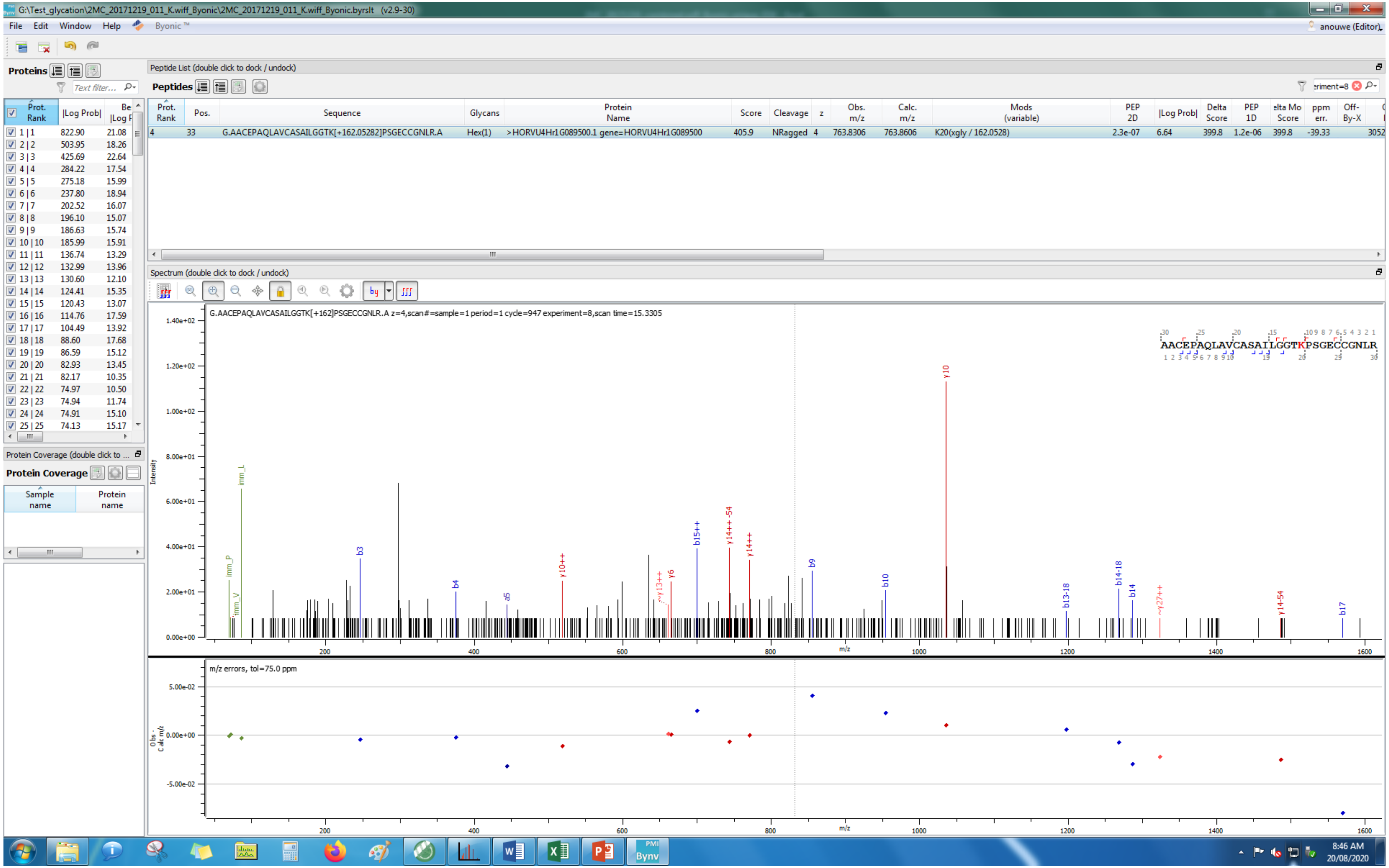
G:\Test_glycation\2MC_20171219_011_K.wiff_Byonic\2MC_20171219_011_K.wiff_Byonic.byrslt (v2.9-30)
File Edit Window Help Byonic ™
anouwe (Editor)
Proteins
Peptide List (double click to dock / undock)
Peptides
Prot. Rank
Pos.
Sequence
Glycans
Protein Name
Score
Cleavage
z
Obs. m/z
Calc. m/z
Mods (variable)
PEP 2D
|Log Prob|
Delta Score
PEP 1D
Delta Mod Score
ppm err.
Off-By-X
4 33 G.AACEPAQLAVCASAILGGTK[+162.05282]PSGECCGNLR.A Hex(1) >HORVU4Hr1G089500.1 gene=HORVU4Hr1G089500 405.9 NRagged 4 763.8306 763.8606 K20(xgly / 162.0528) 2.3e-07 6.64 399.8 1.2e-06 399.8 -39.33
Spectrum (double click to dock / undock)
G.AACEPAQLAVCASAILGGTK[+162]PSGECCGNLR.A z=4,scan#=sample=1 period=1 cycle=947 experiment=8,scan time=15.3305
AACEPAQLAVCASAILGGTKPSGECCGNLR
Intensity
m/z
y10
b15++
y14++-54
y14++
y10++
y13++
y6
b3
b4
a5
b9
b10
b13-18
b14-18
b14
y27++
y14-54
b17
imm_L
imm_P
imm_V
m/z errors, tol=75.0 ppm
Obs - Calc m/z
Protein Coverage
Sample name
Protein name
8:46 AM
20/08/2020

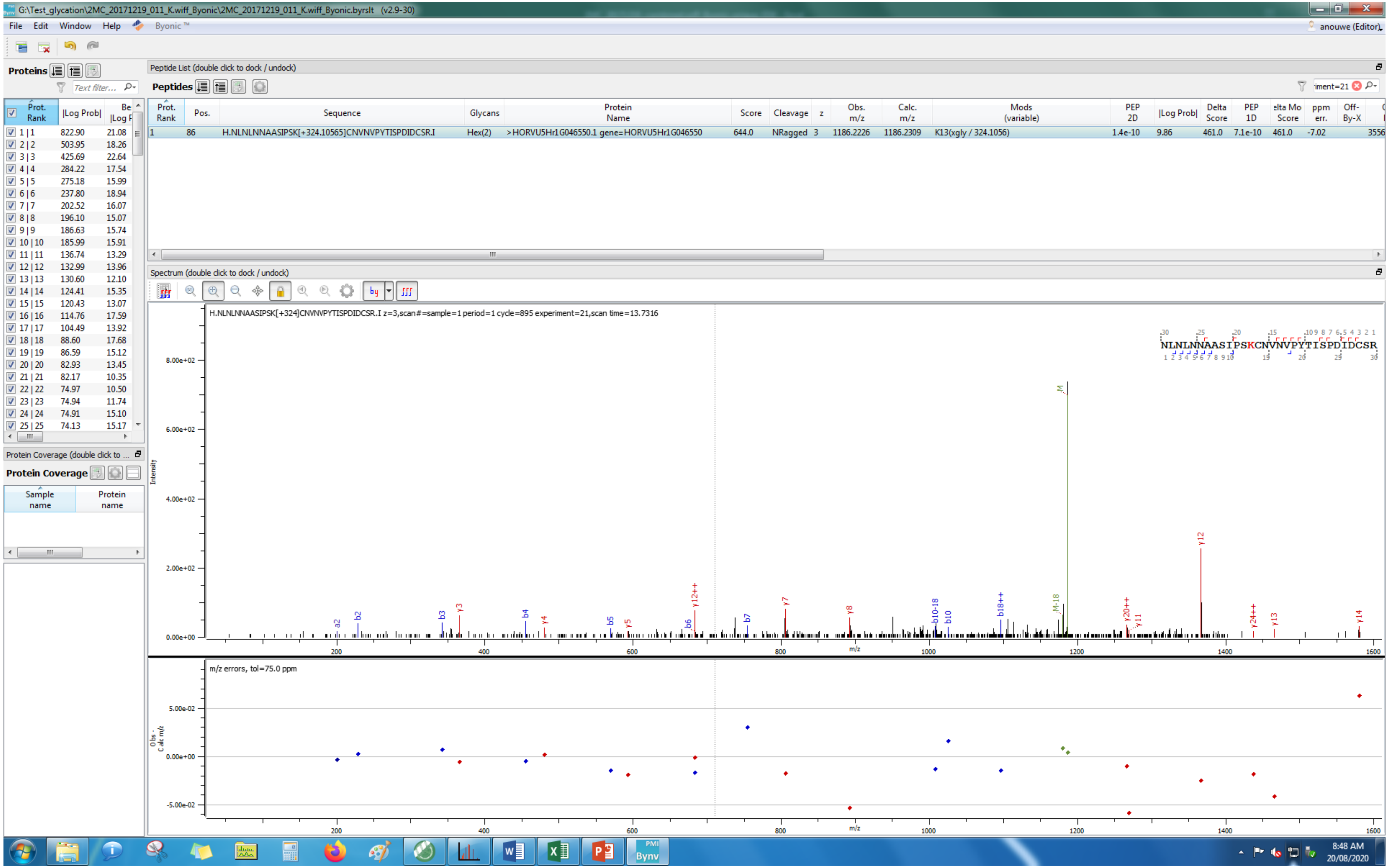
G:\Test_glycation\2MC_20171219_011_K.wiff_Byonic\2MC_20171219_011_K.wiff_Byonic.byrslt (v2.9-30)
File Edit Window Help Byonic™
Proteins
Peptide List (double click to dock / undock)
Peptides
H.NLNLNNAASIPSK[+324.10565]CNVNVPYTISPDIDCSR.I
Hex(2)
>HORVU5Hr1G046550.1 gene=HORVU5Hr1G046550
644.0
NRagged
1186.2226
1186.2309
K13(xgly / 324.1056)
Spectrum (double click to dock / undock)
H.NLNLNNAASIPSK[+324]CNVNVPYTISPDIDCSR.I z=3,scan#=sample=1 period=1 cycle=895 experiment=21,scan time=13.7316
NLNLNNAASIPSKCNVNVPYTISPDIDCSR
Intensity
m/z
m/z errors, tol=75.0 ppm
Protein Coverage
8:48 AM
20/08/2020

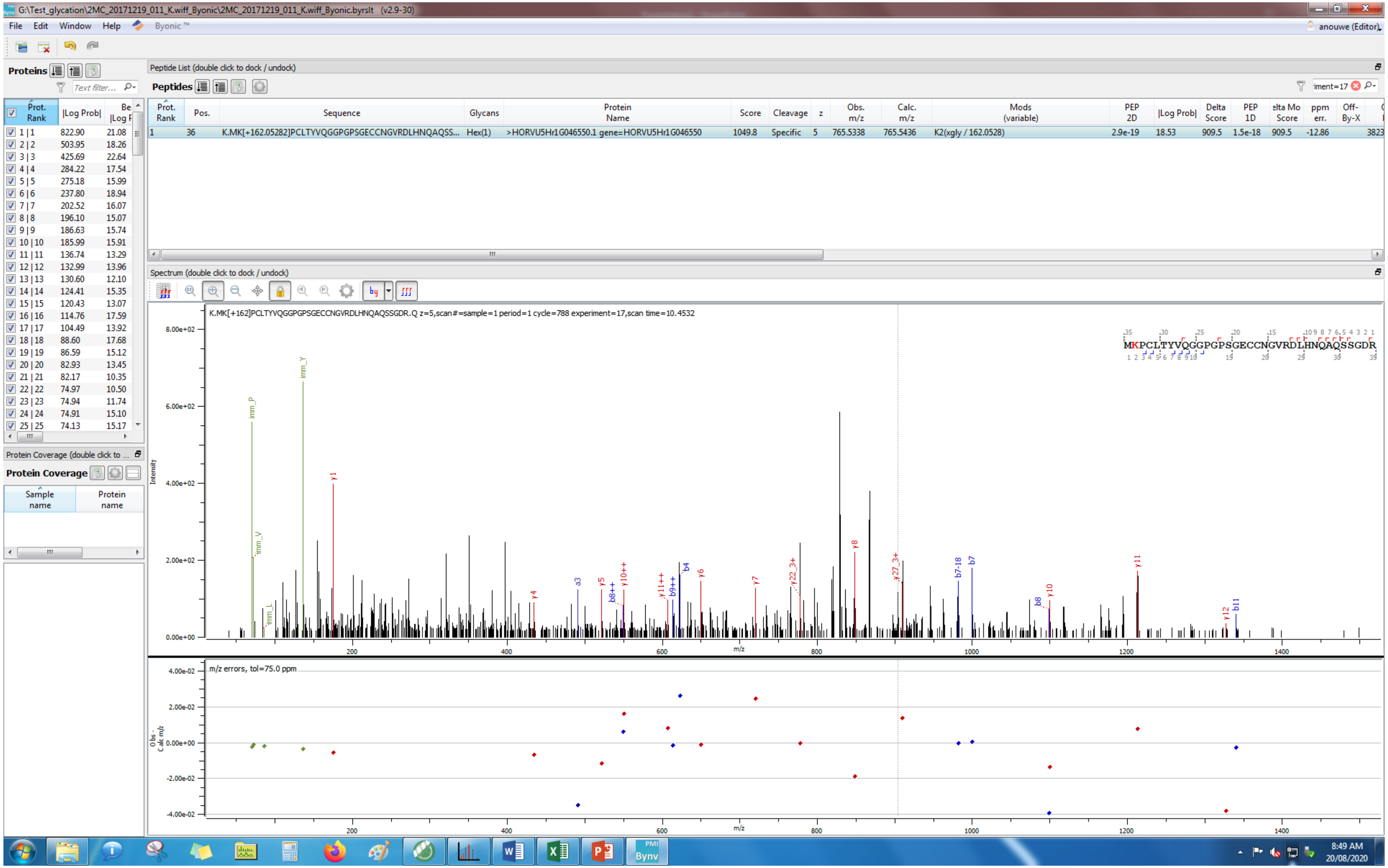
G:\Test_glycation\2MC_20171219_011_K.wiff_Byonic\2MC_20171219_011_K.wiff_Byonic.byrslt (v2.9-30)
File Edit Window Help Byonic™
anouwe (Editor)
Proteins
Peptide List (double click to dock / undock)
Peptides
Prot. Rank | Pos. | Sequence | Glycans | Protein Name | Score | Cleavage | z | Obs. m/z | Calc. m/z | Mods (variable) | PEP 2D | |Log Prob| | Delta Score | PEP 1D | Delta Mod Score | ppm err. | Off-By-X
1 | 36 | K.MK[+162.05282]PCLTYVQGGPGPSGECCNGVRDLHNQAQSS... | Hex(1) | >HORVU5Hr1G046550.1 gene=HORVU5Hr1G046550 | 1049.8 | Specific | 5 | 765.5338 | 765.5436 | K2(xgly / 162.0528) | 2.9e-19 | 18.53 | 909.5 | 1.5e-18 | 909.5 | -12.86
Spectrum (double click to dock / undock)
K.MK[+162]PCLTYVQGGPGPSGECCNGVRDLHNQAQSSGDR.Q z=5,scan#=sample=1 period=1 cycle=788 experiment=17,scan time=10.4532
MKPCLTYVQGGPGPSGECCNGVRDLHNQAQSSGDR
Intensity
m/z
m/z errors, tol=75.0 ppm
Protein Coverage
Sample name
Protein name
8:49 AM
20/08/2020

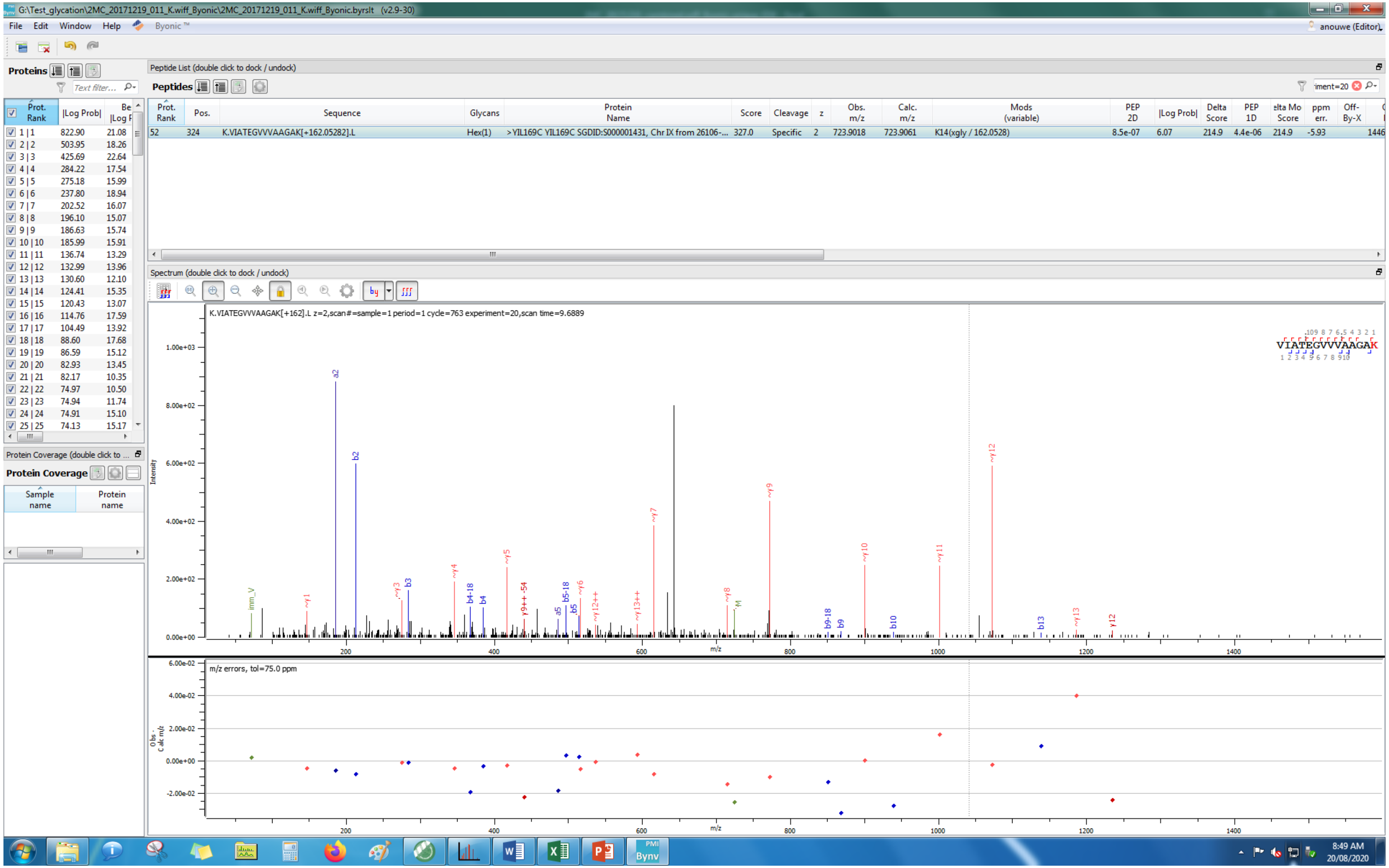
G:\Test_glycation\2MC_20171219_011_K.wiff_Byonic\2MC_20171219_011_K.wiff_Byonic.byrslt (v2.9-30)
File Edit Window Help Byonic™
anouwe (Editor)
Proteins
Peptide List (double click to dock / undock)
Peptides
Prot. Rank | Pos. | Sequence | Glycans | Protein Name | Score | Cleavage | z | Obs. m/z | Calc. m/z | Mods (variable) | PEP 2D | |Log Prob| | Delta Score | PEP 1D | Delta Mod Score | ppm err. | Off-By-X
52 | 324 | K.VIATEGVVVAAGAK[+162.05282].L | Hex(1) | >YIL169C YIL169C SGDID:S000001431, Chr IX from 26106-... | 327.0 | Specific | 2 | 723.9018 | 723.9061 | K14(xgly / 162.0528) | 8.5e-07 | 6.07 | 214.9 | 4.4e-06 | 214.9 | -5.93 | 1446
Spectrum (double click to dock / undock)
K.VIATEGVVVAAGAK[+162].L z=2,scan#=sample=1 period=1 cycle=763 experiment=20,scan time=9.6889
VIATEGVVVAAGAK
Intensity
m/z
m/z errors, tol=75.0 ppm
Protein Coverage (double click to ...
Protein Coverage
Sample name
Protein name
8:49 AM
20/08/2020

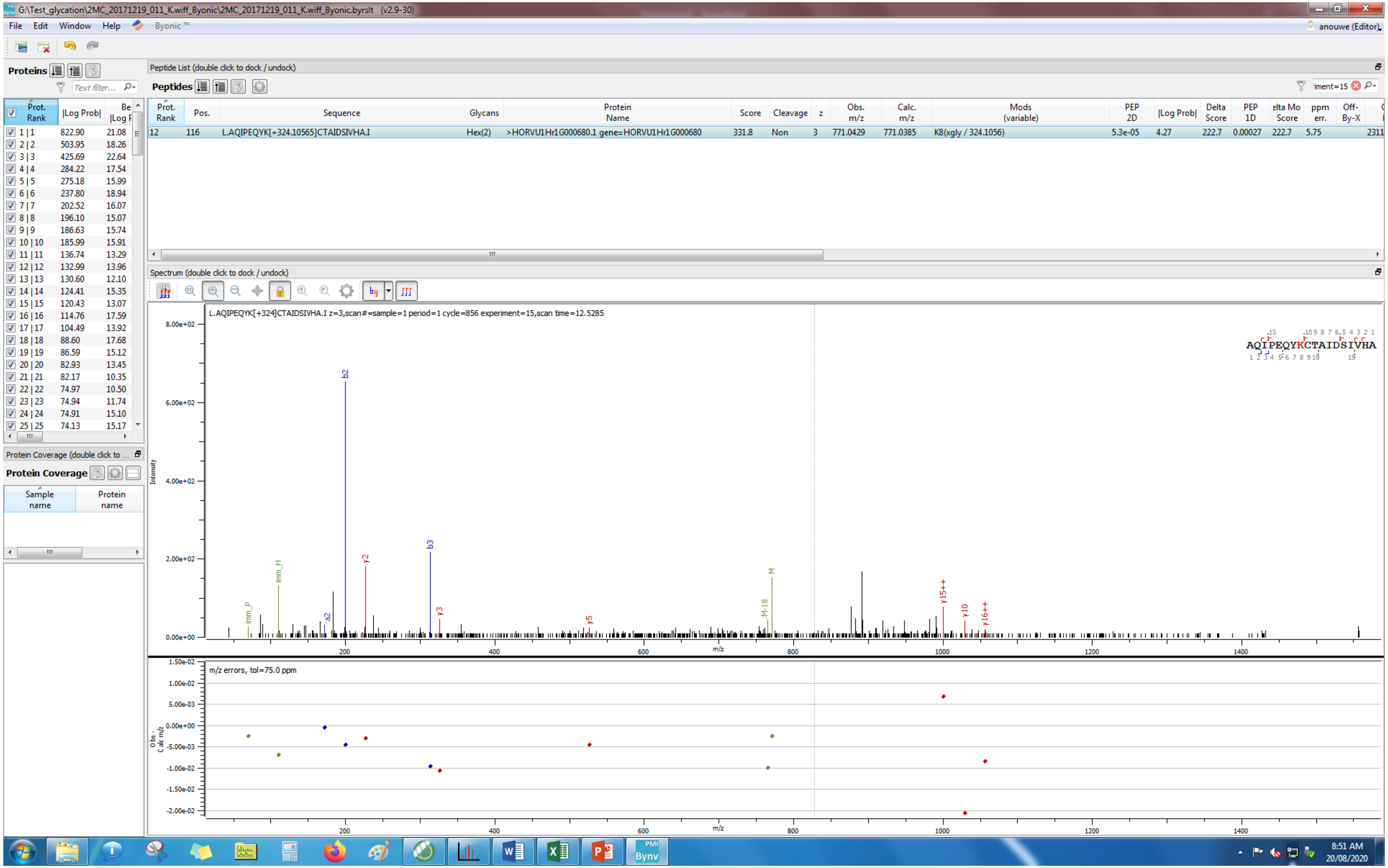
G:\Test_glycation\2MC_20171219_011_K.wiff_Byonic\2MC_20171219_011_K.wiff_Byonic.byrslt (v2.9-30)
File Edit Window Help Byonic ™
anouwe (Editor)
Proteins
Peptide List (double click to dock / undock)
Peptides
Prot. Rank | Pos. | Sequence | Glycans | Protein Name | Score | Cleavage | z | Obs. m/z | Calc. m/z | Mods (variable) | PEP 2D | |Log Prob| | Delta Score | PEP 1D | Delta Mod Score | ppm err. | Off-By-X
12 | 116 | L.AQIPEQYK[+324.10565]CTAIDSIVHA.I | Hex(2) | >HORVU1Hr1G000680.1 gene=HORVU1Hr1G000680 | 331.8 | Non | 3 | 771.0429 | 771.0385 | K8(xgly / 324.1056) | 5.3e-05 | 4.27 | 222.7 | 0.00027 | 222.7 | 5.75
Spectrum (double click to dock / undock)
L.AQIPEQYK[+324]CTAIDSIVHA.I z=3,scan#=sample=1 period=1 cycle=856 experiment=15,scan time=12.5285
AQIPEQYKCTAIDSIVHA
Intensity
m/z
m/z errors, tol=75.0 ppm
Protein Coverage (double click to ...
Protein Coverage
Sample name
Protein name
8:51 AM
20/08/2020

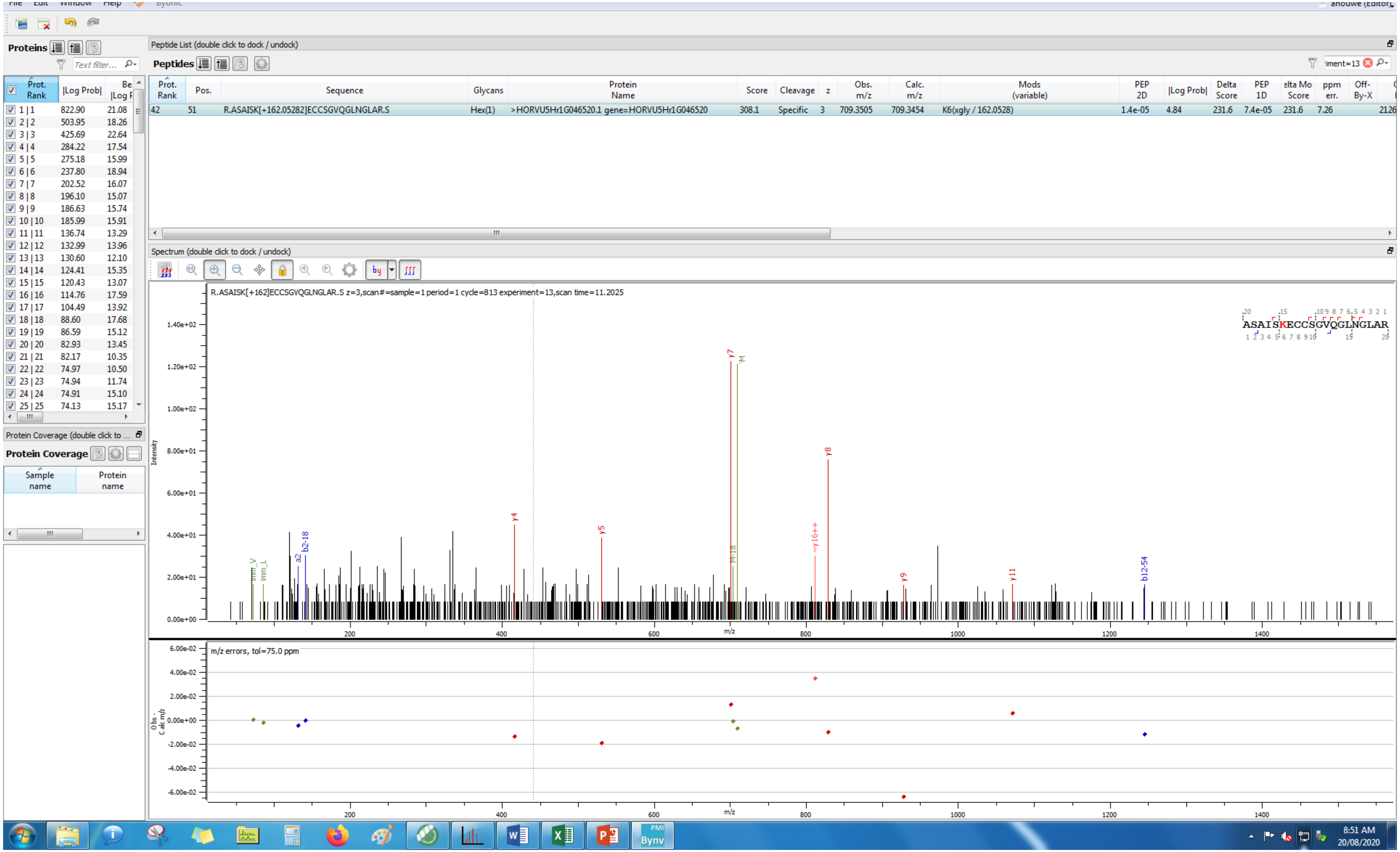
Proteins
Peptide List (double click to dock / undock)
Peptides
Prot. Rank | Pos. | Sequence | Glycans | Protein Name | Score | Cleavage | z | Obs. m/z | Calc. m/z | Mods (variable) | PEP 2D | |Log Prob| | Delta Score | PEP 1D | Delta Mod Score | ppm err. | Off-By-X
42 | 51 | R.ASAISK[+162.05282]ECCSGVQGLNGLAR.S | Hex(1) | >HORVU5Hr1G046520.1 gene=HORVU5Hr1G046520 | 308.1 | Specific | 3 | 709.3505 | 709.3454 | K6(xgly / 162.0528) | 1.4e-05 | 4.84 | 231.6 | 7.4e-05 | 231.6 | 7.26
Spectrum (double click to dock / undock)
R.ASAISK[+162]ECCSGVQGLNGLAR.S z=3,scan#=sample=1 period=1 cycle=813 experiment=13,scan time=11.2025
ASAISKECCSGVQGLNGLAR
Intensity
m/z
y4
y5
y7
y8
y9
y11
b2-18
a2
b12-54
~y16++
M-18
m/z errors, tol=75.0 ppm
Obs - Calc m/z
Protein Coverage (double click to dock / undock)
Protein Coverage
Sample name
Protein name
8:51 AM
20/08/2020

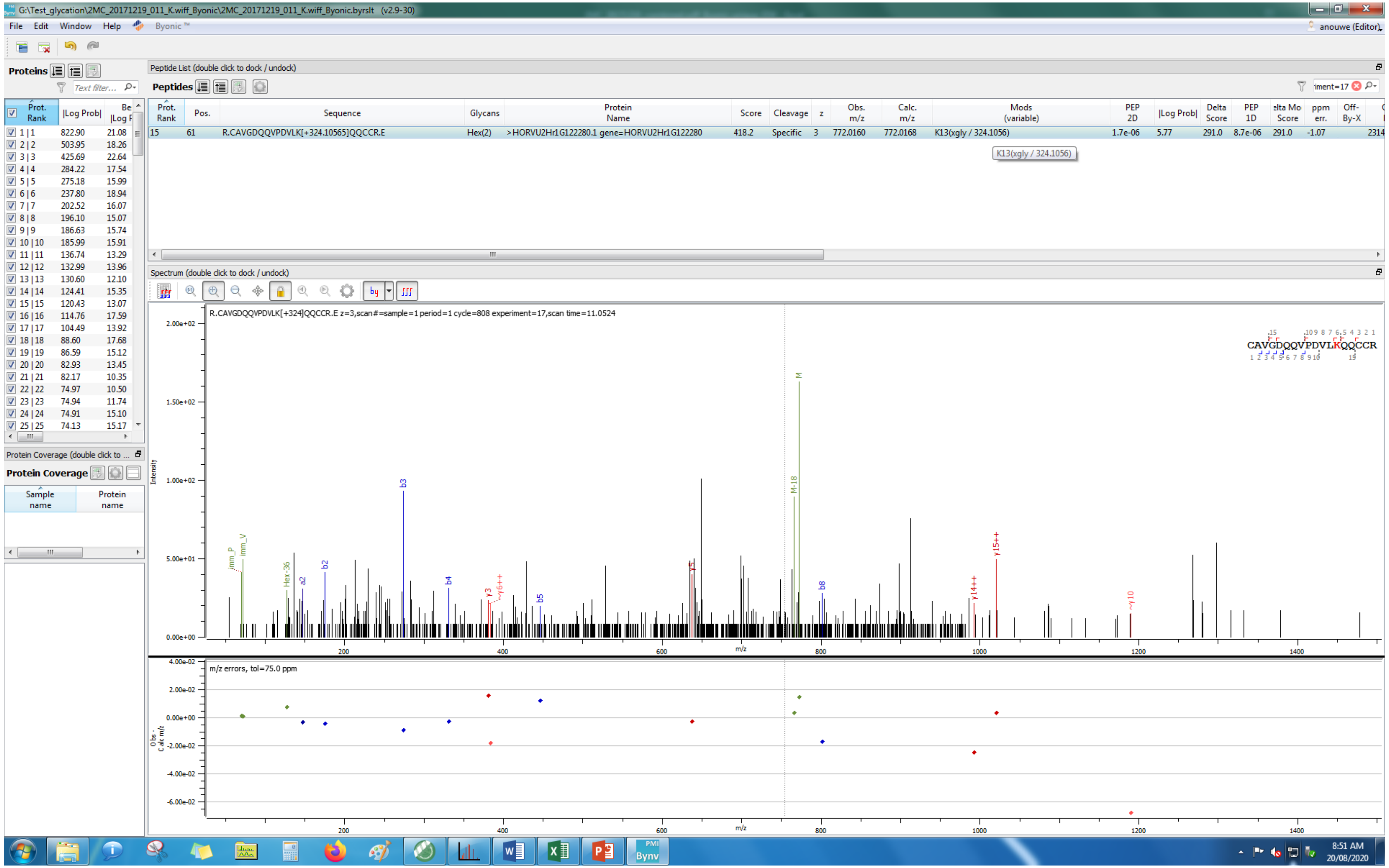

G:\Test_glycation\2MC_20171219_011_K.wiff_Byonic\2MC_20171219_011_K.wiff_Byonic.byrslt (v2.9-30)
File Edit Window Help Byonic ™
anouwe (Editor)
Proteins
Peptide List (double click to dock / undock)
Peptides
Prot. Rank | Pos. | Sequence | Glycans | Protein Name | Score | Cleavage | z | Obs. m/z | Calc. m/z | Mods (variable) | PEP 2D | |Log Prob| | Delta Score | PEP 1D | elta Mo Score | ppm err. | Off-By-X
15 | 61 | R.CAVGDQQVPDVLK[+324.10565]QQCCR.E | Hex(2) | >HORVU2Hr1G122280.1 gene=HORVU2Hr1G122280 | 418.2 | Specific | 3 | 772.0160 | 772.0168 | K13(xgly / 324.1056) | 1.7e-06 | 5.77 | 291.0 | 8.7e-06 | 291.0 | -1.07 | 2314
K13(xgly / 324.1056)
Spectrum (double click to dock / undock)
R.CAVGDQQVPDVLK[+324]QQCCR.E z=3,scan#=sample=1 period=1 cycle=808 experiment=17,scan time=11.0524
CAVGDQQVPDVLKQQCCR
Intensity
m/z
m/z errors, tol=75.0 ppm
Protein Coverage (double click to ...
Protein Coverage
Sample name
Protein name
8:51 AM
20/08/2020

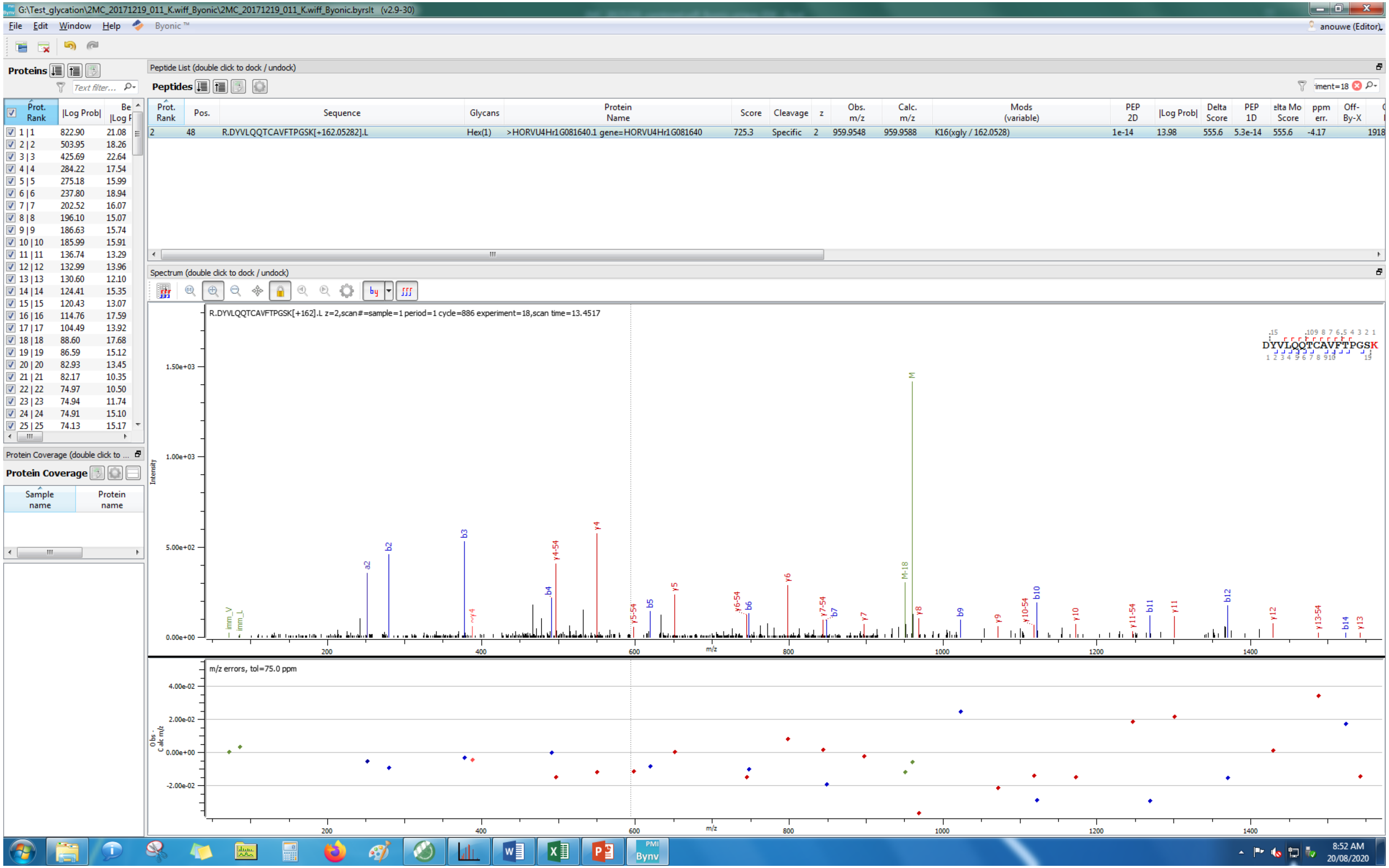
G:\Test_glycation\2MC_20171219_011_K.wiff_Byonic\2MC_20171219_011_K.wiff_Byonic.byrslt (v2.9-30)
File Edit Window Help Byonic™
anouwe (Editor)
Proteins
Peptide List (double click to dock / undock)
Peptides
Prot. Rank | Pos. | Sequence | Glycans | Protein Name | Score | Cleavage | z | Obs. m/z | Calc. m/z | Mods (variable) | PEP 2D | |Log Prob| | Delta Score | PEP 1D | Delta Mod Score | ppm err. | Off-By-X
2 | 48 | R.DYVLQQTCAVFTPGSK[+162.05282].L | Hex(1) | >HORVU4Hr1G081640.1 gene=HORVU4Hr1G081640 | 725.3 | Specific | 2 | 959.9548 | 959.9588 | K16(xgly / 162.0528) | 1e-14 | 13.98 | 555.6 | 5.3e-14 | 555.6 | -4.17 | 1918
Spectrum (double click to dock / undock)
R.DYVLQQTCAVFTPGSK[+162].L z=2,scan#=sample=1 period=1 cycle=886 experiment=18,scan time=13.4517
DYVLQQTCAVFTPGSK
Intensity
m/z
m/z errors, tol=75.0 ppm
Obs - Calc m/z
Protein Coverage (double click to ...
Protein Coverage
Sample name
Protein name
8:52 AM
20/08/2020

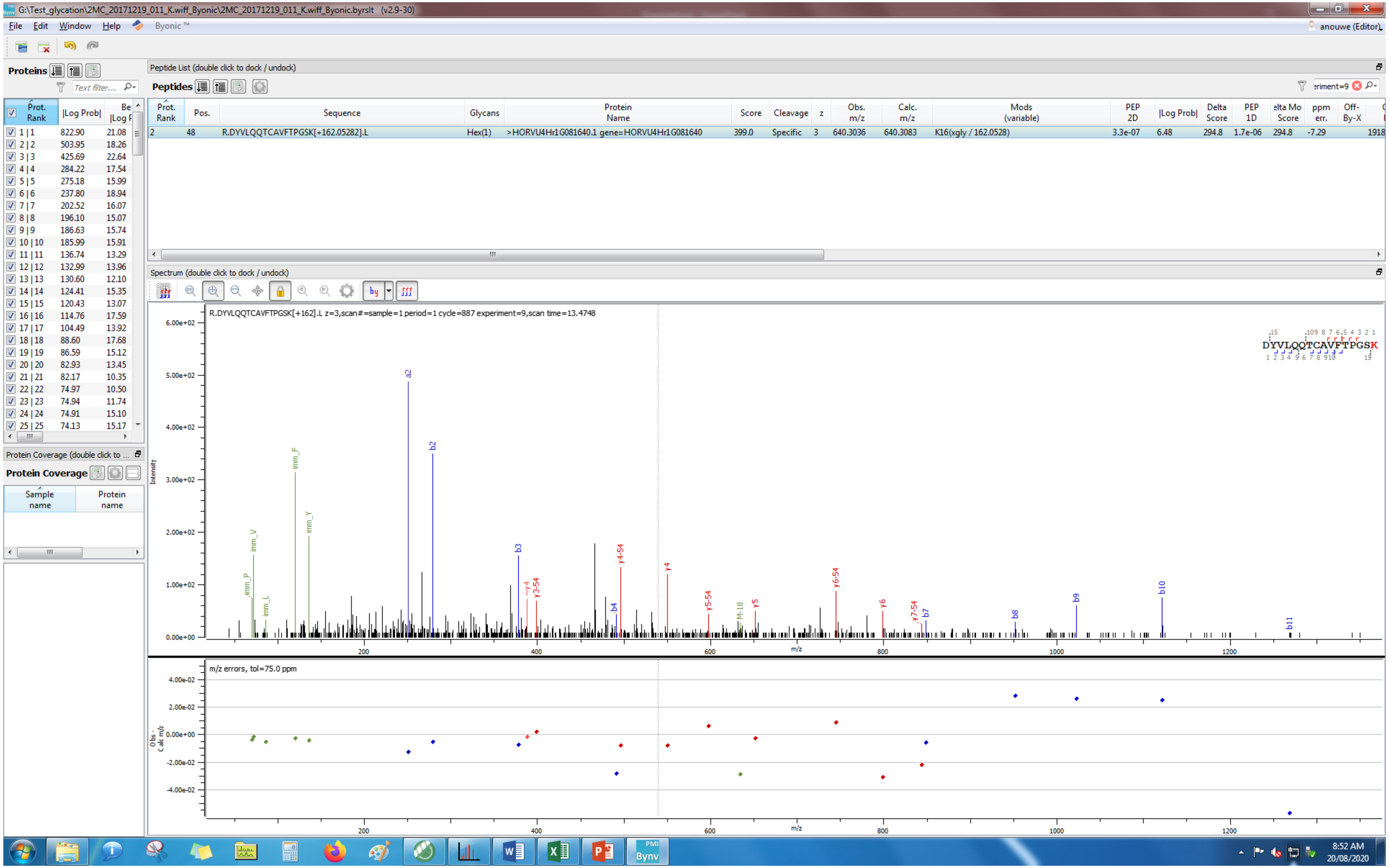
G:\Test_glycation\2MC_20171219_011_K.wiff_Byonic\2MC_20171219_011_K.wiff_Byonic.byrslt (v2.9-30)
File Edit Window Help Byonic ™
anouwe (Editor)
Proteins
Peptide List (double click to dock / undock)
Peptides
Prot. Rank | |Log Prob|
1 | 1 822.90 21.08
2 | 2 503.95 18.26
3 | 3 425.69 22.64
4 | 4 284.22 17.54
5 | 5 275.18 15.99
6 | 6 237.80 18.94
7 | 7 202.52 16.07
8 | 8 196.10 15.07
9 | 9 186.63 15.74
10 | 10 185.99 15.91
11 | 11 136.74 13.29
12 | 12 132.99 13.96
13 | 13 130.60 12.10
14 | 14 124.41 15.35
15 | 15 120.43 13.07
16 | 16 114.76 17.59
17 | 17 104.49 13.92
18 | 18 88.60 17.68
19 | 19 86.59 15.12
20 | 20 82.93 13.45
21 | 21 82.17 10.35
22 | 22 74.97 10.50
23 | 23 74.94 11.74
24 | 24 74.91 15.10
25 | 25 74.13 15.17
Prot. Rank Pos. Sequence Glycans Protein Name Score Cleavage z Obs. m/z Calc. m/z Mods (variable) PEP 2D |Log Prob| Delta Score PEP 1D Delta Mod Score ppm err. Off-By-X
2 48 R.DYVLQQTCAVFTPGSK[+162.05282].L Hex(1) >HORVU4Hr1G081640.1 gene=HORVU4Hr1G081640 399.0 Specific 3 640.3036 640.3083 K16(xgly / 162.0528) 3.3e-07 6.48 294.8 1.7e-06 294.8 -7.29 1918
Protein Coverage (double click to ...
Protein Coverage
Sample name
Protein name
Spectrum (double click to dock / undock)
R.DYVLQQTCAVFTPGSK[+162].L z=3,scan#=sample=1 period=1 cycle=887 experiment=9,scan time=13.4748
DYVLQQTCAVFTPGSK
Intensity
m/z
imm_P imm_V imm_L imm_F imm_Y a2 b2 b3 y4 y3-54 b4 y4-54 y4 y5-54 M-18 y5 y6-54 y6 y7-54 b7 b8 b9 b10 b11
m/z errors, tol=75.0 ppm
Obs - Calc m/z
8:52 AM
20/08/2020

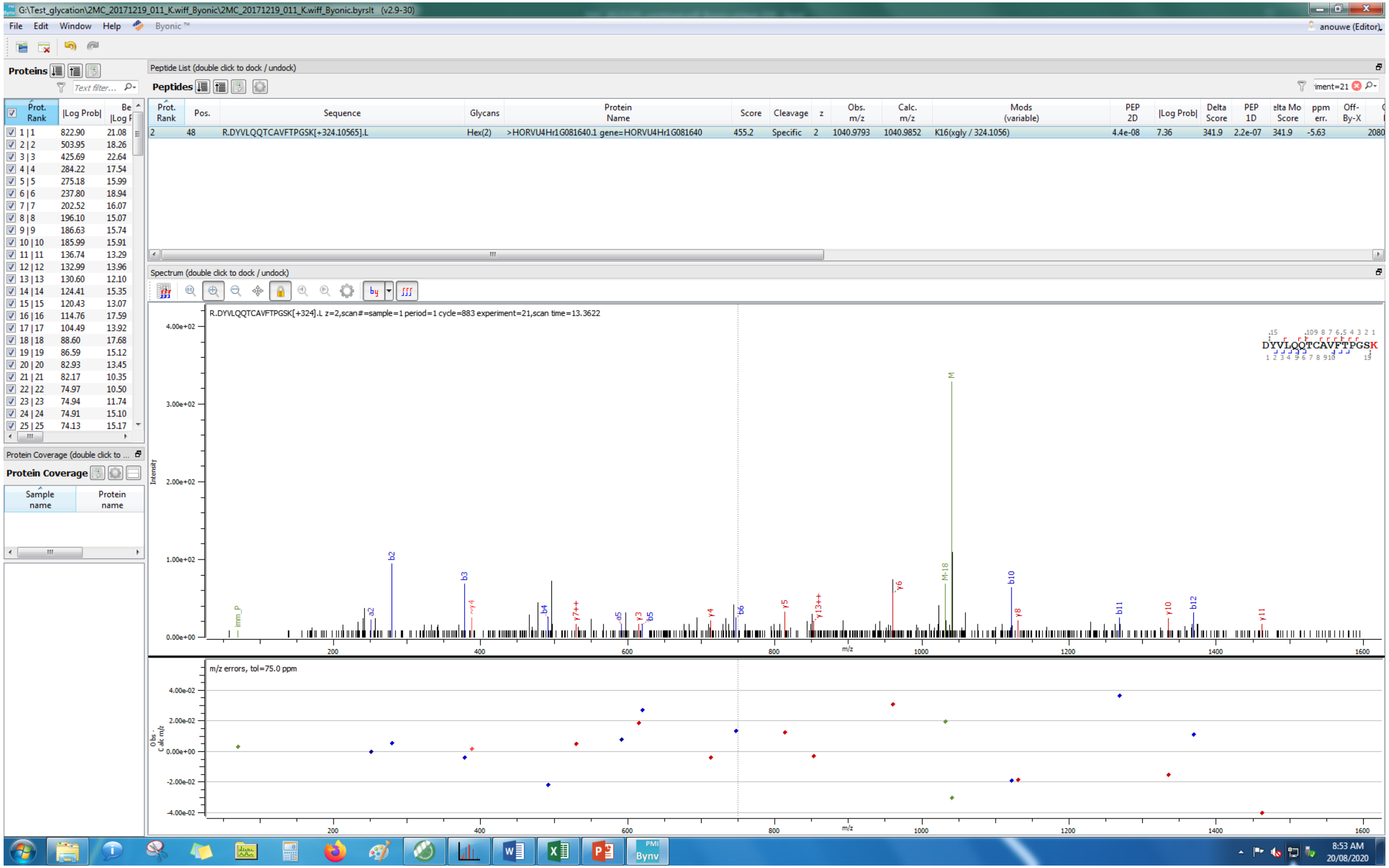

G:\Test_glycation\2MC_20171219_011_K.wiff_Byonic\2MC_20171219_011_K.wiff_Byonic.byrslt (v2.9-30)
File Edit Window Help Byonic
Proteins
Peptide List (double click to dock / undock)
Peptides
Prot. Rank | Pos. | Sequence | Glycans | Protein Name | Score | Cleavage | z | Obs. m/z | Calc. m/z | Mods (variable) | PEP 2D | |Log Prob| | Delta Score | PEP 1D | Delta Mod Score | ppm err.
2 | 48 | R.DYVLQQTCAVFTPGSK[+324.10565].L | Hex(2) | >HORVU4Hr1G081640.1 gene=HORVU4Hr1G081640 | 455.2 | Specific | 2 | 1040.9793 | 1040.9852 | K16(xgly / 324.1056) | 4.4e-08 | 7.36 | 341.9 | 2.2e-07 | 341.9 | -5.63
Spectrum (double click to dock / undock)
R.DYVLQQTCAVFTPGSK[+324].L z=2,scan#=sample=1 period=1 cycle=883 experiment=21,scan time=13.3622
DYVLQQTCAVFTPGSK
Intensity
m/z
m/z errors, tol=75.0 ppm
Protein Coverage (double click to ...
Protein Coverage
Sample name
Protein name
anouwe (Editor)
8:53 AM
20/08/2020

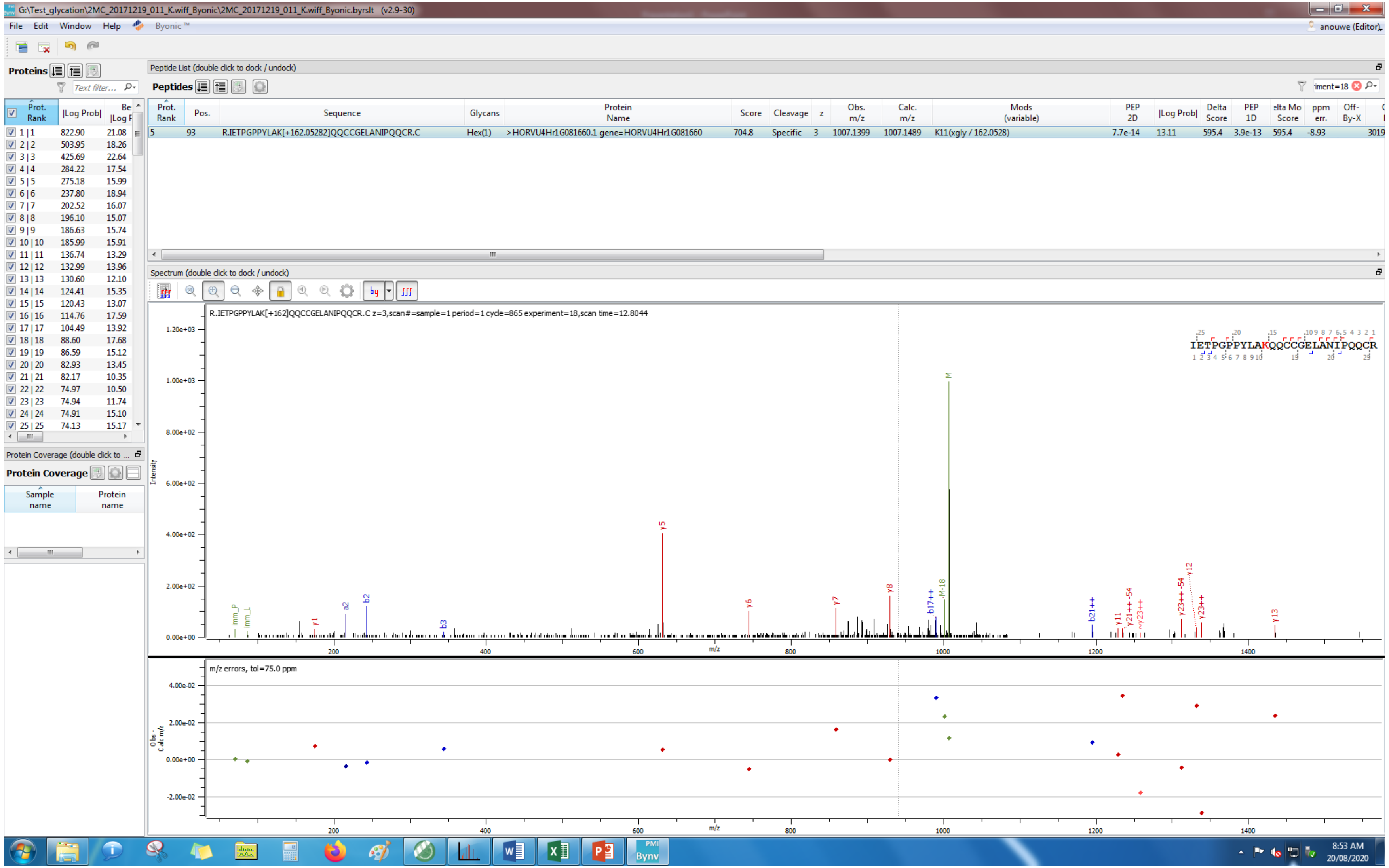
G:\Test_glycation\2MC_20171219_011_K.wiff_Byonic\2MC_20171219_011_K.wiff_Byonic.byrslt (v2.9-30)
File Edit Window Help Byonic ™
anouwe (Editor)
Proteins
Peptide List (double click to dock / undock)
Peptides
R.IETPGPPYLAK[+162.05282]QQCCGELANIPQQCR.C
Hex(1)
>HORVU4Hr1G081660.1 gene=HORVU4Hr1G081660
K11(xgly / 162.0528)
Spectrum (double click to dock / undock)
R.IETPGPPYLAK[+162]QQCCGELANIPQQCR.C z=3,scan#=sample=1 period=1 cycle=865 experiment=18,scan time=12.8044
IETPGPPYLAKQQCCGELANIPQQCR
Intensity
m/z
m/z errors, tol=75.0 ppm
Protein Coverage (double click to ...
Protein Coverage
Sample name
Protein name
8:53 AM
20/08/2020

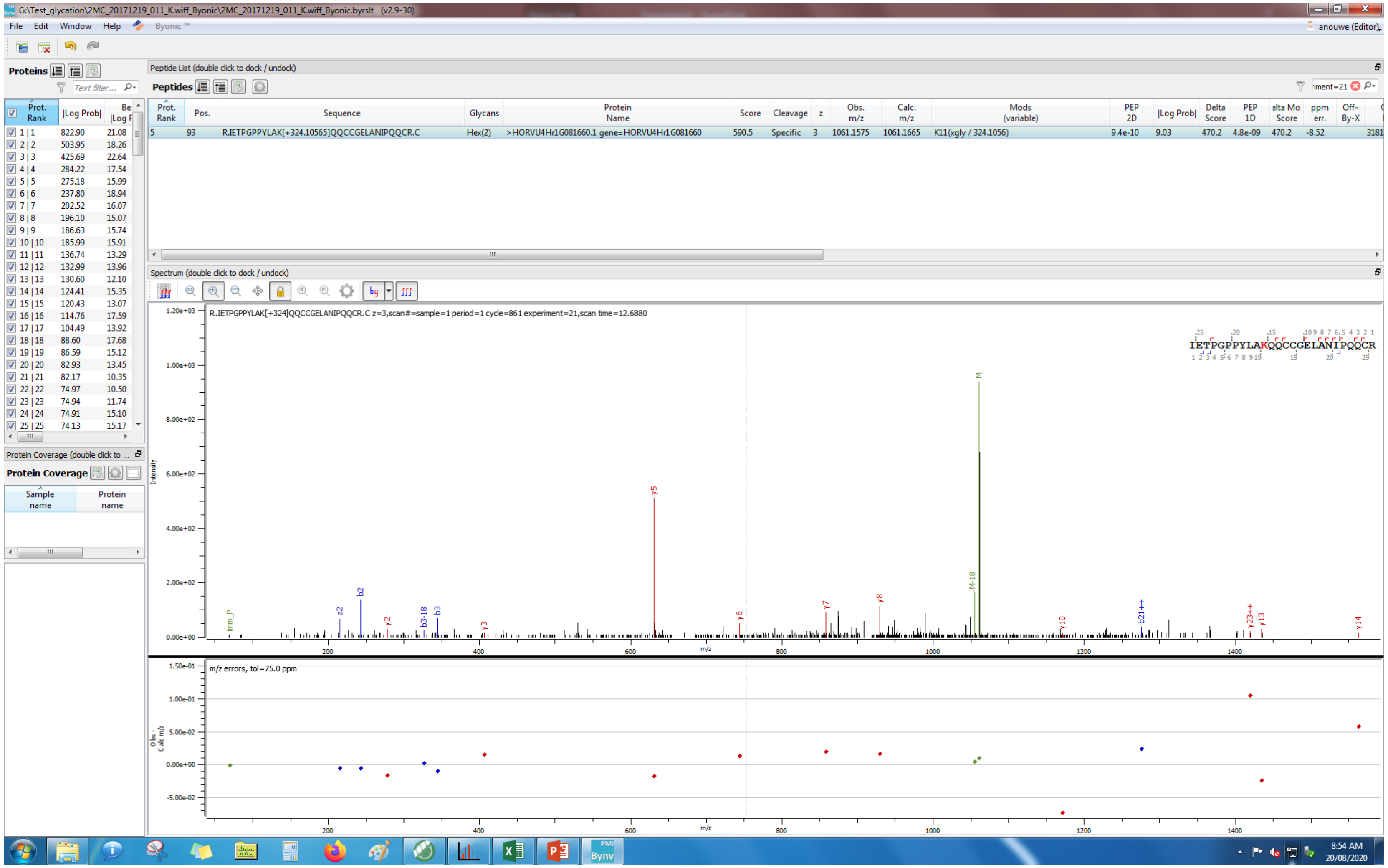

G:\Test_glycation\2MC_20171219_011_K.wiff_Byonic\2MC_20171219_011_K.wiff_Byonic.byrslt (v2.9-30)
File Edit Window Help Byonic
Proteins
Peptide List (double click to dock / undock)
Peptides
Prot. Rank | Pos. | Sequence | Glycans | Protein Name | Score | Cleavage | z | Obs. m/z | Calc. m/z | Mods (variable) | PEP 2D | |Log Prob| | Delta Score | PEP 1D | Delta Mod Score | ppm err. | Off-By-X
5 | 93 | R.IETPGPPYLAK[+324.10565]QQCCGELANIPQQCR.C | Hex(2) | >HORVU4Hr1G081660.1 gene=HORVU4Hr1G081660 | 590.5 | Specific | 3 | 1061.1575 | 1061.1665 | K11(xgly / 324.1056) | 9.4e-10 | 9.03 | 470.2 | 4.8e-09 | 470.2 | -8.52
Spectrum (double click to dock / undock)
R.IETPGPPYLAK[+324]QQCCGELANIPQQCR.C z=3,scan#=sample=1 period=1 cycle=861 experiment=21,scan time=12.6880
IETPGPPYLAKQQCCGELANIPQQCR
Intensity
m/z
m/z errors, tol=75.0 ppm
Protein Coverage (double click to ...
Protein Coverage
Sample name
Protein name
8:54 AM
20/08/2020

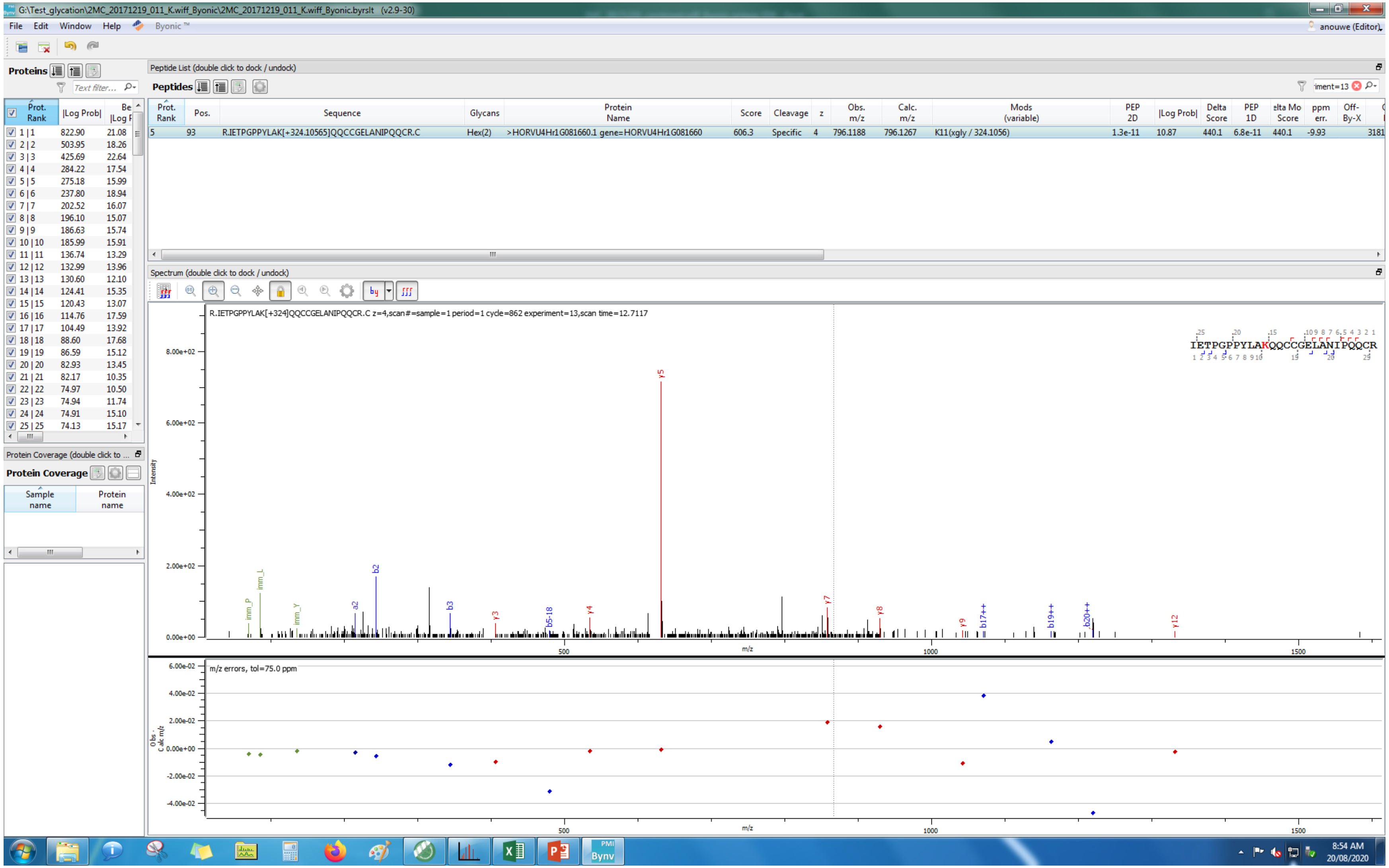
G:\Test_glycation\2MC_20171219_011_K.wiff_Byonic\2MC_20171219_011_K.wiff_Byonic.byrslt (v2.9-30)
File Edit Window Help Byonic™
anouwe (Editor)
Proteins
Peptide List (double click to dock / undock)
Peptides
Prot. Rank | Pos. | Sequence | Glycans | Protein Name | Score | Cleavage | z | Obs. m/z | Calc. m/z | Mods (variable) | PEP 2D | |Log Prob| | Delta Score | PEP 1D | elta Mo Score | ppm err. | Off-By-X
5 | 93 | R.IETPGPPYLAK[+324.10565]QQCCGELANIPQQCR.C | Hex(2) | >HORVU4Hr1G081660.1 gene=HORVU4Hr1G081660 | 606.3 | Specific | 4 | 796.1188 | 796.1267 | K11(xgly / 324.1056) | 1.3e-11 | 10.87 | 440.1 | 6.8e-11 | 440.1 | -9.93 | 3181
Prot. Rank | |Log Prob| | Be |Log F
1 / 1 822.90 21.08
2 / 2 503.95 18.26
3 / 3 425.69 22.64
4 / 4 284.22 17.54
5 / 5 275.18 15.99
6 / 6 237.80 18.94
7 / 7 202.52 16.07
8 / 8 196.10 15.07
9 / 9 186.63 15.74
10 / 10 185.99 15.91
11 / 11 136.74 13.29
12 / 12 132.99 13.96
13 / 13 130.60 12.10
14 / 14 124.41 15.35
15 / 15 120.43 13.07
16 / 16 114.76 17.59
17 / 17 104.49 13.92
18 / 18 88.60 17.68
19 / 19 86.59 15.12
20 / 20 82.93 13.45
21 / 21 82.17 10.35
22 / 22 74.97 10.50
23 / 23 74.94 11.74
24 / 24 74.91 15.10
25 / 25 74.13 15.17
Protein Coverage (double click to ...
Protein Coverage
Sample name
Protein name
Spectrum (double click to dock / undock)
R.IETPGPPYLAK[+324]QQCCGELANIPQQCR.C z=4,scan#=sample=1 period=1 cycle=862 experiment=13,scan time=12.7117
IETPGPPYLAKQQCCGELANIPQQCR
Intensity
m/z
y5
imm_P
imm_L
imm_Y
a2
b2
b3
y3
b5-18
y4
y7
y8
y9
b17++
b19++
b20++
y12
m/z errors, tol=75.0 ppm
Obs - Calc m/z
8:54 AM
20/08/2020

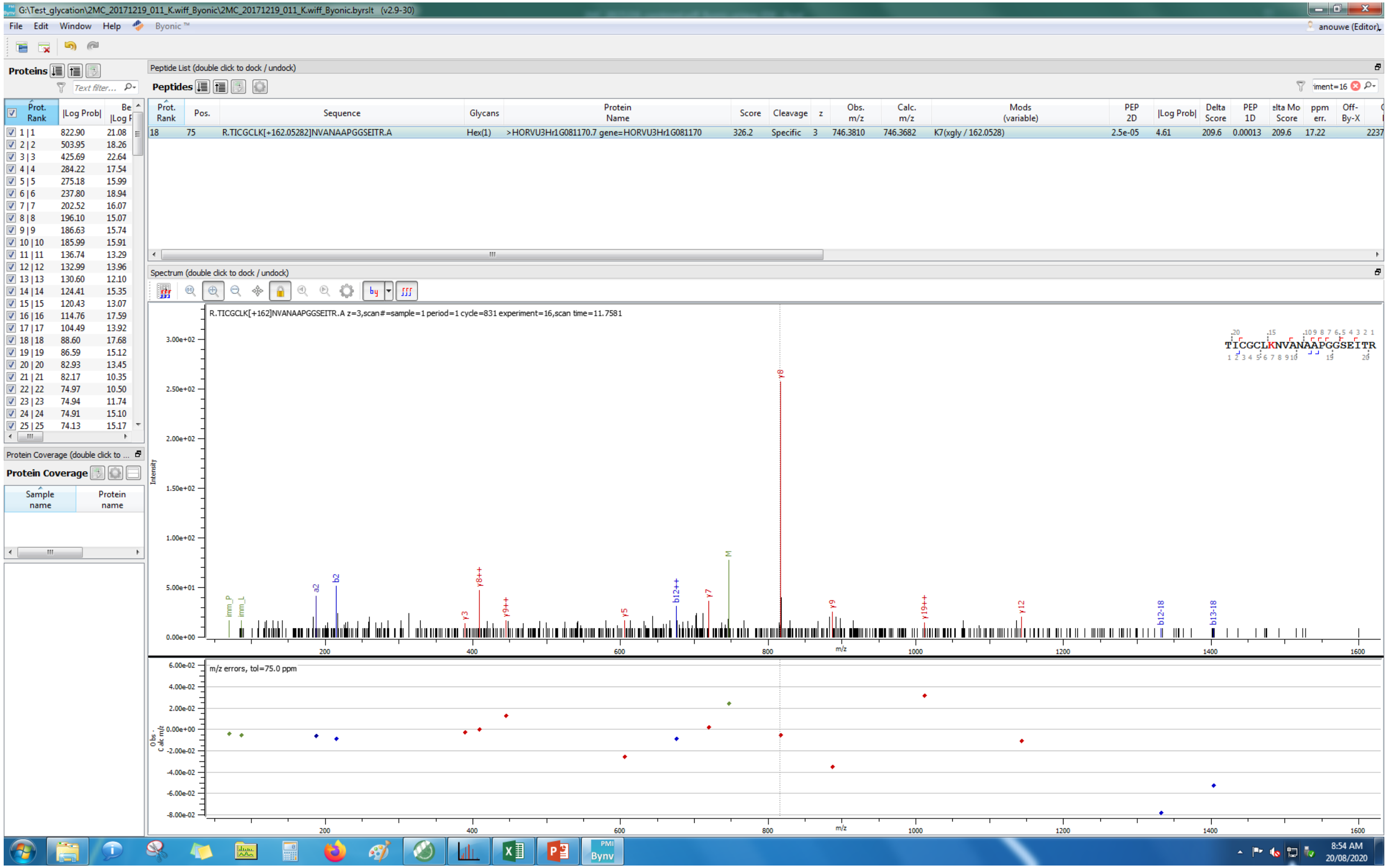

G:\Test_glycation\2MC_20171219_011_K.wiff_Byonic\2MC_20171219_011_K.wiff_Byonic.byrslt (v2.9-30)
File Edit Window Help Byonic™
Proteins
Peptide List (double click to dock / undock)
Peptides
18 75 R.TICGCLK[+162.05282]NVANAAPGGSEITR.A Hex(1) >HORVU3Hr1G081170.7 gene=HORVU3Hr1G081170 326.2 Specific 3 746.3810 746.3682 K7(xgly / 162.0528) 2.5e-05 4.61 209.6 0.00013 209.6 17.22
Spectrum (double click to dock / undock)
R.TICGCLK[+162]NVANAAPGGSEITR.A z=3,scan#=sample=1 period=1 cycle=831 experiment=16,scan time=11.7581
TICGCLKNVANAAPGGSEITR
Intensity
m/z
m/z errors, tol=75.0 ppm
Protein Coverage
Sample name
Protein name
8:54 AM
20/08/2020

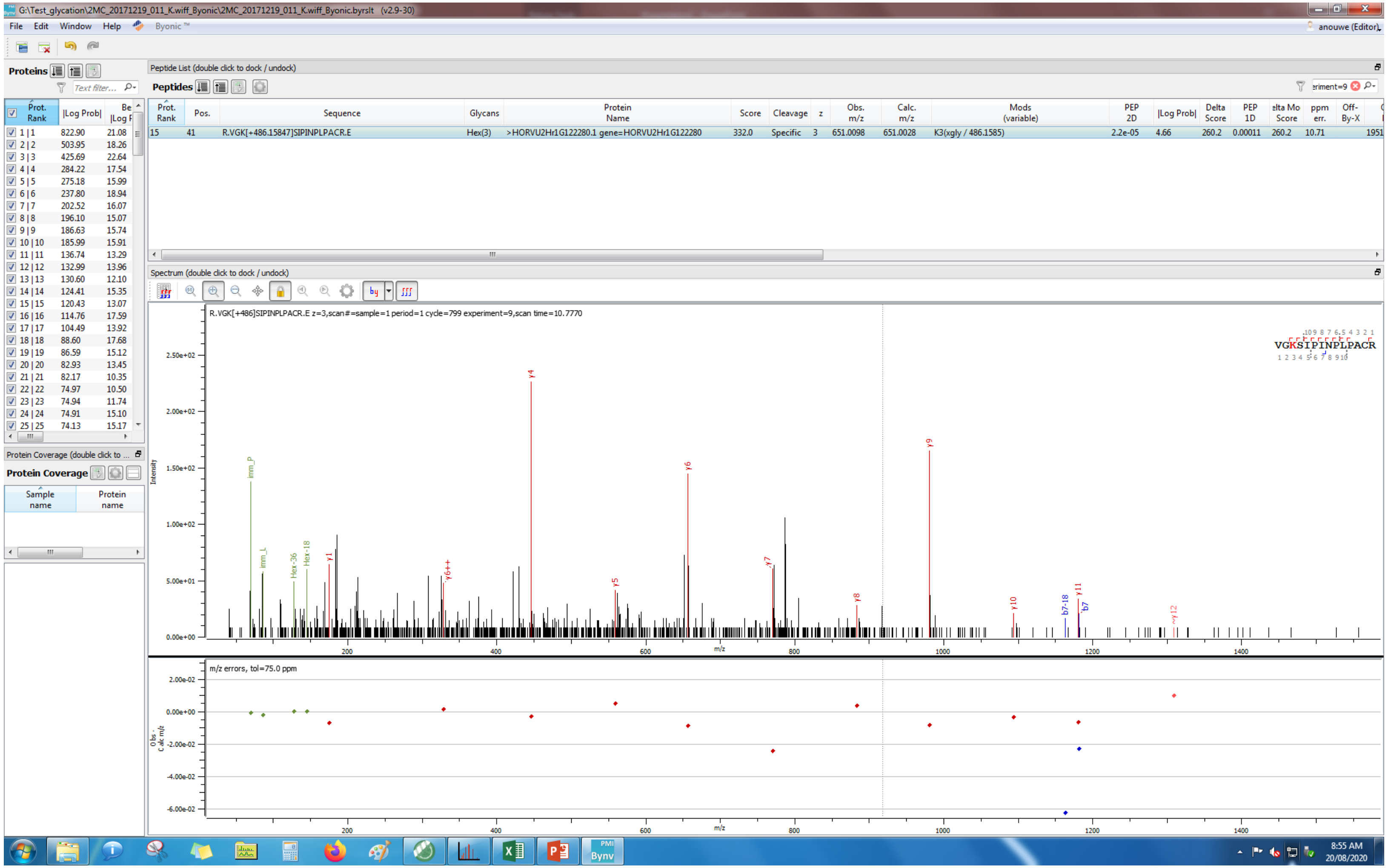
G:\Test_glycation\2MC_20171219_011_K.wiff_Byonic\2MC_20171219_011_K.wiff_Byonic.byrslt (v2.9-30)
File Edit Window Help Byonic
Proteins
Peptide List (double click to dock / undock)
Peptides
15 41 R.VGK[+486.15847]SIPINPLPACR.E Hex(3) >HORVU2Hr1G122280.1 gene=HORVU2Hr1G122280 332.0 Specific 3 651.0098 651.0028 K3(xgly / 486.1585) 2.2e-05 4.66 260.2 0.00011 260.2 10.71
Spectrum (double click to dock / undock)
R.VGK[+486]SIPINPLPACR.E z=3,scan#=sample=1 period=1 cycle=799 experiment=9,scan time=10.7770
VGKSIPINPLPACR
m/z errors, tol=75.0 ppm
Protein Coverage
8:55 AM
20/08/2020

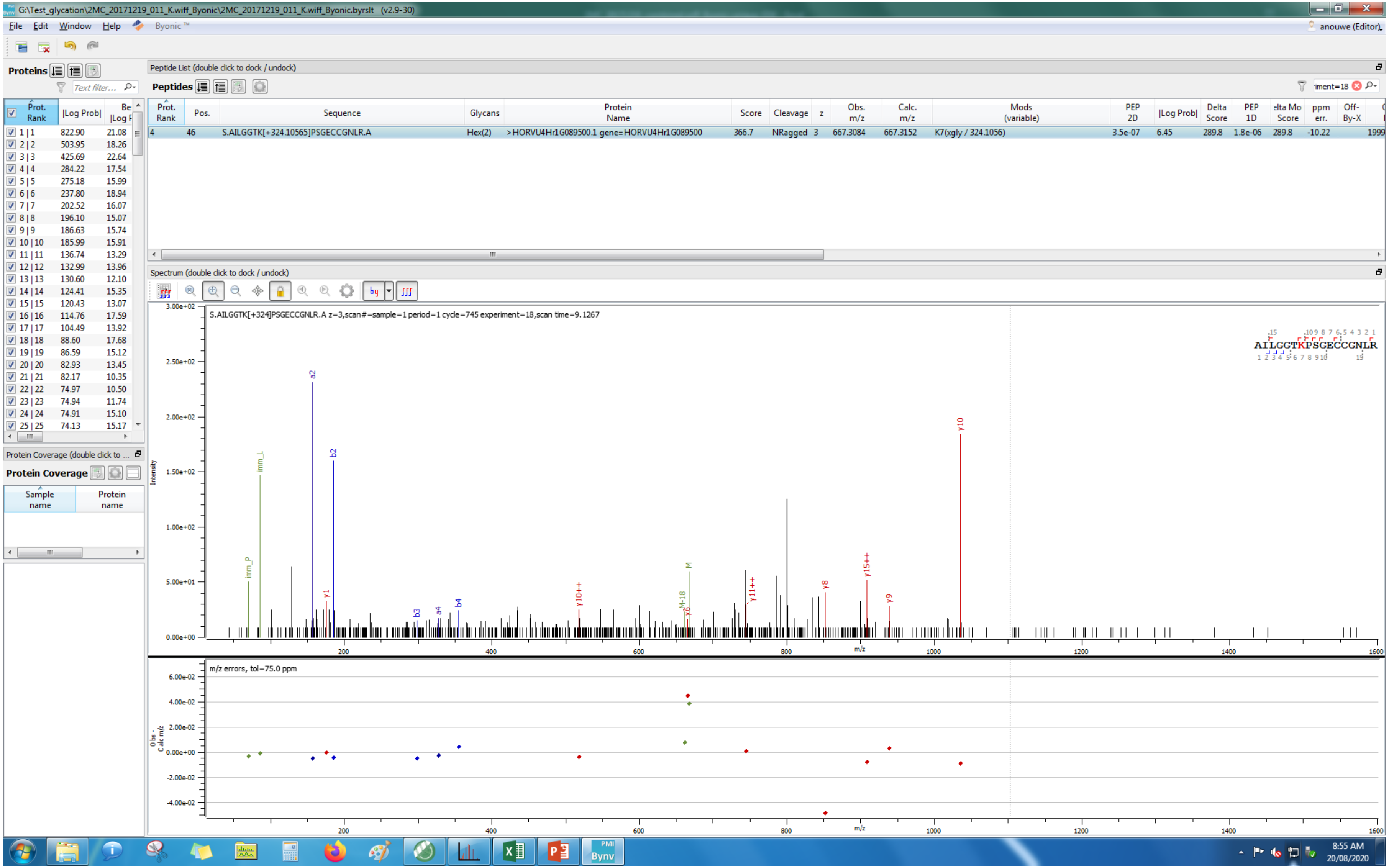
G:\Test_glycation\2MC_20171219_011_K.wiff_Byonic\2MC_20171219_011_K.wiff_Byonic.byrslt (v2.9-30)
File Edit Window Help Byonic ™
anouwe (Editor)
Proteins
Peptide List (double click to dock / undock)
Peptides
Prot. Rank | Pos. | Sequence | Glycans | Protein Name | Score | Cleavage | z | Obs. m/z | Calc. m/z | Mods (variable) | PEP 2D | |Log Prob| | Delta Score | PEP 1D | elta Mo Score | ppm err. | Off-By-X
4 | 46 | S.AILGGTK[+324.10565]PSGECCGNLR.A | Hex(2) | >HORVU4Hr1G089500.1 gene=HORVU4Hr1G089500 | 366.7 | NRagged | 3 | 667.3084 | 667.3152 | K7(xgly / 324.1056) | 3.5e-07 | 6.45 | 289.8 | 1.8e-06 | 289.8 | -10.22 | 1999
Spectrum (double click to dock / undock)
S.AILGGTK[+324]PSGECCGNLR.A z=3,scan#=sample=1 period=1 cycle=745 experiment=18,scan time=9.1267
AILGGTKPSGECCGNLR
Intensity
m/z
m/z errors, tol=75.0 ppm
Protein Coverage
Sample name
Protein name
8:55 AM
20/08/2020

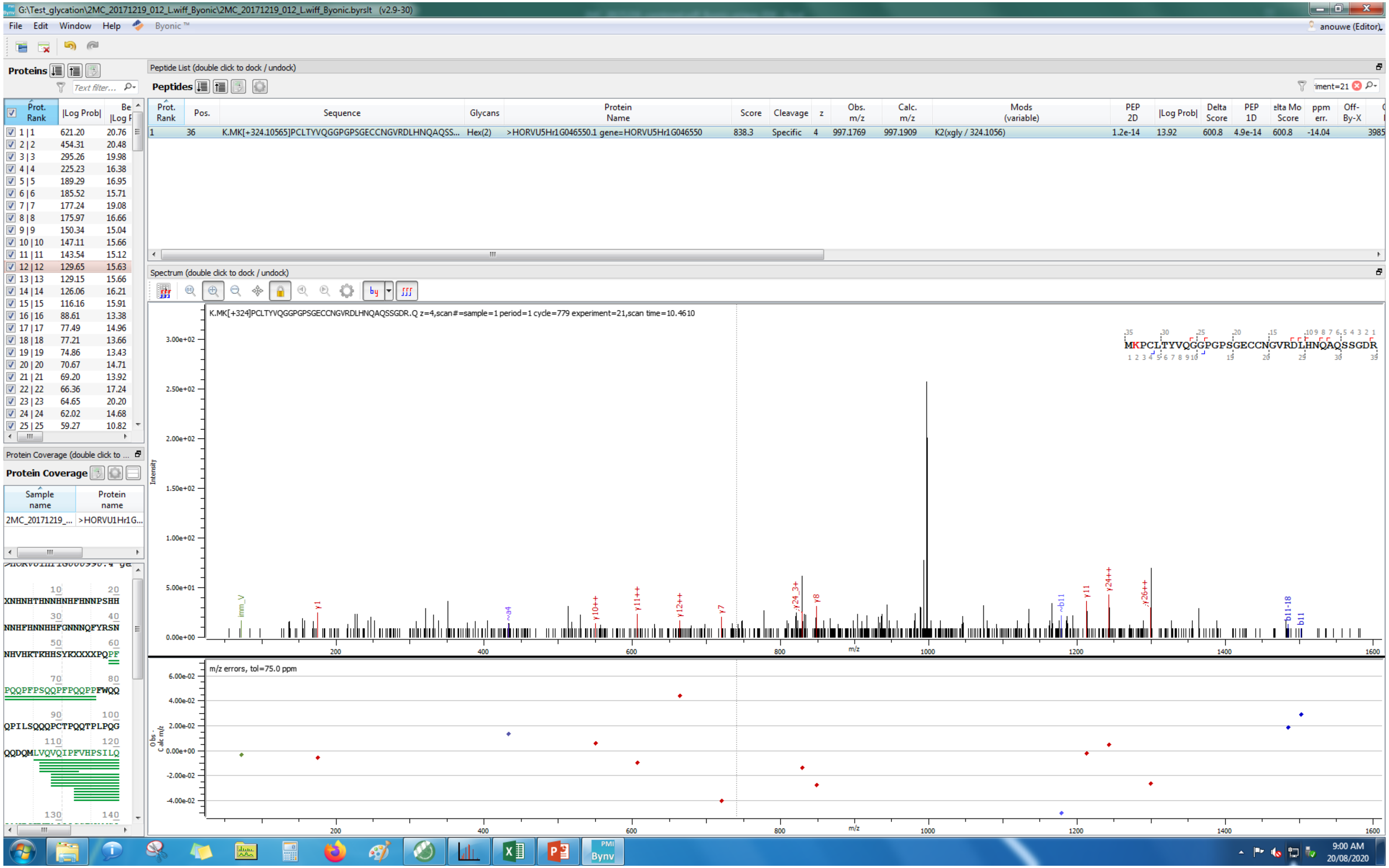
G:\Test_glycation\2MC_20171219_012_L.wiff_Byonic\2MC_20171219_012_L.wiff_Byonic.byrslt (v2.9-30)
File Edit Window Help Byonic ™
anouwe (Editor)
Proteins
Peptide List (double click to dock / undock)
Peptides
Prot. Rank | Pos. | Sequence | Glycans | Protein Name | Score | Cleavage | z | Obs. m/z | Calc. m/z | Mods (variable) | PEP 2D | |Log Prob| | Delta Score | PEP 1D | Delta Mod Score | ppm err. | Off-By-X
1 | 36 | K.MK[+324.10565]PCLTYVQGGPGPSGECCNGVRDLHNQAQSS... | Hex(2) | >HORVU5Hr1G046550.1 gene=HORVU5Hr1G046550 | 838.3 | Specific | 4 | 997.1769 | 997.1909 | K2(xgly / 324.1056) | 1.2e-14 | 13.92 | 600.8 | 4.9e-14 | 600.8 | -14.04
Spectrum (double click to dock / undock)
K.MK[+324]PCLTYVQGGPGPSGECCNGVRDLHNQAQSSGDR.Q z=4,scan#=sample=1 period=1 cycle=779 experiment=21,scan time=10.4610
MKPCLTYVQGGPGPSGECCNGVRDLHNQAQSSGDR
Intensity
m/z
m/z errors, tol=75.0 ppm
Protein Coverage (double click to ...
Protein Coverage
Sample name | Protein name
2MC_20171219_... | >HORVU1Hr1G...
XNHNHTHNNHNHFHNNPSHH
NNHFHNNHHFGNNNQFYRSN
NHVHKTKHHSYKXXXXXPQPF
PQQPFPSQQPFPQQPPFWQQ
QPILSQQQPCTPQQTPLPQG
QQDQMLVQVQIPFVHPSILQ
9:00 AM
20/08/2020

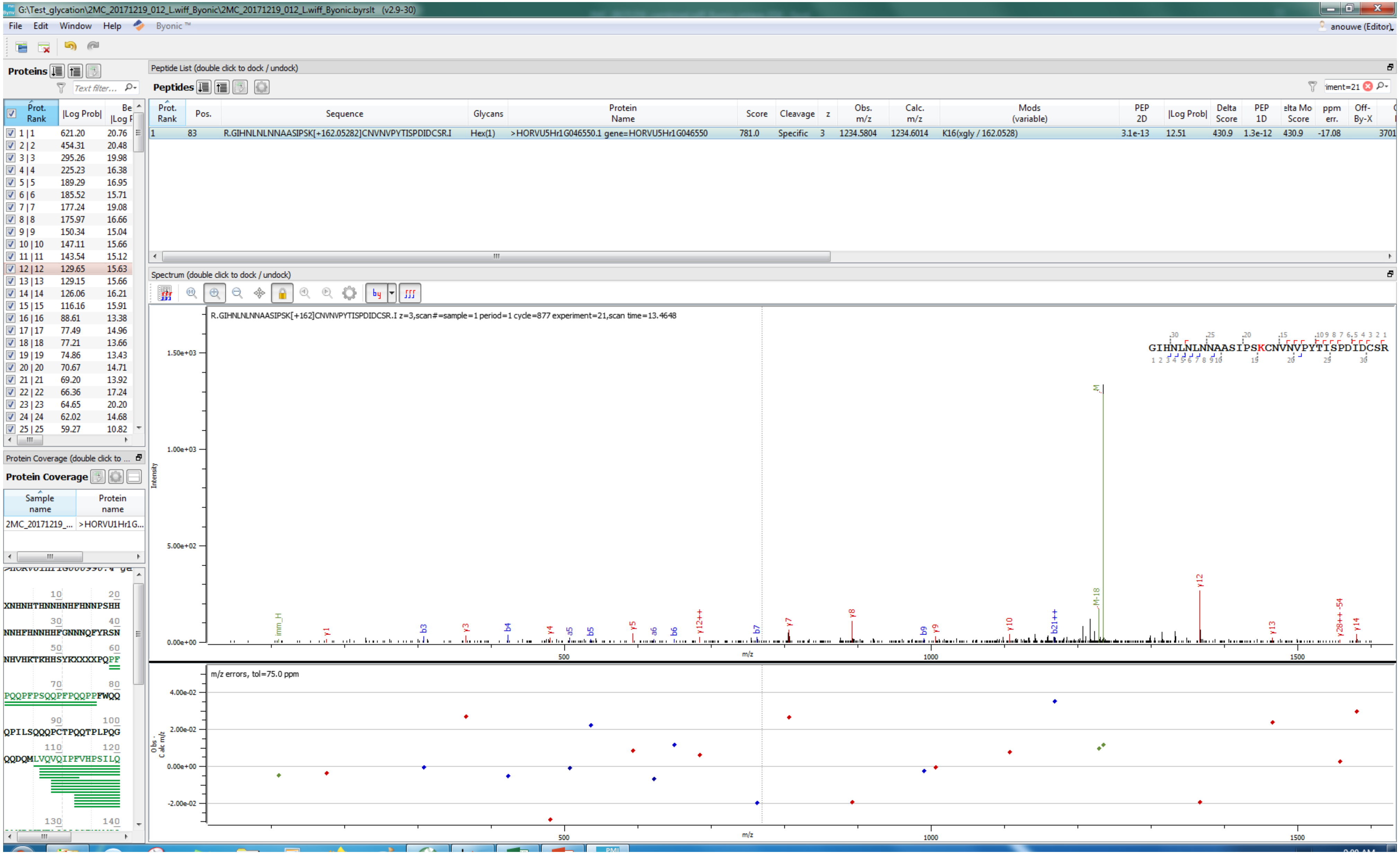
G:\Test_glycation\2MC_20171219_012_L.wiff_Byonic\2MC_20171219_012_L.wiff_Byonic.byrslt (v2.9-30)
File Edit Window Help Byonic ™
anouwe (Editor)
Proteins
Peptide List (double click to dock / undock)
Peptides
Prot. Rank | Pos. | Sequence | Glycans | Protein Name | Score | Cleavage | z | Obs. m/z | Calc. m/z | Mods (variable) | PEP 2D | |Log Prob| | Delta Score | PEP 1D | Delta Mod Score | ppm err. | Off-By-X
1 | 83 | R.GIHNLNLNNAASIPSK[+162.05282]CNVNVPYTISPDIDCSR.I | Hex(1) | >HORVU5Hr1G046550.1 gene=HORVU5Hr1G046550 | 781.0 | Specific | 3 | 1234.5804 | 1234.6014 | K16(xgly / 162.0528) | 3.1e-13 | 12.51 | 430.9 | 1.3e-12 | 430.9 | -17.08 | 3701
Spectrum (double click to dock / undock)
R.GIHNLNLNNAASIPSK[+162]CNVNVPYTISPDIDCSR.I z=3,scan#=sample=1 period=1 cycle=877 experiment=21,scan time=13.4648
GIHNLNLNNAASIPSKCNVNVPYTISPDIDCSR
Intensity
m/z
m/z errors, tol=75.0 ppm
Protein Coverage
Sample name | Protein name
2MC_20171219_... | >HORVU1Hr1G...
XNHNHTHNNHNHFHNNPSHH
NNHFHNNHHFGNNNQFYRSN
NHVHKTKHHSYKXXXXPQPF
PQQPFPSQQPFPQQPPFWQQ
QPILSQQQPCTPQQTPLPQG
QQDQMLVQVQIPFVHPSILQ

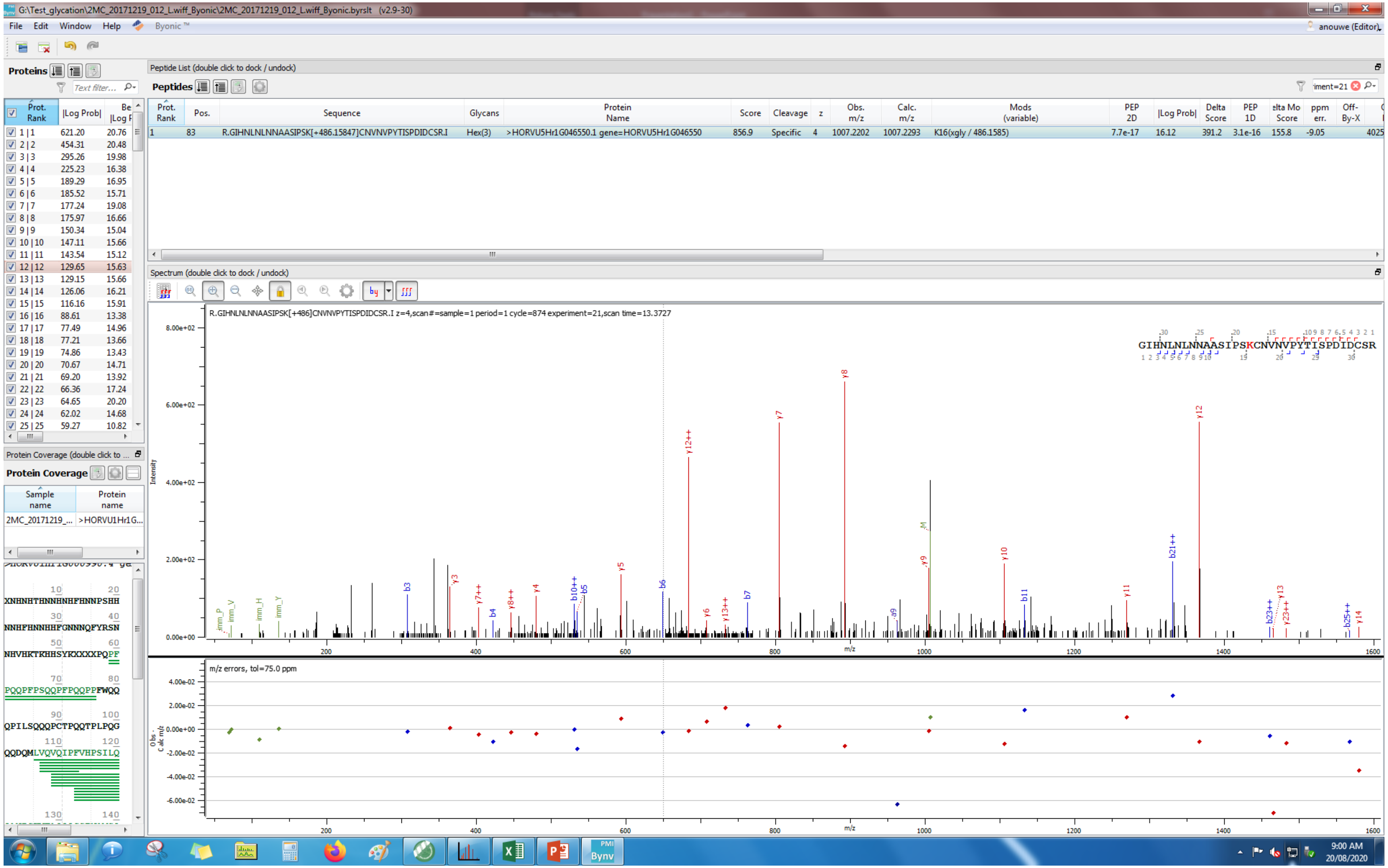
G:\Test_glycation\2MC_20171219_012_L.wiff_Byonic\2MC_20171219_012_L.wiff_Byonic.byrslt (v2.9-30)
File Edit Window Help Byonic ™
anouwe (Editor)
Proteins
Peptide List (double click to dock / undock)
Peptides
Prot. Rank | Pos. | Sequence | Glycans | Protein Name | Score | Cleavage | z | Obs. m/z | Calc. m/z | Mods (variable) | PEP 2D | |Log Prob| | Delta Score | PEP 1D | Delta Mod Score | ppm err. | Off-By-X
1 | 83 | R.GIHNLNLNNAASIPSK[+486.15847]CNVNVPYTISPDIDCSR.I | Hex(3) | >HORVU5Hr1G046550.1 gene=HORVU5Hr1G046550 | 856.9 | Specific | 4 | 1007.2202 | 1007.2293 | K16(xgly / 486.1585) | 7.7e-17 | 16.12 | 391.2 | 3.1e-16 | 155.8 | -9.05 | 4025
Spectrum (double click to dock / undock)
R.GIHNLNLNNAASIPSK[+486]CNVNVPYTISPDIDCSR.I z=4,scan#=sample=1 period=1 cycle=874 experiment=21,scan time=13.3727
GIHNLNLNNAASIPSKCNVNVPYTISPDIDCSR
Intensity
m/z
m/z errors, tol=75.0 ppm
Obs. - Calc m/z
Protein Coverage (double click to dock / undock)
Protein Coverage
Sample name | Protein name
2MC_20171219_... | >HORVU1Hr1G...
9:00 AM
20/08/2020

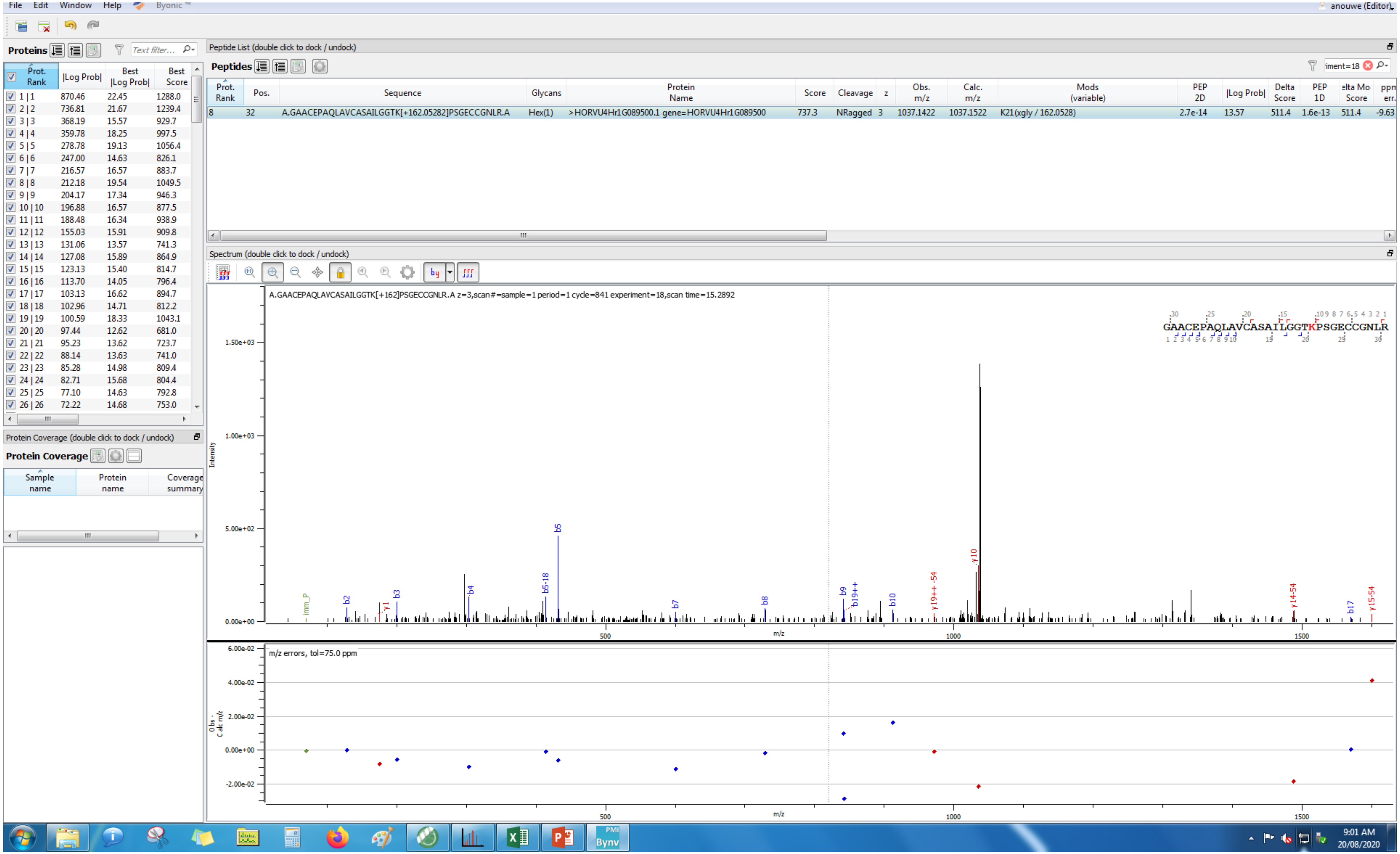

File Edit Window Help Byonic
anouwe (Editor)
Proteins
Peptide List (double click to dock / undock)
Peptides
Prot. Rank | Pos. | Sequence | Glycans | Protein Name | Score | Cleavage | z | Obs. m/z | Calc. m/z | Mods (variable) | PEP 2D | |Log Prob| | Delta Score | PEP 1D | Delta Mod Score | ppm err.
8 | 32 | A.GAACEPAQLAVCASAILGGTK[+162.05282]PSGECCGNLR.A | Hex(1) | >HORVU4Hr1G089500.1 gene=HORVU4Hr1G089500 | 737.3 | NRagged | 3 | 1037.1422 | 1037.1522 | K21(xgly / 162.0528) | 2.7e-14 | 13.57 | 511.4 | 1.6e-13 | 511.4 | -9.63
Spectrum (double click to dock / undock)
A.GAACEPAQLAVCASAILGGTK[+162]PSGECCGNLR.A z=3,scan#=sample=1 period=1 cycle=841 experiment=18,scan time=15.2892
GAACEPAQLAVCASAILGGTKPSGECCGNLR
Intensity
m/z
m/z errors, tol=75.0 ppm
Obs - Calc m/z
Protein Coverage (double click to dock / undock)
Protein Coverage
Sample name | Protein name | Coverage summary
Prot. Rank | |Log Prob| | Best |Log Prob| | Best Score
1 | 1 | 870.46 | 22.45 | 1288.0
2 | 2 | 736.81 | 21.67 | 1239.4
3 | 3 | 368.19 | 15.57 | 929.7
4 | 4 | 359.78 | 18.25 | 997.5
5 | 5 | 278.78 | 19.13 | 1056.4
6 | 6 | 247.00 | 14.63 | 826.1
7 | 7 | 216.57 | 16.57 | 883.7
8 | 8 | 212.18 | 19.54 | 1049.5
9 | 9 | 204.17 | 17.34 | 946.3
10 | 10 | 196.88 | 16.57 | 877.5
11 | 11 | 188.48 | 16.34 | 938.9
12 | 12 | 155.03 | 15.91 | 909.8
13 | 13 | 131.06 | 13.57 | 741.3
14 | 14 | 127.08 | 15.89 | 864.9
15 | 15 | 123.13 | 15.40 | 814.7
16 | 16 | 113.70 | 14.05 | 796.4
17 | 17 | 103.13 | 16.62 | 894.7
18 | 18 | 102.96 | 14.71 | 812.2
19 | 19 | 100.59 | 18.33 | 1043.1
20 | 20 | 97.44 | 12.62 | 681.0
21 | 21 | 95.23 | 13.62 | 723.7
22 | 22 | 88.14 | 13.63 | 741.0
23 | 23 | 85.28 | 14.98 | 809.4
24 | 24 | 82.71 | 15.68 | 804.4
25 | 25 | 77.10 | 14.63 | 792.8
26 | 26 | 72.22 | 14.68 | 753.0
9:01 AM
20/08/2020

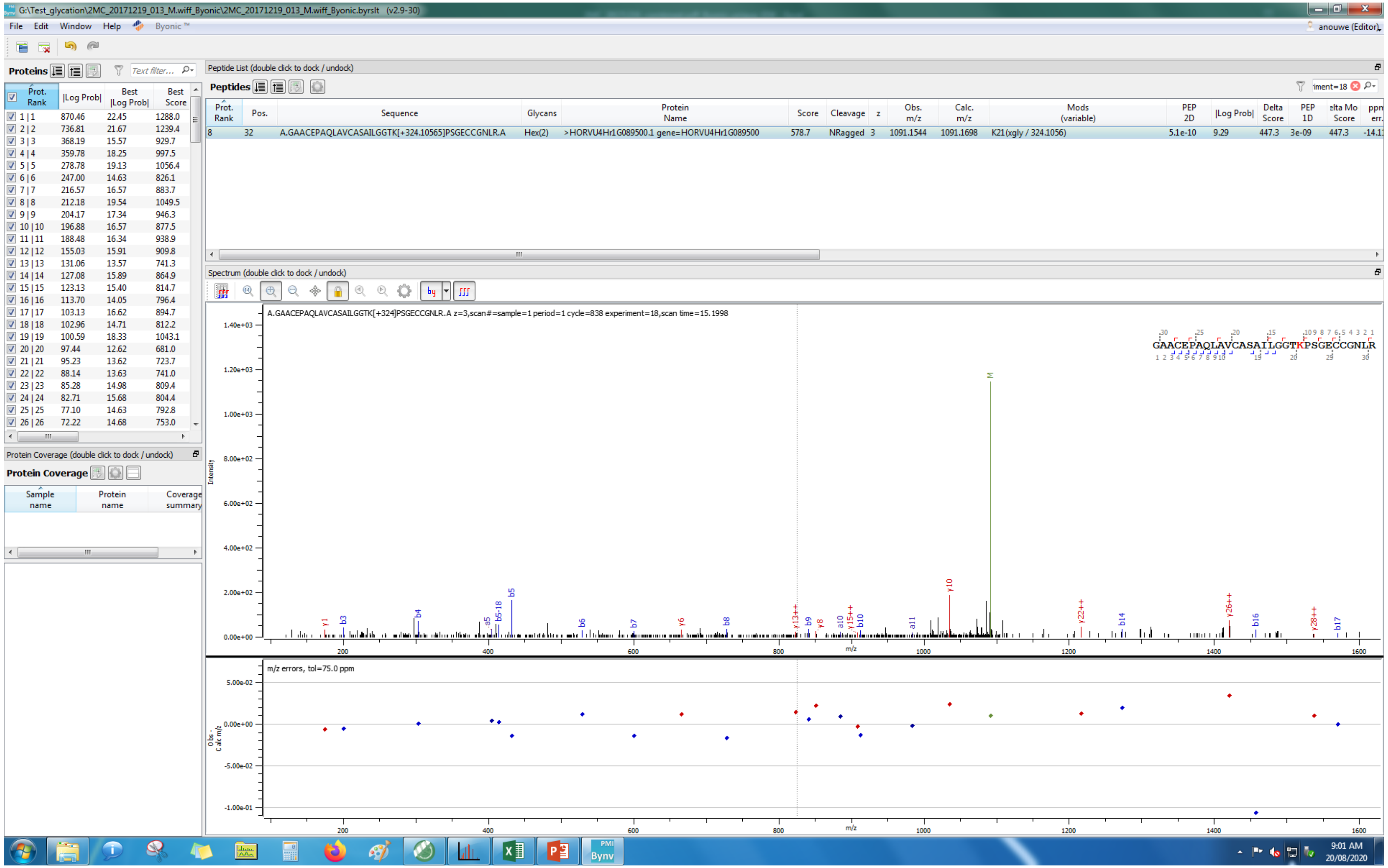
G:\Test_glycation\2MC_20171219_013_M.wiff_Byonic\2MC_20171219_013_M.wiff_Byonic.byrslt (v2.9-30)
File Edit Window Help Byonic™
anouwe (Editor)
Proteins
Peptide List (double click to dock / undock)
Peptides
Protein Coverage (double click to dock / undock)
Protein Coverage
Spectrum (double click to dock / undock)
A.GAACEPAQLAVCASAILGGTK[+324]PSGECCGNLR.A z=3,scan#=sample=1 period=1 cycle=838 experiment=18,scan time=15.1998
GAACEPAQLAVCASAILGGTKPSGECCGNLR
Intensity
m/z
m/z errors, tol=75.0 ppm
9:01 AM
20/08/2020

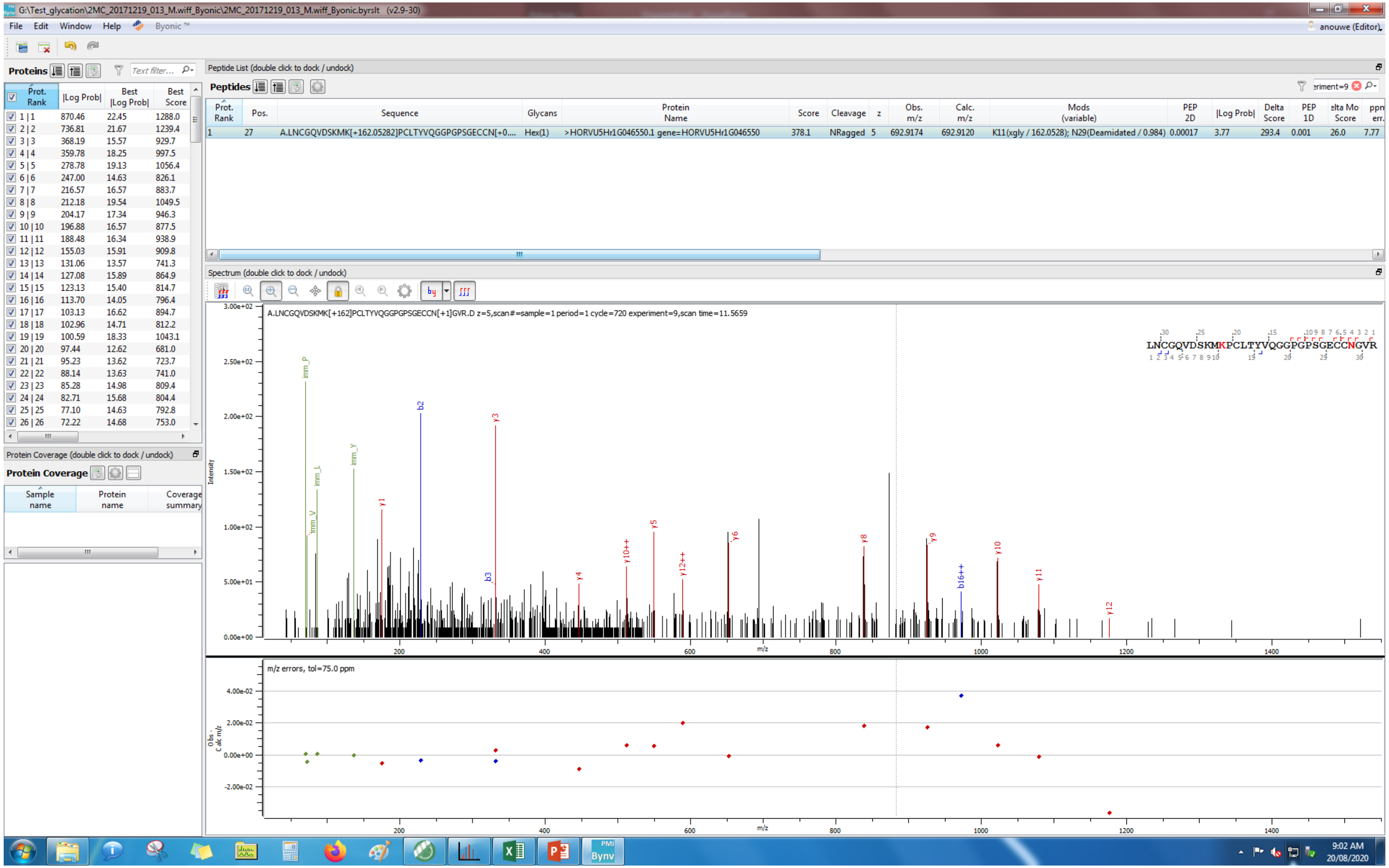
G:\Test_glycation\2MC_20171219_013_M.wiff_Byonic\2MC_20171219_013_M.wiff_Byonic.byrslt (v2.9-30)
File Edit Window Help Byonic™
anouwe (Editor)
Proteins
Peptide List (double click to dock / undock)
Peptides
Prot. Rank | Pos. | Sequence | Glycans | Protein Name | Score | Cleavage | z | Obs. m/z | Calc. m/z | Mods (variable) | PEP 2D | |Log Prob| | Delta Score | PEP 1D | Delta Mod Score | ppm err.
1 | 27 | A.LNCGQVDSKMK[+162.05282]PCLTYVQGGPGPSGECCN[+0.... | Hex(1) | >HORVU5Hr1G046550.1 gene=HORVU5Hr1G046550 | 378.1 | NRagged | 5 | 692.9174 | 692.9120 | K11(xgly / 162.0528); N29(Deamidated / 0.984) | 0.00017 | 3.77 | 293.4 | 0.001 | 26.0 | 7.77
Spectrum (double click to dock / undock)
A.LNCGQVDSKMK[+162]PCLTYVQGGPGPSGECCN[+1]GVR.D z=5,scan#=sample=1 period=1 cycle=720 experiment=9,scan time=11.5659
LNCGQVDSKMKPCLTYVQGGPGPSGECCNGVR
Intensity
m/z
m/z errors, tol=75.0 ppm
Protein Coverage (double click to dock / undock)
Protein Coverage
Sample name
Protein name
Coverage summary
9:02 AM
20/08/2020

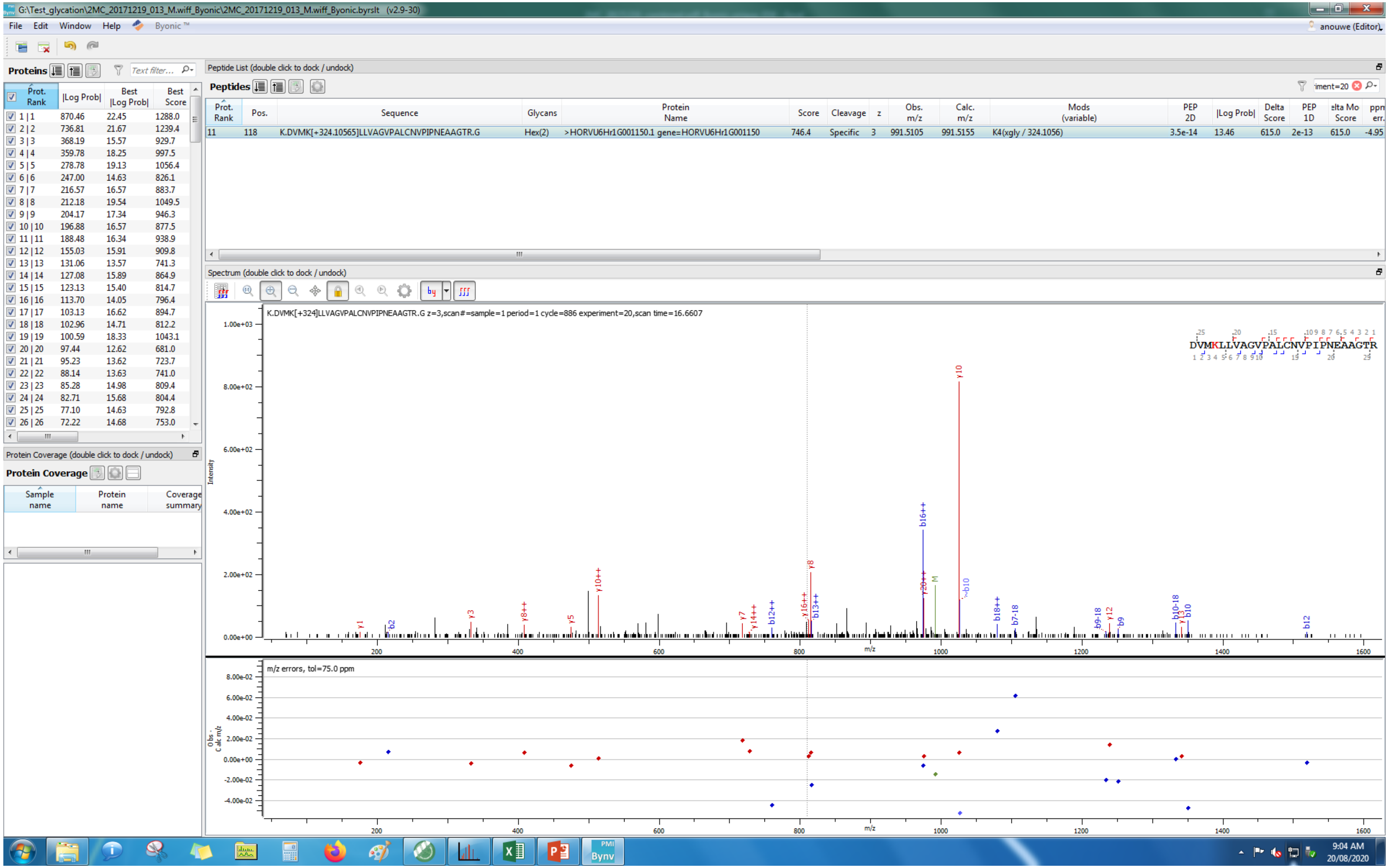
G:\Test_glycation\2MC_20171219_013_M.wiff_Byonic\2MC_20171219_013_M.wiff_Byonic.byrslt (v2.9-30)
File Edit Window Help Byonic ™
Proteins
Peptide List (double click to dock / undock)
Peptides
Prot. Rank | Pos. | Sequence | Glycans | Protein Name | Score | Cleavage | z | Obs. m/z | Calc. m/z | Mods (variable) | PEP 2D | |Log Prob| | Delta Score | PEP 1D | Delta Mod Score | ppm err.
11 | 118 | K.DVMK[+324.10565]LLVAGVPALCNVPIPNEAAGTR.G | Hex(2) | >HORVU6Hr1G001150.1 gene=HORVU6Hr1G001150 | 746.4 | Specific | 3 | 991.5105 | 991.5155 | K4(xgly / 324.1056) | 3.5e-14 | 13.46 | 615.0 | 2e-13 | 615.0 | -4.95
Spectrum (double click to dock / undock)
K.DVMK[+324]LLVAGVPALCNVPIPNEAAGTR.G z=3,scan#=sample=1 period=1 cycle=886 experiment=20,scan time=16.6607
DVMKLLVAGVPALCNVPIPNEAAGTR
Intensity
m/z
m/z errors, tol=75.0 ppm
Obs - Calc m/z
Protein Coverage (double click to dock / undock)
Protein Coverage
Sample name
Protein name
Coverage summary
9:04 AM
20/08/2020

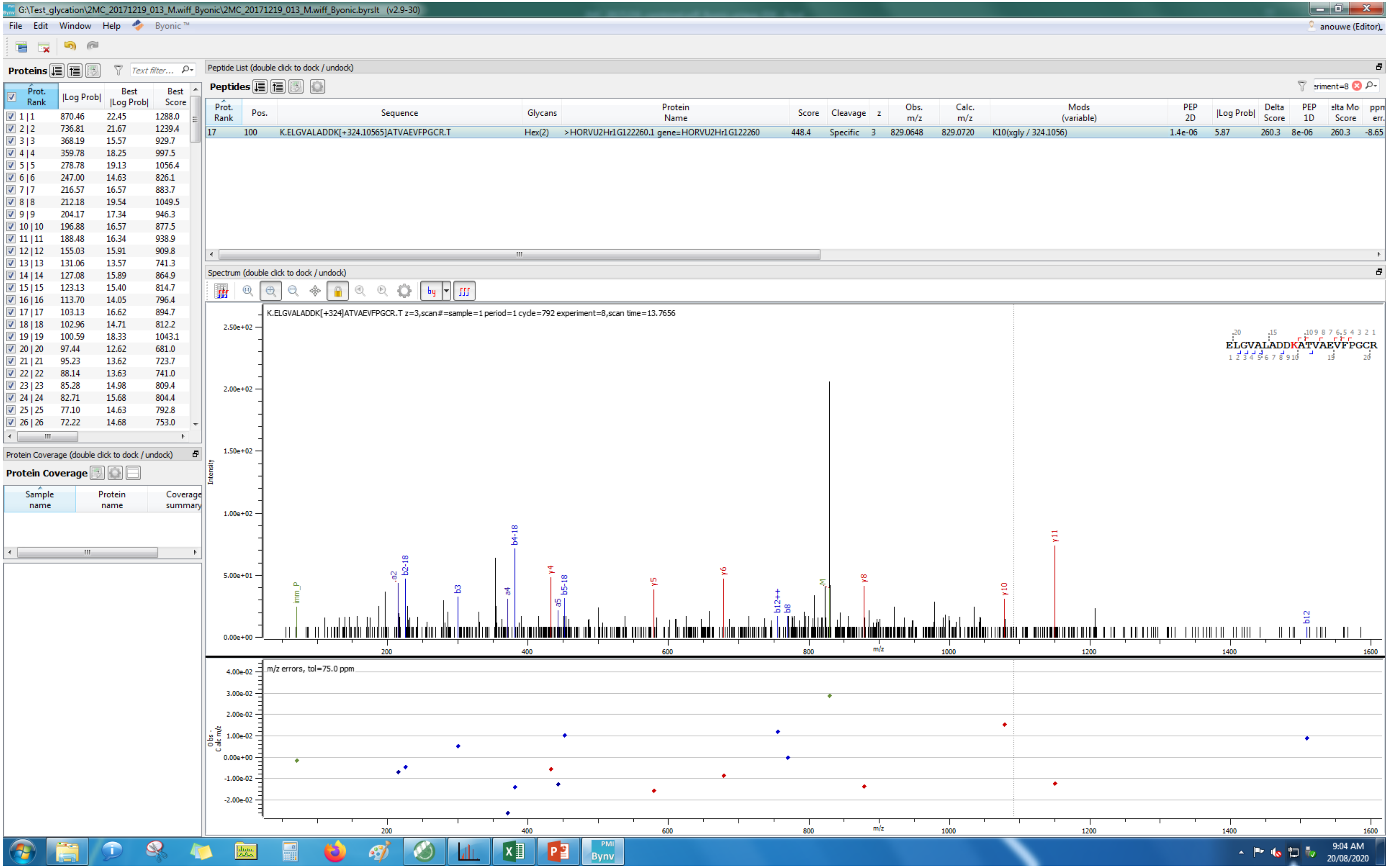

G:\Test_glycation\2MC_20171219_013_M.wiff_Byonic\2MC_20171219_013_M.wiff_Byonic.byrslt (v2.9-30)
File Edit Window Help Byonic™
anouwe (Editor)
Proteins
Peptide List (double click to dock / undock)
Peptides
Prot. Rank | Pos. | Sequence | Glycans | Protein Name | Score | Cleavage | z | Obs. m/z | Calc. m/z | Mods (variable) | PEP 2D | |Log Prob| | Delta Score | PEP 1D | Delta Mod Score | ppm err.
17 | 100 | K.ELGVALADDK[+324.10565]ATVAEVFPGCR.T | Hex(2) | >HORVU2Hr1G122260.1 gene=HORVU2Hr1G122260 | 448.4 | Specific | 3 | 829.0648 | 829.0720 | K10(xgly / 324.1056) | 1.4e-06 | 5.87 | 260.3 | 8e-06 | 260.3 | -8.65
Spectrum (double click to dock / undock)
K.ELGVALADDK[+324]ATVAEVFPGCR.T z=3,scan#=sample=1 period=1 cycle=792 experiment=8,scan time=13.7656
ELGVALADDKATVAEVFPGCR
Intensity
m/z
m/z errors, tol=75.0 ppm
Obs-Calc m/z
Protein Coverage (double click to dock / undock)
Protein Coverage
Sample name
Protein name
Coverage summary
9:04 AM
20/08/2020

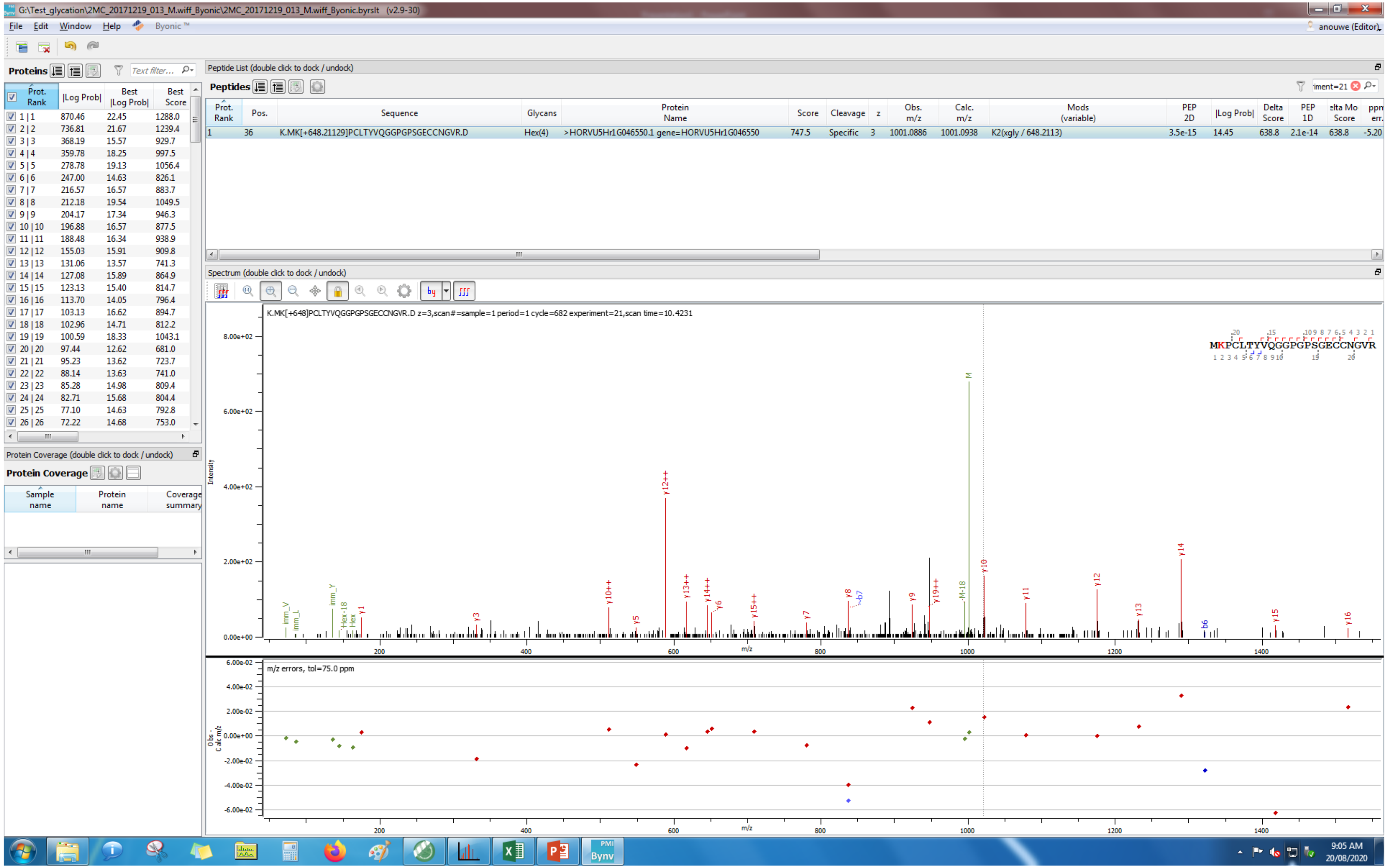
G:\Test_glycation\2MC_20171219_013_M.wiff_Byonic\2MC_20171219_013_M.wiff_Byonic.byrslt (v2.9-30)
File Edit Window Help Byonic
anouwe (Editor)
Proteins
Prot. Rank |Log Prob| Best |Log Prob| Best Score
1 | 1 870.46 22.45 1288.0
2 | 2 736.81 21.67 1239.4
3 | 3 368.19 15.57 929.7
4 | 4 359.78 18.25 997.5
5 | 5 278.78 19.13 1056.4
6 | 6 247.00 14.63 826.1
7 | 7 216.57 16.57 883.7
8 | 8 212.18 19.54 1049.5
9 | 9 204.17 17.34 946.3
10 | 10 196.88 16.57 877.5
11 | 11 188.48 16.34 938.9
12 | 12 155.03 15.91 909.8
13 | 13 131.06 13.57 741.3
14 | 14 127.08 15.89 864.9
15 | 15 123.13 15.40 814.7
16 | 16 113.70 14.05 796.4
17 | 17 103.13 16.62 894.7
18 | 18 102.96 14.71 812.2
19 | 19 100.59 18.33 1043.1
20 | 20 97.44 12.62 681.0
21 | 21 95.23 13.62 723.7
22 | 22 88.14 13.63 741.0
23 | 23 85.28 14.98 809.4
24 | 24 82.71 15.68 804.4
25 | 25 77.10 14.63 792.8
26 | 26 72.22 14.68 753.0
Protein Coverage (double click to dock / undock)
Protein Coverage
Sample name Protein name Coverage summary
Peptide List (double click to dock / undock)
Peptides
Prot. Rank Pos. Sequence Glycans Protein Name Score Cleavage z Obs. m/z Calc. m/z Mods (variable) PEP 2D |Log Prob| Delta Score PEP 1D elta Mo Score ppm err.
1 36 K.MK[+648.21129]PCLTYVQGGPGPSGECCNGVR.D Hex(4) >HORVU5Hr1G046550.1 gene=HORVU5Hr1G046550 747.5 Specific 3 1001.0886 1001.0938 K2(xgly / 648.2113) 3.5e-15 14.45 638.8 2.1e-14 638.8 -5.20
Spectrum (double click to dock / undock)
K.MK[+648]PCLTYVQGGPGPSGECCNGVR.D z=3,scan#=sample=1 period=1 cycle=682 experiment=21,scan time=10.4231
MKPCLTYVQGGPGPSGECCNGVR
Intensity
m/z
m/z errors, tol=75.0 ppm
9:05 AM
20/08/2020

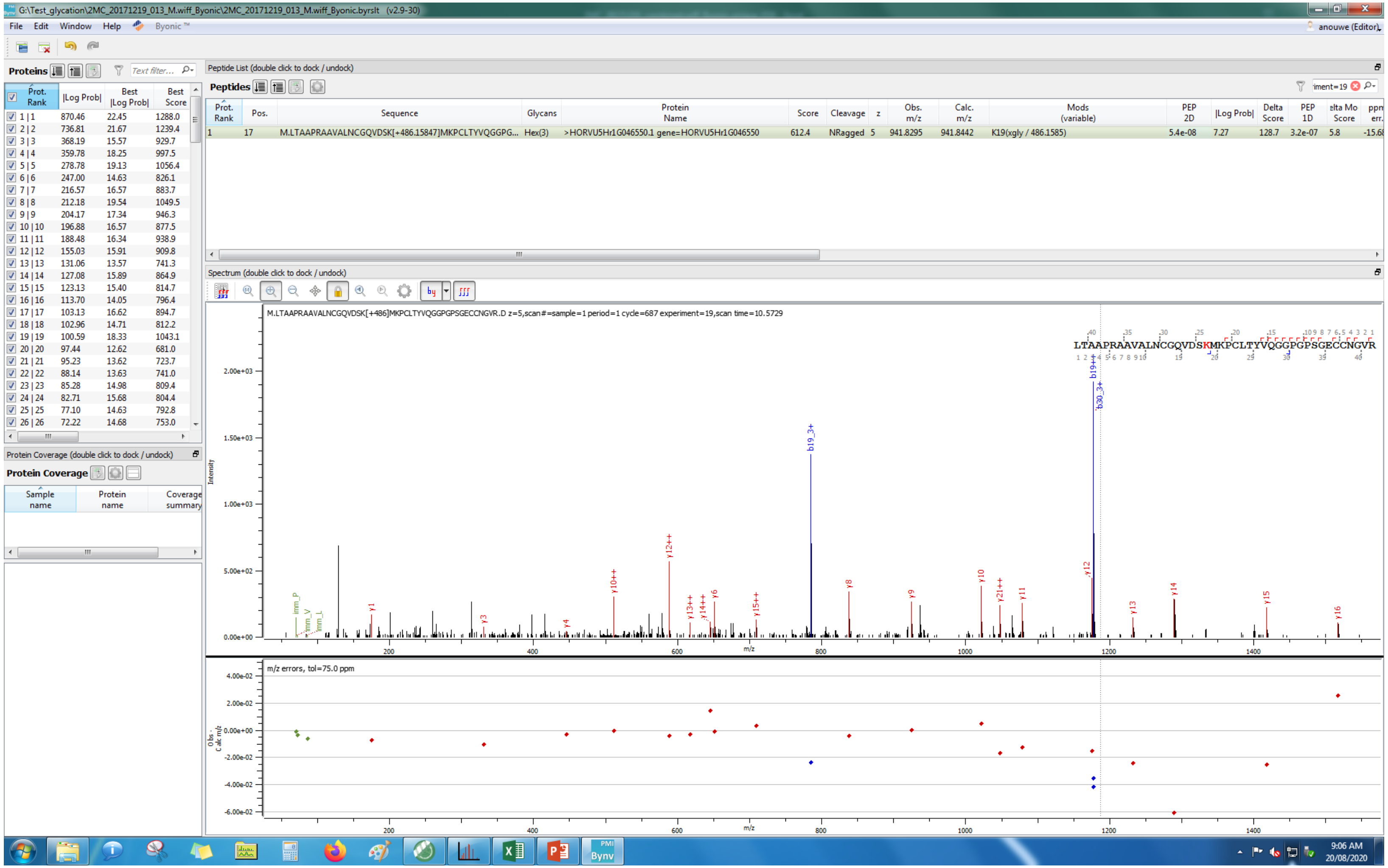
G:\Test_glycation\2MC_20171219_013_M.wiff_Byonic\2MC_20171219_013_M.wiff_Byonic.byrslt (v2.9-30)
File Edit Window Help Byonic ™
anouwe (Editor)
Proteins
Text filter...
Prot. Rank | |Log Prob| | Best |Log Prob| | Best Score
1 | 1 870.46 22.45 1288.0
2 | 2 736.81 21.67 1239.4
3 | 3 368.19 15.57 929.7
4 | 4 359.78 18.25 997.5
5 | 5 278.78 19.13 1056.4
6 | 6 247.00 14.63 826.1
7 | 7 216.57 16.57 883.7
8 | 8 212.18 19.54 1049.5
9 | 9 204.17 17.34 946.3
10 | 10 196.88 16.57 877.5
11 | 11 188.48 16.34 938.9
12 | 12 155.03 15.91 909.8
13 | 13 131.06 13.57 741.3
14 | 14 127.08 15.89 864.9
15 | 15 123.13 15.40 814.7
16 | 16 113.70 14.05 796.4
17 | 17 103.13 16.62 894.7
18 | 18 102.96 14.71 812.2
19 | 19 100.59 18.33 1043.1
20 | 20 97.44 12.62 681.0
21 | 21 95.23 13.62 723.7
22 | 22 88.14 13.63 741.0
23 | 23 85.28 14.98 809.4
24 | 24 82.71 15.68 804.4
25 | 25 77.10 14.63 792.8
26 | 26 72.22 14.68 753.0
Protein Coverage (double click to dock / undock)
Protein Coverage
Sample name | Protein name | Coverage summary
Peptide List (double click to dock / undock)
Peptides
iment=19
Prot. Rank | Pos. | Sequence | Glycans | Protein Name | Score | Cleavage | z | Obs. m/z | Calc. m/z | Mods (variable) | PEP 2D | |Log Prob| | Delta Score | PEP 1D | elta Mo Score | ppm err.
1 17 M.LTAAPRAAVALNCGQVDSK[+486.15847]MKPCLTYVQGGPG... Hex(3) >HORVU5Hr1G046550.1 gene=HORVU5Hr1G046550 612.4 NRagged 5 941.8295 941.8442 K19(xgly / 486.1585) 5.4e-08 7.27 128.7 3.2e-07 5.8 -15.68
Spectrum (double click to dock / undock)
M.LTAAPRAAVALNCGQVDSK[+486]MKPCLTYVQGGPGPSGECCNGVR.D z=5,scan#=sample=1 period=1 cycle=687 experiment=19,scan time=10.5729
LTAAPRAAVALNCGQVDSKMKPCLTYVQGGPGPSGECCNGVR
Intensity
m/z
b19_3+
b30_3+
y12++
y10++
y8
y9
y10
y21++
y11
y12
y13
y14
y15
y16
y1
y3
y4
y6
y13++
y14++
y15++
m/z errors, tol=75.0 ppm
Obs - Calc m/z
9:06 AM
20/08/2020

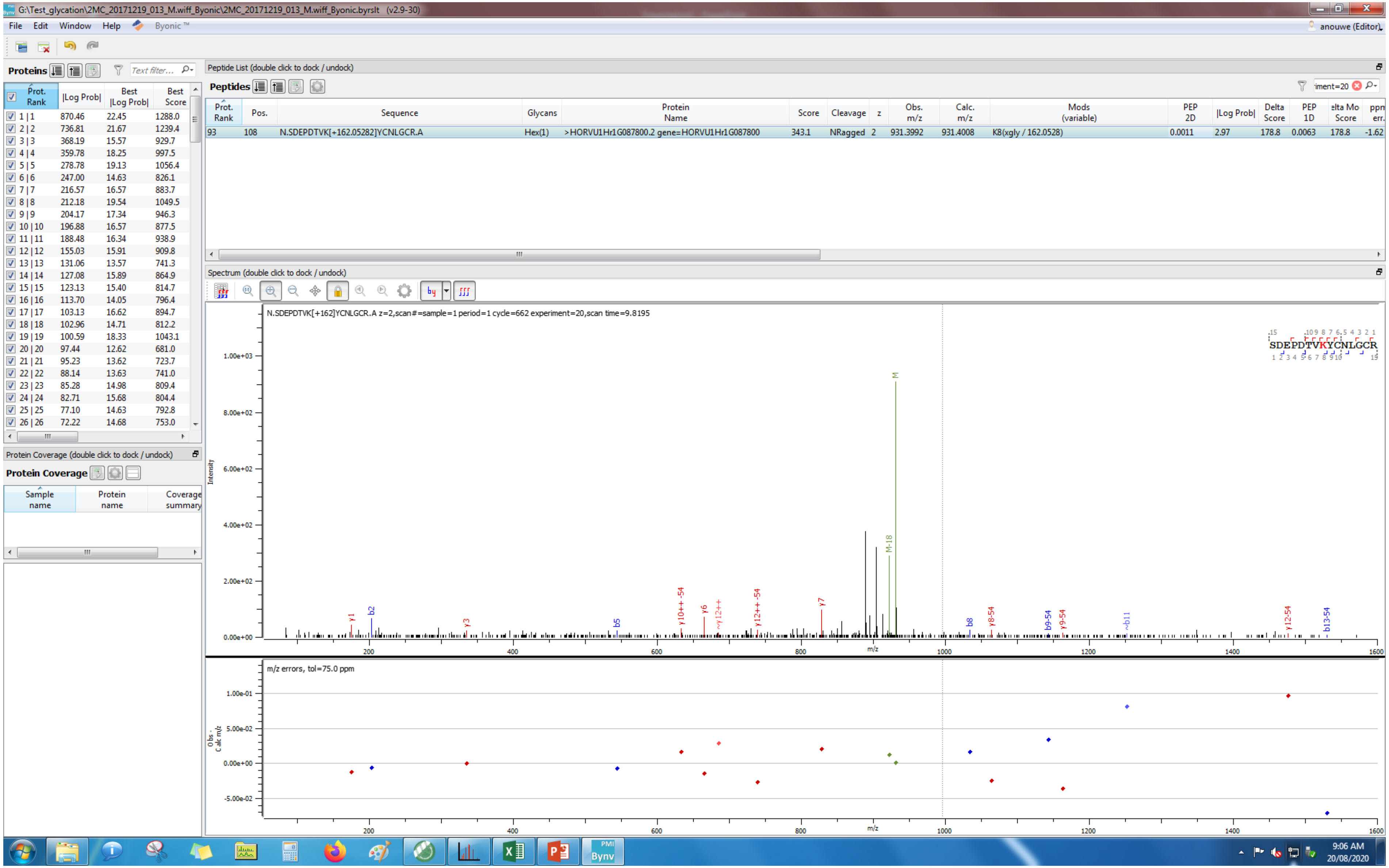
G:\Test_glycation\2MC_20171219_013_M.wiff_Byonic\2MC_20171219_013_M.wiff_Byonic.byrslt (v2.9-30)
File Edit Window Help Byonic
anouwe (Editor)
Proteins
Peptide List (double click to dock / undock)
Peptides
Prot. Rank | Pos. | Sequence | Glycans | Protein Name | Score | Cleavage | z | Obs. m/z | Calc. m/z | Mods (variable) | PEP 2D | |Log Prob| | Delta Score | PEP 1D | Delta Mod Score | ppm err.
93 | 108 | N.SDEPDTVK[+162.05282]YCNLGCR.A | Hex(1) | >HORVU1Hr1G087800.2 gene=HORVU1Hr1G087800 | 343.1 | NRagged | 2 | 931.3992 | 931.4008 | K8(xgly / 162.0528) | 0.0011 | 2.97 | 178.8 | 0.0063 | 178.8 | -1.62
Protein Coverage (double click to dock / undock)
Protein Coverage
Sample name | Protein name | Coverage summary
Spectrum (double click to dock / undock)
N.SDEPDTVK[+162]YCNLGCR.A z=2,scan#=sample=1 period=1 cycle=662 experiment=20,scan time=9.8195
SDEPDTVKYCNLGCR
Intensity
m/z
m/z errors, tol=75.0 ppm
9:06 AM
20/08/2020

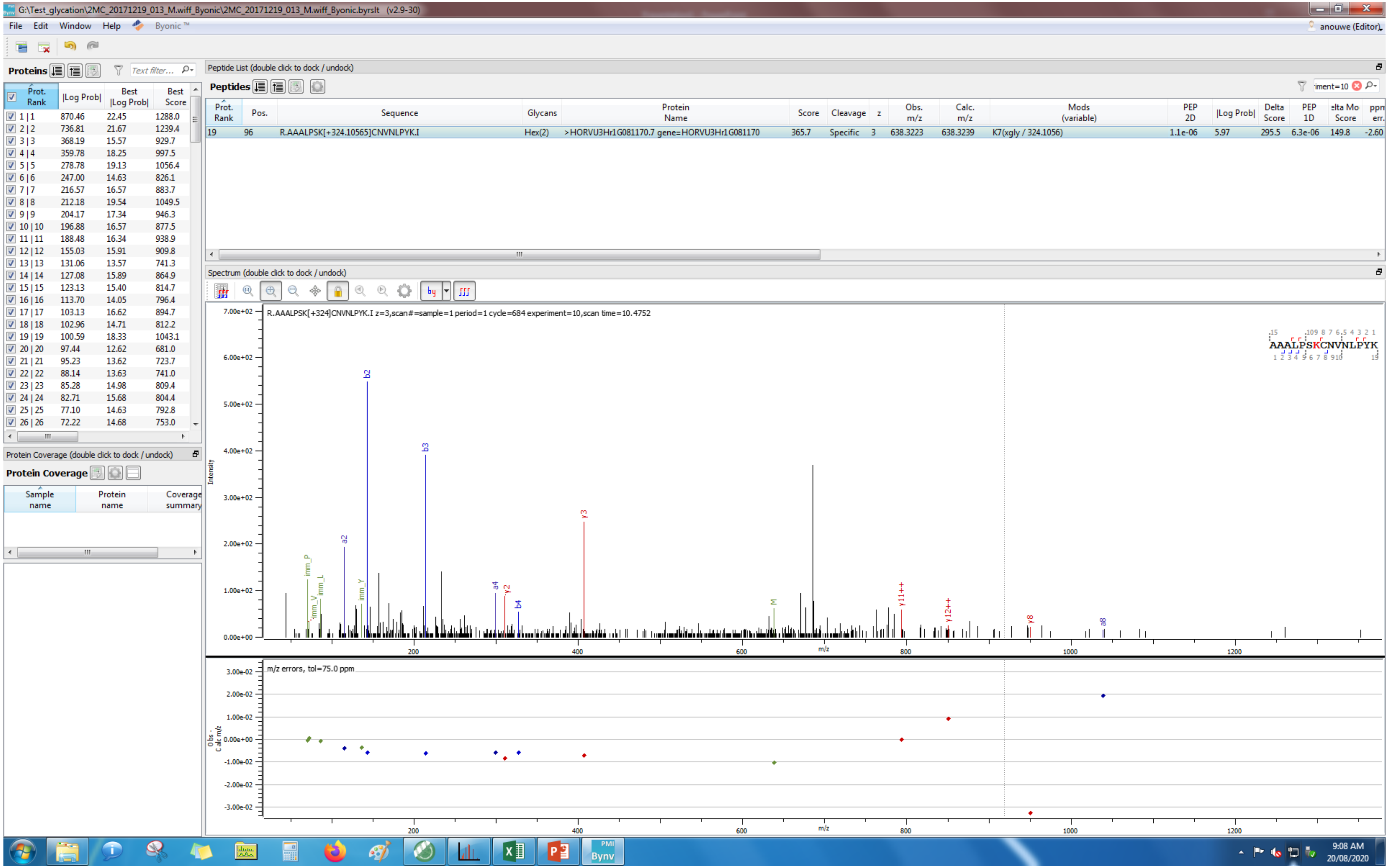
G:\Test_glycation\2MC_20171219_013_M.wiff_Byonic\2MC_20171219_013_M.wiff_Byonic.byrslt (v2.9-30)
File Edit Window Help Byonic ™
anouwe (Editor)
Proteins
Text filter...
Prot. Rank |Log Prob| Best |Log Prob| Best Score
1 | 1 870.46 22.45 1288.0
2 | 2 736.81 21.67 1239.4
3 | 3 368.19 15.57 929.7
4 | 4 359.78 18.25 997.5
5 | 5 278.78 19.13 1056.4
6 | 6 247.00 14.63 826.1
7 | 7 216.57 16.57 883.7
8 | 8 212.18 19.54 1049.5
9 | 9 204.17 17.34 946.3
10 | 10 196.88 16.57 877.5
11 | 11 188.48 16.34 938.9
12 | 12 155.03 15.91 909.8
13 | 13 131.06 13.57 741.3
14 | 14 127.08 15.89 864.9
15 | 15 123.13 15.40 814.7
16 | 16 113.70 14.05 796.4
17 | 17 103.13 16.62 894.7
18 | 18 102.96 14.71 812.2
19 | 19 100.59 18.33 1043.1
20 | 20 97.44 12.62 681.0
21 | 21 95.23 13.62 723.7
22 | 22 88.14 13.63 741.0
23 | 23 85.28 14.98 809.4
24 | 24 82.71 15.68 804.4
25 | 25 77.10 14.63 792.8
26 | 26 72.22 14.68 753.0
Protein Coverage (double click to dock / undock)
Protein Coverage
Sample name Protein name Coverage summary
Peptide List (double click to dock / undock)
Peptides
iment=10
Prot. Rank Pos. Sequence Glycans Protein Name Score Cleavage z Obs. m/z Calc. m/z Mods (variable) PEP 2D |Log Prob| Delta Score PEP 1D elta Mo Score ppm err.
19 96 R.AAALPSK[+324.10565]CNVNLPYK.I Hex(2) >HORVU3Hr1G081170.7 gene=HORVU3Hr1G081170 365.7 Specific 3 638.3223 638.3239 K7(xgly / 324.1056) 1.1e-06 5.97 295.5 6.3e-06 149.8 -2.60
Spectrum (double click to dock / undock)
R.AAALPSK[+324]CNVNLPYK.I z=3,scan#=sample=1 period=1 cycle=684 experiment=10,scan time=10.4752
AAALPSKCNVNLPYK
Intensity
b2 b3 a2 y3 a4 y2 b4 imm_P imm_L imm_V imm_Y M y11++ y12++ y8 a8
m/z
m/z errors, tol=75.0 ppm
Obs - Calc m/z
9:08 AM
20/08/2020

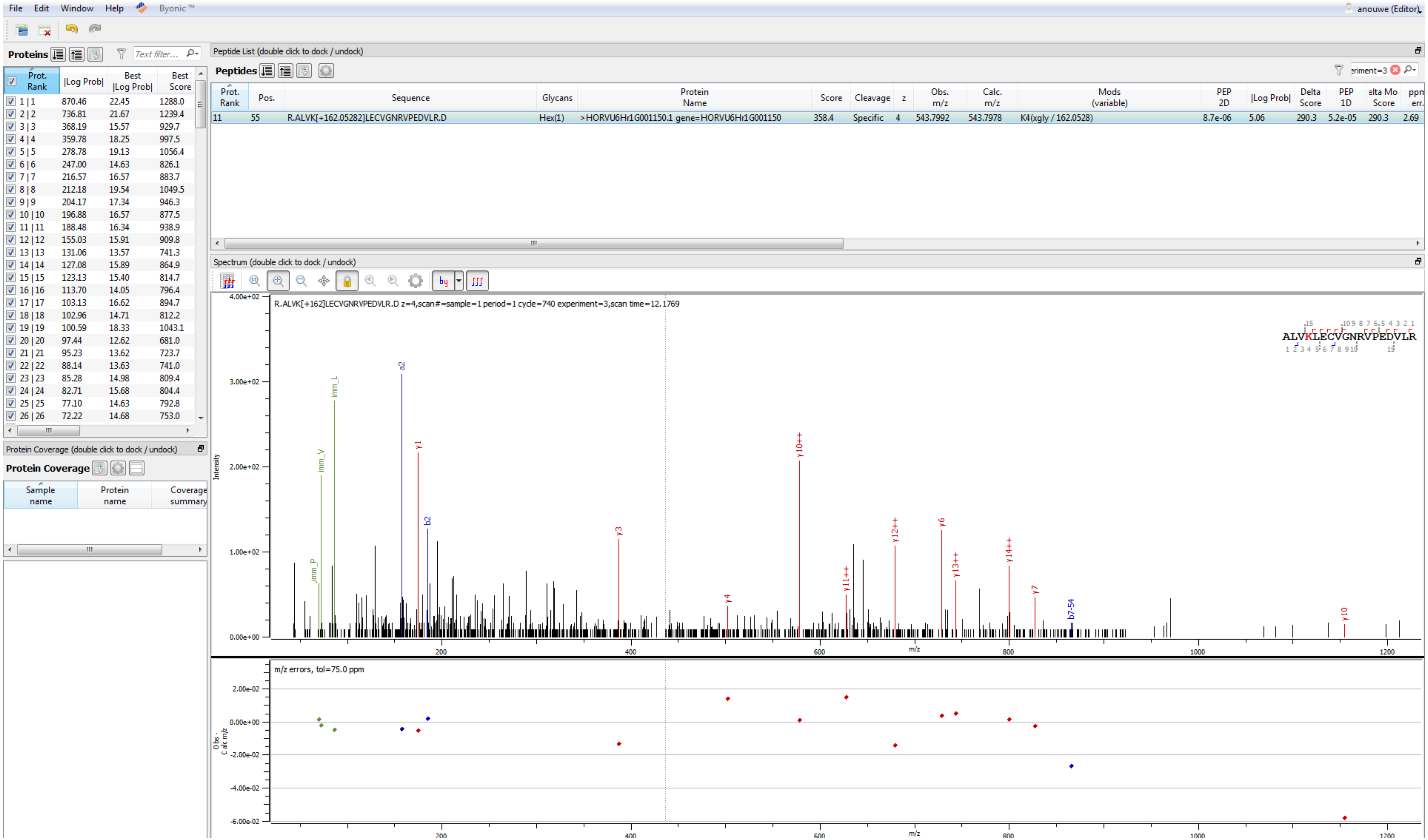

File Edit Window Help Byonic ™
anouwe (Editor)
Proteins
Prot. Rank | |Log Prob| | Best |Log Prob| | Best Score
1 | 1 870.46 22.45 1288.0
2 | 2 736.81 21.67 1239.4
3 | 3 368.19 15.57 929.7
4 | 4 359.78 18.25 997.5
5 | 5 278.78 19.13 1056.4
6 | 6 247.00 14.63 826.1
7 | 7 216.57 16.57 883.7
8 | 8 212.18 19.54 1049.5
9 | 9 204.17 17.34 946.3
10 | 10 196.88 16.57 877.5
11 | 11 188.48 16.34 938.9
12 | 12 155.03 15.91 909.8
13 | 13 131.06 13.57 741.3
14 | 14 127.08 15.89 864.9
15 | 15 123.13 15.40 814.7
16 | 16 113.70 14.05 796.4
17 | 17 103.13 16.62 894.7
18 | 18 102.96 14.71 812.2
19 | 19 100.59 18.33 1043.1
20 | 20 97.44 12.62 681.0
21 | 21 95.23 13.62 723.7
22 | 22 88.14 13.63 741.0
23 | 23 85.28 14.98 809.4
24 | 24 82.71 15.68 804.4
25 | 25 77.10 14.63 792.8
26 | 26 72.22 14.68 753.0
Peptide List (double click to dock / undock)
Peptides
Prot. Rank | Pos. | Sequence | Glycans | Protein Name | Score | Cleavage | z | Obs. m/z | Calc. m/z | Mods (variable) | PEP 2D | |Log Prob| | Delta Score | PEP 1D | elta Mo Score | ppm err.
11 | 55 | R.ALVK[+162.05282]LECVGNRVPEDVLR.D | Hex(1) | >HORVU6Hr1G001150.1 gene=HORVU6Hr1G001150 | 358.4 | Specific | 4 | 543.7992 | 543.7978 | K4(xgly / 162.0528) | 8.7e-06 | 5.06 | 290.3 | 5.2e-05 | 290.3 | 2.69
Spectrum (double click to dock / undock)
R.ALVK[+162]LECVGNRVPEDVLR.D z=4,scan#=sample=1 period=1 cycle=740 experiment=3,scan time=12.1769
ALVKLECVGNRVPEDVLR
Intensity
m/z
imm_L imm_V imm_P a2 y1 b2 y3 y4 y10++ y11++ y12++ y6 y13++ y14++ y7 b7-54 y10
m/z errors, tol=75.0 ppm
Obs - Calc m/z
Protein Coverage (double click to dock / undock)
Protein Coverage
Sample name | Protein name | Coverage summary

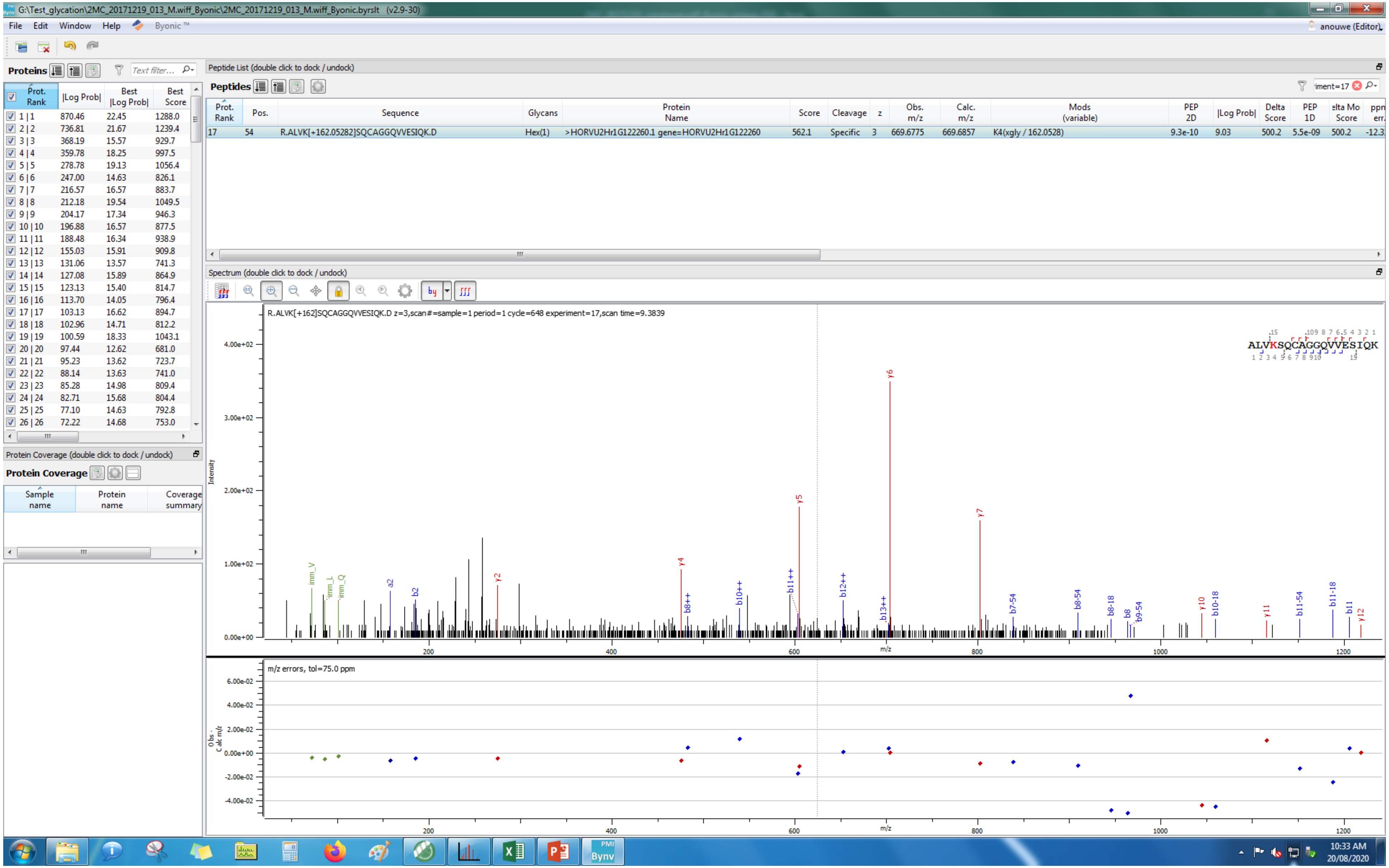
G:\Test_glycation\2MC_20171219_013_M.wiff_Byonic\2MC_20171219_013_M.wiff_Byonic.byrslt (v2.9-30)
File Edit Window Help Byonic
anouwe (Editor)
Proteins
Prot. Rank |Log Prob| Best |Log Prob| Best Score
1 | 1 870.46 22.45 1288.0
2 | 2 736.81 21.67 1239.4
3 | 3 368.19 15.57 929.7
4 | 4 359.78 18.25 997.5
5 | 5 278.78 19.13 1056.4
6 | 6 247.00 14.63 826.1
7 | 7 216.57 16.57 883.7
8 | 8 212.18 19.54 1049.5
9 | 9 204.17 17.34 946.3
10 | 10 196.88 16.57 877.5
11 | 11 188.48 16.34 938.9
12 | 12 155.03 15.91 909.8
13 | 13 131.06 13.57 741.3
14 | 14 127.08 15.89 864.9
15 | 15 123.13 15.40 814.7
16 | 16 113.70 14.05 796.4
17 | 17 103.13 16.62 894.7
18 | 18 102.96 14.71 812.2
19 | 19 100.59 18.33 1043.1
20 | 20 97.44 12.62 681.0
21 | 21 95.23 13.62 723.7
22 | 22 88.14 13.63 741.0
23 | 23 85.28 14.98 809.4
24 | 24 82.71 15.68 804.4
25 | 25 77.10 14.63 792.8
26 | 26 72.22 14.68 753.0
Protein Coverage (double click to dock / undock)
Protein Coverage
Sample name Protein name Coverage summary
Peptide List (double click to dock / undock)
Peptides
Prot. Rank Pos. Sequence Glycans Protein Name Score Cleavage z Obs. m/z Calc. m/z Mods (variable) PEP 2D |Log Prob| Delta Score PEP 1D Delta Mod Score ppm err.
17 54 R.ALVK[+162.05282]SQCAGGQVVESIQK.D Hex(1) >HORVU2Hr1G122260.1 gene=HORVU2Hr1G122260 562.1 Specific 3 669.6775 669.6857 K4(xgly / 162.0528) 9.3e-10 9.03 500.2 5.5e-09 500.2 -12.3
Spectrum (double click to dock / undock)
R.ALVK[+162]SQCAGGQVVESIQK.D z=3,scan#=sample=1 period=1 cycle=648 experiment=17,scan time=9.3839
ALVKSQCAGGQVVESIQK
Intensity
m/z
m/z errors, tol=75.0 ppm
10:33 AM
20/08/2020

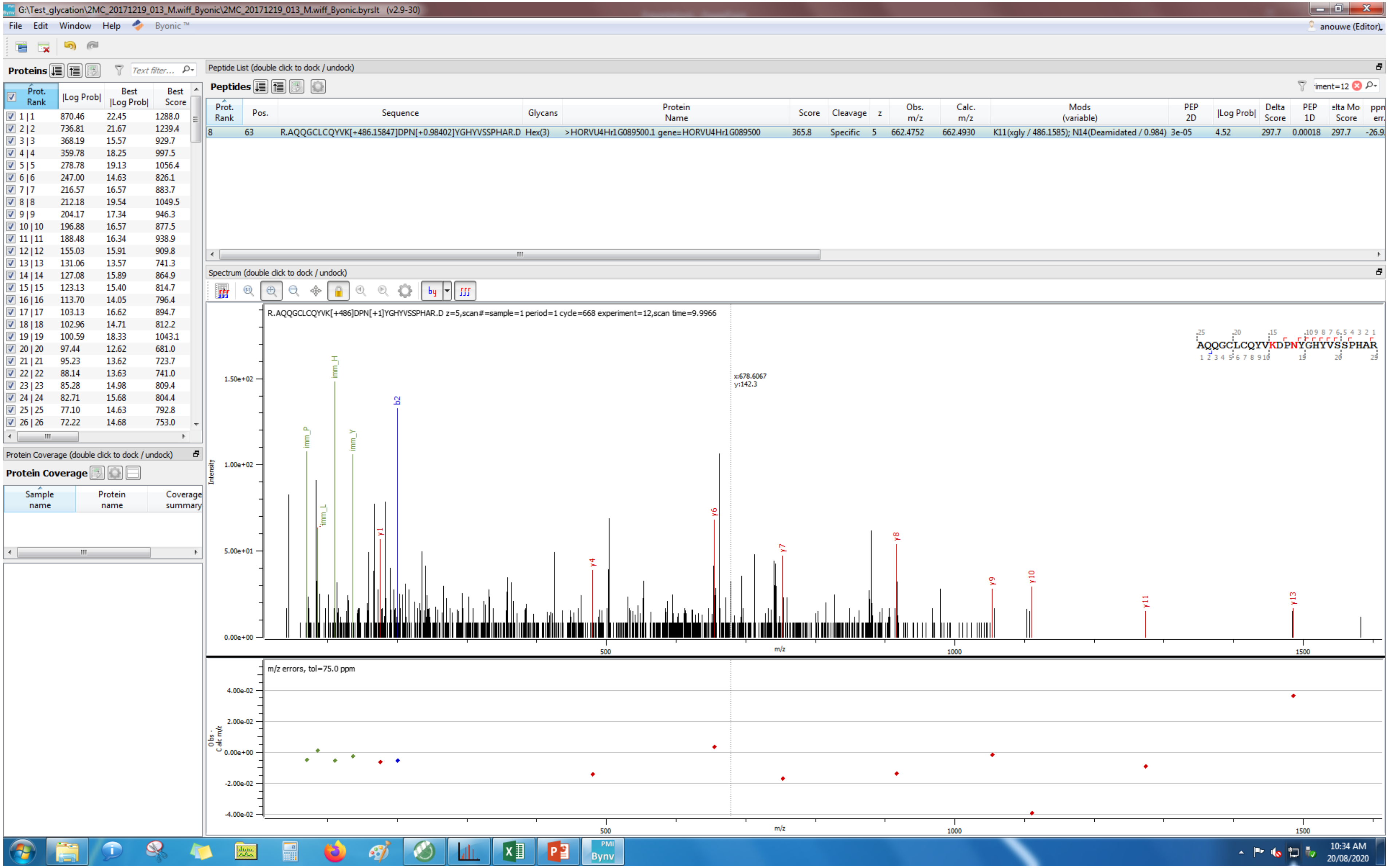
G:\Test_glycation\2MC_20171219_013_M.wiff_Byonic\2MC_20171219_013_M.wiff_Byonic.byrslt (v2.9-30)
File Edit Window Help Byonic™
anouwe (Editor)
Proteins
Peptide List (double click to dock / undock)
Peptides
Prot. Rank | Pos. | Sequence | Glycans | Protein Name | Score | Cleavage | z | Obs. m/z | Calc. m/z | Mods (variable) | PEP 2D | |Log Prob| | Delta Score | PEP 1D | elta Mo Score | ppm err.
8 | 63 | R.AQQGCLCQYVK[+486.15847]DPN[+0.98402]YGHYVSSPHAR.D | Hex(3) | >HORVU4Hr1G089500.1 gene=HORVU4Hr1G089500 | 365.8 | Specific | 5 | 662.4752 | 662.4930 | K11(xgly / 486.1585); N14(Deamidated / 0.984) | 3e-05 | 4.52 | 297.7 | 0.00018 | 297.7 | -26.9
Spectrum (double click to dock / undock)
R.AQQGCLCQYVK[+486]DPN[+1]YGHYVSSPHAR.D z=5,scan#=sample=1 period=1 cycle=668 experiment=12,scan time=9.9966
AQQGCLCQYVKDPNYGHYVSSPHAR
x:678.6067
y:142.3
Intensity
m/z
m/z errors, tol=75.0 ppm
Obs - Calc m/z
Protein Coverage (double click to dock / undock)
Protein Coverage
Sample name | Protein name | Coverage summary
10:34 AM
20/08/2020

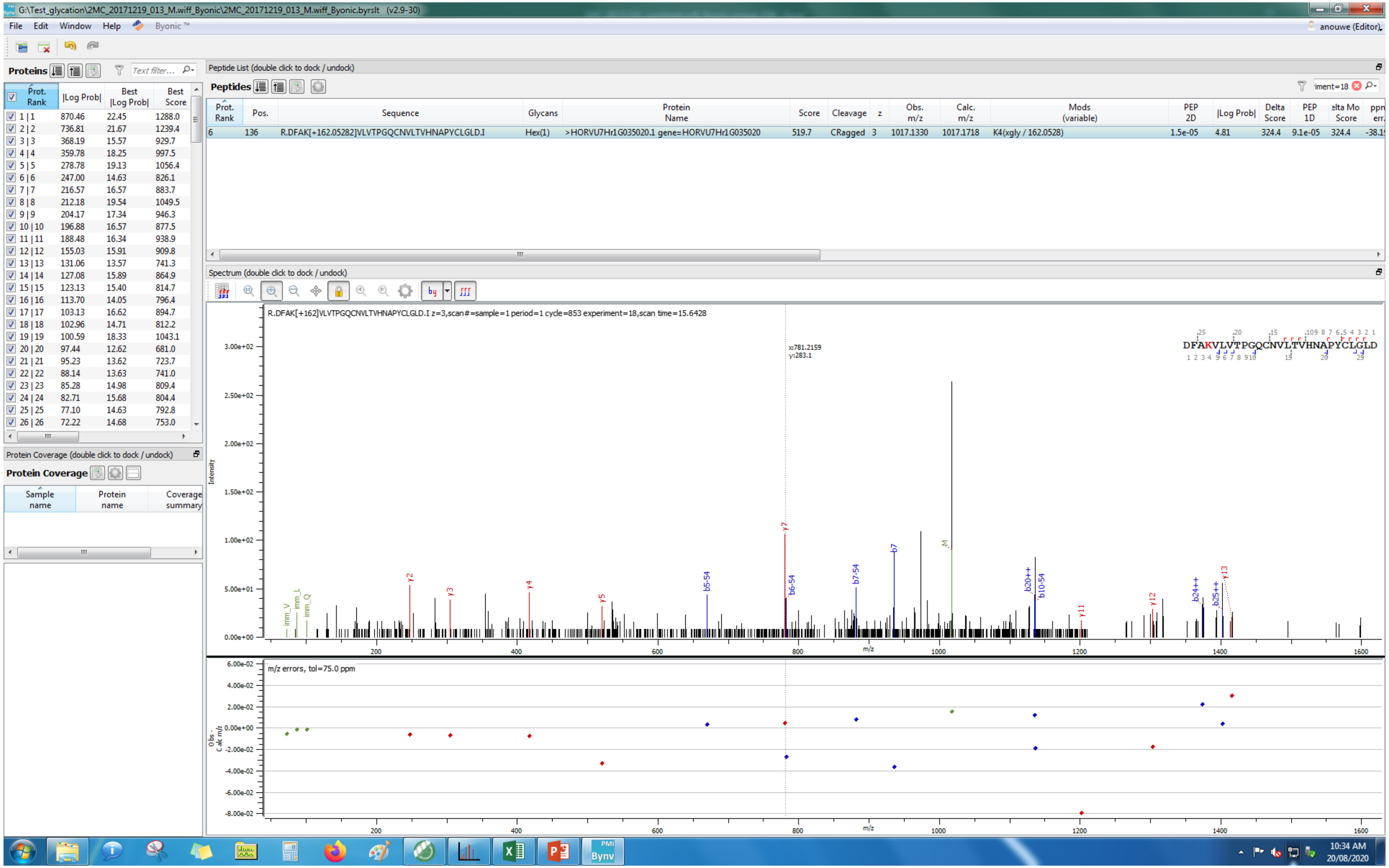
G:\Test_glycation\2MC_20171219_013_M.wiff_Byonic\2MC_20171219_013_M.wiff_Byonic.byrslt (v2.9-30)
File Edit Window Help Byonic
anouwe (Editor)
Proteins
Prot. Rank | |Log Prob| | Best |Log Prob| | Best Score
1 | 1 870.46 22.45 1288.0
2 | 2 736.81 21.67 1239.4
3 | 3 368.19 15.57 929.7
4 | 4 359.78 18.25 997.5
5 | 5 278.78 19.13 1056.4
6 | 6 247.00 14.63 826.1
7 | 7 216.57 16.57 883.7
8 | 8 212.18 19.54 1049.5
9 | 9 204.17 17.34 946.3
10 | 10 196.88 16.57 877.5
11 | 11 188.48 16.34 938.9
12 | 12 155.03 15.91 909.8
13 | 13 131.06 13.57 741.3
14 | 14 127.08 15.89 864.9
15 | 15 123.13 15.40 814.7
16 | 16 113.70 14.05 796.4
17 | 17 103.13 16.62 894.7
18 | 18 102.96 14.71 812.2
19 | 19 100.59 18.33 1043.1
20 | 20 97.44 12.62 681.0
21 | 21 95.23 13.62 723.7
22 | 22 88.14 13.63 741.0
23 | 23 85.28 14.98 809.4
24 | 24 82.71 15.68 804.4
25 | 25 77.10 14.63 792.8
26 | 26 72.22 14.68 753.0
Protein Coverage (double click to dock / undock)
Protein Coverage
Sample name | Protein name | Coverage summary
Peptide List (double click to dock / undock)
Peptides
Prot. Rank | Pos. | Sequence | Glycans | Protein Name | Score | Cleavage | z | Obs. m/z | Calc. m/z | Mods (variable) | PEP 2D | |Log Prob| | Delta Score | PEP 1D | Delta Mod Score
6 | 136 | R.DFAK[+162.05282]VLVTPGQCNVLTVHNAPYCLGLD.I | Hex(1) | >HORVU7Hr1G035020.1 gene=HORVU7Hr1G035020 | 519.7 | CRagged | 3 | 1017.1330 | 1017.1718 | K4(xgly / 162.0528) | 1.5e-05 | 4.81 | 324.4 | 9.1e-05 | 324.4
Spectrum (double click to dock / undock)
R.DFAK[+162]VLVTPGQCNVLTVHNAPYCLGLD.I z=3,scan#=sample=1 period=1 cycle=853 experiment=18,scan time=15.6428
x:781.2159
y:283.1
DFAKVLVTPGQCNVLTVHNAPYCLGLD
Intensity
m/z
m/z errors, tol=75.0 ppm
Obs - Calc m/z
10:34 AM
20/08/2020

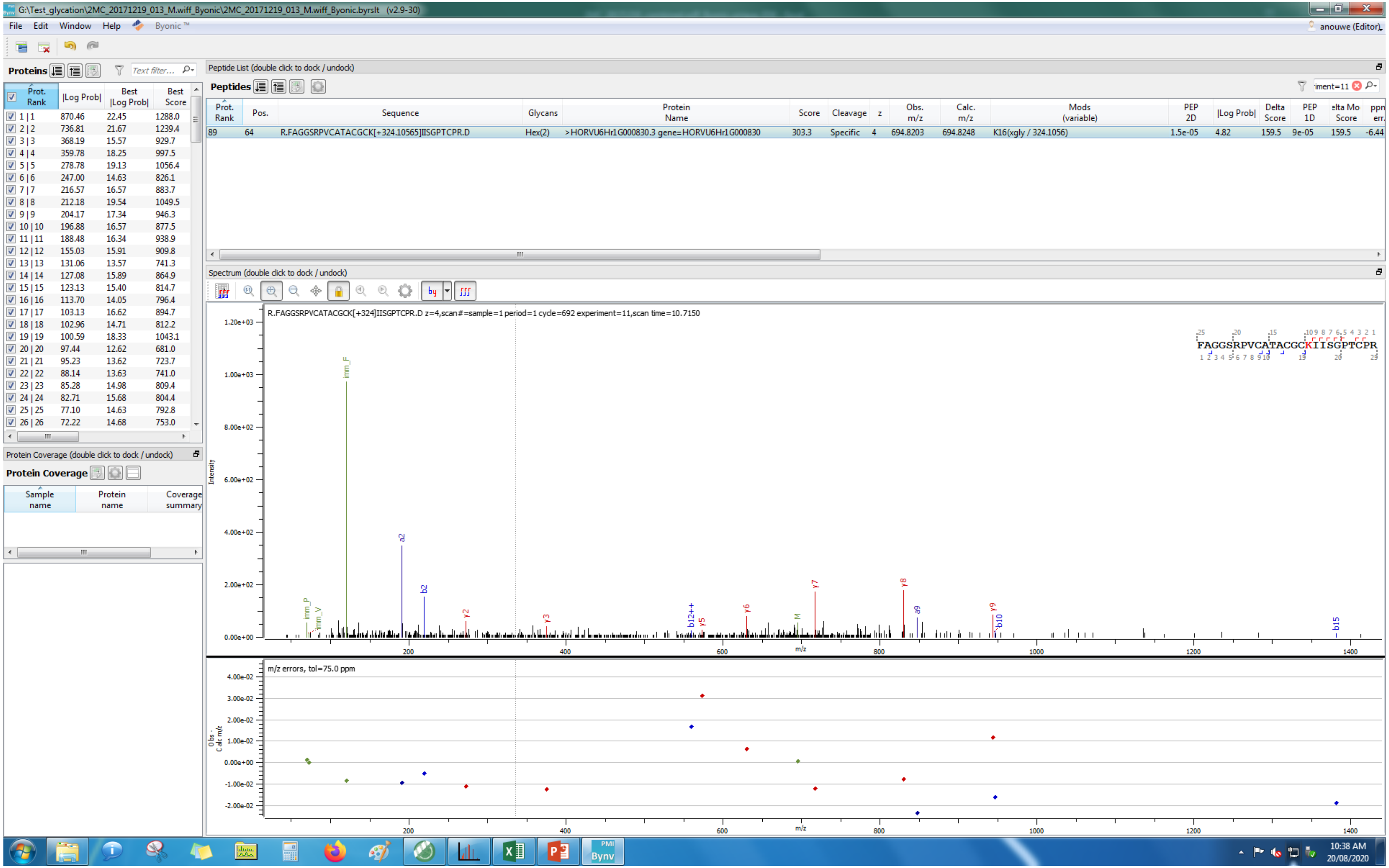
G:\Test_glycation\2MC_20171219_013_M.wiff_Byonic\2MC_20171219_013_M.wiff_Byonic.byrslt (v2.9-30)
File Edit Window Help Byonic™
anouwe (Editor)
Proteins
Prot. Rank | |Log Prob| | Best |Log Prob| | Best Score
1 | 1 870.46 22.45 1288.0
2 | 2 736.81 21.67 1239.4
3 | 3 368.19 15.57 929.7
4 | 4 359.78 18.25 997.5
5 | 5 278.78 19.13 1056.4
6 | 6 247.00 14.63 826.1
7 | 7 216.57 16.57 883.7
8 | 8 212.18 19.54 1049.5
9 | 9 204.17 17.34 946.3
10 | 10 196.88 16.57 877.5
11 | 11 188.48 16.34 938.9
12 | 12 155.03 15.91 909.8
13 | 13 131.06 13.57 741.3
14 | 14 127.08 15.89 864.9
15 | 15 123.13 15.40 814.7
16 | 16 113.70 14.05 796.4
17 | 17 103.13 16.62 894.7
18 | 18 102.96 14.71 812.2
19 | 19 100.59 18.33 1043.1
20 | 20 97.44 12.62 681.0
21 | 21 95.23 13.62 723.7
22 | 22 88.14 13.63 741.0
23 | 23 85.28 14.98 809.4
24 | 24 82.71 15.68 804.4
25 | 25 77.10 14.63 792.8
26 | 26 72.22 14.68 753.0
Protein Coverage (double click to dock / undock)
Protein Coverage
Sample name | Protein name | Coverage summary
Peptide List (double click to dock / undock)
Peptides
Prot. Rank | Pos. | Sequence | Glycans | Protein Name | Score | Cleavage | z | Obs. m/z | Calc. m/z | Mods (variable) | PEP 2D | |Log Prob| | Delta Score | PEP 1D | elta Mo Score | ppm err.
89 | 64 | R.FAGGSRPVCATACGCK[+324.10565]IISGPTCPR.D | Hex(2) | >HORVU6Hr1G000830.3 gene=HORVU6Hr1G000830 | 303.3 | Specific | 4 | 694.8203 | 694.8248 | K16(xgly / 324.1056) | 1.5e-05 | 4.82 | 159.5 | 9e-05 | 159.5 | -6.44
Spectrum (double click to dock / undock)
R.FAGGSRPVCATACGCK[+324]IISGPTCPR.D z=4,scan#=sample=1 period=1 cycle=692 experiment=11,scan time=10.7150
FAGGSRPVCATACGCKIISGPTCPR
Intensity
m/z
imm_F
imm_P
imm_V
a2
b2
y2
y3
b12++
y5
y6
M
y7
y8
a9
y9
b10
b15
m/z errors, tol=75.0 ppm
Obs - Calc m/z
10:38 AM
20/08/2020

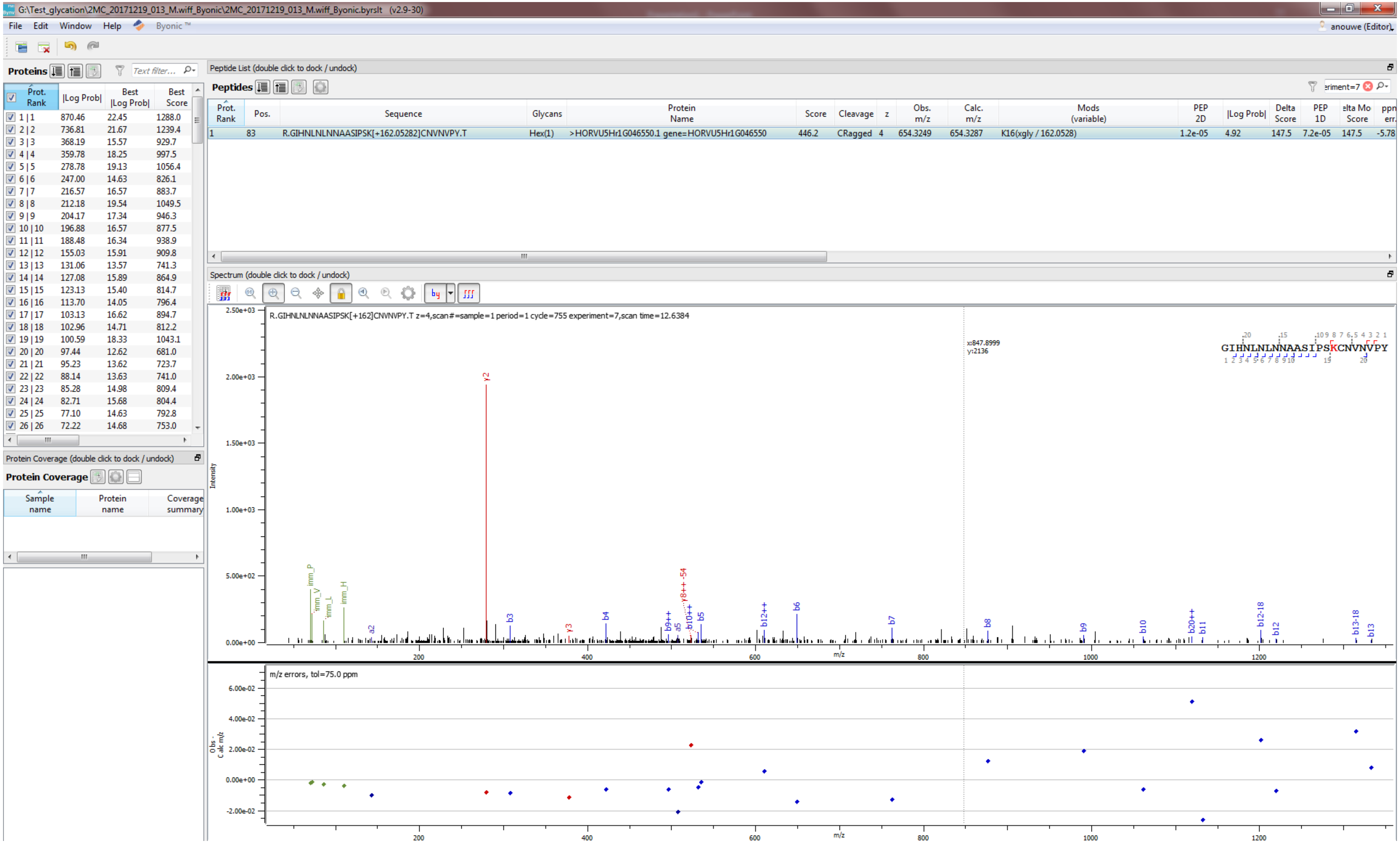
G:\Test_glycation\2MC_20171219_013_M.wiff_Byonic\2MC_20171219_013_M.wiff_Byonic.byrslt (v2.9-30)
File Edit Window Help Byonic ™
anouwe (Editor)
Proteins
Text filter...
Prot. Rank | |Log Prob| | Best |Log Prob| | Best Score
1 | 1 870.46 22.45 1288.0
2 | 2 736.81 21.67 1239.4
3 | 3 368.19 15.57 929.7
4 | 4 359.78 18.25 997.5
5 | 5 278.78 19.13 1056.4
6 | 6 247.00 14.63 826.1
7 | 7 216.57 16.57 883.7
8 | 8 212.18 19.54 1049.5
9 | 9 204.17 17.34 946.3
10 | 10 196.88 16.57 877.5
11 | 11 188.48 16.34 938.9
12 | 12 155.03 15.91 909.8
13 | 13 131.06 13.57 741.3
14 | 14 127.08 15.89 864.9
15 | 15 123.13 15.40 814.7
16 | 16 113.70 14.05 796.4
17 | 17 103.13 16.62 894.7
18 | 18 102.96 14.71 812.2
19 | 19 100.59 18.33 1043.1
20 | 20 97.44 12.62 681.0
21 | 21 95.23 13.62 723.7
22 | 22 88.14 13.63 741.0
23 | 23 85.28 14.98 809.4
24 | 24 82.71 15.68 804.4
25 | 25 77.10 14.63 792.8
26 | 26 72.22 14.68 753.0
Protein Coverage (double click to dock / undock)
Protein Coverage
Sample name | Protein name | Coverage summary
Peptide List (double click to dock / undock)
Peptides
eriment=7
Prot. Rank | Pos. | Sequence | Glycans | Protein Name | Score | Cleavage | z | Obs. m/z | Calc. m/z | Mods (variable) | PEP 2D | |Log Prob| | Delta Score | PEP 1D | elta Mo Score | ppm err.
1 | 83 | R.GIHNLNLNNAASIPSK[+162.05282]CNVNVPY.T | Hex(1) | >HORVU5Hr1G046550.1 gene=HORVU5Hr1G046550 | 446.2 | CRagged | 4 | 654.3249 | 654.3287 | K16(xgly / 162.0528) | 1.2e-05 | 4.92 | 147.5 | 7.2e-05 | 147.5 | -5.78
Spectrum (double click to dock / undock)
R.GIHNLNLNNAASIPSK[+162]CNVNVPY.T z=4,scan#=sample=1 period=1 cycle=755 experiment=7,scan time=12.6384
x:847.8999
y:2136
GIHNLNLNNAASIPSKCNVNVPY
Intensity
2.50e+03
2.00e+03
1.50e+03
1.00e+03
5.00e+02
0.00e+00
imm_P imm_V imm_L imm_H a2 y2 b3 y3 b4 b9++ a5 b10++ y8++ -54 b5 b12++ b6 b7 b8 b9 b10 b20++ b11 b12-18 b12 b13-18 b13
200 400 600 800 1000 1200
m/z
m/z errors, tol=75.0 ppm
Obs - Calc m/z
6.00e-02
4.00e-02
2.00e-02
0.00e+00
-2.00e-02
200 400 600 800 1000 1200
m/z

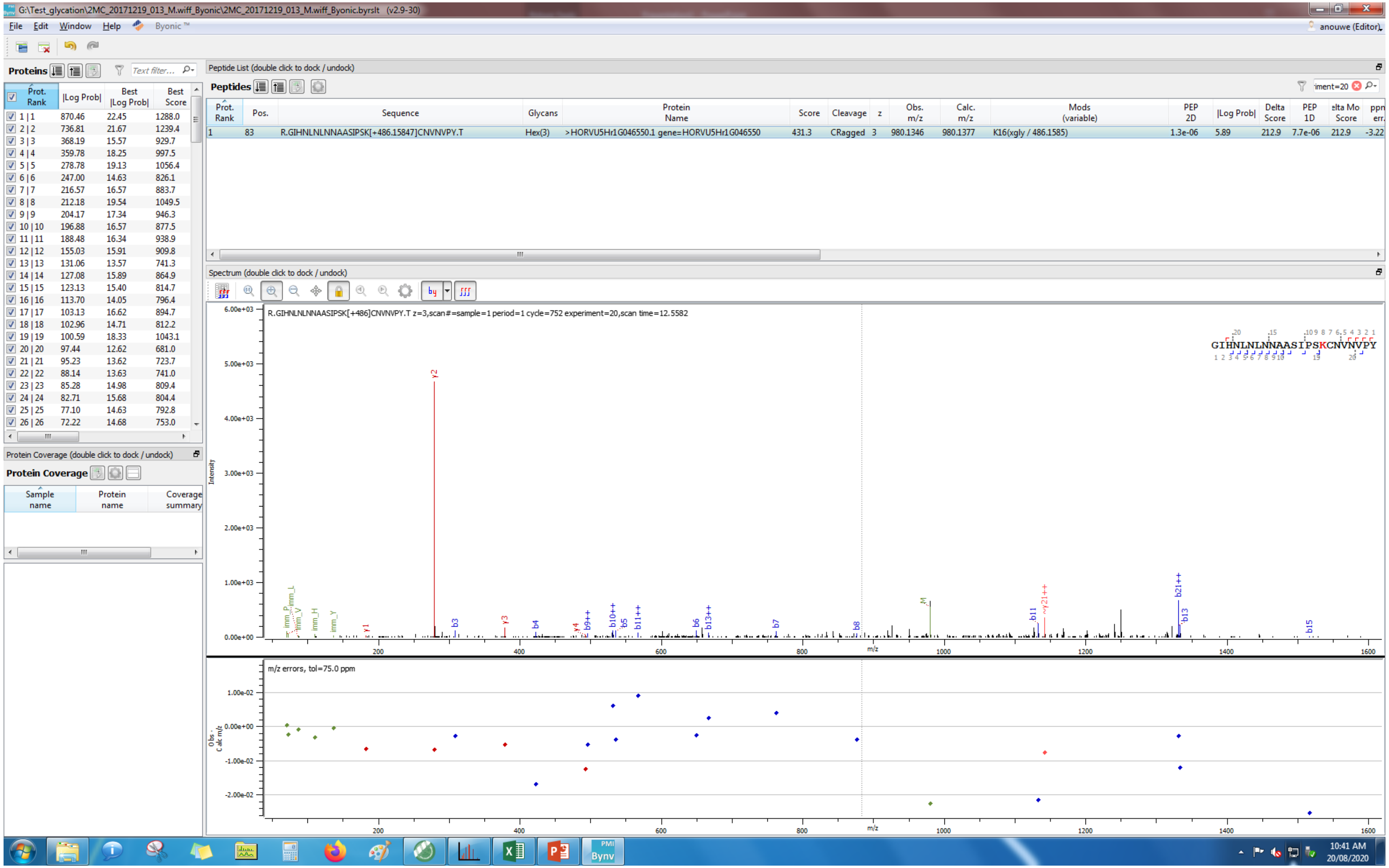
G:\Test_glycation\2MC_20171219_013_M.wiff_Byonic\2MC_20171219_013_M.wiff_Byonic.byrslt (v2.9-30)
File Edit Window Help Byonic ™
anouwe (Editor)
Proteins
Text filter...
Prot. Rank | |Log Prob| | Best |Log Prob| | Best Score
1 | 1 870.46 22.45 1288.0
2 | 2 736.81 21.67 1239.4
3 | 3 368.19 15.57 929.7
4 | 4 359.78 18.25 997.5
5 | 5 278.78 19.13 1056.4
6 | 6 247.00 14.63 826.1
7 | 7 216.57 16.57 883.7
8 | 8 212.18 19.54 1049.5
9 | 9 204.17 17.34 946.3
10 | 10 196.88 16.57 877.5
11 | 11 188.48 16.34 938.9
12 | 12 155.03 15.91 909.8
13 | 13 131.06 13.57 741.3
14 | 14 127.08 15.89 864.9
15 | 15 123.13 15.40 814.7
16 | 16 113.70 14.05 796.4
17 | 17 103.13 16.62 894.7
18 | 18 102.96 14.71 812.2
19 | 19 100.59 18.33 1043.1
20 | 20 97.44 12.62 681.0
21 | 21 95.23 13.62 723.7
22 | 22 88.14 13.63 741.0
23 | 23 85.28 14.98 809.4
24 | 24 82.71 15.68 804.4
25 | 25 77.10 14.63 792.8
26 | 26 72.22 14.68 753.0
Protein Coverage (double click to dock / undock)
Protein Coverage
Sample name | Protein name | Coverage summary
Peptide List (double click to dock / undock)
Peptides
iment=20
Prot. Rank | Pos. | Sequence | Glycans | Protein Name | Score | Cleavage | z | Obs. m/z | Calc. m/z | Mods (variable) | PEP 2D | |Log Prob| | Delta Score | PEP 1D | elta Mo Score | ppn err.
1 | 83 | R.GIHNLNLNNAASIPSK[+486.15847]CNVNVPY.T | Hex(3) | >HORVU5Hr1G046550.1 gene=HORVU5Hr1G046550 | 431.3 | CRagged | 3 | 980.1346 | 980.1377 | K16(xgly / 486.1585) | 1.3e-06 | 5.89 | 212.9 | 7.7e-06 | 212.9 | -3.22
Spectrum (double click to dock / undock)
R.GIHNLNLNNAASIPSK[+486]CNVNVPY.T z=3,scan#=sample=1 period=1 cycle=752 experiment=20,scan time=12.5582
GIHNLNLNNAASIPSKCNVNVPY
Intensity
6.00e+03
5.00e+03
4.00e+03
3.00e+03
2.00e+03
1.00e+03
0.00e+00
y2 y1 b3 y3 b4 y4 b9++ b10++ b5 b11++ b6 b13++ b7 b8 M b11 y21++ b21++ b13 b15
imm_L imm_P imm_V imm_H imm_Y
m/z
200 400 600 800 1000 1200 1400 1600
m/z errors, tol=75.0 ppm
Obs - Calc m/z
1.00e-02
0.00e+00
-1.00e-02
-2.00e-02
10:41 AM
20/08/2020

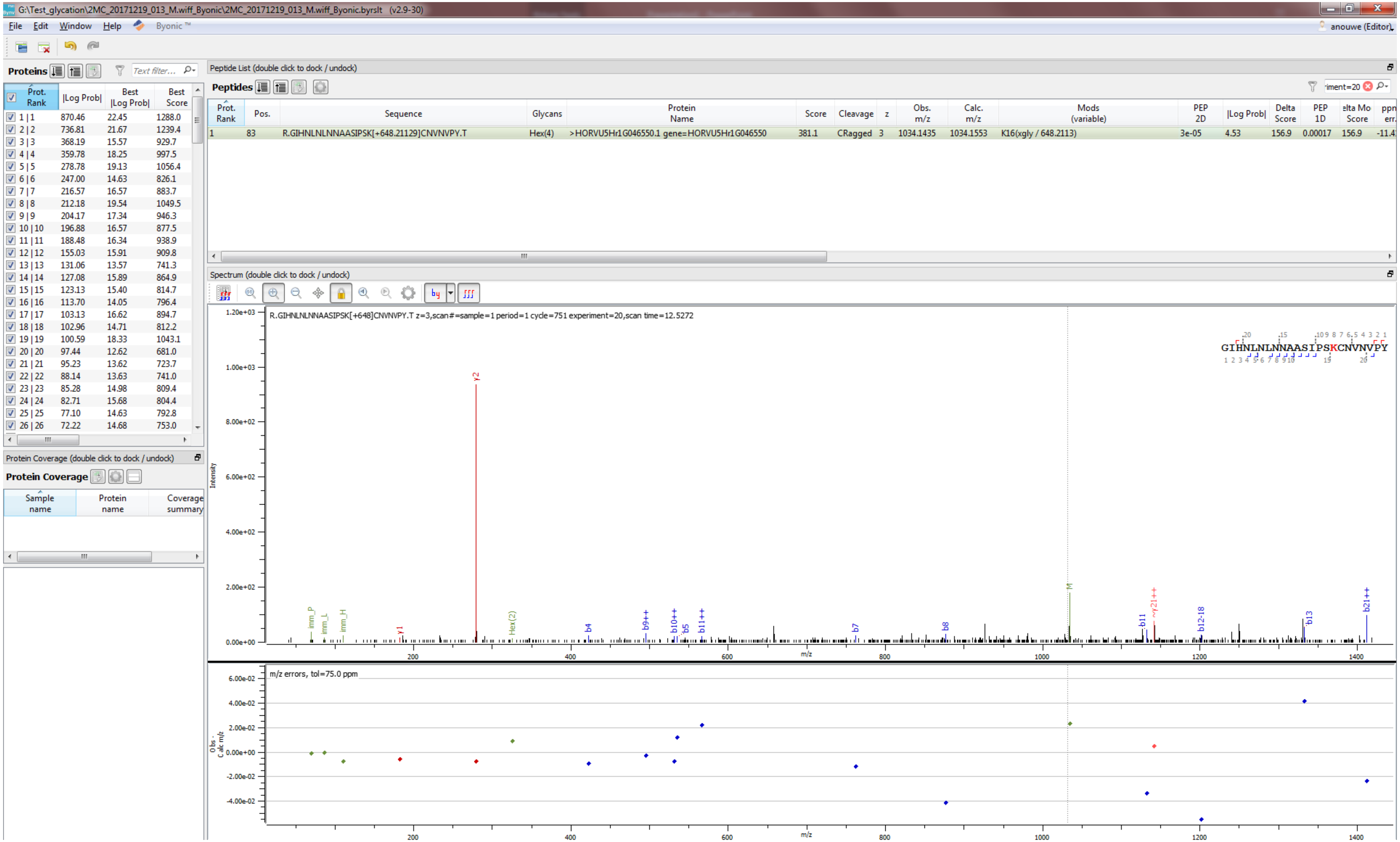
G:\Test_glycation\2MC_20171219_013_M.wiff_Byonic\2MC_20171219_013_M.wiff_Byonic.byrslt (v2.9-30)
File Edit Window Help Byonic™
anouwe (Editor)
Proteins
Peptide List (double click to dock / undock)
Peptides
Prot. Rank | Pos. | Sequence | Glycans | Protein Name | Score | Cleavage | z | Obs. m/z | Calc. m/z | Mods (variable) | PEP 2D | |Log Prob| | Delta Score | PEP 1D | Delta Mod Score | ppm err.
1 | 83 | R.GIHNLNLNNAASIPSK[+648.21129]CNVNVPY.T | Hex(4) | >HORVU5Hr1G046550.1 gene=HORVU5Hr1G046550 | 381.1 | CRagged | 3 | 1034.1435 | 1034.1553 | K16(xgly / 648.2113) | 3e-05 | 4.53 | 156.9 | 0.00017 | 156.9 | -11.4
Prot. Rank | |Log Prob| | Best |Log Prob| | Best Score
1 | 1 | 870.46 | 22.45 | 1288.0
2 | 2 | 736.81 | 21.67 | 1239.4
3 | 3 | 368.19 | 15.57 | 929.7
4 | 4 | 359.78 | 18.25 | 997.5
5 | 5 | 278.78 | 19.13 | 1056.4
6 | 6 | 247.00 | 14.63 | 826.1
7 | 7 | 216.57 | 16.57 | 883.7
8 | 8 | 212.18 | 19.54 | 1049.5
9 | 9 | 204.17 | 17.34 | 946.3
10 | 10 | 196.88 | 16.57 | 877.5
11 | 11 | 188.48 | 16.34 | 938.9
12 | 12 | 155.03 | 15.91 | 909.8
13 | 13 | 131.06 | 13.57 | 741.3
14 | 14 | 127.08 | 15.89 | 864.9
15 | 15 | 123.13 | 15.40 | 814.7
16 | 16 | 113.70 | 14.05 | 796.4
17 | 17 | 103.13 | 16.62 | 894.7
18 | 18 | 102.96 | 14.71 | 812.2
19 | 19 | 100.59 | 18.33 | 1043.1
20 | 20 | 97.44 | 12.62 | 681.0
21 | 21 | 95.23 | 13.62 | 723.7
22 | 22 | 88.14 | 13.63 | 741.0
23 | 23 | 85.28 | 14.98 | 809.4
24 | 24 | 82.71 | 15.68 | 804.4
25 | 25 | 77.10 | 14.63 | 792.8
26 | 26 | 72.22 | 14.68 | 753.0
Protein Coverage (double click to dock / undock)
Protein Coverage
Sample name | Protein name | Coverage summary
Spectrum (double click to dock / undock)
R.GIHNLNLNNAASIPSK[+648]CNVNVPY.T z=3,scan#=sample=1 period=1 cycle=751 experiment=20,scan time=12.5272
GIHNLNLNNAASIPSKCNVNVPY
Intensity
m/z
y2
y1
imm_P
imm_L
imm_H
Hex(2)
b4
b9++
b10++
b5
b11++
b7
b8
b11
y21++
b12-18
b13
b21++
M
m/z errors, tol=75.0 ppm
Obs - Calc m/z

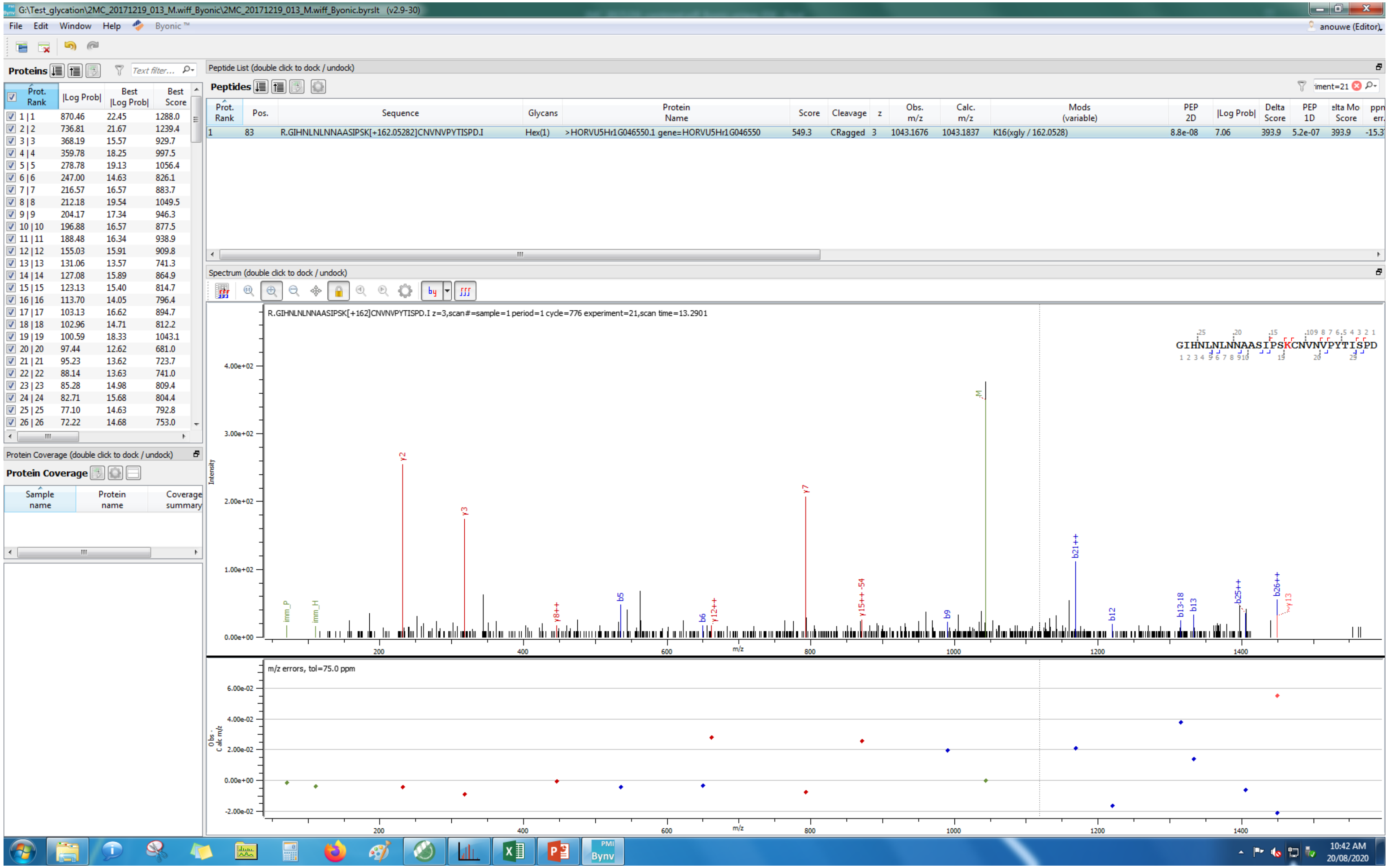

G:\Test_glycation\2MC_20171219_013_M.wiff_Byonic\2MC_20171219_013_M.wiff_Byonic.byrslt (v2.9-30)
File Edit Window Help Byonic ™
anouwe (Editor)
Proteins
Text filter...
Prot. Rank | |Log Prob| | Best |Log Prob| | Best Score
1 | 1 870.46 22.45 1288.0
2 | 2 736.81 21.67 1239.4
3 | 3 368.19 15.57 929.7
4 | 4 359.78 18.25 997.5
5 | 5 278.78 19.13 1056.4
6 | 6 247.00 14.63 826.1
7 | 7 216.57 16.57 883.7
8 | 8 212.18 19.54 1049.5
9 | 9 204.17 17.34 946.3
10 | 10 196.88 16.57 877.5
11 | 11 188.48 16.34 938.9
12 | 12 155.03 15.91 909.8
13 | 13 131.06 13.57 741.3
14 | 14 127.08 15.89 864.9
15 | 15 123.13 15.40 814.7
16 | 16 113.70 14.05 796.4
17 | 17 103.13 16.62 894.7
18 | 18 102.96 14.71 812.2
19 | 19 100.59 18.33 1043.1
20 | 20 97.44 12.62 681.0
21 | 21 95.23 13.62 723.7
22 | 22 88.14 13.63 741.0
23 | 23 85.28 14.98 809.4
24 | 24 82.71 15.68 804.4
25 | 25 77.10 14.63 792.8
26 | 26 72.22 14.68 753.0
Protein Coverage (double click to dock / undock)
Protein Coverage
Sample name | Protein name | Coverage summary
Peptide List (double click to dock / undock)
Peptides
iment=21
Prot. Rank | Pos. | Sequence | Glycans | Protein Name | Score | Cleavage | z | Obs. m/z | Calc. m/z | Mods (variable) | PEP 2D | |Log Prob| | Delta Score | PEP 1D | elta Mo Score | ppn err.
1 | 83 | R.GIHNLNLNNAASIPSK[+162.05282]CNVNVPYTISPD.I | Hex(1) | >HORVU5Hr1G046550.1 gene=HORVU5Hr1G046550 | 549.3 | CRagged | 3 | 1043.1676 | 1043.1837 | K16(xgly / 162.0528) | 8.8e-08 | 7.06 | 393.9 | 5.2e-07 | 393.9 | -15.3
Spectrum (double click to dock / undock)
R.GIHNLNLNNAASIPSK[+162]CNVNVPYTISPD.I z=3,scan#=sample=1 period=1 cycle=776 experiment=21,scan time=13.2901
GIHNLNLNNAASIPSKCNVNVPYTISPD
Intensity
m/z
y2
y3
y7
M
b21++
b5
b6
y8++
y12++
y15++-54
b9
b12
b13-18
b13
b25++
b26++
y13
imm_P
imm_H
m/z errors, tol=75.0 ppm
Obs - Calc m/z
10:42 AM
20/08/2020

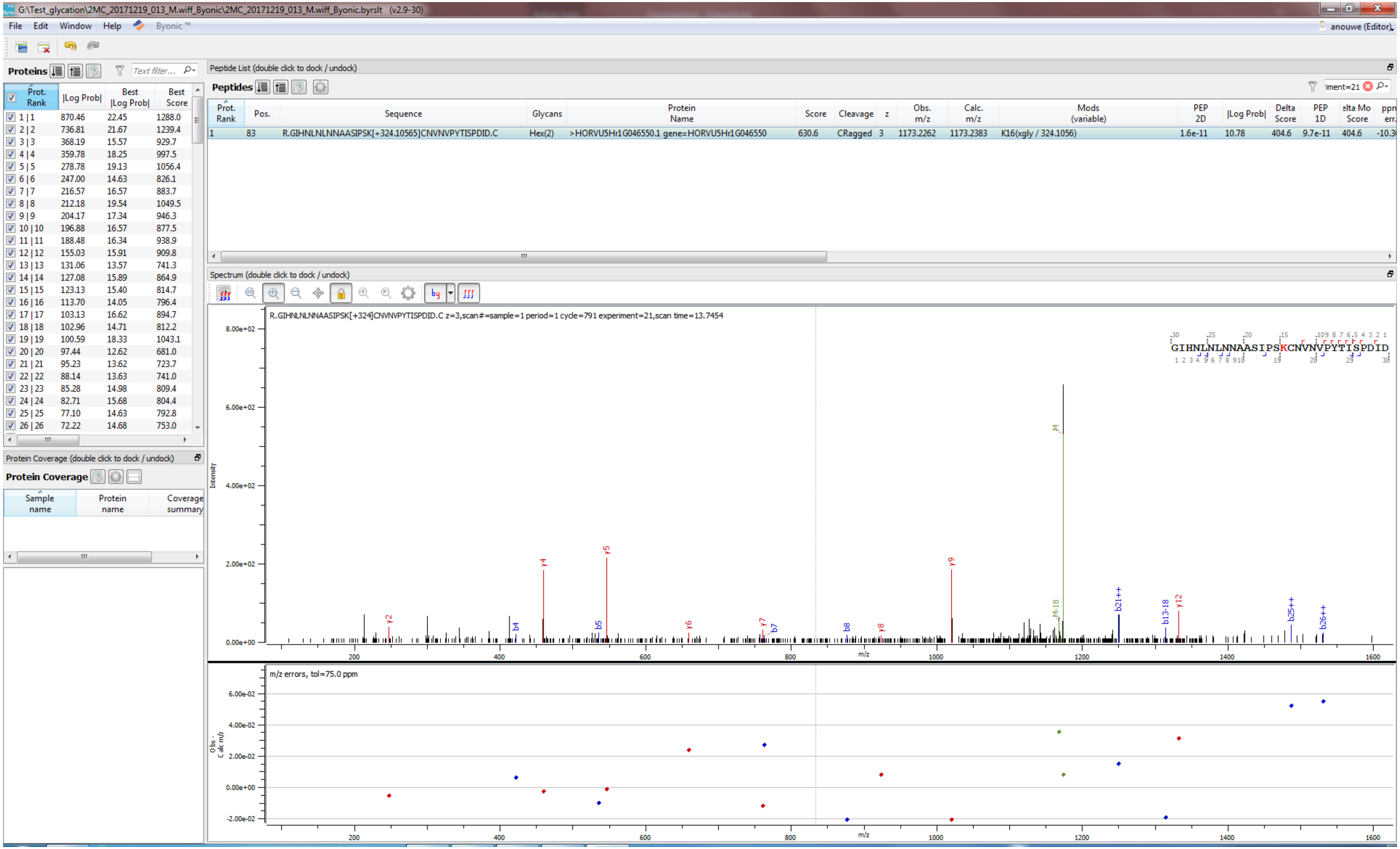
G:\Test_glycation\2MC_20171219_013_M.wiff_Byonic\2MC_20171219_013_M.wiff_Byonic.byrslt (v2.9-30)
File Edit Window Help Byonic ™
anouwe (Editor)
Proteins
Peptide List (double click to dock / undock)
Peptides
Prot. Rank | Pos. | Sequence | Glycans | Protein Name | Score | Cleavage | z | Obs. m/z | Calc. m/z | Mods (variable) | PEP 2D | |Log Prob| | Delta Score | PEP 1D
1 | 83 | R.GIHNLNLNNAASIPSK[+324.10565]CNVNVPYTISPDID.C | Hex(2) | >HORVU5Hr1G046550.1 gene=HORVU5Hr1G046550 | 630.6 | CRagged | 3 | 1173.2262 | 1173.2383 | K16(xgly / 324.1056) | 1.6e-11 | 10.78 | 404.6 | 9.7e-11
Spectrum (double click to dock / undock)
R.GIHNLNLNNAASIPSK[+324]CNVNVPYTISPDID.C z=3,scan#=sample=1 period=1 cycle=791 experiment=21,scan time=13.7454
GIHNLNLNNAASIPSKCNVNVPYTISPDID
Intensity
m/z
m/z errors, tol=75.0 ppm
Protein Coverage (double click to dock / undock)
Protein Coverage
Sample name
Protein name
Coverage summary

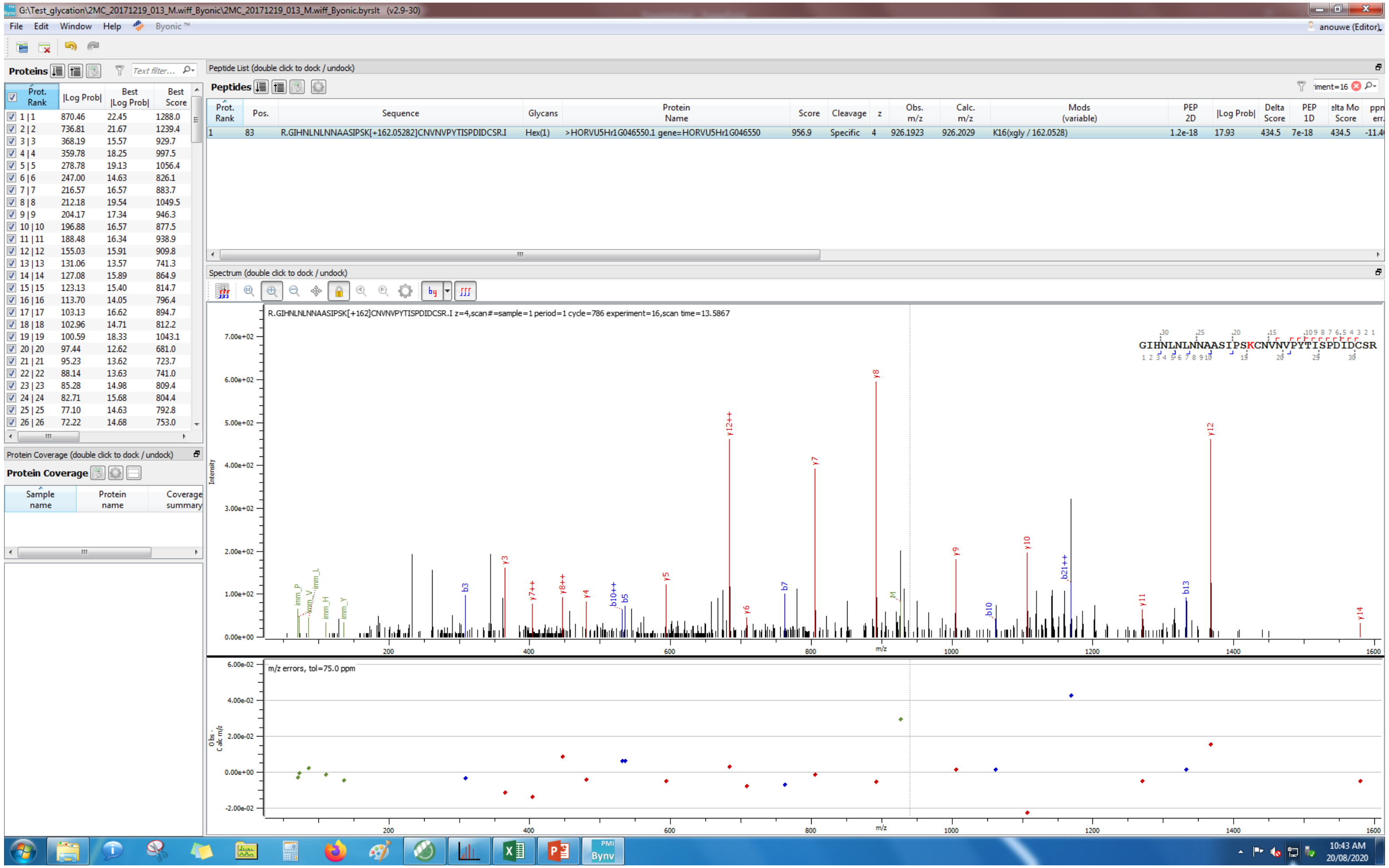
G:\Test_glycation\2MC_20171219_013_M.wiff_Byonic\2MC_20171219_013_M.wiff_Byonic.byrslt (v2.9-30)
File Edit Window Help Byonic™
Proteins
Peptide List (double click to dock / undock)
Peptides
R.GIHNLNLNNAASIPSK[+162.05282]CNVNVPYTISPDIDCSR.I
Hex(1)
>HORVU5Hr1G046550.1 gene=HORVU5Hr1G046550
K16(xgly / 162.0528)
Spectrum (double click to dock / undock)
R.GIHNLNLNNAASIPSK[+162]CNVNVPYTISPDIDCSR.I z=4,scan#=sample=1 period=1 cycle=786 experiment=16,scan time=13.5867
GIHNLNLNNAASIPSKCNVNVPYTISPDIDCSR
Protein Coverage (double click to dock / undock)
Protein Coverage
Sample name
Protein name
Coverage summary
m/z errors, tol=75.0 ppm
m/z
Intensity
10:43 AM
20/08/2020

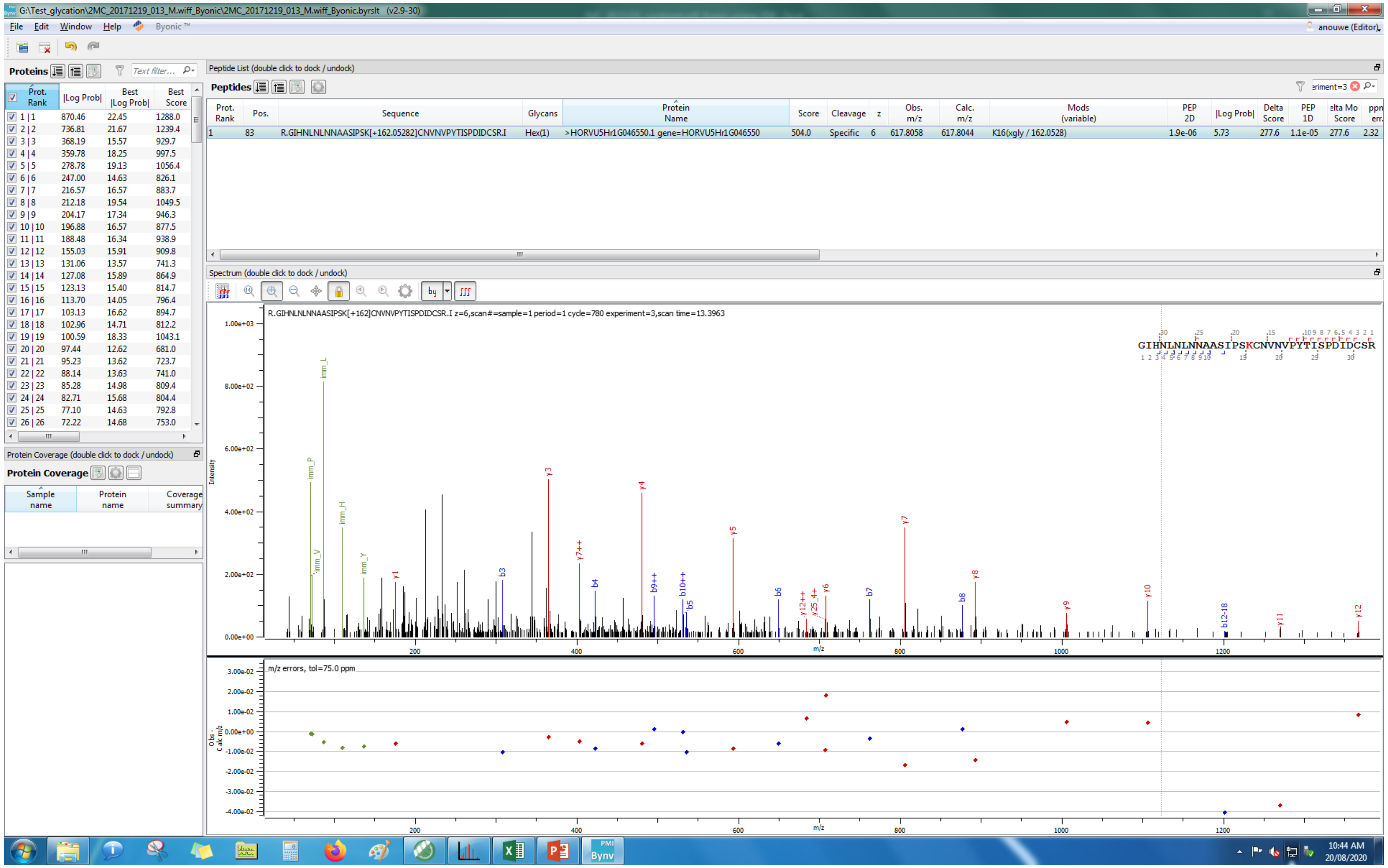
G:\Test_glycation\2MC_20171219_013_M.wiff_Byonic\2MC_20171219_013_M.wiff_Byonic.byrslt (v2.9-30)
File Edit Window Help Byonic™
Proteins
Peptide List (double click to dock / undock)
Peptides
R.GIHNLNLNNAASIPSK[+162.05282]CNVNPYTISPDIDCSR.I
Hex(1)
>HORVU5Hr1G046550.1 gene=HORVU5Hr1G046550
504.0 Specific 6 617.8058 617.8044
K16(xglγ / 162.0528)
Spectrum (double click to dock / undock)
R.GIHNLNLNNAASIPSK[+162]CNVNPYTISPDIDCSR.I z=6,scan#=sample=1 period=1 cycle=780 experiment=3,scan time=13.3963
GIHNLNLNNAASIPSKCNVNPYTISPDIDCSR
Intensity
m/z
m/z errors, tol=75.0 ppm
Protein Coverage (double click to dock / undock)
Protein Coverage
Sample name
Protein name
Coverage summary
10:44 AM
20/08/2020

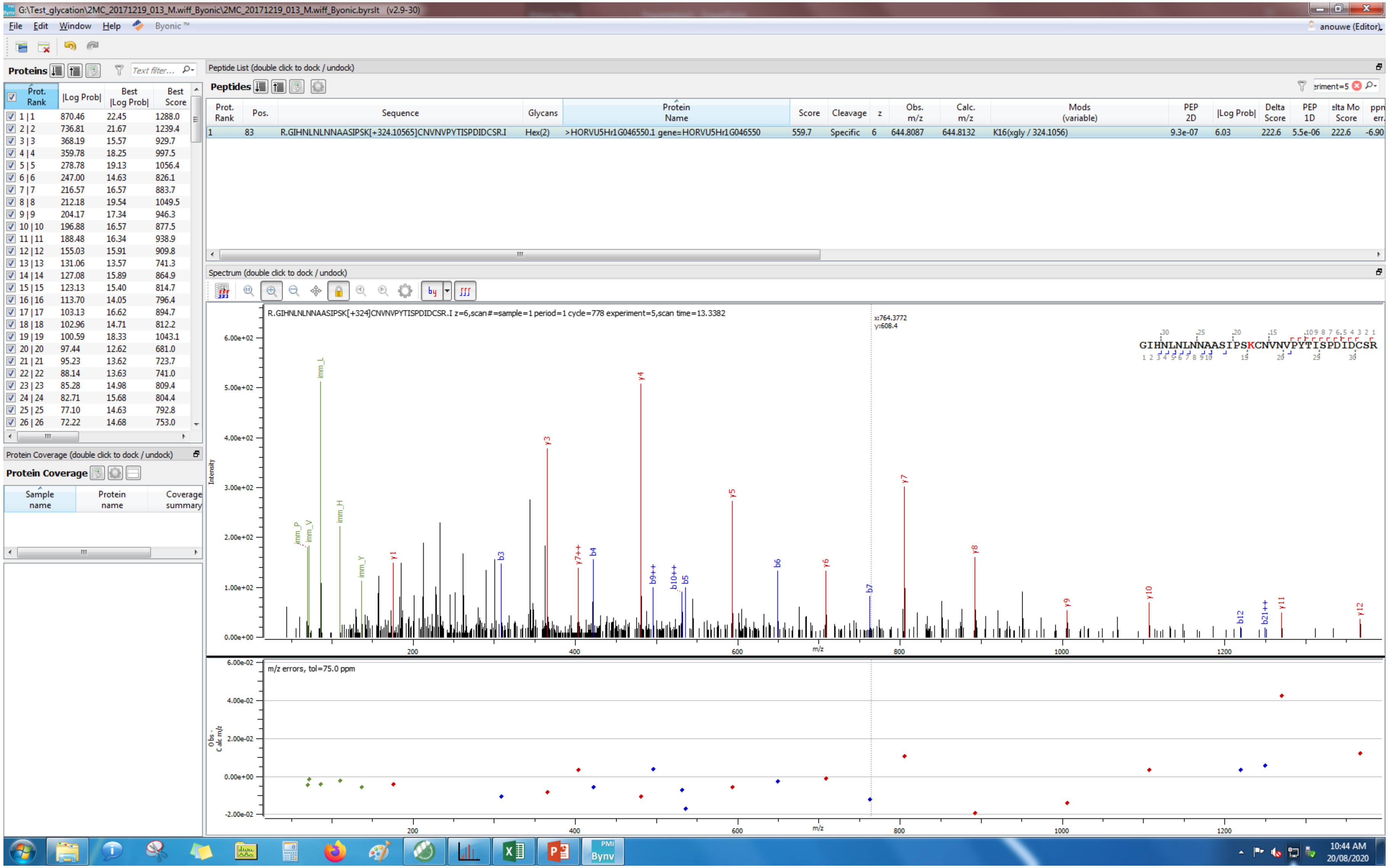
G:\Test_glycation\2MC_20171219_013_M.wiff_Byonic\2MC_20171219_013_M.wiff_Byonic.byrslt (v2.9-30)
File Edit Window Help Byonic™
anouwe (Editor)
Proteins
Peptide List (double click to dock / undock)
Peptides
Prot. Rank | Pos. | Sequence | Glycans | Protein Name | Score | Cleavage | z | Obs. m/z | Calc. m/z | Mods (variable) | PEP 2D | |Log Prob| | Delta Score | PEP 1D | Delta Mod Score | ppm err.
1 | 83 | R.GIHNLNLNNAASIPSK[+324.10565]CNVNVPYTISPDIDCSR.I | Hex(2) | >HORVU5Hr1G046550.1 gene=HORVU5Hr1G046550 | 559.7 | Specific | 6 | 644.8087 | 644.8132 | K16(xgly / 324.1056) | 9.3e-07 | 6.03 | 222.6 | 5.5e-06 | 222.6 | -6.90
Prot. Rank | |Log Prob| | Best |Log Prob| | Best Score
1 | 1 870.46 22.45 1288.0
2 | 2 736.81 21.67 1239.4
3 | 3 368.19 15.57 929.7
4 | 4 359.78 18.25 997.5
5 | 5 278.78 19.13 1056.4
6 | 6 247.00 14.63 826.1
7 | 7 216.57 16.57 883.7
8 | 8 212.18 19.54 1049.5
9 | 9 204.17 17.34 946.3
10 | 10 196.88 16.57 877.5
11 | 11 188.48 16.34 938.9
12 | 12 155.03 15.91 909.8
13 | 13 131.06 13.57 741.3
14 | 14 127.08 15.89 864.9
15 | 15 123.13 15.40 814.7
16 | 16 113.70 14.05 796.4
17 | 17 103.13 16.62 894.7
18 | 18 102.96 14.71 812.2
19 | 19 100.59 18.33 1043.1
20 | 20 97.44 12.62 681.0
21 | 21 95.23 13.62 723.7
22 | 22 88.14 13.63 741.0
23 | 23 85.28 14.98 809.4
24 | 24 82.71 15.68 804.4
25 | 25 77.10 14.63 792.8
26 | 26 72.22 14.68 753.0
Protein Coverage (double click to dock / undock)
Protein Coverage
Sample name | Protein name | Coverage summary
Spectrum (double click to dock / undock)
R.GIHNLNLNNAASIPSK[+324]CNVNVPYTISPDIDCSR.I z=6,scan#=sample=1 period=1 cycle=778 experiment=5,scan time=13.3382
x:764.3772
y:608.4
GIHNLNLNNAASIPSKCNVNVPYTISPDIDCSR
Intensity
m/z
m/z errors, tol=75.0 ppm
Obs - Calc m/z
10:44 AM
20/08/2020

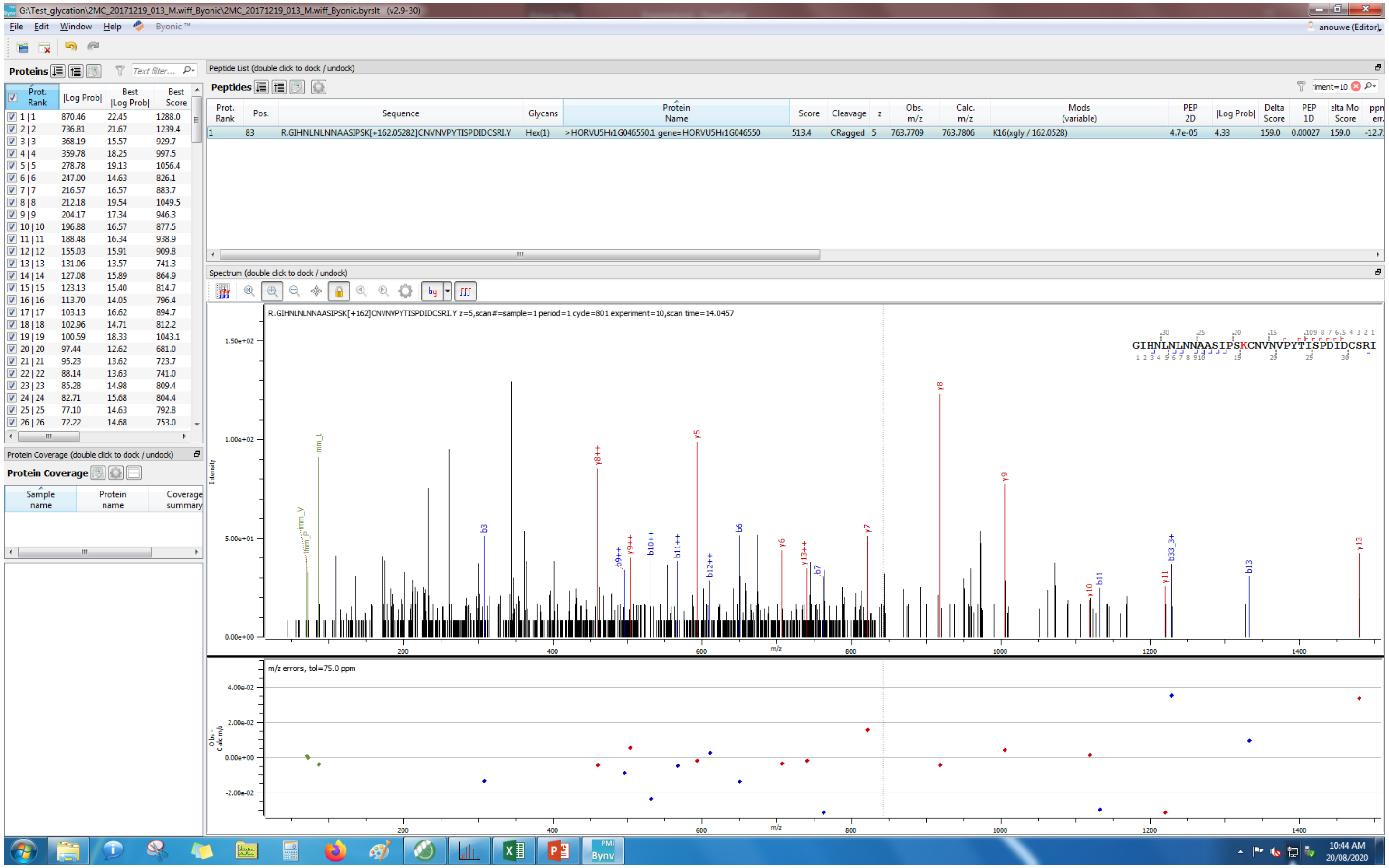
G:\Test_glycation\2MC_20171219_013_M.wiff_Byonic\2MC_20171219_013_M.wiff_Byonic.byrslt (v2.9-30)
File Edit Window Help Byonic
Proteins
Peptide List (double click to dock / undock)
Peptides
R.GIHNLNLNNAASIPSK[+162.05282]CNVNVPYTISPDIDCSRI.Y
Hex(1)
>HORVU5Hr1G046550.1 gene=HORVU5Hr1G046550
K16(xgly / 162.0528)
Spectrum (double click to dock / undock)
R.GIHNLNLNNAASIPSK[+162]CNVNVPYTISPDIDCSRI.Y z=5,scan#=sample=1 period=1 cycle=801 experiment=10,scan time=14.0457
GIHNLNLNNAASIPSKCNVNVPYTISPDIDCSRI
Intensity
m/z
m/z errors, tol=75.0 ppm
Protein Coverage (double click to dock / undock)
Protein Coverage

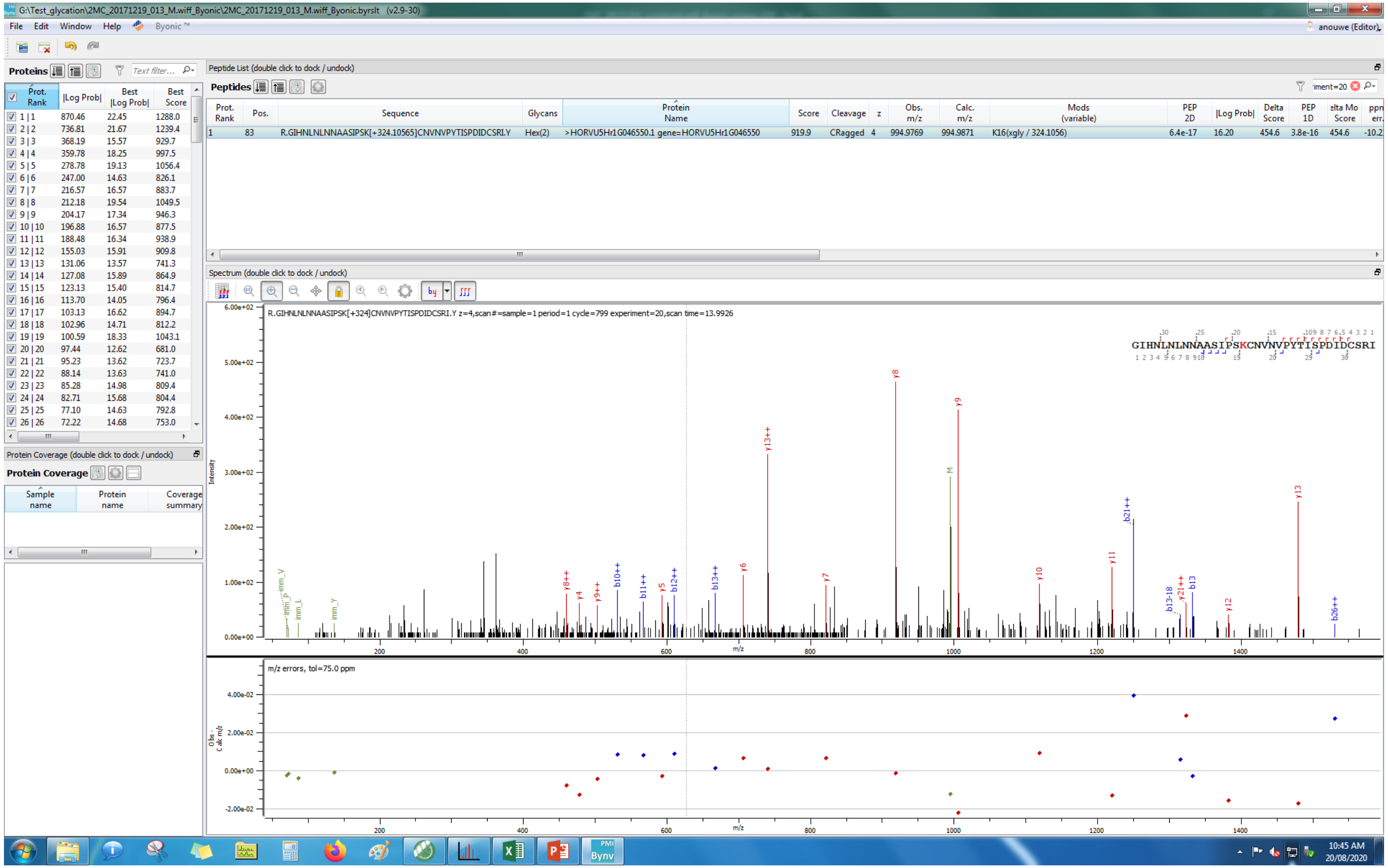

G:\Test_glycation\2MC_20171219_013_M.wiff_Byonic\2MC_20171219_013_M.wiff_Byonic.byrslt (v2.9-30)
File Edit Window Help Byonic
Proteins
Peptide List (double click to dock / undock)
Peptides
R.GIHNLNLNNAASIPSK[+324.10565]CNVNVPYTISPDIDCSRI.Y
Hex(2)
>HORVU5Hr1G046550.1 gene=HORVU5Hr1G046550
K16(xgly / 324.1056)
Spectrum (double click to dock / undock)
R.GIHNLNLNNAASIPSK[+324]CNVNVPYTISPDIDCSRI.Y z=4,scan#=sample=1 period=1 cycle=799 experiment=20,scan time=13.9926
GIHNLNLNNAASIPSKCNVNVPYTISPDIDCSRI
Protein Coverage (double click to dock / undock)
Protein Coverage
m/z errors, tol=75.0 ppm
10:45 AM
20/08/2020

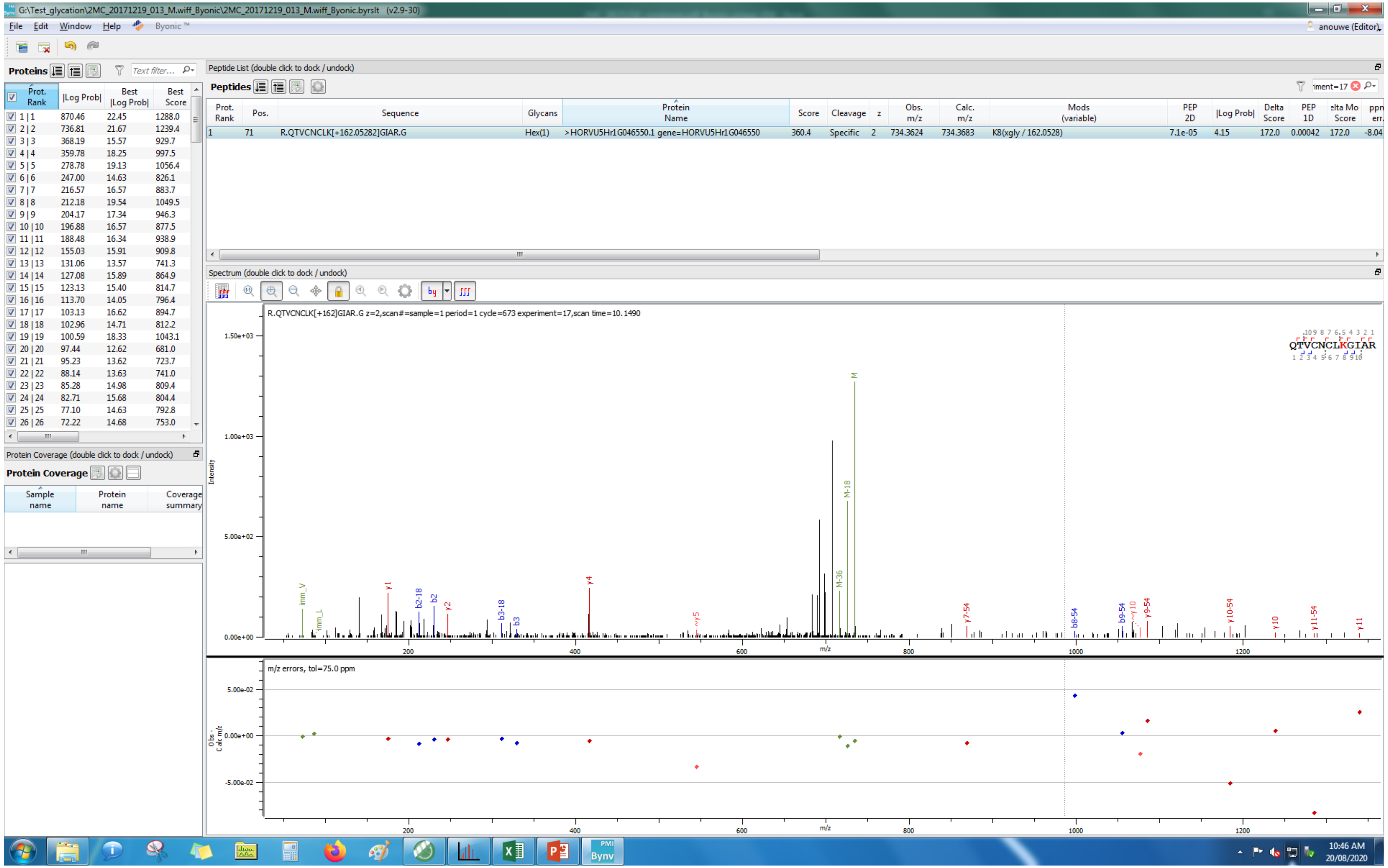
G:\Test_glycation\2MC_20171219_013_M.wiff_Byonic\2MC_20171219_013_M.wiff_Byonic.byrslt (v2.9-30)
File Edit Window Help Byonic™
anouwe (Editor)
Proteins
Peptide List (double click to dock / undock)
Peptides
Prot. Rank | Pos. | Sequence | Glycans | Protein Name | Score | Cleavage | z | Obs. m/z | Calc. m/z | Mods (variable) | PEP 2D | |Log Prob| | Delta Score | PEP 1D | elta Mo Score | ppm err.
1 | 71 | R.QTVCNCLK[+162.05282]GIAR.G | Hex(1) | >HORVU5Hr1G046550.1 gene=HORVU5Hr1G046550 | 360.4 | Specific | 2 | 734.3624 | 734.3683 | K8(xgly / 162.0528) | 7.1e-05 | 4.15 | 172.0 | 0.00042 | 172.0 | -8.04
Spectrum (double click to dock / undock)
R.QTVCNCLK[+162]GIAR.G z=2,scan#=sample=1 period=1 cycle=673 experiment=17,scan time=10.1490
QTVCNCLKGIAR
Intensity
m/z
m/z errors, tol=75.0 ppm
Protein Coverage (double click to dock / undock)
Protein Coverage
Sample name | Protein name | Coverage summary
10:46 AM
20/08/2020

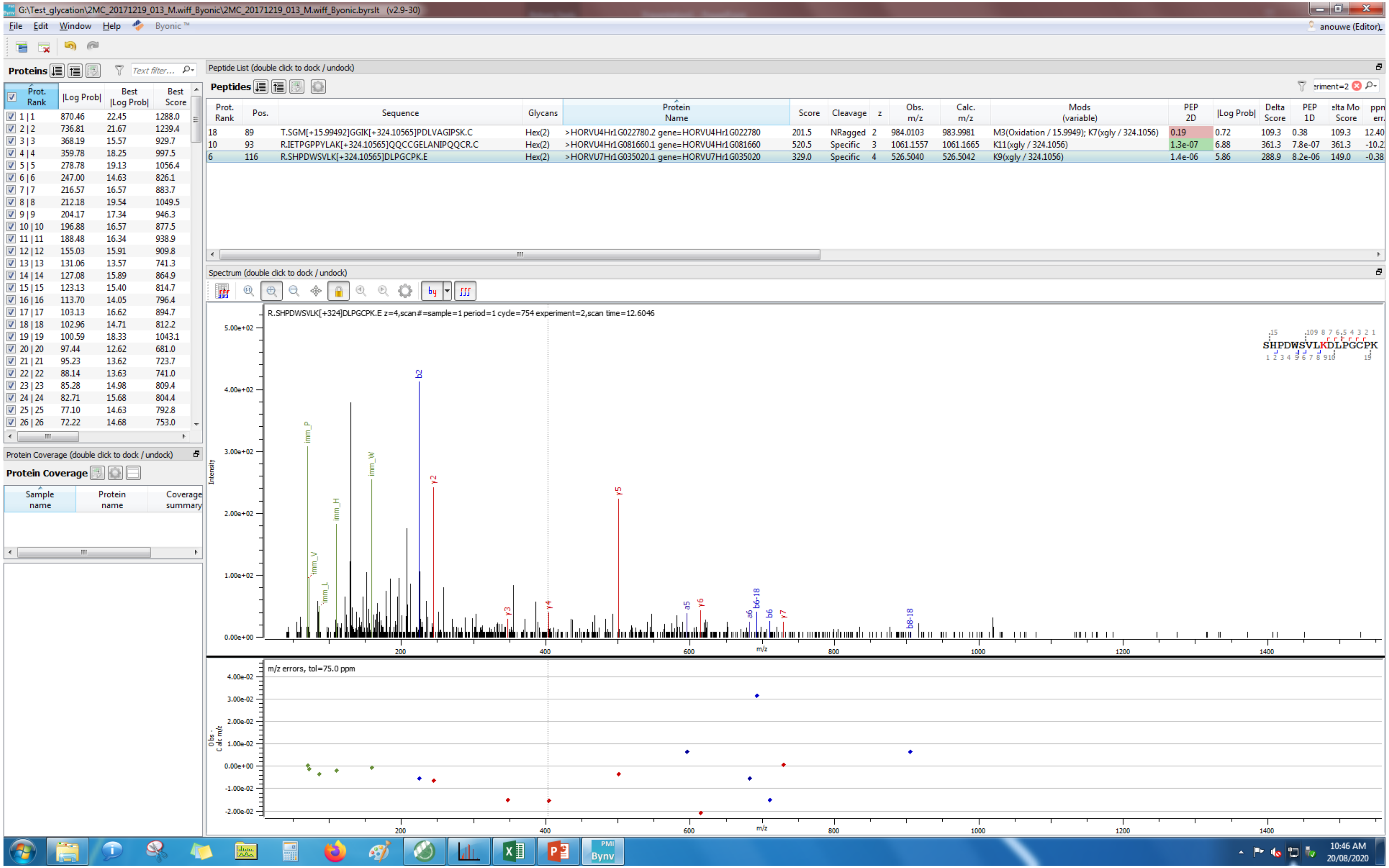
G:\Test_glycation\2MC_20171219_013_M.wiff_Byonic\2MC_20171219_013_M.wiff_Byonic.byrslt (v2.9-30)
File Edit Window Help Byonic™
anouwe (Editor)
Proteins
Peptide List (double click to dock / undock)
Peptides
Prot. Rank | Pos. | Sequence | Glycans | Protein Name | Score | Cleavage | z | Obs. m/z | Calc. m/z | Mods (variable) | PEP 2D | |Log Prob| | Delta Score | PEP 1D | Delta Mod Score | ppm err.
18 | 89 | T.SGM[+15.99492]GGIK[+324.10565]PDLVAGIPSK.C | Hex(2) | >HORVU4Hr1G022780.2 gene=HORVU4Hr1G022780 | 201.5 | NRagged | 2 | 984.0103 | 983.9981 | M3(Oxidation / 15.9949); K7(xgly / 324.1056) | 0.19 | 0.72 | 109.3 | 0.38 | 109.3 | 12.40
10 | 93 | R.IETPGPPYLAK[+324.10565]QQCCGELANIPQQCR.C | Hex(2) | >HORVU4Hr1G081660.1 gene=HORVU4Hr1G081660 | 520.5 | Specific | 3 | 1061.1557 | 1061.1665 | K11(xgly / 324.1056) | 1.3e-07 | 6.88 | 361.3 | 7.8e-07 | 361.3 | -10.2
6 | 116 | R.SHPDWSVLK[+324.10565]DLPGCPK.E | Hex(2) | >HORVU7Hr1G035020.1 gene=HORVU7Hr1G035020 | 329.0 | Specific | 4 | 526.5040 | 526.5042 | K9(xgly / 324.1056) | 1.4e-06 | 5.86 | 288.9 | 8.2e-06 | 149.0 | -0.38
Spectrum (double click to dock / undock)
R.SHPDWSVLK[+324]DLPGCPK.E z=4,scan#=sample=1 period=1 cycle=754 experiment=2,scan time=12.6046
SHPDWSVLKDLPGCPK
Intensity
m/z
m/z errors, tol=75.0 ppm
Obs - Calc m/z
Protein Coverage (double click to dock / undock)
Protein Coverage
Sample name | Protein name | Coverage summary
10:46 AM
20/08/2020

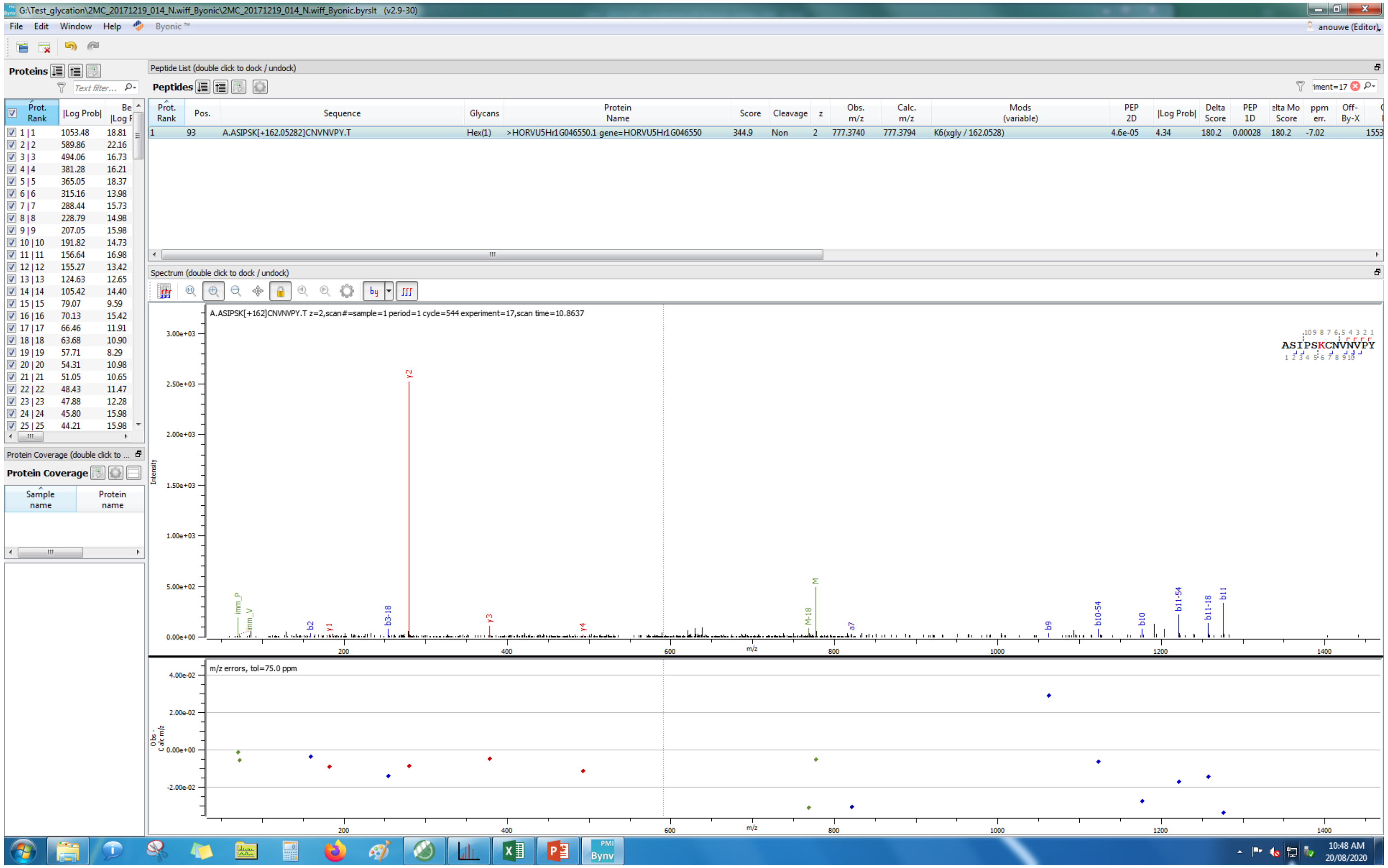
G:\Test_glycation\2MC_20171219_014_N.wiff_Byonic\2MC_20171219_014_N.wiff_Byonic.byrslt (v2.9-30)
File Edit Window Help Byonic
Proteins
Peptide List (double click to dock / undock)
Peptides
A.ASIPSK[+162.05282]CNVNVPY.T
Hex(1)
>HORVU5Hr1G046550.1 gene=HORVU5Hr1G046550
Spectrum (double click to dock / undock)
A.ASIPSK[+162]CNVNVPY.T z=2,scan#=sample=1 period=1 cycle=544 experiment=17,scan time=10.8637
m/z errors, tol=75.0 ppm
Protein Coverage

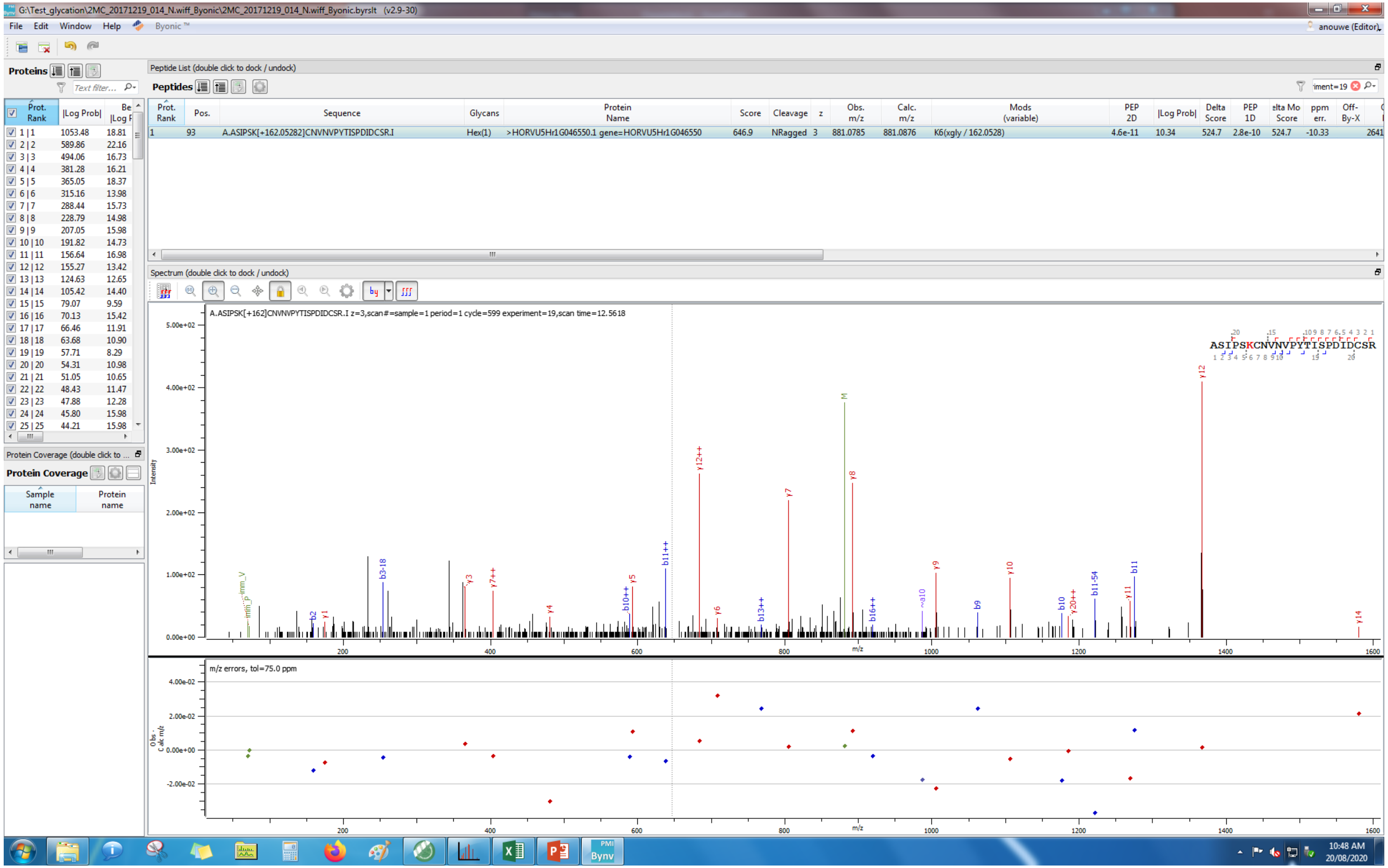
G:\Test_glycation\2MC_20171219_014_N.wiff_Byonic\2MC_20171219_014_N.wiff_Byonic.byrslt (v2.9-30)
File Edit Window Help Byonic ™
anouwe (Editor)
Proteins
Peptide List (double click to dock / undock)
Peptides
Prot. Rank | Pos. | Sequence | Glycans | Protein Name | Score | Cleavage | z | Obs. m/z | Calc. m/z | Mods (variable) | PEP 2D | |Log Prob| | Delta Score | PEP 1D | elta Mo Score | ppm err. | Off-By-X
1 | 93 | A.ASIPSK[+162.05282]CNVNVPYTISPDIDCSR.I | Hex(1) | >HORVU5Hr1G046550.1 gene=HORVU5Hr1G046550 | 646.9 | NRagged | 3 | 881.0785 | 881.0876 | K6(xgly / 162.0528) | 4.6e-11 | 10.34 | 524.7 | 2.8e-10 | 524.7 | -10.33
Spectrum (double click to dock / undock)
A.ASIPSK[+162]CNVNVPYTISPDIDCSR.I z=3,scan#=sample=1 period=1 cycle=599 experiment=19,scan time=12.5618
ASIPSKCNVNVPYTISPDIDCSR
Intensity
m/z
m/z errors, tol=75.0 ppm
Obs-Calc m/z
Protein Coverage (double click to ...
Protein Coverage
Sample name
Protein name
10:48 AM
20/08/2020

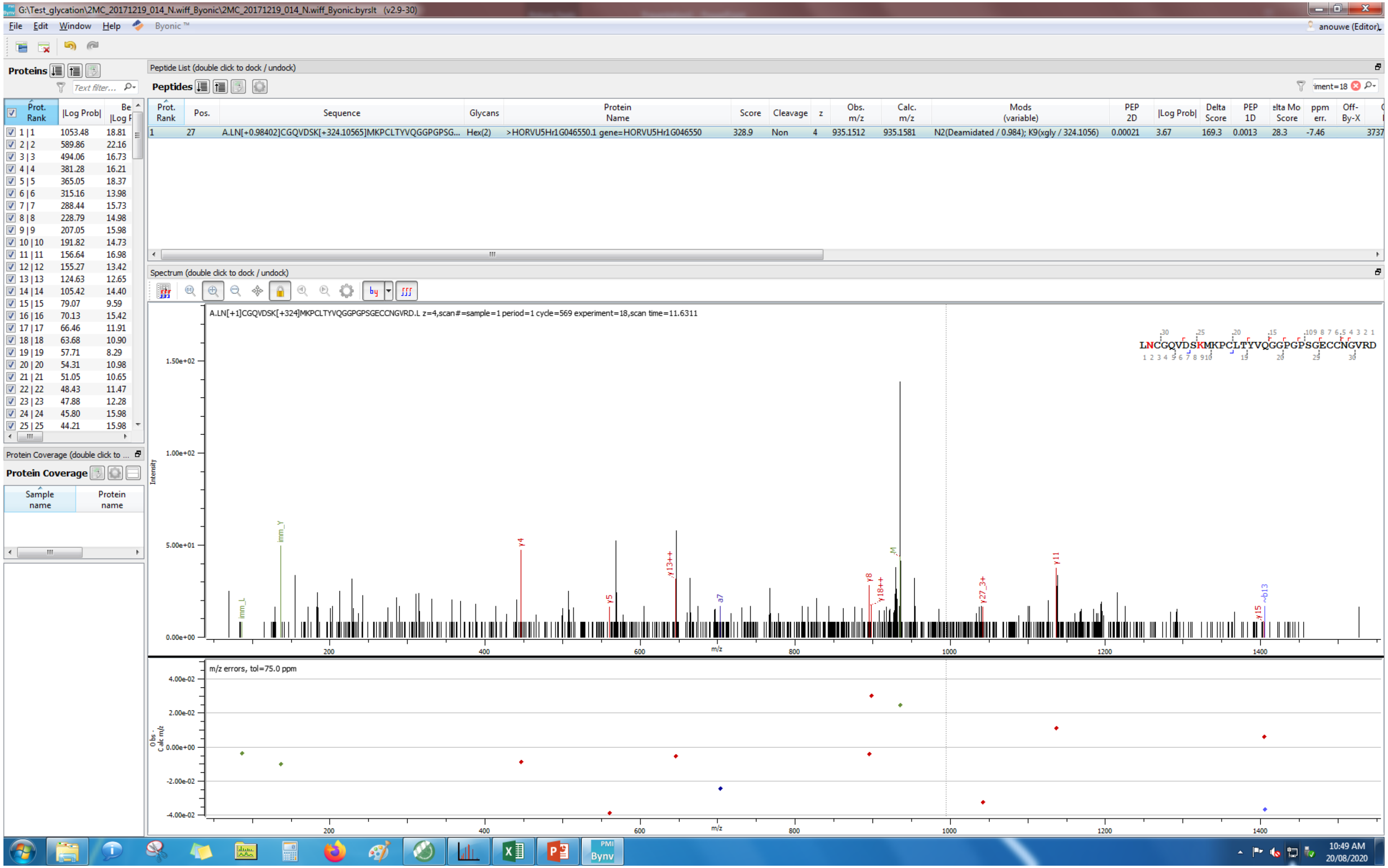
G:\Test_glycation\2MC_20171219_014_N.wiff_Byonic\2MC_20171219_014_N.wiff_Byonic.byrslt (v2.9-30)
File Edit Window Help Byonic™
anouwe (Editor)
Proteins
Peptide List (double click to dock / undock)
Peptides
Prot. Rank | Pos. | Sequence | Glycans | Protein Name | Score | Cleavage | z | Obs. m/z | Calc. m/z | Mods (variable) | PEP 2D | |Log Prob| | Delta Score | PEP 1D | Delta Mod Score | ppm err. | Off-By-X
1 | 27 | A.LN[+0.98402]CGQVDSK[+324.10565]MKPCLTYVQGGPGPSG... | Hex(2) | >HORVU5Hr1G046550.1 gene=HORVU5Hr1G046550 | 328.9 | Non | 4 | 935.1512 | 935.1581 | N2(Deamidated / 0.984); K9(xgly / 324.1056) | 0.00021 | 3.67 | 169.3 | 0.0013 | 28.3 | -7.46
Spectrum (double click to dock / undock)
A.LN[+1]CGQVDSK[+324]MKPCLTYVQGGPGPSGECCNGVRD.L z=4,scan#=sample=1 period=1 cycle=569 experiment=18,scan time=11.6311
LNCGQVDSKMKPCLTYVQGGPGPSGECCNGVRD
Intensity
m/z
m/z errors, tol=75.0 ppm
Obs - Calc m/z
Protein Coverage
Sample name
Protein name
10:49 AM
20/08/2020

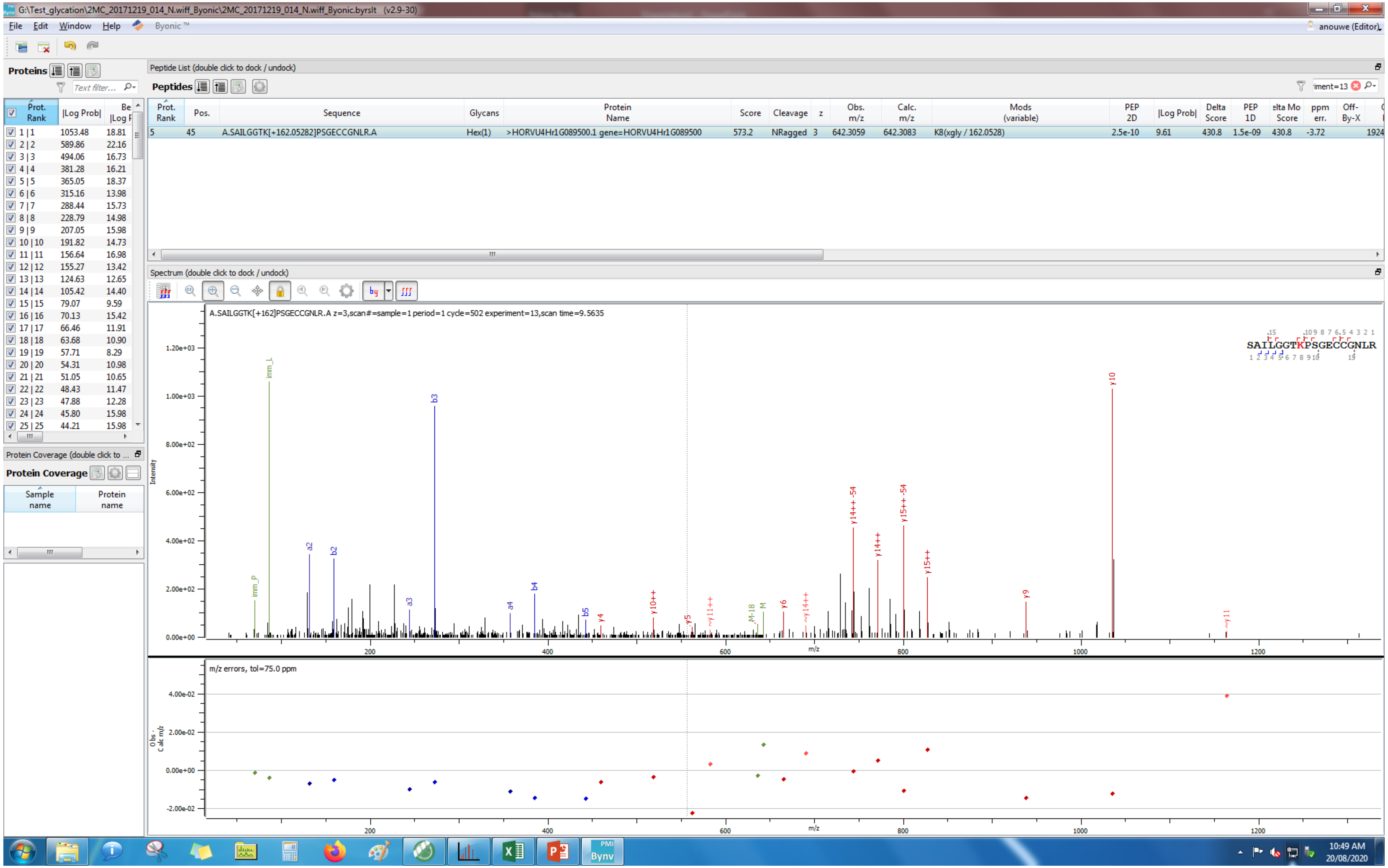
G:\Test_glycation\2MC_20171219_014_N.wiff_Byonic\2MC_20171219_014_N.wiff_Byonic.byrslt (v2.9-30)
File Edit Window Help Byonic™
anouwe (Editor)
Proteins
Peptide List (double click to dock / undock)
Peptides
Prot. Rank | Pos. | Sequence | Glycans | Protein Name | Score | Cleavage | z | Obs. m/z | Calc. m/z | Mods (variable) | PEP 2D | |Log Prob| | Delta Score | PEP 1D | ppm err. | Off-By-X
5 | 45 | A.SAILGGTK[+162.05282]PSGECCGNLR.A | Hex(1) | >HORVU4Hr1G089500.1 gene=HORVU4Hr1G089500 | 573.2 | NRagged | 3 | 642.3059 | 642.3083 | K8(xgly / 162.0528) | 2.5e-10 | 9.61 | 430.8 | 1.5e-09 | 430.8 | -3.72 | 1924
Spectrum (double click to dock / undock)
A.SAILGGTK[+162]PSGECCGNLR.A z=3,scan#=sample=1 period=1 cycle=502 experiment=13,scan time=9.5635
SAILGGTKPSGECCGNLR
Intensity
m/z
m/z errors, tol=75.0 ppm
Protein Coverage (double click to ...
Protein Coverage
Sample name
Protein name
10:49 AM
20/08/2020

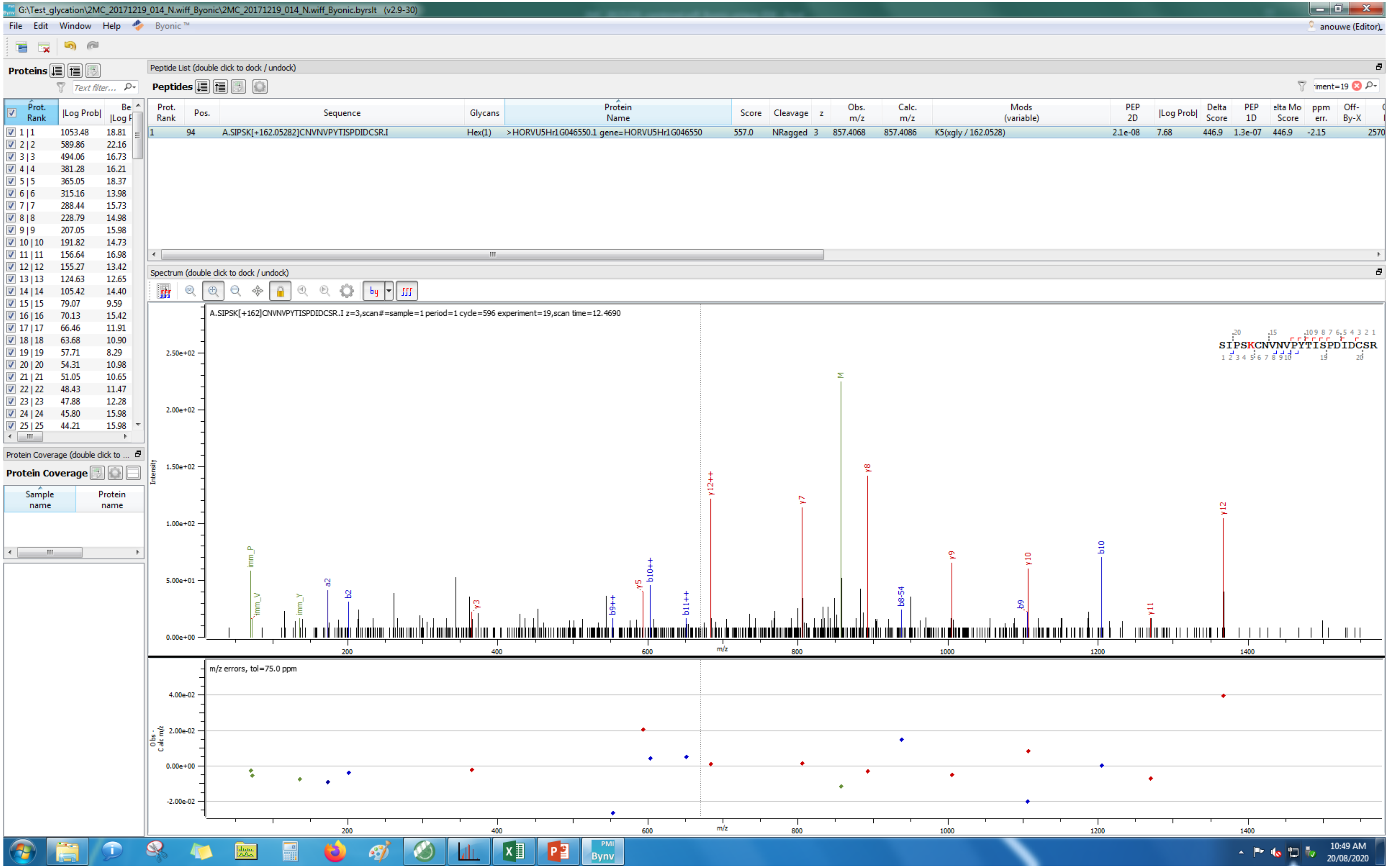
G:\Test_glycation\2MC_20171219_014_N.wiff_Byonic\2MC_20171219_014_N.wiff_Byonic.byrslt (v2.9-30)
File Edit Window Help Byonic ™
anouwe (Editor)
Proteins
Peptide List (double click to dock / undock)
Peptides
Prot. Rank | Pos. | Sequence | Glycans | Protein Name | Score | Cleavage | z | Obs. m/z | Calc. m/z | Mods (variable) | PEP 2D | |Log Prob| | Delta Score | PEP 1D | Delta Mod Score | ppm err. | Off-By-X
1 | 94 | A.SIPSK[+162.05282]CNVNVPYTISPDIDCSR.I | Hex(1) | >HORVU5Hr1G046550.1 gene=HORVU5Hr1G046550 | 557.0 | NRagged | 3 | 857.4068 | 857.4086 | K5(xgly / 162.0528) | 2.1e-08 | 7.68 | 446.9 | 1.3e-07 | 446.9 | -2.15
Spectrum (double click to dock / undock)
A.SIPSK[+162]CNVNVPYTISPDIDCSR.I z=3,scan#=sample=1 period=1 cycle=596 experiment=19,scan time=12.4690
SIPSKCNVNVPYTISPDIDCSR
Intensity
m/z
m/z errors, tol=75.0 ppm
Obs. - Calc m/z
Protein Coverage (double click to ...
Protein Coverage
Sample name
Protein name
10:49 AM
20/08/2020

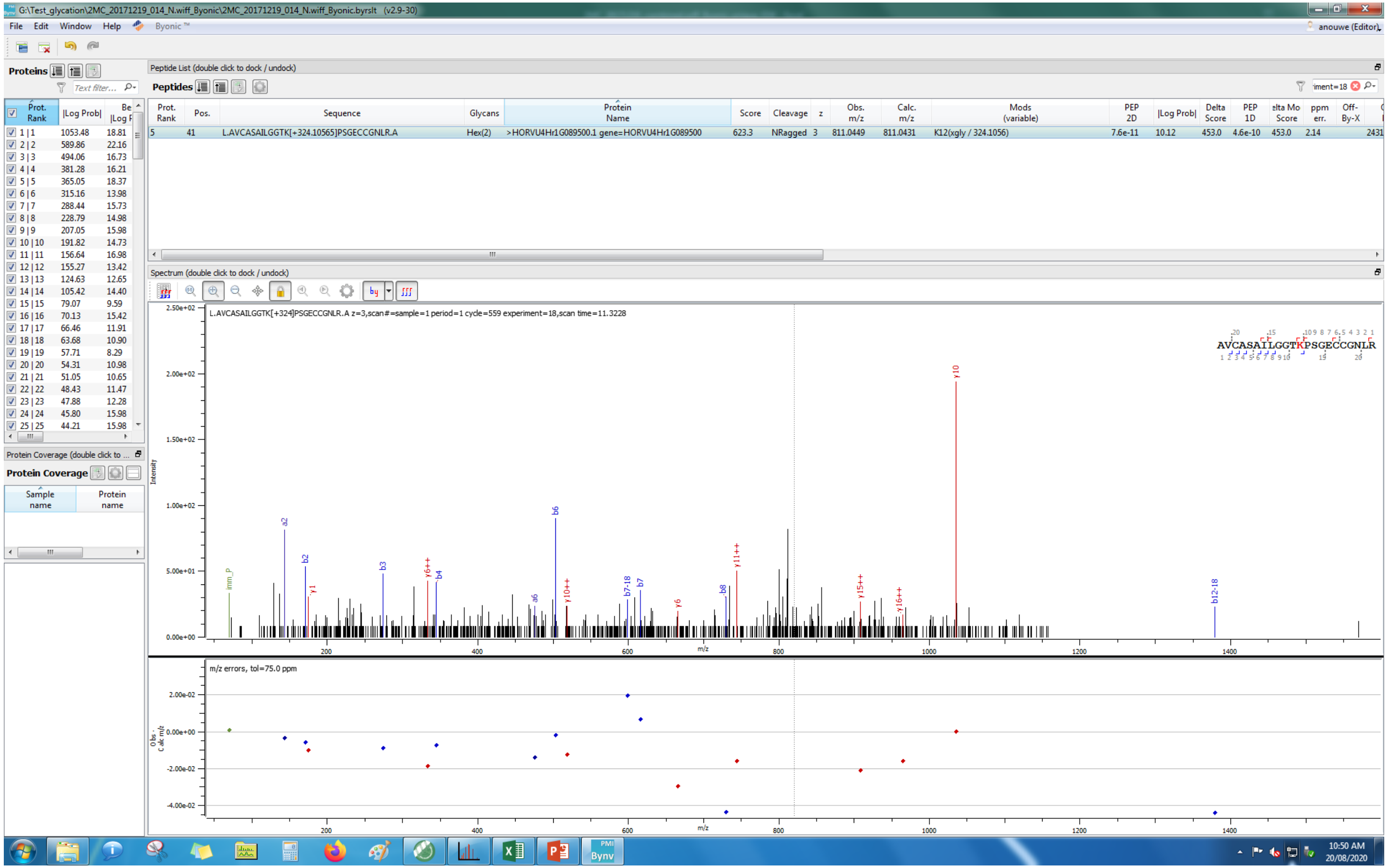
G:\Test_glycation\2MC_20171219_014_N.wiff_Byonic\2MC_20171219_014_N.wiff_Byonic.byrslt (v2.9-30)
File Edit Window Help Byonic
Proteins
Peptide List (double click to dock / undock)
Peptides
Prot. Rank | Pos. | Sequence | Glycans | Protein Name | Score | Cleavage | z | Obs. m/z | Calc. m/z | Mods (variable) | PEP 2D | |Log Prob| | Delta Score | PEP 1D | Delta Mod Score | ppm err. | Off-By-X
5 | 41 | L.AVCASAILGGTK[+324.10565]PSGECCGNLR.A | Hex(2) | >HORVU4Hr1G089500.1 gene=HORVU4Hr1G089500 | 623.3 | NRagged | 3 | 811.0449 | 811.0431 | K12(xgly / 324.1056) | 7.6e-11 | 10.12 | 453.0 | 4.6e-10 | 453.0 | 2.14
Spectrum (double click to dock / undock)
L.AVCASAILGGTK[+324]PSGECCGNLR.A z=3,scan#=sample=1 period=1 cycle=559 experiment=18,scan time=11.3228
AVCASAILGGTKPSGECCGNLR
m/z errors, tol=75.0 ppm
Protein Coverage
Sample name
Protein name
10:50 AM
20/08/2020

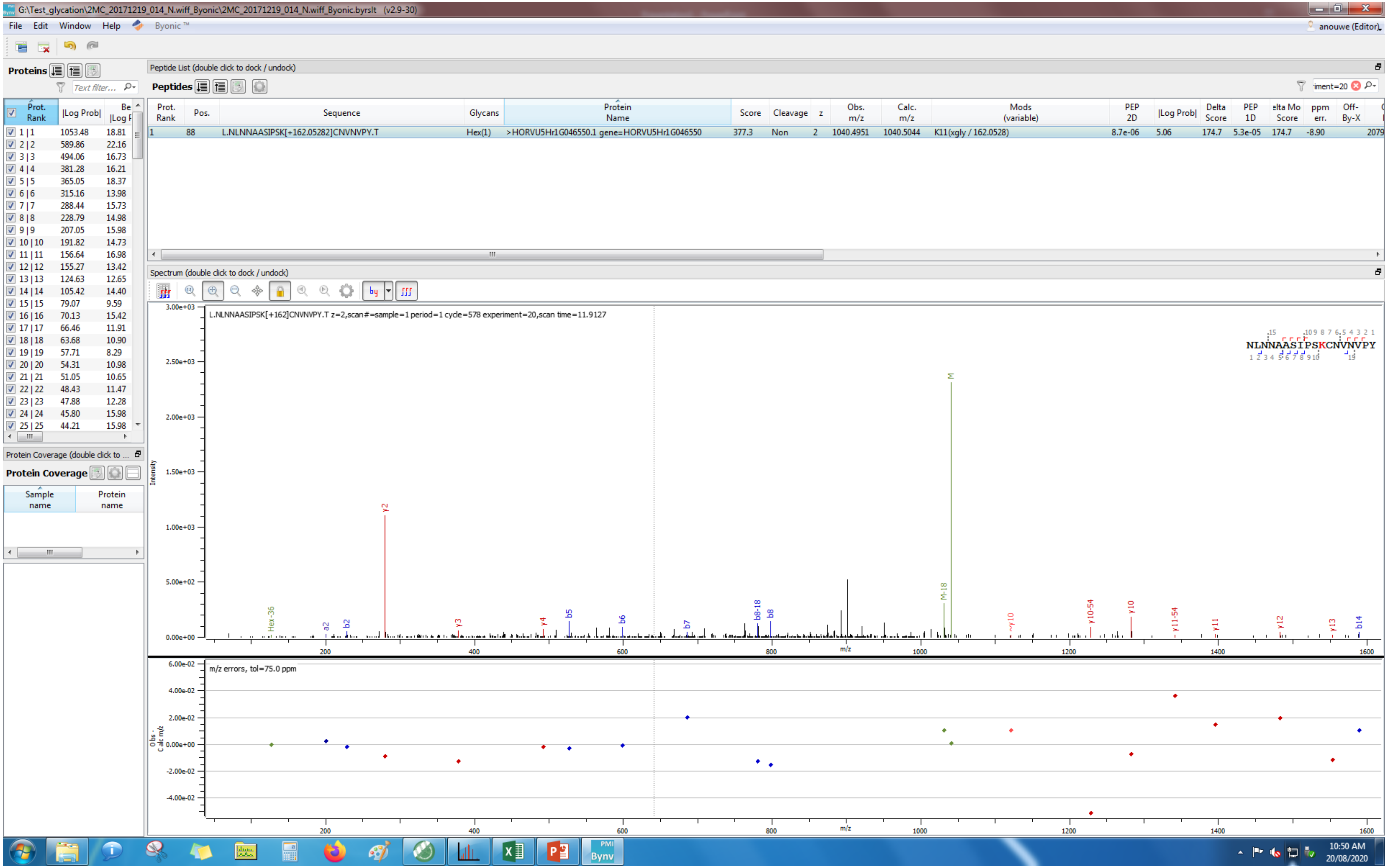
G:\Test_glycation\2MC_20171219_014_N.wiff_Byonic\2MC_20171219_014_N.wiff_Byonic.byrslt (v2.9-30)
File Edit Window Help Byonic ™
anouwe (Editor)
Proteins
Peptide List (double click to dock / undock)
Peptides
Prot. Rank | Pos. | Sequence | Glycans | Protein Name | Score | Cleavage | z | Obs. m/z | Calc. m/z | Mods (variable) | PEP 2D | |Log Prob| | Delta Score | PEP 1D | Delta Mod Score | ppm err. | Off-By-X
1 | 88 | L.NLNNAASIPSK[+162.05282]CNVNVPY.T | Hex(1) | >HORVU5Hr1G046550.1 gene=HORVU5Hr1G046550 | 377.3 | Non | 2 | 1040.4951 | 1040.5044 | K11(xgly / 162.0528) | 8.7e-06 | 5.06 | 174.7 | 5.3e-05 | 174.7 | -8.90 | 2079
Spectrum (double click to dock / undock)
L.NLNNAASIPSK[+162]CNVNVPY.T z=2,scan#=sample=1 period=1 cycle=578 experiment=20,scan time=11.9127
NLNNAASIPSKCNVNVPY
Intensity
m/z
m/z errors, tol=75.0 ppm
Obs - Calc m/z
Protein Coverage (double click to dock / undock)
Protein Coverage
Sample name
Protein name
10:50 AM
20/08/2020

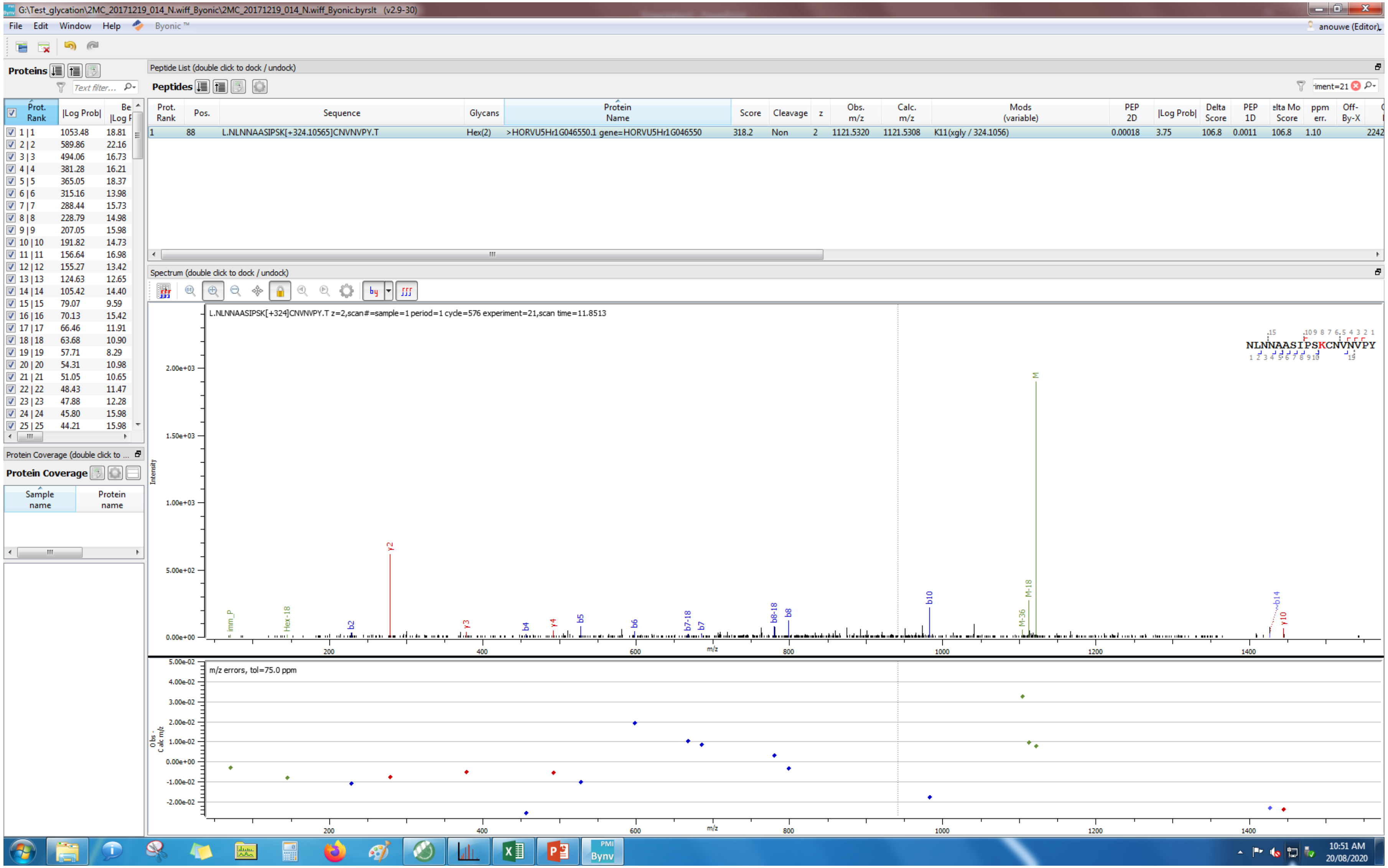

G:\Test_glycation\2MC_20171219_014_N.wiff_Byonic\2MC_20171219_014_N.wiff_Byonic.byrslt (v2.9-30)
File Edit Window Help Byonic ™
anouwe (Editor)
Proteins
Peptide List (double click to dock / undock)
Peptides
Prot. Rank | Pos. | Sequence | Glycans | Protein Name | Score | Cleavage | z | Obs. m/z | Calc. m/z | Mods (variable) | PEP 2D | |Log Prob| | Delta Score | PEP 1D | Delta Mod Score | ppm err. | Off-By-X
1 | 88 | L.NLNNAASIPSK[+324.10565]CNVNVPY.T | Hex(2) | >HORVU5Hr1G046550.1 gene=HORVU5Hr1G046550 | 318.2 | Non | 2 | 1121.5320 | 1121.5308 | K11(xgly / 324.1056) | 0.00018 | 3.75 | 106.8 | 0.0011 | 106.8 | 1.10
Spectrum (double click to dock / undock)
L.NLNNAASIPSK[+324]CNVNVPY.T z=2,scan#=sample=1 period=1 cycle=576 experiment=21,scan time=11.8513
NLNNAASIPSKCNVNVPY
Intensity
m/z
m/z errors, tol=75.0 ppm
Obs - Calc m/z
Protein Coverage (double click to ...
Protein Coverage
Sample name
Protein name
10:51 AM
20/08/2020

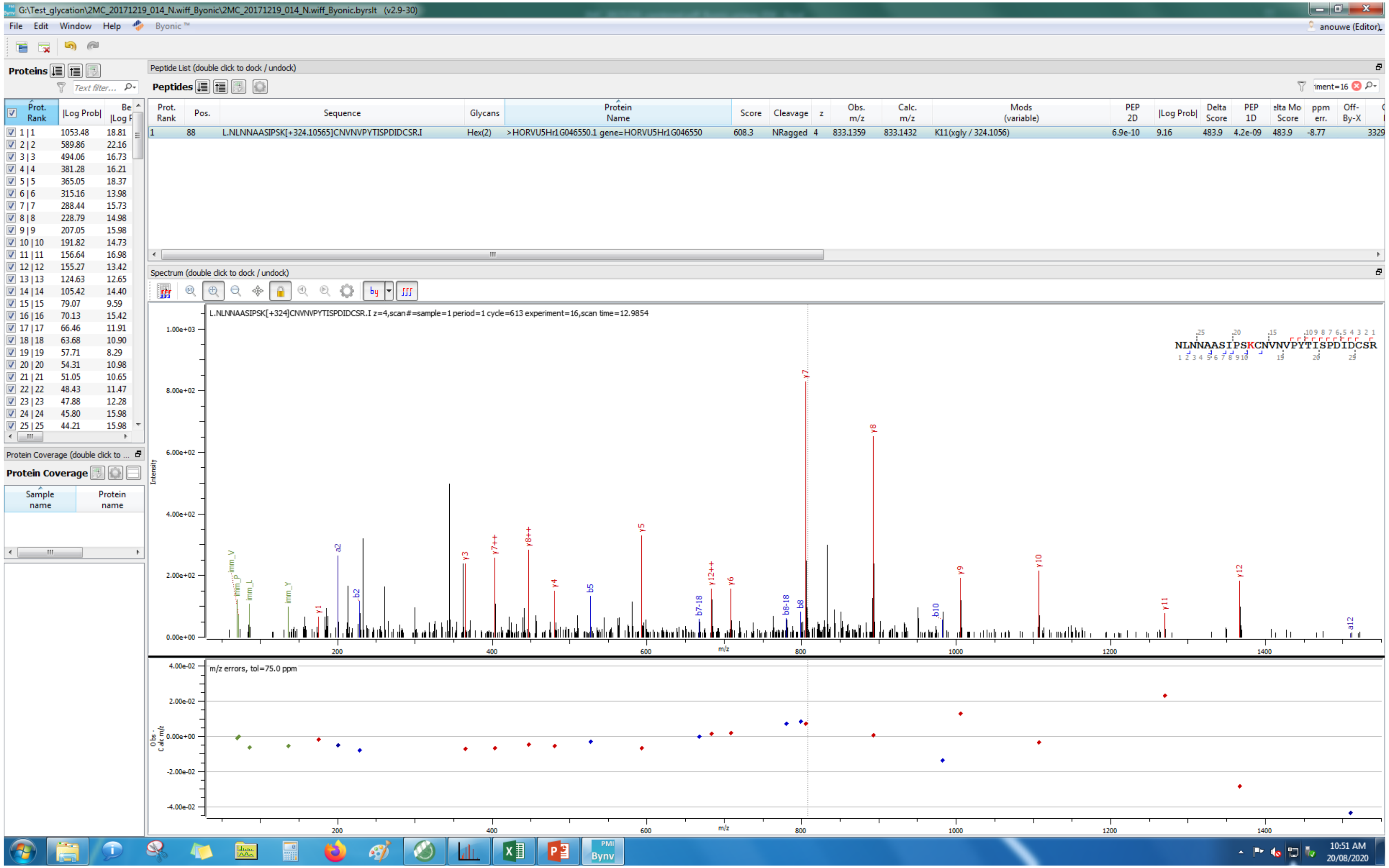
G:\Test_glycation\2MC_20171219_014_N.wiff_Byonic\2MC_20171219_014_N.wiff_Byonic.byrslt (v2.9-30)
File Edit Window Help Byonic™
anouwe (Editor)
Proteins
Peptide List (double click to dock / undock)
Peptides
Prot. Rank | Pos. | Sequence | Glycans | Protein Name | Score | Cleavage | z | Obs. m/z | Calc. m/z | Mods (variable) | PEP 2D | |Log Prob| | Delta Score | PEP 1D
1 | 88 | L.NLNNAASIPSK[+324.10565]CNVNVPYTISPDIDCSR.I | Hex(2) | >HORVU5Hr1G046550.1 gene=HORVU5Hr1G046550 | 608.3 | NRagged | 4 | 833.1359 | 833.1432 | K11(xgly / 324.1056) | 6.9e-10 | 9.16 | 483.9 | 4.2e-09
Spectrum (double click to dock / undock)
L.NLNNAASIPSK[+324]CNVNVPYTISPDIDCSR.I z=4,scan#=sample=1 period=1 cycle=613 experiment=16,scan time=12.9854
NLNNAASIPSKCNVNVPYTISPDIDCSR
Intensity
m/z
m/z errors, tol=75.0 ppm
Protein Coverage
Sample name
Protein name
10:51 AM
20/08/2020

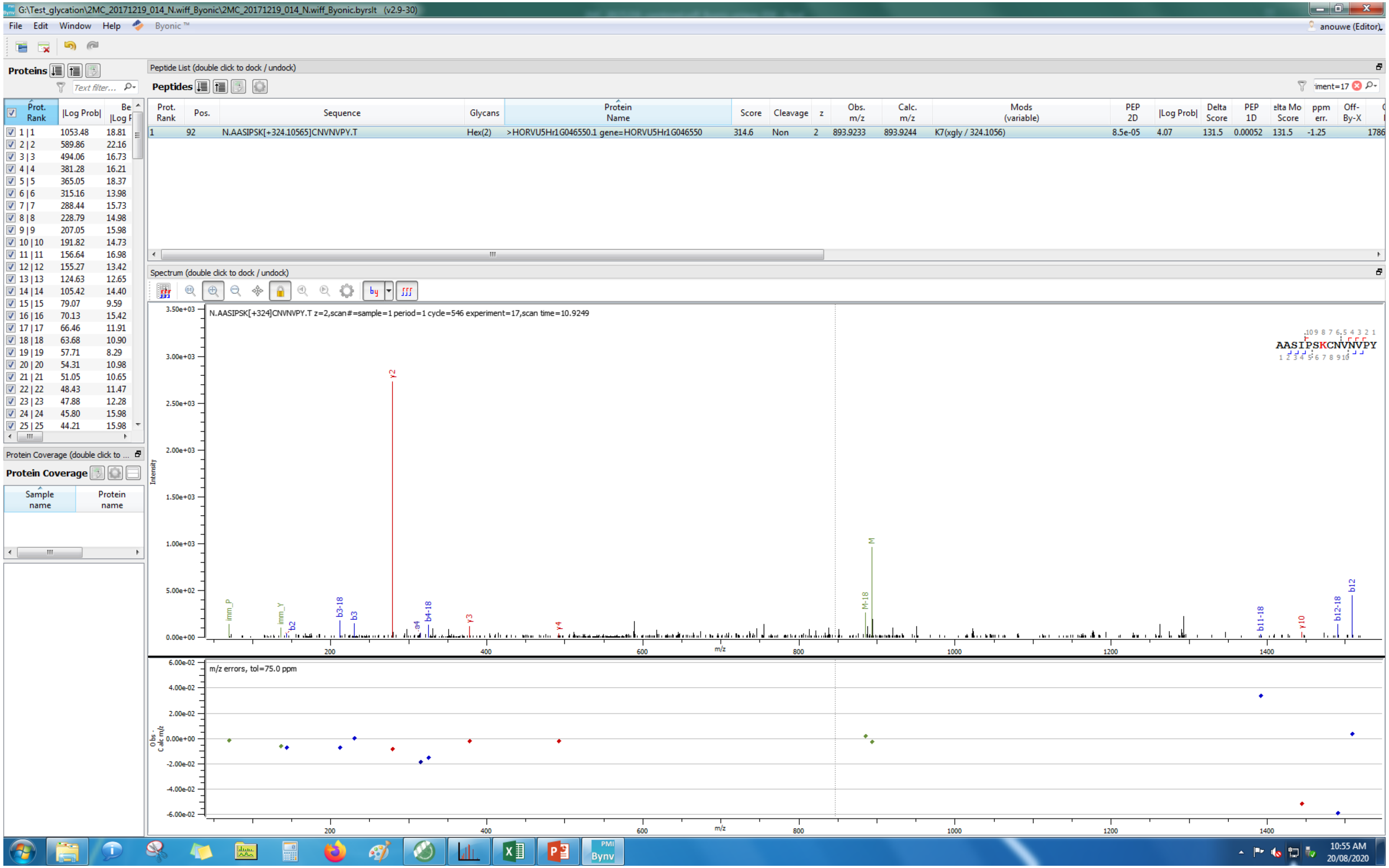
G:\Test_glycation\2MC_20171219_014_N.wiff_Byonic\2MC_20171219_014_N.wiff_Byonic.byrslt (v2.9-30)
File Edit Window Help Byonic™
Proteins
Peptide List (double click to dock / undock)
Peptides
Prot. Rank | Pos. | Sequence | Glycans | Protein Name | Score | Cleavage | z | Obs. m/z | Calc. m/z | Mods (variable) | PEP 2D | |Log Prob| | Delta Score | PEP 1D | ppm err.
1 | 92 | N.AASIPSK[+324.10565]CNVNVPY.T | Hex(2) | >HORVU5Hr1G046550.1 gene=HORVU5Hr1G046550 | 314.6 | Non | 2 | 893.9233 | 893.9244 | K7(xgly / 324.1056) | 8.5e-05 | 4.07 | 131.5 | 0.00052 | -1.25
Spectrum (double click to dock / undock)
N.AASIPSK[+324]CNVNVPY.T z=2,scan#=sample=1 period=1 cycle=546 experiment=17,scan time=10.9249
AASIPSKCNVNVPY
Intensity
m/z
m/z errors, tol=75.0 ppm
Protein Coverage
Sample name
Protein name
10:55 AM
20/08/2020

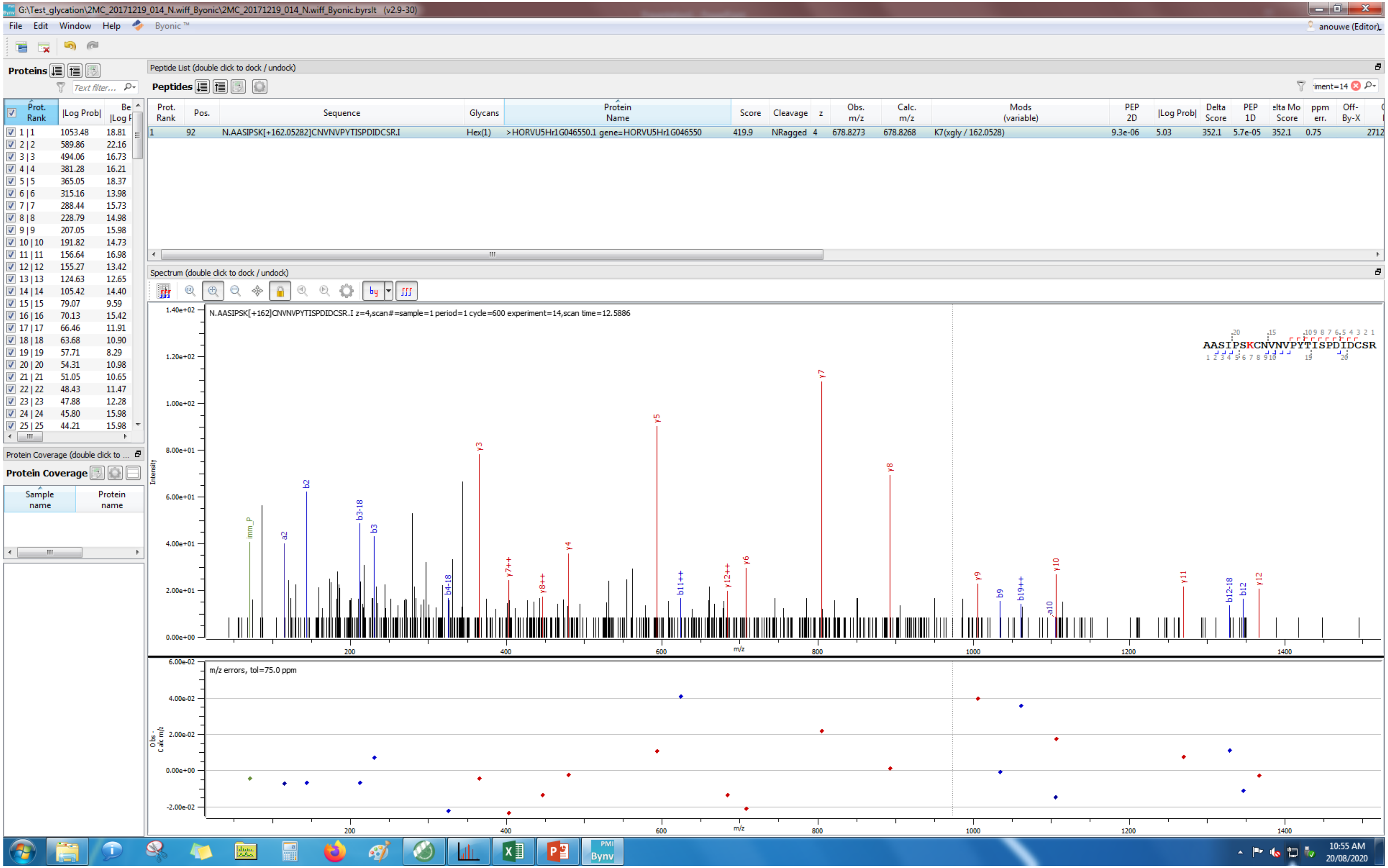

G:\Test_glycation\2MC_20171219_014_N.wiff_Byonic\2MC_20171219_014_N.wiff_Byonic.byrslt (v2.9-30)
File Edit Window Help Byonic ™
anouwe (Editor)
Proteins
Peptide List (double click to dock / undock)
Peptides
Prot. Rank | Pos. | Sequence | Glycans | Protein Name | Score | Cleavage | z | Obs. m/z | Calc. m/z | Mods (variable) | PEP 2D | |Log Prob| | Delta Score | PEP 1D | Delta Mod Score | ppm err. | Off-By-X
1 | 92 | N.AASIPSK[+162.05282]CNVNVPYTISPDIDCSR.I | Hex(1) | >HORVU5Hr1G046550.1 gene=HORVU5Hr1G046550 | 419.9 | NRagged | 4 | 678.8273 | 678.8268 | K7(xgly / 162.0528) | 9.3e-06 | 5.03 | 352.1 | 5.7e-05 | 352.1 | 0.75 | 2712
Spectrum (double click to dock / undock)
N.AASIPSK[+162]CNVNVPYTISPDIDCSR.I z=4,scan#=sample=1 period=1 cycle=600 experiment=14,scan time=12.5886
AASIPSKCNVNVPYTISPDIDCSR
Intensity
m/z
m/z errors, tol=75.0 ppm
Obs - Calc m/z
Protein Coverage (double click to ...
Protein Coverage
Sample name
Protein name
10:55 AM
20/08/2020

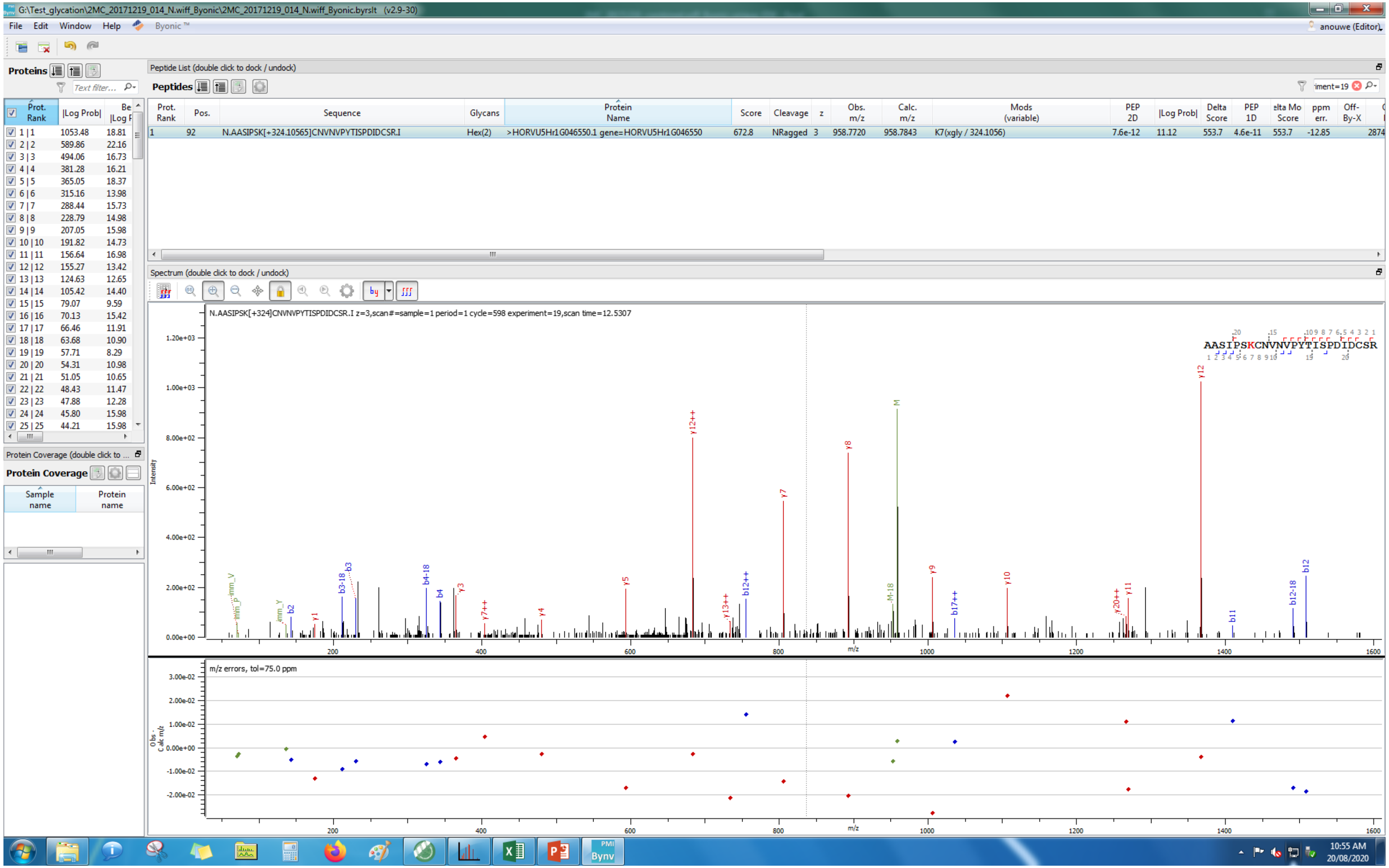
G:\Test_glycation\2MC_20171219_014_N.wiff_Byonic\2MC_20171219_014_N.wiff_Byonic.byrslt (v2.9-30)
File Edit Window Help Byonic
Proteins
Peptide List (double click to dock / undock)
Peptides
Prot. Rank | Pos. | Sequence | Glycans | Protein Name | Score | Cleavage | z | Obs. m/z | Calc. m/z | Mods (variable) | PEP 2D | |Log Prob| | Delta Score | PEP 1D | Delta Mod Score | ppm err. | Off-By-X
1 | 92 | N.AASIPSK[+324.10565]CNVNVPYTISPDIDCSR.I | Hex(2) | >HORVU5Hr1G046550.1 gene=HORVU5Hr1G046550 | 672.8 | NRagged | 3 | 958.7720 | 958.7843 | K7(xgly / 324.1056) | 7.6e-12 | 11.12 | 553.7 | 4.6e-11 | 553.7 | -12.85
Spectrum (double click to dock / undock)
N.AASIPSK[+324]CNVNVPYTISPDIDCSR.I z=3,scan#=sample=1 period=1 cycle=598 experiment=19,scan time=12.5307
AASIPSKCNVNVPYTISPDIDCSR
Intensity
m/z
m/z errors, tol=75.0 ppm
Protein Coverage (double click to ...
Protein Coverage
Sample name
Protein name
10:55 AM
20/08/2020

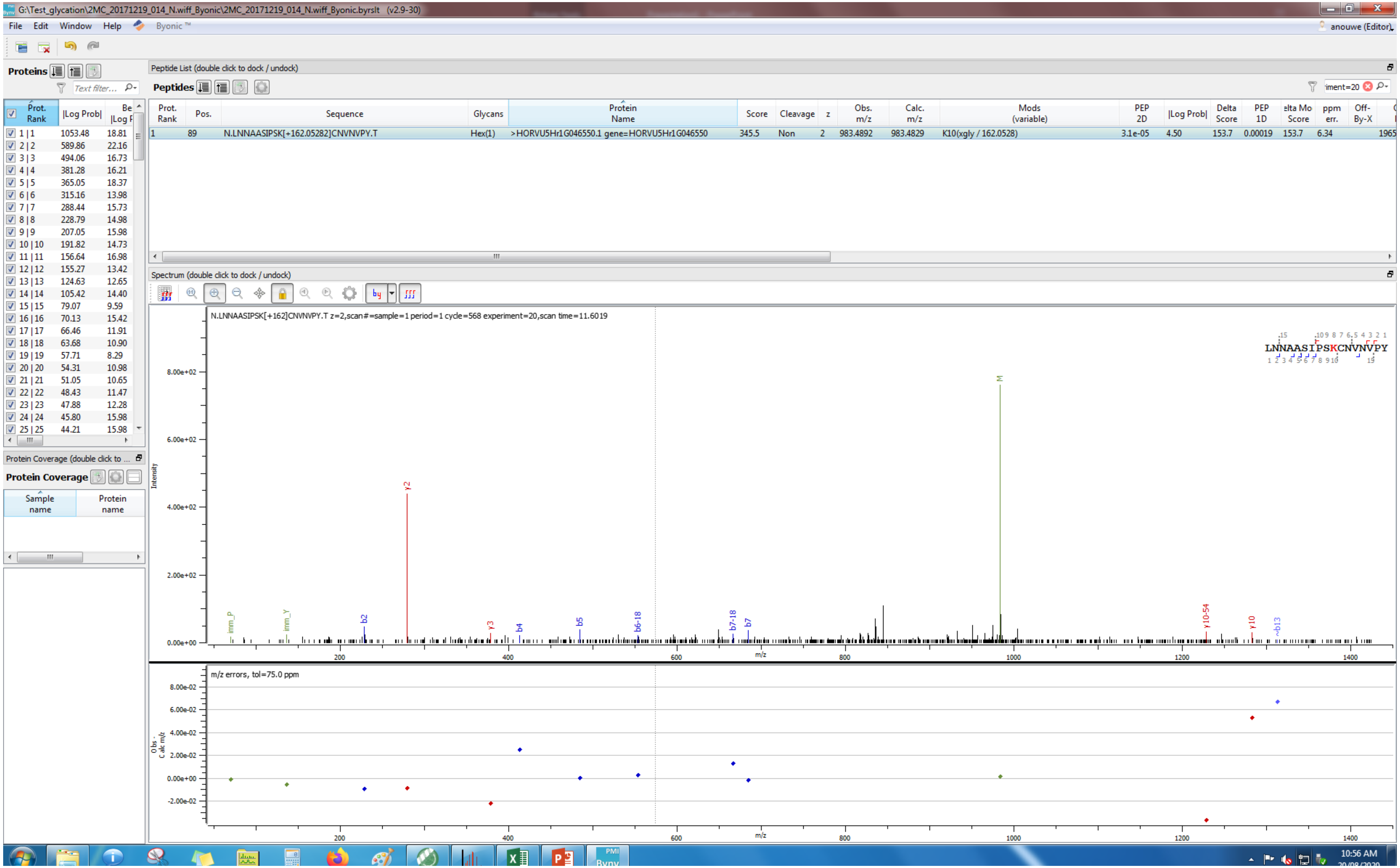
G:\Test_glycation\2MC_20171219_014_N.wiff_Byonic\2MC_20171219_014_N.wiff_Byonic.byrslt (v2.9-30)
File Edit Window Help Byonic™
anouwe (Editor)
Proteins
Peptide List (double click to dock / undock)
Peptides
Prot. Rank | Pos. | Sequence | Glycans | Protein Name | Score | Cleavage | z | Obs. m/z | Calc. m/z | Mods (variable) | PEP 2D | |Log Prob| | Delta Score | PEP 1D | elta Mo Score | ppm err. | Off-By-X
1 | 89 | N.LNNAASIPSK[+162.05282]CNVNVPY.T | Hex(1) | >HORVU5Hr1G046550.1 gene=HORVU5Hr1G046550 | 345.5 | Non | 2 | 983.4892 | 983.4829 | K10(xgly / 162.0528) | 3.1e-05 | 4.50 | 153.7 | 0.00019 | 153.7 | 6.34
Prot. Rank | |Log Prob| | |Log P
1 | 1 1053.48 18.81
2 | 2 589.86 22.16
3 | 3 494.06 16.73
4 | 4 381.28 16.21
5 | 5 365.05 18.37
6 | 6 315.16 13.98
7 | 7 288.44 15.73
8 | 8 228.79 14.98
9 | 9 207.05 15.98
10 | 10 191.82 14.73
11 | 11 156.64 16.98
12 | 12 155.27 13.42
13 | 13 124.63 12.65
14 | 14 105.42 14.40
15 | 15 79.07 9.59
16 | 16 70.13 15.42
17 | 17 66.46 11.91
18 | 18 63.68 10.90
19 | 19 57.71 8.29
20 | 20 54.31 10.98
21 | 21 51.05 10.65
22 | 22 48.43 11.47
23 | 23 47.88 12.28
24 | 24 45.80 15.98
25 | 25 44.21 15.98
Protein Coverage
Sample name
Protein name
Spectrum (double click to dock / undock)
N.LNNAASIPSK[+162]CNVNVPY.T z=2,scan#=sample=1 period=1 cycle=568 experiment=20,scan time=11.6019
LNNAASIPSKCNVNVPY
Intensity
8.00e+02
6.00e+02
4.00e+02
2.00e+02
0.00e+00
imm_P
imm_Y
b2
y2
y3
b4
b5
b6-18
b7-18
b7
M
y10-54
y10
~b13
m/z
200
400
600
800
1000
1200
1400
m/z errors, tol=75.0 ppm
Obs - Calc m/z
8.00e-02
6.00e-02
4.00e-02
2.00e-02
0.00e+00
-2.00e-02
10:56 AM

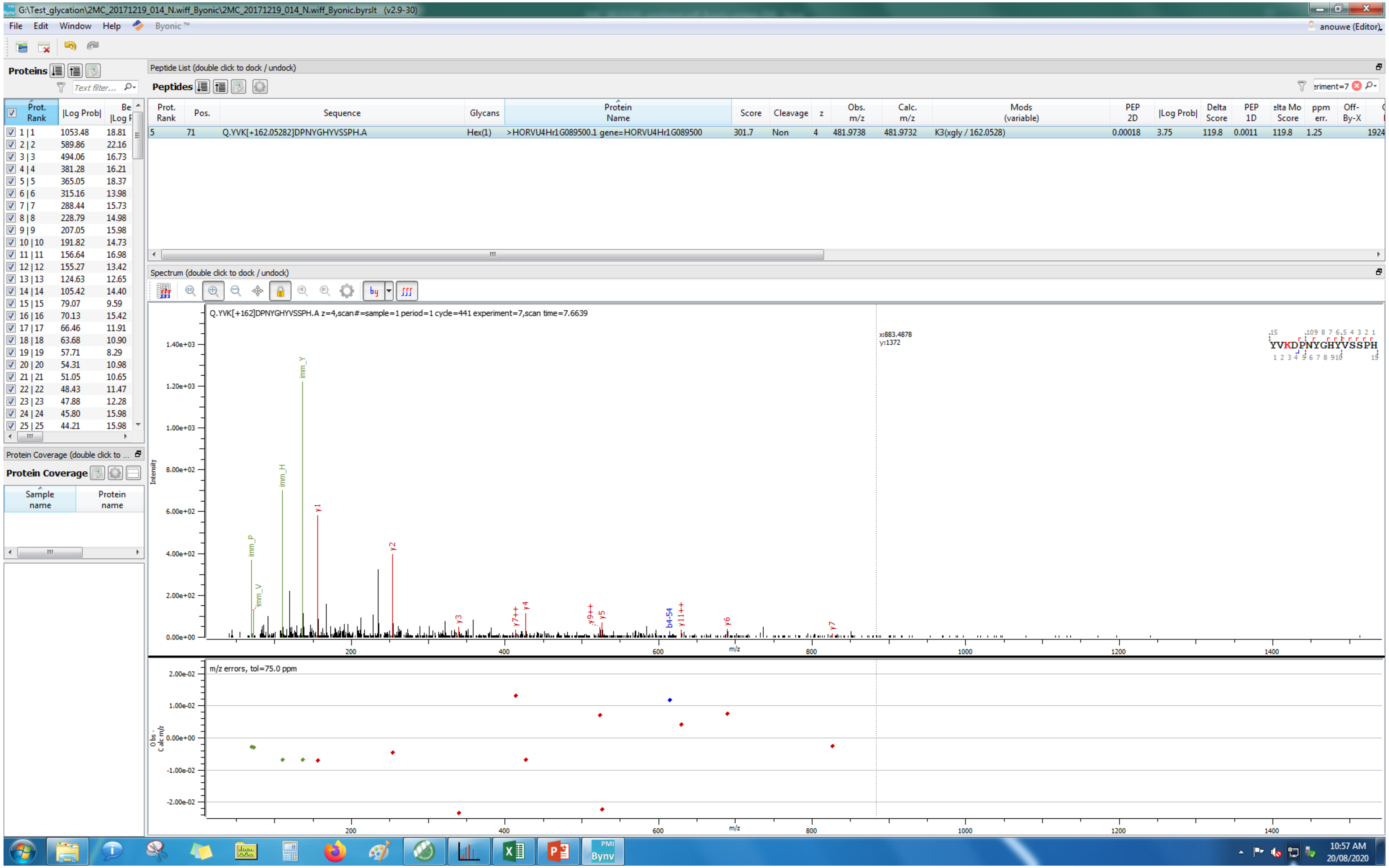
G:\Test_glycation\2MC_20171219_014_N.wiff_Byonic\2MC_20171219_014_N.wiff_Byonic.byrslt (v2.9-30)
File Edit Window Help Byonic™
anouwe (Editor)
Proteins
Peptide List (double click to dock / undock)
Peptides
Prot. Rank | Pos. | Sequence | Glycans | Protein Name | Score | Cleavage | z | Obs. m/z | Calc. m/z | Mods (variable) | PEP 2D | |Log Prob| | Delta Score | PEP 1D | Delta Mod Score | ppm err. | Off-By-X
5 | 71 | Q.YVK[+162.05282]DPNYGHYVSSPH.A | Hex(1) | >HORVU4Hr1G089500.1 gene=HORVU4Hr1G089500 | 301.7 | Non | 4 | 481.9738 | 481.9732 | K3(xgly / 162.0528) | 0.00018 | 3.75 | 119.8 | 0.0011 | 119.8 | 1.25 | 1924
Prot. Rank | |Log Prob| | Best |Log Prob|
1 | 1 1053.48 18.81
2 | 2 589.86 22.16
3 | 3 494.06 16.73
4 | 4 381.28 16.21
5 | 5 365.05 18.37
6 | 6 315.16 13.98
7 | 7 288.44 15.73
8 | 8 228.79 14.98
9 | 9 207.05 15.98
10 | 10 191.82 14.73
11 | 11 156.64 16.98
12 | 12 155.27 13.42
13 | 13 124.63 12.65
14 | 14 105.42 14.40
15 | 15 79.07 9.59
16 | 16 70.13 15.42
17 | 17 66.46 11.91
18 | 18 63.68 10.90
19 | 19 57.71 8.29
20 | 20 54.31 10.98
21 | 21 51.05 10.65
22 | 22 48.43 11.47
23 | 23 47.88 12.28
24 | 24 45.80 15.98
25 | 25 44.21 15.98
Protein Coverage (double click to ...
Protein Coverage
Sample name | Protein name
Spectrum (double click to dock / undock)
Q.YVK[+162]DPNYGHYVSSPH.A z=4,scan#=sample=1 period=1 cycle=441 experiment=7,scan time=7.6639
x:883.4878
y:1372
YVKDPNYGHYVSSPH
Intensity
imm_Y
imm_H
imm_P
imm_V
y1
y2
y3
y7++
y4
y9++
y5
b4-54
y11++
y6
y7
m/z
m/z errors, tol=75.0 ppm
Obs - Calc m/z
10:57 AM
20/08/2020

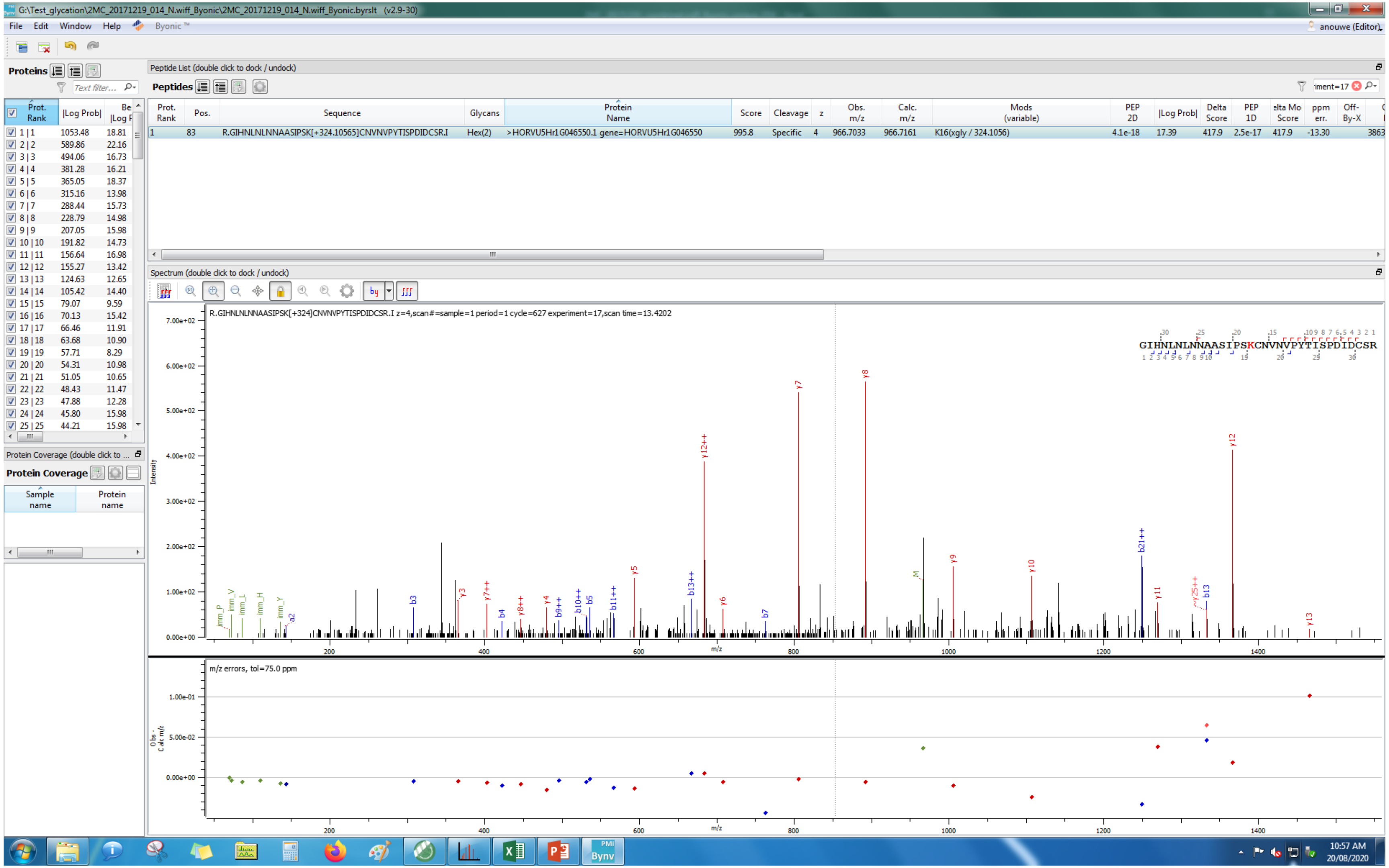
G:\Test_glycation\2MC_20171219_014_N.wiff_Byonic\2MC_20171219_014_N.wiff_Byonic.byrslt (v2.9-30)
File Edit Window Help Byonic
Proteins
Peptide List (double click to dock / undock)
Peptides
Prot. Rank Pos. Sequence Glycans Protein Name Score Cleavage z Obs. m/z Calc. m/z Mods (variable) PEP 2D |Log Prob| Delta Score PEP 1D Delta Mod Score ppm err. Off-By-X
1 83 R.GIHNLNLNNAASIPSK[+324.10565]CNVNVPYTISPDIDCSR.I Hex(2) >HORVU5Hr1G046550.1 gene=HORVU5Hr1G046550 995.8 Specific 4 966.7033 966.7161 K16(xgly / 324.1056) 4.1e-18 17.39 417.9 2.5e-17 417.9 -13.30
Spectrum (double click to dock / undock)
R.GIHNLNLNNAASIPSK[+324]CNVNVPYTISPDIDCSR.I z=4,scan#=sample=1 period=1 cycle=627 experiment=17,scan time=13.4202
GIHNLNLNNAASIPSKCNVNVPYTISPDIDCSR
Intensity
m/z
m/z errors, tol=75.0 ppm
Protein Coverage
Sample name
Protein name
10:57 AM
20/08/2020

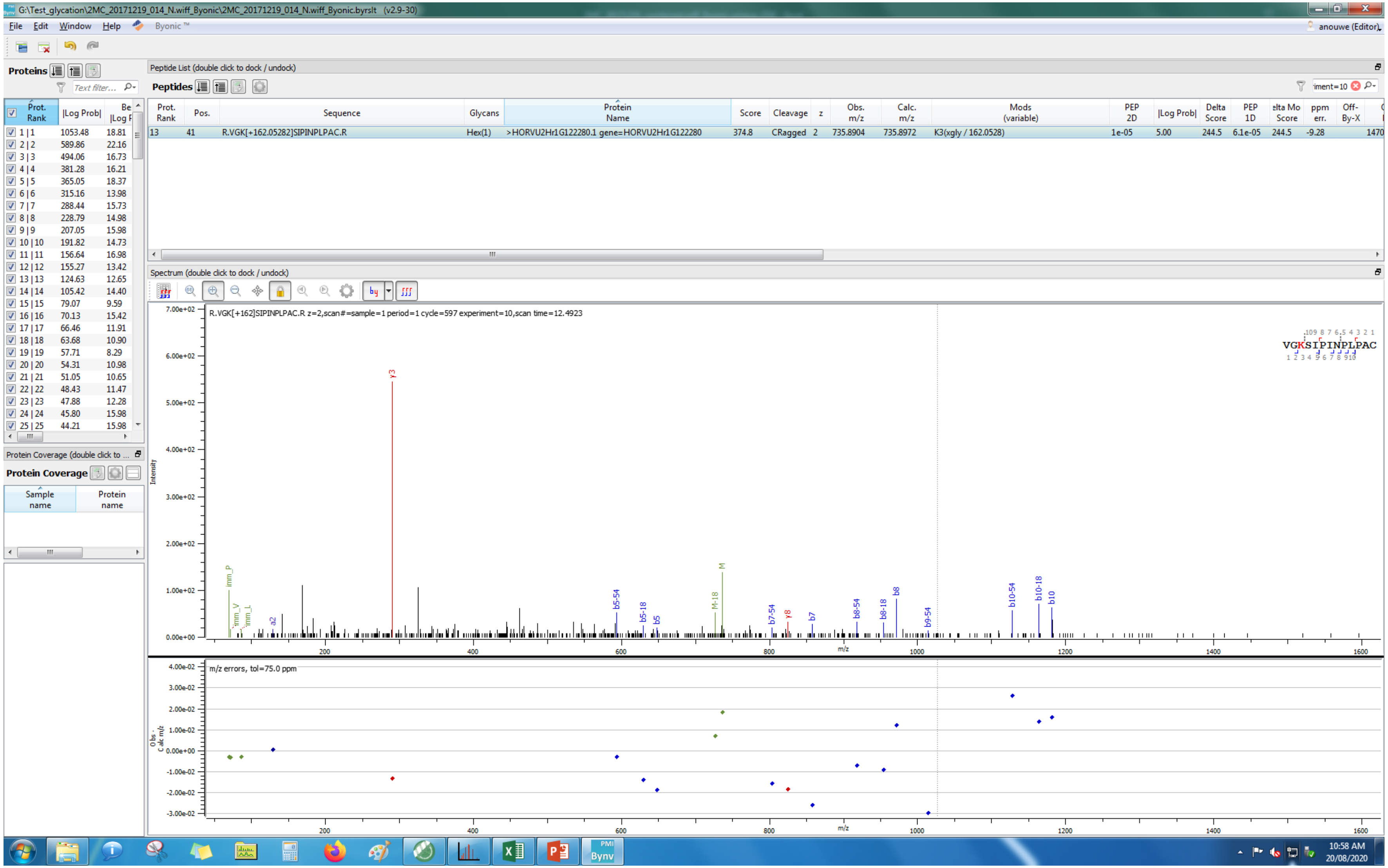
G:\Test_glycation\2MC_20171219_014_N.wiff_Byonic\2MC_20171219_014_N.wiff_Byonic.byrslt (v2.9-30)
File Edit Window Help Byonic
anouwe (Editor)
Proteins
Peptide List (double click to dock / undock)
Peptides
Prot. Rank
|Log Prob|
Pos.
Sequence
Glycans
Protein Name
Score
Cleavage
z
Obs. m/z
Calc. m/z
Mods (variable)
PEP 2D
Delta Score
PEP 1D
ppm err.
Off-By-X
13 41 R.VGK[+162.05282]SIPINPLPAC.R Hex(1) >HORVU2Hr1G122280.1 gene=HORVU2Hr1G122280 374.8 CRagged 2 735.8904 735.8972 K3(xgly / 162.0528) 1e-05 5.00 244.5 6.1e-05 244.5 -9.28 1470
Spectrum (double click to dock / undock)
R.VGK[+162]SIPINPLPAC.R z=2,scan#=sample=1 period=1 cycle=597 experiment=10,scan time=12.4923
VGKSIPINPLPAC
Intensity
m/z
m/z errors, tol=75.0 ppm
Protein Coverage
Sample name
Protein name
10:58 AM
20/08/2020

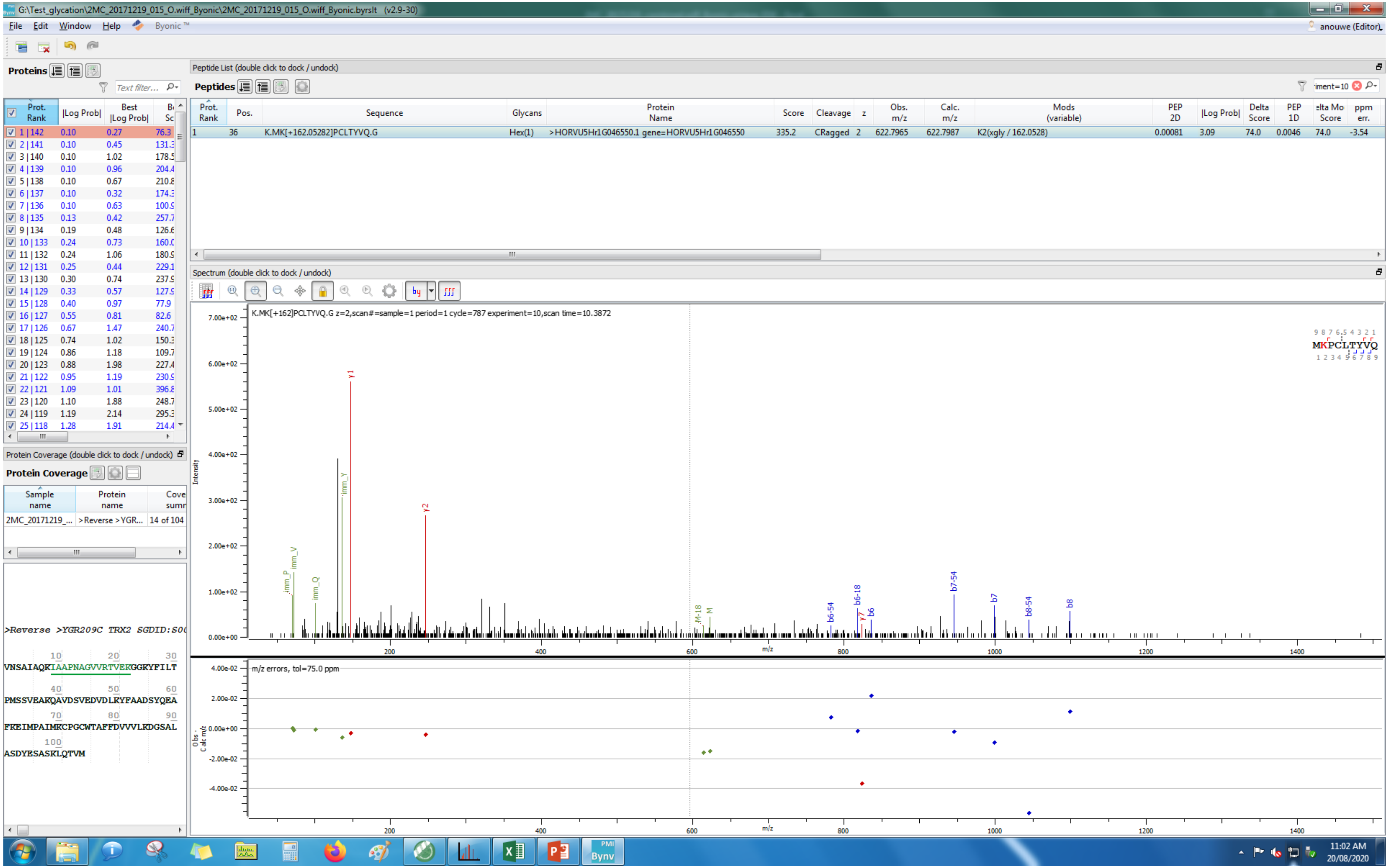
G:\Test_glycation\2MC_20171219_015_O.wiff_Byonic\2MC_20171219_015_O.wiff_Byonic.byrslt (v2.9-30)
File Edit Window Help Byonic™
anouwe (Editor)
Proteins
Peptide List (double click to dock / undock)
Peptides
Prot. Rank | Pos. | Sequence | Glycans | Protein Name | Score | Cleavage | z | Obs. m/z | Calc. m/z | Mods (variable) | PEP 2D | |Log Prob| | Delta Score | PEP 1D | Delta Mod Score | ppm err.
1 | 36 | K.MK[+162.05282]PCLTYVQ.G | Hex(1) | >HORVU5Hr1G046550.1 gene=HORVU5Hr1G046550 | 335.2 | CRagged | 2 | 622.7965 | 622.7987 | K2(xgly / 162.0528) | 0.00081 | 3.09 | 74.0 | 0.0046 | 74.0 | -3.54
Spectrum (double click to dock / undock)
K.MK[+162]PCLTYVQ.G z=2,scan#=sample=1 period=1 cycle=787 experiment=10,scan time=10.3872
MKPCLTYVQ
Intensity
m/z
m/z errors, tol=75.0 ppm
Protein Coverage (double click to dock / undock)
Protein Coverage
Sample name | Protein name | Coverage summary
2MC_20171219_... | >Reverse >YGR... | 14 of 104
>Reverse >YGR209C TRX2 SGDID:S0
VNSAIAQKIAAPNAGVVRTVEKGGKYFILT
PMSSVEAKQAVDSVEDVLRYFAADSYQEA
FKEIMPAIMKCPGCWTAFFDVVVLKDGSAL
ASDYESASKLQTVM
11:02 AM
20/08/2020

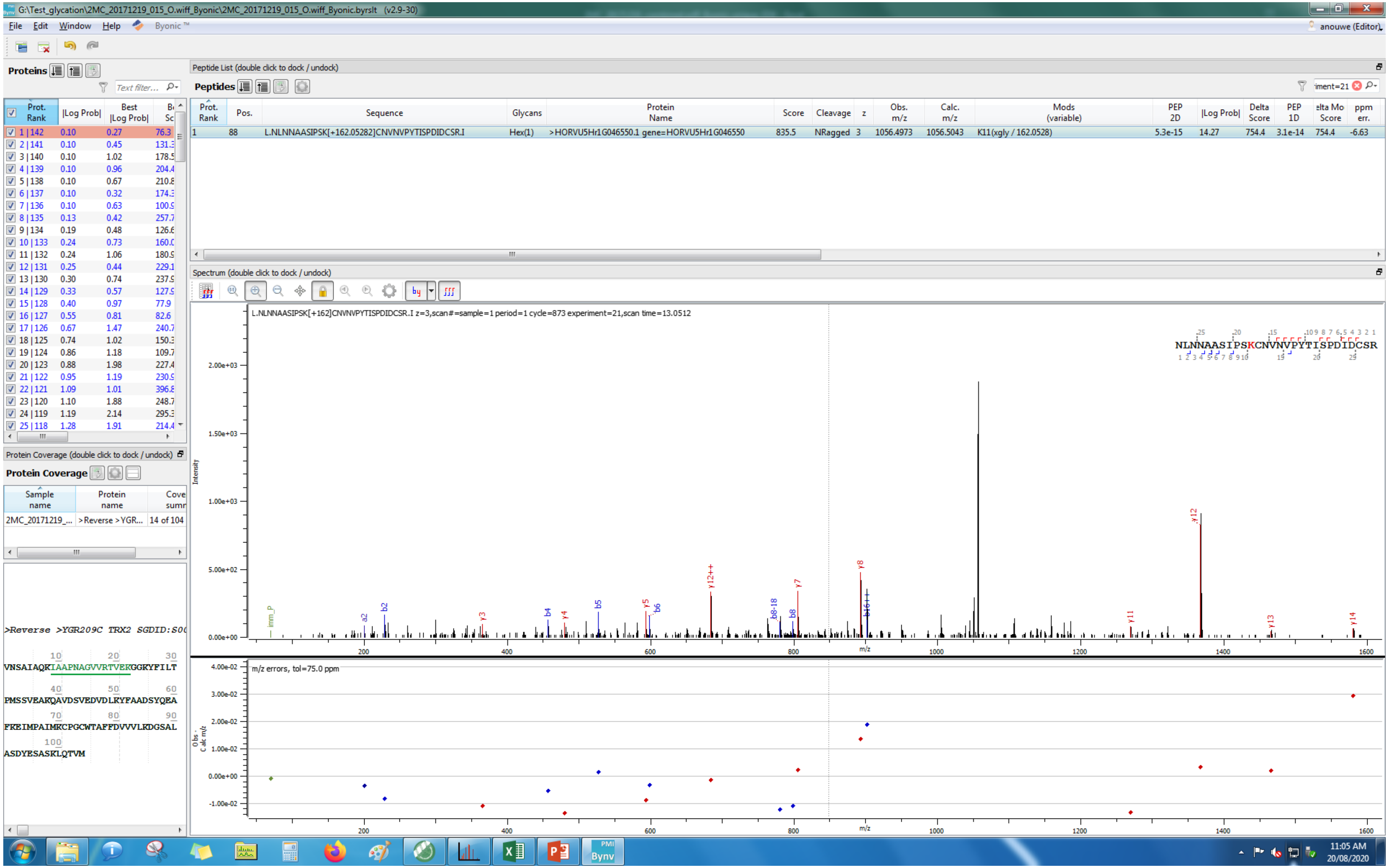

G:\Test_glycation\2MC_20171219_015_O.wiff_Byonic\2MC_20171219_015_O.wiff_Byonic.byrslt (v2.9-30)
File Edit Window Help Byonic
Proteins
Peptide List (double click to dock / undock)
Peptides
Prot. Rank | Pos. | Sequence | Glycans | Protein Name | Score | Cleavage | z | Obs. m/z | Calc. m/z | Mods (variable) | PEP 2D | |Log Prob| | Delta Score | PEP 1D | ppm err.
1 | 88 | L.NLNNAASIPSK[+162.05282]CNVNVPYTISPDIDCSR.I | Hex(1) | >HORVU5Hr1G046550.1 gene=HORVU5Hr1G046550 | 835.5 | NRagged | 3 | 1056.4973 | 1056.5043 | K11(xgly / 162.0528) | 5.3e-15 | 14.27 | 754.4 | 3.1e-14 | 754.4 | -6.63
Spectrum (double click to dock / undock)
L.NLNNAASIPSK[+162]CNVNVPYTISPDIDCSR.I z=3,scan#=sample=1 period=1 cycle=873 experiment=21,scan time=13.0512
NLNNAASIPSKCNVNVPYTISPDIDCSR
m/z errors, tol=75.0 ppm
Protein Coverage (double click to dock / undock)
Protein Coverage
Sample name | Protein name
2MC_20171219_... | >Reverse >YGR...
>Reverse >YGR209C TRX2 SGDID:S00
VNSAIAQKIAAPNAGVVRTVEKGGKYFILT
PMSSVEAKQAVDSVEDVDLKYFAADSYQEA
FKEIMPAIMKCPGCWTAFFDVVVLKDGSAL
ASDYESASKLQTVM

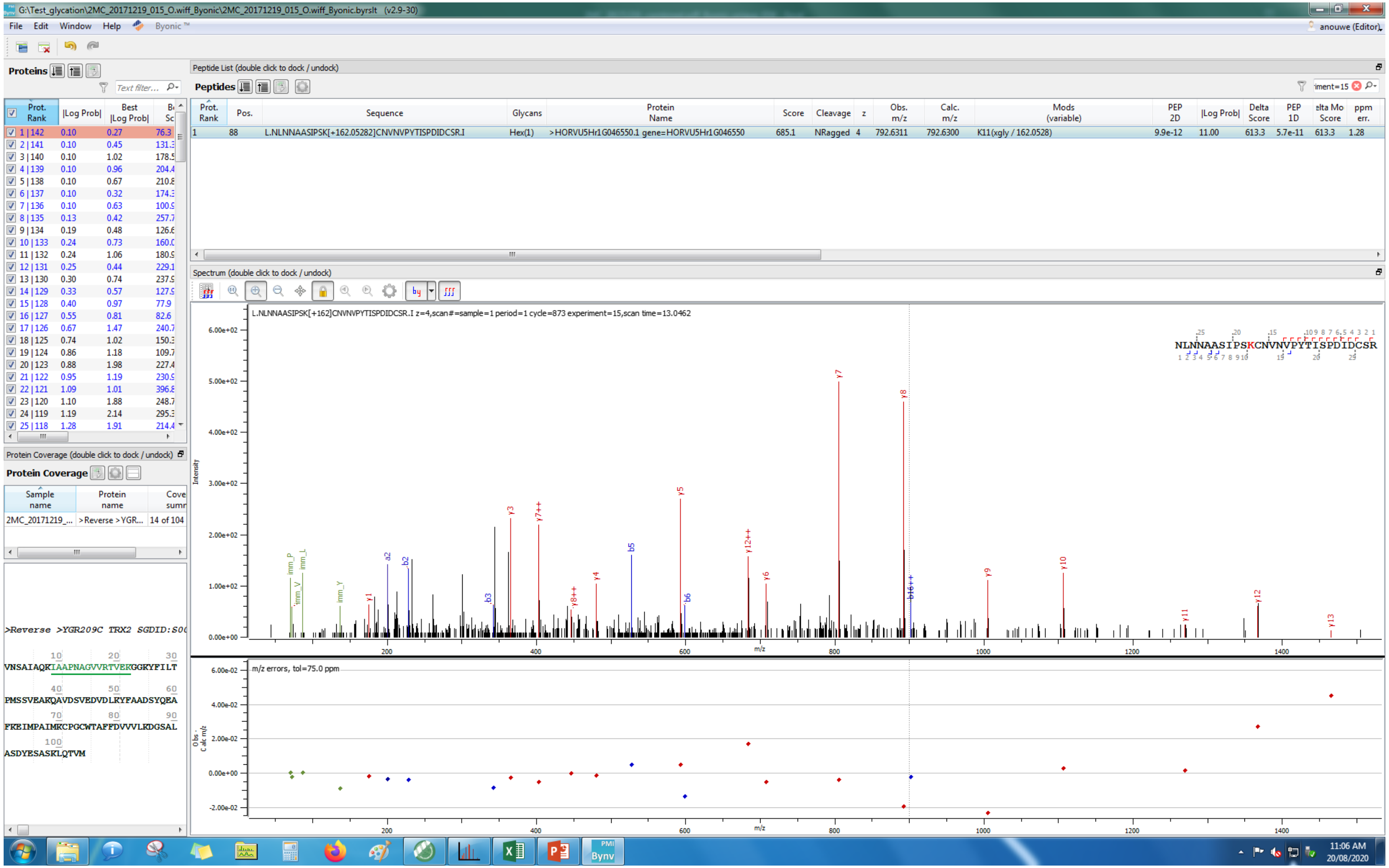
G:\Test_glycation\2MC_20171219_015_O.wiff_Byonic\2MC_20171219_015_O.wiff_Byonic.byrslt (v2.9-30)
File Edit Window Help Byonic™
anouwe (Editor)
Proteins
Peptide List (double click to dock / undock)
Peptides
Prot. Rank | Pos. | Sequence | Glycans | Protein Name | Score | Cleavage | z | Obs. m/z | Calc. m/z | Mods (variable) | PEP 2D | |Log Prob| | Delta Score | PEP 1D | Delta Mod Score | ppm err.
1 | 88 | L.NLNNAASIPSK[+162.05282]CNVNVPYTISPDIDCSR.I | Hex(1) | >HORVU5Hr1G046550.1 gene=HORVU5Hr1G046550 | 685.1 | NRagged | 4 | 792.6311 | 792.6300 | K11(xgly / 162.0528) | 9.9e-12 | 11.00 | 613.3 | 5.7e-11 | 613.3 | 1.28
Spectrum (double click to dock / undock)
L.NLNNAASIPSK[+162]CNVNVPYTISPDIDCSR.I z=4,scan#=sample=1 period=1 cycle=873 experiment=15,scan time=13.0462
NLNNAASIPSKCNVNVPYTISPDIDCSR
Intensity
m/z
m/z errors, tol=75.0 ppm
Protein Coverage (double click to dock / undock)
Protein Coverage
Sample name | Protein name | Coverage summary
2MC_20171219_... | >Reverse >YGR... | 14 of 104
>Reverse >YGR209C TRX2 SGDID:S00
VNSAIAQKIAAPNAGVVRTVERGGKYFILT
PMSSVEAKQAVDSVEDVDLKYFAADSYQEA
FKEIMPAIMKCPGCWTAPFDVVVLKDGSAL
ASDYESASKLQTVM
11:06 AM
20/08/2020

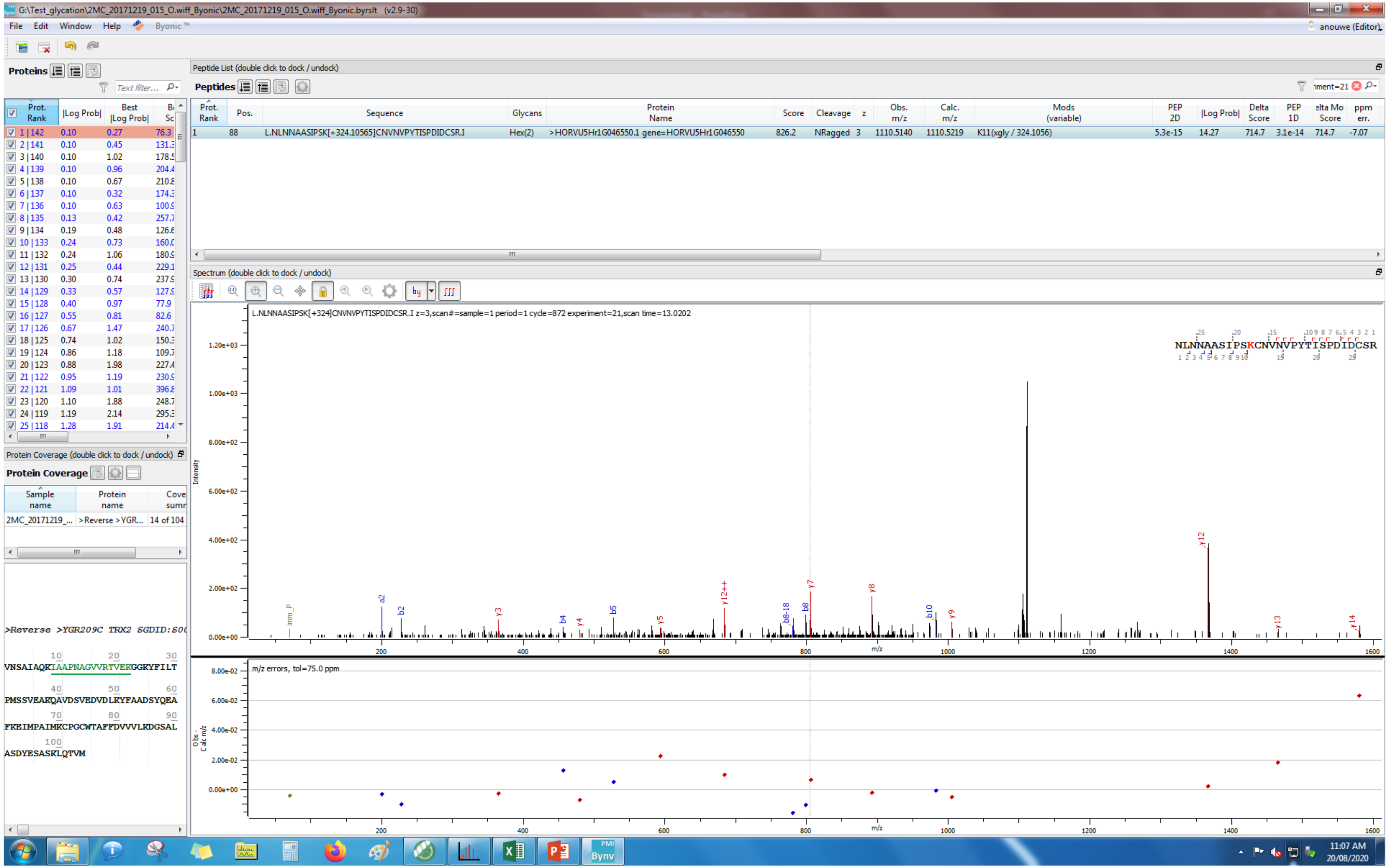
G:\Test_glycation\2MC_20171219_015_O.wiff_Byonic\2MC_20171219_015_O.wiff_Byonic.byrslt (v2.9-30)
File Edit Window Help Byonic ™
anouwe (Editor)
Proteins
Peptide List (double click to dock / undock)
Peptides
Prot. Rank | Pos. | Sequence | Glycans | Protein Name | Score | Cleavage | z | Obs. m/z | Calc. m/z | Mods (variable) | PEP 2D | |Log Prob| | Delta Score | PEP 1D | elta Mo Score | ppm err.
1 | 88 | L.NLNNAASIPSK[+324.10565]CNVNVPYTISPDIDCSR.I | Hex(2) | >HORVU5Hr1G046550.1 gene=HORVU5Hr1G046550 | 826.2 | NRagged | 3 | 1110.5140 | 1110.5219 | K11(xgly / 324.1056) | 5.3e-15 | 14.27 | 714.7 | 3.1e-14 | 714.7 | -7.07
Spectrum (double click to dock / undock)
L.NLNNAASIPSK[+324]CNVNVPYTISPDIDCSR.I z=3,scan#=sample=1 period=1 cycle=872 experiment=21,scan time=13.0202
NLNNAASIPSKCNVNVPYTISPDIDCSR
Intensity
m/z
m/z errors, tol=75.0 ppm
Protein Coverage (double click to dock / undock)
Protein Coverage
Sample name | Protein name | Cove summ
2MC_20171219_... | >Reverse >YGR... | 14 of 104
>Reverse >YGR209C TRX2 SGDID:S0
VNSAIAQKIAAPNAGVVRTVEKGGKYFILT
PMSSVEARQAVDSVEDVDLKYFAADSYQEA
FKEIMPAIMKCPGCWTAFFDVVVLKDGSAL
ASDYESASKLQTVM
11:07 AM
20/08/2020

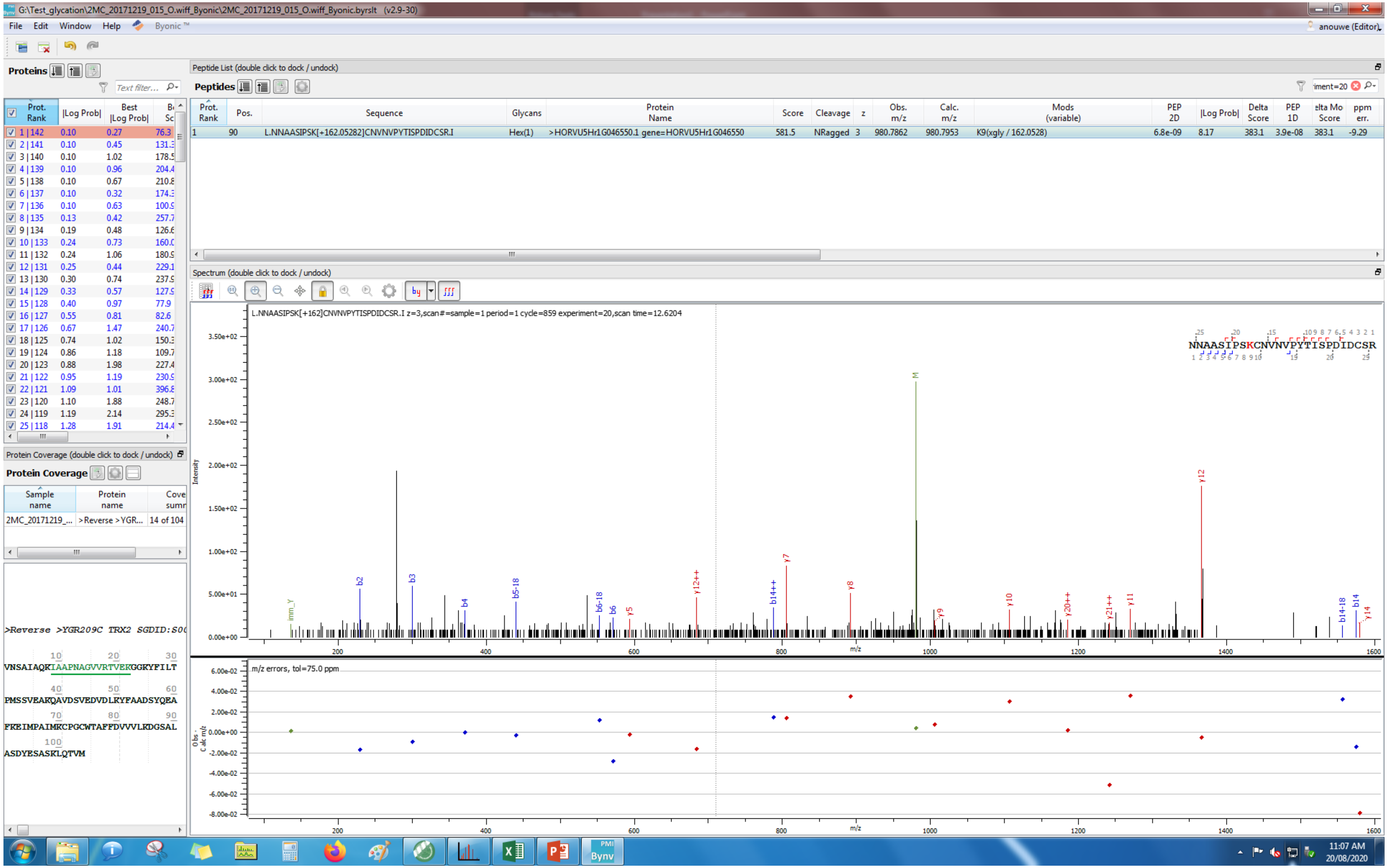
G:\Test_glycation\2MC_20171219_015_O.wiff_Byonic\2MC_20171219_015_O.wiff_Byonic.byrslt (v2.9-30)
File Edit Window Help Byonic™
anouwe (Editor)
Proteins
Peptide List (double click to dock / undock)
Peptides
Prot. Rank | Pos. | Sequence | Glycans | Protein Name | Score | Cleavage | z | Obs. m/z | Calc. m/z | Mods (variable) | PEP 2D | |Log Prob| | Delta Score | PEP 1D | Delta Mod Score | ppm err.
1 | 90 | L.NNAASIPSK[+162.05282]CNVNVPYTISPDIDCSR.I | Hex(1) | >HORVU5Hr1G046550.1 gene=HORVU5Hr1G046550 | 581.5 | NRagged | 3 | 980.7862 | 980.7953 | K9(xgly / 162.0528) | 6.8e-09 | 8.17 | 383.1 | 3.9e-08 | 383.1 | -9.29
Spectrum (double click to dock / undock)
L.NNAASIPSK[+162]CNVNVPYTISPDIDCSR.I z=3,scan#=sample=1 period=1 cycle=859 experiment=20,scan time=12.6204
NNAASIPSKCNVNVPYTISPDIDCSR
Intensity
m/z
m/z errors, tol=75.0 ppm
Obs - Calc m/z
Protein Coverage (double click to dock / undock)
Protein Coverage
Sample name | Protein name | Coverage summary
2MC_20171219_... | >Reverse >YGR... | 14 of 104
>Reverse >YGR209C TRX2 SGDID:S00
VNSAIAQKIAAPNAGVVRTVEKGGKYPILT
PMSSVEAKQAVDSVEDVDLKYFAADSYQEA
FKEIMPAIMKCPGCWTAFFDVVVLKDGSAL
ASDYESASKLQTVM
11:07 AM
20/08/2020

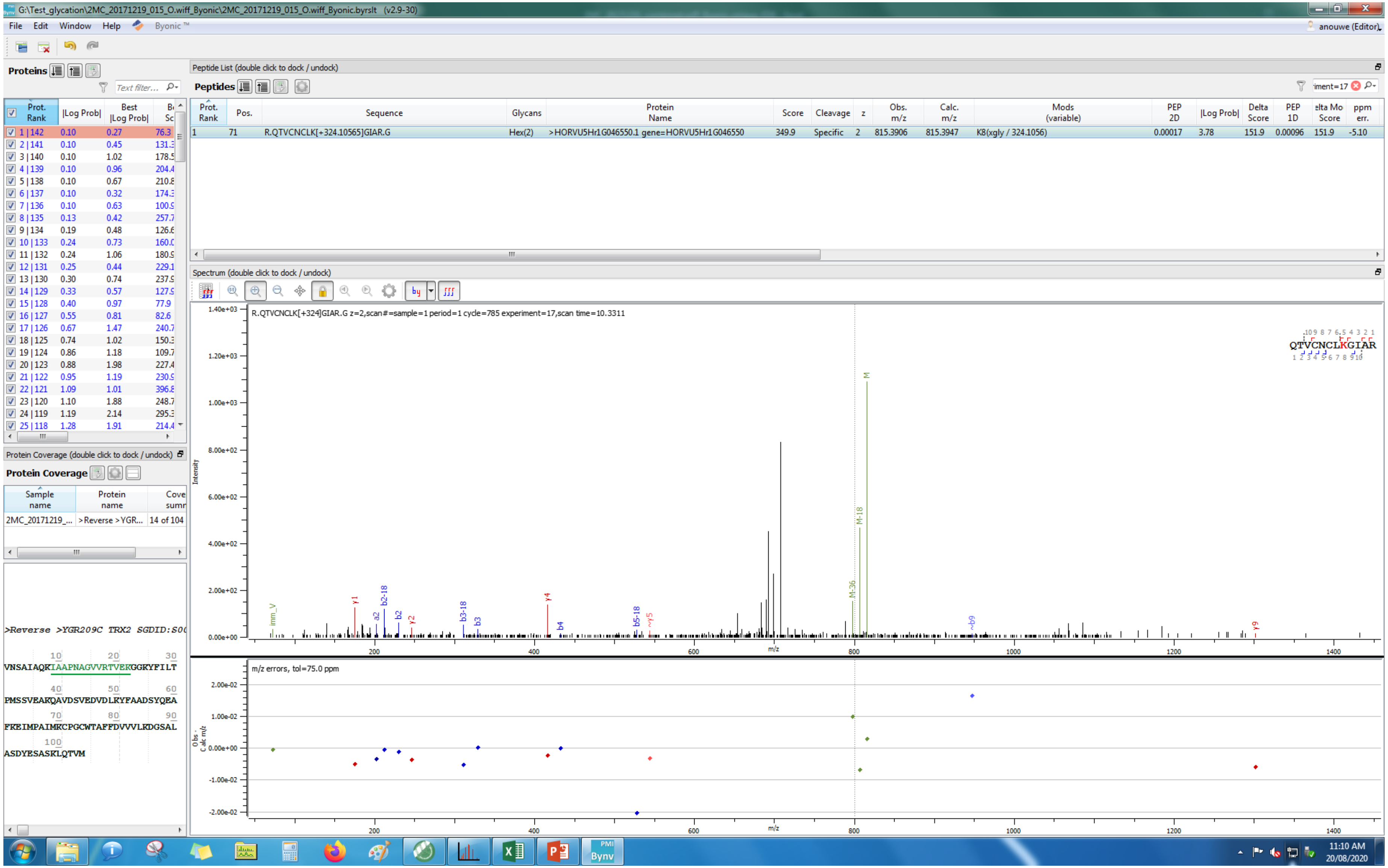
G:\Test_glycation\2MC_20171219_015_O.wiff_Byonic\2MC_20171219_015_O.wiff_Byonic.byrslt (v2.9-30)
File Edit Window Help Byonic™
Proteins
Peptide List (double click to dock / undock)
Peptides
Prot. Rank | Pos. | Sequence | Glycans | Protein Name | Score | Cleavage | z | Obs. m/z | Calc. m/z | Mods (variable) | PEP 2D | |Log Prob| | Delta Score | PEP 1D | Delta Mod Score | ppm err.
1 | 71 | R.QTVCNCLK[+324.10565]GIAR.G | Hex(2) | >HORVU5Hr1G046550.1 gene=HORVU5Hr1G046550 | 349.9 | Specific | 2 | 815.3906 | 815.3947 | K8(xgly / 324.1056) | 0.00017 | 3.78 | 151.9 | 0.00096 | 151.9 | -5.10
Spectrum (double click to dock / undock)
R.QTVCNCLK[+324]GIAR.G z=2,scan#=sample=1 period=1 cycle=785 experiment=17,scan time=10.3311
QTVCNCLKGIAR
m/z errors, tol=75.0 ppm
Protein Coverage (double click to dock / undock)
Protein Coverage
Sample name | Protein name | Coverage summary
2MC_20171219_... | >Reverse >YGR... | 14 of 104
>Reverse >YGR209C TRX2 SGDID:S0
VNSAIAQKIAAPNAGVVRTVEKGGKYFILT
PMSSVEAKQAVDSVEDVDLKYFAADSYQEA
FKEIMPAIMKCPGCWTAFFDVVVLKDGSAL
ASDYESASKLQTVM
anouwe (Editor)
11:10 AM
20/08/2020

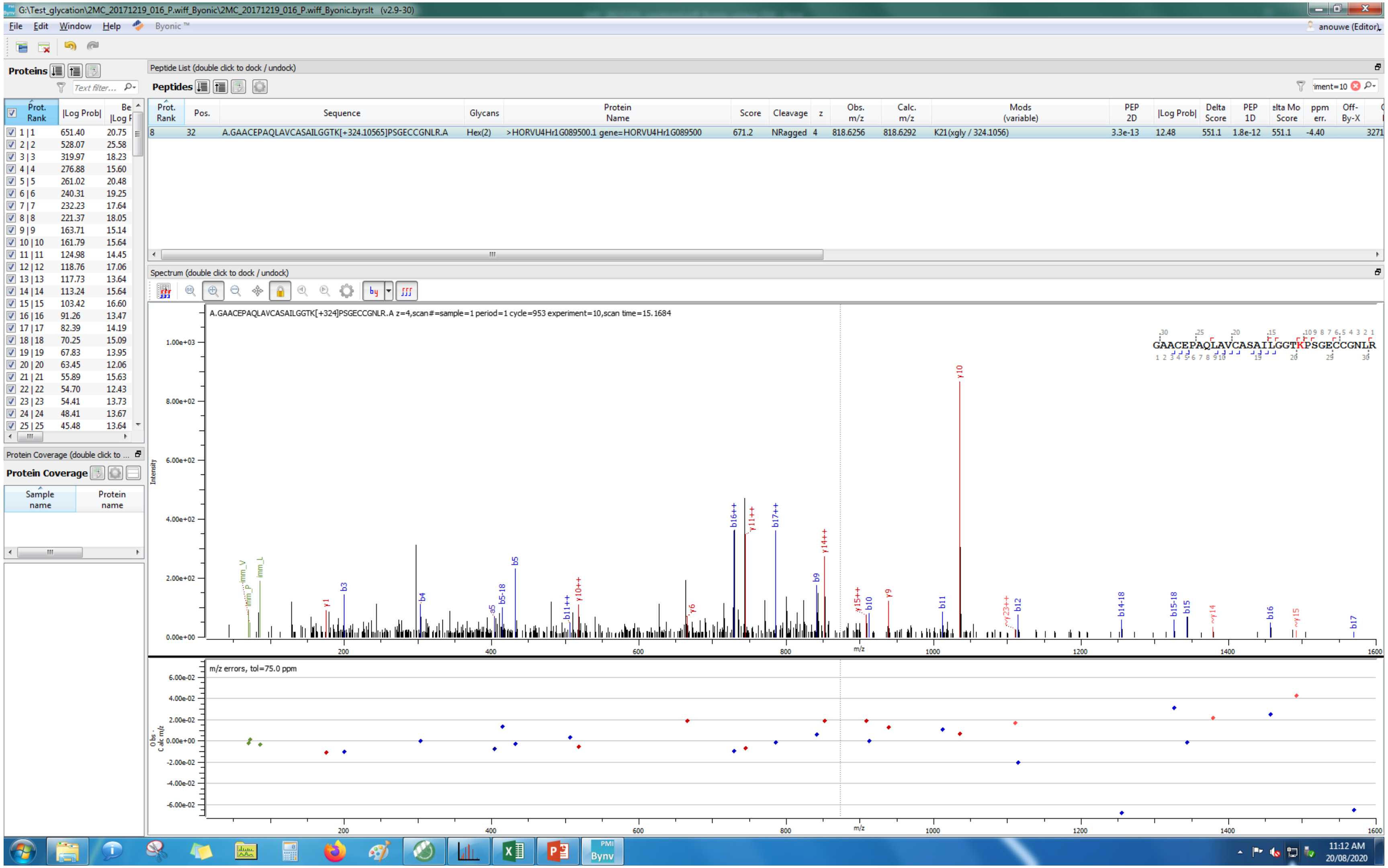
G:\Test_glycation\2MC_20171219_016_P.wiff_Byonic\2MC_20171219_016_P.wiff_Byonic.byrslt (v2.9-30)
File Edit Window Help Byonic ™
anouwe (Editor)
Proteins
Peptide List (double click to dock / undock)
Peptides
Prot. Rank | Pos. | Sequence | Glycans | Protein Name | Score | Cleavage | z | Obs. m/z | Calc. m/z | Mods (variable) | PEP 2D | |Log Prob| | Delta Score | PEP 1D | Delta Mod Score | ppm err.
8 | 32 | A.GAACEPAQLAVCASAILGGTK[+324.10565]PSGECCGNLR.A | Hex(2) | >HORVU4Hr1G089500.1 gene=HORVU4Hr1G089500 | 671.2 | NRagged | 4 | 818.6256 | 818.6292 | K21(xgly / 324.1056) | 3.3e-13 | 12.48 | 551.1 | 1.8e-12 | 551.1 | -4.40
Spectrum (double click to dock / undock)
A.GAACEPAQLAVCASAILGGTK[+324]PSGECCGNLR.A z=4,scan#=sample=1 period=1 cycle=953 experiment=10,scan time=15.1684
GAACEPAQLAVCASAILGGTKPSGECCGNLR
Intensity
m/z
m/z errors, tol=75.0 ppm
Protein Coverage
Sample name
Protein name
11:12 AM
20/08/2020

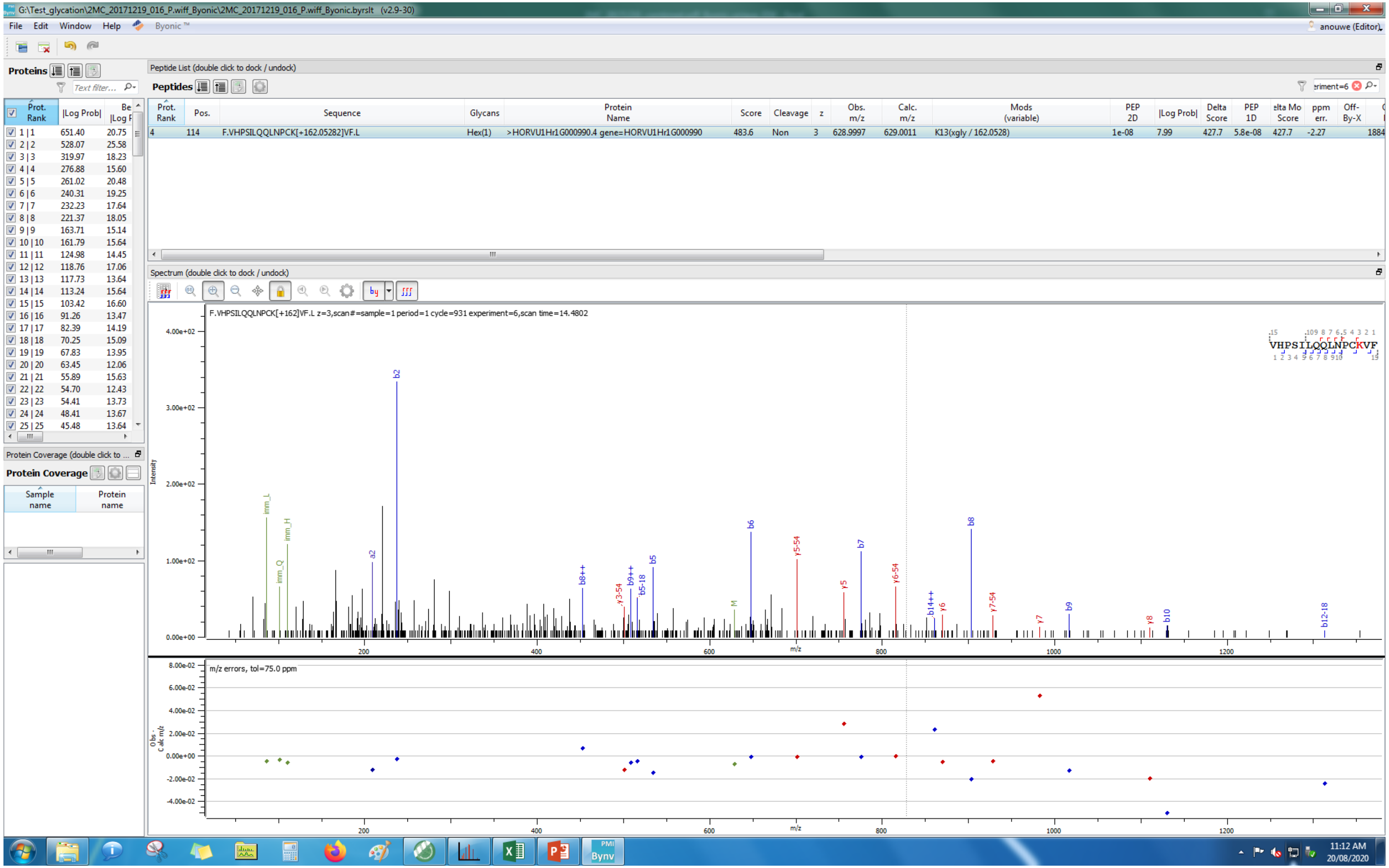
G:\Test_glycation\2MC_20171219_016_P.wiff_Byonic\2MC_20171219_016_P.wiff_Byonic.byrslt (v2.9-30)
File Edit Window Help Byonic ™
anouwe (Editor)
Proteins
Peptide List (double click to dock / undock)
Peptides
Prot. Rank | Pos. | Sequence | Glycans | Protein Name | Score | Cleavage | z | Obs. m/z | Calc. m/z | Mods (variable) | PEP 2D | |Log Prob| | Delta Score | PEP 1D | Delta Mod Score | ppm err. | Off-By-X
4 | 114 | F.VHPSILQQLNPCK[+162.05282]VF.L | Hex(1) | >HORVU1Hr1G000990.4 gene=HORVU1Hr1G000990 | 483.6 | Non | 3 | 628.9997 | 629.0011 | K13(xgly / 162.0528) | 1e-08 | 7.99 | 427.7 | 5.8e-08 | 427.7 | -2.27
Prot. Rank | |Log Prob| | Best |Log Prob|
1 | 1 651.40 20.75
2 | 2 528.07 25.58
3 | 3 319.97 18.23
4 | 4 276.88 15.60
5 | 5 261.02 20.48
6 | 6 240.31 19.25
7 | 7 232.23 17.64
8 | 8 221.37 18.05
9 | 9 163.71 15.14
10 | 10 161.79 15.64
11 | 11 124.98 14.45
12 | 12 118.76 17.06
13 | 13 117.73 13.64
14 | 14 113.24 15.64
15 | 15 103.42 16.60
16 | 16 91.26 13.47
17 | 17 82.39 14.19
18 | 18 70.25 15.09
19 | 19 67.83 13.95
20 | 20 63.45 12.06
21 | 21 55.89 15.63
22 | 22 54.70 12.43
23 | 23 54.41 13.73
24 | 24 48.41 13.67
25 | 25 45.48 13.64
Protein Coverage (double click to ...
Protein Coverage
Sample name
Protein name
Spectrum (double click to dock / undock)
F.VHPSILQQLNPCK[+162]VF.L z=3,scan#=sample=1 period=1 cycle=931 experiment=6,scan time=14.4802
VHPSILQQLNPCKVF
Intensity
m/z
m/z errors, tol=75.0 ppm
Obs - Calc m/z
11:12 AM
20/08/2020

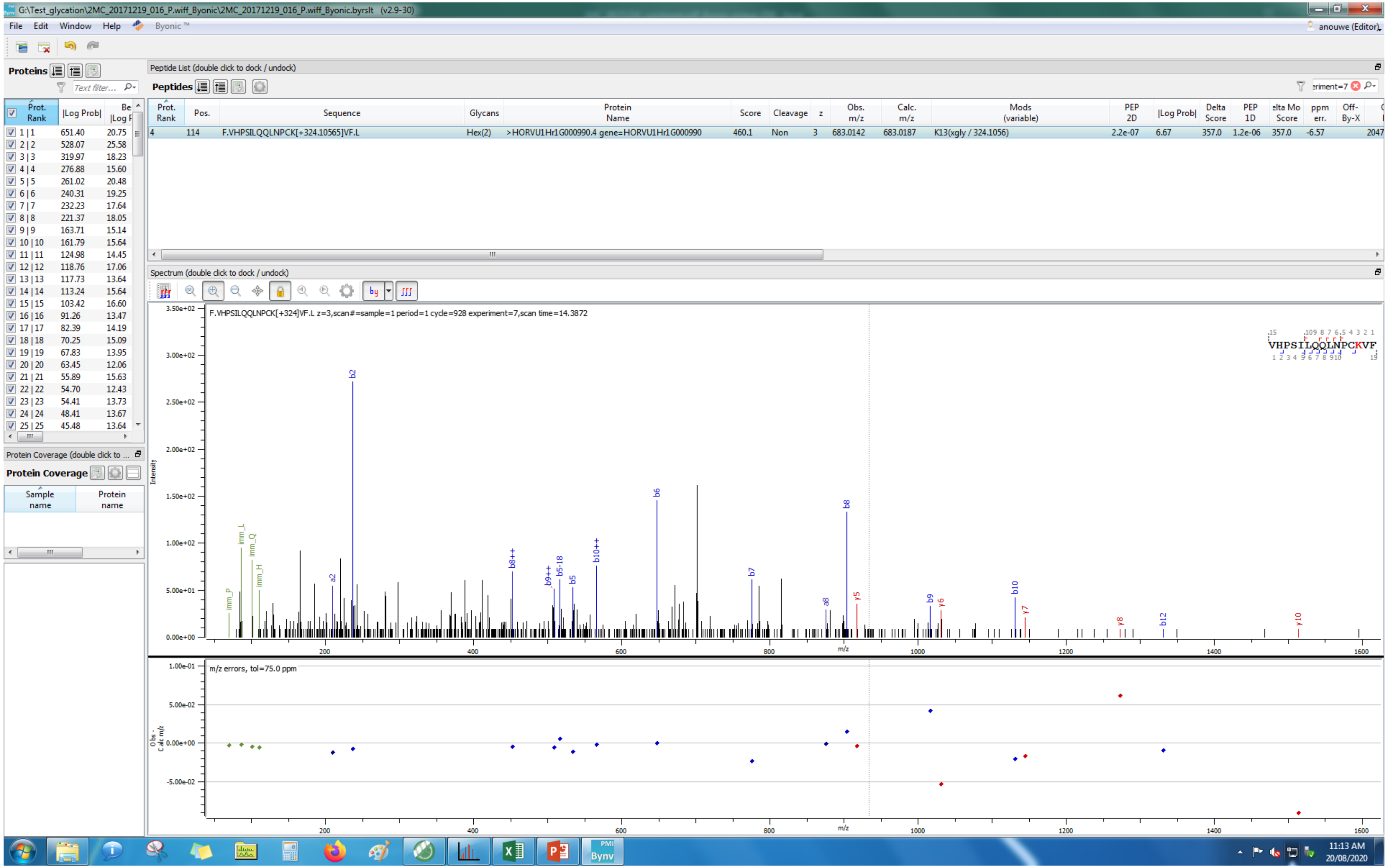
G:\Test_glycation\2MC_20171219_016_P.wiff_Byonic\2MC_20171219_016_P.wiff_Byonic.byrslt (v2.9-30)
File Edit Window Help Byonic
Proteins
Peptide List (double click to dock / undock)
Peptides
F.VHPSILQQLNPCK[+324.10565]VF.L
Hex(2)
>HORVU1Hr1G000990.4 gene=HORVU1Hr1G000990
K13(xgly / 324.1056)
Spectrum (double click to dock / undock)
F.VHPSILQQLNPCK[+324]VF.L z=3,scan#=sample=1 period=1 cycle=928 experiment=7,scan time=14.3872
VHPSILQQLNPCKVF
m/z errors, tol=75.0 ppm
Protein Coverage
11:13 AM
20/08/2020

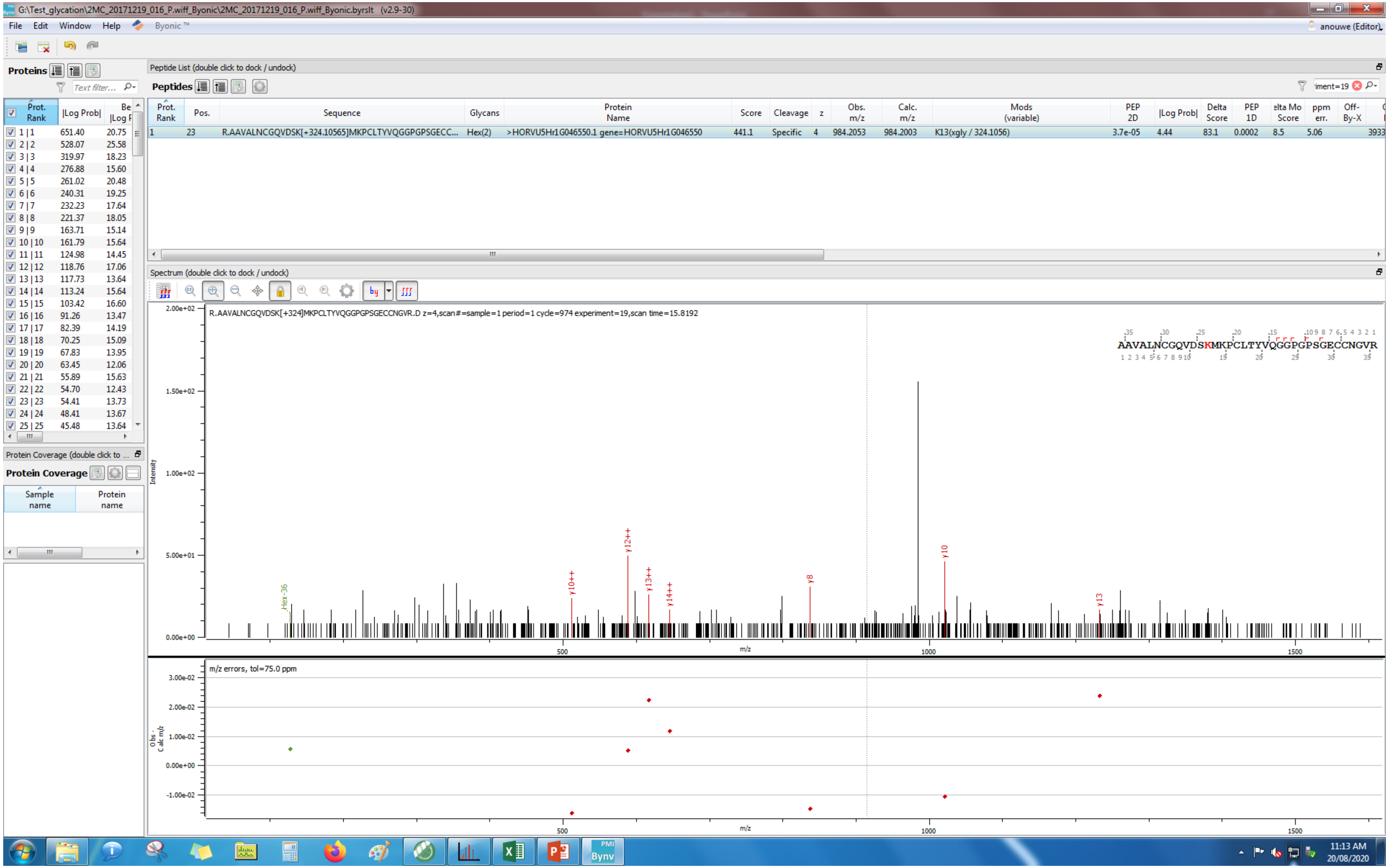
G:\Test_glycation\2MC_20171219_016_P.wiff_Byonic\2MC_20171219_016_P.wiff_Byonic.byrslt (v2.9-30)
File Edit Window Help Byonic ™
anouwe (Editor)
Proteins
Text filter...
Prot. Rank |Log Prob|
1 | 1 651.40 20.75
2 | 2 528.07 25.58
3 | 3 319.97 18.23
4 | 4 276.88 15.60
5 | 5 261.02 20.48
6 | 6 240.31 19.25
7 | 7 232.23 17.64
8 | 8 221.37 18.05
9 | 9 163.71 15.14
10 | 10 161.79 15.64
11 | 11 124.98 14.45
12 | 12 118.76 17.06
13 | 13 117.73 13.64
14 | 14 113.24 15.64
15 | 15 103.42 16.60
16 | 16 91.26 13.47
17 | 17 82.39 14.19
18 | 18 70.25 15.09
19 | 19 67.83 13.95
20 | 20 63.45 12.06
21 | 21 55.89 15.63
22 | 22 54.70 12.43
23 | 23 54.41 13.73
24 | 24 48.41 13.67
25 | 25 45.48 13.64
Protein Coverage (double click to ...
Protein Coverage
Sample name
Protein name
Peptide List (double click to dock / undock)
Peptides
Prot. Rank | Pos. | Sequence | Glycans | Protein Name | Score | Cleavage | z | Obs. m/z | Calc. m/z | Mods (variable) | PEP 2D | |Log Prob| | Delta Score | PEP 1D | Delta Mod Score | ppm err. | Off-By-X
1 | 23 | R.AAVALNCGQVDSK[+324.10565]MKPCLTYVQGGPGPSGECC... | Hex(2) | >HORVU5Hr1G046550.1 gene=HORVU5Hr1G046550 | 441.1 | Specific | 4 | 984.2053 | 984.2003 | K13(xgly / 324.1056) | 3.7e-05 | 4.44 | 83.1 | 0.0002 | 8.5 | 5.06 | 3933
Spectrum (double click to dock / undock)
R.AAVALNCGQVDSK[+324]MKPCLTYVQGGPGPSGECCNGVR z=4,scan#=sample=1 period=1 cycle=974 experiment=19,scan time=15.8192
AAVALNCGQVDSKMKPCLTYVQGGPGPSGECCNGVR
Intensity
2.00e+02
1.50e+02
1.00e+02
5.00e+01
0.00e+00
Hex-36
y10++
y12++
y13++
y14++
y8
y10
y13
500
m/z
1000
1500
m/z errors, tol=75.0 ppm
Obs - Calc m/z
3.00e-02
2.00e-02
1.00e-02
0.00e+00
-1.00e-02
500
m/z
1000
1500
11:13 AM
20/08/2020

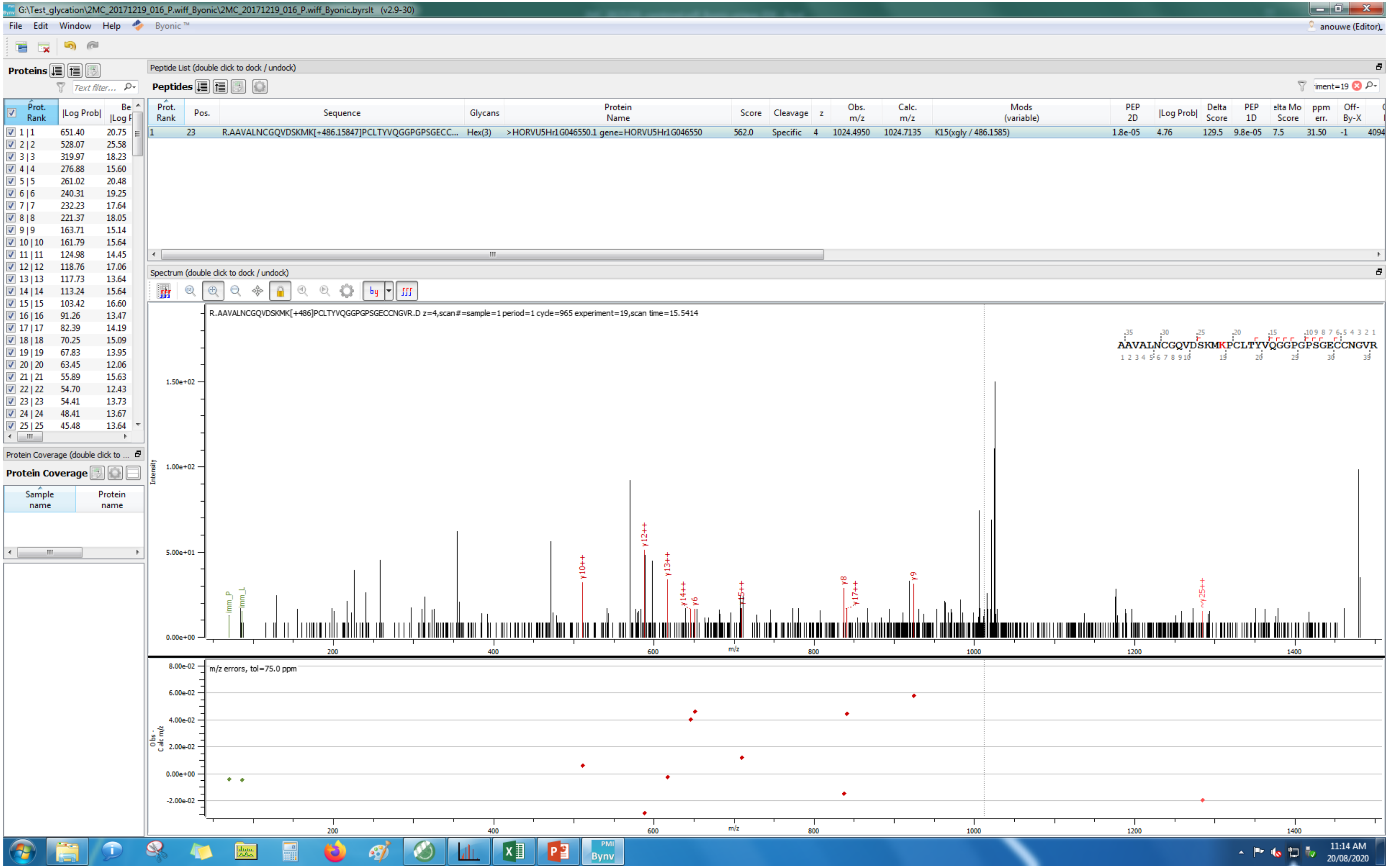
G:\Test_glycation\2MC_20171219_016_P.wiff_Byonic\2MC_20171219_016_P.wiff_Byonic.byrslt (v2.9-30)
Peptide List (double click to dock / undock)
1 23 R.AAVALNCGQVDSKMK[+486.15847]PCLTYVQGGPGPSGECC... Hex(3) >HORVU5Hr1G046550.1 gene=HORVU5Hr1G046550 562.0 Specific 4 1024.4950 1024.7135 K15(xgly / 486.1585) 1.8e-05 4.76 129.5 9.8e-05 7.5 31.50 -1
Spectrum (double click to dock / undock)
R.AAVALNCGQVDSKMK[+486]PCLTYVQGGPGPSGECCNGVR.D z=4,scan#=sample=1 period=1 cycle=965 experiment=19,scan time=15.5414
AAVALNCGQVDSKMKPCLTYVQGGPGPSGECCNGVR
m/z errors, tol=75.0 ppm

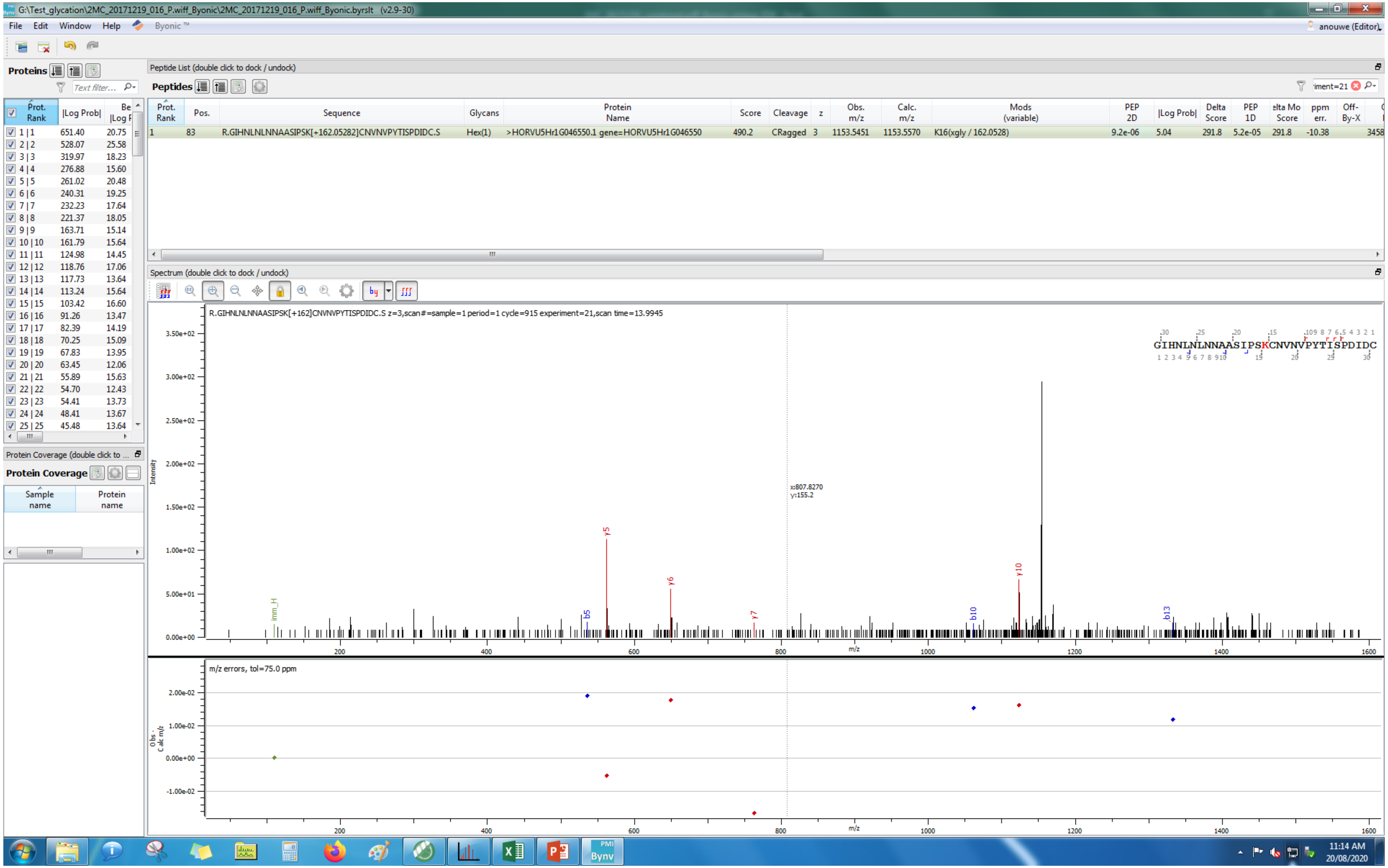
G:\Test_glycation\2MC_20171219_016_P.wiff_Byonic\2MC_20171219_016_P.wiff_Byonic.byrslt (v2.9-30)
File Edit Window Help Byonic™
anouwe (Editor)
Proteins
Peptide List (double click to dock / undock)
Peptides
Prot. Rank | Pos. | Sequence | Glycans | Protein Name | Score | Cleavage | z | Obs. m/z | Calc. m/z | Mods (variable) | PEP 2D | |Log Prob| | Delta Score | PEP 1D | Delta Mod Score | ppm err. | Off-By-X
1 | 83 | R.GIHNLNLNNAASIPSK[+162.05282]CNVNVPYTISPDIDC.S | Hex(1) | >HORVU5Hr1G046550.1 gene=HORVU5Hr1G046550 | 490.2 | CRagged | 3 | 1153.5451 | 1153.5570 | K16(xgly / 162.0528) | 9.2e-06 | 5.04 | 291.8 | 5.2e-05 | 291.8 | -10.38
Spectrum (double click to dock / undock)
R.GIHNLNLNNAASIPSK[+162]CNVNVPYTISPDIDC.S z=3,scan#=sample=1 period=1 cycle=915 experiment=21,scan time=13.9945
GIHNLNLNNAASIPSKCNVNVPYTISPDIDC
x:807.8270
y:155.2
Intensity
m/z
m/z errors, tol=75.0 ppm
Obs - Calc m/z
Protein Coverage (double click to ...
Protein Coverage
Sample name
Protein name
11:14 AM
20/08/2020

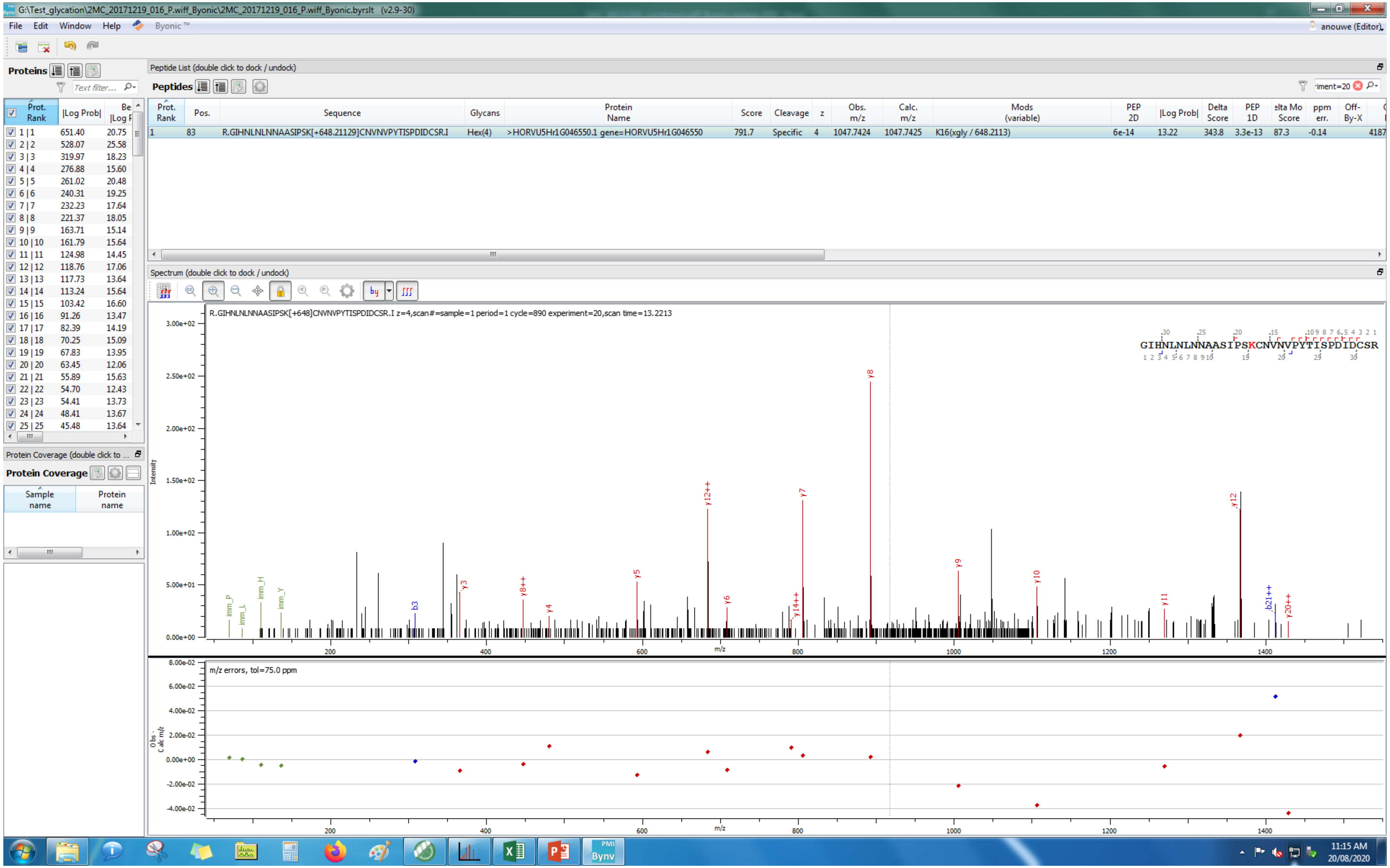
G:\Test_glycation\2MC_20171219_016_P.wiff_Byonic\2MC_20171219_016_P.wiff_Byonic.byrslt (v2.9-30)
File Edit Window Help Byonic™
anouwe (Editor)
Proteins
Peptide List (double click to dock / undock)
Peptides
Prot. Rank | Pos. | Sequence | Glycans | Protein Name | Score | Cleavage | z | Obs. m/z | Calc. m/z | Mods (variable) | PEP 2D | |Log Prob| | Delta Score | PEP 1D | ppm err.
1 | 83 | R.GIHNLNLNNAASIPSK[+648.21129]CNVNPYTISPDIDCSR.I | Hex(4) | >HORVU5Hr1G046550.1 gene=HORVU5Hr1G046550 | 791.7 | Specific | 4 | 1047.7424 | 1047.7425 | K16(xgly / 648.2113) | 6e-14 | 13.22 | 343.8 | 3.3e-13 | -0.14
Spectrum (double click to dock / undock)
R.GIHNLNLNNAASIPSK[+648]CNVNPYTISPDIDCSR.I z=4,scan#=sample=1 period=1 cycle=890 experiment=20,scan time=13.2213
GIHNLNLNNAASIPSKCNVNPYTISPDIDCSR
Intensity
m/z
m/z errors, tol=75.0 ppm
Obs - Calc m/z
Protein Coverage
Sample name
Protein name
11:15 AM
20/08/2020

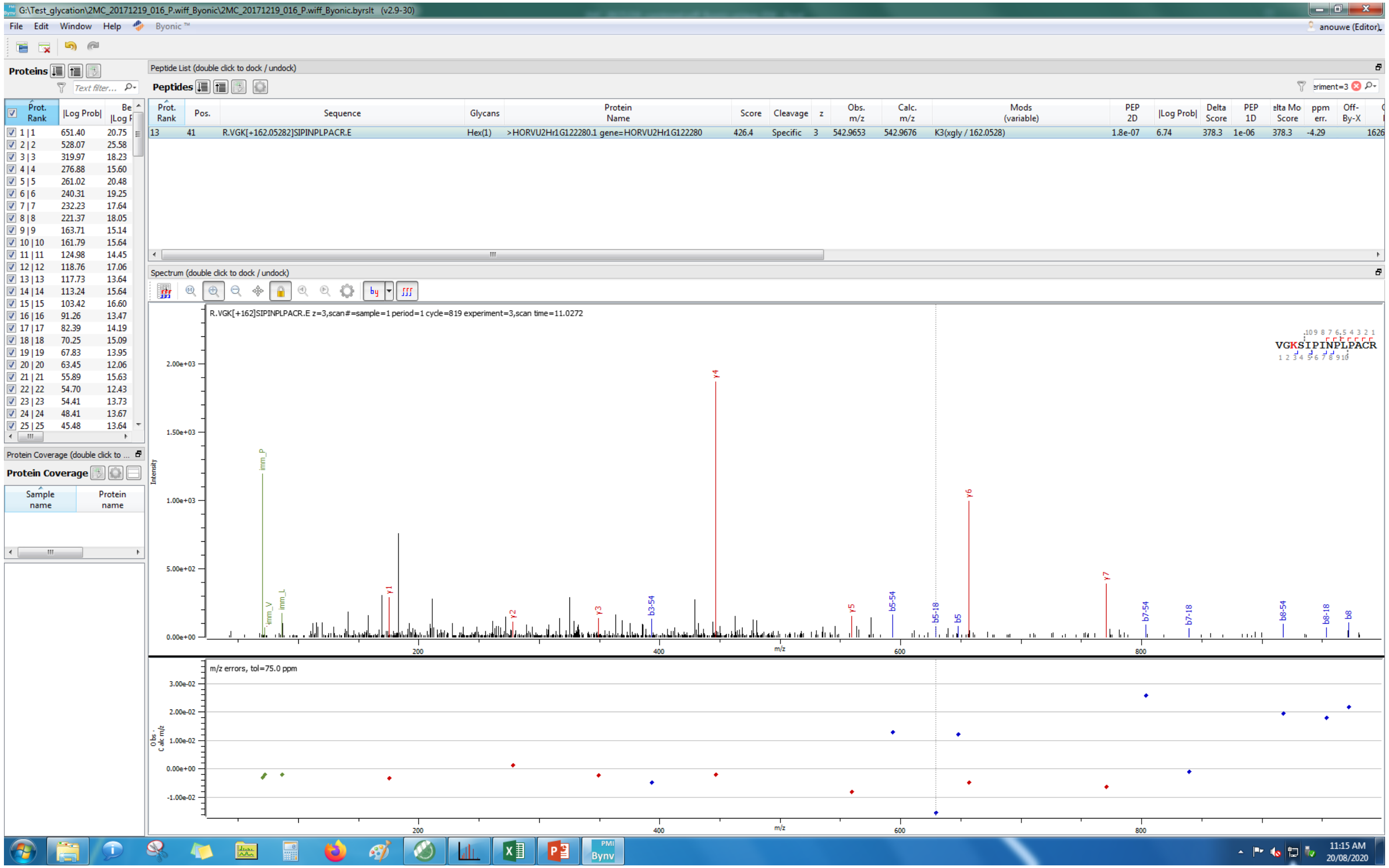
G:\Test_glycation\2MC_20171219_016_P.wiff_Byonic\2MC_20171219_016_P.wiff_Byonic.byrslt (v2.9-30)
File Edit Window Help Byonic ™
Proteins
Peptide List (double click to dock / undock)
Peptides
Prot. Rank | Pos. | Sequence | Glycans | Protein Name | Score | Cleavage | z | Obs. m/z | Calc. m/z | Mods (variable) | PEP 2D | |Log Prob| | Delta Score | PEP 1D | ppm err. | Off-By-X
13 | 41 | R.VGK[+162.05282]SIPINPLPACR.E | Hex(1) | >HORVU2Hr1G122280.1 gene=HORVU2Hr1G122280 | 426.4 | Specific | 3 | 542.9653 | 542.9676 | K3(xgly / 162.0528) | 1.8e-07 | 6.74 | 378.3 | 1e-06 | 378.3 | -4.29 | 1626
Spectrum (double click to dock / undock)
R.VGK[+162]SIPINPLPACR.E z=3,scan#=sample=1 period=1 cycle=819 experiment=3,scan time=11.0272
VGKSIPINPLPACR
Intensity
m/z
m/z errors, tol=75.0 ppm
Protein Coverage
Sample name
Protein name
anouwe (Editor)
11:15 AM
20/08/2020

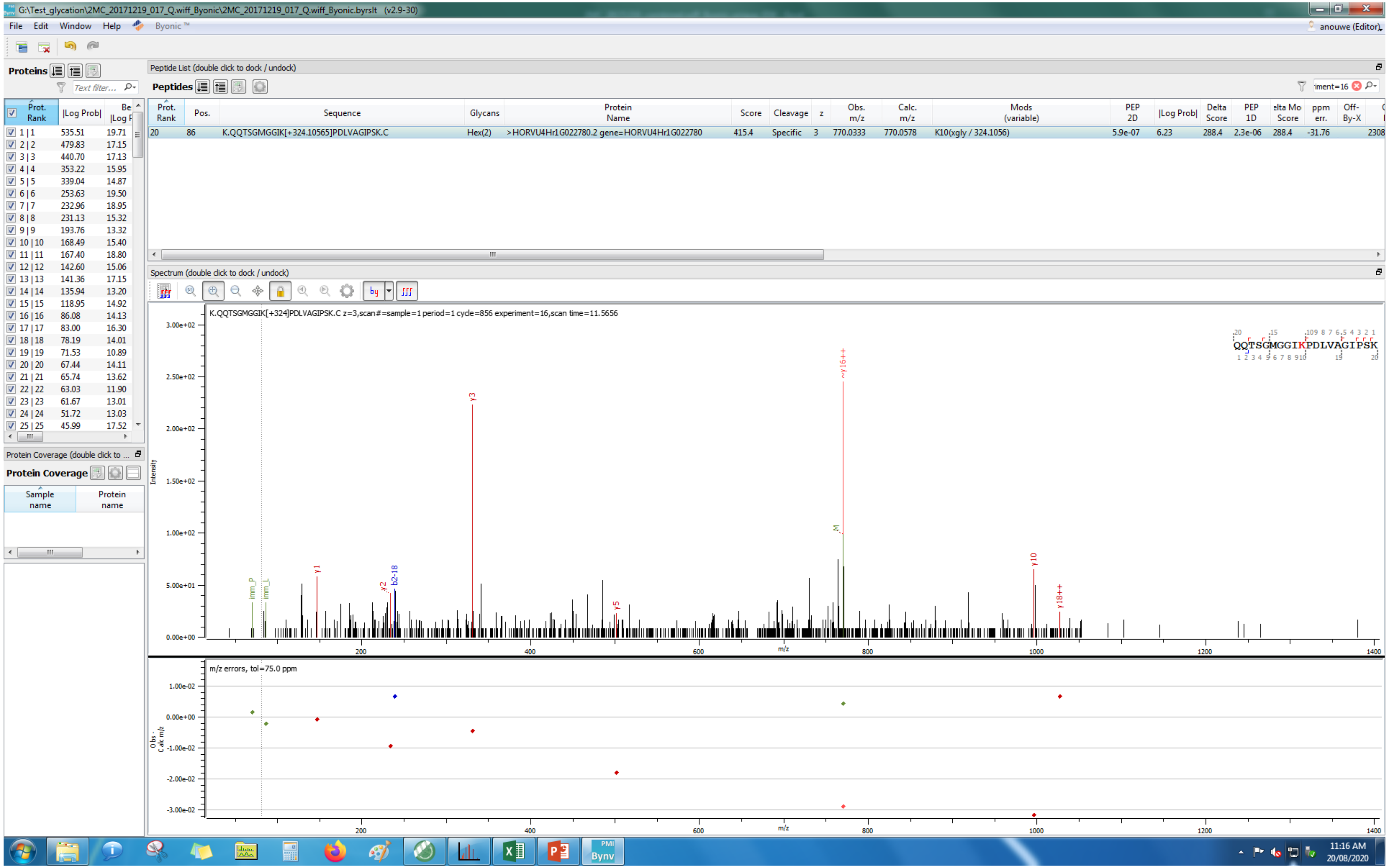
G:\Test_glycation\2MC_20171219_017_Q.wiff_Byonic\2MC_20171219_017_Q.wiff_Byonic.byrslt (v2.9-30)
File Edit Window Help Byonic™
anouwe (Editor)
Proteins
Peptide List (double click to dock / undock)
Peptides
Prot. Rank | Pos. | Sequence | Glycans | Protein Name | Score | Cleavage | z | Obs. m/z | Calc. m/z | Mods (variable) | PEP 2D | |Log Prob| | Delta Score | PEP 1D | ppm err. | Off-By-X
20 | 86 | K.QQTSGMGGIK[+324.10565]PDLVAGIPSK.C | Hex(2) | >HORVU4Hr1G022780.2 gene=HORVU4Hr1G022780 | 415.4 | Specific | 3 | 770.0333 | 770.0578 | K10(xgly / 324.1056) | 5.9e-07 | 6.23 | 288.4 | 2.3e-06 | 288.4 | -31.76
Spectrum (double click to dock / undock)
K.QQTSGMGGIK[+324]PDLVAGIPSK.C z=3,scan#=sample=1 period=1 cycle=856 experiment=16,scan time=11.5656
QQTSGMGGIKPDLVAGIPSK
Intensity
m/z
m/z errors, tol=75.0 ppm
Protein Coverage (double click to ...
Protein Coverage
Sample name
Protein name
11:16 AM
20/08/2020

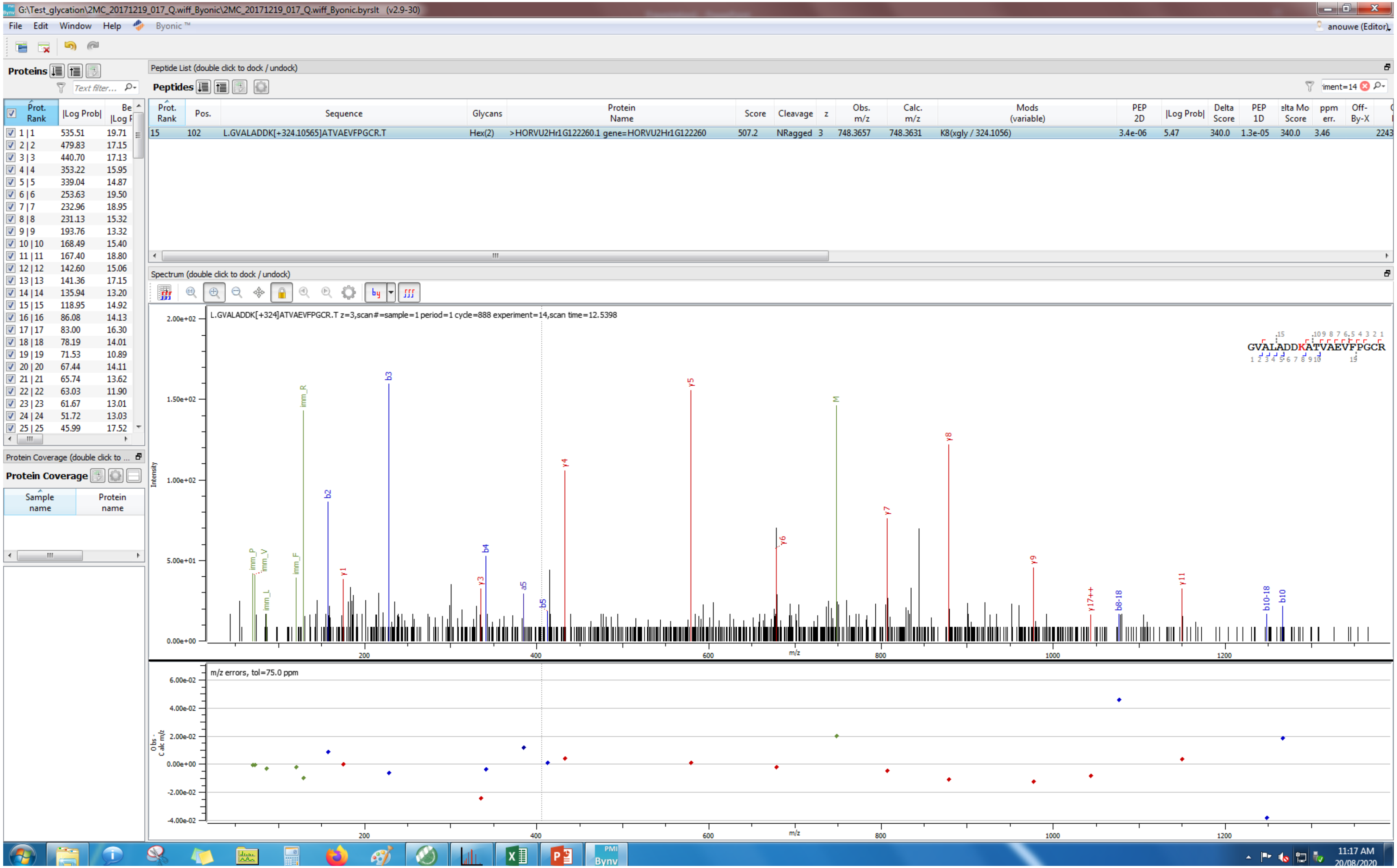
G:\Test_glycation\2MC_20171219_017_Q.wiff_Byonic\2MC_20171219_017_Q.wiff_Byonic.byrslt (v2.9-30)
File Edit Window Help Byonic ™
anouwe (Editor)
Proteins
Peptide List (double click to dock / undock)
Peptides
Prot. Rank |Log Prob| Be |Log F
Prot. Rank Pos. Sequence Glycans Protein Name Score Cleavage z Obs. m/z Calc. m/z Mods (variable) PEP 2D |Log Prob| Delta Score PEP 1D ppm err. Off-By-X
15 102 L.GVALADDK[+324.10565]ATVAEVFPGCR.T Hex(2) >HORVU2Hr1G122260.1 gene=HORVU2Hr1G122260 507.2 NRagged 3 748.3657 748.3631 K8(xgly / 324.1056) 3.4e-06 5.47 340.0 1.3e-05 340.0 3.46 2243
1 | 1 535.51 19.71
2 | 2 479.83 17.15
3 | 3 440.70 17.13
4 | 4 353.22 15.95
5 | 5 339.04 14.87
6 | 6 253.63 19.50
7 | 7 232.96 18.95
8 | 8 231.13 15.32
9 | 9 193.76 13.32
10 | 10 168.49 15.40
11 | 11 167.40 18.80
12 | 12 142.60 15.06
13 | 13 141.36 17.15
14 | 14 135.94 13.20
15 | 15 118.95 14.92
16 | 16 86.08 14.13
17 | 17 83.00 16.30
18 | 18 78.19 14.01
19 | 19 71.53 10.89
20 | 20 67.44 14.11
21 | 21 65.74 13.62
22 | 22 63.03 11.90
23 | 23 61.67 13.01
24 | 24 51.72 13.03
25 | 25 45.99 17.52
Protein Coverage (double click to ...
Protein Coverage
Sample name
Protein name
Spectrum (double click to dock / undock)
L.GVALADDK[+324]ATVAEVFPGCR.T z=3,scan#=sample=1 period=1 cycle=888 experiment=14,scan time=12.5398
GVALADDKATVAEVFPGCR
Intensity
m/z
imm_R b3 y5 M y8 y4 b2 y7 y6 b4 y9 imm_P imm_V imm_F y1 y3 a5 b5 imm_L y17++ b8-18 y11 b10-18 b10
m/z errors, tol=75.0 ppm
Obs - Calc m/z
11:17 AM
20/08/2020

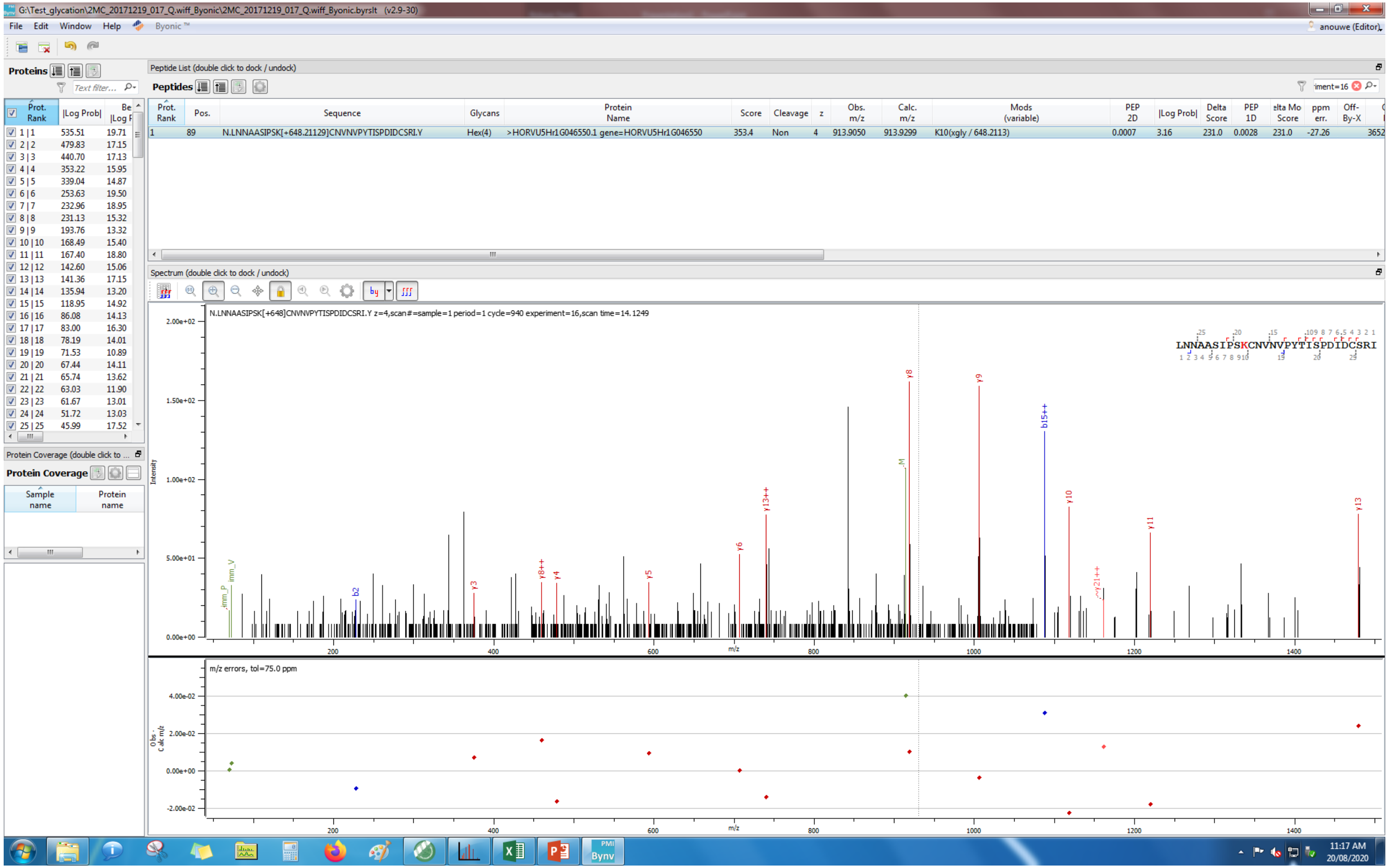
G:\Test_glycation\2MC_20171219_017_Q.wiff_Byonic\2MC_20171219_017_Q.wiff_Byonic.byrslt (v2.9-30)
File Edit Window Help Byonic ™
anouwe (Editor)
Proteins
Peptide List (double click to dock / undock)
Peptides
Prot. Rank | Pos. | Sequence | Glycans | Protein Name | Score | Cleavage | z | Obs. m/z | Calc. m/z | Mods (variable) | PEP 2D | |Log Prob| | Delta Score | PEP 1D | Delta Mod Score | ppm err.
1 | 89 | N.LNNAASIPSK[+648.21129]CNVNVPYTISPDIDCSRI.Y | Hex(4) | >HORVU5Hr1G046550.1 gene=HORVU5Hr1G046550 | 353.4 | Non | 4 | 913.9050 | 913.9299 | K10(xgly / 648.2113) | 0.0007 | 3.16 | 231.0 | 0.0028 | 231.0 | -27.26
Spectrum (double click to dock / undock)
N.LNNAASIPSK[+648]CNVNVPYTISPDIDCSRI.Y z=4,scan#=sample=1 period=1 cycle=940 experiment=16,scan time=14.1249
LNNAASIPSKCNVNVPYTISPDIDCSRI
Intensity
m/z
m/z errors, tol=75.0 ppm
Protein Coverage
Sample name
Protein name

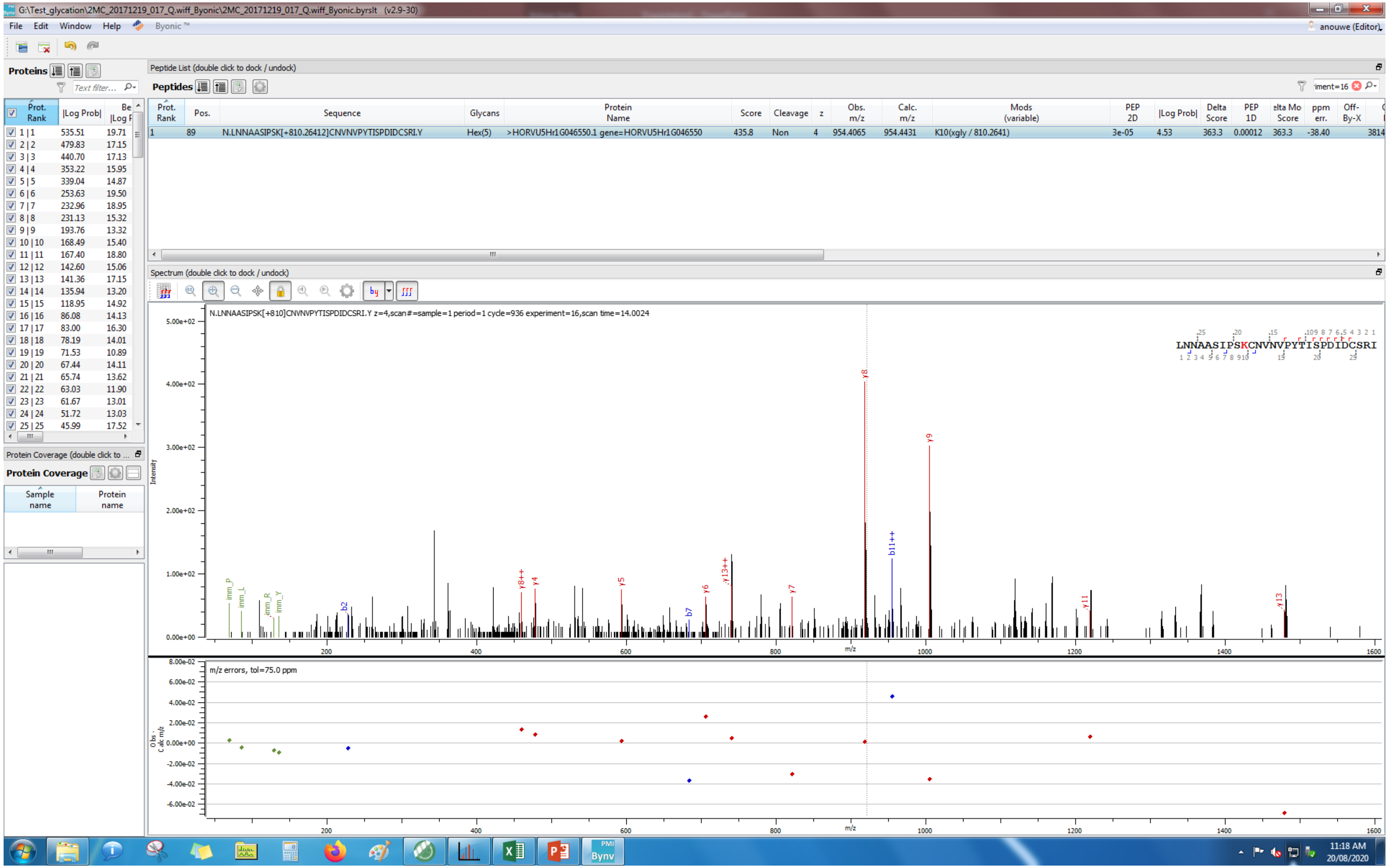
G:\Test_glycation\2MC_20171219_017_Q.wiff_Byonic\2MC_20171219_017_Q.wiff_Byonic.byrslt (v2.9-30)
File Edit Window Help Byonic™
Proteins
Peptide List (double click to dock / undock)
Peptides
Prot. Rank
Pos.
Sequence
Glycans
Protein Name
Score
Cleavage
z
Obs. m/z
Calc. m/z
Mods (variable)
PEP 2D
|Log Prob|
Delta Score
PEP 1D
ppm err.
Off-By-X
1 89 N.LNNAASIPSK[+810.26412]CNVNVPYTISPDIDCSRI.Y Hex(5) >HORVU5Hr1G046550.1 gene=HORVU5Hr1G046550 435.8 Non 4 954.4065 954.4431 K10(xgly / 810.2641) 3e-05 4.53 363.3 0.00012 363.3 -38.40
Spectrum (double click to dock / undock)
N.LNNAASIPSK[+810]CNVNVPYTISPDIDCSRI.Y z=4,scan#=sample=1 period=1 cycle=936 experiment=16,scan time=14.0024
LNNAASIPSKCNVNVPYTISPDIDCSRI
Intensity
m/z
m/z errors, tol=75.0 ppm
Protein Coverage
Sample name
Protein name
11:18 AM
20/08/2020

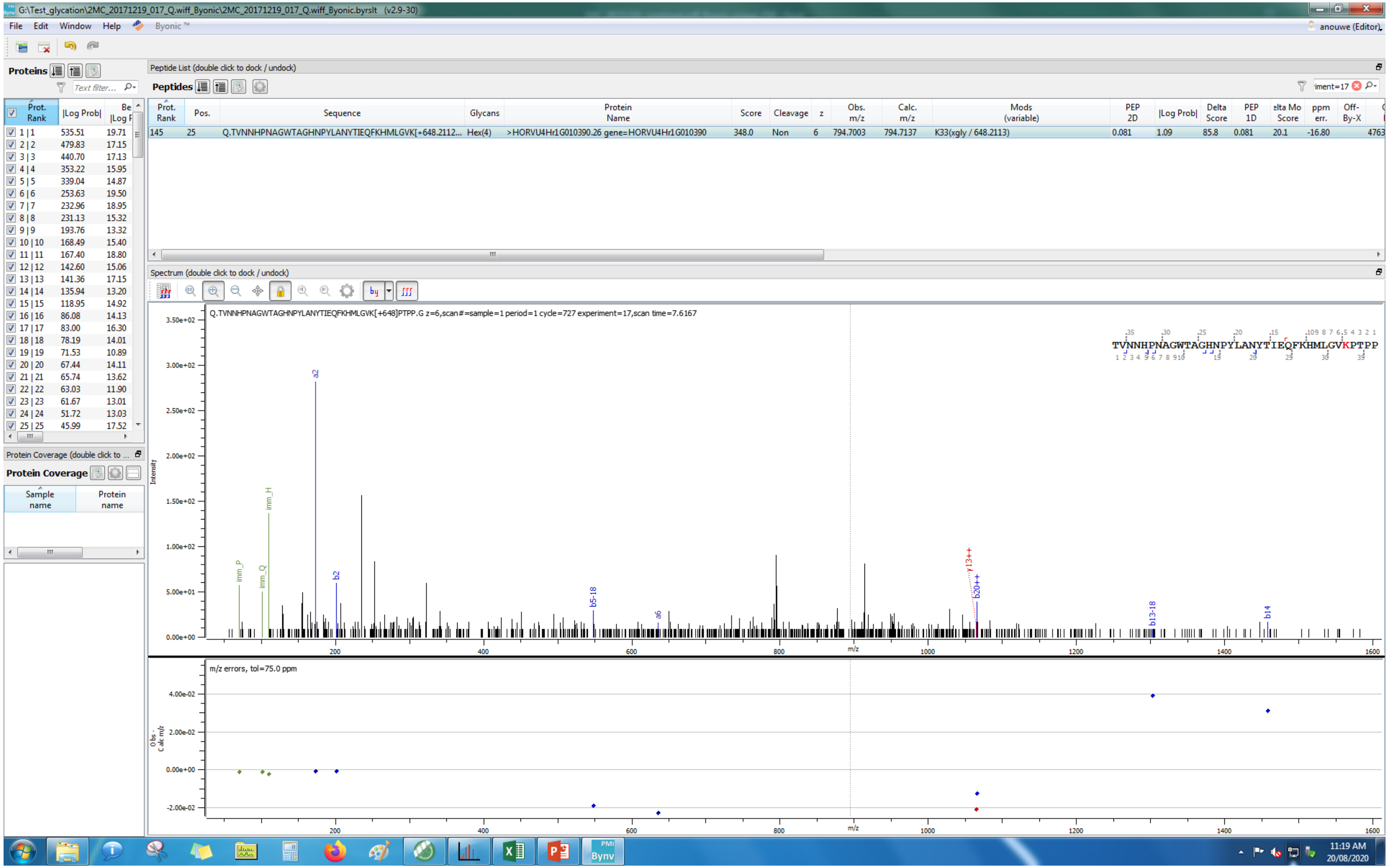

G:\Test_glycation\2MC_20171219_017_Q.wiff_Byonic\2MC_20171219_017_Q.wiff_Byonic.byrslt (v2.9-30)
File Edit Window Help Byonic™
anouwe (Editor)
Proteins
Peptide List (double click to dock / undock)
Peptides
Prot. Rank | Pos. | Sequence | Glycans | Protein Name | Score | Cleavage | z | Obs. m/z | Calc. m/z | Mods (variable) | PEP 2D | |Log Prob| | Delta Score | PEP 1D | elta Mo Score | ppm err. | Off-By-X
145 | 25 | Q.TVNNHPNAGWTAGHNPYLANYTIEQFKHMLGVK[+648.2112... | Hex(4) | >HORVU4Hr1G010390.26 gene=HORVU4Hr1G010390 | 348.0 | Non | 6 | 794.7003 | 794.7137 | K33(xgly / 648.2113) | 0.081 | 1.09 | 85.8 | 0.081 | 20.1 | -16.80 | 4763
Protein Coverage (double click to ...
Protein Coverage
Sample name
Protein name
Spectrum (double click to dock / undock)
Q.TVNNHPNAGWTAGHNPYLANYTIEQFKHMLGVK[+648]PTPP.G z=6,scan#=sample=1 period=1 cycle=727 experiment=17,scan time=7.6167
TVNNHPNAGWTAGHNPYLANYTIEQFKHMLGVKPTPP
Intensity
imm_P
imm_Q
imm_H
a2
b2
b5-18
a6
y13++
b20++
b13-18
b14
m/z
m/z errors, tol=75.0 ppm
Obs - Calc m/z
11:19 AM
20/08/2020

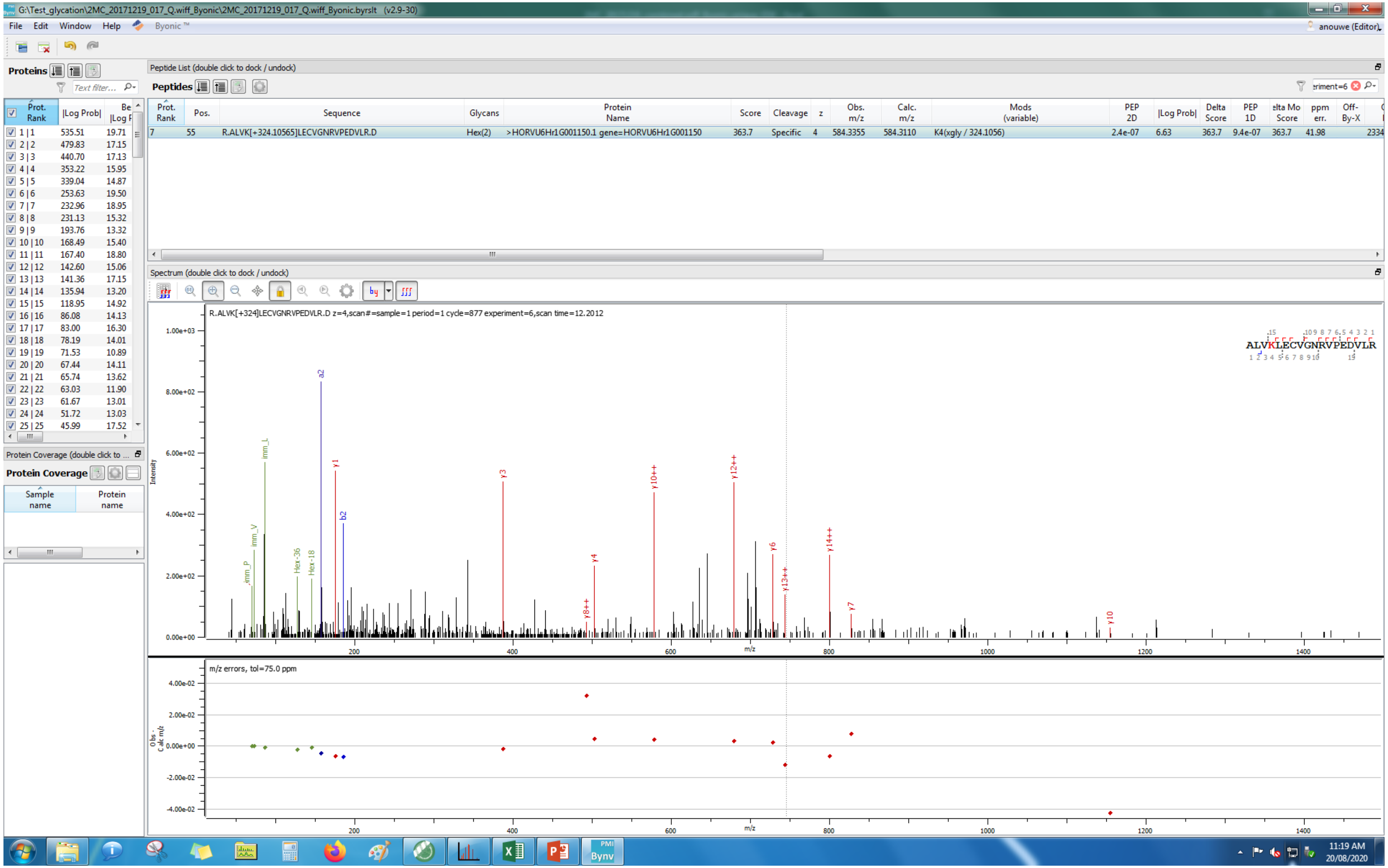
G:\Test_glycation\2MC_20171219_017_Q.wiff_Byonic\2MC_20171219_017_Q.wiff_Byonic.byrslt (v2.9-30)
File Edit Window Help Byonic™
anouwe (Editor)
Proteins
Peptide List (double click to dock / undock)
Peptides
Prot. Rank | Pos. | Sequence | Glycans | Protein Name | Score | Cleavage | z | Obs. m/z | Calc. m/z | Mods (variable) | PEP 2D | |Log Prob| | Delta Score | PEP 1D | Delta Mod Score | ppm err. | Off-By-X
7 | 55 | R.ALVK[+324.10565]LECVGNRVPEDVLR.D | Hex(2) | >HORVU6Hr1G001150.1 gene=HORVU6Hr1G001150 | 363.7 | Specific | 4 | 584.3355 | 584.3110 | K4(xgly / 324.1056) | 2.4e-07 | 6.63 | 363.7 | 9.4e-07 | 363.7 | 41.98 | 2334
Prot. Rank | |Log Prob| | Best |Log P|
1 | 1 535.51 19.71
2 | 2 479.83 17.15
3 | 3 440.70 17.13
4 | 4 353.22 15.95
5 | 5 339.04 14.87
6 | 6 253.63 19.50
7 | 7 232.96 18.95
8 | 8 231.13 15.32
9 | 9 193.76 13.32
10 | 10 168.49 15.40
11 | 11 167.40 18.80
12 | 12 142.60 15.06
13 | 13 141.36 17.15
14 | 14 135.94 13.20
15 | 15 118.95 14.92
16 | 16 86.08 14.13
17 | 17 83.00 16.30
18 | 18 78.19 14.01
19 | 19 71.53 10.89
20 | 20 67.44 14.11
21 | 21 65.74 13.62
22 | 22 63.03 11.90
23 | 23 61.67 13.01
24 | 24 51.72 13.03
25 | 25 45.99 17.52
Protein Coverage (double click to ...
Protein Coverage
Sample name
Protein name
Spectrum (double click to dock / undock)
R.ALVK[+324]LECVGNRVPEDVLR.D z=4,scan#=sample=1 period=1 cycle=877 experiment=6,scan time=12.2012
ALVKLECVGNRVPEDVLR
Intensity
m/z
imm_P
imm_V
imm_L
Hex-36
Hex-18
a2
b2
y1
y3
y4
y8++
y10++
y12++
y6
y13++
y14++
y7
y10
m/z errors, tol=75.0 ppm
Obs - Calc m/z
11:19 AM
20/08/2020

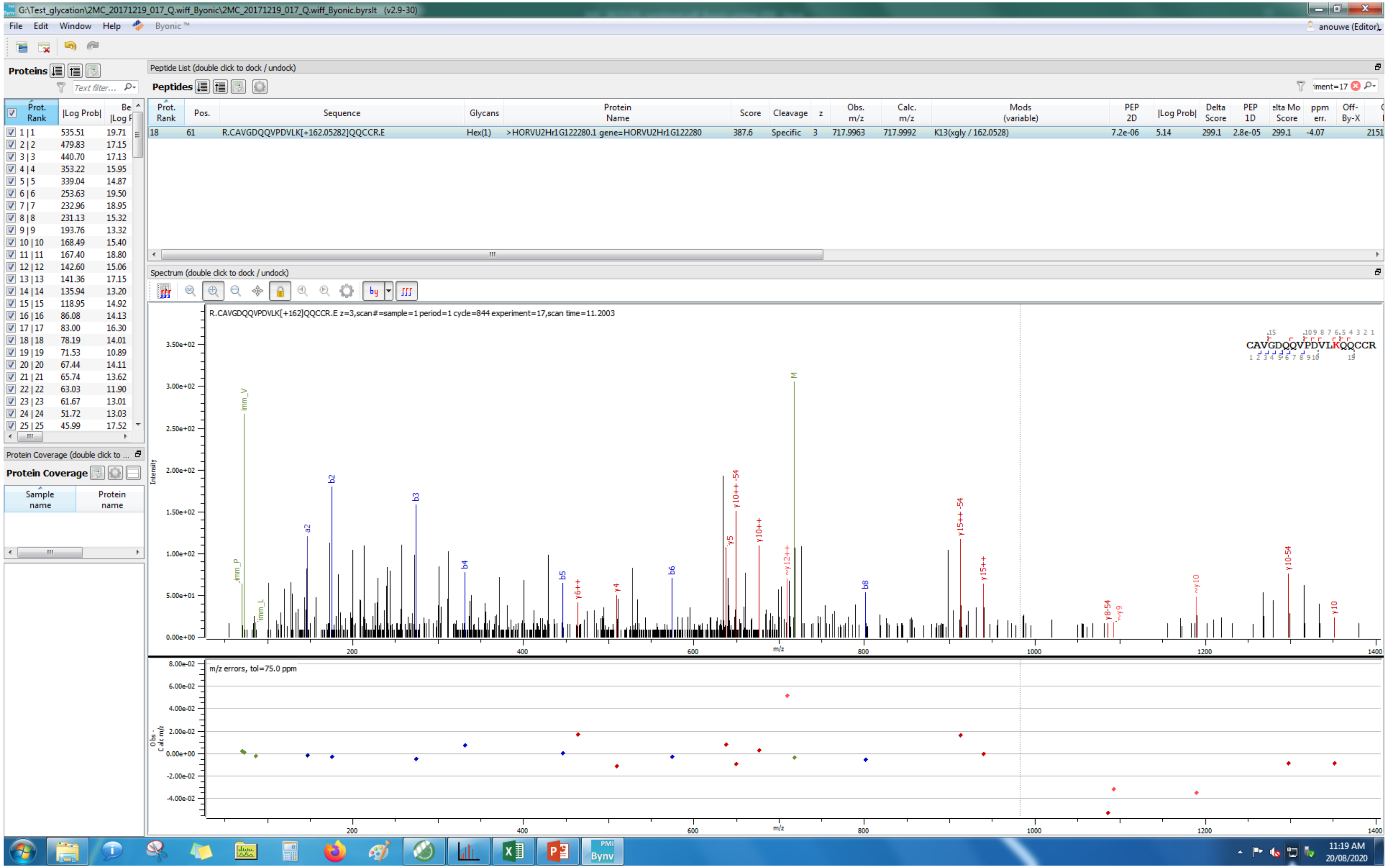
G:\Test_glycation\2MC_20171219_017_Q.wiff_Byonic\2MC_20171219_017_Q.wiff_Byonic.byrslt (v2.9-30)
File Edit Window Help Byonic™
anouwe (Editor)
Proteins
Peptide List (double click to dock / undock)
Peptides
Prot. Rank | Pos. | Sequence | Glycans | Protein Name | Score | Cleavage | z | Obs. m/z | Calc. m/z | Mods (variable) | PEP 2D | |Log Prob| | Delta Score | PEP 1D | Delta Mod Score | ppm err. | Off-By-X
18 | 61 | R.CAVGDQQVPDVLK[+162.05282]QQCCR.E | Hex(1) | >HORVU2Hr1G122280.1 gene=HORVU2Hr1G122280 | 387.6 | Specific | 3 | 717.9963 | 717.9992 | K13(xgly / 162.0528) | 7.2e-06 | 5.14 | 299.1 | 2.8e-05 | 299.1 | -4.07 | 2151
Spectrum (double click to dock / undock)
R.CAVGDQQVPDVLK[+162]QQCCR.E z=3,scan#=sample=1 period=1 cycle=844 experiment=17,scan time=11.2003
CAVGDQQVPDVLKQQCCR
m/z errors, tol=75.0 ppm
Protein Coverage (double click to ...
Protein Coverage
Sample name
Protein name
11:19 AM
20/08/2020

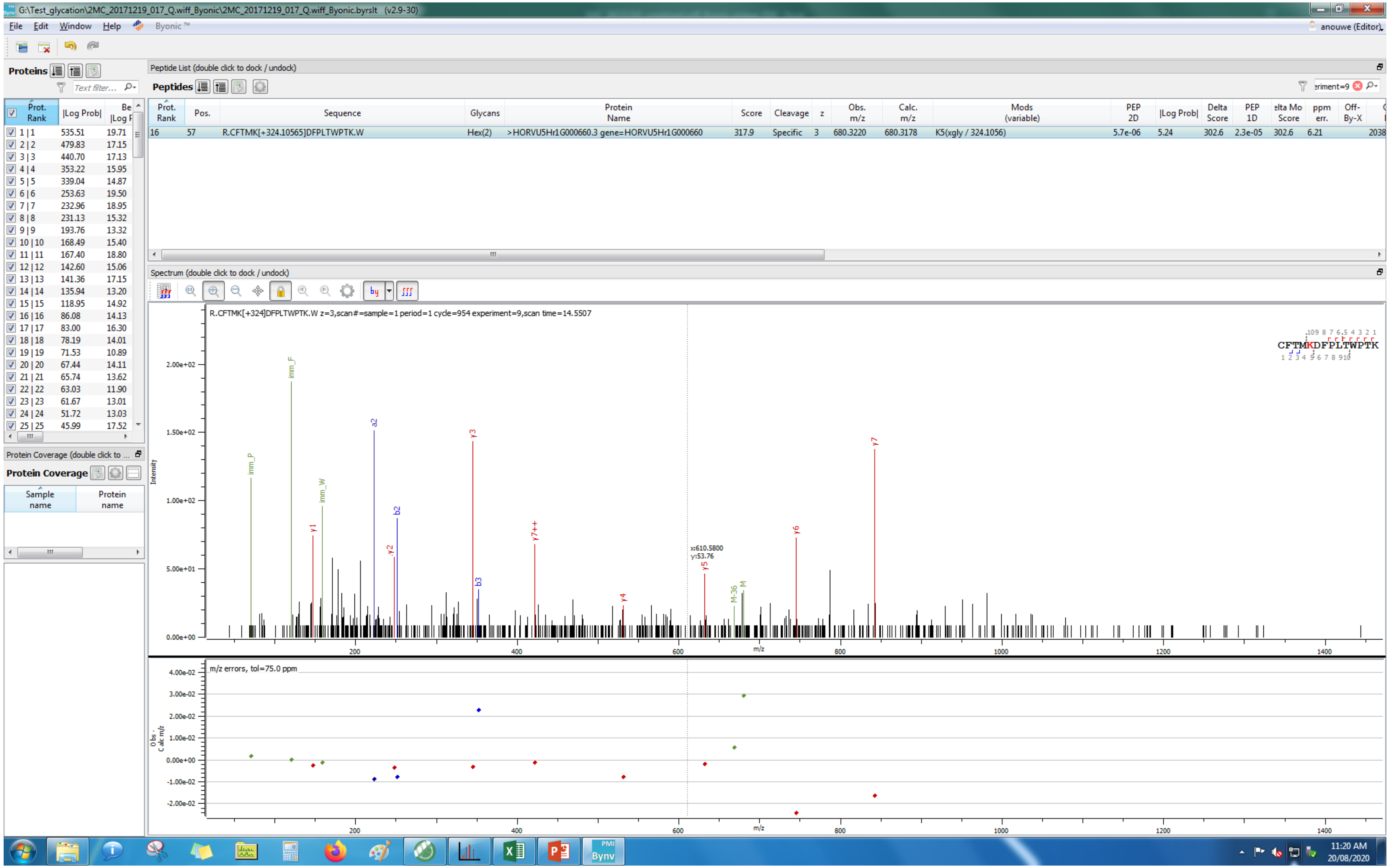
G:\Test_glycation\2MC_20171219_017_Q.wiff_Byonic\2MC_20171219_017_Q.wiff_Byonic.byrslt (v2.9-30)
File Edit Window Help Byonic™
anouwe (Editor)
Proteins
Peptide List (double click to dock / undock)
Peptides
Prot. Rank | Pos. | Sequence | Glycans | Protein Name | Score | Cleavage | z | Obs. m/z | Calc. m/z | Mods (variable) | PEP 2D | |Log Prob| | Delta Score | PEP 1D | Delta Mod Score | ppm err. | Off-By-X
16 | 57 | R.CFTMK[+324.10565]DFPLTWPTK.W | Hex(2) | >HORVU5Hr1G000660.3 gene=HORVU5Hr1G000660 | 317.9 | Specific | 3 | 680.3220 | 680.3178 | K5(xgly / 324.1056) | 5.7e-06 | 5.24 | 302.6 | 2.3e-05 | 302.6 | 6.21
Spectrum (double click to dock / undock)
R.CFTMK[+324]DFPLTWPTK.W z=3,scan#=sample=1 period=1 cycle=954 experiment=9,scan time=14.5507
CFTMKDFPLTWPTK
Intensity
m/z
m/z errors, tol=75.0 ppm
Obs - Calc m/z
Protein Coverage (double click to ...
Protein Coverage
Sample name
Protein name
11:20 AM
20/08/2020

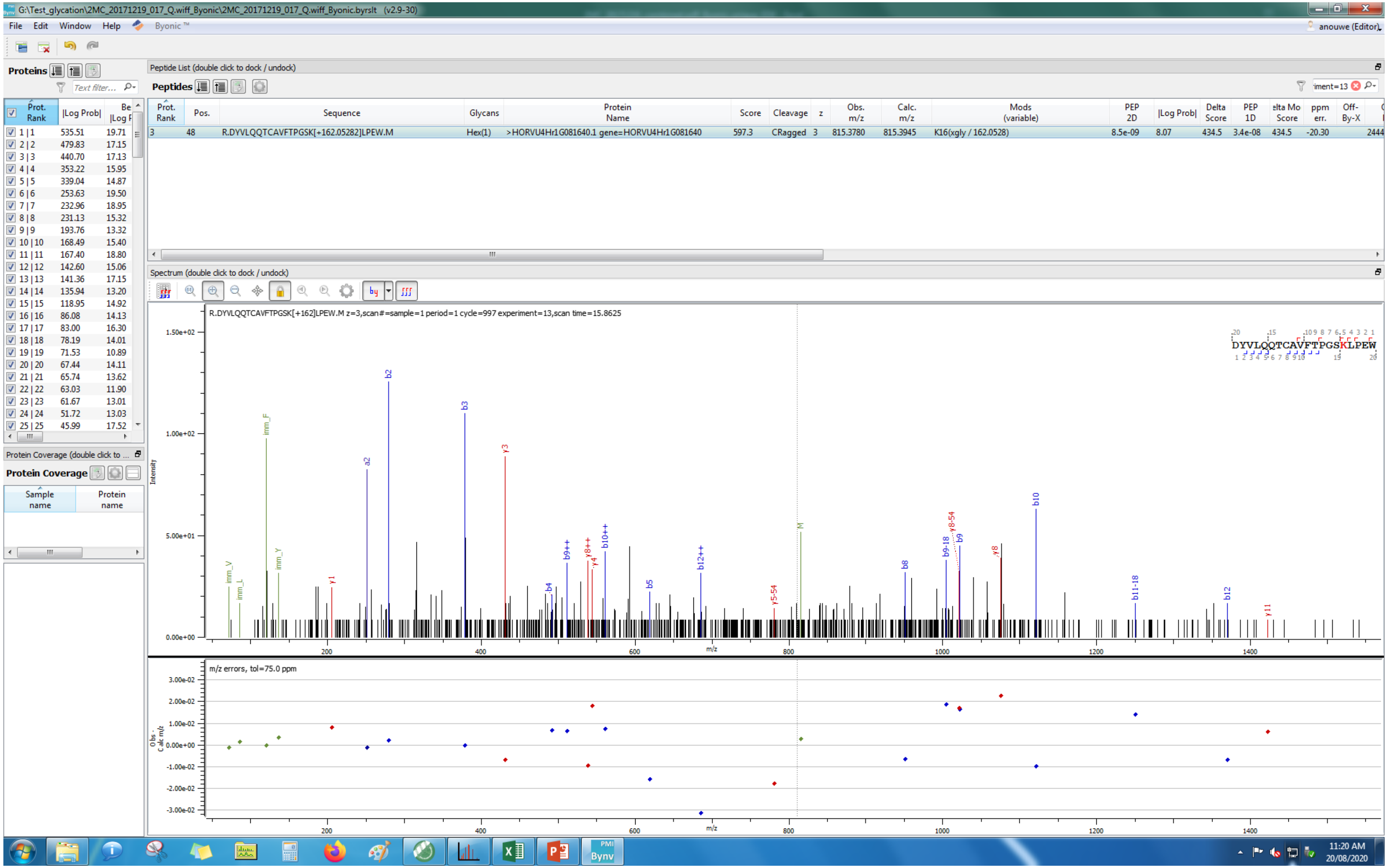
G:\Test_glycation\2MC_20171219_017_Q.wiff_Byonic\2MC_20171219_017_Q.wiff_Byonic.byrslt (v2.9-30)
File Edit Window Help Byonic™
Proteins
Peptide List (double click to dock / undock)
Peptides
Prot. Rank | Pos. | Sequence | Glycans | Protein Name | Score | Cleavage | z | Obs. m/z | Calc. m/z | Mods (variable) | PEP 2D | |Log Prob| | Delta Score | PEP 1D | Delta Mod Score | ppm err. | Off-By-X
3 | 48 | R.DYVLQQTCAVFTPGSK[+162.05282]LPEW.M | Hex(1) | >HORVU4Hr1G081640.1 gene=HORVU4Hr1G081640 | 597.3 | CRagged | 3 | 815.3780 | 815.3945 | K16(xgly / 162.0528) | 8.5e-09 | 8.07 | 434.5 | 3.4e-08 | 434.5 | -20.30
Spectrum (double click to dock / undock)
R.DYVLQQTCAVFTPGSK[+162]LPEW.M z=3,scan#=sample=1 period=1 cycle=997 experiment=13,scan time=15.8625
DYVLQQTCAVFTPGSKLPEW
Intensity
m/z
m/z errors, tol=75.0 ppm
Protein Coverage (double click to ...
Protein Coverage
Sample name
Protein name
11:20 AM
20/08/2020

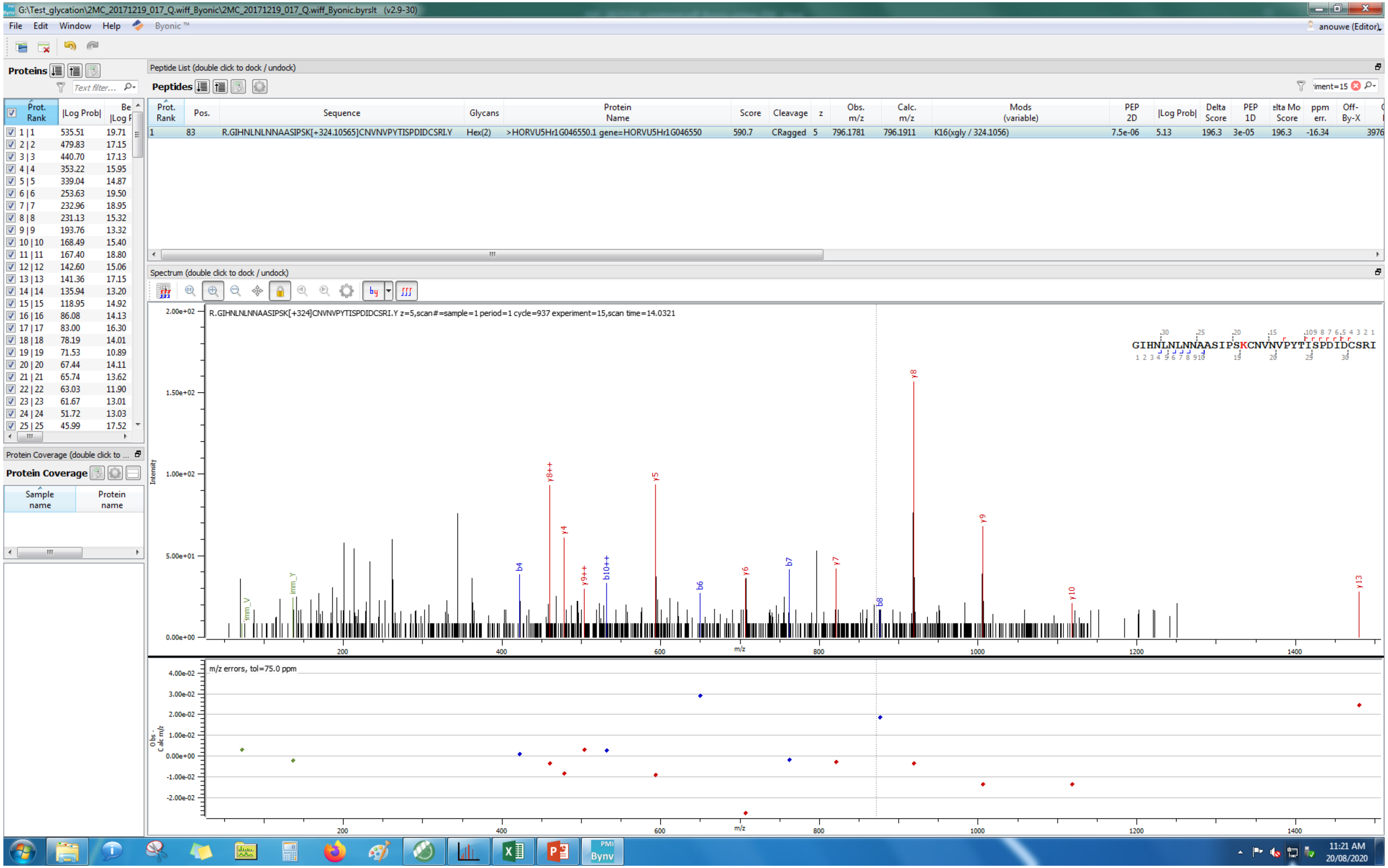

G:\Test_glycation\2MC_20171219_017_Q.wiff_Byonic\2MC_20171219_017_Q.wiff_Byonic.byrslt (v2.9-30)
File Edit Window Help Byonic™
anouwe (Editor)
Proteins
Text filter...
Prot. Rank |Log Prob| Best |Log P
1 | 1 535.51 19.71
2 | 2 479.83 17.15
3 | 3 440.70 17.13
4 | 4 353.22 15.95
5 | 5 339.04 14.87
6 | 6 253.63 19.50
7 | 7 232.96 18.95
8 | 8 231.13 15.32
9 | 9 193.76 13.32
10 | 10 168.49 15.40
11 | 11 167.40 18.80
12 | 12 142.60 15.06
13 | 13 141.36 17.15
14 | 14 135.94 13.20
15 | 15 118.95 14.92
16 | 16 86.08 14.13
17 | 17 83.00 16.30
18 | 18 78.19 14.01
19 | 19 71.53 10.89
20 | 20 67.44 14.11
21 | 21 65.74 13.62
22 | 22 63.03 11.90
23 | 23 61.67 13.01
24 | 24 51.72 13.03
25 | 25 45.99 17.52
Protein Coverage (double click to ...
Protein Coverage
Sample name Protein name
Peptide List (double click to dock / undock)
Peptides
iment=15
Prot. Rank | Pos. | Sequence | Glycans | Protein Name | Score | Cleavage | z | Obs. m/z | Calc. m/z | Mods (variable) | PEP 2D | |Log Prob| | Delta Score | PEP 1D | elta Mo Score | ppm err. | Off-By-X
1 | 83 | R.GIHNLNLNNAASIPSK[+324.10565]CNVNVPYTISPDIDCSRI.Y | Hex(2) | >HORVU5Hr1G046550.1 gene=HORVU5Hr1G046550 | 590.7 | CRagged | 5 | 796.1781 | 796.1911 | K16(xgly / 324.1056) | 7.5e-06 | 5.13 | 196.3 | 3e-05 | 196.3 | -16.34 | | 3976
Spectrum (double click to dock / undock)
R.GIHNLNLNNAASIPSK[+324]CNVNVPYTISPDIDCSRI.Y z=5,scan#=sample=1 period=1 cycle=937 experiment=15,scan time=14.0321
GIHNLNLNNAASIPSKCNVNVPYTISPDIDCSRI
Intensity
2.00e+02
1.50e+02
1.00e+02
5.00e+01
0.00e+00
m/z
200 400 600 800 1000 1200 1400
y8 y8++ y5 y9 y4 y9++ b10++ b4 b6 y6 b7 y7 b8 y10 y13 Imm_Y Imm_V
m/z errors, tol=75.0 ppm
Obs - Calc m/z
4.00e-02
3.00e-02
2.00e-02
1.00e-02
0.00e+00
-1.00e-02
-2.00e-02
11:21 AM
20/08/2020

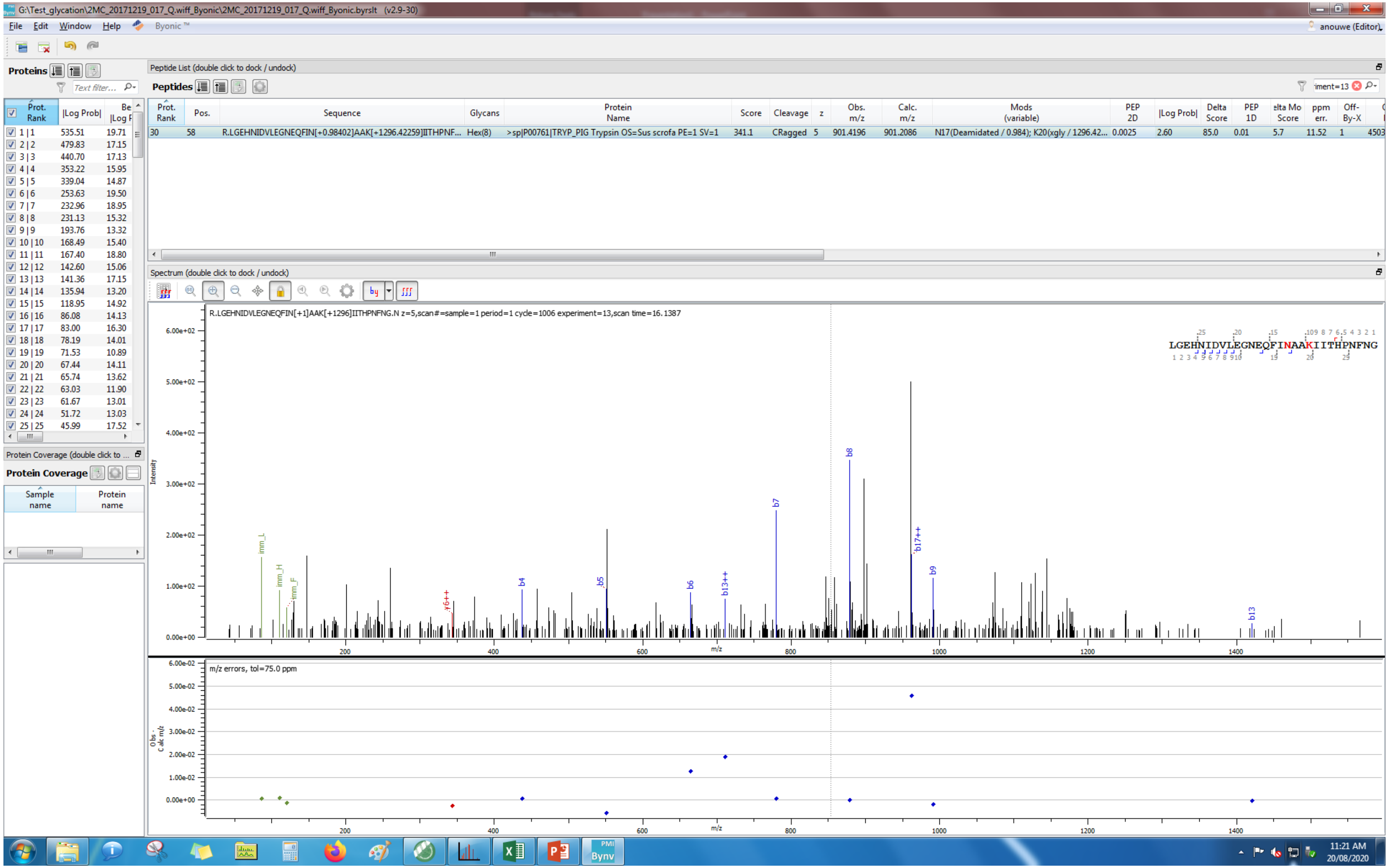
G:\Test_glycation\2MC_20171219_017_Q.wiff_Byonic\2MC_20171219_017_Q.wiff_Byonic.byrslt (v2.9-30)
File Edit Window Help Byonic ™
anouwe (Editor)
Proteins
Peptide List (double click to dock / undock)
Peptides
Prot. Rank | Pos. | Sequence | Glycans | Protein Name | Score | Cleavage | z | Obs. m/z | Calc. m/z | Mods (variable) | PEP 2D | |Log Prob| | Delta Score | PEP 1D | Delta Mod Score | ppm err. | Off-By-X
30 | 58 | R.LGEHNIDVLEGNEQFIN[+0.98402]AAK[+1296.42259]IITHPNF... | Hex(8) | >sp|P00761|TRYP_PIG Trypsin OS=Sus scrofa PE=1 SV=1 | 341.1 | CRagged | 5 | 901.4196 | 901.2086 | N17(Deamidated / 0.984); K20(xgly / 1296.42... | 0.0025 | 2.60 | 85.0 | 0.01 | 5.7 | 11.52 | 1
Prot. Rank | |Log Prob|
1 | 1 535.51 19.71
2 | 2 479.83 17.15
3 | 3 440.70 17.13
4 | 4 353.22 15.95
5 | 5 339.04 14.87
6 | 6 253.63 19.50
7 | 7 232.96 18.95
8 | 8 231.13 15.32
9 | 9 193.76 13.32
10 | 10 168.49 15.40
11 | 11 167.40 18.80
12 | 12 142.60 15.06
13 | 13 141.36 17.15
14 | 14 135.94 13.20
15 | 15 118.95 14.92
16 | 16 86.08 14.13
17 | 17 83.00 16.30
18 | 18 78.19 14.01
19 | 19 71.53 10.89
20 | 20 67.44 14.11
21 | 21 65.74 13.62
22 | 22 63.03 11.90
23 | 23 61.67 13.01
24 | 24 51.72 13.03
25 | 25 45.99 17.52
Protein Coverage (double click to ...
Protein Coverage
Sample name
Protein name
Spectrum (double click to dock / undock)
R.LGEHNIDVLEGNEQFIN[+1]AAK[+1296]IITHPNFNG.N z=5,scan#=sample=1 period=1 cycle=1006 experiment=13,scan time=16.1387
LGEHNIDVLEGNEQFINAAKIITHPNFNG
Intensity
6.00e+02
5.00e+02
4.00e+02
3.00e+02
2.00e+02
1.00e+02
0.00e+00
imm_L
imm_H
imm_F
y6++
b4
b5
b6
b13++
b7
b8
b17++
b9
b13
200
400
600
800
1000
1200
1400
m/z
m/z errors, tol=75.0 ppm
Obs - Calc m/z
6.00e-02
5.00e-02
4.00e-02
3.00e-02
2.00e-02
1.00e-02
0.00e+00
11:21 AM
20/08/2020

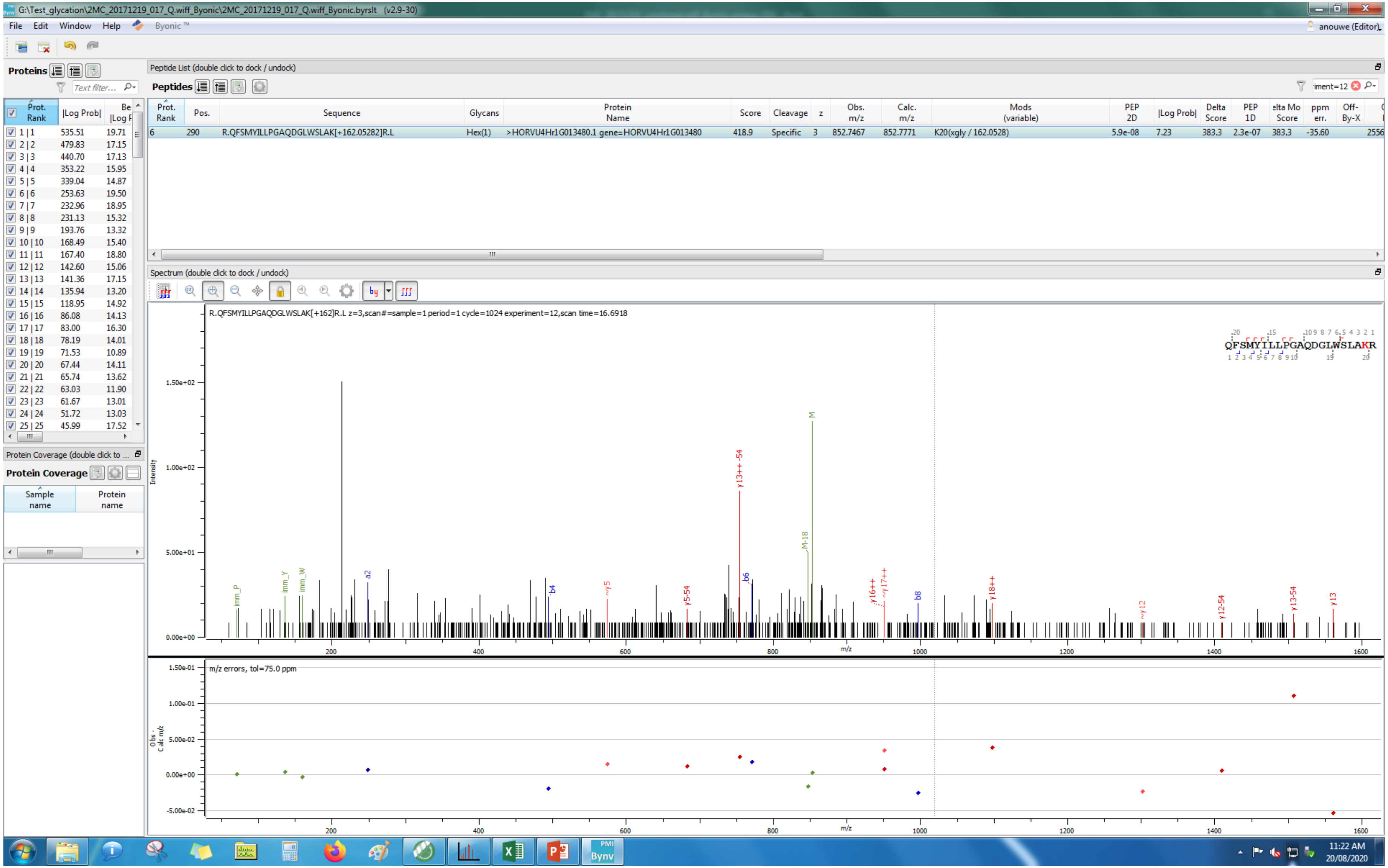
G:\Test_glycation\2MC_20171219_017_Q.wiff_Byonic\2MC_20171219_017_Q.wiff_Byonic.byrslt (v2.9-30)
File Edit Window Help Byonic™
anouwe (Editor)
Proteins
Peptide List (double click to dock / undock)
Peptides
Prot. Rank | Pos. | Sequence | Glycans | Protein Name | Score | Cleavage | z | Obs. m/z | Calc. m/z | Mods (variable) | PEP 2D | |Log Prob| | Delta Score | PEP 1D | Delta Mod Score | ppm err. | Off-By-X
6 | 290 | R.QFSMYILLPGAQDGLWSLAK[+162.05282]R.L | Hex(1) | >HORVU4Hr1G013480.1 gene=HORVU4Hr1G013480 | 418.9 | Specific | 3 | 852.7467 | 852.7771 | K20(xgly / 162.0528) | 5.9e-08 | 7.23 | 383.3 | 2.3e-07 | 383.3 | -35.60 | 2556
Spectrum (double click to dock / undock)
R.QFSMYILLPGAQDGLWSLAK[+162]R.L z=3,scan#=sample=1 period=1 cycle=1024 experiment=12,scan time=16.6918
QFSMYILLPGAQDGLWSLAKR
Intensity
m/z
m/z errors, tol=75.0 ppm
Protein Coverage
Sample name
Protein name
11:22 AM
20/08/2020

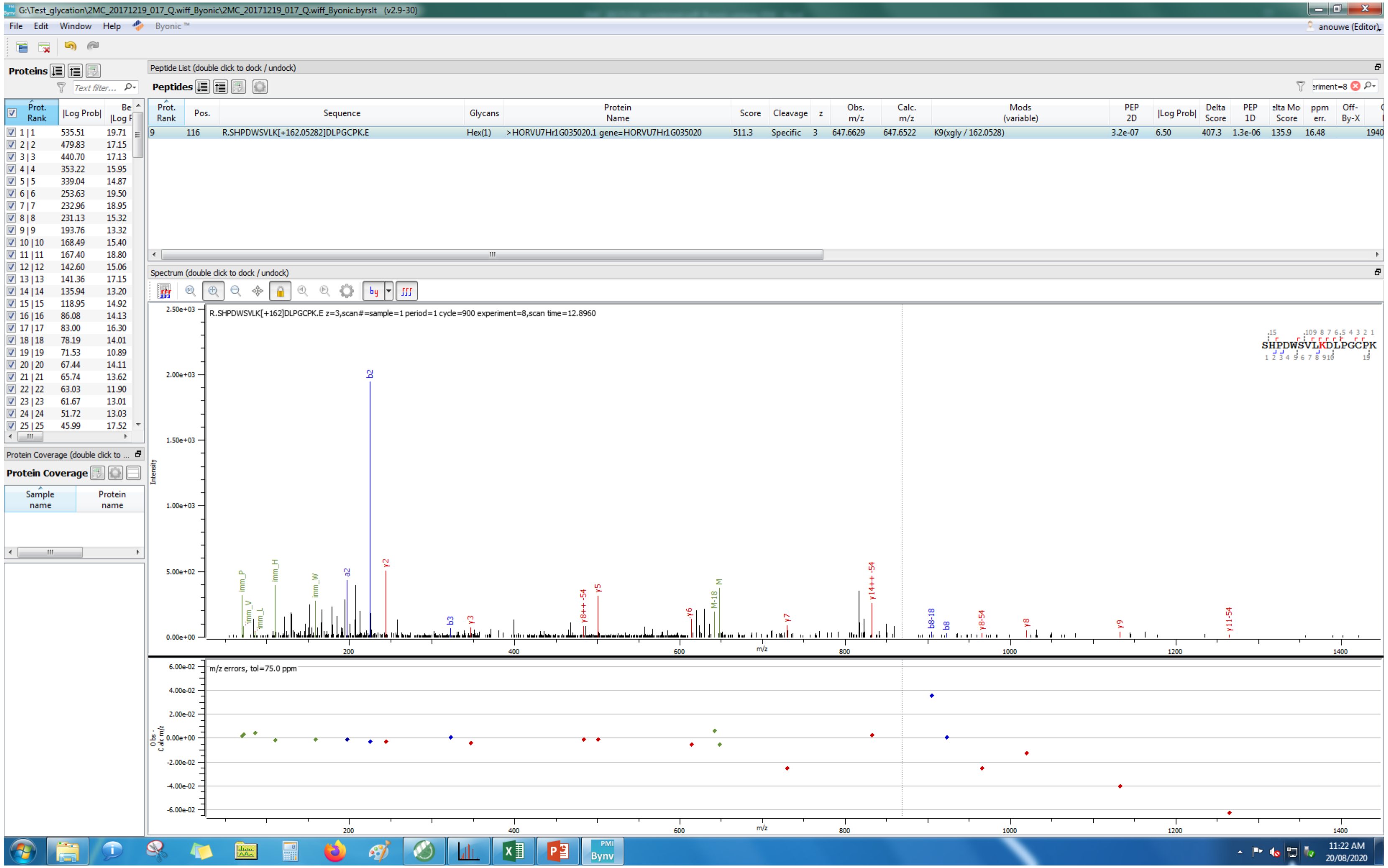
G:\Test_glycation\2MC_20171219_017_Q.wiff_Byonic\2MC_20171219_017_Q.wiff_Byonic.byrslt (v2.9-30)
File Edit Window Help Byonic ™
anouwe (Editor)
Proteins
Peptide List (double click to dock / undock)
Peptides
Prot. Rank | Pos. | Sequence | Glycans | Protein Name | Score | Cleavage | z | Obs. m/z | Calc. m/z | Mods (variable) | PEP 2D | |Log Prob| | Delta Score | PEP 1D | Delta Mod Score | ppm err. | Off-By-X
9 | 116 | R.SHPDWSVLK[+162.05282]DLPGCPK.E | Hex(1) | >HORVU7Hr1G035020.1 gene=HORVU7Hr1G035020 | 511.3 | Specific | 3 | 647.6629 | 647.6522 | K9(xgly / 162.0528) | 3.2e-07 | 6.50 | 407.3 | 1.3e-06 | 135.9 | 16.48 | 1940
Spectrum (double click to dock / undock)
R.SHPDWSVLK[+162]DLPGCPK.E z=3,scan#=sample=1 period=1 cycle=900 experiment=8,scan time=12.8960
SHPDWSVLKDLPGCPK
m/z errors, tol=75.0 ppm
Protein Coverage
Sample name
Protein name
11:22 AM
20/08/2020

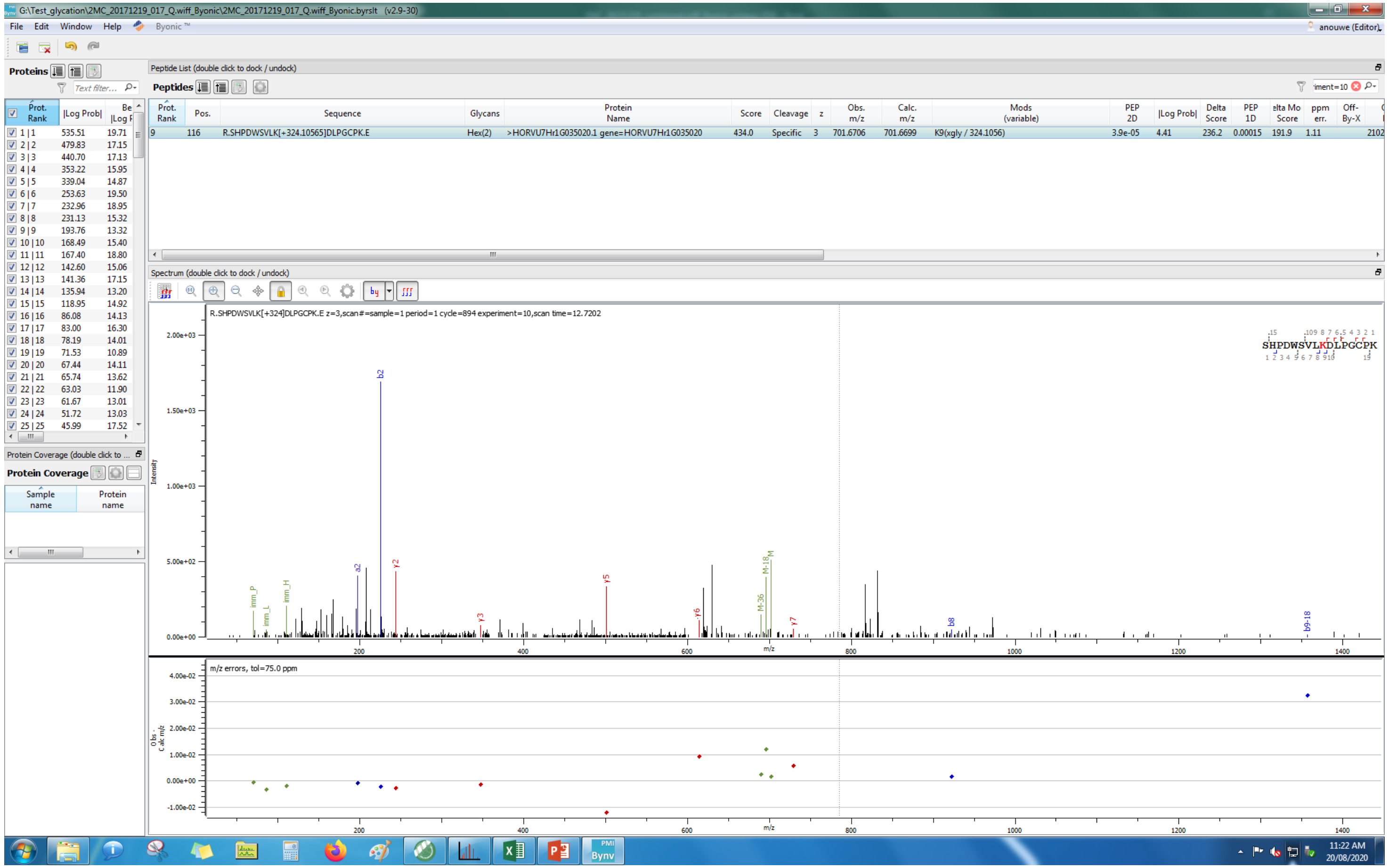

G:\Test_glycation\2MC_20171219_017_Q.wiff_Byonic\2MC_20171219_017_Q.wiff_Byonic.byrslt (v2.9-30)
File Edit Window Help Byonic™
anouwe (Editor)
Proteins
Peptide List (double click to dock / undock)
Peptides
Prot. Rank | Pos. | Sequence | Glycans | Protein Name | Score | Cleavage | z | Obs. m/z | Calc. m/z | Mods (variable) | PEP 2D | |Log Prob| | Delta Score | PEP 1D | Delta Mod Score | ppm err. | Off-By-X
9 | 116 | R.SHPDWSVLK[+324.10565]DLPGCPK.E | Hex(2) | >HORVU7Hr1G035020.1 gene=HORVU7Hr1G035020 | 434.0 | Specific | 3 | 701.6706 | 701.6699 | K9(xgly / 324.1056) | 3.9e-05 | 4.41 | 236.2 | 0.00015 | 191.9 | 1.11 | 2102
Prot. Rank | |Log Prob| | Best |Log P|
1 | 1 535.51 19.71
2 | 2 479.83 17.15
3 | 3 440.70 17.13
4 | 4 353.22 15.95
5 | 5 339.04 14.87
6 | 6 253.63 19.50
7 | 7 232.96 18.95
8 | 8 231.13 15.32
9 | 9 193.76 13.32
10 | 10 168.49 15.40
11 | 11 167.40 18.80
12 | 12 142.60 15.06
13 | 13 141.36 17.15
14 | 14 135.94 13.20
15 | 15 118.95 14.92
16 | 16 86.08 14.13
17 | 17 83.00 16.30
18 | 18 78.19 14.01
19 | 19 71.53 10.89
20 | 20 67.44 14.11
21 | 21 65.74 13.62
22 | 22 63.03 11.90
23 | 23 61.67 13.01
24 | 24 51.72 13.03
25 | 25 45.99 17.52
Protein Coverage
Sample name
Protein name
Spectrum (double click to dock / undock)
R.SHPDWSVLK[+324]DLPGCPK.E z=3,scan#=sample=1 period=1 cycle=894 experiment=10,scan time=12.7202
SHPDWSVLKDLPGCPK
Intensity
b2
a2
y2
y3
y5
y6
y7
M-18
M
M-36
b8
b9-18
imm_P
imm_L
imm_H
m/z
m/z errors, tol=75.0 ppm
Obs - Calc m/z
11:22 AM
20/08/2020

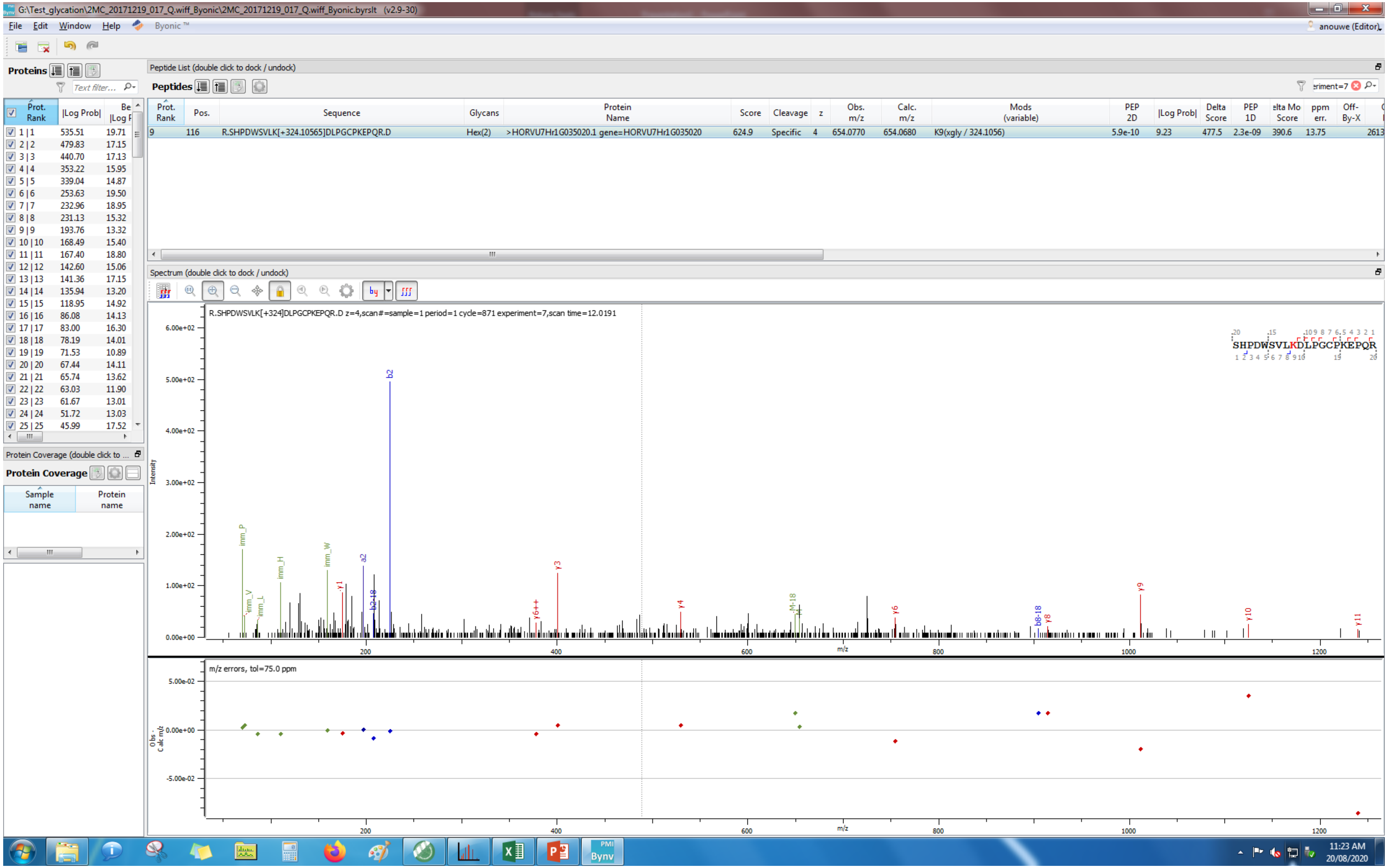
G:\Test_glycation\2MC_20171219_017_Q.wiff_Byonic\2MC_20171219_017_Q.wiff_Byonic.byrslt (v2.9-30)
File Edit Window Help Byonic™
anouwe (Editor)
Proteins
Peptide List (double click to dock / undock)
Peptides
Prot. Rank | Pos. | Sequence | Glycans | Protein Name | Score | Cleavage | z | Obs. m/z | Calc. m/z | Mods (variable) | PEP 2D | |Log Prob| | Delta Score | PEP 1D | Delta Mod Score | ppm err. | Off-By-X
9 | 116 | R.SHPDWSVLK[+324.10565]DLPGCPKEPQR.D | Hex(2) | >HORVU7Hr1G035020.1 gene=HORVU7Hr1G035020 | 624.9 | Specific | 4 | 654.0770 | 654.0680 | K9(xgly / 324.1056) | 5.9e-10 | 9.23 | 477.5 | 2.3e-09 | 390.6 | 13.75 | 2613
Prot. Rank | |Log Prob|
1 | 1 535.51 19.71
2 | 2 479.83 17.15
3 | 3 440.70 17.13
4 | 4 353.22 15.95
5 | 5 339.04 14.87
6 | 6 253.63 19.50
7 | 7 232.96 18.95
8 | 8 231.13 15.32
9 | 9 193.76 13.32
10 | 10 168.49 15.40
11 | 11 167.40 18.80
12 | 12 142.60 15.06
13 | 13 141.36 17.15
14 | 14 135.94 13.20
15 | 15 118.95 14.92
16 | 16 86.08 14.13
17 | 17 83.00 16.30
18 | 18 78.19 14.01
19 | 19 71.53 10.89
20 | 20 67.44 14.11
21 | 21 65.74 13.62
22 | 22 63.03 11.90
23 | 23 61.67 13.01
24 | 24 51.72 13.03
25 | 25 45.99 17.52
Protein Coverage (double click to ...
Protein Coverage
Sample name
Protein name
Spectrum (double click to dock / undock)
R.SHPDWSVLK[+324]DLPGCPKEPQR.D z=4,scan#=sample=1 period=1 cycle=871 experiment=7,scan time=12.0191
SHPDWSVLKDLPGCPKEPQR
Intensity
m/z
m/z errors, tol=75.0 ppm
Obs - Calc m/z
11:23 AM
20/08/2020

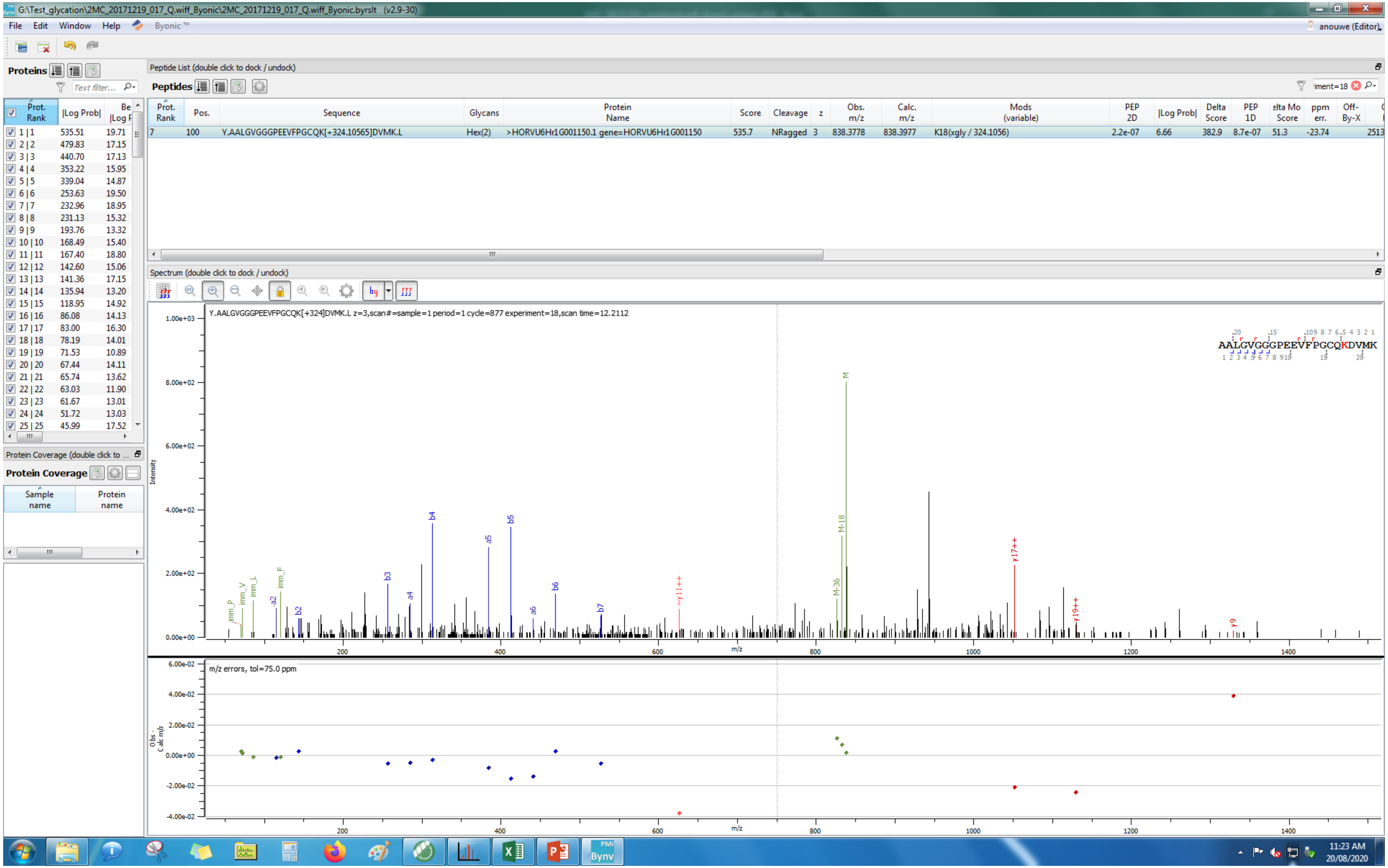
G:\Test_glycation\2MC_20171219_017_Q.wiff_Byonic\2MC_20171219_017_Q.wiff_Byonic.byrslt (v2.9-30)
File Edit Window Help Byonic™
anouwe (Editor)
Proteins
Peptide List (double click to dock / undock)
Peptides
Prot. Rank | Pos. | Sequence | Glycans | Protein Name | Score | Cleavage | z | Obs. m/z | Calc. m/z | Mods (variable) | PEP 2D | |Log Prob| | Delta Score | PEP 1D | Delta Mod Score | ppm err. | Off-By-X
7 | 100 | Y.AALGVGGGPEEVFPGCQK[+324.10565]DVMK.L | Hex(2) | >HORVU6Hr1G001150.1 gene=HORVU6Hr1G001150 | 535.7 | NRagged | 3 | 838.3778 | 838.3977 | K18(xgly / 324.1056) | 2.2e-07 | 6.66 | 382.9 | 8.7e-07 | 51.3 | -23.74 | 2513
Prot. Rank | |Log Prob| | Best |Log P|
1 | 1 535.51 19.71
2 | 2 479.83 17.15
3 | 3 440.70 17.13
4 | 4 353.22 15.95
5 | 5 339.04 14.87
6 | 6 253.63 19.50
7 | 7 232.96 18.95
8 | 8 231.13 15.32
9 | 9 193.76 13.32
10 | 10 168.49 15.40
11 | 11 167.40 18.80
12 | 12 142.60 15.06
13 | 13 141.36 17.15
14 | 14 135.94 13.20
15 | 15 118.95 14.92
16 | 16 86.08 14.13
17 | 17 83.00 16.30
18 | 18 78.19 14.01
19 | 19 71.53 10.89
20 | 20 67.44 14.11
21 | 21 65.74 13.62
22 | 22 63.03 11.90
23 | 23 61.67 13.01
24 | 24 51.72 13.03
25 | 25 45.99 17.52
Protein Coverage
Sample name
Protein name
Spectrum (double click to dock / undock)
Y.AALGVGGGPEEVFPGCQK[+324]DVMK.L z=3,scan#=sample=1 period=1 cycle=877 experiment=18,scan time=12.2112
AALGVGGGPEEVFPGCQKDVMK
Intensity
m/z
M
M-18
M-36
b2 b3 b4 b5 b6 b7
a2 a4 a5 a6
imm_P imm_V imm_L imm_F
y11++ y17++ y19++ y9
m/z errors, tol=75.0 ppm
Obs - Calc m/z
11:23 AM
20/08/2020

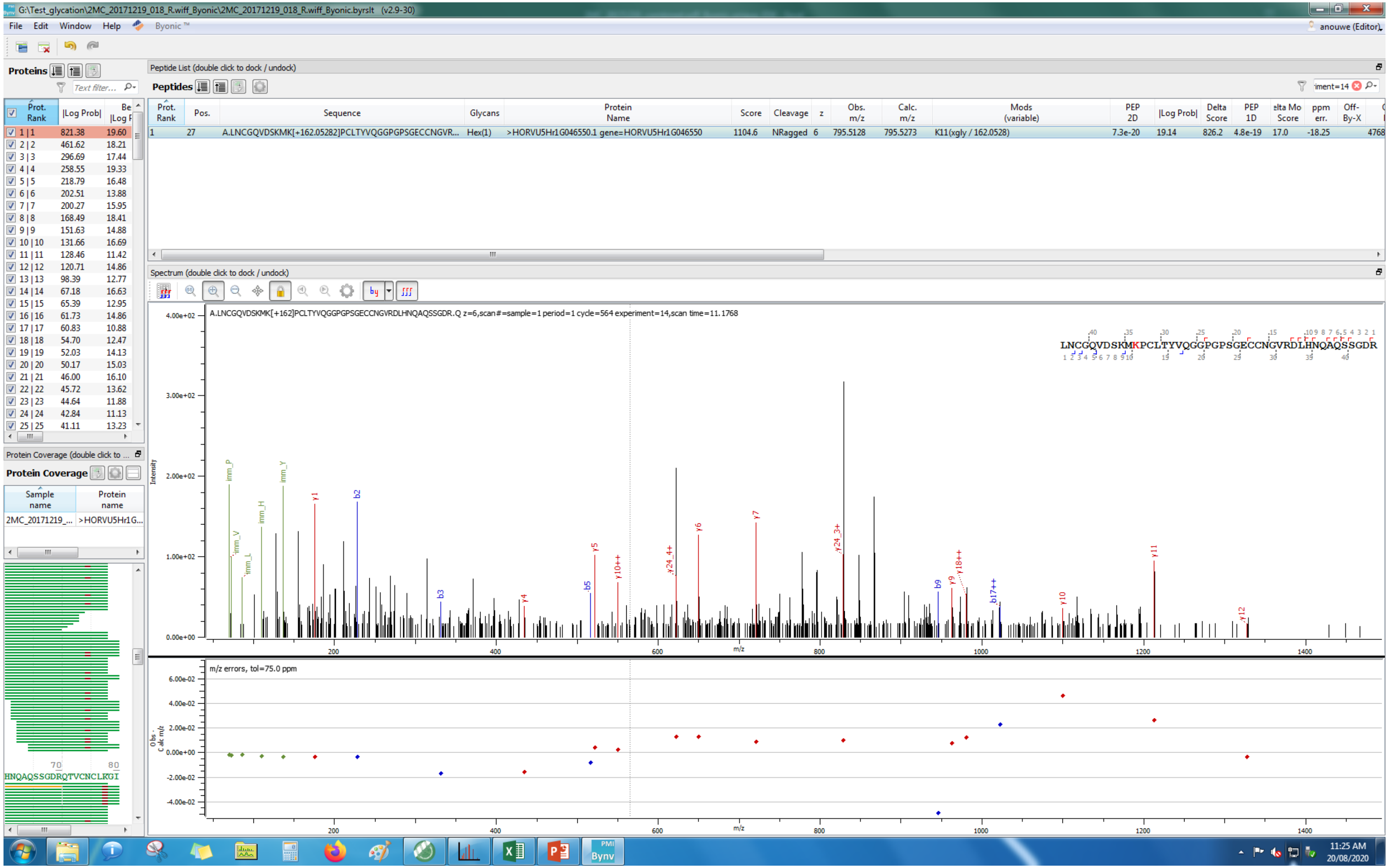
G:\Test_glycation\2MC_20171219_018_R.wiff_Byonic\2MC_20171219_018_R.wiff_Byonic.byrslt (v2.9-30)
File Edit Window Help Byonic™
anouwe (Editor)
Proteins
Peptide List (double click to dock / undock)
Peptides
Prot. Rank | Pos. | Sequence | Glycans | Protein Name | Score | Cleavage | z | Obs. m/z | Calc. m/z | Mods (variable) | PEP 2D | |Log Prob| | Delta Score | PEP 1D | Delta Mod Score | ppm err. | Off-By-X
1 | 27 | A.LNCGQVDSKMK[+162.05282]PCLTYVQGGPGPSGECCNGVR... | Hex(1) | >HORVU5Hr1G046550.1 gene=HORVU5Hr1G046550 | 1104.6 | NRagged | 6 | 795.5128 | 795.5273 | K11(xgly / 162.0528) | 7.3e-20 | 19.14 | 826.2 | 4.8e-19 | 17.0 | -18.25 | 4768
Spectrum (double click to dock / undock)
A.LNCGQVDSKMK[+162]PCLTYVQGGPGPSGECCNGVRDLHNQAQSSGDR.Q z=6,scan#=sample=1 period=1 cycle=564 experiment=14,scan time=11.1768
LNCGQVDSKMKPCLTYVQGGPGPSGECCNGVRDLHNQAQSSGDR
m/z errors, tol=75.0 ppm
Protein Coverage
Sample name | Protein name
2MC_20171219_... | >HORVU5Hr1G...
HNQAQSSGDRQTVCNCLKGI
11:25 AM
20/08/2020

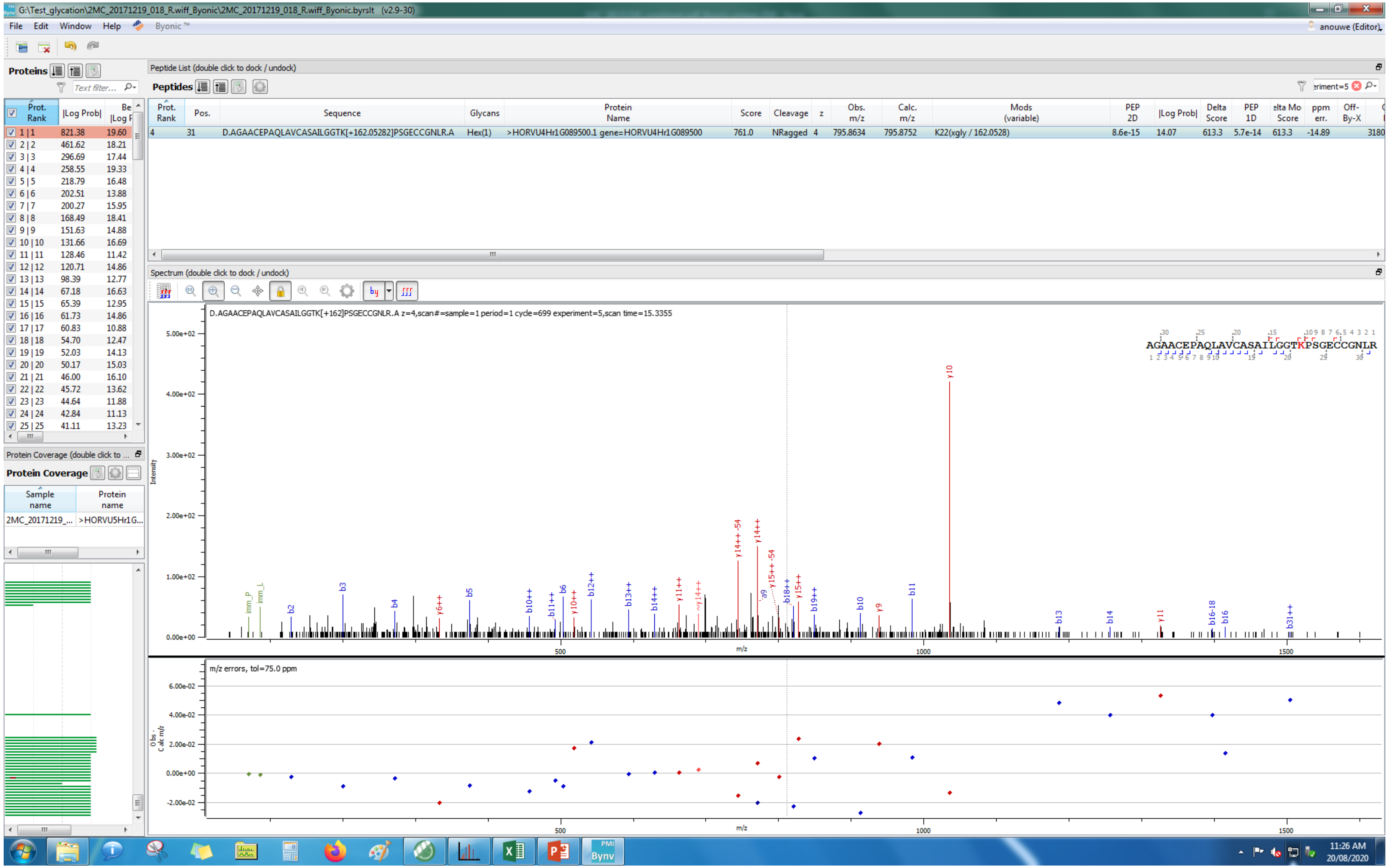
G:\Test_glycation\2MC_20171219_018_R.wiff_Byonic\2MC_20171219_018_R.wiff_Byonic.byrslt (v2.9-30)
File Edit Window Help Byonic
Proteins
Peptide List (double click to dock / undock)
Peptides
Prot. Rank | Pos. | Sequence | Glycans | Protein Name | Score | Cleavage | z | Obs. m/z | Calc. m/z | Mods (variable) | PEP 2D | |Log Prob| | Delta Score | PEP 1D | Delta Mod Score | ppm err. | Off-By-X
4 | 31 | D.AGAACEPAQLAVCASAILGGTK[+162.05282]PSGECCGNLR.A | Hex(1) | >HORVU4Hr1G089500.1 gene=HORVU4Hr1G089500 | 761.0 | NRagged | 4 | 795.8634 | 795.8752 | K22(xgly / 162.0528) | 8.6e-15 | 14.07 | 613.3 | 5.7e-14 | 613.3 | -14.89
Spectrum (double click to dock / undock)
D.AGAACEPAQLAVCASAILGGTK[+162]PSGECCGNLR.A z=4,scan#=sample=1 period=1 cycle=699 experiment=5,scan time=15.3355
AGAACEPAQLAVCASAILGGTKPSGECCGNLR
Intensity
m/z
m/z errors, tol=75.0 ppm
Obs - Calc m/z
Protein Coverage
Sample name | Protein name
2MC_20171219_... | >HORVU5Hr1G...
11:26 AM
20/08/2020

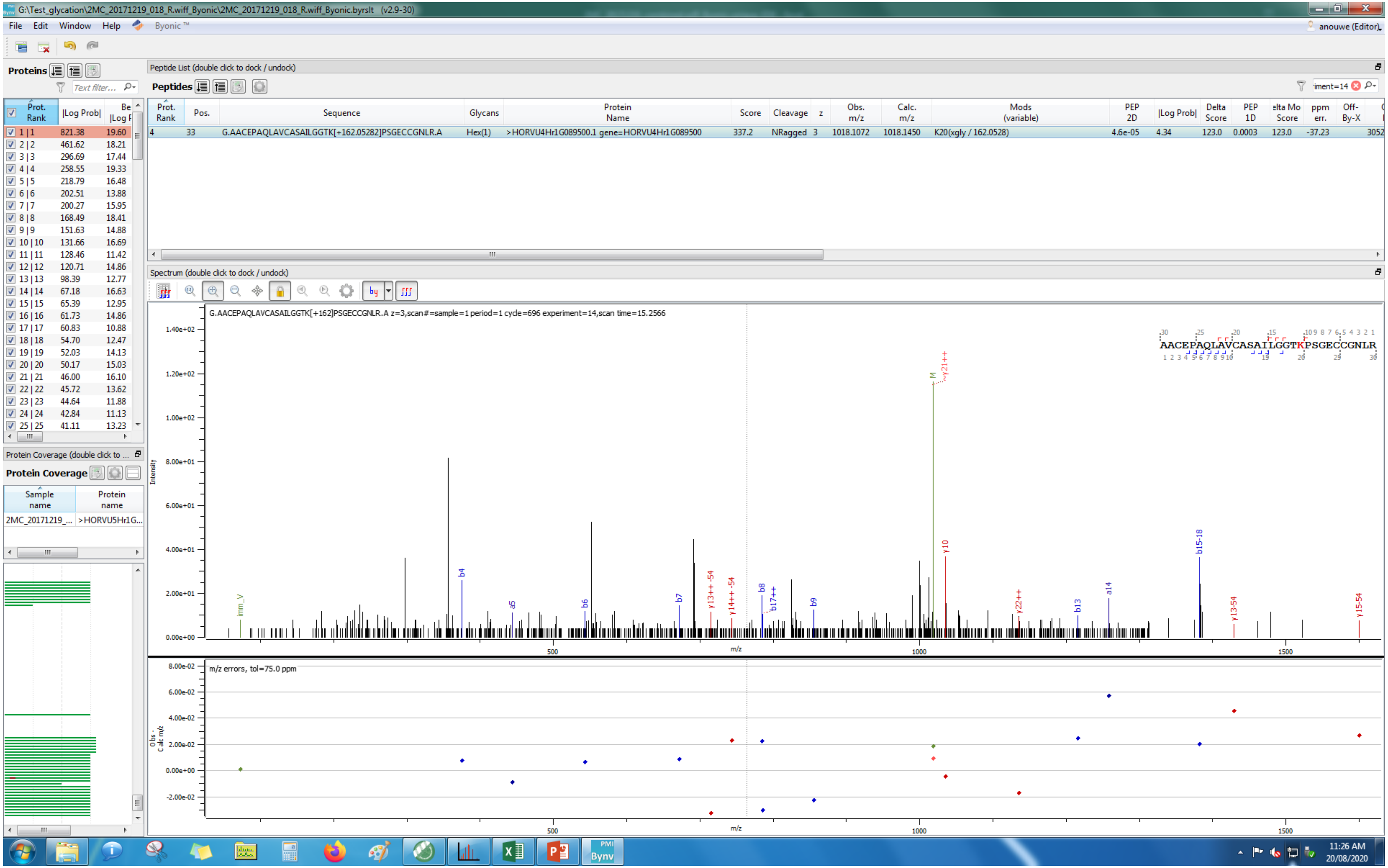
G:\Test_glycation\2MC_20171219_018_R.wiff_Byonic\2MC_20171219_018_R.wiff_Byonic.byrslt (v2.9-30)
File Edit Window Help Byonic™
Proteins
Peptide List (double click to dock / undock)
Peptides
Prot. Rank | Pos. | Sequence | Glycans | Protein Name | Score | Cleavage | z | Obs. m/z | Calc. m/z | Mods (variable) | PEP 2D | |Log Prob| | Delta Score | PEP 1D | Delta Mod Score | ppm err. | Off-By-X
4 | 33 | G.AACEPAQLAVCASAILGGTK[+162.05282]PSGECCGNLR.A | Hex(1) | >HORVU4Hr1G089500.1 gene=HORVU4Hr1G089500 | 337.2 | NRagged | 3 | 1018.1072 | 1018.1450 | K20(xgly / 162.0528) | 4.6e-05 | 4.34 | 123.0 | 0.0003 | 123.0 | -37.23
Spectrum (double click to dock / undock)
G.AACEPAQLAVCASAILGGTK[+162]PSGECCGNLR.A z=3,scan#=sample=1 period=1 cycle=696 experiment=14,scan time=15.2566
AACEPAQLAVCASAILGGTKPSGECCGNLR
Intensity
m/z
m/z errors, tol=75.0 ppm
Protein Coverage
Sample name | Protein name
2MC_20171219_... | >HORVU5Hr1G...
anouwe (Editor)
11:26 AM
20/08/2020

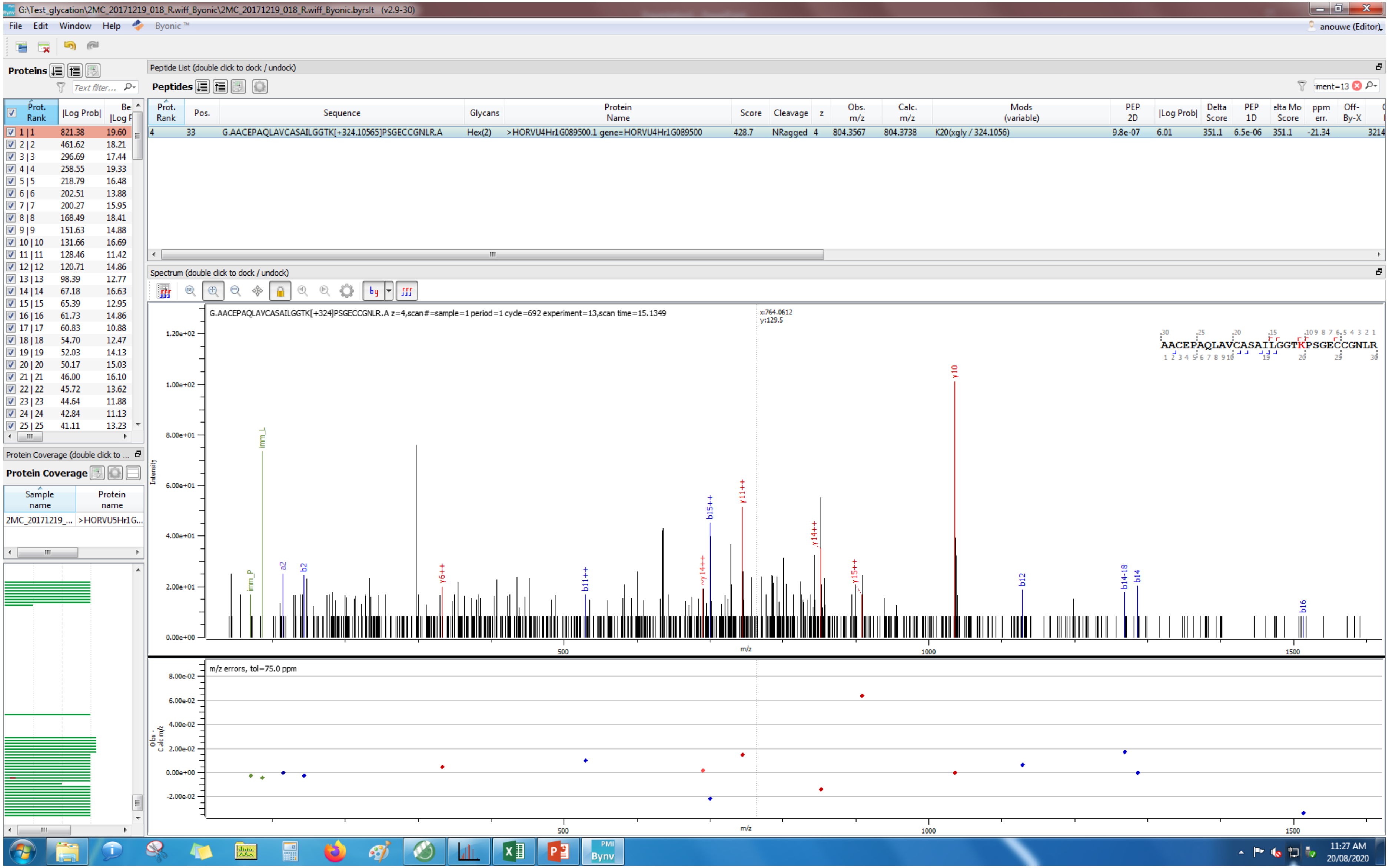
G:\Test_glycation\2MC_20171219_018_R.wiff_Byonic\2MC_20171219_018_R.wiff_Byonic.byrslt (v2.9-30)
File Edit Window Help Byonic ™
anouwe (Editor)
Proteins
Peptide List (double click to dock / undock)
Peptides
Prot. Rank | Pos. | Sequence | Glycans | Protein Name | Score | Cleavage | z | Obs. m/z | Calc. m/z | Mods (variable) | PEP 2D | |Log Prob| | Delta Score | PEP 1D | elta Mo Score | ppm err. | Off-By-X
4 | 33 | G.AACEPAQLAVCASAILGGTK[+324.10565]PSGECCGNLR.A | Hex(2) | >HORVU4Hr1G089500.1 gene=HORVU4Hr1G089500 | 428.7 | NRagged | 4 | 804.3567 | 804.3738 | K20(xgly / 324.1056) | 9.8e-07 | 6.01 | 351.1 | 6.5e-06 | 351.1 | -21.34 | 3214
Prot. Rank | |Log Prob| | Be |Log P
1 | 1 821.38 19.60
2 | 2 461.62 18.21
3 | 3 296.69 17.44
4 | 4 258.55 19.33
5 | 5 218.79 16.48
6 | 6 202.51 13.88
7 | 7 200.27 15.95
8 | 8 168.49 18.41
9 | 9 151.63 14.88
10 | 10 131.66 16.69
11 | 11 128.46 11.42
12 | 12 120.71 14.86
13 | 13 98.39 12.77
14 | 14 67.18 16.63
15 | 15 65.39 12.95
16 | 16 61.73 14.86
17 | 17 60.83 10.88
18 | 18 54.70 12.47
19 | 19 52.03 14.13
20 | 20 50.17 15.03
21 | 21 46.00 16.10
22 | 22 45.72 13.62
23 | 23 44.64 11.88
24 | 24 42.84 11.13
25 | 25 41.11 13.23
Protein Coverage (double click to ...
Protein Coverage
Sample name | Protein name
2MC_20171219_... | >HORVU5Hr1G...
Spectrum (double click to dock / undock)
G.AACEPAQLAVCASAILGGTK[+324]PSGECCGNLR.A z=4,scan#=sample=1 period=1 cycle=692 experiment=13,scan time=15.1349
x:764.0612
y:129.5
AACEPAQLAVCASAILGGTKPSGECCGNLR
Intensity
m/z
m/z errors, tol=75.0 ppm
Obs - Calc m/z
11:27 AM
20/08/2020

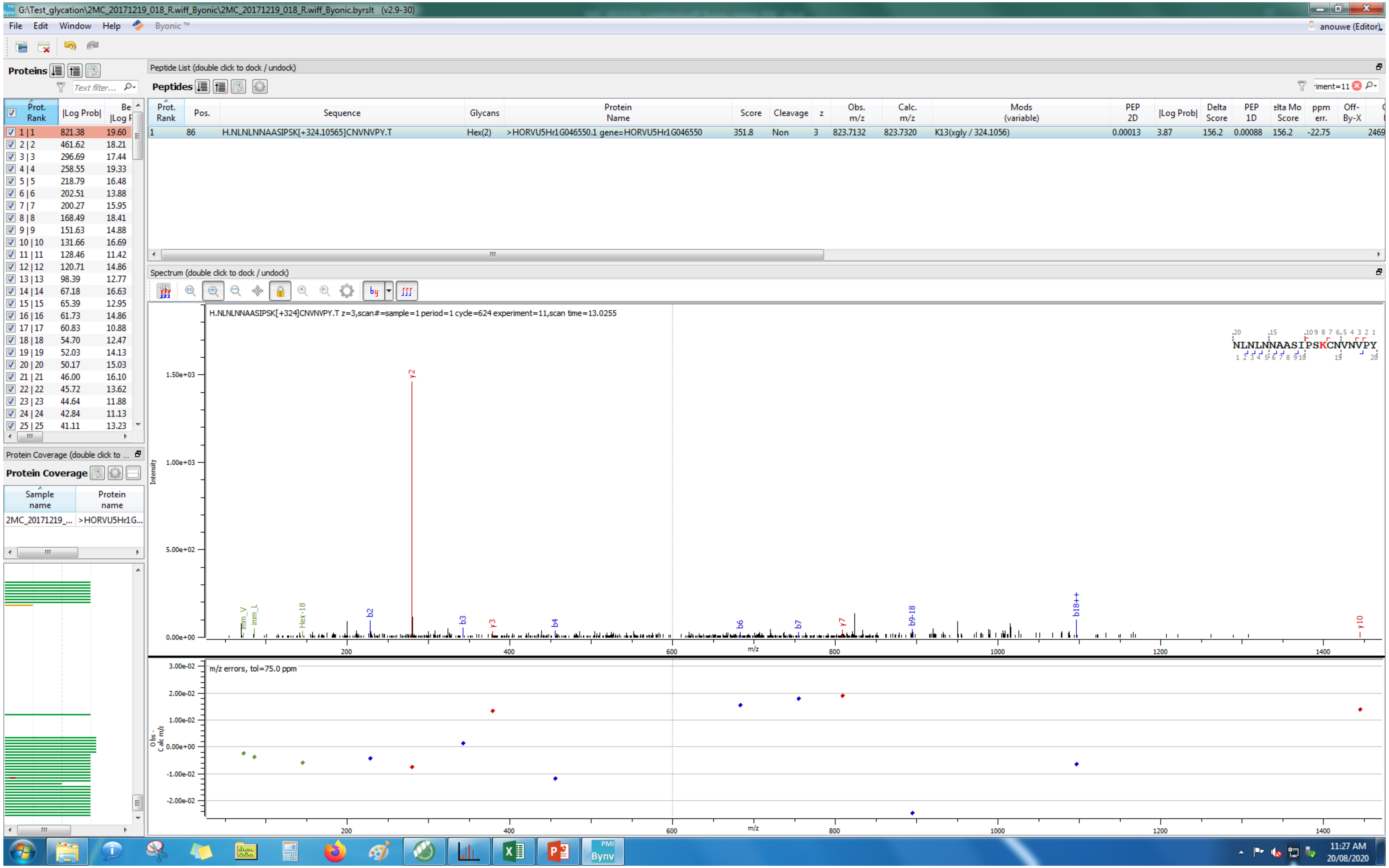
G:\Test_glycation\2MC_20171219_018_R.wiff_Byonic\2MC_20171219_018_R.wiff_Byonic.byrslt (v2.9-30)
File Edit Window Help Byonic ™
Proteins
Peptide List (double click to dock / undock)
Peptides
Prot. Rank | Pos. | Sequence | Glycans | Protein Name | Score | Cleavage | z | Obs. m/z | Calc. m/z | Mods (variable) | PEP 2D | |Log Prob| | Delta Score | PEP 1D | Delta Mod Score | ppm err. | Off-By-X
1 | 86 | H.NLNLNNAASIPSK[+324.10565]CNVNVPY.T | Hex(2) | >HORVU5Hr1G046550.1 gene=HORVU5Hr1G046550 | 351.8 | Non | 3 | 823.7132 | 823.7320 | K13(xgly / 324.1056) | 0.00013 | 3.87 | 156.2 | 0.00088 | 156.2 | -22.75 | 2469
Spectrum (double click to dock / undock)
H.NLNLNNAASIPSK[+324]CNVNVPY.T z=3,scan#=sample=1 period=1 cycle=624 experiment=11,scan time=13.0255
NLNLNNAASIPSKCNVNVPY
Intensity
m/z
m/z errors, tol=75.0 ppm
Obs - Calc m/z
Protein Coverage
Sample name | Protein name
2MC_20171219_... | >HORVU5Hr1G...
11:27 AM
20/08/2020

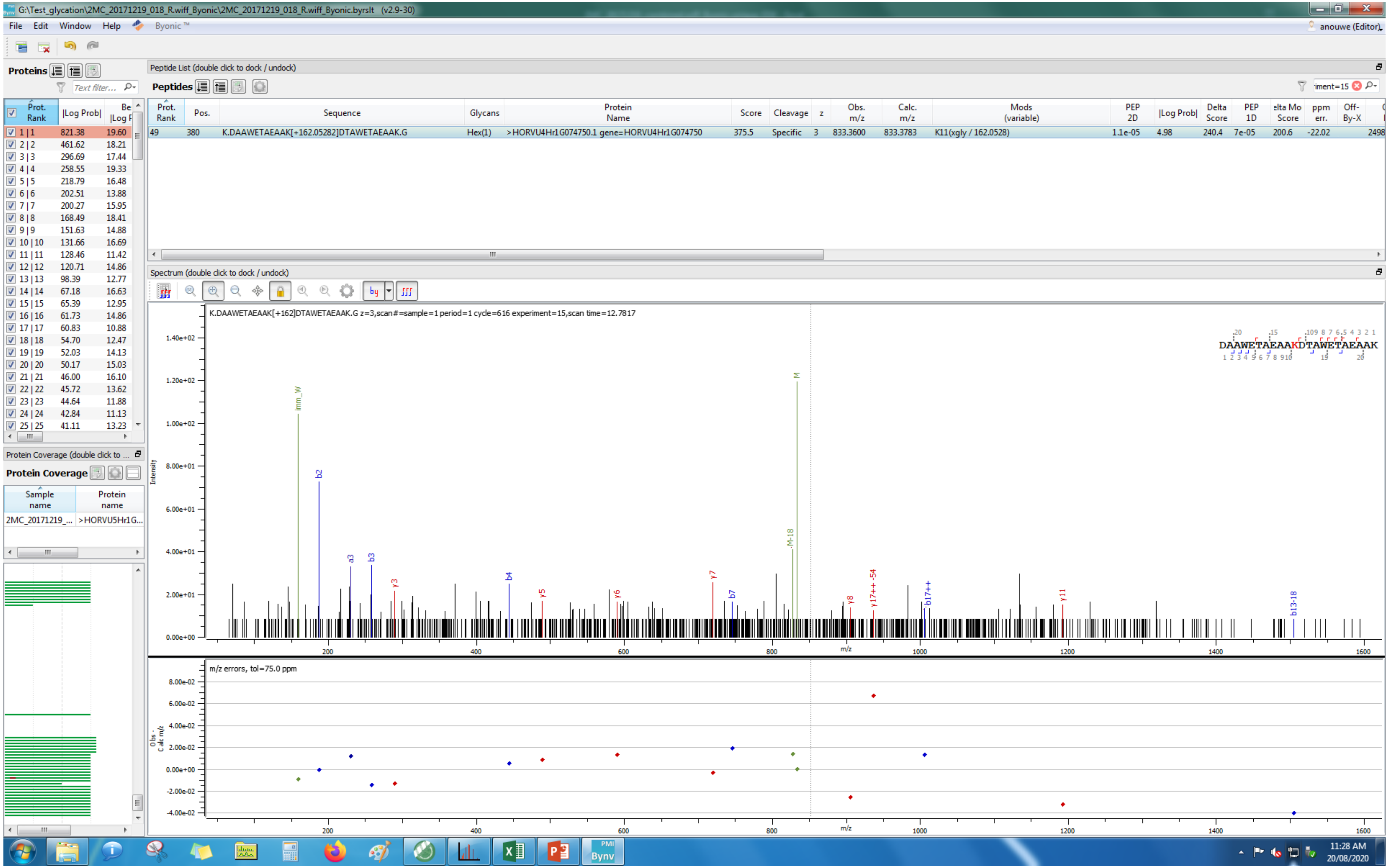

G:\Test_glycation\2MC_20171219_018_R.wiff_Byonic\2MC_20171219_018_R.wiff_Byonic.byrslt (v2.9-30)
File Edit Window Help Byonic™
anouwe (Editor)
Proteins
Peptide List (double click to dock / undock)
Peptides
Prot. Rank | Pos. | Sequence | Glycans | Protein Name | Score | Cleavage | z | Obs. m/z | Calc. m/z | Mods (variable) | PEP 2D | |Log Prob| | Delta Score | PEP 1D | Delta Mod Score | ppm err. | Off-By-X
49 | 380 | K.DAAWETAEAAK[+162.05282]DTAWETAEAAK.G | Hex(1) | >HORVU4Hr1G074750.1 gene=HORVU4Hr1G074750 | 375.5 | Specific | 3 | 833.3600 | 833.3783 | K11(xgly / 162.0528) | 1.1e-05 | 4.98 | 240.4 | 7e-05 | 200.6 | -22.02 | 2498
Spectrum (double click to dock / undock)
K.DAAWETAEAAK[+162]DTAWETAEAAK.G z=3,scan#=sample=1 period=1 cycle=616 experiment=15,scan time=12.7817
DAAWETAEAAKDTAWETAEAAK
Intensity
m/z
m/z errors, tol=75.0 ppm
Protein Coverage (double click to ...
Protein Coverage
Sample name | Protein name
2MC_20171219_... | >HORVU5Hr1G...
11:28 AM
20/08/2020

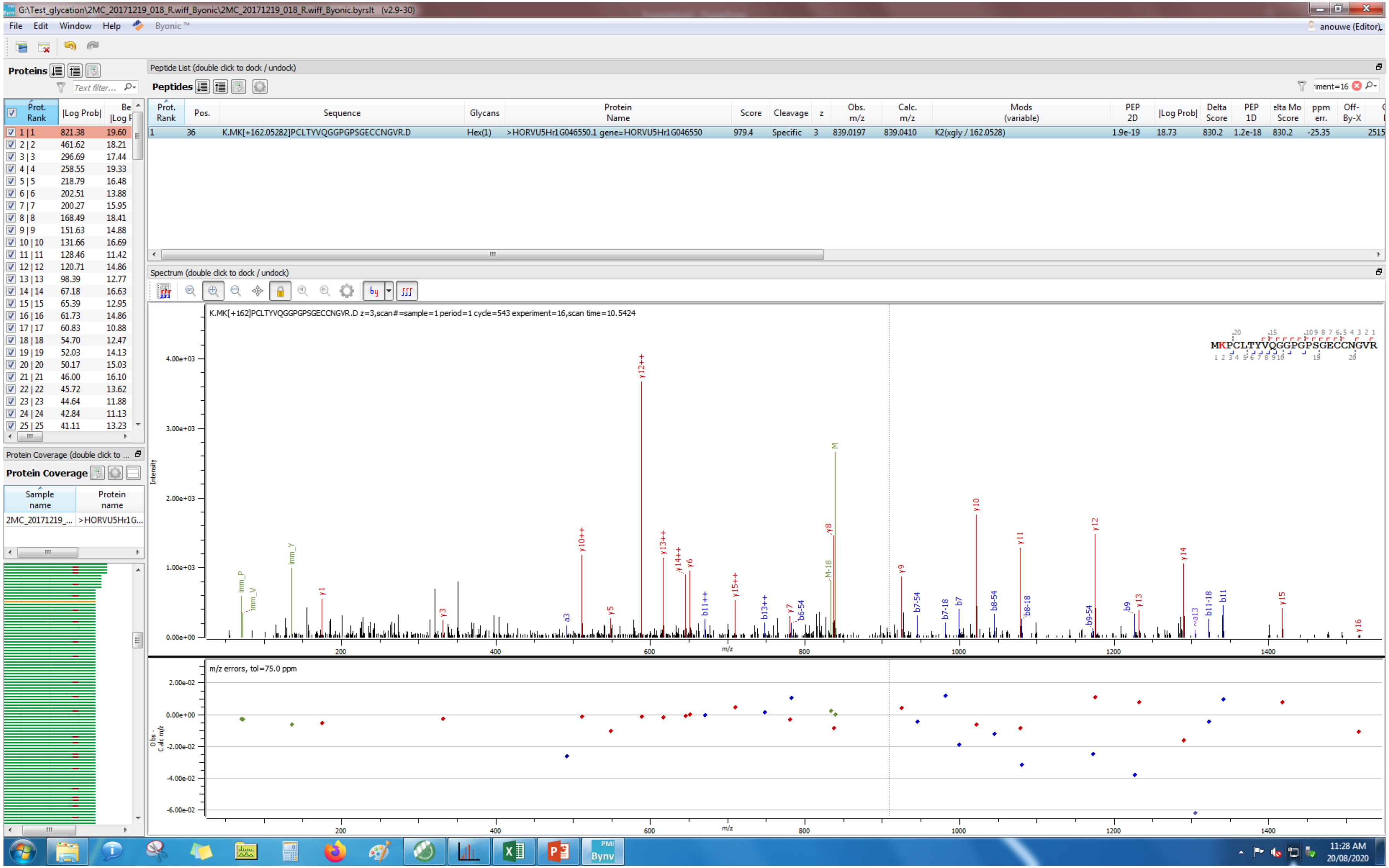
G:\Test_glycation\2MC_20171219_018_R.wiff_Byonic\2MC_20171219_018_R.wiff_Byonic.byrslt (v2.9-30)
File Edit Window Help Byonic™
Proteins
Peptide List (double click to dock / undock)
Peptides
K.MK[+162.05282]PCLTYVQGGPGPSGECCNGVR.D
Hex(1)
>HORVU5Hr1G046550.1 gene=HORVU5Hr1G046550
979.4
Specific
839.0197
839.0410
K2(xgly / 162.0528)
Spectrum (double click to dock / undock)
K.MK[+162]PCLTYVQGGPGPSGECCNGVR.D z=3,scan#=sample=1 period=1 cycle=543 experiment=16,scan time=10.5424
MKPCLTYVQGGPGPSGECCNGVR
m/z errors, tol=75.0 ppm
Protein Coverage
2MC_20171219_...
>HORVU5Hr1G...
11:28 AM
20/08/2020

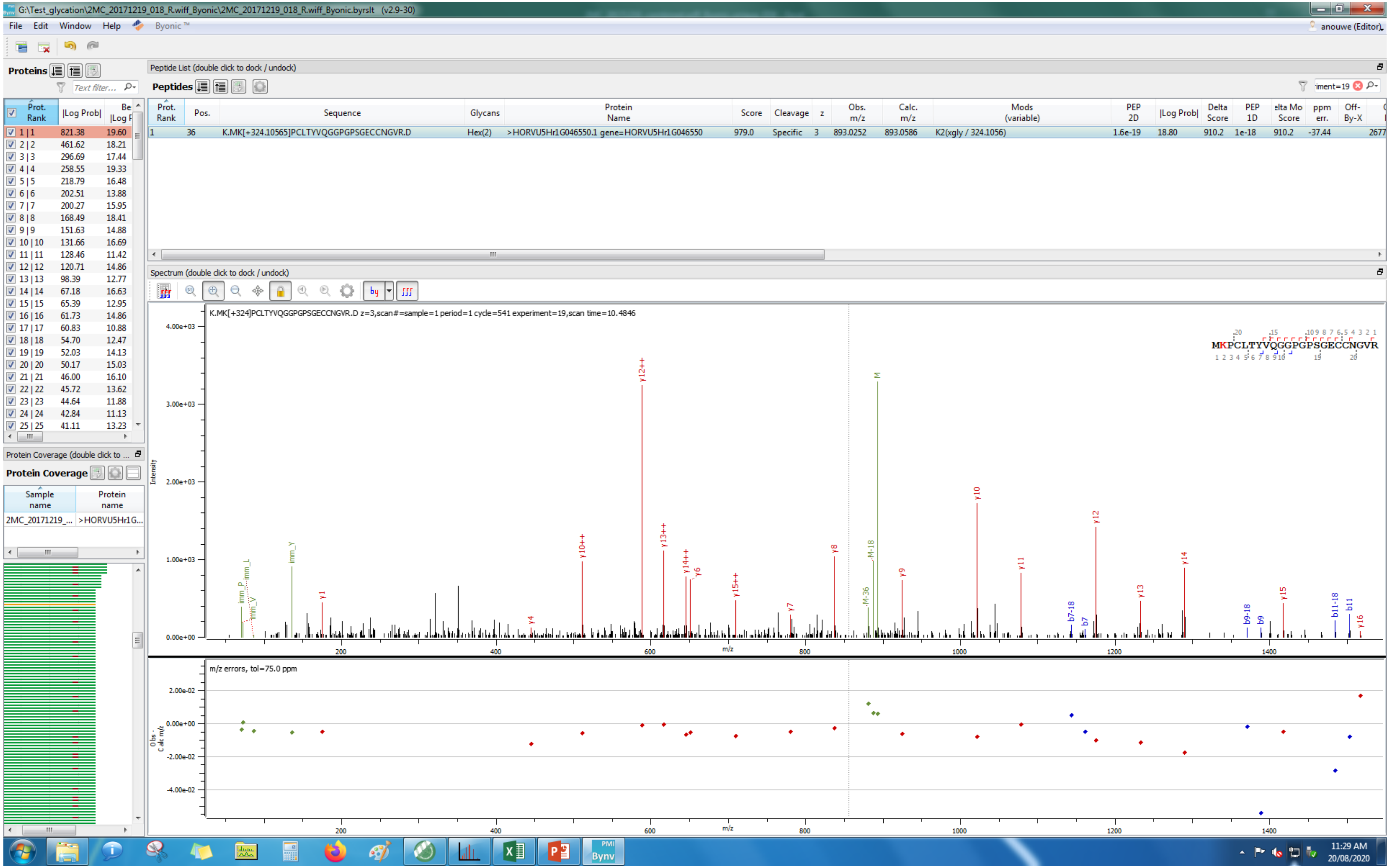
G:\Test_glycation\2MC_20171219_018_R.wiff_Byonic\2MC_20171219_018_R.wiff_Byonic.byrslt (v2.9-30)
File Edit Window Help Byonic ™
Proteins
Peptide List (double click to dock / undock)
Peptides
Prot. Rank | Pos. | Sequence | Glycans | Protein Name | Score | Cleavage | z | Obs. m/z | Calc. m/z | Mods (variable) | PEP 2D | |Log Prob| | Delta Score | PEP 1D | Delta Mod Score | ppm err. | Off-By-X
1 | 36 | K.MK[+324.10565]PCLTYVQGGPGPSGECCNGVR.D | Hex(2) | >HORVU5Hr1G046550.1 gene=HORVU5Hr1G046550 | 979.0 | Specific | 3 | 893.0252 | 893.0586 | K2(xgly / 324.1056) | 1.6e-19 | 18.80 | 910.2 | 1e-18 | 910.2 | -37.44
Spectrum (double click to dock / undock)
K.MK[+324]PCLTYVQGGPGPSGECCNGVR.D z=3,scan#=sample=1 period=1 cycle=541 experiment=19,scan time=10.4846
MKPCLTYVQGGPGPSGECCNGVR
Intensity
m/z
m/z errors, tol=75.0 ppm
Obs - Calc m/z
Protein Coverage
Sample name | Protein name
2MC_20171219_... | >HORVU5Hr1G...
11:29 AM
20/08/2020

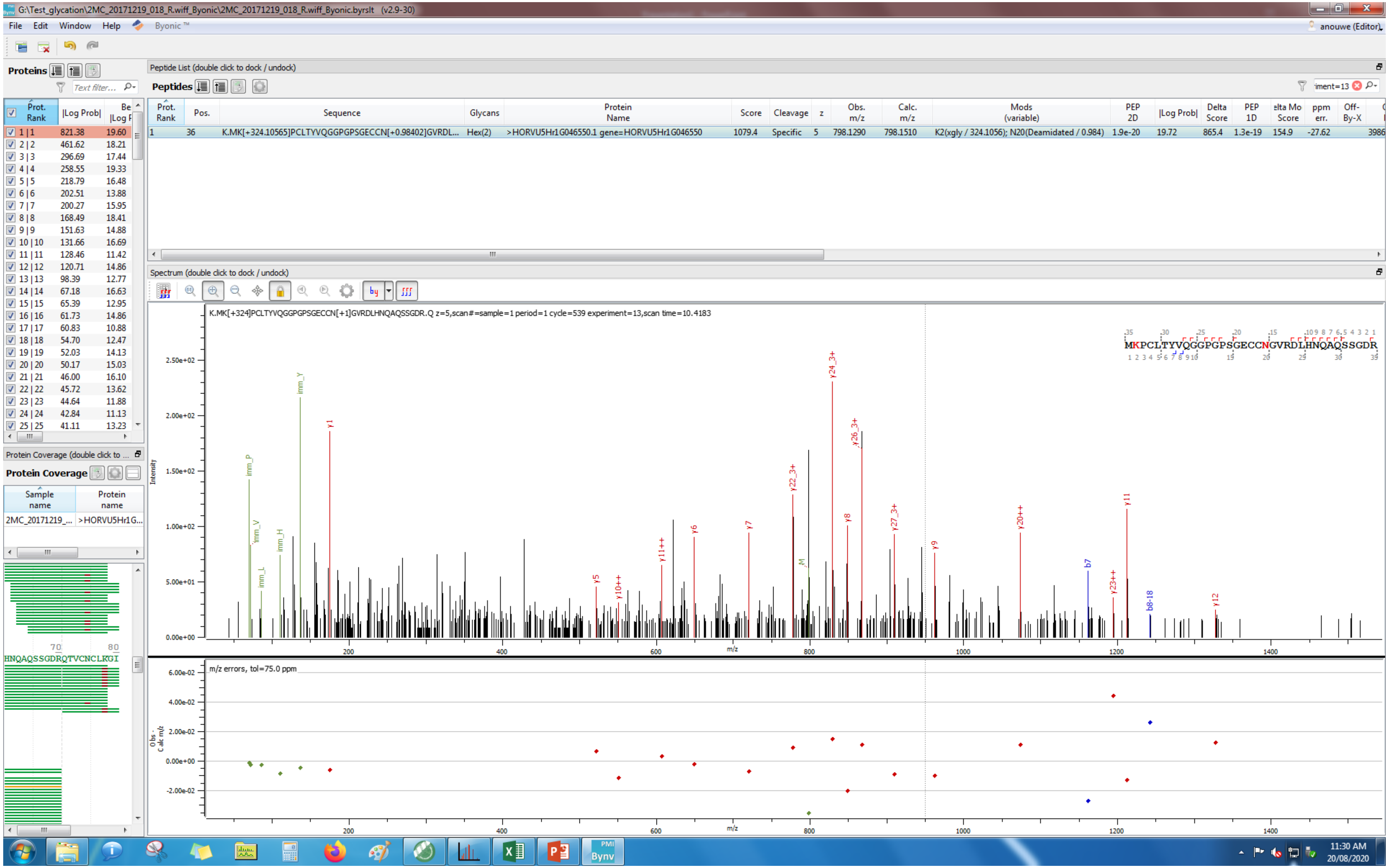
G:\Test_glycation\2MC_20171219_018_R.wiff_Byonic\2MC_20171219_018_R.wiff_Byonic.byrslt (v2.9-30)
File Edit Window Help Byonic™
Proteins
Peptide List (double click to dock / undock)
Peptides
1 36 K.MK[+324.10565]PCLTYVQGGPGPSGECCN[+0.98402]GVRDL... Hex(2) >HORVU5Hr1G046550.1 gene=HORVU5Hr1G046550 1079.4 Specific 5 798.1290 798.1510 K2(xgly / 324.1056); N20(Deamidated / 0.984) 1.9e-20 19.72 865.4 1.3e-19 154.9 -27.62
Spectrum (double click to dock / undock)
K.MK[+324]PCLTYVQGGPGPSGECCN[+1]GVRDLHNQAQSSGDR.Q z=5,scan#=sample=1 period=1 cycle=539 experiment=13,scan time=10.4183
MKPCLTYVQGGPGPSGECCNGVRDLHNQAQSSGDR
m/z errors, tol=75.0 ppm
Protein Coverage
HNQAQSSGDRQTVCNCLKGI
11:30 AM
20/08/2020

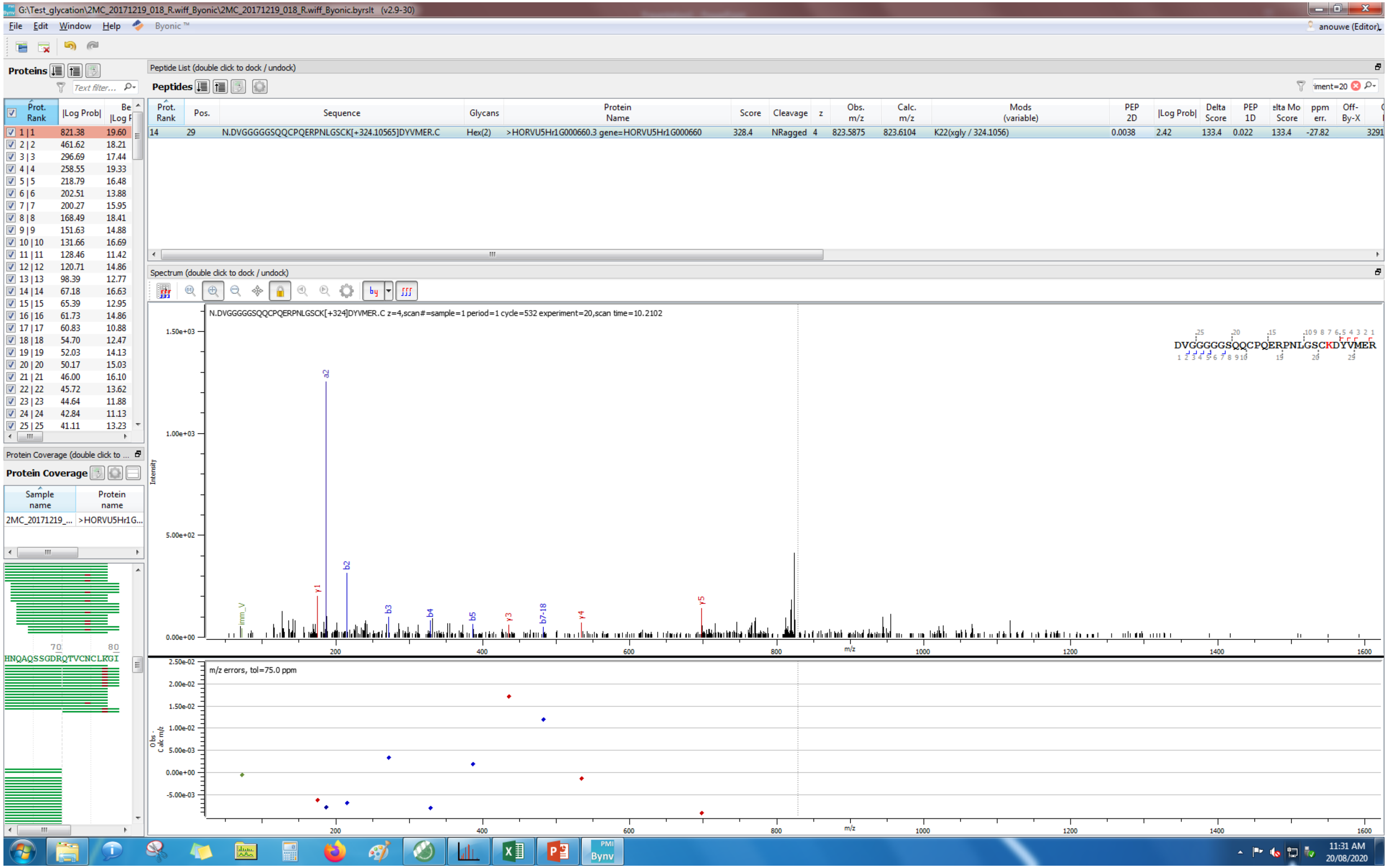
G:\Test_glycation\2MC_20171219_018_R.wiff_Byonic\2MC_20171219_018_R.wiff_Byonic.byrslt (v2.9-30)
File Edit Window Help Byonic ™
anouwe (Editor)
Proteins
Peptide List (double click to dock / undock)
Peptides
Prot. Rank Pos. Sequence Glycans Protein Name Score Cleavage z Obs. m/z Calc. m/z Mods (variable) PEP 2D |Log Prob| Delta Score PEP 1D Delta Mod Score ppm err. Off-By-X
14 29 N.DVGGGGGSQQCPQERPNLGSCK[+324.10565]DYVMER.C Hex(2) >HORVU5Hr1G000660.3 gene=HORVU5Hr1G000660 328.4 NRagged 4 823.5875 823.6104 K22(xgly / 324.1056) 0.0038 2.42 133.4 0.022 133.4 -27.82
Spectrum (double click to dock / undock)
N.DVGGGGGSQQCPQERPNLGSCK[+324]DYVMER.C z=4,scan#=sample=1 period=1 cycle=532 experiment=20,scan time=10.2102
DVGGGGGSQQCPQERPNLGSCKDYVMER
Intensity
m/z
m/z errors, tol=75.0 ppm
Protein Coverage
Sample name
Protein name
2MC_20171219_...
>HORVU5Hr1G...
HNQAQSSGDRQTVCNCLKGI
11:31 AM
20/08/2020

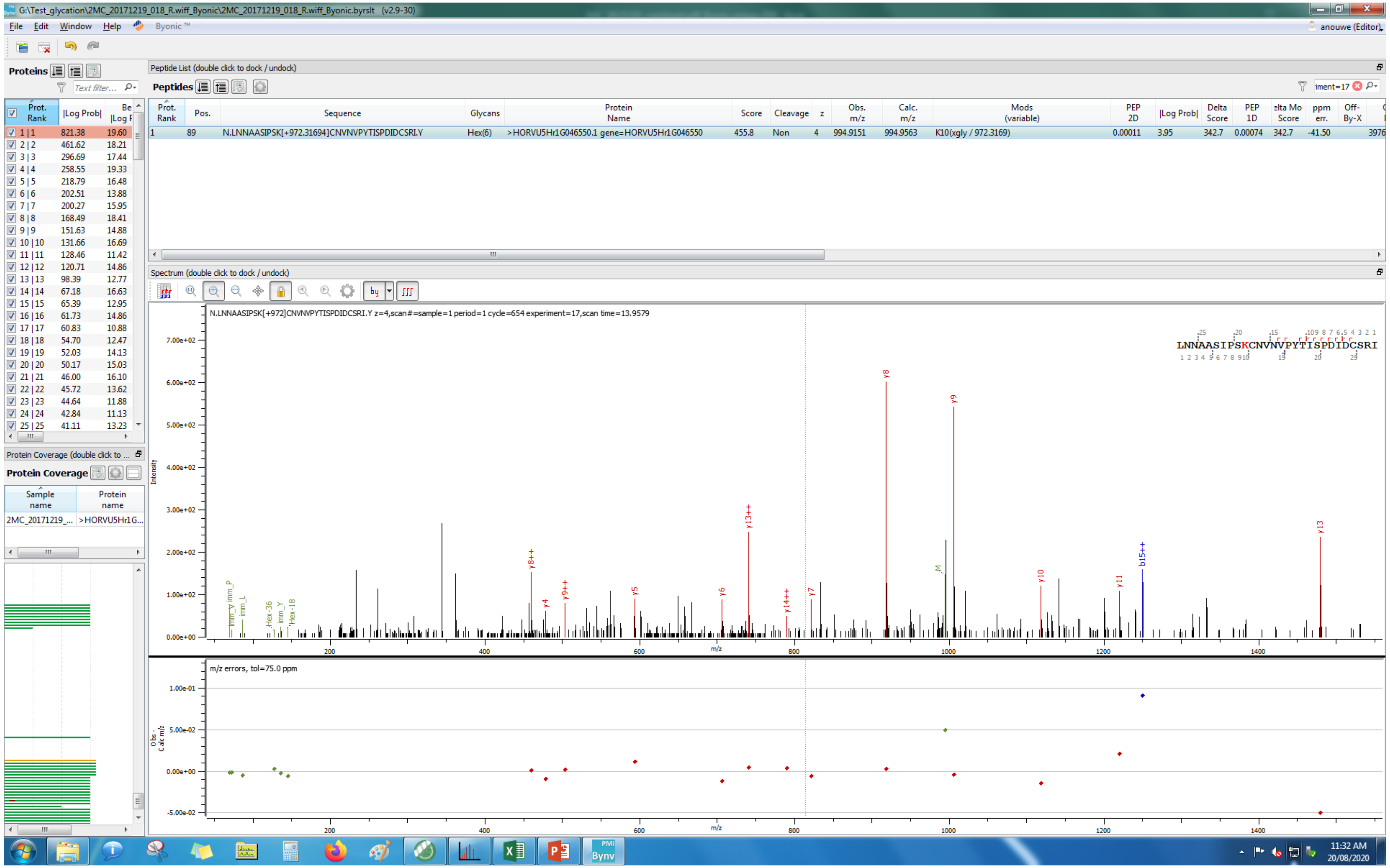
G:\Test_glycation\2MC_20171219_018_R.wiff_Byonic\2MC_20171219_018_R.wiff_Byonic.byrslt (v2.9-30)
File Edit Window Help Byonic ™
anouwe (Editor)
Proteins
Peptide List (double click to dock / undock)
Peptides
Prot. Rank | Pos. | Sequence | Glycans | Protein Name | Score | Cleavage | z | Obs. m/z | Calc. m/z | Mods (variable) | PEP 2D | |Log Prob| | Delta Score | PEP 1D | Delta Mod Score | ppm err. | Off-By-X
1 | 89 | N.LNNAASIPSK[+972.31694]CNVNVPYTISPDIDCSRI.Y | Hex(6) | >HORVU5Hr1G046550.1 gene=HORVU5Hr1G046550 | 455.8 | Non | 4 | 994.9151 | 994.9563 | K10(xgly / 972.3169) | 0.00011 | 3.95 | 342.7 | 0.00074 | 342.7 | -41.50 | 3976
Spectrum (double click to dock / undock)
N.LNNAASIPSK[+972]CNVNVPYTISPDIDCSRI.Y z=4,scan#=sample=1 period=1 cycle=654 experiment=17,scan time=13.9579
LNNAASIPSKCNVNVPYTISPDIDCSRI
Intensity
m/z
y8
y9
y13++
y13
b15++
y8++
y9++
y4
y5
y6
y7
y14++
y10
y11
m/z errors, tol=75.0 ppm
Obs - Calc m/z
Protein Coverage (double click to ...
Protein Coverage
Sample name | Protein name
2MC_20171219_... | >HORVU5Hr1G...
11:32 AM
20/08/2020

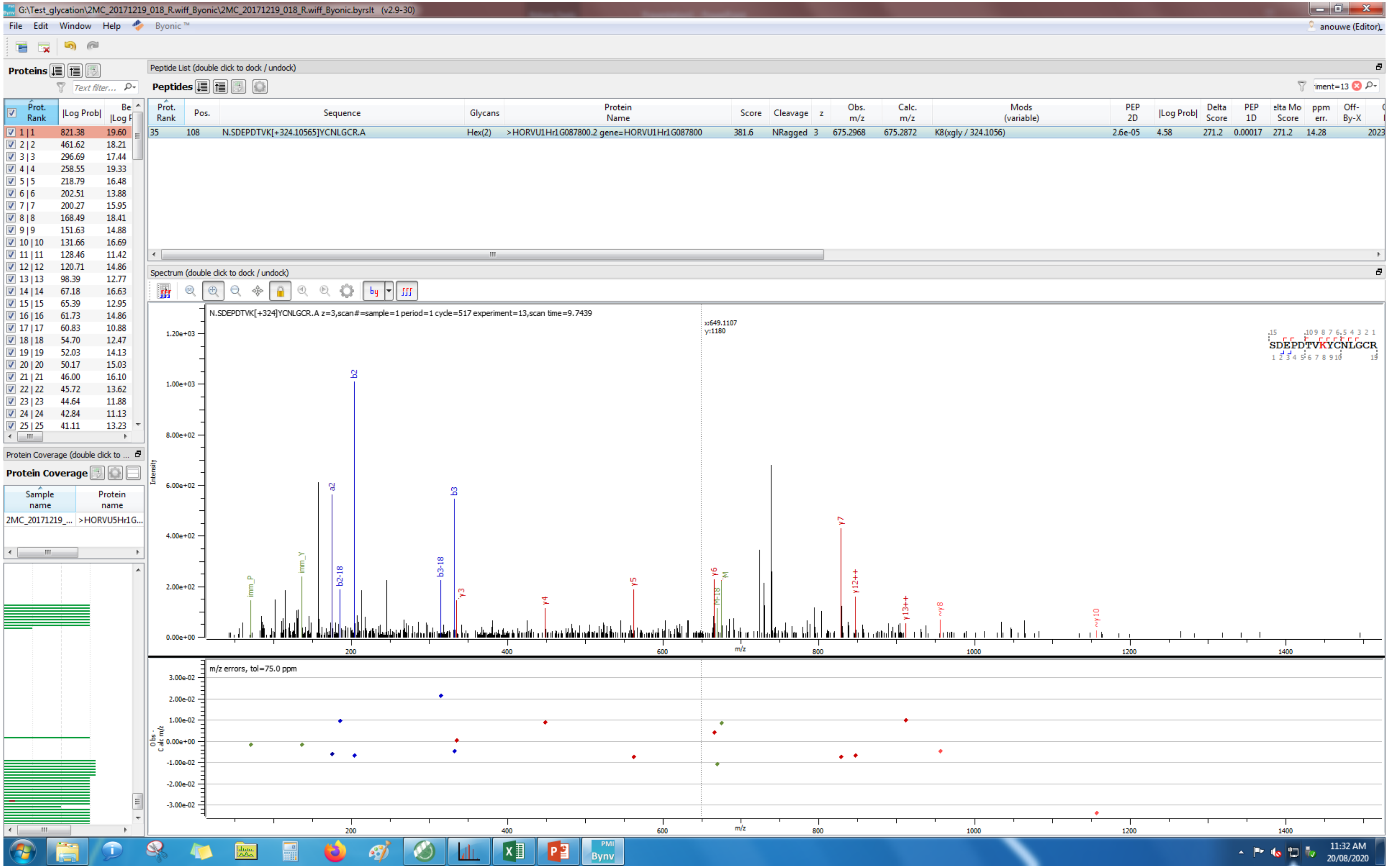
G:\Test_glycation\2MC_20171219_018_R.wiff_Byonic\2MC_20171219_018_R.wiff_Byonic.byrslt (v2.9-30)
File Edit Window Help Byonic ™
Proteins
Peptide List (double click to dock / undock)
Peptides
Prot. Rank | Pos. | Sequence | Glycans | Protein Name | Score | Cleavage | z | Obs. m/z | Calc. m/z | Mods (variable) | PEP 2D | |Log Prob| | Delta Score | PEP 1D | ppm err.
35 | 108 | N.SDEPDTVK[+324.10565]YCNLGCR.A | Hex(2) | >HORVU1Hr1G087800.2 gene=HORVU1Hr1G087800 | 381.6 | NRagged | 3 | 675.2968 | 675.2872 | K8(xgly / 324.1056) | 2.6e-05 | 4.58 | 271.2 | 0.00017 | 271.2 | 14.28
Spectrum (double click to dock / undock)
N.SDEPDTVK[+324]YCNLGCR.A z=3,scan#=sample=1 period=1 cycle=517 experiment=13,scan time=9.7439
x:649.1107
y:1180
SDEPDTVKYCNLGCR
Intensity
m/z
m/z errors, tol=75.0 ppm
Protein Coverage
Sample name
Protein name
2MC_20171219_...
>HORVU5Hr1G...
anouwe (Editor)
11:32 AM
20/08/2020

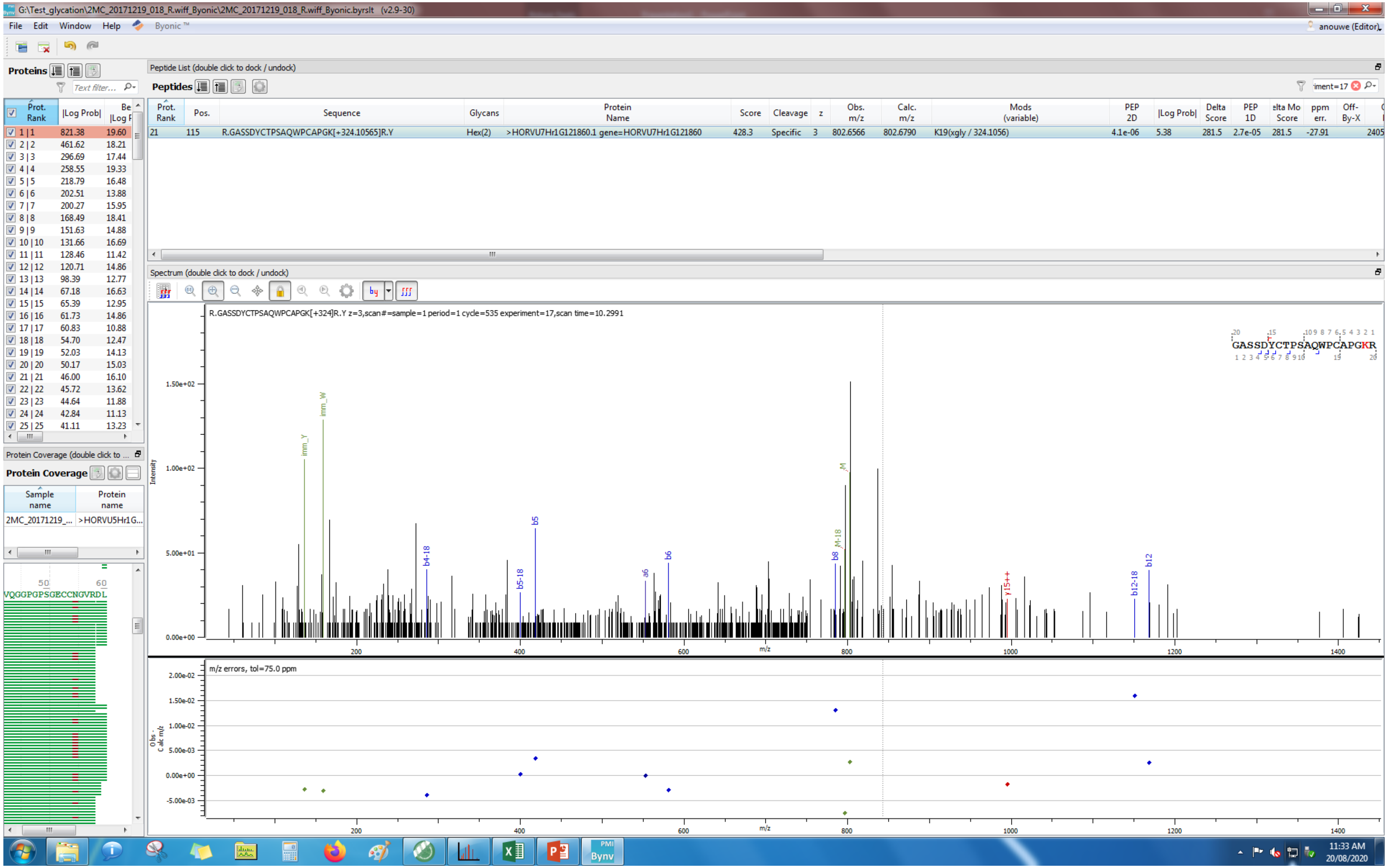

G:\Test_glycation\2MC_20171219_018_R.wiff_Byonic\2MC_20171219_018_R.wiff_Byonic.byrslt (v2.9-30)
File Edit Window Help Byonic™
anouwe (Editor)
Proteins
Peptide List (double click to dock / undock)
Peptides
Prot. Rank | Pos. | Sequence | Glycans | Protein Name | Score | Cleavage | z | Obs. m/z | Calc. m/z | Mods (variable) | PEP 2D | |Log Prob| | Delta Score | PEP 1D | Delta Mod Score | ppm err. | Off-By-X
21 | 115 | R.GASSDYCTPSAQWPCAPGK[+324.10565]R.Y | Hex(2) | >HORVU7Hr1G121860.1 gene=HORVU7Hr1G121860 | 428.3 | Specific | 3 | 802.6566 | 802.6790 | K19(xgly / 324.1056) | 4.1e-06 | 5.38 | 281.5 | 2.7e-05 | 281.5 | -27.91 | 2405
Spectrum (double click to dock / undock)
R.GASSDYCTPSAQWPCAPGK[+324]R.Y z=3,scan#=sample=1 period=1 cycle=535 experiment=17,scan time=10.2991
GASSDYCTPSAQWPCAPGKR
Intensity
m/z
m/z errors, tol=75.0 ppm
Obs - Calc m/z
Protein Coverage
Sample name
Protein name
2MC_20171219_...
>HORVU5Hr1G...
VQGGPGPSGECCNGVRDL
11:33 AM
20/08/2020

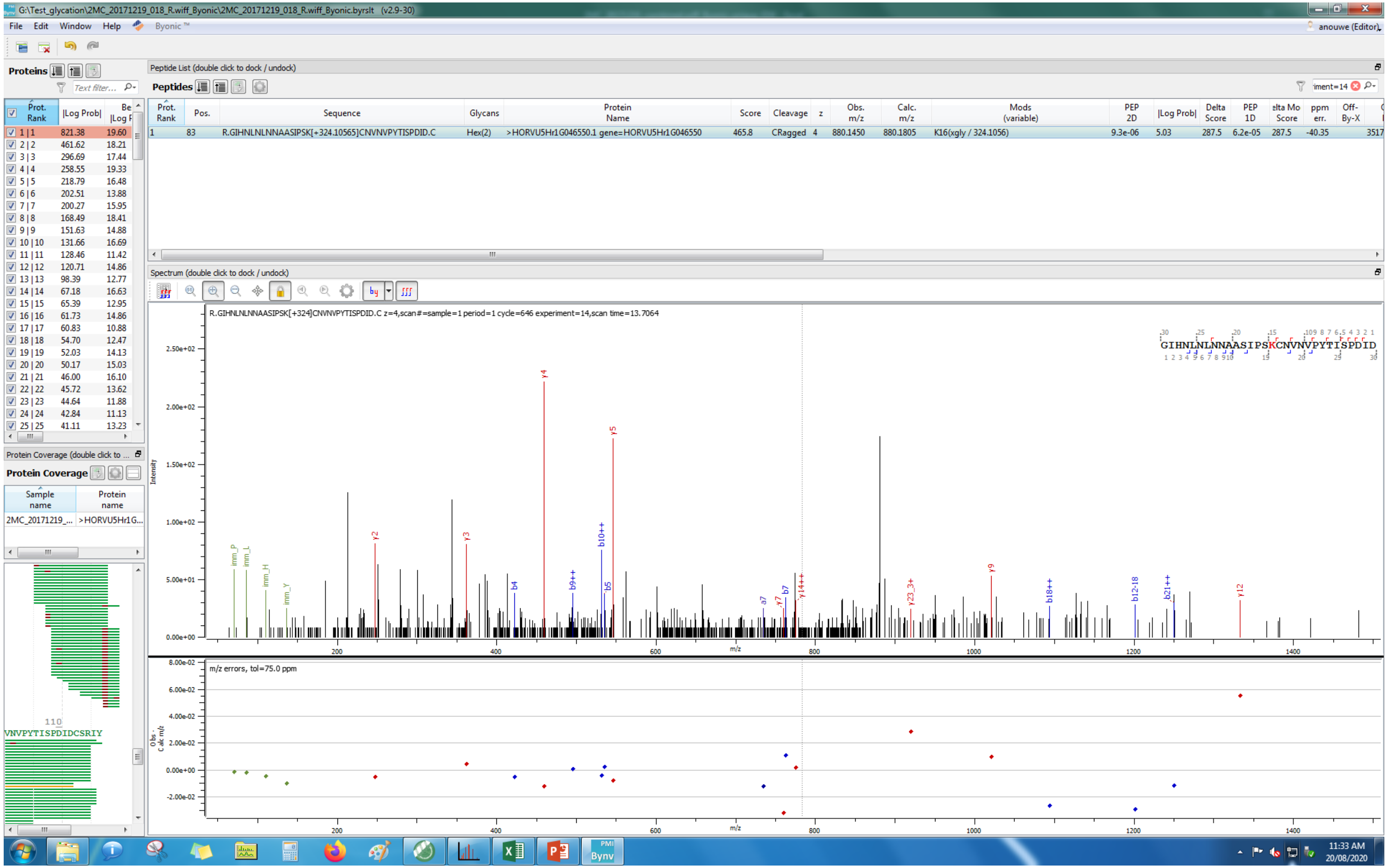
G:\Test_glycation\2MC_20171219_018_R.wiff_Byonic\2MC_20171219_018_R.wiff_Byonic.byrslt (v2.9-30)
File Edit Window Help Byonic™
anouwe (Editor)
Proteins
Peptide List (double click to dock / undock)
Peptides
Prot. Rank
Pos.
Sequence
Glycans
Protein Name
Score
Cleavage
z
Obs. m/z
Calc. m/z
Mods (variable)
PEP 2D
|Log Prob|
Delta Score
PEP 1D
ppm err.
Off-By-X
1 83 R.GIHNLNLNNAASIPSK[+324.10565]CNVNPYTISPDID.C Hex(2) >HORVU5Hr1G046550.1 gene=HORVU5Hr1G046550 465.8 CRagged 4 880.1450 880.1805 K16(xgly / 324.1056) 9.3e-06 5.03 287.5 6.2e-05 287.5 -40.35
Spectrum (double click to dock / undock)
R.GIHNLNLNNAASIPSK[+324]CNVNPYTISPDID.C z=4,scan#=sample=1 period=1 cycle=646 experiment=14,scan time=13.7064
GIHNLNLNNAASIPSKCNVNPYTISPDID
Intensity
m/z
m/z errors, tol=75.0 ppm
Protein Coverage (double click to ...
Protein Coverage
Sample name
Protein name
2MC_20171219_...
>HORVU5Hr1G...
VNPYTISPDIDCSRIY
11:33 AM
20/08/2020

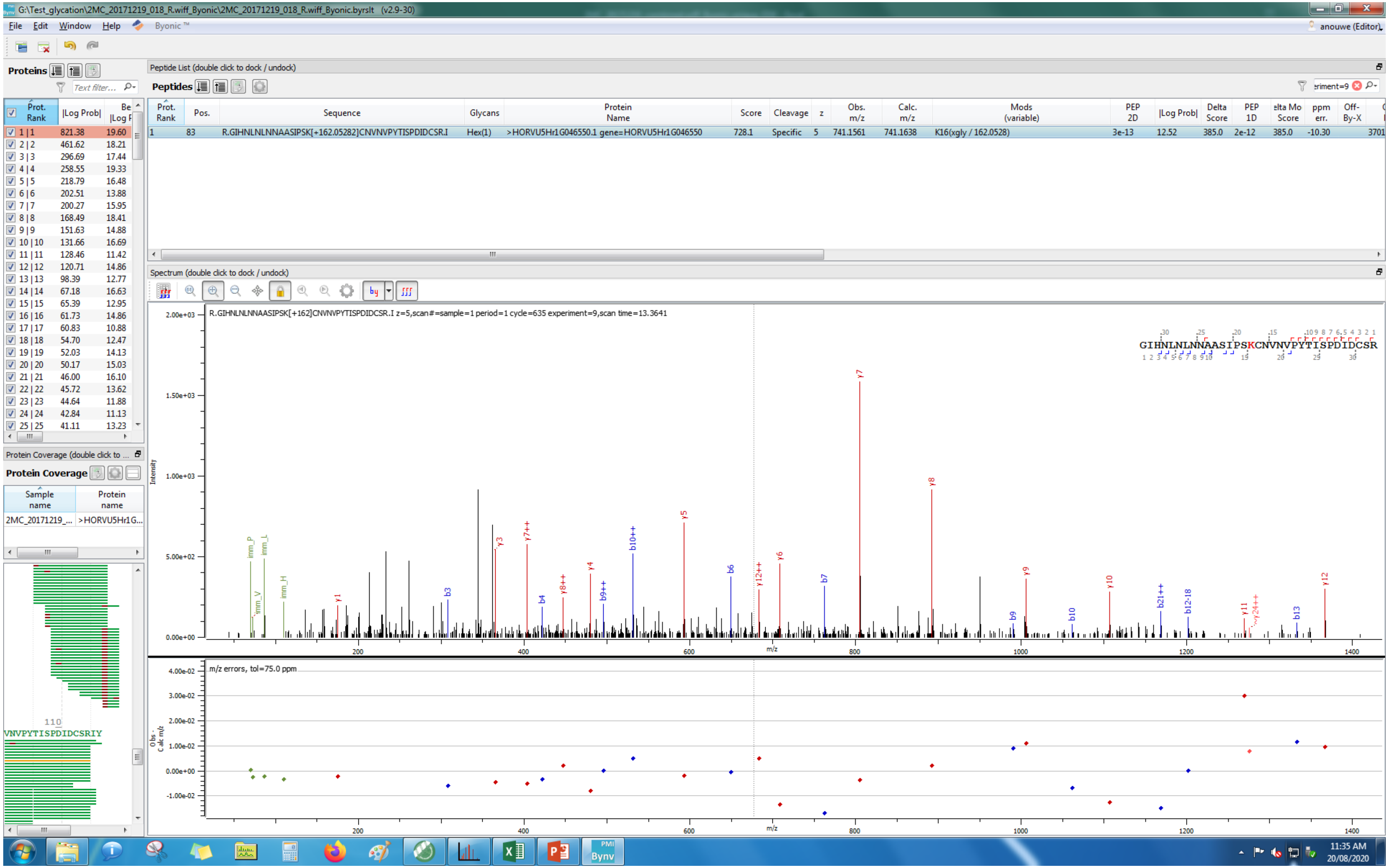

G:\Test_glycation\2MC_20171219_018_R.wiff_Byonic\2MC_20171219_018_R.wiff_Byonic.byrslt (v2.9-30)
File Edit Window Help Byonic ™
anouwe (Editor)
Proteins
Peptide List (double click to dock / undock)
Peptides
Prot. Rank | Pos. | Sequence | Glycans | Protein Name | Score | Cleavage | z | Obs. m/z | Calc. m/z | Mods (variable) | PEP 2D | |Log Prob| | Delta Score | PEP 1D | Delta Mod Score | ppm err. | Off-By-X
1 | 83 | R.GIHNLNLNNAASIPSK[+162.05282]CNVNVPYTISPDIDCSR.I | Hex(1) | >HORVU5Hr1G046550.1 gene=HORVU5Hr1G046550 | 728.1 | Specific | 5 | 741.1561 | 741.1638 | K16(xgly / 162.0528) | 3e-13 | 12.52 | 385.0 | 2e-12 | 385.0 | -10.30
Prot. Rank | |Log Prob| | Best |Log Prob|
1 | 1 821.38 19.60
2 | 2 461.62 18.21
3 | 3 296.69 17.44
4 | 4 258.55 19.33
5 | 5 218.79 16.48
6 | 6 202.51 13.88
7 | 7 200.27 15.95
8 | 8 168.49 18.41
9 | 9 151.63 14.88
10 | 10 131.66 16.69
11 | 11 128.46 11.42
12 | 12 120.71 14.86
13 | 13 98.39 12.77
14 | 14 67.18 16.63
15 | 15 65.39 12.95
16 | 16 61.73 14.86
17 | 17 60.83 10.88
18 | 18 54.70 12.47
19 | 19 52.03 14.13
20 | 20 50.17 15.03
21 | 21 46.00 16.10
22 | 22 45.72 13.62
23 | 23 44.64 11.88
24 | 24 42.84 11.13
25 | 25 41.11 13.23
Protein Coverage (double click to ...
Protein Coverage
Sample name | Protein name
2MC_20171219_... | >HORVU5Hr1G...
110
VNVPYTISPDIDCSRIY
Spectrum (double click to dock / undock)
R.GIHNLNLNNAASIPSK[+162]CNVNVPYTISPDIDCSR.I z=5,scan#=sample=1 period=1 cycle=635 experiment=9,scan time=13.3641
GIHNLNLNNAASIPSKCNVNVPYTISPDIDCSR
Intensity
m/z
m/z errors, tol=75.0 ppm
Obs - Calc m/z
11:35 AM
20/08/2020

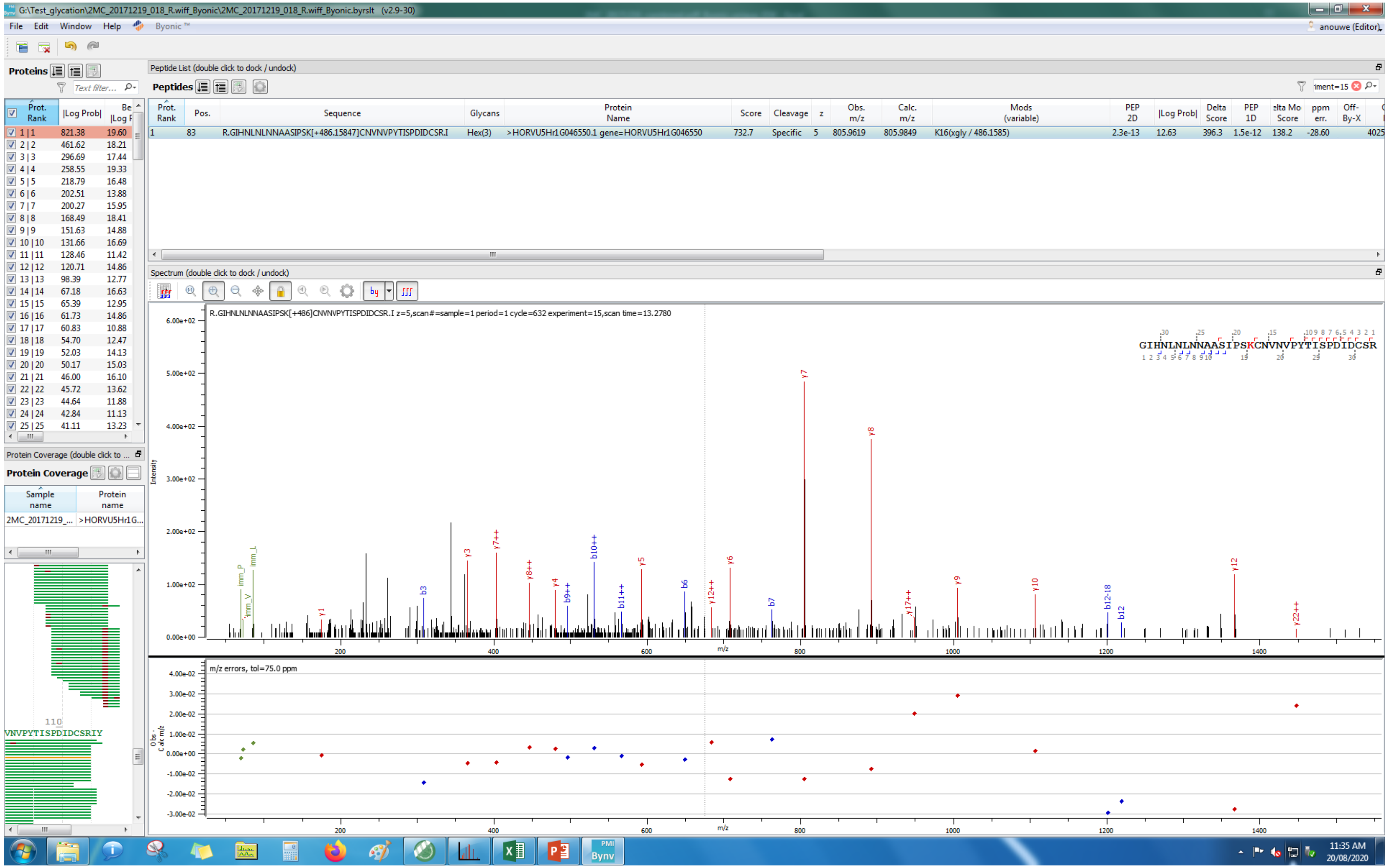
G:\Test_glycation\2MC_20171219_018_R.wiff_Byonic\2MC_20171219_018_R.wiff_Byonic.byrslt (v2.9-30)
File Edit Window Help Byonic™
Proteins
Peptide List (double click to dock / undock)
Peptides
R.GIHNLNLNNAASIPSK[+486.15847]CNVNPYTISPDIDCSR.I
Hex(3)
>HORVU5Hr1G046550.1 gene=HORVU5Hr1G046550
732.7
Specific
805.9619
805.9849
K16(xgly / 486.1585)
Spectrum (double click to dock / undock)
R.GIHNLNLNNAASIPSK[+486]CNVNPYTISPDIDCSR.I z=5,scan#=sample=1 period=1 cycle=632 experiment=15,scan time=13.2780
GIHNLNLNNAASIPSKCNVNPYTISPDIDCSR
m/z errors, tol=75.0 ppm
Protein Coverage
VNVPYTISPDIDCSRIY
11:35 AM
20/08/2020

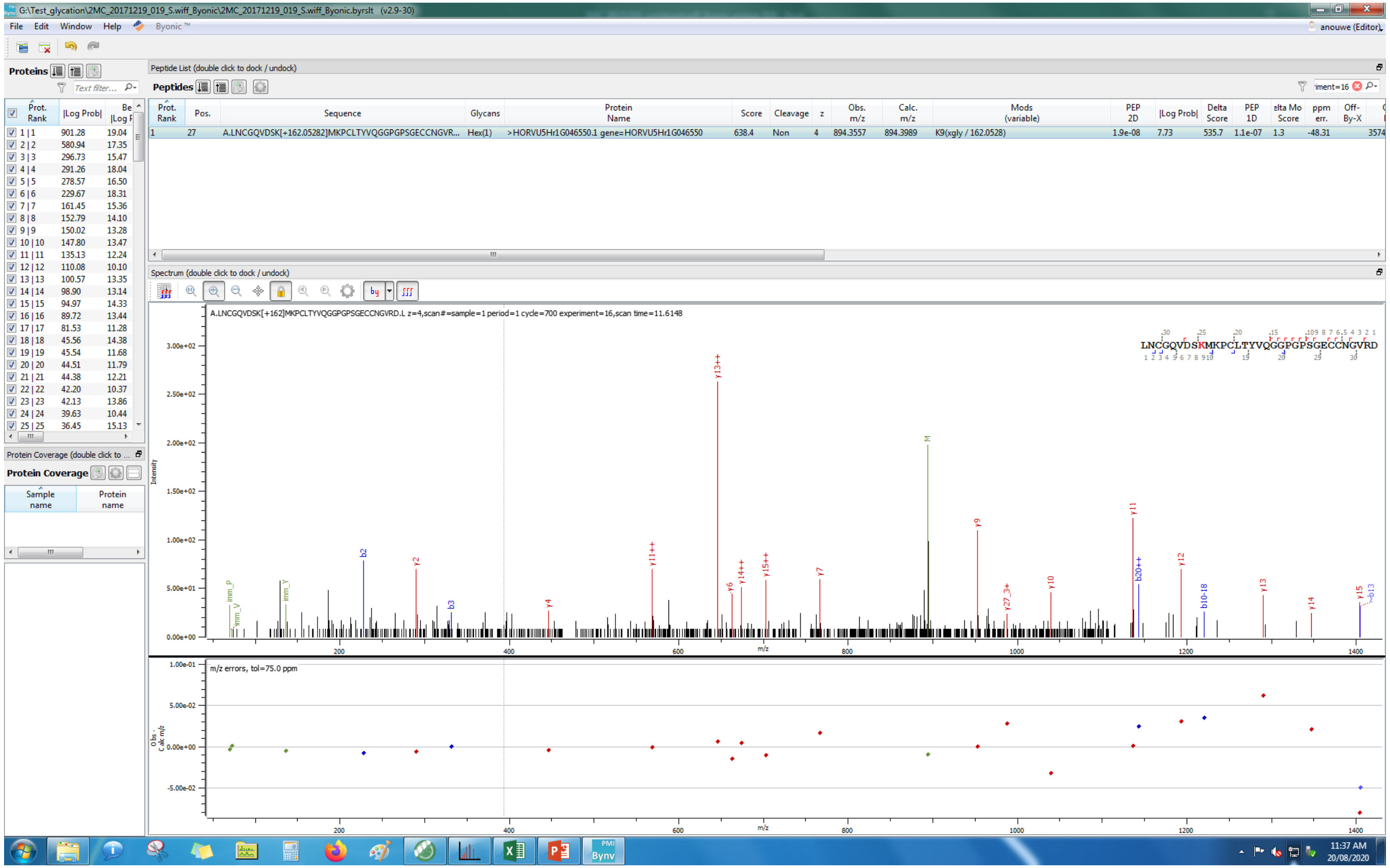
G:\Test_glycation\2MC_20171219_019_S.wiff_Byonic\2MC_20171219_019_S.wiff_Byonic.byrslt (v2.9-30)
File Edit Window Help Byonic ™
anouwe (Editor)
Proteins
Peptide List (double click to dock / undock)
Peptides
Prot. Rank | Pos. | Sequence | Glycans | Protein Name | Score | Cleavage | z | Obs. m/z | Calc. m/z | Mods (variable) | PEP 2D | |Log Prob| | Delta Score | PEP 1D | elta Mo Score | ppm err. | Off-By-X
1 | 27 | A.LNCGQVDSK[+162.05282]MKPCLTYVQGGPGPSGECCNGVR... | Hex(1) | >HORVU5Hr1G046550.1 gene=HORVU5Hr1G046550 | 638.4 | Non | 4 | 894.3557 | 894.3989 | K9(xgly / 162.0528) | 1.9e-08 | 7.73 | 535.7 | 1.1e-07 | 1.3 | -48.31 | 3574
Spectrum (double click to dock / undock)
A.LNCGQVDSK[+162]MKPCLTYVQGGPGPSGECCNGVRD.L z=4,scan#=sample=1 period=1 cycle=700 experiment=16,scan time=11.6148
LNCGQVDSKMKPCLTYVQGGPGPSGECCNGVRD
Intensity
m/z
m/z errors, tol=75.0 ppm
Protein Coverage (double click to ...
Protein Coverage
Sample name
Protein name
11:37 AM
20/08/2020

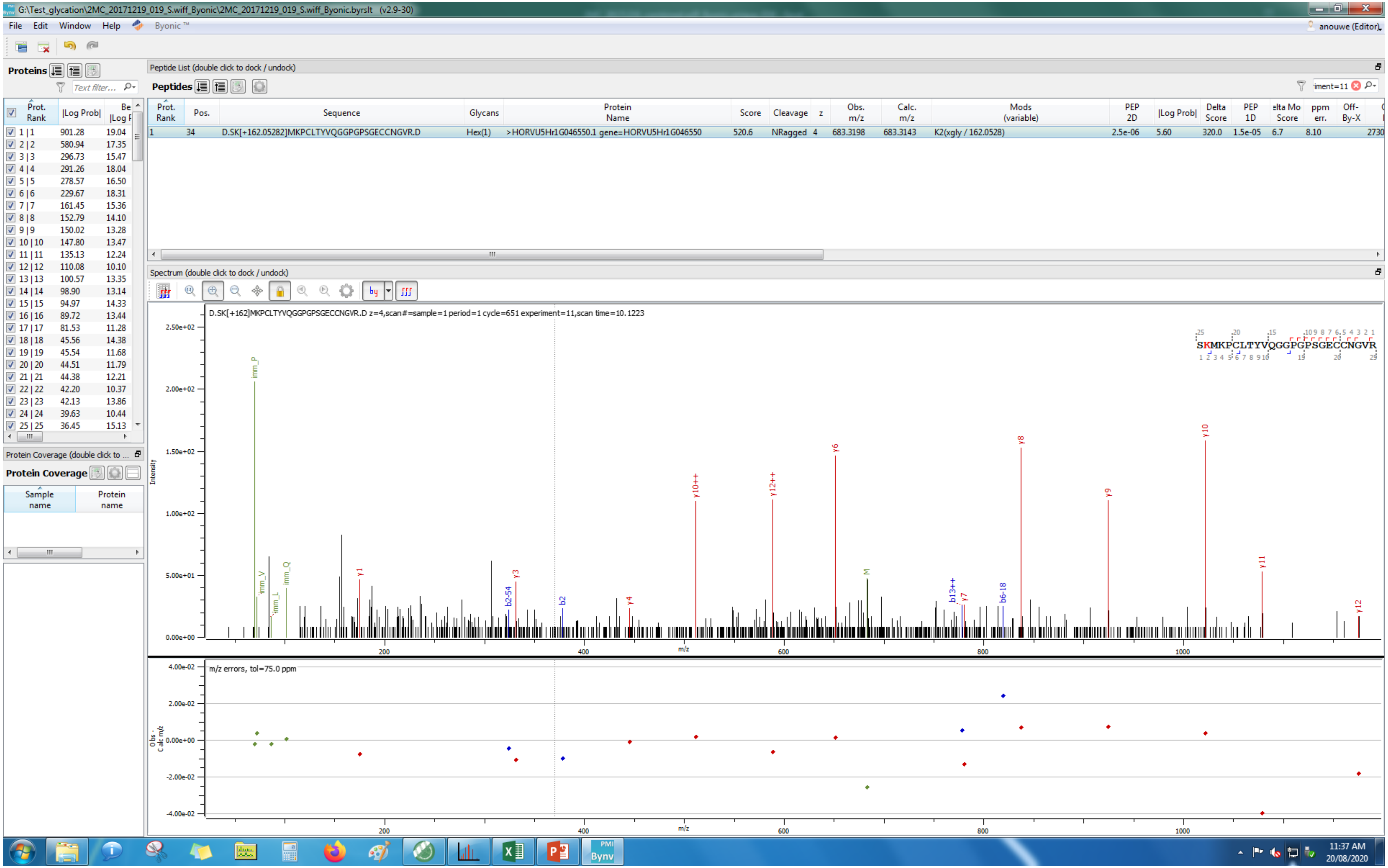
G:\Test_glycation\2MC_20171219_019_S.wiff_Byonic\2MC_20171219_019_S.wiff_Byonic.byrslt (v2.9-30)
File Edit Window Help Byonic™
anouwe (Editor)
Proteins
Peptide List (double click to dock / undock)
Peptides
Prot. Rank | Pos. | Sequence | Glycans | Protein Name | Score | Cleavage | z | Obs. m/z | Calc. m/z | Mods (variable) | PEP 2D | |Log Prob| | Delta Score | PEP 1D | elta Mo Score | ppm err. | Off-By-X
1 | 34 | D.SK[+162.05282]MKPCLTYVQGGPGPSGECCNGVR.D | Hex(1) | >HORVU5Hr1G046550.1 gene=HORVU5Hr1G046550 | 520.6 | NRagged | 4 | 683.3198 | 683.3143 | K2(xgly / 162.0528) | 2.5e-06 | 5.60 | 320.0 | 1.5e-05 | 6.7 | 8.10 | 2730
Prot. Rank | |Log Prob| | Be |Log P
1 | 1 901.28 19.04
2 | 2 580.94 17.35
3 | 3 296.73 15.47
4 | 4 291.26 18.04
5 | 5 278.57 16.50
6 | 6 229.67 18.31
7 | 7 161.45 15.36
8 | 8 152.79 14.10
9 | 9 150.02 13.28
10 | 10 147.80 13.47
11 | 11 135.13 12.24
12 | 12 110.08 10.10
13 | 13 100.57 13.35
14 | 14 98.90 13.14
15 | 15 94.97 14.33
16 | 16 89.72 13.44
17 | 17 81.53 11.28
18 | 18 45.56 14.38
19 | 19 45.54 11.68
20 | 20 44.51 11.79
21 | 21 44.38 12.21
22 | 22 42.20 10.37
23 | 23 42.13 13.86
24 | 24 39.63 10.44
25 | 25 36.45 15.13
Protein Coverage (double click to ...
Protein Coverage
Sample name | Protein name
Spectrum (double click to dock / undock)
D.SK[+162]MKPCLTYVQGGPGPSGECCNGVR.D z=4,scan#=sample=1 period=1 cycle=651 experiment=11,scan time=10.1223
SKMKPCLTYVQGGPGPSGECCNGVR
Intensity
m/z
imm_P imm_V imm_L imm_Q y1 b2-54 y3 b2 y4 y10++ y12++ y6 M b13++ y7 b6-18 y8 y9 y10 y11 y12
m/z errors, tol=75.0 ppm
Obs - Calc m/z
11:37 AM
20/08/2020

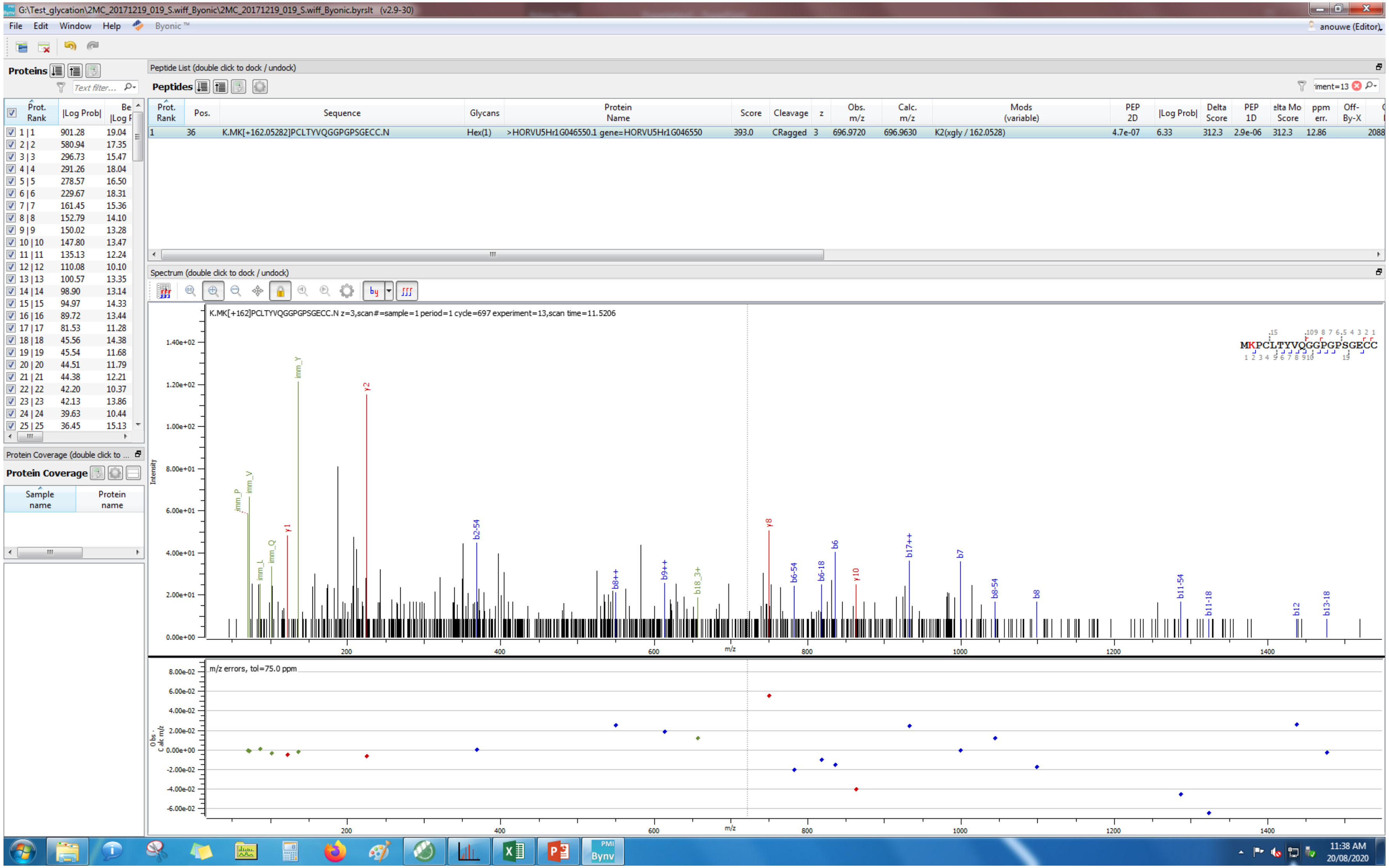
G:\Test_glycation\2MC_20171219_019_S.wiff_Byonic\2MC_20171219_019_S.wiff_Byonic.byrslt (v2.9-30)
File Edit Window Help Byonic™
anouwe (Editor)
Proteins
Peptide List (double click to dock / undock)
Peptides
Prot. Rank
Pos.
Sequence
Glycans
Protein Name
Score
Cleavage
z
Obs. m/z
Calc. m/z
Mods (variable)
PEP 2D
|Log Prob|
Delta Score
PEP 1D
ppm err.
Off-By-X
1 36 K.MK[+162.05282]PCLTYVQGGPGPSGECC.N Hex(1) >HORVU5Hr1G046550.1 gene=HORVU5Hr1G046550 393.0 CRagged 3 696.9720 696.9630 K2(xgly / 162.0528) 4.7e-07 6.33 312.3 2.9e-06 312.3 12.86 2088
Spectrum (double click to dock / undock)
K.MK[+162]PCLTYVQGGPGPSGECC.N z=3,scan#=sample=1 period=1 cycle=697 experiment=13,scan time=11.5206
MKPCLTYVQGGPGPSGECC
Intensity
m/z
m/z errors, tol=75.0 ppm
Protein Coverage (double click to ...
Protein Coverage
Sample name
Protein name
11:38 AM
20/08/2020

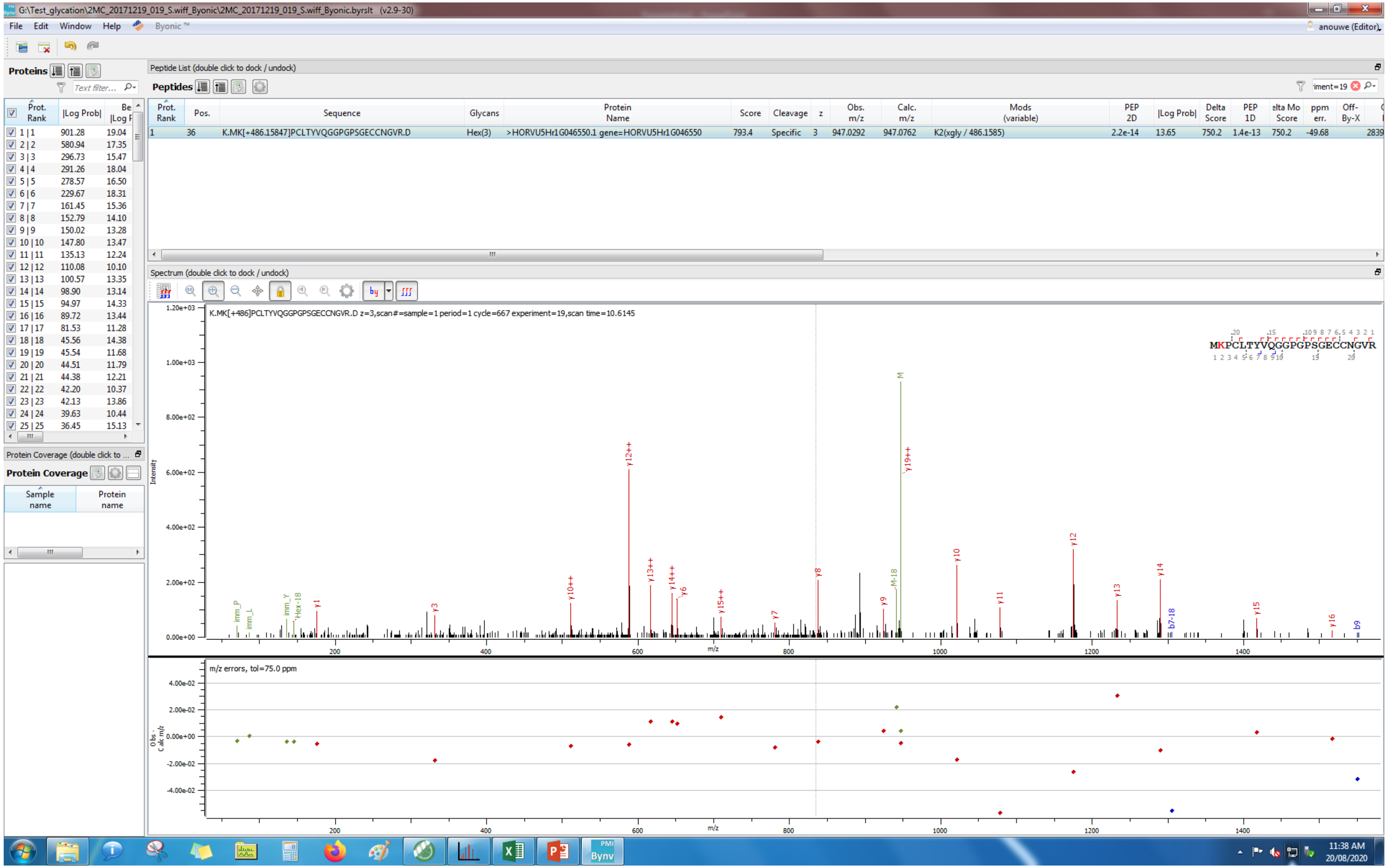

G:\Test_glycation\2MC_20171219_019_S.wiff_Byonic\2MC_20171219_019_S.wiff_Byonic.byrslt (v2.9-30)
File Edit Window Help Byonic™
Proteins
Peptide List (double click to dock / undock)
Peptides
K.MK[+486.15847]PCLTYVQGGPGPSGECCNGVR.D
Hex(3)
>HORVU5Hr1G046550.1 gene=HORVU5Hr1G046550
793.4
Specific
947.0292
947.0762
K2(xgly / 486.1585)
Spectrum (double click to dock / undock)
K.MK[+486]PCLTYVQGGPGPSGECCNGVR.D z=3,scan#=sample=1 period=1 cycle=667 experiment=19,scan time=10.6145
MKPCLTYVQGGPGPSGECCNGVR
m/z errors, tol=75.0 ppm
Protein Coverage

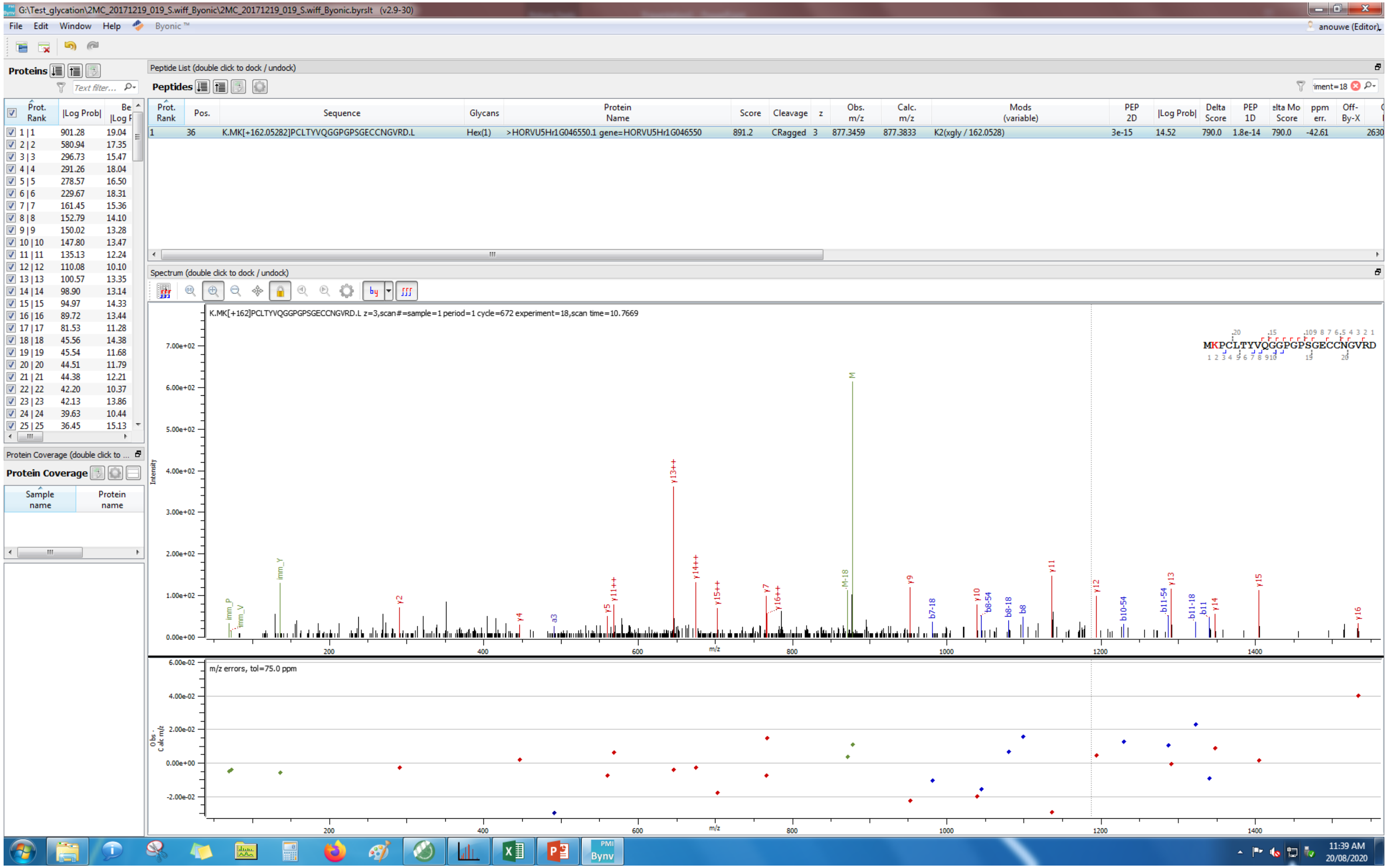

G:\Test_glycation\2MC_20171219_019_S.wiff_Byonic\2MC_20171219_019_S.wiff_Byonic.byrslt (v2.9-30)
File Edit Window Help Byonic™
anouwe (Editor)
Proteins
Peptide List (double click to dock / undock)
Peptides
Prot. Rank | Pos. | Sequence | Glycans | Protein Name | Score | Cleavage | z | Obs. m/z | Calc. m/z | Mods (variable) | PEP 2D | |Log Prob| | Delta Score | PEP 1D | elta Mo Score | ppm err. | Off-By-X
1 | 36 | K.MK[+162.05282]PCLTYVQGGPGPSGECCNGVRD.L | Hex(1) | >HORVU5Hr1G046550.1 gene=HORVU5Hr1G046550 | 891.2 | CRagged | 3 | 877.3459 | 877.3833 | K2(xgly / 162.0528) | 3e-15 | 14.52 | 790.0 | 1.8e-14 | 790.0 | -42.61 | 2630
Prot. Rank | |Log Prob| | Be |Log P
1 | 1 901.28 19.04
2 | 2 580.94 17.35
3 | 3 296.73 15.47
4 | 4 291.26 18.04
5 | 5 278.57 16.50
6 | 6 229.67 18.31
7 | 7 161.45 15.36
8 | 8 152.79 14.10
9 | 9 150.02 13.28
10 | 10 147.80 13.47
11 | 11 135.13 12.24
12 | 12 110.08 10.10
13 | 13 100.57 13.35
14 | 14 98.90 13.14
15 | 15 94.97 14.33
16 | 16 89.72 13.44
17 | 17 81.53 11.28
18 | 18 45.56 14.38
19 | 19 45.54 11.68
20 | 20 44.51 11.79
21 | 21 44.38 12.21
22 | 22 42.20 10.37
23 | 23 42.13 13.86
24 | 24 39.63 10.44
25 | 25 36.45 15.13
Protein Coverage (double click to ...
Protein Coverage
Sample name | Protein name
Spectrum (double click to dock / undock)
K.MK[+162]PCLTYVQGGPGPSGECCNGVRD.L z=3,scan#=sample=1 period=1 cycle=672 experiment=18,scan time=10.7669
MKPCLTYVQGGPGPSGECCNGVRD
Intensity
m/z
m/z errors, tol=75.0 ppm
Obs - Calc m/z
11:39 AM
20/08/2020

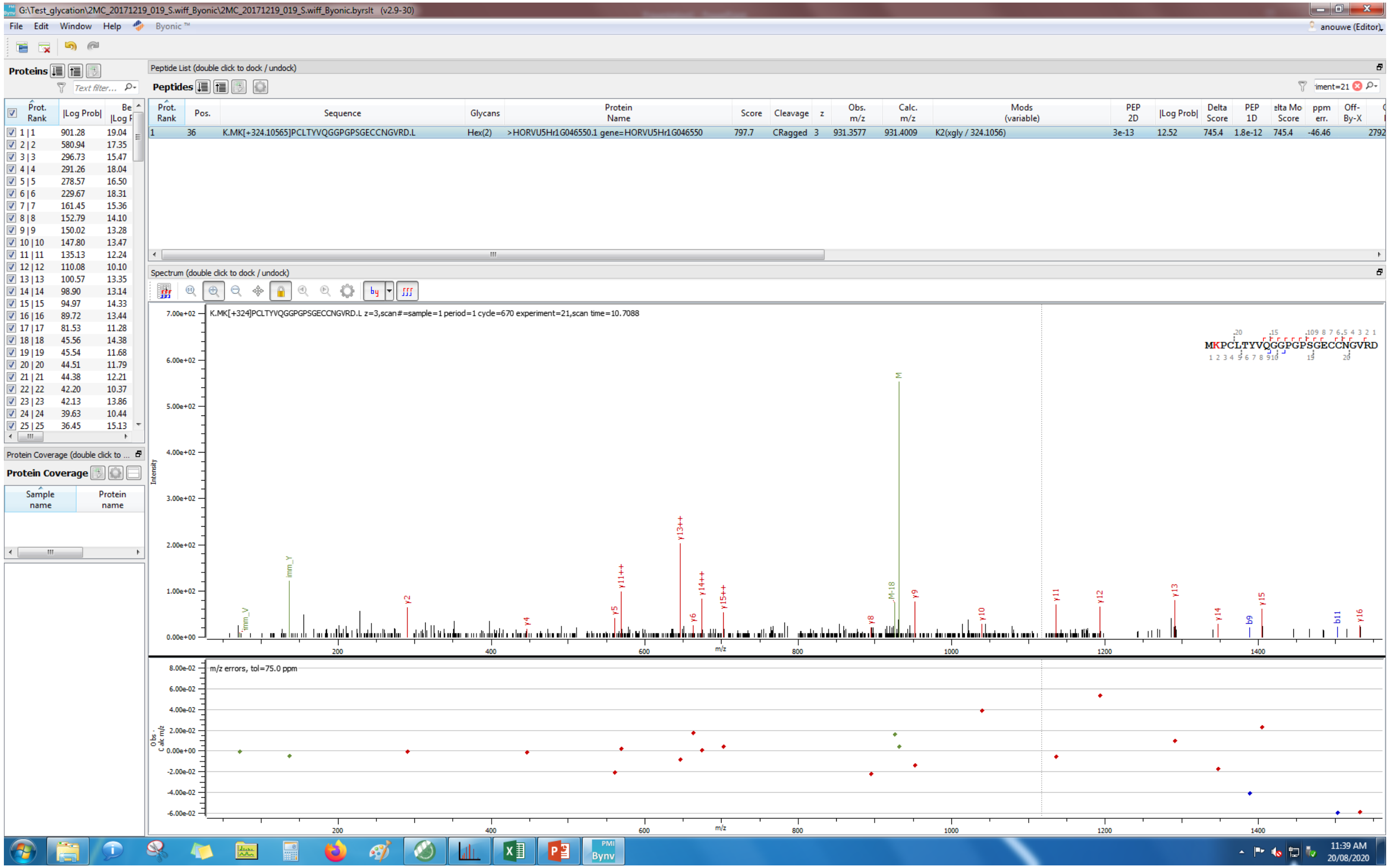
G:\Test_glycation\2MC_20171219_019_S.wiff_Byonic\2MC_20171219_019_S.wiff_Byonic.byrslt (v2.9-30)
File Edit Window Help Byonic
Proteins
Peptide List (double click to dock / undock)
Peptides
Prot. Rank | Pos. | Sequence | Glycans | Protein Name | Score | Cleavage | z | Obs. m/z | Calc. m/z | Mods (variable) | PEP 2D | |Log Prob| | Delta Score | PEP 1D | Delta Mod Score | ppm err. | Off-By-X
1 | 36 | K.MK[+324.10565]PCLTYVQGGPGPSGECCNGVRD.L | Hex(2) | >HORVU5Hr1G046550.1 gene=HORVU5Hr1G046550 | 797.7 | CRagged | 3 | 931.3577 | 931.4009 | K2(xgly / 324.1056) | 3e-13 | 12.52 | 745.4 | 1.8e-12 | 745.4 | -46.46
Spectrum (double click to dock / undock)
K.MK[+324]PCLTYVQGGPGPSGECCNGVRD.L z=3,scan#=sample=1 period=1 cycle=670 experiment=21,scan time=10.7088
MKPCLTYVQGGPGPSGECCNGVRD
Intensity
m/z
m/z errors, tol=75.0 ppm
Protein Coverage
Sample name
Protein name
11:39 AM
20/08/2020

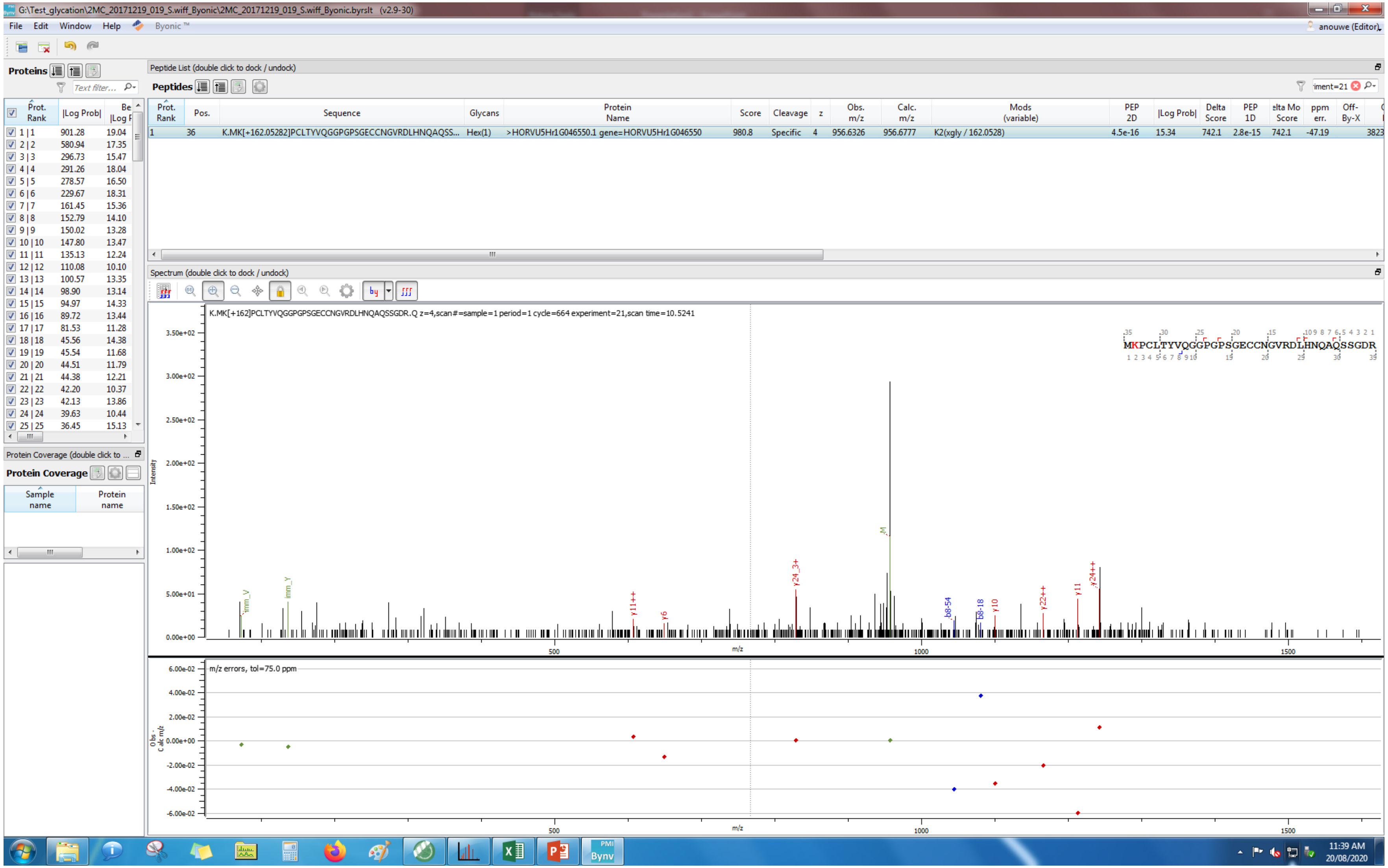
G:\Test_glycation\2MC_20171219_019_S.wiff_Byonic\2MC_20171219_019_S.wiff_Byonic.byrslt (v2.9-30)
File Edit Window Help Byonic ™
anouwe (Editor)
Proteins
Text filter...
Prot. Rank | |Log Prob| | Be |Log F
1 | 1 901.28 19.04
2 | 2 580.94 17.35
3 | 3 296.73 15.47
4 | 4 291.26 18.04
5 | 5 278.57 16.50
6 | 6 229.67 18.31
7 | 7 161.45 15.36
8 | 8 152.79 14.10
9 | 9 150.02 13.28
10 | 10 147.80 13.47
11 | 11 135.13 12.24
12 | 12 110.08 10.10
13 | 13 100.57 13.35
14 | 14 98.90 13.14
15 | 15 94.97 14.33
16 | 16 89.72 13.44
17 | 17 81.53 11.28
18 | 18 45.56 14.38
19 | 19 45.54 11.68
20 | 20 44.51 11.79
21 | 21 44.38 12.21
22 | 22 42.20 10.37
23 | 23 42.13 13.86
24 | 24 39.63 10.44
25 | 25 36.45 15.13
Protein Coverage (double click to ...
Protein Coverage
Sample name | Protein name
Peptide List (double click to dock / undock)
Peptides
iment=21
Prot. Rank | Pos. | Sequence | Glycans | Protein Name | Score | Cleavage | z | Obs. m/z | Calc. m/z | Mods (variable) | PEP 2D | |Log Prob| | Delta Score | PEP 1D | elta Mo Score | ppm err. | Off-By-X
1 | 36 | K.MK[+162.05282]PCLTYVQGGPGPSGECCNGVRDLHNQAQSS... | Hex(1) | >HORVU5Hr1G046550.1 gene=HORVU5Hr1G046550 | 980.8 | Specific | 4 | 956.6326 | 956.6777 | K2(xgly / 162.0528) | 4.5e-16 | 15.34 | 742.1 | 2.8e-15 | 742.1 | -47.19 | 3823
Spectrum (double click to dock / undock)
K.MK[+162]PCLTYVQGGPGPSGECCNGVRDLHNQAQSSGDR.Q z=4,scan#=sample=1 period=1 cycle=664 experiment=21,scan time=10.5241
MKPCLTYVQGGPGPSGECCNGVRDLHNQAQSSGDR
Intensity
m/z
m/z errors, tol=75.0 ppm
Obs - Calc m/z
11:39 AM
20/08/2020

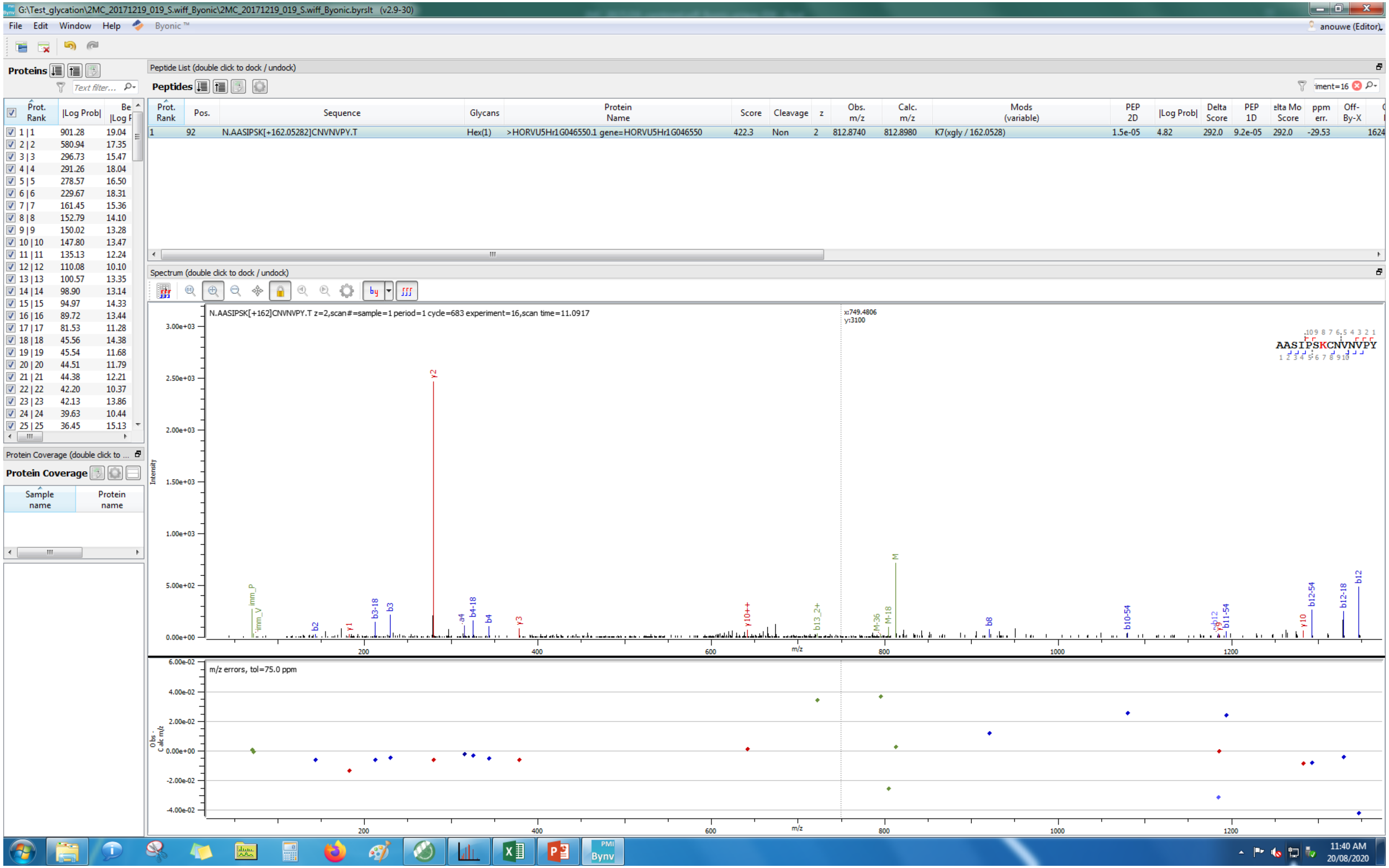
G:\Test_glycation\2MC_20171219_019_S.wiff_Byonic\2MC_20171219_019_S.wiff_Byonic.byrslt (v2.9-30)
File Edit Window Help Byonic™
anouwe (Editor)
Proteins
Peptide List (double click to dock / undock)
Peptides
Prot. Rank | Pos. | Sequence | Glycans | Protein Name | Score | Cleavage | z | Obs. m/z | Calc. m/z | Mods (variable) | PEP 2D | |Log Prob| | Delta Score | PEP 1D | Delta Mod Score | ppm err. | Off-By-X
1 | 92 | N.AASIPSK[+162.05282]CNVNVPY.T | Hex(1) | >HORVU5Hr1G046550.1 gene=HORVU5Hr1G046550 | 422.3 | Non | 2 | 812.8740 | 812.8980 | K7(xgly / 162.0528) | 1.5e-05 | 4.82 | 292.0 | 9.2e-05 | 292.0 | -29.53 | 1624
Spectrum (double click to dock / undock)
N.AASIPSK[+162]CNVNVPY.T z=2,scan#=sample=1 period=1 cycle=683 experiment=16,scan time=11.0917
x:749.4806
y:3100
AASIPSKCNVNVPY
Intensity
m/z
m/z errors, tol=75.0 ppm
Protein Coverage (double click to ...
Protein Coverage
Sample name
Protein name
11:40 AM
20/08/2020

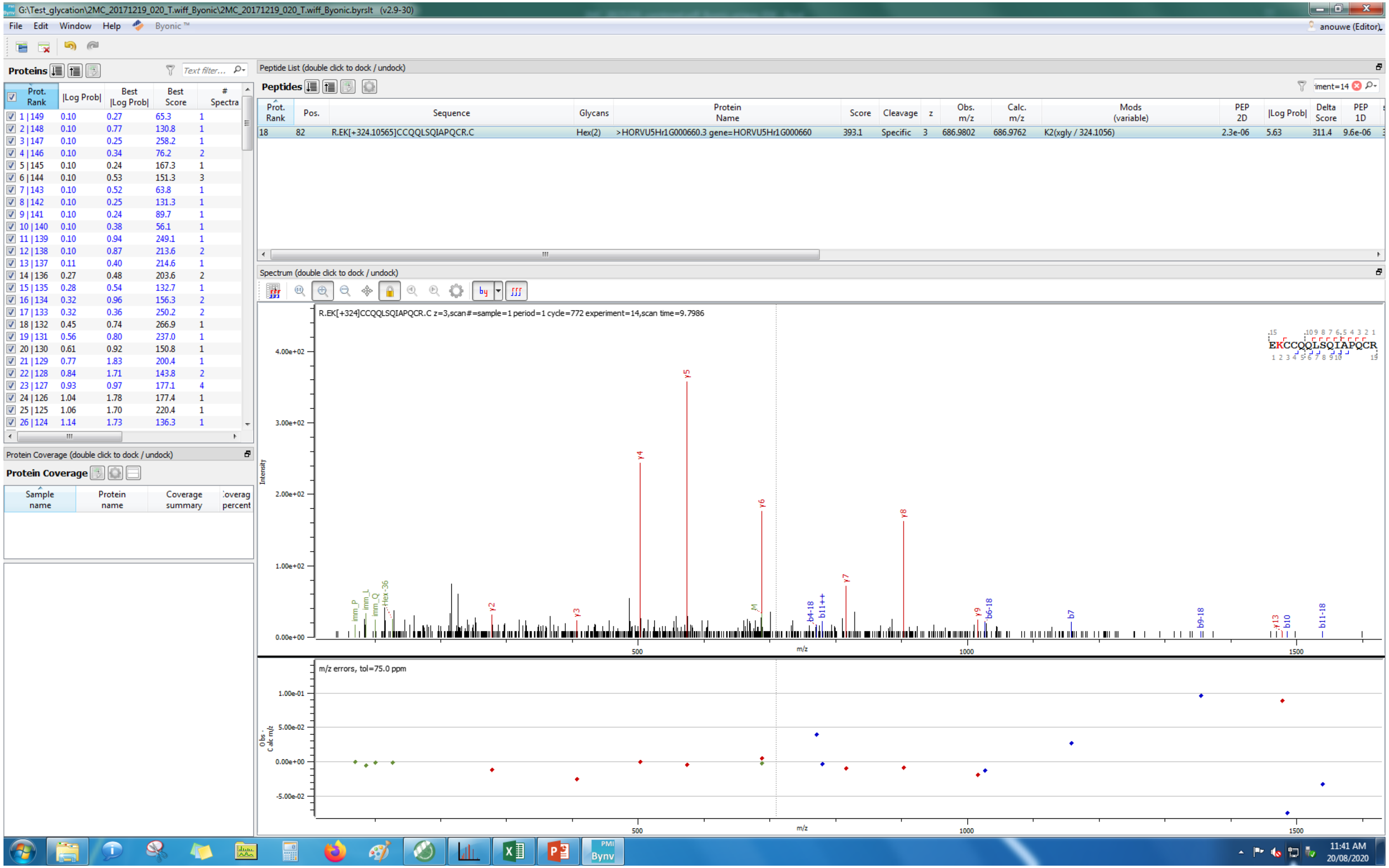
G:\Test_glycation\2MC_20171219_020_T.wiff_Byonic\2MC_20171219_020_T.wiff_Byonic.byrslt (v2.9-30)
File Edit Window Help Byonic ™
anouwe (Editor)
Proteins
Text filter...
Prot. Rank | |Log Prob| | Best |Log Prob| | Best Score | # Spectra
1 | 149 0.10 0.27 65.3 1
2 | 148 0.10 0.77 130.8 1
3 | 147 0.10 0.25 258.2 1
4 | 146 0.10 0.34 76.2 2
5 | 145 0.10 0.24 167.3 1
6 | 144 0.10 0.53 151.3 3
7 | 143 0.10 0.52 63.8 1
8 | 142 0.10 0.25 131.3 1
9 | 141 0.10 0.24 89.7 1
10 | 140 0.10 0.38 56.1 1
11 | 139 0.10 0.94 249.1 1
12 | 138 0.10 0.87 213.6 2
13 | 137 0.11 0.40 214.6 1
14 | 136 0.27 0.48 203.6 2
15 | 135 0.28 0.54 132.7 1
16 | 134 0.32 0.96 156.3 2
17 | 133 0.32 0.36 250.2 2
18 | 132 0.45 0.74 266.9 1
19 | 131 0.56 0.80 237.0 1
20 | 130 0.61 0.92 150.8 1
21 | 129 0.77 1.83 200.4 1
22 | 128 0.84 1.71 143.8 2
23 | 127 0.93 0.97 177.1 4
24 | 126 1.04 1.78 177.4 1
25 | 125 1.06 1.70 220.4 1
26 | 124 1.14 1.73 136.3 1
Protein Coverage (double click to dock / undock)
Protein Coverage
Sample name | Protein name | Coverage summary | Coverag percent
Peptide List (double click to dock / undock)
Peptides
iment=14
Prot. Rank | Pos. | Sequence | Glycans | Protein Name | Score | Cleavage | z | Obs. m/z | Calc. m/z | Mods (variable) | PEP 2D | |Log Prob| | Delta Score | PEP 1D
18 82 R.EK[+324.10565]CCQQLSQIAPQCR.C Hex(2) >HORVU5Hr1G000660.3 gene=HORVU5Hr1G000660 393.1 Specific 3 686.9802 686.9762 K2(xgly / 324.1056) 2.3e-06 5.63 311.4 9.6e-06
Spectrum (double click to dock / undock)
R.EK[+324]CCQQLSQIAPQCR.C z=3,scan#=sample=1 period=1 cycle=772 experiment=14,scan time=9.7986
EKCCQQLSQIAPQCR
4.00e+02
3.00e+02
2.00e+02
1.00e+02
0.00e+00
Intensity
y5
y4
y6
y8
y7
y2
y3
y9
b6-18
b7
b9-18
y13
b10
b11-18
b4-18
b11++
imm_P
imm_L
imm_Q
Hex-36
M
500
1000
1500
m/z
m/z errors, tol=75.0 ppm
1.00e-01
5.00e-02
0.00e+00
-5.00e-02
Obs - Calc m/z
11:41 AM
20/08/2020

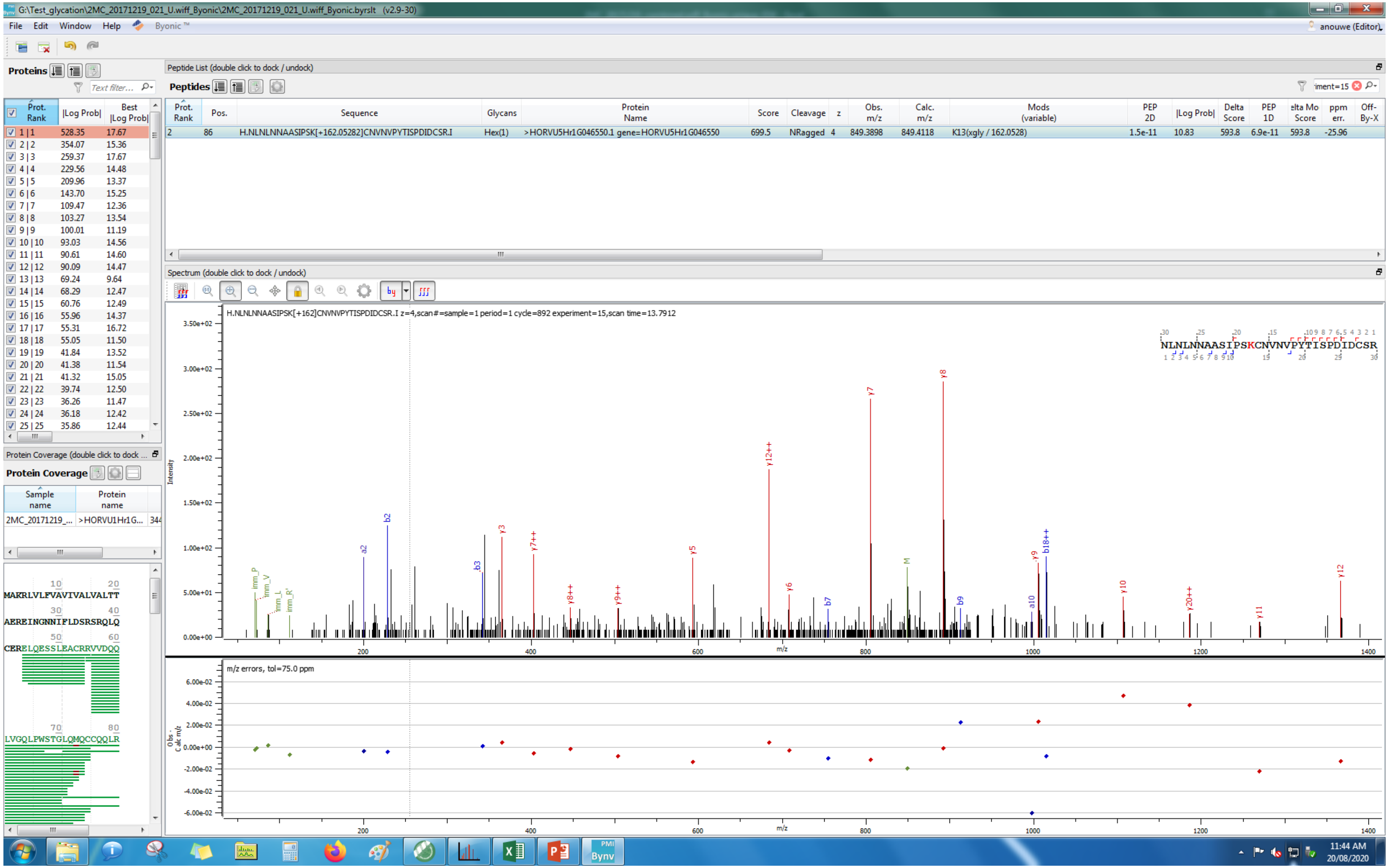
G:\Test_glycation\2MC_20171219_021_U.wiff_Byonic\2MC_20171219_021_U.wiff_Byonic.byrslt (v2.9-30)
File Edit Window Help Byonic
anouwe (Editor)
Proteins
Peptide List (double click to dock / undock)
Peptides
Prot. Rank | Pos. | Sequence | Glycans | Protein Name | Score | Cleavage | z | Obs. m/z | Calc. m/z | Mods (variable) | PEP 2D | |Log Prob| | Delta Score | PEP 1D | Delta Mod Score | ppm err. | Off-By-X
2 | 86 | H.NLNLNNAASIPSK[+162.05282]CNVNVPYTISPDIDCSR.I | Hex(1) | >HORVU5Hr1G046550.1 gene=HORVU5Hr1G046550 | 699.5 | NRagged | 4 | 849.3898 | 849.4118 | K13(xgly / 162.0528) | 1.5e-11 | 10.83 | 593.8 | 6.9e-11 | 593.8 | -25.96
Spectrum (double click to dock / undock)
H.NLNLNNAASIPSK[+162]CNVNVPYTISPDIDCSR.I z=4,scan#=sample=1 period=1 cycle=892 experiment=15,scan time=13.7912
NLNLNNAASIPSKCNVNVPYTISPDIDCSR
Intensity
m/z
m/z errors, tol=75.0 ppm
Obs - Calc m/z
Protein Coverage (double click to dock / undock)
Protein Coverage
Sample name | Protein name
2MC_20171219_... | >HORVU1Hr1G...
MAKRLVLFVAVIVALVALTT
AERENGNNIFLDSRSRQLQ
CERELQESSLEACRRVVDQQ
LVGQLPWSTGLQMQCCQQLR
11:44 AM
20/08/2020

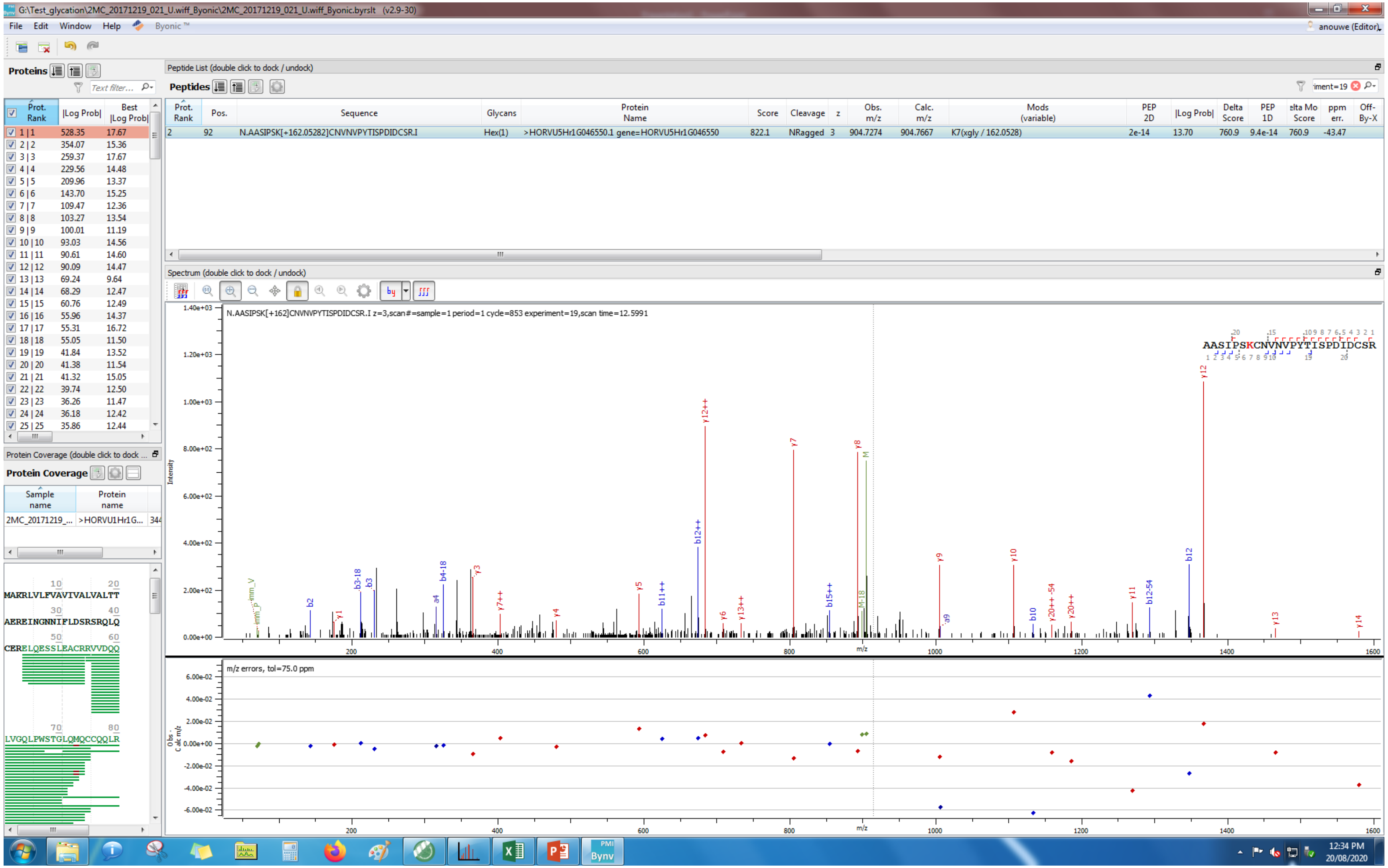
G:\Test_glycation\2MC_20171219_021_U.wiff_Byonic\2MC_20171219_021_U.wiff_Byonic.byrslt (v2.9-30)
File Edit Window Help Byonic ™
Proteins
Peptide List (double click to dock / undock)
Peptides
Prot. Rank | Pos. | Sequence | Glycans | Protein Name | Score | Cleavage | z | Obs. m/z | Calc. m/z | Mods (variable) | PEP 2D | |Log Prob| | Delta Score | PEP 1D | Delta Mod Score | ppm err. | Off-By-X
2 | 92 | N.AASIPSK[+162.05282]CNVNVPYTISPDIDCSR.I | Hex(1) | >HORVU5Hr1G046550.1 gene=HORVU5Hr1G046550 | 822.1 | NRagged | 3 | 904.7274 | 904.7667 | K7(xgly / 162.0528) | 2e-14 | 13.70 | 760.9 | 9.4e-14 | 760.9 | -43.47
Spectrum (double click to dock / undock)
N.AASIPSK[+162]CNVNVPYTISPDIDCSR.I z=3,scan#=sample=1 period=1 cycle=853 experiment=19,scan time=12.5991
AASIPSKCNVNVPYTISPDIDCSR
m/z errors, tol=75.0 ppm
Protein Coverage (double click to dock
Protein Coverage
Sample name
Protein name
MAKRLVLFVAVIVALVALTT
AEREINGNNIFLDSRSRQLQ
CERELQESSLEACRRVVDQQ
LVGQLPWSTGLQMQCCQQLR

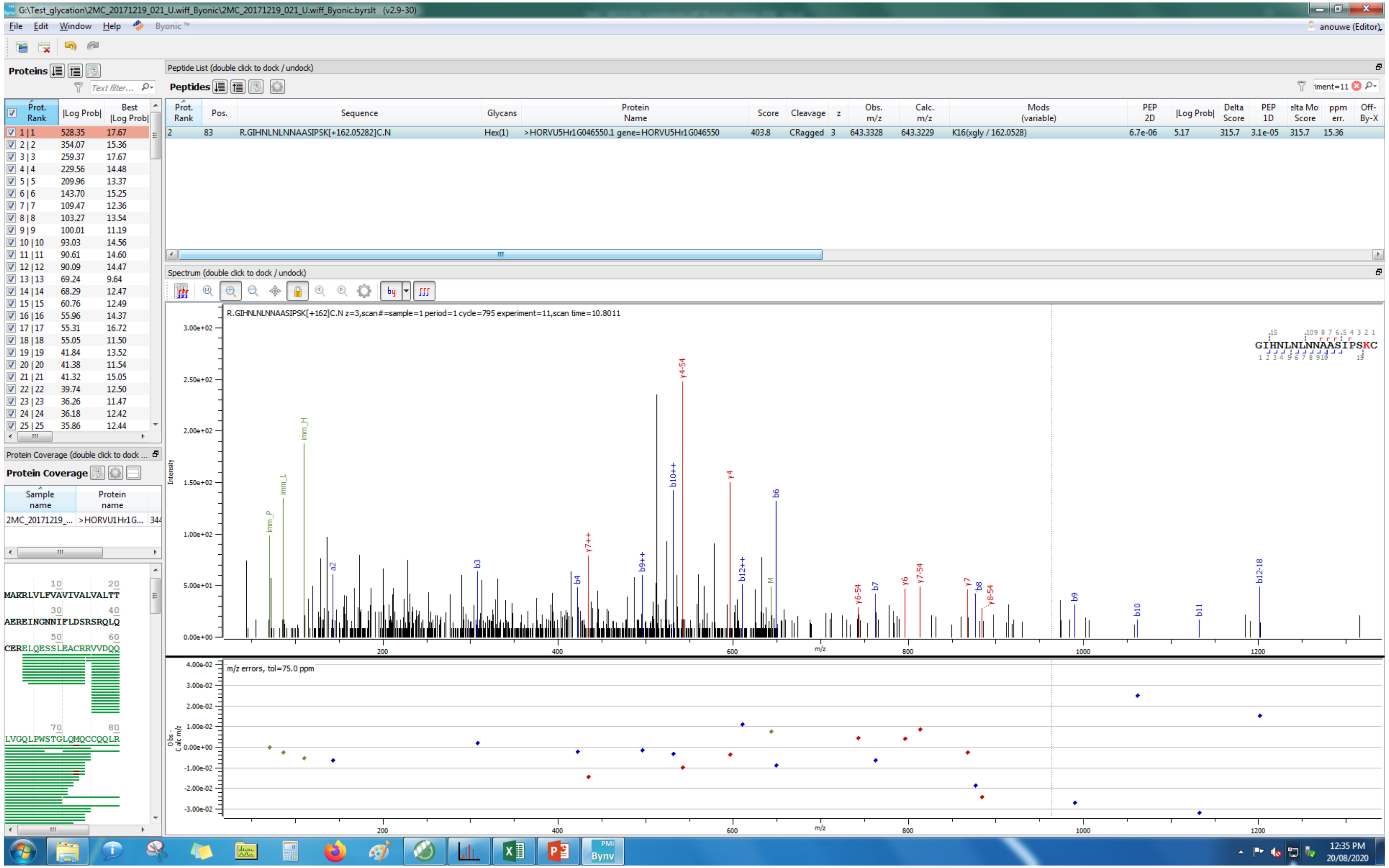
G:\Test_glycation\2MC_20171219_021_U.wiff_Byonic\2MC_20171219_021_U.wiff_Byonic.byrslt (v2.9-30)
File Edit Window Help Byonic ™
Proteins
Peptide List (double click to dock / undock)
Peptides
Prot. Rank | Pos. | Sequence | Glycans | Protein Name | Score | Cleavage | z | Obs. m/z | Calc. m/z | Mods (variable) | PEP 2D | |Log Prob| | Delta Score | PEP 1D | Delta Mod Score | ppm err. | Off-By-X
2 | 83 | R.GIHNLNLNNAASIPSK[+162.05282]C.N | Hex(1) | >HORVU5Hr1G046550.1 gene=HORVU5Hr1G046550 | 403.8 | CRagged | 3 | 643.3328 | 643.3229 | K16(xgly / 162.0528) | 6.7e-06 | 5.17 | 315.7 | 3.1e-05 | 315.7 | 15.36
Spectrum (double click to dock / undock)
R.GIHNLNLNNAASIPSK[+162]C.N z=3,scan#=sample=1 period=1 cycle=795 experiment=11,scan time=10.8011
m/z errors, tol=75.0 ppm
Protein Coverage
MAKRLVLFVAVIVALVALTT
AEREINGNNIFLDSRSRQLQ
CERELQESSLEACRRVVDQQ
LVGQLPWSTGLQMQCCQQLR

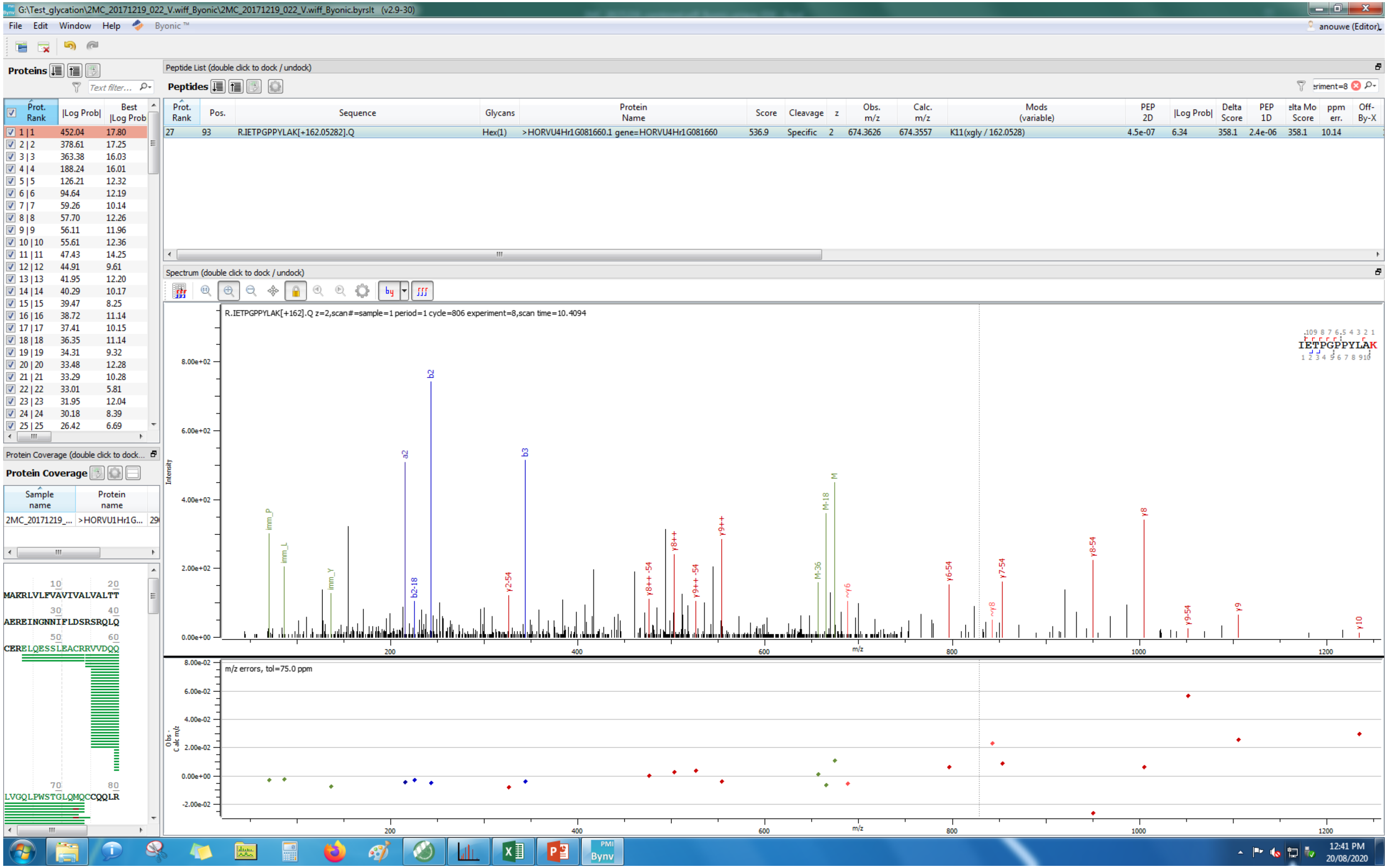
G:\Test_glycation\2MC_20171219_022_V.wiff_Byonic\2MC_20171219_022_V.wiff_Byonic.byrslt (v2.9-30)
File Edit Window Help Byonic™
anouwe (Editor)
Proteins
Peptide List (double click to dock / undock)
Peptides
Prot. Rank | Pos. | Sequence | Glycans | Protein Name | Score | Cleavage | z | Obs. m/z | Calc. m/z | Mods (variable) | PEP 2D | |Log Prob| | Delta Score | PEP 1D | Delta Mod Score | ppm err. | Off-By-X
27 | 93 | R.IETPGPPYLAK[+162.05282].Q | Hex(1) | >HORVU4Hr1G081660.1 gene=HORVU4Hr1G081660 | 536.9 | Specific | 2 | 674.3626 | 674.3557 | K11(xgly / 162.0528) | 4.5e-07 | 6.34 | 358.1 | 2.4e-06 | 358.1 | 10.14
Spectrum (double click to dock / undock)
R.IETPGPPYLAK[+162].Q z=2,scan#=sample=1 period=1 cycle=806 experiment=8,scan time=10.4094
IETPGPPYLAK
Intensity
m/z
m/z errors, tol=75.0 ppm
Protein Coverage (double click to dock...
Protein Coverage
Sample name
Protein name
2MC_20171219_...
>HORVU1Hr1G...
MAKRLVLFVAVIVALVALTT
AEREINGNNIFLDSRSRQLQ
CERELQESSLEACRRVVDQQ
LVGQLPWSTGLQMQCCQQLR
12:41 PM
20/08/2020

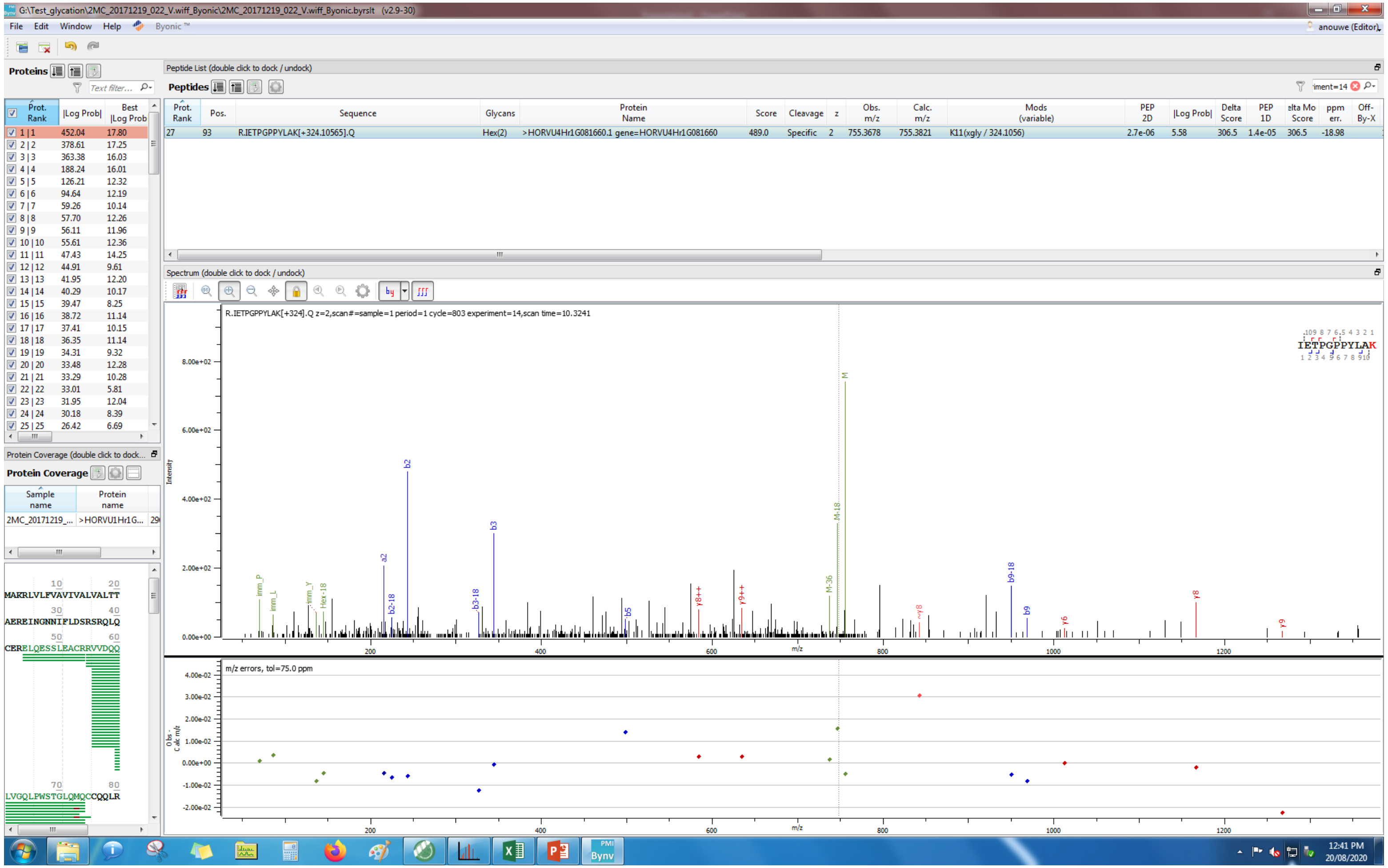

G:\Test_glycation\2MC_20171219_022_V.wiff_Byonic\2MC_20171219_022_V.wiff_Byonic.byrslt (v2.9-30)
File Edit Window Help Byonic™
anouwe (Editor)
Proteins
Peptide List (double click to dock / undock)
Peptides
Prot. Rank | Pos. | Sequence | Glycans | Protein Name | Score | Cleavage | z | Obs. m/z | Calc. m/z | Mods (variable) | PEP 2D | |Log Prob| | Delta Score | PEP 1D | elta Mo Score | ppm err. | Off-By-X
27 | 93 | R.IETPGPPYLAK[+324.10565].Q | Hex(2) | >HORVU4Hr1G081660.1 gene=HORVU4Hr1G081660 | 489.0 | Specific | 2 | 755.3678 | 755.3821 | K11(xgly / 324.1056) | 2.7e-06 | 5.58 | 306.5 | 1.4e-05 | 306.5 | -18.98
Prot. Rank | |Log Prob| | Best |Log Prob
1 | 1 | 452.04 | 17.80
2 | 2 | 378.61 | 17.25
3 | 3 | 363.38 | 16.03
4 | 4 | 188.24 | 16.01
5 | 5 | 126.21 | 12.32
6 | 6 | 94.64 | 12.19
7 | 7 | 59.26 | 10.14
8 | 8 | 57.70 | 12.26
9 | 9 | 56.11 | 11.96
10 | 10 | 55.61 | 12.36
11 | 11 | 47.43 | 14.25
12 | 12 | 44.91 | 9.61
13 | 13 | 41.95 | 12.20
14 | 14 | 40.29 | 10.17
15 | 15 | 39.47 | 8.25
16 | 16 | 38.72 | 11.14
17 | 17 | 37.41 | 10.15
18 | 18 | 36.35 | 11.14
19 | 19 | 34.31 | 9.32
20 | 20 | 33.48 | 12.28
21 | 21 | 33.29 | 10.28
22 | 22 | 33.01 | 5.81
23 | 23 | 31.95 | 12.04
24 | 24 | 30.18 | 8.39
25 | 25 | 26.42 | 6.69
Protein Coverage (double click to dock...
Protein Coverage
Sample name | Protein name
2MC_20171219_... | >HORVU1Hr1G...
MAKRLVLFVAVIVALVALTT
AEREINGNNIFLDSRSRQLQ
CERELQESSLEACRRVVDQQ
LVGQLPWSTGLQMQCCQQLR
Spectrum (double click to dock / undock)
R.IETPGPPYLAK[+324].Q z=2,scan#=sample=1 period=1 cycle=803 experiment=14,scan time=10.3241
IETPGPPYLAK
Intensity
m/z
m/z errors, tol=75.0 ppm
Obs - Calc m/z
12:41 PM
20/08/2020

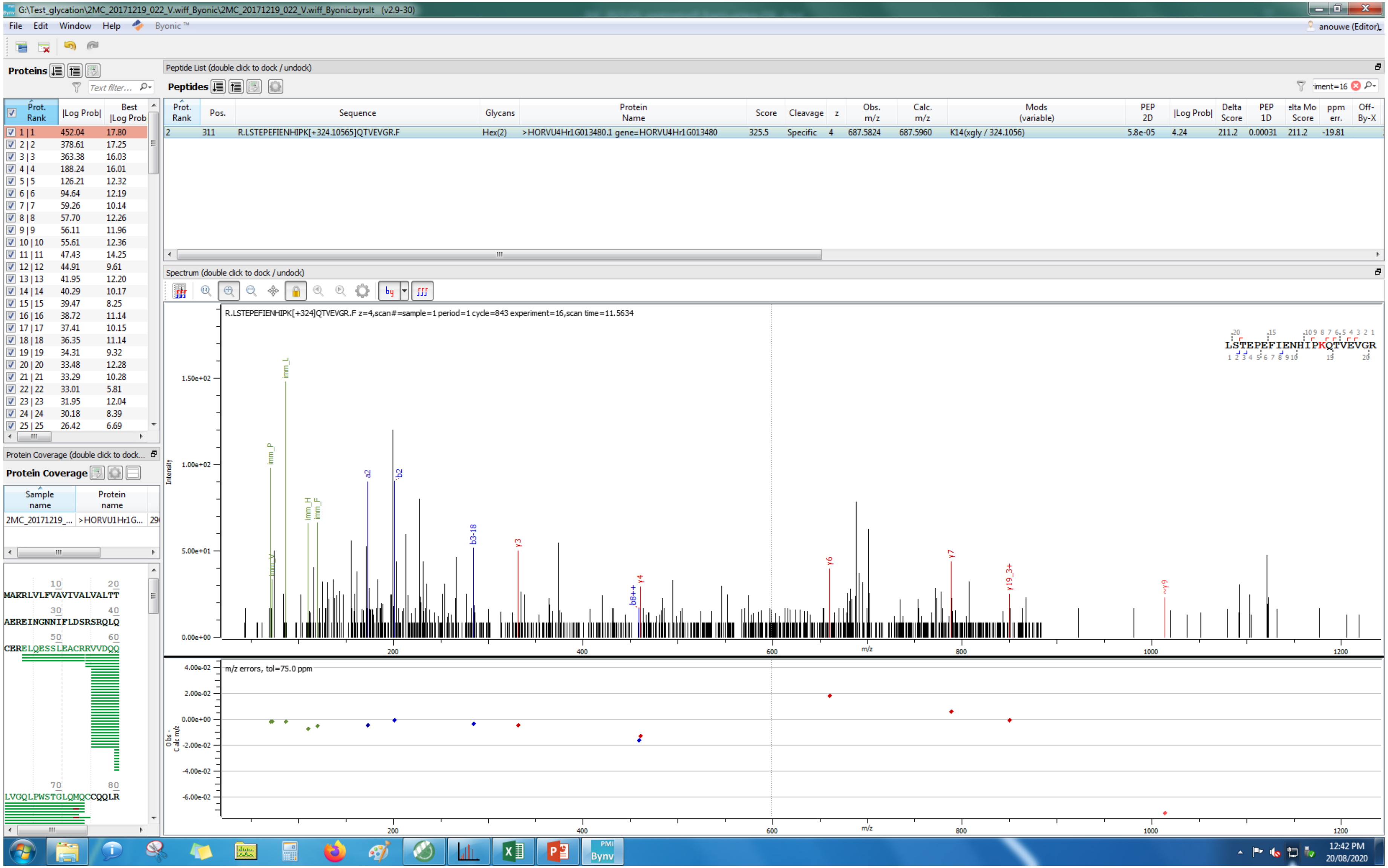
G:\Test_glycation\2MC_20171219_022_V.wiff_Byonic\2MC_20171219_022_V.wiff_Byonic.byrslt (v2.9-30)
File Edit Window Help Byonic™
anouwe (Editor)
Proteins
Peptide List (double click to dock / undock)
Peptides
Prot. Rank | Pos. | Sequence | Glycans | Protein Name | Score | Cleavage | z | Obs. m/z | Calc. m/z | Mods (variable) | PEP 2D | |Log Prob| | Delta Score | PEP 1D | elta Mo Score | ppm err. | Off-By-X
2 | 311 | R.LSTEPEFIENHIPK[+324.10565]QTVEVGR.F | Hex(2) | >HORVU4Hr1G013480.1 gene=HORVU4Hr1G013480 | 325.5 | Specific | 4 | 687.5824 | 687.5960 | K14(xgly / 324.1056) | 5.8e-05 | 4.24 | 211.2 | 0.00031 | 211.2 | -19.81
Spectrum (double click to dock / undock)
R.LSTEPEFIENHIPK[+324]QTVEVGR.F z=4,scan#=sample=1 period=1 cycle=843 experiment=16,scan time=11.5634
LSTEPEFIENHIPKQTVEVGR
Intensity
m/z
m/z errors, tol=75.0 ppm
Obs. - Calc m/z
Protein Coverage (double click to dock...
Protein Coverage
Sample name | Protein name
2MC_20171219_... | >HORVU1Hr1G...
MAKRLVLFVAVIVALVALTT
AEREINGNNIFLDSRSRQLQ
CERELQESSLEACRRVVDQQ
LVGQLPWSTGLQMQCCQQLR
12:42 PM
20/08/2020

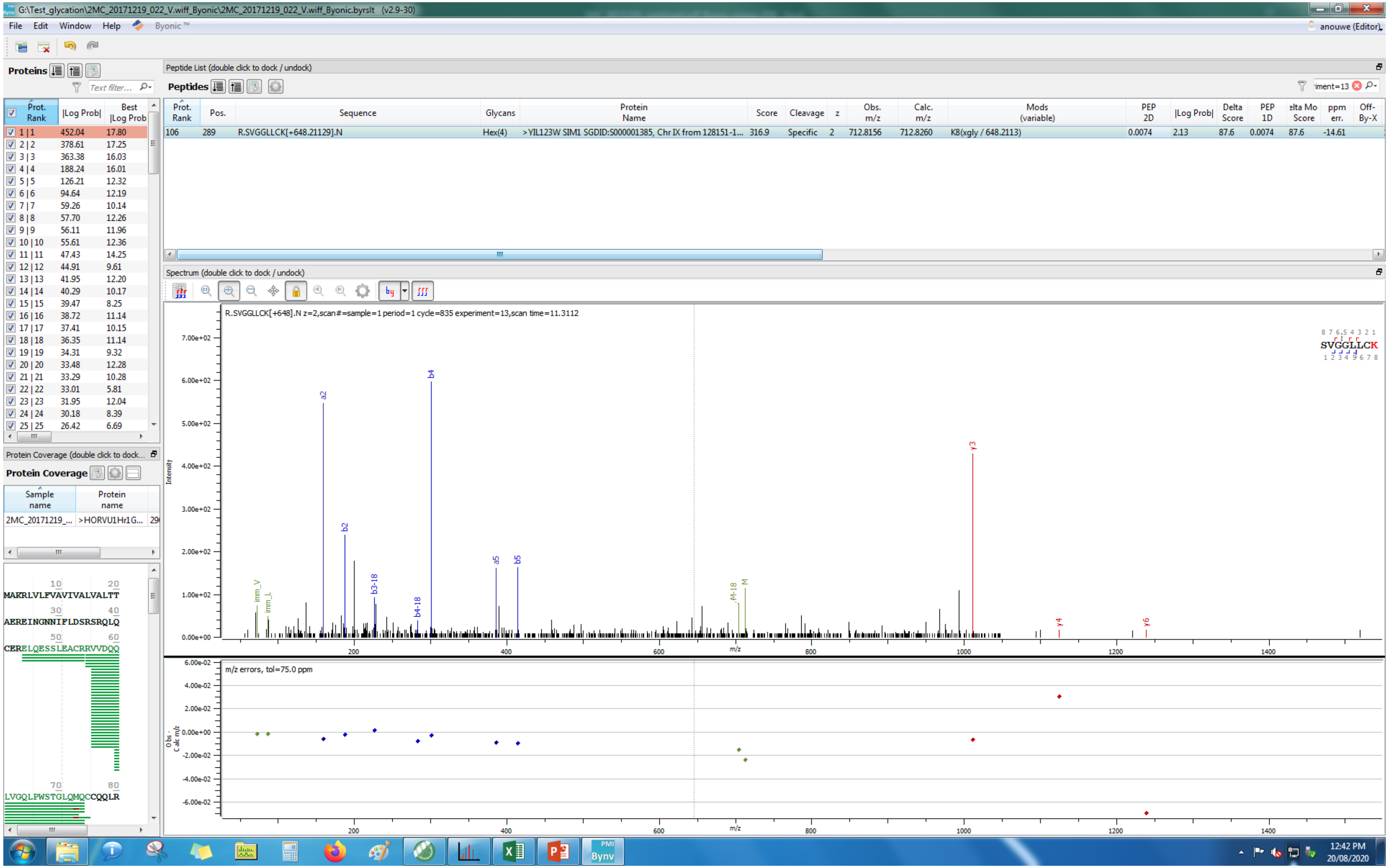
G:\Test_glycation\2MC_20171219_022_V.wiff_Byonic\2MC_20171219_022_V.wiff_Byonic.byrslt (v2.9-30)
File Edit Window Help Byonic™
anouwe (Editor)
Proteins
Peptide List (double click to dock / undock)
Peptides
Prot. Rank | Pos. | Sequence | Glycans | Protein Name | Score | Cleavage | z | Obs. m/z | Calc. m/z | Mods (variable) | PEP 2D | |Log Prob| | Delta Score | PEP 1D | elta Mo Score | ppm err. | Off-By-X
106 | 289 | R.SVGGLLCK[+648.21129].N | Hex(4) | >YIL123W SIM1 SGDID:S000001385, Chr IX from 128151-1... | 316.9 | Specific | 2 | 712.8156 | 712.8260 | K8(xgly / 648.2113) | 0.0074 | 2.13 | 87.6 | 0.0074 | 87.6 | -14.61
Spectrum (double click to dock / undock)
R.SVGGLLCK[+648].N z=2,scan#=sample=1 period=1 cycle=835 experiment=13,scan time=11.3112
SVGGLLCK
Intensity
m/z
m/z errors, tol=75.0 ppm
Protein Coverage (double click to dock...
Protein Coverage
Sample name | Protein name
2MC_20171219_... | >HORVU1Hr1G...
MAKRLVLFVAVIVALVALTT
AEREINGNNIFLDSRSRQLQ
CERELQESSLEACRRVVDQQ
LVGQLPWSTGLQMQCCQQLR
12:42 PM
20/08/2020

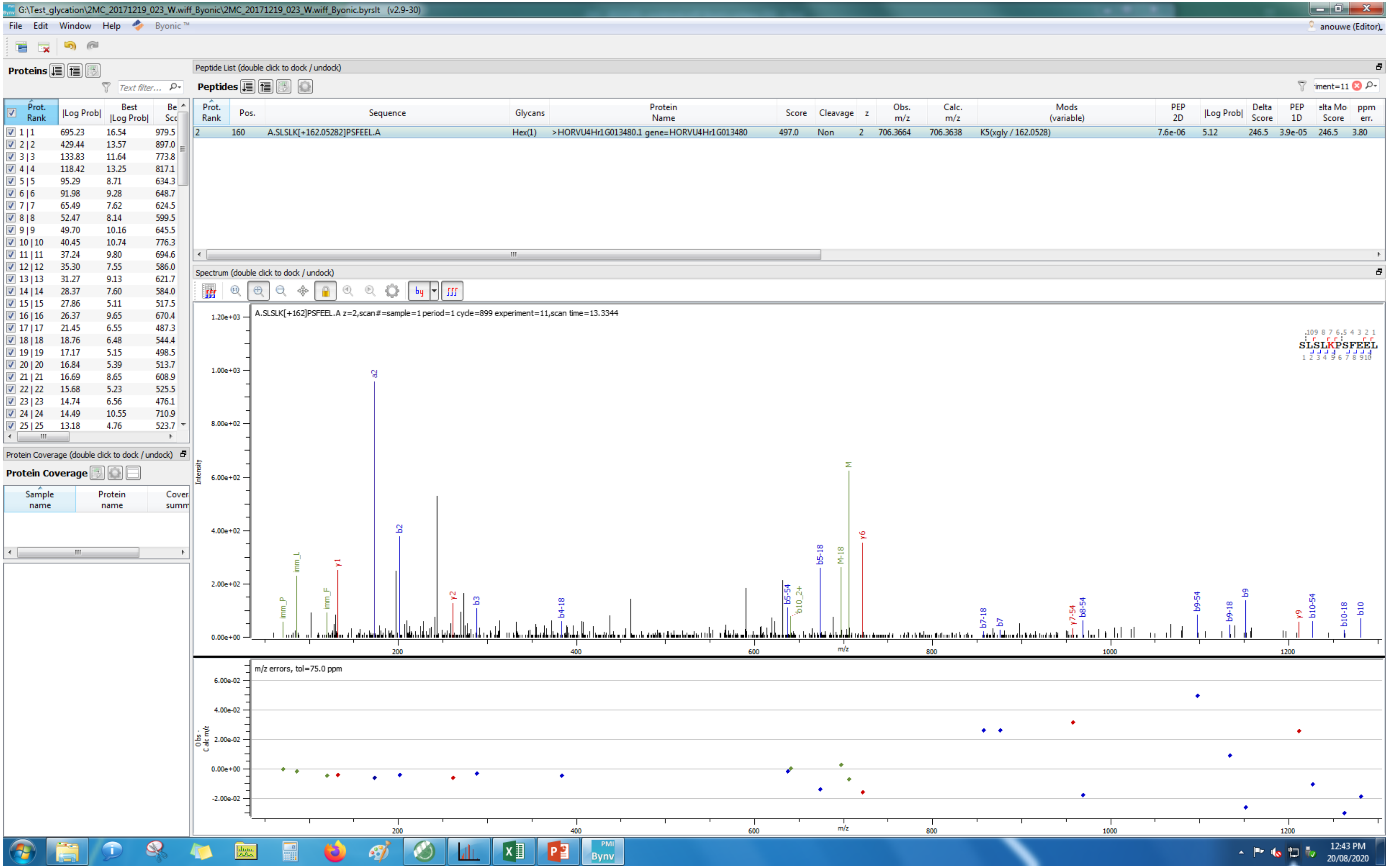
G:\Test_glycation\2MC_20171219_023_W.wiff_Byonic\2MC_20171219_023_W.wiff_Byonic.byrslt (v2.9-30)
File Edit Window Help Byonic™
anouwe (Editor)
Proteins
Text filter...
Prot. Rank |Log Prob| Best |Log Prob|
1 | 1 695.23 16.54 979.5
2 | 2 429.44 13.57 897.0
3 | 3 133.83 11.64 773.8
4 | 4 118.42 13.25 817.1
5 | 5 95.29 8.71 634.3
6 | 6 91.98 9.28 648.7
7 | 7 65.49 7.62 624.5
8 | 8 52.47 8.14 599.5
9 | 9 49.70 10.16 645.5
10 | 10 40.45 10.74 776.3
11 | 11 37.24 9.80 694.6
12 | 12 35.30 7.55 586.0
13 | 13 31.27 9.13 621.7
14 | 14 28.37 7.60 584.0
15 | 15 27.86 5.11 517.5
16 | 16 26.37 9.65 670.4
17 | 17 21.45 6.55 487.3
18 | 18 18.76 6.48 544.4
19 | 19 17.17 5.15 498.5
20 | 20 16.84 5.39 513.7
21 | 21 16.69 8.65 608.9
22 | 22 15.68 5.23 525.5
23 | 23 14.74 6.56 476.1
24 | 24 14.49 10.55 710.9
25 | 25 13.18 4.76 523.7
Protein Coverage (double click to dock / undock)
Protein Coverage
Sample name
Protein name
Peptide List (double click to dock / undock)
Peptides
Prot. Rank Pos. Sequence Glycans Protein Name Score Cleavage z Obs. m/z Calc. m/z Mods (variable) PEP 2D |Log Prob| Delta Score PEP 1D Delta Mod Score ppm err.
2 160 A.SLSLK[+162.05282]PSFEEL.A Hex(1) >HORVU4Hr1G013480.1 gene=HORVU4Hr1G013480 497.0 Non 2 706.3664 706.3638 K5(xgly / 162.0528) 7.6e-06 5.12 246.5 3.9e-05 246.5 3.80
Spectrum (double click to dock / undock)
A.SLSLK[+162]PSFEEL.A z=2,scan#=sample=1 period=1 cycle=899 experiment=11,scan time=13.3344
SLSLKPSFEEL
Intensity
1.20e+03
1.00e+03
8.00e+02
6.00e+02
4.00e+02
2.00e+02
0.00e+00
imm_P imm_L imm_F y1 a2 b2 y2 b3 b4-18 b5-54 b10_2+ b5-18 M-18 M y6 b7-18 b7 y7-54 b8-54 b9-54 b9-18 b9 y9 b10-54 b10-18 b10
200 400 600 800 1000 1200
m/z
m/z errors, tol=75.0 ppm
6.00e-02
4.00e-02
2.00e-02
0.00e+00
-2.00e-02
Obs - Calc m/z
12:43 PM
20/08/2020

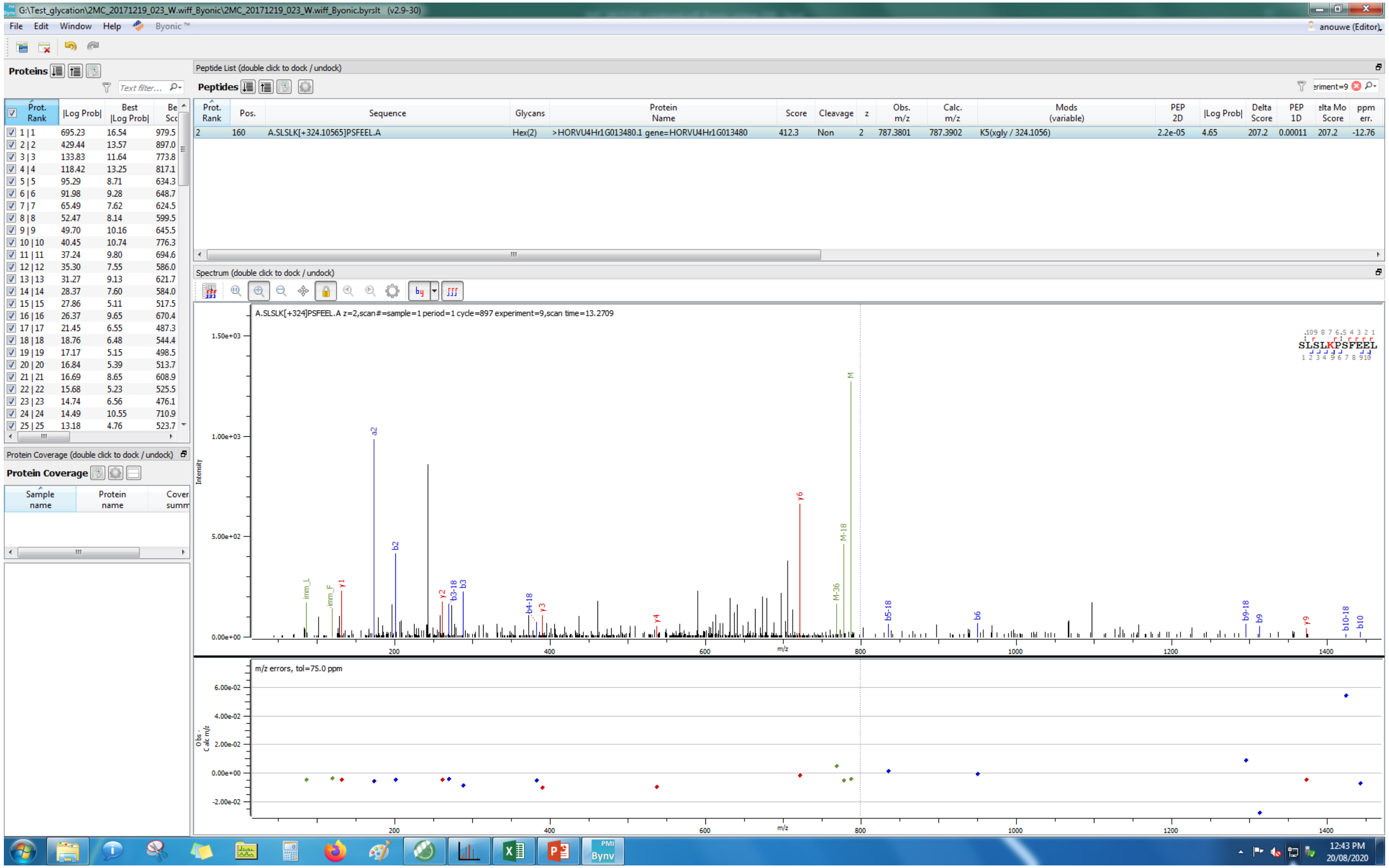
G:\Test_glycation\2MC_20171219_023_W.wiff_Byonic\2MC_20171219_023_W.wiff_Byonic.byrslt (v2.9-30)
File Edit Window Help Byonic™
Proteins
Peptide List (double click to dock / undock)
Peptides
Prot. Rank | Pos. | Sequence | Glycans | Protein Name | Score | Cleavage | z | Obs. m/z | Calc. m/z | Mods (variable) | PEP 2D | |Log Prob| | Delta Score | PEP 1D | Delta Mod Score | ppm err.
2 | 160 | A.SLSLK[+324.10565]PSFEEL.A | Hex(2) | >HORVU4Hr1G013480.1 gene=HORVU4Hr1G013480 | 412.3 | Non | 2 | 787.3801 | 787.3902 | K5(xgly / 324.1056) | 2.2e-05 | 4.65 | 207.2 | 0.00011 | 207.2 | -12.76
Spectrum (double click to dock / undock)
A.SLSLK[+324]PSFEEL.A z=2,scan#=sample=1 period=1 cycle=897 experiment=9,scan time=13.2709
SLSLKPSFEEL
Intensity
m/z
m/z errors, tol=75.0 ppm
Protein Coverage (double click to dock / undock)
Protein Coverage
Sample name
Protein name
12:43 PM
20/08/2020

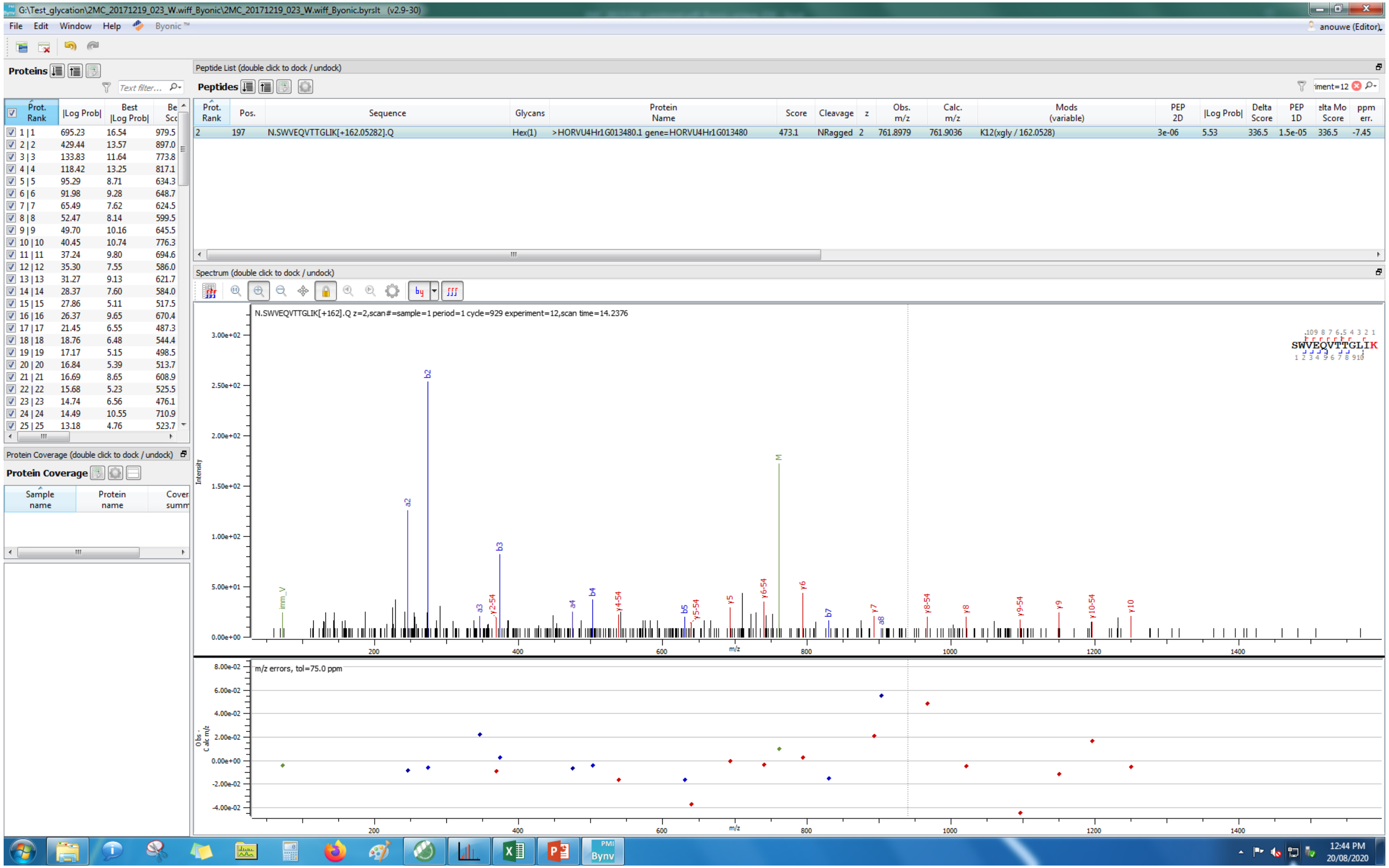
G:\Test_glycation\2MC_20171219_023_W.wiff_Byonic\2MC_20171219_023_W.wiff_Byonic.byrslt (v2.9-30)
File Edit Window Help Byonic™
anouwe (Editor)
Proteins
Peptide List (double click to dock / undock)
Peptides
Prot. Rank | Pos. | Sequence | Glycans | Protein Name | Score | Cleavage | z | Obs. m/z | Calc. m/z | Mods (variable) | PEP 2D | |Log Prob| | Delta Score | PEP 1D | Delta Mod Score | ppm err.
2 | 197 | N.SWVEQVTTGLIK[+162.05282].Q | Hex(1) | >HORVU4Hr1G013480.1 gene=HORVU4Hr1G013480 | 473.1 | NRagged | 2 | 761.8979 | 761.9036 | K12(xgly / 162.0528) | 3e-06 | 5.53 | 336.5 | 1.5e-05 | 336.5 | -7.45
Spectrum (double click to dock / undock)
N.SWVEQVTTGLIK[+162].Q z=2,scan#=sample=1 period=1 cycle=929 experiment=12,scan time=14.2376
SWVEQVTTGLIK
Intensity
m/z
m/z errors, tol=75.0 ppm
Obs - Calc m/z
Protein Coverage (double click to dock / undock)
Protein Coverage
Sample name
Protein name
12:44 PM
20/08/2020

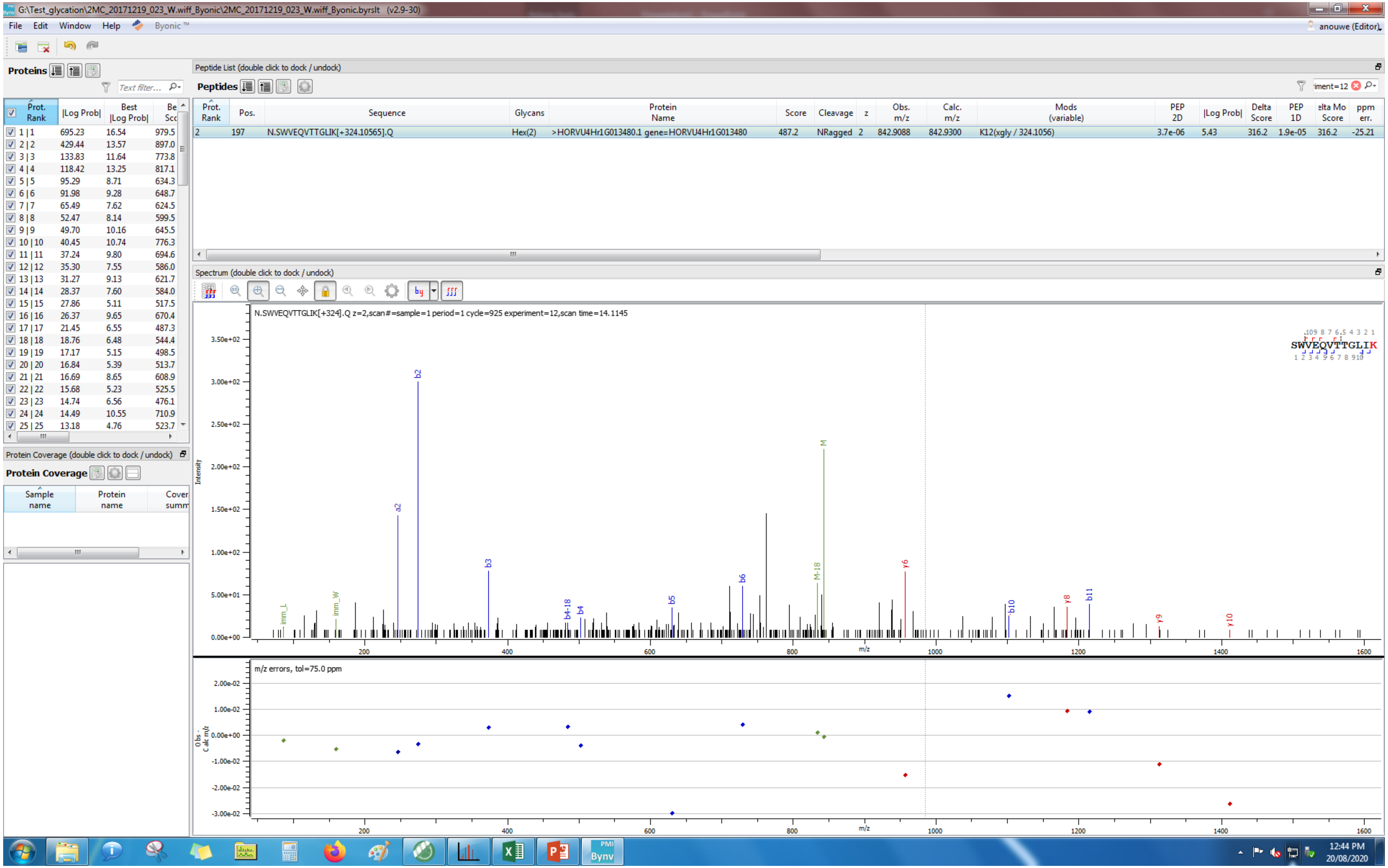

G:\Test_glycation\2MC_20171219_023_W.wiff_Byonic\2MC_20171219_023_W.wiff_Byonic.byrslt (v2.9-30)
File Edit Window Help Byonic™
anouwe (Editor)
Proteins
Peptide List (double click to dock / undock)
Peptides
Prot. Rank | Pos. | Sequence | Glycans | Protein Name | Score | Cleavage | z | Obs. m/z | Calc. m/z | Mods (variable) | PEP 2D | |Log Prob| | Delta Score | PEP 1D | Delta Mod Score | ppm err.
2 | 197 | N.SWVEQVTTGLIK[+324.10565].Q | Hex(2) | >HORVU4Hr1G013480.1 gene=HORVU4Hr1G013480 | 487.2 | NRagged | 2 | 842.9088 | 842.9300 | K12(xgly / 324.1056) | 3.7e-06 | 5.43 | 316.2 | 1.9e-05 | 316.2 | -25.21
Prot. Rank | |Log Prob| | Best |Log Prob| | Best Score
1 | 1 | 695.23 | 16.54 | 979.5
2 | 2 | 429.44 | 13.57 | 897.0
3 | 3 | 133.83 | 11.64 | 773.8
4 | 4 | 118.42 | 13.25 | 817.1
5 | 5 | 95.29 | 8.71 | 634.3
6 | 6 | 91.98 | 9.28 | 648.7
7 | 7 | 65.49 | 7.62 | 624.5
8 | 8 | 52.47 | 8.14 | 599.5
9 | 9 | 49.70 | 10.16 | 645.5
10 | 10 | 40.45 | 10.74 | 776.3
11 | 11 | 37.24 | 9.80 | 694.6
12 | 12 | 35.30 | 7.55 | 586.0
13 | 13 | 31.27 | 9.13 | 621.7
14 | 14 | 28.37 | 7.60 | 584.0
15 | 15 | 27.86 | 5.11 | 517.5
16 | 16 | 26.37 | 9.65 | 670.4
17 | 17 | 21.45 | 6.55 | 487.3
18 | 18 | 18.76 | 6.48 | 544.4
19 | 19 | 17.17 | 5.15 | 498.5
20 | 20 | 16.84 | 5.39 | 513.7
21 | 21 | 16.69 | 8.65 | 608.9
22 | 22 | 15.68 | 5.23 | 525.5
23 | 23 | 14.74 | 6.56 | 476.1
24 | 24 | 14.49 | 10.55 | 710.9
25 | 25 | 13.18 | 4.76 | 523.7
Protein Coverage (double click to dock / undock)
Protein Coverage
Sample name | Protein name | Coverage summary
Spectrum (double click to dock / undock)
N.SWVEQVTTGLIK[+324].Q z=2,scan#=sample=1 period=1 cycle=925 experiment=12,scan time=14.1145
SWVEQVTTGLIK
Intensity
m/z
m/z errors, tol=75.0 ppm
Obs - Calc m/z
12:44 PM
20/08/2020

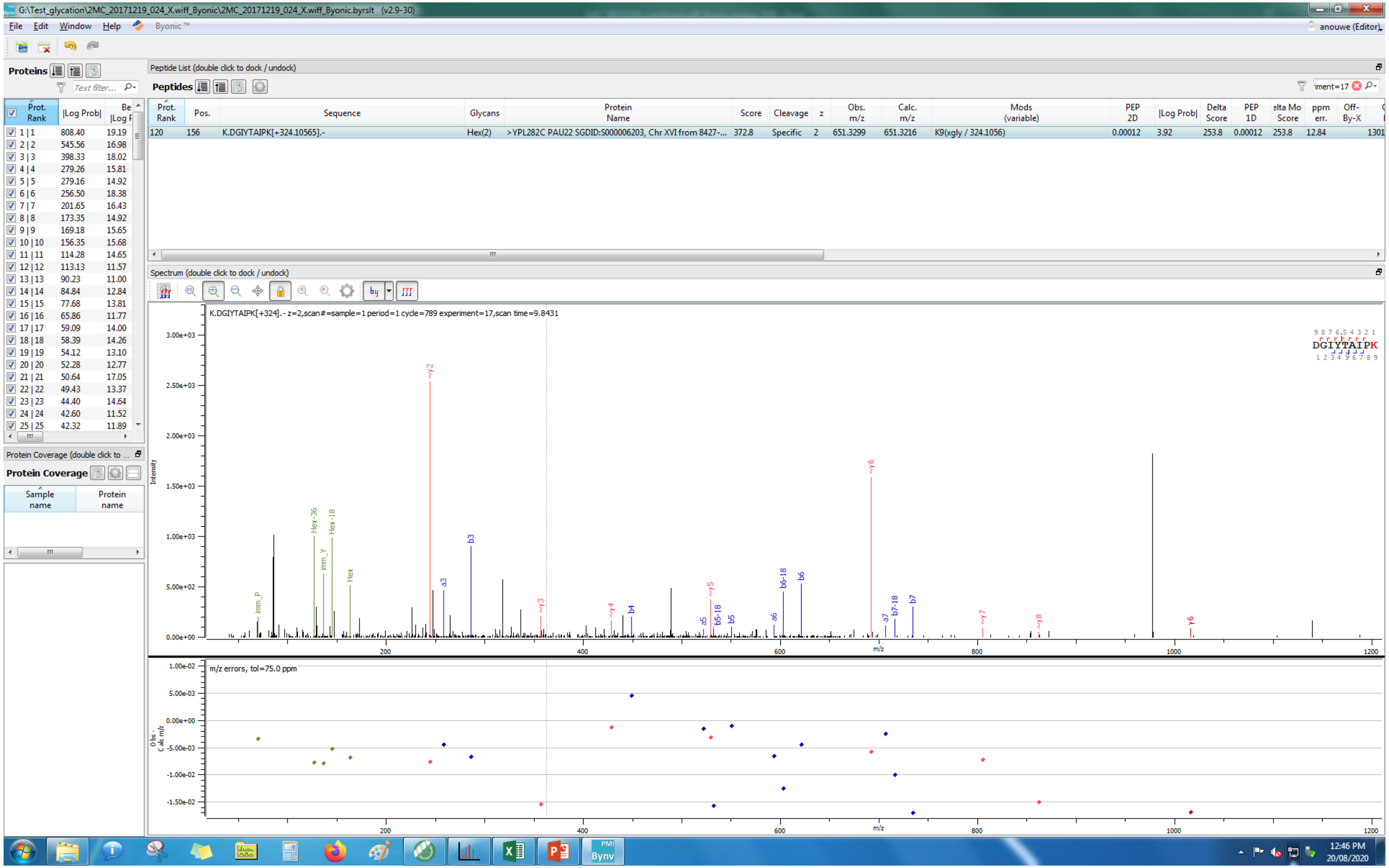
G:\Test_glycation\2MC_20171219_024_X.wiff_Byonic\2MC_20171219_024_X.wiff_Byonic.byrslt (v2.9-30)
File Edit Window Help Byonic™
anouwe (Editor)
Proteins
Peptide List (double click to dock / undock)
Peptides
Prot. Rank | Pos. | Sequence | Glycans | Protein Name | Score | Cleavage | z | Obs. m/z | Calc. m/z | Mods (variable) | PEP 2D | |Log Prob| | Delta Score | PEP 1D | Delta Mod Score | ppm err. | Off-By-X
120 | 156 | K.DGIYTAIPK[+324.10565].- | Hex(2) | >YPL282C PAU22 SGDID:S000006203, Chr XVI from 8427-... | 372.8 | Specific | 2 | 651.3299 | 651.3216 | K9(xgly / 324.1056) | 0.00012 | 3.92 | 253.8 | 0.00012 | 253.8 | 12.84 | 1301
Spectrum (double click to dock / undock)
K.DGIYTAIPK[+324].- z=2,scan#=sample=1 period=1 cycle=789 experiment=17,scan time=9.8431
DGIYTAIPK
Intensity
m/z
m/z errors, tol=75.0 ppm
Obs - Calc m/z
Protein Coverage (double click to ...
Protein Coverage
Sample name
Protein name
12:46 PM
20/08/2020

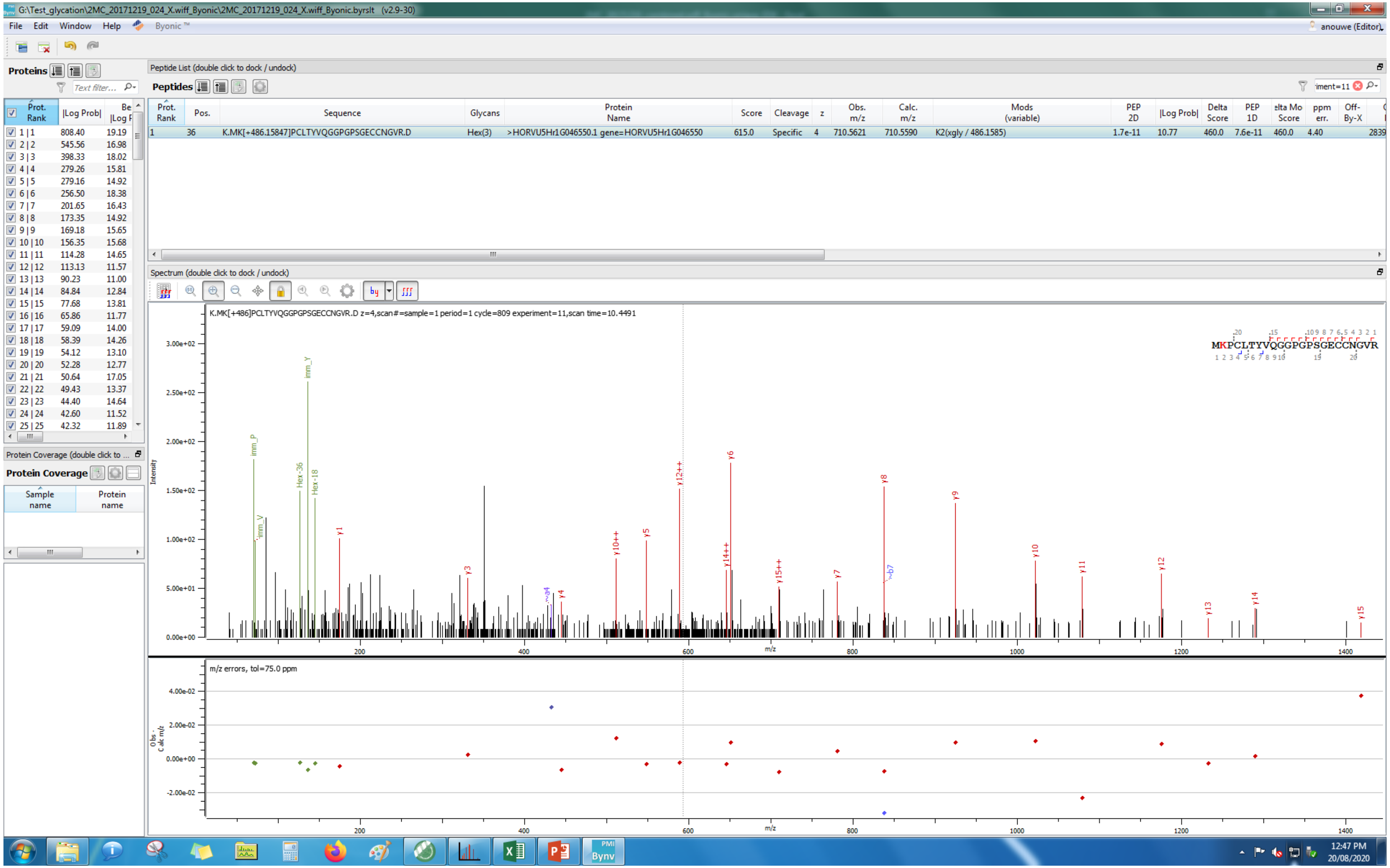
G:\Test_glycation\2MC_20171219_024_X.wiff_Byonic\2MC_20171219_024_X.wiff_Byonic.byrslt (v2.9-30)
File Edit Window Help Byonic™
anouwe (Editor)
Proteins
Peptide List (double click to dock / undock)
Peptides
Prot. Rank | Pos. | Sequence | Glycans | Protein Name | Score | Cleavage | z | Obs. m/z | Calc. m/z | Mods (variable) | PEP 2D | |Log Prob| | Delta Score | PEP 1D | ppm err.
1 36 K.MK[+486.15847]PCLTYVQGGPGPSGECCNGVR.D Hex(3) >HORVU5Hr1G046550.1 gene=HORVU5Hr1G046550 615.0 Specific 4 710.5621 710.5590 K2(xgly / 486.1585) 1.7e-11 10.77 460.0 7.6e-11 460.0 4.40
Spectrum (double click to dock / undock)
K.MK[+486]PCLTYVQGGPGPSGECCNGVR.D z=4,scan#=sample=1 period=1 cycle=809 experiment=11,scan time=10.4491
MKPCLTYVQGGPGPSGECCNGVR
Intensity
m/z
m/z errors, tol=75.0 ppm
Protein Coverage (double click to ...
Protein Coverage
Sample name
Protein name
12:47 PM
20/08/2020

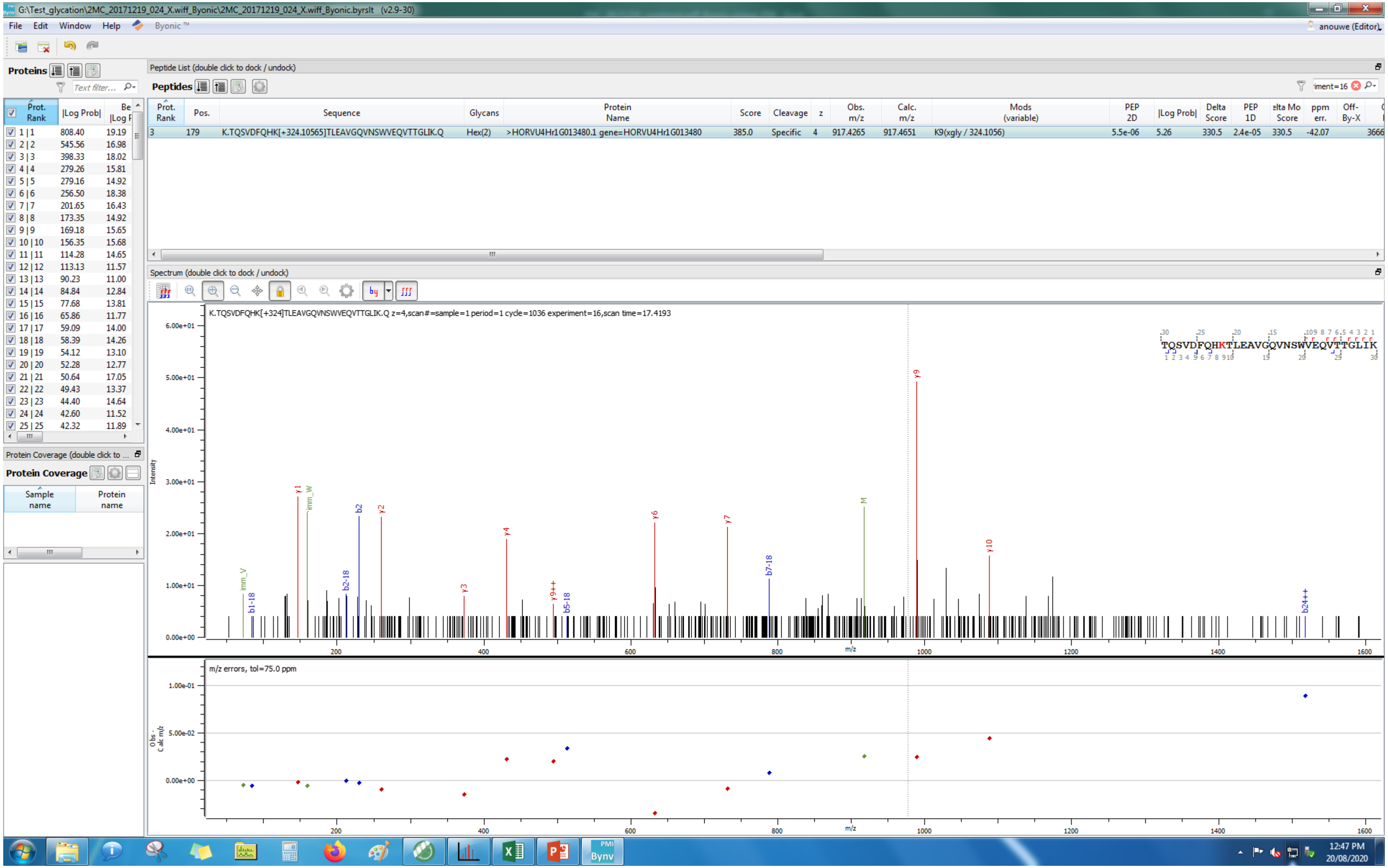
G:\Test_glycation\2MC_20171219_024_X.wiff_Byonic\2MC_20171219_024_X.wiff_Byonic.byrslt (v2.9-30)
File Edit Window Help Byonic™
anouwe (Editor)
Proteins
Peptide List (double click to dock / undock)
Peptides
Prot. Rank | Pos. | Sequence | Glycans | Protein Name | Score | Cleavage | z | Obs. m/z | Calc. m/z | Mods (variable) | PEP 2D | |Log Prob| | Delta Score | PEP 1D | Delta Mod Score | ppm err. | Off-By-X
3 | 179 | K.TQSVDFQHK[+324.10565]TLEAVGQVNSWVEQVTTGLIK.Q | Hex(2) | >HORVU4Hr1G013480.1 gene=HORVU4Hr1G013480 | 385.0 | Specific | 4 | 917.4265 | 917.4651 | K9(xgly / 324.1056) | 5.5e-06 | 5.26 | 330.5 | 2.4e-05 | 330.5 | -42.07
Spectrum (double click to dock / undock)
K.TQSVDFQHK[+324]TLEAVGQVNSWVEQVTTGLIK.Q z=4,scan#=sample=1 period=1 cycle=1036 experiment=16,scan time=17.4193
TQSVDFQHKTLEAVGQVNSWVEQVTTGLIK
Intensity
m/z
m/z errors, tol=75.0 ppm
Obs - Calc m/z
Protein Coverage
Sample name
Protein name
12:47 PM
20/08/2020

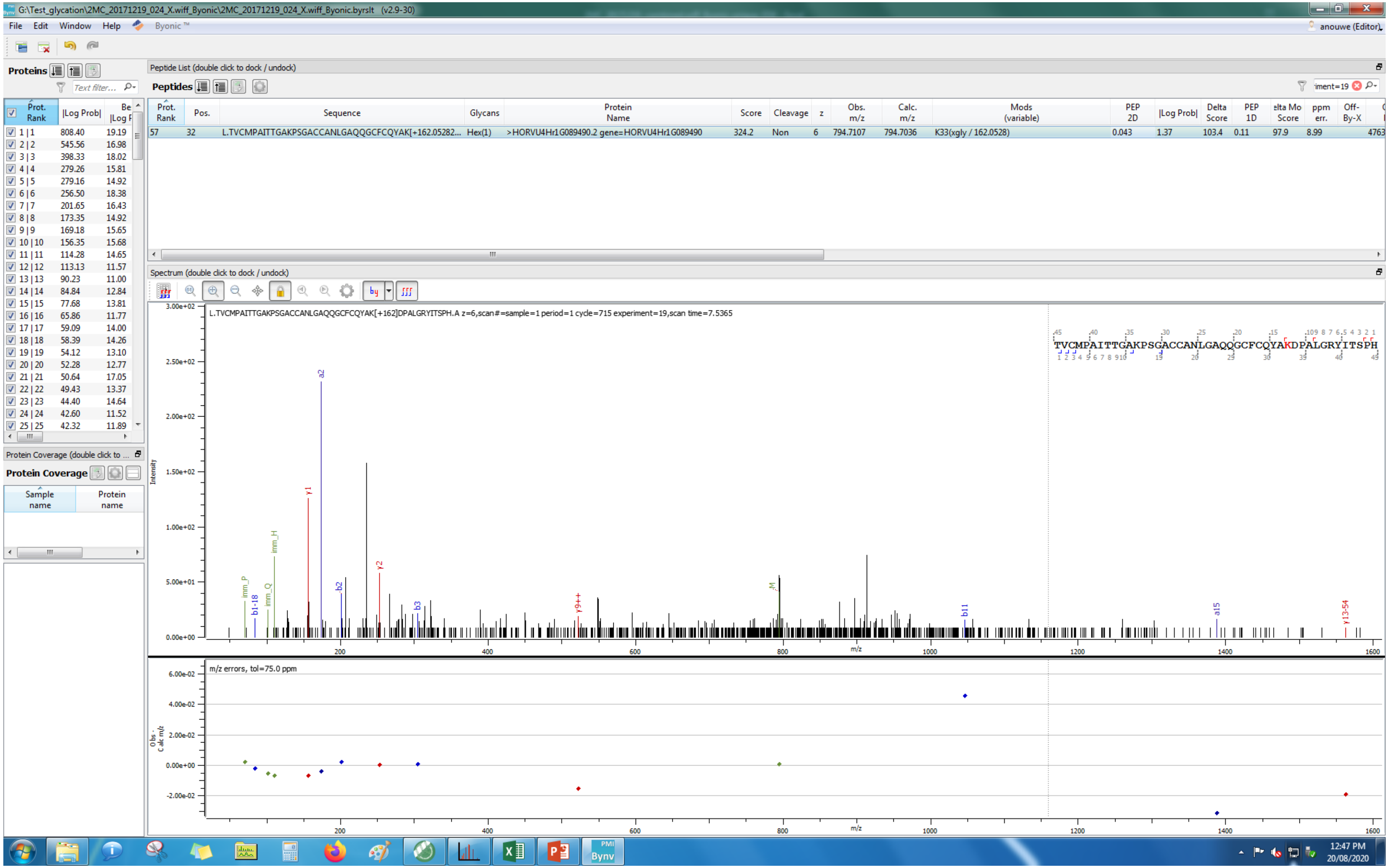
G:\Test_glycation\2MC_20171219_024_X.wiff_Byonic\2MC_20171219_024_X.wiff_Byonic.byrslt (v2.9-30)
File Edit Window Help Byonic™
anouwe (Editor)
Proteins
Peptide List (double click to dock / undock)
Peptides
Prot. Rank | Pos. | Sequence | Glycans | Protein Name | Score | Cleavage | z | Obs. m/z | Calc. m/z | Mods (variable) | PEP 2D | |Log Prob| | Delta Score | PEP 1D | Delta Mod Score | ppm err. | Off-By-X
57 | 32 | L.TVCMPAITTGAKPSGACCANLGAQQGCFCQYAK[+162.05282... | Hex(1) | >HORVU4Hr1G089490.2 gene=HORVU4Hr1G089490 | 324.2 | Non | 6 | 794.7107 | 794.7036 | K33(xgly / 162.0528) | 0.043 | 1.37 | 103.4 | 0.11 | 97.9 | 8.99
Spectrum (double click to dock / undock)
L.TVCMPAITTGAKPSGACCANLGAQQGCFCQYAK[+162]DPALGRYITSPH.A z=6,scan#=sample=1 period=1 cycle=715 experiment=19,scan time=7.5365
TVCMPAITTGAKPSGACCANLGAQQGCFCQYAKDPALGRYITSPH
Intensity
m/z
m/z errors, tol=75.0 ppm
Obs - Calc m/z
Protein Coverage
Sample name
Protein name
12:47 PM
20/08/2020

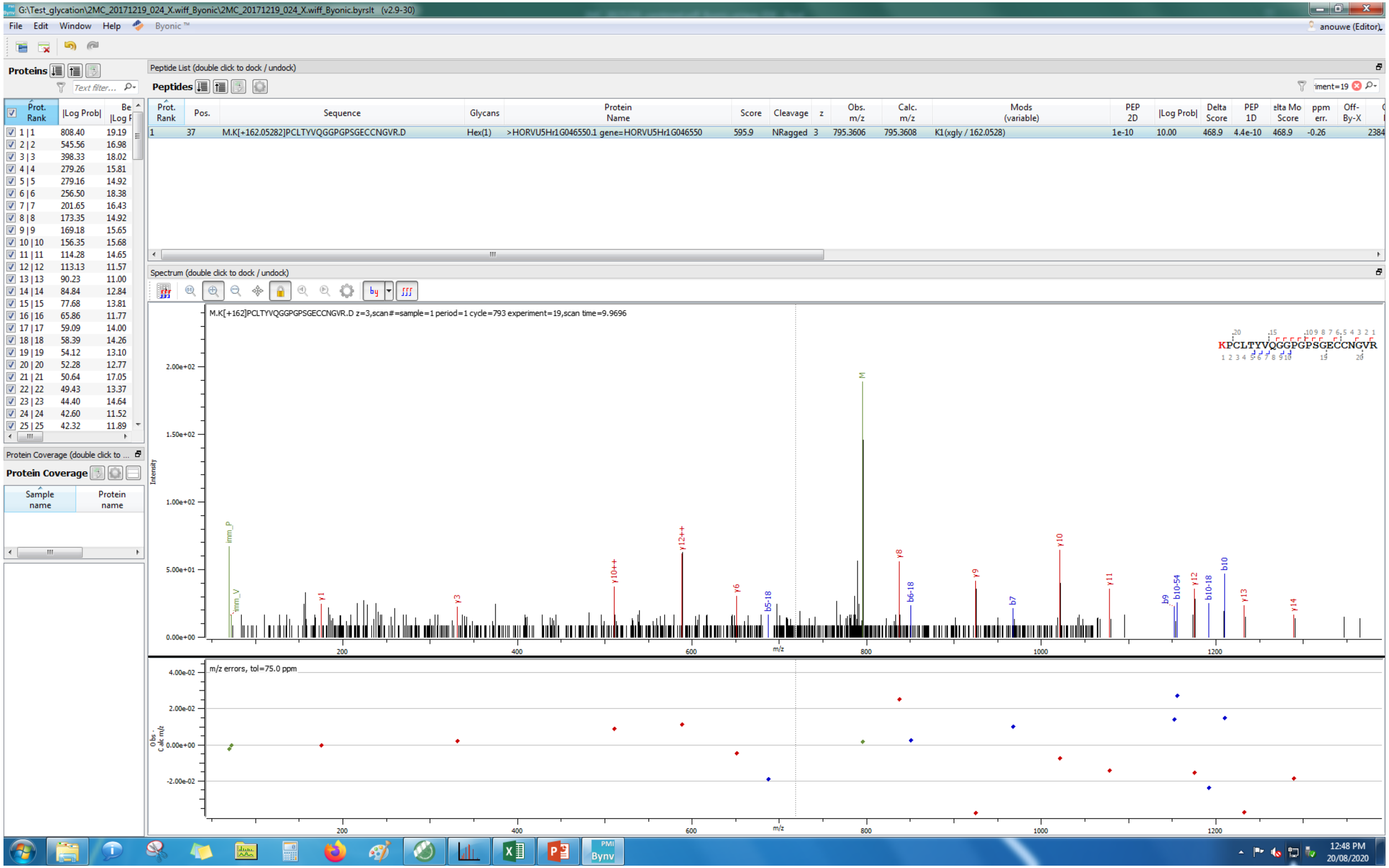
G:\Test_glycation\2MC_20171219_024_X.wiff_Byonic\2MC_20171219_024_X.wiff_Byonic.byrslt (v2.9-30)
File Edit Window Help Byonic™
anouwe (Editor)
Proteins
Peptide List (double click to dock / undock)
Peptides
Prot. Rank | Pos. | Sequence | Glycans | Protein Name | Score | Cleavage | z | Obs. m/z | Calc. m/z | Mods (variable) | PEP 2D | |Log Prob| | Delta Score | PEP 1D | ppm err.
1 | 37 | M.K[+162.05282]PCLTYVQGGPGPSGECCNGVR.D | Hex(1) | >HORVU5Hr1G046550.1 gene=HORVU5Hr1G046550 | 595.9 | NRagged | 3 | 795.3606 | 795.3608 | K1(xgly / 162.0528) | 1e-10 | 10.00 | 468.9 | 4.4e-10 | -0.26
Spectrum (double click to dock / undock)
M.K[+162]PCLTYVQGGPGPSGECCNGVR.D z=3,scan#=sample=1 period=1 cycle=793 experiment=19,scan time=9.9696
KPCLTYVQGGPGPSGECCNGVR
Intensity
m/z
m/z errors, tol=75.0 ppm
Protein Coverage
Sample name
Protein name
12:48 PM
20/08/2020

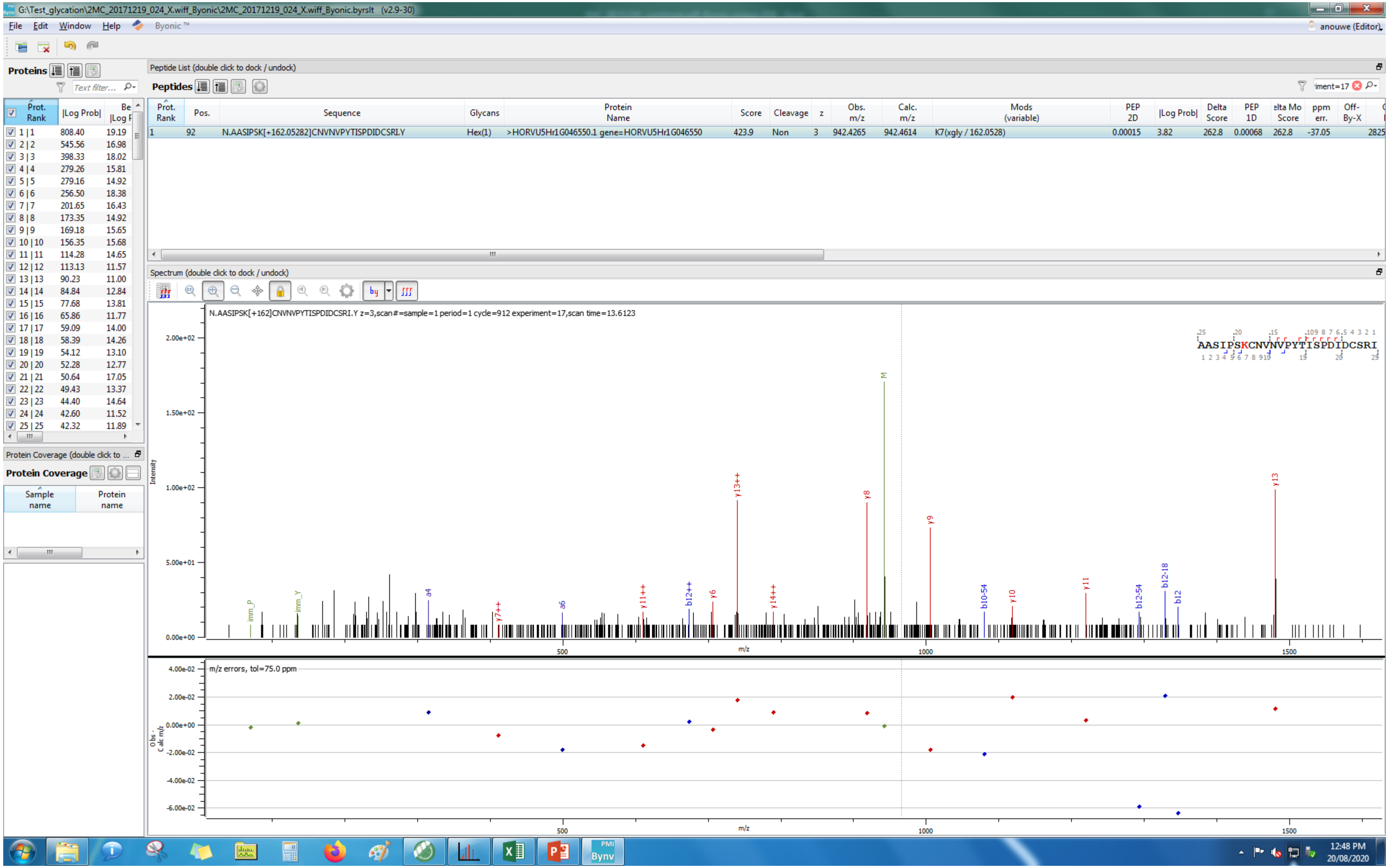
G:\Test_glycation\2MC_20171219_024_X.wiff_Byonic\2MC_20171219_024_X.wiff_Byonic.byrslt (v2.9-30)
File Edit Window Help Byonic™
anouwe (Editor)
Proteins
Peptide List (double click to dock / undock)
Peptides
Prot. Rank | Pos. | Sequence | Glycans | Protein Name | Score | Cleavage | z | Obs. m/z | Calc. m/z | Mods (variable) | PEP 2D | |Log Prob| | Delta Score | PEP 1D | Delta Mod Score | ppm err. | Off-By-X
1 | 92 | N.AASIPSK[+162.05282]CNVNVPYTISPDIDCSRI.Y | Hex(1) | >HORVU5Hr1G046550.1 gene=HORVU5Hr1G046550 | 423.9 | Non | 3 | 942.4265 | 942.4614 | K7(xgly / 162.0528) | 0.00015 | 3.82 | 262.8 | 0.00068 | 262.8 | -37.05
Protein Coverage (double click to ...
Protein Coverage
Sample name
Protein name
Spectrum (double click to dock / undock)
N.AASIPSK[+162]CNVNVPYTISPDIDCSRI.Y z=3,scan#=sample=1 period=1 cycle=912 experiment=17,scan time=13.6123
AASIPSKCNVNVPYTISPDIDCSRI
Intensity
m/z
m/z errors, tol=75.0 ppm
Obs - Calc m/z
12:48 PM
20/08/2020
