## Supplementary Material S3 for "The post-translational modification landscape of commercial beers": main.py

```
import argparse
import pandas as pd
parser = argparse.ArgumentParser(description="Automated normalization of proteolysis products in PeakView SWATH output")
parser.add_argument("-s",
                    "-swath",
                    type=str,
                    help="Filepath to PeakView peptide output file in csv format", dest="s")
parser.add_argument("-f",
                    "-fasta",
                    type=str,
                    help="Filepath to fasta file containing original sequences of the protein before being digested", dest="f")
parser.add_argument("-o",
                    "-output",
                    type=str,
                    help="Filepath to output", dest="o")

args = parser.parse_args()

input_file = args.s
input_fasta = args.f
output_file = args.o

def fasta_reader(fasta):
    df = []
    with open(fasta, "rt") as fasta_file:
        id = ""
        seq = ""
        for line in fasta_file:
            line = line.strip()
            if line.startswith(">"):

                if id:
                    df.append([id[:], seq[:]])
                stop = line.find(" ")
                id = line[1:stop+1].strip()
                seq = ""
            else:
                seq += line
        if id:
            df.append([id[:], seq[:]])
        return pd.DataFrame(df, columns=["id", "sequence"])


def digest(seq):
    tryptic = []
    length = len(seq)
    current_position = 0
    for i in range(length):
        if seq[i] in "KR":
            tryptic.append([current_position, seq[current_position:i+1], i+1 - current_position])
            current_position = i+1
    final_seq = seq[current_position:length]
    if final_seq[0] != "*":
        tryptic.append([current_position, seq[current_position:length-1], length-1-current_position])
    if not tryptic:
        tryptic.append([current_position, seq[current_position:length - 1], length - 1 - current_position])
    return pd.DataFrame(tryptic, columns=["position", "seq", "length"])


if __name__ == "__main__":

    df = pd.read_csv(input_file)
    fasta = fasta_reader(input_fasta)
    not_found = []
    df = df[~(df["Peptide"].str.contains("[", regex=False))]
    samples = df.columns[5:]
    data = []
    for i, g in df.groupby("Protein", sort=False):
        print(i)
        d = fasta[fasta["id"] == i]

        print(d)
        if d.empty:
            print(i)
            not_found.append(i)
        else:
            tryptic = digest(d["sequence"].values[0])

            for i2, r in g.iterrows():
                tryptic_locate = tryptic[tryptic["seq"].str.contains(r["Peptide"], regex=False)]
                if not tryptic_locate.empty:
                    g.at[i2, "tryptic_seq"] = tryptic_locate["seq"].values[0]
                    g.at[i2, "tryptic_location"] = tryptic_locate["position"].values[0]
                else:
                    print(r["Peptide"])
            if "tryptic_location" in g.columns:
                for i3, g2 in g.groupby("tryptic_location"):
                    total_dict = {k: g2[k].sum() for k in samples}
                    for i4, r2 in g2.iterrows():
                        for c in total_dict:
                            g2.at[i4, c] = g2.at[i4, c]/total_dict[c]
                    data.append(g2)
            else:
                print(d["sequence"].values[0])
    #columns = [df.columns[0], "tryptic_seq", "tryptic_location"] + [c for c in df.columns[1:]]
    #print(columns)

    result = pd.concat(data, ignore_index=True)
    result = result.astype({"tryptic_location": "int32"})
    result = result.set_index(["Protein", "tryptic_seq", "tryptic_location", "Peptide"])
    with pd.ExcelWriter(output_file) as writer:
        result.to_excel(writer)
```
