## Supplementary Material S3 for "The post-translational modification landscape of commercial beers": README.md

## Normalization of proteolysis products from SWATH analysis

# Requirements

Python 3.4+ with `pandas` and `openpyxl` package installed

# Usage

The program can be run from the location of the script using `python main.py`. The following parameters can be used for operating the script within the commandline.

|Paremeters|Descriptions|
|---------:|-----------:|
|`-s`| Filepath for PeakView SWATH output in csv format|
|`-f`| Filepath for FASTA file containing sequences of undigested proteins in the sample|
|`-o`| Filepath for output file in Excel format|
